# SHAPPEing agent-based models to spatial data: Sequential Hooke’s law Algorithm for Point Pattern reconstruction with Environmental constraints

**DOI:** 10.64898/2026.10.14.759208

**Authors:** Blanche Mongeon, Morgan Craig

## Abstract

Agent-based models (ABMs) are widely used to study biological systems consisting of interacting species, such as tumour and immune cells, but their predictions depend on how agents are initially placed. High-resolution imaging techniques can capture the spatial configurations of various biological systems, but their use is costly. Furthermore, they only provide small regions of interest that may be too small to initialise most ABMs directly. Here, we present the Sequential Hooke’s law Algorithm for Point Pattern reconstruction with Environmental constraints, or SHAPPE, a multitype point pattern reconstruction algorithm that generates synthetic reconstructions of arbitrary size and shape by moving points under spring-like forces with data-derived target distances. As it enforces a first-order structure through environmental constraints estimated with density grids, SHAPPE can be applied to both homogeneous and inhomogeneous point patterns. Applied to 87 glioblastoma IMC samples, we show that SHAPPE reduces dissimilarity in density distributions by a median of 53% and errors in inter-cell-type exposure levels by 34% relative to a random baseline. We then initialise a glioblastoma ABM with SHAPPE reconstructions instead of random configurations and find reduced tumour-CD8+ T cell interaction frequency with SHAPPE. This decrease translates into an over 20% decrease in the average number of tumour cells killed by CD8+ T cells over 50 days. Altogether, SHAPPE shows the potential to ease the integration of high-resolution imaging techniques into computational biology ABMs and ground initial conditions in data.

## Introduction

Biological systems, from those within cells to the ecosystem level, function through networks of interacting species. Recent advances in experimental techniques have enabled more granular spatiotemporal data at both the cell-to-cell level^1,2^ and across vast distances^3–5^. With the increased availability of such data comes a need for new analytical approaches. Mathematical models are frequently used to describe biological systems, helping to unravel the complex interactions within them. These models are largely developed using deterministic ordinary differential equations (ODEs) because they are analytically tractable, computationally efficient, and flexible. While robust and cost-effective, ODE models rely on the assumption of well-mixedness and are thus spatially homogenous, assuming uniform distributions of state variables over space. However, as we continue to generate new spatially granular data, biological processes often have non-uniform spatial characteristics that must be accounted for. For example, the growth rate of tumour cells could be impacted by clustering dynamics due to competition for nutrients^6–8^ whereas physical barriers, like blood vessels or mountain ranges, can create regions of density^9–11^.

Though partial differential equations relax the assumption of spatial uniformity, there are limited techniques for their analysis, including numerically. Accordingly, there is a growing interest in using agent-based models (ABMs) to describe multiscale spatial interactions in biological systems. ABMs are defined by individual agents (e.g., cells, trees, animals, etc.) interacting with each other over time and space^12,13^. Importantly, unlike in ODE models, the behaviour of each agent is driven by rules that may depend on their local neighbourhood. A key aspect of spatial ABMs is the initial placement of agents (spatial initialisation). Thanks to advancements in high-resolution imaging techniques, we can obtain realistic spatial configurations of various biological processes. For instance, imaging mass cytometry (IMC) offers single-cell level information about the position and the phenotype of cells in the tumour microenvironment^1,14^ while terrestrial light detection and ranging (TLiDAR) provides tree-level details of the spatial organisation of a forest^15^. In theory, the data obtained from these imaging techniques could be used to initialise ABMs. In practice, integrating new, high resolution spatial data is costly^16^. Further, they may only provide a window onto a small region of interest within the biological system; for example, IMC data is frequently limited to a 1 mm^2^ area^17^ (versus a solid tumour that can exceed 10 cm in diameter^18^). Thus, initialising an ABM directly from such data may potentially limit the variability of the system, particularly if the measurement was taken in an atypical region of the system. Further, biological systems operate at multiple time scales. For example, both microscopic within-cell cellular division processes and distal cell-to-cell immune interactions influence tumour growth. Given these, insufficient scaling of an ABM can lead to erroneous conclusions^19^. Unfortunately, it may not always be possible to satisfy this constraint using the previously mentioned imaging techniques due to e.g., high computational costs or resources limitations^20^. As such, previous studies have relied on random initialisations when using ABMs^19,21^, regrettably discarding the observed spatial information.

In response, the use of synthetic data configured to have similar spatial characteristics and an increased domain size has been proposed for initialising ABMs. Popular in ecology, multitype point pattern reconstruction methods^20,22,23^ are such an approach and use points, each characterised by a categorical type, to reconstruct and conserve spatial motifs in observed data. Pommerening and Stoyan^20^ developed a reconstruction method for tree point patterns that accounts for different tree species. Based on simulated annealing, their method minimises an energy function quantifying the difference in nearest neighbour summary statistics between the reconstructed and the reference patterns. Wudel et al. ^22^ extended this work to plant ecosystems to better capture inter-type spatial correlations by including in their energy function different spatial summary statistics (e.g., pair-correlation function, mark correlation function, *K*-function, *F*-function, etc.). These point pattern reconstruction methods require repeated evaluations of summary statistics for their optimisation schemes, the choice of which is arbitrary. Hence, if the domain is large and/or the configuration contains a high number of points, this optimisation can be computationally costly. Also, this approach may not be suitable for highly inhomogeneous point patterns, as distinct point processes can exhibit similar second-order statistics, yet different first-order intensity structures^23^.

To address these limitations, here we propose *Sequential Hooke’s law Algorithm for Point Pattern reconstruction with Environmental constraints*, or SHAPPE, a multitype point pattern reconstruction algorithm that preserves mechanistic meaning. In SHAPPE, points (i.e., agents) act as springs and interact with one another through repulsive and attractive forces that depend on target distances estimated from the original data. While reconstruction methods have been extended to inhomogeneous point patterns by including differences in estimated intensity functions in the energy function^24^, they rely on an accurate estimate of the first-order intensity function. In contrast, SHAPPE identifies low-density regions explicitly and enforces them as environmental constraints using density grids. To illustrate SHAPPE and evaluate its fidelity, we use a set of 87 IMC samples from glioblastoma patients^14^ as a large-scale case study. We compare our framework to a baseline random placement and show that SHAPPE better recovers first- and second-order spatial statistics for homo- and inhomogeneous point patterns, without requiring iterative evaluation during point placement steps as other methods do^20,22,24^. We further apply SHAPPE to generate synthetic configurations with a large initial number of cells to evaluate its potential for initialising ABMs. Our results show that SHAPPE succeeds at preserving spatial statistics even when the domain area is substantially increased. We then proceed to initialise an ABM describing tumour growth with inhomogeneous tumour microenvironment (TME) synthetic reconstructions. Our results demonstrate that predictions obtained from realistic initialisations recover immunosuppressive mechanisms and tumour aggressivity, thus underlining the need for improved spatial reconstruction approaches, such as SHAPPE. Although we apply SHAPPE to TME spatial configurations, the workflow itself is completely independent of the context. By simply adjusting the parametrisation, it could be adapted to the reconstruction of point patterns emerging from numerous other biological processes.

## Methods

### Sequential Hooke’s Law Algorithm for Point Pattern Reconstruction with Environmental Constraints (SHAPPE)

SHAPPE generates an artificial point pattern with similar spatial statistics as an observed point pattern by extending the idea introduced by Jenner et al.^25^, who used Hooke’s law to place points in the domain of an ABM. Consider a set of points *x*_*i*_ (*i* = 1, …, *N*) on a domain *D* ⊂ ℝ^2^. A *multitype point pattern* is a set *X* where each *x*_*i*_ is attributed a categorical type 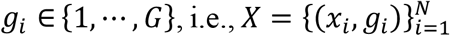. We have *X*_*g*_ = {*x*_*i*_ ∈ *X*|*g*_*i*_ = *g*}, with *N*_*g*_ the number of points of type *g*. Notations supplemented by the superscripts (*Obs*) and (*Art*) refer respectively to the attributes of an observed point pattern, *X*^(*Obs*)^, and an artificial point pattern, *X*^(*Art*)^, reconstructed from *X*^(*Obs*)^. For example, *N*^(*Art*)^ and *N*^(*Obs*)^ are the number of points in *X*^(*Art*)^ and *X*^(*Obs*)^, respectively. *X*^(*Art*)^ contains 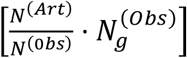 points of type *g*, for *g* ∈ {1, ⋯, *G*}, where [*x*] is the nearest-integer function. Note that the area, *A*(·) (Table S2), of *D*^(*Art*)^ must satisfy 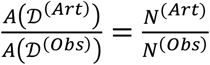.

SHAPPE begins with three preliminary steps. Step 0.1 analyses and quantifies inhomogeneity, Step 0.2 detects significant clustering patterns in *X*^(*Obs*)^, and Step 0.3 generates a baseline configuration under complete spatial randomness (CSR; Supplementary Section M1) or environmentally constrained spatial randomness (ECSR; Supplementary Section M4). Here, CSR is defined as a conditional Poisson process in which every point is at least *h* units from its closest neighbour, i.e., a Gibbs hard-core point process with exclusion distance *h* that reduces to a traditional Poisson process if *h* = 0. ECSR is defined as a Gibbs hard-core point process with exclusion distance *h* and sampled repulsive or attractive zones. The value of *h* can be inferred from the nature of the points (e.g., minimal realistic distance between two cells) or be set to 0 if there is no prior information on its value. SHAPPE then proceeds in six steps that are repeated for every agent type *g*, during which the positions of points 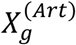 are iteratively modified to recreate significant clustering patterns observed in *X*^(*Obs*)^.

#### SHAPPE steps overview and pseudocode

The three preliminary steps of SHAPPE proceed as follows.

**Step 0.1**:Inhomogeneous environmental constraints over a domain can create first-order inhomogeneity with some regions being denser than others while some others remain empty. We begin by classifying *X*^(*Obs*)^ as homogeneous or inhomogeneous by assuming that points of all types share the same underlying first-order intensity function (i.e., the expected density of each cell type varies across the domain in the same way, with types differing only in their overall abundance). To identify inhomogeneity associated with an inhomogeneous intensity function, we follow a two-step process by ignoring point types (Figure 1). First, we test whether *X*^(*Obs*)^ has empty regions larger than what is expected under CSR using an envelope test based on the *F*-function, also known as the empty-space function, that measures the cumulative distribution of the distance from an arbitrary location within the domain to the nearest point of a point pattern. Following Myllymäki et al.^26^, we build a null envelope of first-order homogeneity by performing 1,000 simulations under CSR and estimating their respective *F*-function. To test for statistically significantly larger empty regions, we use a one-sided global envelope test based on maximum absolute deviation below the null model with significance at the *α* = 0.05 level (Supplementary Section M2). Second, we quantify empty regions in *X*^(*Obs*)^ from an intensity grid by searching for regions with no points and whose surface areas meet a minimum size criterion *A*_2_. These are classified as *zero-density regions*. Finally, a point pattern is classified as inhomogeneous if 1) it has an empty space distribution that is statistically significantly different than what is expected under CSR and 2) it contains at least one valid zero-density region; otherwise, it is deemed to be homogeneous. This classification process is designed to prevent false detections of inhomogeneity, as the first step could falsely identify inhomogeneous dense point patterns in regions with small holes created by local clustering rather than by variation in the first-order intensity function, while the second could pick out inhomogeneous point patterns in empty regions caused by low global density. Complete details on the construction of the intensity grid and the identification zero-density regions can be found in Supplementary Section M3.

**Figure 1.**
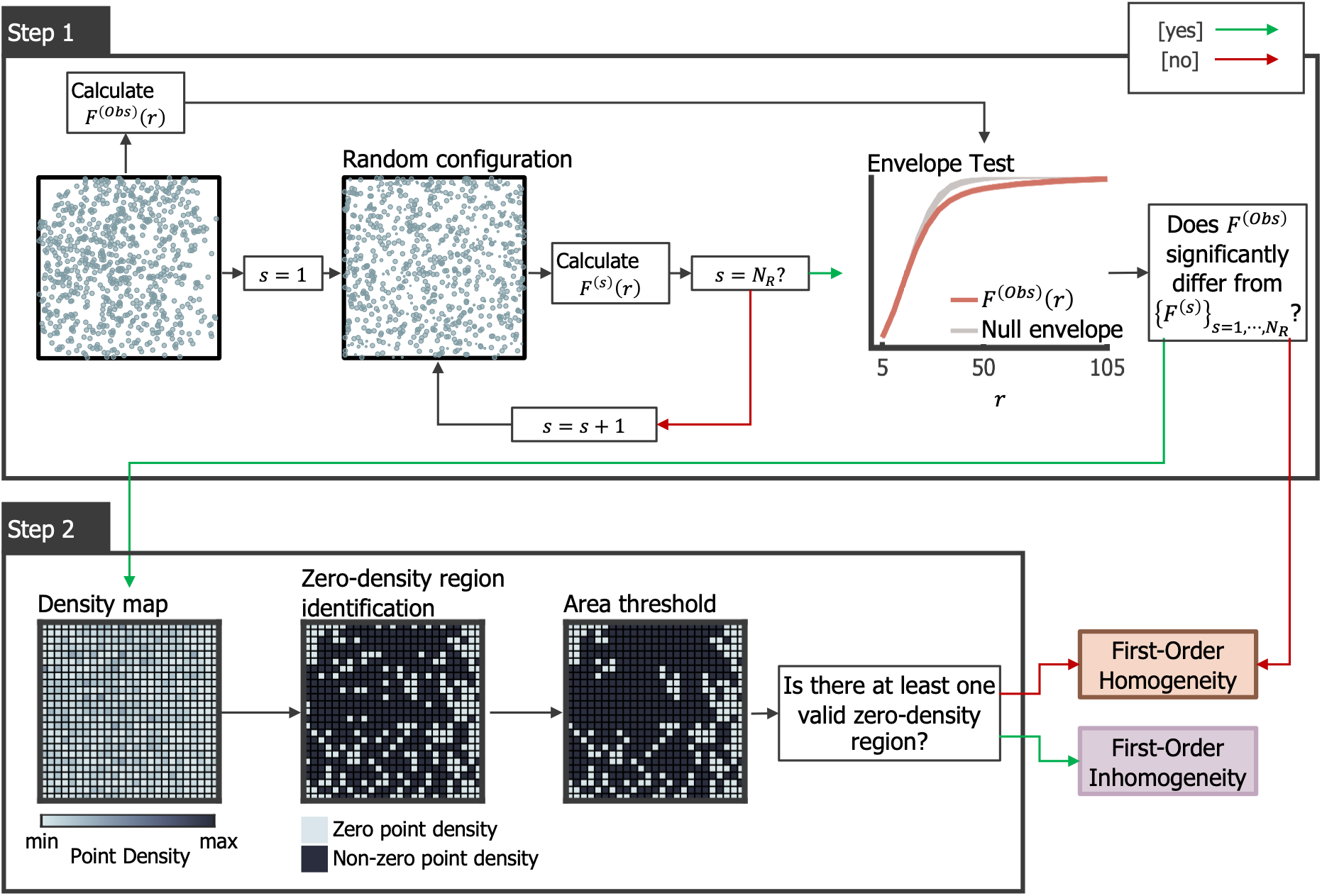
Two-step process for first-order inhomogeneity identification. **Step 1** consists of an envelope test based on the ***F***-function, also known as the empty-space function. If ***X*** has empty regions significantly larger than what is expected under complete spatial randomness (one-sided global deviation test, ***α*** = **0. 05**), we move on to Step 2. Otherwise, ***X*** is automatically classified as homogeneous. In **Step 2**, we build a density grid and label bins with zero-point density as *zero-density bins*. Groups of connected bins (Moore’s neighbourhood) satisfying a minimal area threshold are then labeled as zero-density regions. If ***X*** has at least one valid zero-density region, it is classified as inhomogeneous. Otherwise, it is classified as homogeneous.

**Step 0.2**: Next, we check whether there are differences in observed pairwise occurrences versus what is expected under randomness. For this, we evaluate Ripley’s univariate and multivariate *K*-functions on a relevant set of distances ℛ. The *K*-function, *K*_*gk*_(*r*), measures the expected number of points of type *k* within a distance *r* of a typical point of type *g*, normalised by the intensity of *X*_*k*_. A random labelling envelope for *K*_*gk*_(*r*) is defined using Monte Carlo simulations in which point locations are preserved from *X*^(*Obs*)^ and types are randomly permuted^27^, which allows for the evaluation of *K*_*gk*_(*r*) in *X*^(*Obs*)^ for all *r* ∈ ℛ and all pairs of types (*g, k*) ∈ [1, ⋯, *G*] × [1, ⋯, *G*] (Figure S3). We then compare *K*_*gk*_(*r*) to the observed function in *X*^(*Obs*)^, 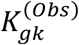, using a two-sided global test based on maximum absolute deviation from the random labelling null, following Myllymäki et al.^26^ (Supplementary Section M5). We test the null hypothesis that type assignment is independent of spatial location after conditioning on the observed underlying point pattern, consistent with our assumption that spatial inhomogeneity is not type-specific. If 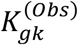 is significantly outside of its corresponding envelope, then points of type *g* encounter points of type *k* more(less) often than expected under intensity-constrained randomness. If that is the case, we set Ω_*gk*_ = 1; otherwise, Ω_*gk*_ = 0.

**Step 0.3**: If *X*^(*Obs*)^ is homogeneous, we generate a baseline configuration *X*^(*R*)^ under CSR; if *X*^(*Obs*)^ is inhomogeneous, *X*^(*R*)^ is generated under ECSR.

Once the baseline configuration is generated, the main steps of SHAPPE are performed (Figure 2). In what follows, we describe these steps for the placement of agents of type 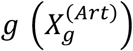, proceeding in the order [1,2, …, *g*, …, *G*].

**Figure 2.**
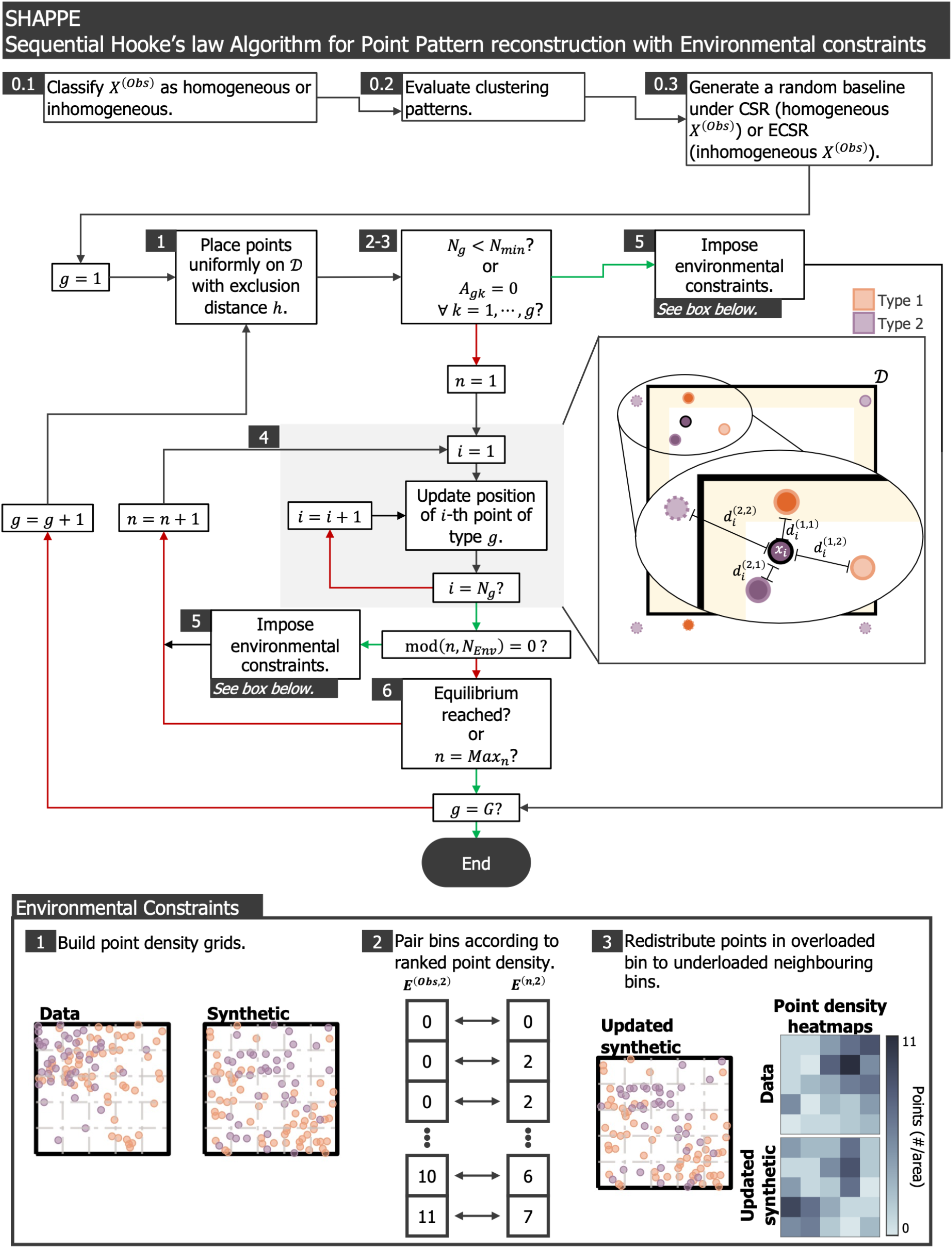
Sequential Hooke’s law Algorithm for Point Pattern reconstruction with Environmental constraints (SHAPPE). SHAPPE starts with three preliminary steps consisting of a first-order inhomogeneity identification (Step 0.1), a clustering pattern assessment (Step 0.2) and a random baseline generation (Step 0.3). Then, for each SHAPPE replicate, points are moved iteratively one type at a time. Points act as springs, repulsing or attracting each other based on target distances estimated from the data (if significant clustering is observed) or from the random baseline. Positions of points of a type are fixed once an equilibrium is reached. Environmental constraints are enforced every *n*_*Env*_ iterations through a comparison between density grids (bottom box).

**Step 1**: <u>Place *N*^(*Art*)^points of type *g* uniformly on *D*^(*Art*)^ with an exclusion distance *h*</u>. For this, sample a position *p*_*i*_ for the *i*-th point *x*_*i*_ ∈ *X*_*g*_ following a uniform distribution on *D*^(*Art*)^. The acceptance criterion for *p*_*i*_ depends on its distance to points already placed: if another point is less than *h* units from *p*_*i*_, reject *p*_*i*_ and resample another position. Otherwise, fix *x*_*i*_ at position *p*_*i*_ and move on to *x*_*i*+1_. If no valid position for *x*_*i*_ is found within 1,000 resampling tries, accept the best position out of the 1,000 tried positions to avoid infinite looping.

**Step 2**: <u>Assess whether the data are sufficient to provide information about the spatial distribution of *X*_*g*_</u>. If 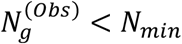, assume the data offer no conclusion relative to the spatial distribution of points of type *g*. Move on to **Step 5**.

**Step 3**: <u>Verify encounter probabilities for agents of type *g* with agents of other types</u>. If Ω_*gk*_ = 0 for all *k* ∈ {1, ⋯, *g*}, then agents of type *g* encounter other agents of all types *k* ∈ {1, ⋯, *g*} as often as what is expected under intensity-constrained randomness. In that case, move on to **Step 5**. Otherwise, set the iteration number *n* to 1 and move to **Step 4**.

**Step 4**: <u>Update point positions</u>. Point positions are updated per the Jacobi method, i.e., point displacements at iteration *n* only consider the positions of points at the start of the iteration. In what follows, we use subscripts to refer to the point being updated (reference point) and superscripts to refer to its neighbours (type and rank). To impose boundary conditions and prevent agents from moving out of the domain, *D*^(*Art*)^, we include ghost points in the search of neighbours (Supplementary Section M7.1). Each point can have at least one corresponding ghost on an opposite side of the domain. To avoids ghost points outnumbering real points when updating positions, we only consider ghosts of points within a distance *d*_*B*_ of the boundary of *D*^(*Art*)^. Let 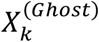 denote the set of valid ghosts of type *k* ∈ {1, ⋯, *G*}.

For every point 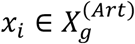:

**Step 4.1**: Calculate its raw displacement from the position of its *L* closest neighbours of each type *k* ∈ {1, ⋯, *g*}, and from target distances that depend on spatial correlations. *p*_*i*_(*n*) and 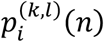, respectively, refer to the position of *x*_*i*_ at iteration *n* and to the position of its *l*^*th*^ closest neighbour among 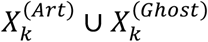. Denote 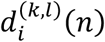 as the distance between the two points. The interaction force between *x*_*i*_ and its *l*^*th*^ neighbour from 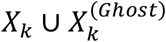 is given by

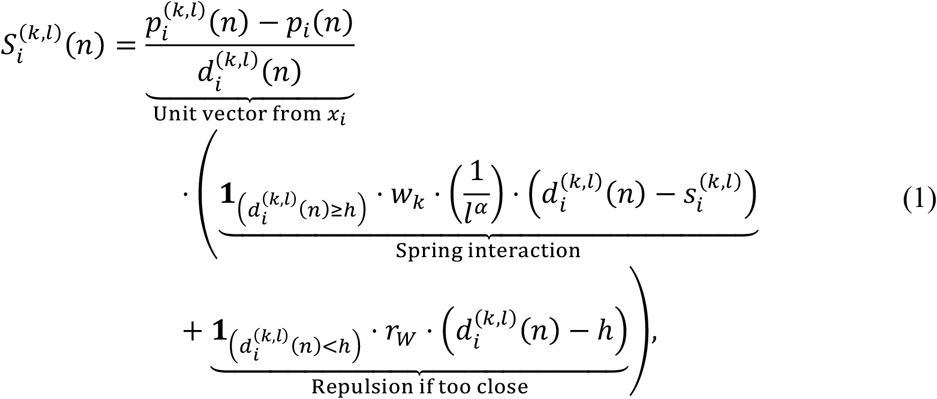

where **1**_(*condition*)_ is the indicator function that evaluates to 1 if the condition is true, and to 0 otherwise. In Eq. (1), *α* weights neighbours of rank *l, r*_*W*_ is the repulsion weight to impose the exclusion distance *h*, and 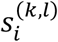 is the target distance assigned to this neighbour. If Ω_*gk*_ = 1, sample 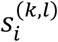 from *X*^(*Obs*)^ and *w*_*k*_ = 1. To preserve correlations observed in *X*^(*Obs*)^, for all *k* ∈ [1, ⋯, *g*] such that Ω_*gk*_ = 1 and all *l* ∈ [1, ⋯, *L*], sample *s*^(*k,l*)^ values jointly using a copula of type *C*. If Ω_*gk*_ = 0, sample 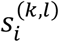 from *X*^(*R*)^ (**Step 0.3**) and *w*_*k*_ = *w*_*NoCorr*_. In this way, pairwise occurrences that do not significantly differ from the random baseline are represented using the random baseline rather than *X*^(*Obs*)^. Supplementary sections M7.2 and M7.3 describe how to sample target distances.

Finally, the raw displacement of *x*_*i*_ at iteration *n* is calculated as

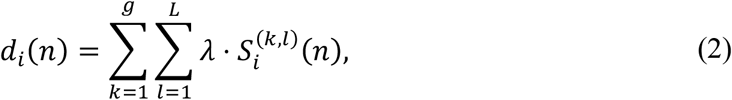

where *λ* dampens strength interactions for numerical stability. For inhomogeneous point patterns, additional environmental constraints are implemented at this step by adding repulsion from repulsive regions in *X*^(*R*)^ (or attraction to attractive regions in *X*^(*R*)^) to Eq. (2), depending on the scenario used to generate *X*^(*R*)^ (Supplementary Section M7.4).

**Step 4.2**: Since *L* is user-defined, *d*_*i*_(*n*) (Eq. (2)) could become quite large so the displacement magnitude of *x*_*i*_ is bounded by an upper step-size limit *Φ*(*n*) that depends on an initial cap *Φ*, which decreases with the iterations until it reaches *Φ*_*min*_:

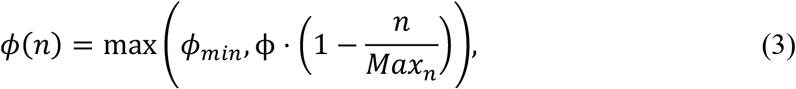

where *Max*_*n*_ is the maximum number of iterations (see **Step 6**).

**Step 4.3**: Calculate the next proposed position for *x*_*i*_ per

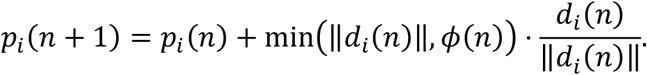

If *p*_*i*_(*n* + 1) falls outside of the domain, we define periodic boundary conditions for a square domain and an approximation of symmetric boundary conditions for a disk (Supplementary Section M7.5). This is why we consider ghost points when looking for neighbours^25^ (**Step 4.1**).

**Step 5**: <u>Enforce environmental constraints</u>. At every *n*_*Env*_ iteration, we define the density grid *E*^(*bs,g*)^ as a proxy for environmental constraints. For this, estimate *E*^(*bs,g*)^ from *X*^(*Obs*)^ while considering only points of type *g* and those placed prior (i.e., types 1, ⋯, *g*), binning such that the side length of a bin in *E*^(*bs,g*)^ is approximately equal to a user-defined quantity *bL*_*Env*_. As with other parameters, *bL*_*Env*_ can be inferred from the nature of the points. Then, compare *E*^(*bs,g*)^ to the corresponding density grid of the current point placement, *E*^(*n,g*)^, by ranking bins based on their point density and pairing same-rank bins (Figure 2). Bins in *E*^(*bs,g*)^ determine the capacity of their paired bin in *E*^(*n,g*)^. Finally, redistribute points from overloaded bins to underloaded neighbouring bins (Supplementary Section M7.6).

**Step 6**: Fix the position of the points *X*^(*Art*)^. When any of the four ending criteria:

1. The median displacement for points in 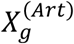 is less than *d*_*med*_ units and the 90^th^ quantile is less than *d*_90_ units,
2. The median net displacement over the last 25 iterations is less than *d*_*Net*_ units and *n* ≥ *n*_*Eq*_,
3. *n* = *Max*_*n*_,
4. *N*^(*Obs*)^ < *N*_*min*_ or Ω_*gk*_ = 0 ∀ *k* = 1, ⋯, *g*,

is reached, fix the positions of points 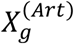. If *g* < *G*, repeat **Steps 1-6** for points 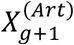, else end the algorithm if *g* = *G*. If no criterion is achieved, set *n* = *n* + 1 and return to **Step 3**.

Hyperparameters and parameters are summarised in Table 1 and Table 2, respectively.

**Table 1.** SHAPPE hyperparameters.

| Hyperparameter | Description | Interval of definition | SHAPPE Step |
| --- | --- | --- | --- |
| $L$ | Number of closest neighbours considered when updating the position of a point. | $[1, +\infty)$ | 4.1 |
| $w_{NoCorr}$ | Weight attributed to uncorrelated types, relative to weight attributed to correlated types | $[0,1]$ | 4.1 |
| $C$ | Copula family to sample target distances for types with significant spatial correlations | {Gaussian, Student-$t$} | 4.1 |
| $\alpha$ | Neighbour influence weight | $[0, +\infty)$ | 4.1 |
| $r_w$ | Repulsion weight to enforce an exclusion distance | $[0, +\infty)$ | 4.1 |
| $\lambda$ | Movement dampening parameter for numerical stability | $(0,1]$ | 4.1 |
| $\phi$ | Baseline step limit | $(0, +\infty)$ | 4.2 |
| $\phi_{min}$ | Minimum step limit | $[0, \phi]$ | 4.2 |

**Table 2.** SHAPPE parameters.

| Parameter | Description | SHAPPE Step |
| --- | --- | --- |
| $N_{min}$ | Inferior limit for $N_g$ under which step 4 is skipped for points $X_g$ | 2 |
| $h$ | Exclusion (i.e. minimum) distance between points | 3 |
| $d_B$ | Maximal distance from the boundary of $\mathcal{D}$ for a point to have a ghost point considered. | 4 |
| $n_{Env}$ | Interval between iterations during which we apply environmental constraints | 5 |
| $bL_{Env}$ | Approximative length of bins in the grid generated to enforce macro-constraints | 5 |
| $d_{Med}$ | Threshold for the median displacement for ending criterion 1 | 6 |
| $d_{90}$ | Threshold for the 90<sup>th</sup> quantile of displacements for ending criterion 1 | 6 |
| $d_{Net}$ | Threshold for the median net displacement over last 25 iterations for ending criterion 2 | 6 |
| $Max_n$ | Maximum number of iterations | 6 |
| $n_{Eq}$ | Minimum number of iterations for ending criterion 2 | 6 |

#### Assessing Density Pattern Preservation in Reproduced Configurations

To assess how SHAPPE preserves density patterns from the reference point pattern, *X*^(*Obs*)^, we consider *N*_*rep*_ generated artificial reconstructions, *X*^(*Art,s*)^ (*s* = 1, ⋯, *N*_*rep*_) and compare the overlap between density grids. Let *X* be a point pattern defined on domain *D* ⊆ [−*R, R*] ⊂ ℝ^2^. Similarly to the environmental constraint step, we divide [−*R, R*] into a grid with bins of length *bL*_*Env*_. For every bin *b* that intersects with *D*, its density, *d*_*b*_, is given by the number of points within its boundaries divided by the area of its intersection with *D*. Since we assume points share the same first-order intensity function, we pool all points together regardless of their types. Therefore, the density distribution is given by

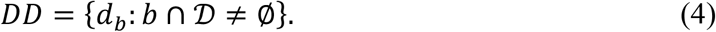

If *D*^(*Obs*)^ and *D*^(*Art*)^ are not of the same size, we use grids with approximately equal bin lengths to build *DD*^(*Obs*)^ and *DD*^(*Art,s*)^. The overlapping index between *DD*^(*Obs*)^ and *DD*^(*Art,s*)^, *ov* (*DD*^(*Obs*)^, *DD*^(*s*)^), measures the density distribution similarity between *X*^(*Obs*)^ and *X*^(*Art,s*)^

(Supplementary Section M8). The dissimilarity in density distributions is thus given by:

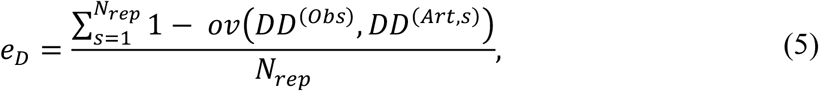

with smaller values of *e*_*D*_ indicating better preservation of density distribution patterns.

#### Assessing Clustering Preservation in Reproduced Configurations

To assess density pattern preservation between *X*^(*Obs*)^ and the artificial reconstructions *X*^(*Art,s*)^, we defined dissimilarity metrics to measure the average difference between 1) pairwise nearest-neighbour distributions, and 2) inter-cell-type clustering patterns at different lengths. For this, we used a pairwise Ripley’s *H*-function, which linearises and stabilises the variance of the *K*-function across different scales^28^, given by

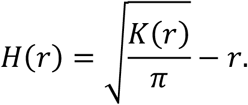

Let 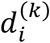 be the distance from point *x*_*i*_ to its closest neighbour of type *k*. Then, the nearest-neighbour distribution from type *g* to *k* is given by

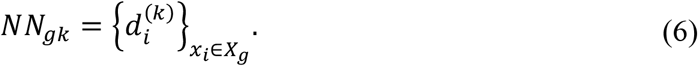

As above, the overlapping index between 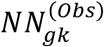 and 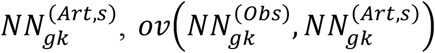,measures the nearest-neighbour similarity between *X*^(*Obs*)^ and *X*^(*Art,s*)^. Accordingly, the dissimilarity in nearest-neighbour distributions is given by:

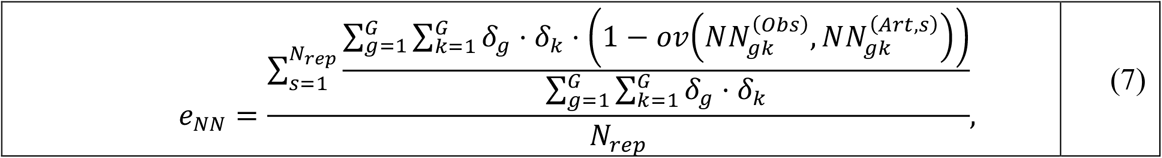

where *δ*_*g*_ (*g* = 1, ⋯, *G*) defines a weight attributed to each type of points. Here, lower values of *e*_*NN*_ indicate better preservation of pairwise nearest-neighbour distributions.

We define the absolute clustering error at distance *r* to be:

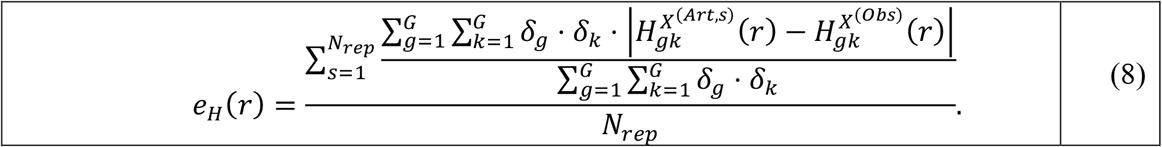

Note that if *X*^(*Obs*)^ is homogeneous, the standard *K*-function is used to estimate the *H*-function. Otherwise, we use the inhomogeneous *K*-function with a moving window to estimate the density function (Supplementary Section M6). The lower the value of *e*_*H*_(*r*), the better SHAPPE preserves pairwise clustering at distance *r*.

A multitype point pattern reconstruction algorithm should also represent observed variability. Thus, we define the variability accuracy score that measures the weighted fraction of pairwise Ripley’s *H*-functions that fall within their corresponding 95% confidence interval bounds over all the *N*_*rep*_ SHAPPE executions:

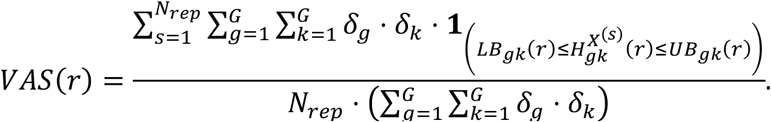

Here, *LB*_*gk*_(*r*) and *UB*_*gk*_(*r*) are the lower bound and upper bounds of the 95% confidence interval of 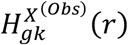, respectively. These are calculated using marked point bootstrapping following the approach by Loh and Stein^29,30^ (Supplementary Section M12).

### Case Study: Tumour Microenvironment Reconstruction from Imaging Mass Cytometry Using SHAPPE

To demonstrate the application of SHAPPE to a heterogeneous set of point patterns, we applied SHAPPE to a reference dataset of IMC images from glioblastoma patients^14^. We previously used this dataset to show that cell-cell interactions on multiple timescales impact the predictive accuracy of agent-based models^19^. In total, we used 87 images of primary glioblastoma or glioma with unknown IDH status from patients who were subsequently treated with standard-of-care post-resection. Each IMC sample is a point pattern, *X*^(*IMC*)^, containing between 486 and 6012 cells classified as tumour, stromal or one of four immune types (CD8+ T cells, M0 TAMs, M1 TAMs and M2 TAMs). Cells are distributed over a square with side length approximately equal to 1 mm. As in Mongeon et al.^19^, microglia and macrophages were grouped under the TAM category. We first reconstructed TMEs of the same size and shape as the observed data and estimated the range of interactions between cells to 50μm^31^, which corresponds to a length scale of two to three cell diameters. We performed SHAPPE ten times for each reference point pattern (i.e., *N*_*rep*_ = 10), with baseline configurations *X*^(*R*)^ generated under CSR for homogeneous samples or ECSR for inhomogeneous samples. We used ECSR with repulsive regions for all but one inhomogeneous IMC point pattern in which cells are visibly organised in a single blob (see SHAPPE Results document on Git hub repository, page 3). For each reconstruction, tumour cells were placed first, followed by stromal cells and immune cell types, in descending order of abundance. Hyperparameters were optimised to minimise the average clustering error at distances *r* = 20, 40 and 60μm across all 87 reference point patterns. Details of the SHAPPE parametrisation and hyperparameter optimisation framework are given in Supplementary Sections M9 and M10. Preliminary steps were performed on an 8-core Apple M3 processor and 16 GB of RAM. Reconstructions (Steps 1-6) were run on the Rorqual cluster (Calcul Québec/Digital Research Alliance of Canada) and each used a single CPU core.

### Using SHAPPE to Initialise Agent-Based Models

We evaluated SHAPPE’s ability to reconstruct larger TME regions of interest by generating ten artificial point patterns per IMC point pattern, each with *N*^(*Art*)^ = 10,000 cells distributed on a circular domain, representing a 1.66 to 20.58 fold-increase in the total number of cells. Synthetic domain radii were calculated to preserve total cell density. Next, we sought to evaluate the differences in ABM predictions depending on whether initial conditions capture first- and second-order structures from the data. For this, we constructed an ABM based on our previous ODE model describing glioblastoma growth and interactions between tumour and immune cells^32^ (Figure 5A). In this model, tumour cells proliferate following logistic growth. Immune cells, including TAMs (innate immune cells) and CD8+ T cells (adaptive immune cells), all die naturally at type-specific death rates. TAMs can be resident (M0) or polarised with the anti-(M1) or pro-tumoural (M2) phenotype. Polarised TAMs can switch phenotype, and M0 TAMs can become activated when they encounter dead tumour cells or polarised TAMs in their local neighborhoods (defined to be the disk of radius 50μm around each cell). TAMs are recruited to the TME at a tumour-mediated rate and recruit CD8+ T cells at a rate dependent on the M2-to-M1 ratio and the PD-1 and PD-L1 receptor concentrations in the TME. The PD-1/PD-L1 signalling axis is an important immunosuppressive mechanism in tumours^33^. CD8+ T cells also proliferate at a tumour- and PD-1/PD-1-mediated rate and can kill tumour cells in their neighborhood. Finally, stromal cells, i.e., connective tissue cells that occupy space in the TME, are recruited to the TME as its radius increases. We defined two types of cell movement in our ABM: 1) random cell movement and 2) spring cell movement, both of which are analogous to how we place points in SHAPPE. In both cases, if two cells become too close, they are repulsed from one another. Further details of the ABM are provided in Supplementary Section M15.

We selected an inhomogeneous IMC sample with a significant negative spatial correlation between tumour and CD8+ T cells, representing less exposure between tumour cells to CD8+ T cells than what is expected under randomness. We then performed three in silico experiments with our ABM:

1. Data-derived experiment: ABM simulated three times per synthetic SHAPPE reconstruction (i.e., 30 simulations) with cells moving only according to spring movement. In this experiment, we considered cells to also be repulsed from ECSR voids defined during the TME reconstructions.
2. ECSR experiment: Same experiment as the data-derived experiment, but with the ABM initialised with ECSR configurations (three runs per initial conditions).
3. Well-mixed model: We generated ten configurations under CSR and used them to initialise our ABM (three runs per configuration) with random cell movement, without repulsion from repulsive regions. This experiment mimics a well-mixed assumption.

In each experiment, we measured the predicted tumour growth over 50 days (i.e., tumour fold, or the number of tumour cells at time *t* divided by the initial number of cells) and the cumulative number of tumour cells killed by CD8+ T cells.

## Results

### Case study: SHAPPE Efficiently Reconstructs Tumour Microenvironments from Imaging Mass Cytometry

We used publicly available IMC data of tumour resections from untreated glioblastoma patients reported in Karimi et al.^14^ (*n* = 87) to demonstrate the application of SHAPPE to a heterogeneous set of point patterns and illustrate its capacities. Each IMC data sample is a reference point pattern *X*^(*IMC*)^. Running Step 0.1 for each sample took a median of 21.63 seconds (interquartile range (IQR): 14.25–41.39), with 69 of the 87 IMC data samples determined to be inhomogeneous, consistent with across- and within-patient heterogeneity in cell density^34^. We estimated that SHAPPE accurately identified inhomogeneity most likely associated with environmental heterogeneity approximately 89% of the time (Supplementary Section R1). Identifying significant spatial correlations (Step 0.2) took a median of 38.61 seconds (IQR: 16.18–90.53) per sample. Additional details on hyperparameter optimisation results and execution times are provided in Supplementary Section R3. Our results showed significant spatial tumour cell autocorrelation and correlations between tumour and immune cells, with over 83% of significant spatial patterns between tumour and immune cells found to be dispersion patterns (Figure S9, two-sided absolute deviation test, *α* = 0.05). While we note that this result does not necessarily uncover a specific inter-cell-type mechanism behind observed repulsion patterns, it nonetheless indicates that tumour cells tend to have fewer immune cells in their neighborhoods than what is observed under a random labelling null. This result thus suggests that ABMs initialised without reconstruction of clustering patterns might overestimate the frequency, and therefore the outcome, of tumour-immune interactions.

### SHAPPE Preserves Density Patterns

From these reconstructions, we next assessed the reliability with which SHAPPE reproduced observed first-order structures in density patterns. The median *e*_*D*_ (i.e., the dissimilarity in density distributions, Eq. (5)) score across all 87 IMC samples was found to be 0.0605 (IQR: 0.0486–0.0780; Figure S12). This implies that for 75% of the reference point patterns, the average overlapping area between the synthetic and reference density distributions was at least 92.2%. Dissimilarity in density distributions remained mostly unchanged at around 89% if we removed repulsion from exclusion zones or attraction to inclusion zones from Step 4.1. However, including these additional environmental constraints was found to better reproduced the size and shape of low-density enclosed regions (Figure 3C). Inhomogeneity identification on synthetic reproductions confirmed that SHAPPE successfully recapitulates the homogeneity/inhomogeneity status of the synthetic point patterns in 88.85% of cases (Figure 3B). To validate the performance of SHAPPE, we compared it to a random baseline, finding a median relative change in *e*_*D*_ (Eq. (S9)) of 53.48% (IQR: 39.55–69.86, Figure 3A, Supplementary Section M14). Overall, these results show that SHAPPE successfully reproduces first-order structures.

**Figure 3.**
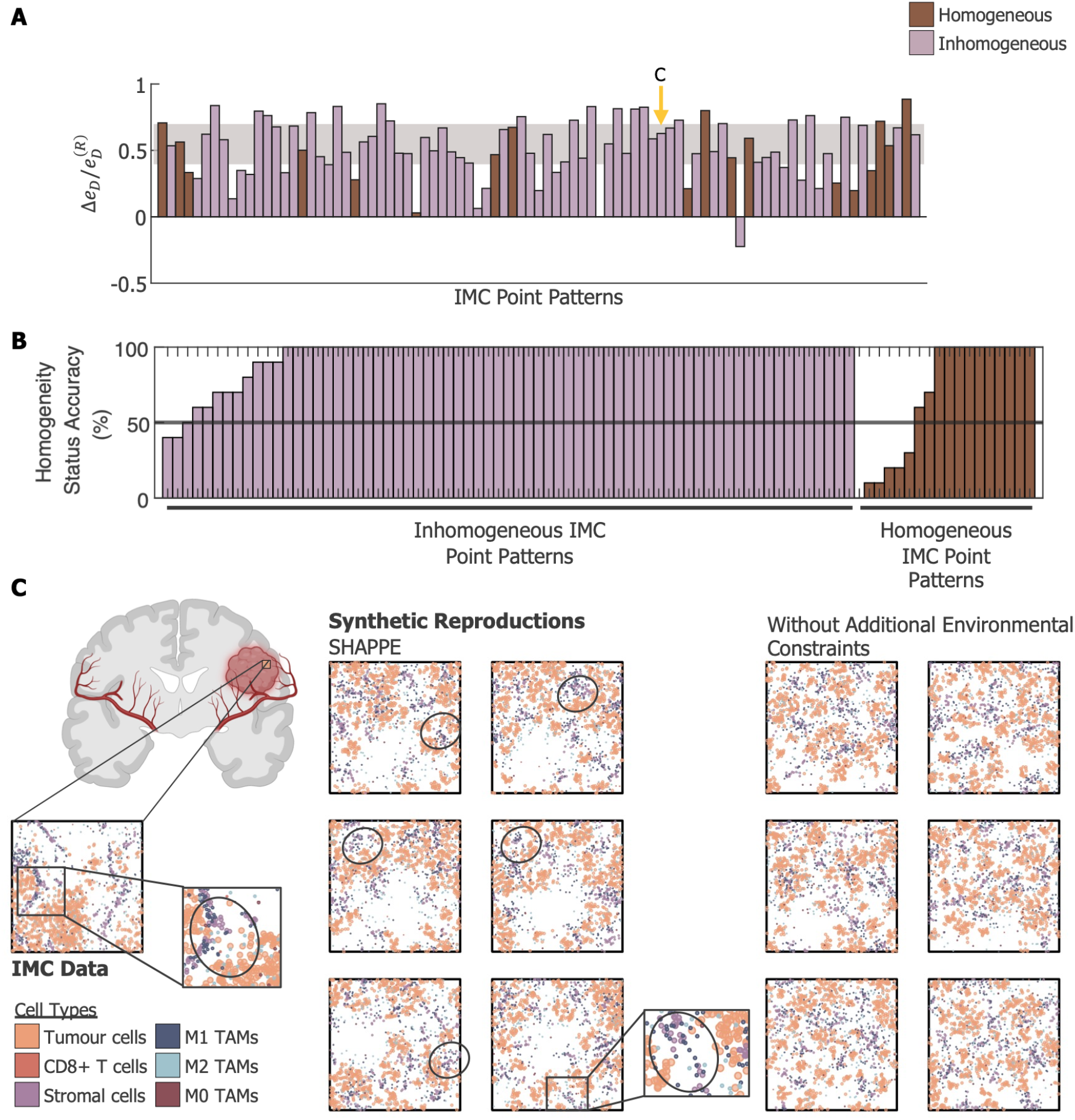
SHAPPE outperforms random point placement at reproducing density patterns. **A**. Relative change in the average dissimilarity with density distributions, *e*_*D*_, with random initialisation. The random baseline was CSR for homogeneous reference point patterns (brown, *n*_*samples*_ = 18) and ECSR for inhomogeneous reference point patterns (pink, *n*_*samples*_ = 69). The interquartile range (gray shaded region) lies almost entirely over 0.5, indicating that SHAPPE decreases the dissimilarity in density distributions obtained with a random point placement (CSR or ECSR) by over 50% in the majority of cases. **B**. For each reference point pattern, we classified its SHAPPE reproduction as either homogeneous of inhomogeneous and calculated the number of times SHAPPE reproduced the same homogeneity status as the reference point pattern. SHAPPE accurately (> 88%) reproduced estimated homogeneity status. **C**. Six randomly selected synthetic configurations obtained for an inhomogeneous reference point pattern (yellow arrow in panel A). SHAPPE reproduces clustering patterns such as the presence of stromal cells and M1/M2 TAMs outside of regions with higher tumour cell densities (black ellipses). With the removal of additional environmental constraints (i.e., introduction of repulsive exclusion zones), clustering patterns were preserved but a visual inspection of the reproductions suggests a loss of the area of lower density regions.

### SHAPPE Preserves Second-Order Structures

Given the observation of consistent density pattern reproduction, we next assessed the efficacy with which SHAPPE reproduced clustering patterns. For this, we first measured the dissimilarity in nearest-neighbour distributions, *e*_*NN*_ (Eq. (7)), for all 87 IMC point patterns. Cell types *g* with at least 15 cells were attributed an equal weight of *δ*_*g*_ = 1, while others had weight *δ*_*g*_ = 0. Overall, the median *e*_*NN*_ score was 0.1964 (IQR: 0.1728–0.2172, Figure S13), translating to an average overlapping area of at least 78.28% for 75% of the tested point patterns (overlap measured between synthetic and reference nearest-neighbour distributions across ten runs of SHAPPE). This implies that SHAPPE is nearest-neighbour-preserving. We further found a median relative change from random baseline (Eq. (S7)) of 18.27% (IQR: 9.91–30.92; Figure 4A), indicating that SHAPPE outperforms a random initialisation when reproducing nearest-neighbour distributions.

**Figure 4.**
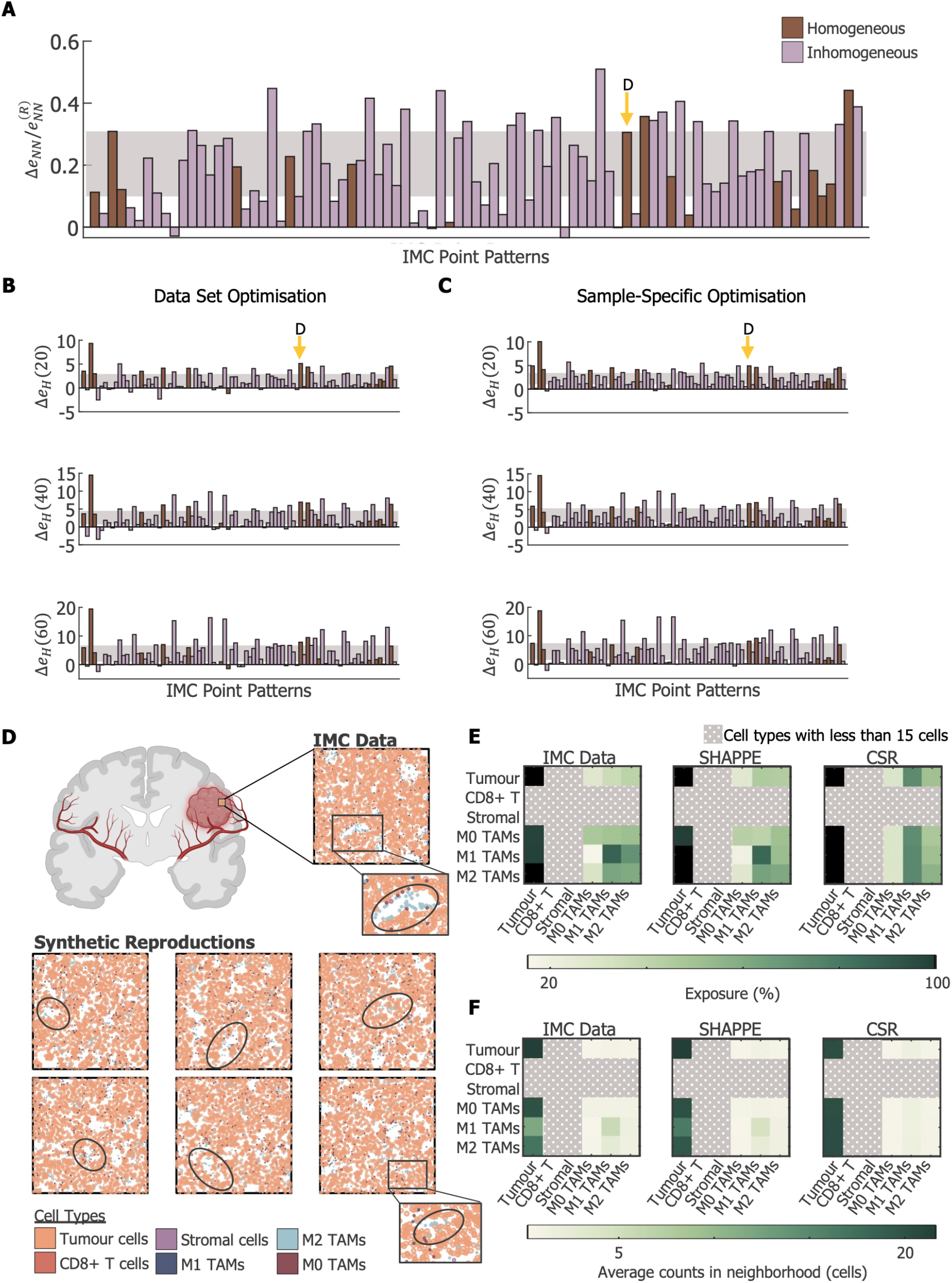
SHAPPE outperforms spatial randomness at reproducing clustering patterns. **A**. Relative change in the dissimilarity with nearest-neighbour distributions, *e*_*NN*_, with random initialisation. The median relative change and its interquartile range (gray shaded region) lie entirely over 0, demonstrating that nearest-neighbour distributions are overall better preserved with SHAPPE than with a random initialisation. **B-C**. Change in the absolute clustering error, *e*_*H*_(*r*), at three distances (*r* = 20, 40, and 60μm) with the hyperparameter set optimised over B. all reference point patterns or C) hyperparameter sets optimised for each reference point pattern. **D**. Six representative synthetic configurations obtained for a homogeneous case, indicated by a yellow arrow in A, B and C. SHAPPE visually reproduces clustering patterns such as the presence of M2 TAMs near lower-density region (black ellipses). **E-F**. Exposure matrices for the reference point pattern shown in panel C, with SHAPPE reproductions and baseline CSR configurations. **E**. SHAPPE better reproduced the exposure status inter-cell-types as well as **F**. the exposure level quantified with the average cell counts in the neighbourhood of a cell. Here, the neighbourhood of a cell was defined as the disk of radius 50μm around a cell.

We also evaluated whether SHAPPE can reproduce pairwise clustering patterns at three interaction-relevant lengths. For this, we calculated the average absolute clustering error, *e*_*H*_(*r*), at distances *r* = 20, 40, and 60 μm, as compared to the random baseline. We found an unnormalised median change in absolute clustering error (Δ*e*_*H*_(*r*), Eq. (S6)) of 1.2214 (IQR: 0.2912–2.9235), 1.9419 (IQR: 0.6727–4.4948), and 3.6810 (IQR: 0.9565–6.7069) for *r* = 20, 40, and 60 μm, respectively (Figure 4B). As with the nearest-neighbour distribution reproduction, these results imply that SHAPPE reproduces pairwise clustering patterns at interaction-relevant scales better than random configurations. For example, SHAPPE successfully recovered clustering of TAMs outside of high tumour cell density zones for point patterns in which tumour cells were found to be significantly negatively correlated with TAMs, whereas this second-order structure was lost in corresponding ECSR/CSR configurations (Figure 4D and Figure 3C). SHAPPE also reproduced exposure status (*e*_Π_) and exposure levels (*e*_Σ_) more accurately than the ECSR/CSR random baselines. Briefly, *e*_Π_ and *e*_Σ_ sum, over all pairs of cell types with at least 15 cells, the absolute differences in the proportion of cells of one type with at least one cell of the other type in its neighbourhood, and in the average number of cells of the other type within the same neighbourhood, respectively, averaged over all replicates (Supplementary Section M13). Median errors were 1.41 vs 2.42 (*e*_Π_) and 11.66 cells vs 19.17 cells (*e*_Σ_) for SHAPPE and the random baselines (Figure 4E-F and Figure S20; IQRs in Supplementary Section R4), corresponding to median relative changes with random baselines of 36.49% (*e*_Π_, IQR: 23.15–49.41) and 33.67% (*e*_Σ_, IQR: 21.60–50.87). Interestingly, clustering reproduction performance was only slightly increased when using point pattern-specific optimised hyperparameters (Figure 4C, Supplementary Section M10).

In addition to recapitulating nearest-neighbour and pairwise interactions, capturing the variability in second-order structures observed in a real-life point pattern is essential for initialising ABMs from point pattern reconstructions. Thus, for each reference point pattern, we calculated the variability accuracy at distances 20, 40 and 60μm, i.e., the fraction of pairwise Ripley’s *H*-functions that fall between the 95% confidence interval bounds for cell types satisfying the minimal number of cells threshold. While the random baseline had a median variability accuracy of 53.53% (IQR: 43.81–64.78) at 20μm, it plateaus to approximately 60% at 40μm (IQR: 49.53–71.97) and 60μm (IQR: 50.30–74.60), respectively (Supplementary Section R5). This suggests that ECR/CSR hardly captures longer range clustering patterns and their variability at larger scales. In contrast, SHAPPE had median variability accuracy scores of 67.56% (IQR: 60.90–76.40), 77.36% (IQR: 72.13–84.30) and 82.33% (IQR: 77.63–87.94) at 20, 40, and 60μm, respectively, demonstrating its preserving structures at both short and long ranges.

### Initialising ABM of Untreated Glioblastoma Growth using SHAPPE Recovers Tumour Aggressivity

Next, we evaluated how SHAPPE performed at generating synthetic point patterns of larger size and different domain shape than reference data. For this, we generated 10 synthetic configurations with 10,000 cells distributed on a disk (the radius of which was calculated to preserve total cell density) for each IMC point pattern. Amongst the 870 synthetic point patterns, we found that SHAPPE performed equally well as same size and domain reconstructions (Figures S11-17), requiring an average execution time of 7.88 minutes. Then, to assess the importance of initialising ABMs with realistic configurations, we evaluated the difference in predicted frequencies of interactions between two cell types. For this, we used an ABM of untreated glioblastoma growth (Figure 5A) derived from a previous ODE model (see Methods). The model includes spatial-dependant mechanisms such as CD8+ T cells being able to kill solely tumour cells that are in their local neighborhood. To estimate the frequency of interactions between tumour and CD8+ T cells, we calculated the cumulative number of tumour cells killed by these immune cells. We selected an inhomogeneous point pattern with a significant (*p*-value: 0.0015) negative spatial correlation between tumour and CD8+ T cells (see Supplementary SHAPPE Results, pages 141-142).

**Figure 5.**
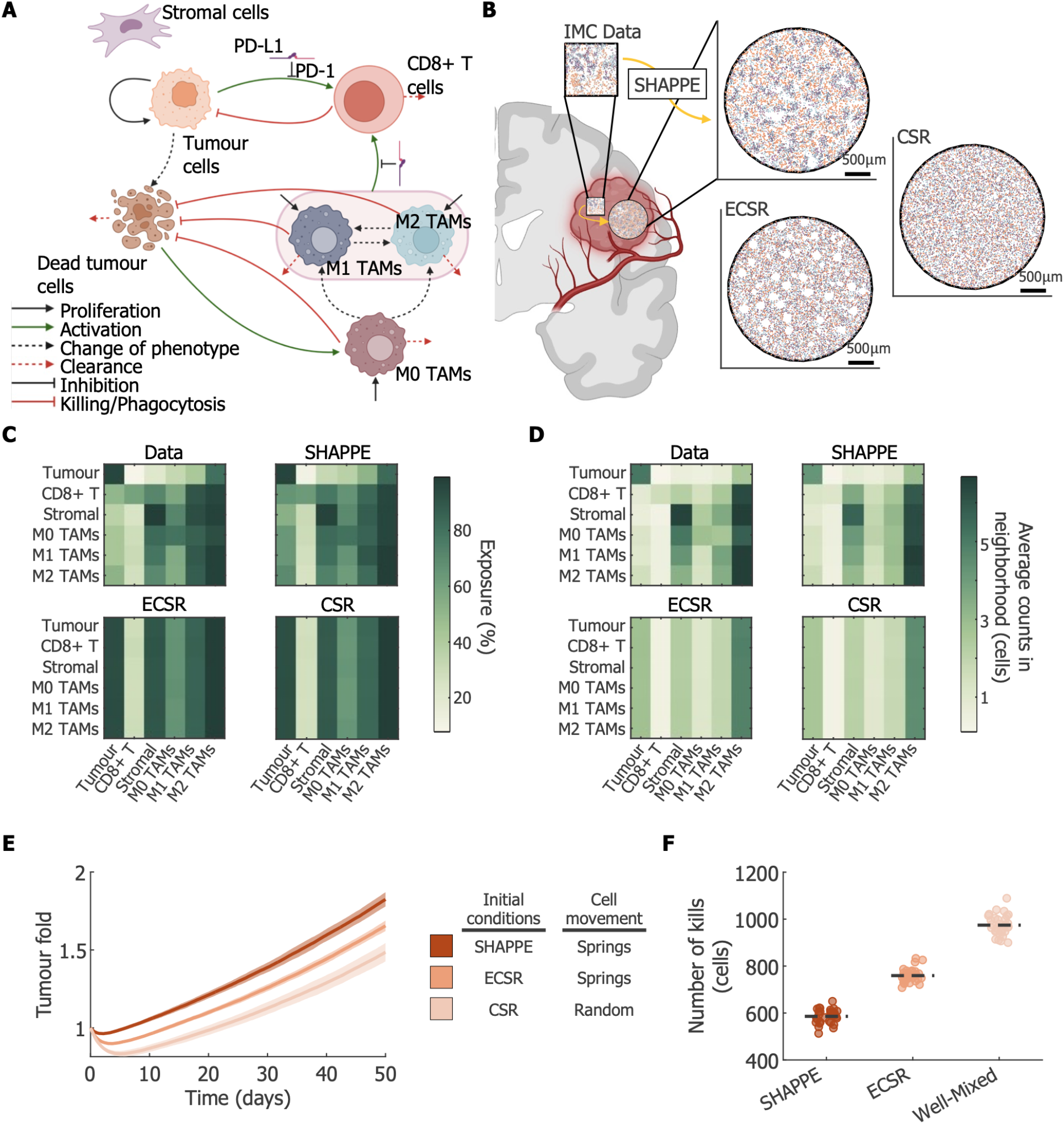
Agent-based modeling of untreated glioblastoma growth over 50 days with inhomogeneous TME reproductions generated using SHAPPE. **A**. Schematic of our ABM describing untreated glioblastoma growth, tumour cell proliferation, and tumour-immune cell (CD8+ T cell and TAM) interactions. **B**. TME visualisations for an inhomogeneous IMC sample and its corresponding SHAPPE, ECSR, and CSR reconstructions. All the visualisations share the same scale. **C-D**. Exposure matrices for the reference point pattern shown in panel B, with SHAPPE reproductions and baseline ECSR and CSR configurations. **C**. SHAPPE better reproduced the exposure status inter-cell-types as well as **D**. the exposure level quantified with the average cell counts in the neighbourhood of a cell. Here, the neighbourhood of a cell was defined as the disk of radius 50μm around a cell. **E**. Fold-increase in tumour cell count and **F**. cumulative number of tumour cells killed by CD8+ T cells over the 50 simulated days for the SHAPPE (dark orange), ECSR (medium orange) and well-mixed (light orange) scenarios. Solid lines in **E**. show the average over the 30 ABM runs (10 different initial configurations, 3 replicates each) and shaded area indicate the standard deviation. Dashed lines in **F**. indicate the average cumulative number over the 30 runs.

We simulated our model over 50 days under three scenarios (Figure 5B): 1) ABM initialised with SHAPPE configurations and with cells moving under spring-like forces, analogous to the point placement in Step 4, 2) ABM initialised with ECSR configurations and the same spring mechanisms for cell movement, or 3) the well-mixed assumption (i.e., ABM initialised with CSR configurations with cells moving around randomly with an exclusion distance to prevent overlap); see Methods for full details. Initial conditions contained 2108 and 256 tumour and CD8+ T cells, respectively. In the original sample, only 7.80% of tumour cells had CD8+ T cells in their neighbourhoods. While SHAPPE slightly increased this exposure (10.89%), random initial conditions overestimated it by approximately 3.5 times (CSR: 27.08%, ECSR: 28.78%) (Figure 5C-D). This difference translates to a 1.66-fold (Well-mixed) and 1.30-fold (ECSR) increase in the average number of tumour cells killed by CD8+ T cells over 50 days, relative to SHAPPE (Figure 5F). Although ECSR slightly overestimated exposure more than CSR initial conditions, it produced the smallest increase in cytotoxic activity. This may occur due to the spring mechanism for cell movement, in which target distances are sampled from the data. Overall, after 50 days, the well-mixed and ECSR scenarios yielded final tumour cell counts that were 18.6% and 9.3% lower on average than under SHAPPE initialisation (Figure 5E). Hence, initialising our ABM with realistic configurations obtained with SHAPPE captured the aggressive, immune-evasive behaviour characteristic of glioblastoma.

## Discussion

Agent-based models offer a flexible framework to study how spatial structures shape biological processes. However, their predictions depend on a variety of factors, including how they are initialised. High-resolution imaging techniques, such as imaging mass cytometry, give insight into spatial arrangements within a region of interest, but their use is costly^16^ and the regions they capture are often too small to adequately initialise an ABM^19^. Thus, synthetic reconstructions of spatial arrangements have been proposed as an alternative^22^. In this vein, we developed SHAPPE, a sequential point placing algorithm that moves points one type at a time, thereby limiting the chasing of moving targets.

To develop SHAPPE, we considered multitype point patterns originating from biological processes that give rise to environmental constraints that create first-order inhomogeneity (e.g., resource availability, blood vessels, etc.). Hence, SHAPPE begins by identifying and quantifying empty regions. Our results show that preliminary steps in SHAPPE effectively identify inhomogeneity arising from changes in the first-order intensity function rather than from clustering (Supplementary Section R1). Further, without an accurate estimate of the first-order intensity function, density patterns in inhomogeneous cases can be difficult to preserve with existing point pattern reconstruction methods that mostly rely on minimising the difference in spatial statistics between a synthetic and a reference point pattern^23^. Thus, we implemented additional constraints by including repulsion or attraction to exclusion or inclusion zones, respectively, to account for inhomogeneity. Integrating these constraints allows for the preservation of density distributions (Figure 3C), making SHAPPE a good fit for both homogeneous and inhomogeneous multitype point pattern reconstruction.

To evaluate how well SHAPPE reconstructs point patterns, we ran it on 87 IMC slides from untreated glioblastoma patients^14^ and compared its outcomes to complete spatial randomness (CSR) and environmentally constrained spatial randomness (ECSR) for homogeneous (18/87) and inhomogeneous (69/87) reference point patterns, respectively. Because ECSR accounts for first-order inhomogeneity by sampling predefined low-density regions, the random baseline density distributions already overlap with about 80% of reference patterns, on average (Figure S12). SHAPPE nonetheless increased the degree of overlap of dense regions to almost 93% (Figure 3A and S12) and reproduced low-density regions with more realistic shapes (Figure 3C). Despite spatial statistics not being directly minimised during the point placement process, our results reveal that SHAPPE recovered most pairwise clustering trends (Figure 4); inter-cell-type exposure status and levels also reconstructed better with SHAPPE than with the random baselines (Figure 4E-F and S14-18). Together, these results suggest that SHAPPE accurately generates synthetic spatial arrangements with similar spatial characteristics to a reference point pattern, without requiring repeated evaluation of second-order statistics.

We also applied SHAPPE to generate synthetic configurations of the glioblastoma tumour microenvironment that were larger and differently shaped than the reference IMC samples and found it to accurately reconstruct these configurations, even when the number of points was increased up to 20-fold (Figure S12-18). We then initialised an ABM of untreated glioblastoma growth using reconstructions of an inhomogeneous TME with significant dispersion between tumour and CD8+ T cells (Figure 5C-D) and simulated it over 50 days. Compared to predictions from initialisations under ECSR and the well-mixed assumption, our ABM predicted less CD8+ T cell cytotoxic activity and faster tumour growth when initialised with data-derived inter-cell-type exposure levels (Figure 5E-F). This result is consistent with the known aggressiveness and immune-evasive nature of glioblastomas.

In SHAPPE, we considered first-order structures (i.e., the previously mentioned environmental constraints) to be shared by all point types. This simplifying assumption is motivated by the updating of the density grids used to enforce environmental constraints (Step 5), which should capture most of the differences in intensity functions between point types. However, some point patterns (e.g., vertical patches of distinct point types) may be harder to recover. Future work could refine our definition of first-order inhomogeneity to make it specific to each point type. Future work could also extend SHAPPE to multi-trait point patterns (i.e., where a point is assigned multiple attributes described by a vector of measurements) by sampling measurements from data-driven distributions and making target distances dependent on a neighbour’s type and attributes.

Altogether, our SHAPPE framework addresses a practical bottleneck in agent-based modelling: while high-resolution imaging techniques detail real spatial arrangements arising from biological processes, they do not provide enough data to initialise ABMs. By reproducing both first-order density structure and second-order clustering, SHAPPE generates configurations of arbitrary size and shape that retain the spatial characteristics of the reference sample. Thus, our framework shows the potential to ease the integration of high-resolution imaging techniques into computational biology ABMs. Our glioblastoma application further underlines the importance of using data-derived synthetic reconstructions to initialise spatial models relying on interactions between agents, such as an ABM studying an immunotherapy. Our results show that predicted tumour-immune interaction frequencies depend on the initial configuration. Hence, what a model says about the potential of a therapy could depend on a configuration chosen before the simulation starts. For example, if one is using an ABM to study a treatment targeting the reactivation of cytotoxic immune-tumour interactions (e.g., anti-PD-1 immune checkpoint inhibition), then overestimating the frequency of these interactions could possibly lead to an erroneous overestimation of the efficacy of the therapy. The initial configuration should be treated as a modelling assumption that needs justification, and reconstruction methods like SHAPPE offer a way to ground it in data. Further, though we assessed the efficacy of SHAPPE on IMC-derived spatial configurations of glioblastoma microenvironments, nothing but the parametrisation in SHAPPE is context-dependent. This makes SHAPPE applicable to multitype point patterns originating from various biological processes, for example, the reconstruction of spatial arrangements in forests with tree species treated as types and repulsion/attraction between trees encoding competition or similar environmental needs. Hence, SHAPPE has broad implications for our understanding of a variety of spatially interacting systems and our ability to accurately predict their temporal behaviour through agent-based modelling.

## Supporting information

Supplementary Methods and Results

Supplementary SHAPPE Results

## Acknowledgements

Computations were made on the supercomputer Rorqual, managed by Calcul Québec and the Digital Research Alliance of Canada (Alliance). The operation of this supercomputer is funded by the Alliance, le ministère de l’Économie, de l’Innovation et de l’Énergie du Québec (MEIE) and le Fonds de recherche du Québec (FRQ). The authors wish to acknowledge helpful conversations and collaborations with Dr. Daniela Quail and Dr. Logan Walsh that led to this work.

## Availability of data and material

The entire computational code for SHAPPE, a user-guide, the datasets resulting from the IMC glioblastoma TME reconstructions and ABM simulations are available on Github: https://github.com/Craig-Lab/SHAPPE. Any further datasets are available from the corresponding authors upon reasonable request.

## Author’s Contributions

BM: Participated in research design, conducted experiments, contributed new analytic tools, performed data analysis, wrote the manuscript. MC: Participated in research design, performed data analysis, wrote the manuscript, secured funding, supervised study.

