## Supplementary Methods and Results for "SHAPPEing agent-based models to spatial data: Sequential Hooke’s law Algorithm for Point Pattern reconstruction with Environmental constraints"

### Mathematical Abbreviations and Functions

| Abbreviation | Definition |
| --- | --- |
| $X$ | Point pattern  $X=\left\{ \left( x_{i},g_{i} \right) \right\}_{i=1,\cdots,N}$ |
| $N$ | Total number of points in $X$, i.e. $N=\left\vert X \right\vert$ |
| $\mathcal{D}$ | Spatial domain on which $X$ is defined, i.e., $\mathcal{D\subseteq}\left[ -R,R \right]\times\left[ -R,R \right]\subset\mathbb{R}^{2}$ |
| $G$ | Number of distinct categorical types in $X$ |
| $X_{g}$ | Set of points of type $g$, i.e. $X_{g}=\left\{ \left( x_{i},g_{i} \right)\in X:g_{i}=g \right\}$ |
| $N_{g}$ | Number of points of type $g$ in $X$, i.e. $N_{g}=\left\vert X_{g} \right\vert$ |
| $X^{\left( Obs \right)}$ | Reference point pattern. Superscript $\left( Obs \right)$ notation refers to the attributes of a reference point pattern $X^{\left( Obs \right)}$. |
| $X^{\left( Art \right)}$ | Artificial point pattern reconstruction from $X^{\left( Obs \right)}$. Superscript $\left( Art \right)$ notation refers to the attributes of an artificial point pattern $X^{\left( Art \right)}$. |
| $X^{\left( R \right)}$ | Random baseline configuration for a reference point pattern $X^{\left( Obs \right)}$. $X^{\left( R \right)}$ follows complete spatial randomness if $X^{\left( Obs \right)}$ is homogeneous, or environmentally constrained spatial randomness (with attractive or repulsive regions) if $X^{\left( Obs \right)}$ is inhomogeneous. |
| $h$ | Exclusion distance in a point pattern $X$, i.e., minimal distance between two points. |
| $A_{0}$ | Minimal area for an empty region to be considered a zero-density region. |
| $\mathcal{R}$ | Set of distances on which a spatial statistic (e.g., Ripleys’ $K$- or $F$- functions) is evaluated. $\mathcal{R=}\left[ r_{min},r_{max} \right]$. |
| $\Omega_{gk}$ | Indication of whether type $g$ is significantly spatially correlated to type $k$ in a reference point pattern $X^{\left( Obs \right)}$. $\Omega_{gk}=1$ if $K_{gk}^{\left( Obs \right)}$ is significantly outside of its random labelling envelope. Otherwise, $\Omega_{gk}=0$. |
| $X_{k}^{\left( Ghost \right)}$ | Set of ghost points of type $k$ in a point pattern reconstruction $X$. |
| $p_{i}\left( n \right)$ | Position of point $x_{i}$ at iteration $n$ |
| $p_{i}^{\left( k,l \right)}\left( n \right)$ | Position of point $x_{i}$’s $l$-th closest neighbour of type $k$ among $X_{k}\cup X_{k}^{\left( Ghost \right)}$. |
| $s_{i}^{\left( k,l \right)}$ | Target distance between a point $x_{i}\in X$ and its $l$-th neighbour of type $k$ |
| $S_{i}^{\left( k,l \right)}\left( n \right)$ | Interaction force between point $x_{i}$ and its $l$-th neighbour of type $k$ at iteration $n$. Calculated from Eq. (1) (Main text). |
| $d_{i}\left( n \right)$ | Raw displacement of point $x_{i}$ at iteration $n$. Calculated from Eq. (2) (Main text). |
| $E^{\left( Obs,g \right)}$ | Density grid built from $X^{\left( Obs \right)}$ and considering points of types $1,\cdots,g$, with $g\leq G$. |
| $E^{\left( n,g \right)}$ | Density grid built from the positions of points at iteration $n$, when placing points of type $g$ to construct $X^{\left( Art \right)}$. Similarly to $E^{\left( Obs,g \right)}$, only points of types $1,\cdots,g$, with $g\leq G$, are considered. |
| $N_{rep}$ | Number of synthetic reconstructions for $X^{\left( Obs \right)}$. |
| $DD^{\left( \cdot\right)}$ | Estimated density distribution obtained from the point pattern indicated by the superscript; ${DD}^{\left( Obs \right)}$ and ${DD}^{\left( Art \right)}$ represent the density distributions from a reference point pattern $X^{\left( Obs \right)}$ and its corresponding reconstruction $X^{\left( Art \right)}$, respectively. |
| $b$ | Bin in a density grid, with density given by $d_{b}$. |
| $d_{i}^{\left( k \right)}$ | Distance from point $x_{i}$ to its closest neighbour of type $k$. |
| $NN_{gk}^{\left( \cdot\right)}$ | Nearest-neighbour distribution from type $g$ to $k$ in point pattern indicated by superscript, i.e., $NN_{gk}=\left\{ d_{i}^{\left( k \right)} \right\}_{x_{i}\in X_{g}}$ (Eq. (6), Main text). $NN_{gk}^{\left( Obs \right)}$ and $NN_{gk}^{\left( Art \right)}$ represent the nearest-neighbour distributions from a reference point pattern $X^{\left( Obs \right)}$ and its corresponding reconstruction $X^{\left( Art \right)}$, respectively. |
| $\delta_{g}$ | Weight attributed to type $g$ ($g\in\left\{ 1,\cdots,G \right\}$). |
| $e_{D}$ | Dissimilarity in density distributions between $X^{\left( Obs \right)}$ and $N_{rep}$ synthetic reconstructions, calculated from Eq. (5) (Main text). |
| $e_{D}^{\left( R \right)}$ | Dissimilarity in density distributions between $X^{\left( Obs \right)}$ and $N_{rep}$ random baseline reconstructions. |
| $e_{NN}$ | Dissimilarity in nearest-neighbour distributions between $X^{\left( Obs \right)}$ and $N_{rep}$ synthetic reconstructions, calculated following Eq. (7) (Main text). |
| $e_{NN}^{\left( R \right)}$ | Dissimilarity in nearest-neighbour distributions between $X^{\left( Obs \right)}$ and $N_{rep}$ random baseline reconstructions. |
| $e_{H}\left( r \right)$ | Absolute clustering error at distance $r$ between $X^{\left( Obs \right)}$ and $N_{rep}$ synthetic reconstructions, calculated following Eq. (8) (Main text). |
| $e_{H}^{\left( R \right)}\left( r \right)$ | Absolute clustering error at distance $r$ between $X^{\left( Obs \right)}$ and $N_{rep}$ random baseline reconstructions. |
| $H_{gk}^{X}\left( r \right)$ | $H_{gk}\left( r \right)$ (Table S2) evaluated on point pattern $X$. |

Table S1. List of abbreviations used in the Main Text and Supplementary Information.

| Function | Definition |
| --- | --- |
| $\begin{matrix} A: & P\left( \mathbb{R}^{2} \right) & \longrightarrow& \mathbb{R}^{+} \\ & \mathcal{D} & ⟼ & \mathrm{area}\left( \mathcal{D} \right) \end{matrix}$ | Area function: $A\left( x \right)$ is the area of $x$, where $x\subset\mathbb{R}^{2}$. $P\left( S \right)$ is the power set of $S$, i.e. the set of all subsets of $S$. |
| $\begin{matrix} F: & \mathbb{R}^{+} & \longrightarrow& \mathbb{R}^{+} \\ & r & ⟼ & \mathbb{P}\left[ d\leq r \right] \end{matrix}$ | Empty-space function for a point pattern $X$ defined on a domain $\mathcal{D}$  Let $d$ be the distance from an arbitrary location within $\mathcal{D}$ to the nearest point of $X$. Then $F\left( x \right)$ is the probability that $d$ is smaller or equal to $x$. |
| $\begin{matrix} K_{gk}: & \mathbb{R}^{+} & \longrightarrow& \mathbb{R}^{+} \\ & x & ⟼ & K_{gk}\left( r \right) \end{matrix}$ | Univariate ($g=k$) or multivariate ($g\neq k$) Ripley’s $K$-function for a point pattern $X$ defined on a domain $\mathcal{D}$  $K_{gk}\left( x \right)$ is defined by the expected number of points of type $k$ within a distance $x$ of a typical point of type $g$, divided by the intensity of type $k$. Formal definitions are given in Eqs. S3, S4 and S5. |
| $\begin{matrix} H_{gk}: & \mathbb{R}^{+} & \longrightarrow& \mathbb{R}^{+} \\ & r & ⟼ & \sqrt{\frac{K_{gk}\left( r \right)}{\pi}}-r \end{matrix}$ | $H$-transformation of Ripley’s $K$-function. |
| $\begin{matrix} ov: & P\left( \mathbb{R} \right)\times P\left( \mathbb{R} \right) & \longrightarrow& \left[ 0,1 \right] \\ & \left( \Phi_{1},\Phi_{2} \right) & ⟼ & \int_{-\infty}^{\infty} \min\left( f_{1}\left( \theta\right),f_{2}\left( \theta\right) \right)d\theta\end{matrix}$ | Overlap function  $\Phi_{1}$ and $\Phi_{2}$ are two distributions defined on $\mathbb{R}$ with probability density functions $f_{1}$ and $f_{2}$. |

Table S2. List of recurring functions in the Main Text and Supplementary Information.

### Supplementary Methods

M1. Generating Complete Spatial Randomness

We define complete spatial randomness (CSR) as the Gibbs hard-core point process with exclusion distance $h$, i.e., the conditional Poisson process where every point is at least $h$ distance-units from its closest neighbour. $h$ can be inferred from the nature of the points (e.g., minimal realistic distance between two trees in a forest) or can be set to $0$ if there is no prior knowledge. In the latter case, our CSR definition reduces to a traditional Poisson process. To generate a configuration with $N$ points under CSR within a domain $\mathcal{D}$, we first sample a position $p_{i}$ for the $i$-th point $x_{i}$ following a uniform distribution on $\mathcal{D}$. Then, if $i>1$, the acceptance criterion for $p_{i}$ depends on its distance to points $x_{1},\cdots,x_{i-1}$; if another point is less than $h$ units from $p_{i}$, we reject $p_{i}$ and resample another position. Otherwise, we fix $x_{i}$ at position $p_{i}$ and we move on to $x_{i+1}$ (Figure S1A). If no valid position for $x_{i}$ is found within 1,000 resampling tries, we accept the best position out of the 1,000 tried positions to avoid infinitely looping. The best position is defined as the position with the largest distance to the closest neighbour. CSR serves as the random baseline for homogeneous point patterns (Figure S1B).

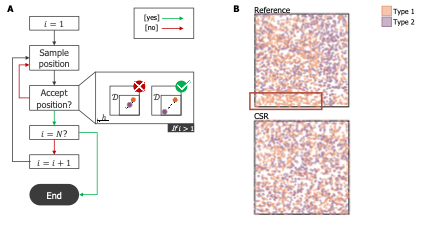

Figure S1. Generating Complete Spatial Randomness (CSR) for homogeneous point processes.

**A.** Points are placed sequentially. When placing a point, a position is sampled form a uniform distribution over the domain. If the sampled position has a closest neighbour less than the exclusion distance, $h$, units away, the position is rejected and resampled. Otherwise, the point is fixed at the sampled position. To avoid infinitely looping, 1,000 resampling tries are allowed, after which the best position (i.e., with the largest distance to the closest neighbour) is accepted. This process is repeated until all the points have been placed. **B.** An example of a reference point pattern (Top) and a corresponding CSR reconstruction (Bottom). Both point patterns have the same number of points of each type, but the CSR reconstruction does not capture the large area without type 2 (purple) points observed at the bottom of the reference configuration (red box).

M2. Testing Inhomogeneity with the $\boldsymbol{F}$-function

The $F$-function $F$, also called the empty-space function, is defined as the cumulative distribution function of the distance from an arbitrary location within $\mathcal{D}$ to the nearest point of $X$^1,2^. More precisely, if $d$ is the distance from an arbitrary location within $\mathcal{D}$ to the nearest point of $X$, then

$$F\left( r \right)\mathbb{=P}\left[ d\leq r \right].$$

(S1)

To estimate $F\left( r \right)$ on $X$, we evaluate it on a set $\mathcal{R}$ of equally spaced distances, $r$, between $r_{min}$ and $r_{max}$. A natural choice for $r_{min}$ and $r_{max}$ could be $0$ and the maximum possible distance within $\mathcal{D}$, respectively. Otherwise, the choice of $r_{min}$ and $r_{max}$ may be guided by the context of the point pattern, such as the nature of the points from $X$. Then, we draw uniformly 10,000 test locations within $\mathcal{D}$ and measure their distance to their respective nearest point of $X$. $F\left( r \right)$ is defined as the fraction of test locations that have a nearest point from $X$ at a distance less or equal to $r$.

To compare $X$ with CSR using the $F$-function (Figure 1 Step 1, Main text), we start by estimating $F\left( r \right)$ on $X$ with 10,000 test locations chosen uniformly and store this as $F^{\left( Obs \right)}\left( r \right)$. Then, we generate 1,000 null configurations $C_{R}^{\left( s \right)}$, $s=1,\cdots,1000$, under CSR on a domain identical to $\mathcal{D}$ and with as many points as $X$. For $s=1,\cdots,1000$, we estimate $F\left( r \right)$ on $C_{R}^{\left( s \right)}$ with 10,000 test locations chosen uniformly and store this as $F^{\left( s \right)}\left( r \right)$. $\left\{ F^{\left( s \right)}\left( r \right) \right\}_{s=1,\cdots,1000}$ becomes our null distribution $F_{Null}\left( r \right)$ for $F\left( r \right)$ under CSR. To test the hypothesis that $X$ has significantly larger empty regions than what is observed under CSR, we perform a one-sided global test based on maximum absolute deviation from the null model, $F_{Null}\left( r \right)$, following Myllymäki et al.^2^:

1. $\mu\left( r \right)=mean\left( F^{\left( s \right)}\left( r \right) \right),$
2. $\sigma\left( r \right)=std\left( F^{\left( s \right)}\left( r \right) \right),$
3. $Z_{Obs}\left( r \right)=\frac{\mu\left( r \right)-F_{Obs}\left( r \right)}{\sigma\left( r \right)},$
4. $Z^{\left( s \right)}\left( r \right)=\frac{\mu\left( r \right)-F^{\left( s \right)}\left( r \right)}{\sigma\left( r \right)},$
5. $T_{Obs}=\max_{r\mathcal{\in R}} Z_{Obs}\left( r \right),$
6. $T^{\left( s \right)}=\max_{r\mathcal{\in R}} Z^{\left( s \right)}\left( r \right),$

| 7. $p=\frac{1+\sum_{s=1}^{1000} \mathbf{1}_{\left( T^{\left( s \right)}\geq T_{obs} \right)}}{1000+1}.$ | (S2) |
| --- | --- |

In point 7, the $\boldsymbol{1}_{\left\{ condition \right\}}$ denotes the indicator function that evaluates to 1 when the condition is true and to $0$, otherwise. To avoid blow up due to very small standard deviation values, and since $F\left( r \right)$ will converge to 1 beyond a certain distance due to point density, we only keep distances $r$ such that

1. $r\leq r^{\star}$, where $r^{\star}$ is the furthest distance at which all curves reach 1 (i.e., the smallest $r\in\left[ r_{min},r_{max} \right]$ such that $F^{\left( Obs \right)}\left( r \right)=1$ and $F^{\left( s \right)}=1\forall s=1,\cdots,1000$), and
2. $\sigma\left( r \right)>{10}^{-6}$.

The $p$-value of the test is given in Eq. (S2). If $p$ is smaller than the significance level $\alpha$, then we conclude that $X$ has empty regions significantly larger than what is observed under CSR.

M3. Identifying and Quantifying Zero-Density Regions

To distinguish deviations from the empty-space function attributed to environmental constraints from those resulting from clustering patterns, we also quantify zero-density regions (Figure 1 Step 2, Main text). First, we divide $\mathcal{D}$ into a $n_{B}\times n_{B}$ grid with bins of side length $bL=\frac{2\cdot R}{n_{B}}$. For each bin, we record the number of points from $X$ within its boundaries and calculate its local point density by dividing this number by the area of the bin that is also within the limits of $\mathcal{D}$. Bins with zero-point density are classified as *zero-density* and others as *non-zero-density*. We we remove empty regions that have an area smaller than threshold $A_{0}$ using the function bwareaopen in MATLAB_R2026a^3^, with connectivity defined as Moore’s neighbourhood. Finally, zero-density regions of appropriate size within $\mathcal{D}$ are found using the function bwconncomp in MATLAB_R2026a^3^ (see the MATLAB function densityGridAnalysis provided with the SHAPPE code). The value for $A_{0}$ can be inferred from the nature of the points. In particular, we suggest choosing $A_{0}$ such that the radius of a disk of equal area is greater or equal to the maximal influence distance between two points. Hence, in a point pattern with an average global point density, an empty region of an area greater or equal to $A_{0}$ is unlikely to be solely attributed to clustering. $n_{B}$, or $bL$, can be chosen from visual inspection of zero-density region identification results depending on different tested values.

M4. Generating Environmentally Constrained Spatial Randomness

Similarly to CSR, environmentally constrained spatial randomness (ECSR) refers to the Gibbs hard-core point process with exclusion distance $h$, but where 1) points are not allowed within some predefined regions (*repulsive* regions), or where 2) points can only be placed within some predefined regions (*attractive* regions) (Figure S2A). Both ECSR scenarios are based on the zero-density region identification and quantification described in Section M3 and implemented in the function densityGridAnalysis (see GitHub code repository). ECSR serves as the random baseline for inhomogeneous point patterns (Figure S2B). The choice of which scenario to choose to generate a ECSR reconstruction $X^{\left( R \right)}$ on a domain $\mathcal{D}^{\left( R \right)}$ is guided by visual inspection of $X^{\left( Obs \right)}$. If $X^{\left( Obs \right)}$ has points scattered on most of the domain with a few distinct empty regions, then ECSR with repulsive regions might be preferred (M4.1). On the other hand, if $X^{\left( Obs \right)}$ contains a few distinct regions containing points surrounded by empty space, then ECSR with attractive regions would be a better choice (M4.2). In what follows, $A\left( \mathcal{D}^{\left( Obs \right)} \right)$ and $A\left( \mathcal{D}^{\left( R \right)} \right)$ refer to the area of $\mathcal{D}^{\left( Obs \right)}$ and $\mathcal{D}^{\left( R \right)}$, respectively (Table S2). Also, $N_{g}$, for $g=1,\cdots,G$, is the number of points of type $g$ in $X$, and $N_{g}^{\left( R \right)}=\left[ N_{g}^{\left( Obs \right)}\cdot\frac{A\left( \mathcal{D}^{\left( R \right)} \right)}{A\left( \mathcal{D}^{\left( Obs \right)} \right)} \right]$, where $f\left( x \right)=\left[ x \right]$ is the nearest-integer function.

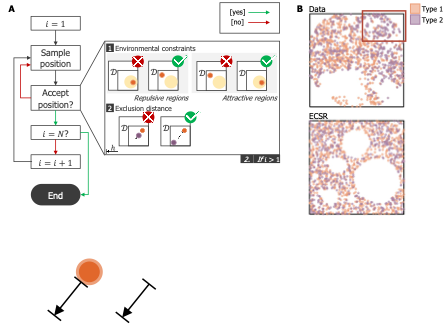

Figure S2. Generating Environmentally Constrained Spatial Randomness (ECSR) for inhomogeneous point processes.

**A.** Points are placed sequentially. When placing a point, a position is sampled from a uniform distribution over the domain. For the repulsive region scenario, the position is rejected if it falls within one of the predefined regions of exclusion (i.e., repulsive), illustrated in yellow. For the attractive region scenario, it is rejected if it *does not* fall within one the predefined inclusion (i.e., attractive) regions (illustrated in yellow). Also, if the sampled position has a closest neighbour less than $h$ (i.e., the exclusion distance) distance units away, the position is rejected and resampled. Otherwise, the point is fixed at the sampled position. To avoid infinitely looping, 1,000 resampling tries are allowed, after which the best position (i.e., with the largest distance to the closest neighbour, and outside of repulsive regions or inside of an attractive regions) is accepted. This process is repeated until all the points are placed. **B.** An example of an inhomogeneous point pattern (Top) and a corresponding ECSR reconstruction with repulsive regions (Bottom). Both point patterns have the same number of points of each type. ECSR lacks regions with higher densities of type 2 points, as can be seen in the upper right corner of the reference point pattern (red box).

M4.1. ECSR with Repulsive Regions

The function densityGridAnalysis returns the set $Z_{Low}$ of zero-density region areas in $X^{\left( Obs \right)}$, with $Z_{Low}^{\left( Obs \right)}=\left\{ Z_{Low}^{\left( 1 \right)},\cdots,Z_{Low}^{\left( n_{Low}^{\left( Obs \right)} \right)} \right\}$ and $n_{Low}^{\left( Obs \right)}=\left| Z_{Low}^{\left( Obs \right)} \right|$. To respect the zero-density region pattern in $X^{\left( Obs \right)}$, $X^{\left( R \right)}$ will contain $n_{Low}^{\left( R \right)}=\left[ \frac{A\left( \mathcal{D}^{\left( R \right)} \right)}{A\left( \mathcal{D}^{\left( Obs \right)} \right)}\cdot n_{Low}^{\left( Obs \right)} \right]$ circular zero-density regions $Z_{i}$ $\left( i=1,\cdots,n_{Low}^{\left( R \right)} \right)$ called repulsive regions, each defined by a center $\left( v_{x}^{\left( i \right)},v_{y}^{\left( i \right)} \right)\in\mathcal{D}^{\left( R \right)}$ and a radius $r^{\left( i \right)}$. $Z_{low}^{\left( R \right)}=\left\{ \left( v_{x}^{\left( i \right)},v_{y}^{\left( i \right)},r^{\left( i \right)} \right) \right\}_{i=1,\cdots,n_{Low}^{\left( R \right)}}$ is the set of repulsive regions in $X^{\left( R \right)}$ that act as exclusion zones. To generate ECSR, we start by sampling the exclusion zones (**Step 1**), then we place points (**Step 2**). The pseudocode to generate a point pattern $X^{\left( R \right)}$ under ECSR with repulsing regions $Z$, is:

**Step 0**: Set $i=1$.

**Step 1**: Set $n_{Try}=1$.

**Step 1.1**: Define $k$:

If $\boldsymbol{n}_{\boldsymbol{Low}}^{\left( \boldsymbol{R} \right)}=\boldsymbol{n}_{\boldsymbol{Low}}^{\left( \boldsymbol{Obs} \right)}$: $k=i$.

If $\boldsymbol{n}_{\boldsymbol{Low}}^{\left( \boldsymbol{R} \right)}>\boldsymbol{n}_{\boldsymbol{Low}}^{\left( \boldsymbol{Obs} \right)}$ and $\boldsymbol{i}\leq\boldsymbol{n}_{\boldsymbol{Low}}^{\left( \boldsymbol{Obs} \right)}$: $k=i$.

**Else**: Sample $k$ from a discrete uniform distribution between $1$ and $n_{Low}^{\left( Obs \right)}$.

**Step 1.2**: Define radius $r^{\left( i \right)}$ for void $V_{i}$ with $r^{\left( i \right)}=\sqrt{\frac{Z_{Low}^{\left( k \right)}}{\pi}}$.

**Step 1.3:** Sample a center $\left( v_{x}^{\left( i \right)},v_{y}^{\left( i \right)} \right)$ uniformly on $\mathcal{D}^{\left( R \right)}$ and within a distance $r^{\left( i \right)}$ from the boundaries.

**Step 1.4**: If there exists $j\in\left\{ 1,\cdots,i-1 \right\}$ such that the circle of radius $r^{\left( i \right)}$ centered at $\left( v_{x}^{\left( i \right)},v_{y}^{\left( i \right)} \right)$ intersects with $Z_{j}$, set $n_{Try}=n_{Try}+1$. If $n_{Try}=100$, go back to **Step 0**. Otherwise, go back to step **1.3**. If there exists no such $j$, then $Z_{i}=\left( v_{x}^{\left( i \right)},v_{y}^{\left( i \right)},r^{\left( i \right)} \right)$ and $i=i+1$. If $i> n_{Low}^{\left( R \right)}$, all the voids have been defined and go to **Step 2**. Otherwise, go to **Step 1** to define the next void.

**Step 2**: Set $i=1$ and $g=1$.

**Step 2.1**: If $i>N_{g}^{\left( R \right)}$, set $g=g+1$.

**Step 2.2**: Create a point $x_{i}$ of type $g$. Set $\mathrm{bes}t_{\mathrm{pos}}=\left( 0,0 \right)$ and $\mathrm{bes}t_{dist-to-neighbour}=-1$.

**Step 2.3**: Set $n_{Try}=1$.

**Step 2.4**: Sample a position $\left( x,y \right)$ uniformly on $\mathcal{D}^{\left( R \right)}$.

**Step 2.5**: [*Check repulsive regions*]

If $\left( x,y \right)$ falls within any of the repulsive regions in $Z_{Low}^{\left( R \right)}$, set $n_{Try}=n_{Try}+1$ and go to **Step** **2.8**. Otherwise, go to **Step 2.6**.

**Step 2.6**: [*Check distance to other points*]

If $\left( x,y \right)$ is within a distance less than $h$ from a point $x_{j}$ ($j=1,\cdots,i-1$), set $n_{Try}=n_{Try}+1$ and go to **Step 2.7**. Otherwise, set $\mathrm{bes}t_{\mathrm{pos}}=\left( x,y \right)$ go to **Step 2.9**.

**Step 2.7**: [*Store best position*]

If the minimal distance between $\left( x,y \right)$ and a point $x_{j}$ ($j=1,\cdots,i-1$) is greater than $\mathrm{bes}t_{dist-to-neighbour}$, set $\mathrm{bes}t_{dist-to-neighbour}$ to that minimal distance and $\mathrm{bes}t_{\mathrm{pos}}=\left( x,y \right)$.

**Step 2.8**: If $n_{Try}\leq1000$, go back to **Step 2.4**. Otherwise, go to **Step 2.9**.

**Step 2.9**: Place point $x_{i}$ at position $\mathrm{bes}t_{\mathrm{pos}}$ and go to **Step 2.10**.

**Step 2.10**: Set $i=i+1$. If $i\leq N^{\left( R \right)}$, go to **Step 2.1**. Otherwise, all points have been placed.

M4.2. ECSR with Attractive Regions

ECSR with attractive regions is generated similarly to ECSR with repulsive regions, but by defining inclusion zones, i.e., regions where points can be placed, instead of exclusion zones. The function densityGridAnalysis also returns the set $Z_{High}^{\left( Obs \right)}$ of non-zero-density region areas in $X^{\left( Obs \right)}$, with $Z_{High}^{\left( Obs \right)}=\left\{ Z_{High}^{\left( 1 \right)},\cdots,Z_{High}^{\left( n_{High}^{\left( Obs \right)} \right)} \right\}$ and $n_{High}^{\left( Obs \right)}=\left| Z_{High}^{\left( Obs \right)} \right|$. To respect the non-zero-density region pattern in $X^{\left( Obs \right)}$, $X^{\left( R \right)}$ will contain $n_{High}^{\left( R \right)}=\left[ \frac{A\left( \mathcal{D}^{\left( R \right)} \right)}{A\left( \mathcal{D}^{\left( Obs \right)} \right)}\cdot n_{High}^{\left( Obs \right)} \right]$ circular non-zero-density regions $Z_{i}$ $\left( i=1,\cdots,n_{High}^{\left( R \right)} \right)$ called *attractive regions*, each defined by a center $\left( v_{x}^{\left( i \right)},v_{y}^{\left( i \right)} \right)\in\mathcal{D}^{\left( R \right)}$ and a radius $r^{\left( i \right)}$. $Z_{High}^{\left( R \right)}=\left\{ \left( v_{x}^{\left( i \right)},v_{y}^{\left( i \right)},r^{\left( i \right)} \right) \right\}_{i=1,\cdots,n_{High}^{\left( R \right)}}$ is the set of attractive regions in $X^{\left( R \right)}$ that will act as inclusion zones. As in **M4.1**, we start by sampling the inclusion zones (**Step 1**), then we place points (**Step 2**). The pseudocode to generate a point pattern $X$ under ECSR with attractive regions $Z$, is:

**Step 0**: Set $i=1$.

**Step 1**: Set $n_{Try}=1$.

**Step 1.1**: Define $k$:

If $\boldsymbol{n}_{\boldsymbol{High}}^{\left( \boldsymbol{R} \right)}=\boldsymbol{n}_{\boldsymbol{High}}^{\left( \boldsymbol{Obs} \right)}$: $k=i$.

If $\boldsymbol{n}_{\boldsymbol{High}}^{\left( \boldsymbol{R} \right)}>\boldsymbol{n}_{\boldsymbol{High}}^{\left( \boldsymbol{Obs} \right)}$ and $\boldsymbol{i}\leq\boldsymbol{n}_{\boldsymbol{High}}^{\left( \boldsymbol{Obs} \right)}$: $k=i$.

**Else**: Sample $k$ from a discrete uniform distribution between $1$ and $n_{High}^{\left( Obs \right)}$.

**Step 1.2**: Define radius $r^{\left( i \right)}$ for attracting region $V_{i}$ with $r^{\left( i \right)}=\sqrt{\frac{Z_{High}^{\left( k \right)}}{\pi}}$.

**Step 1.3**: Sample a center $\left( v_{x}^{\left( i \right)},v_{y}^{\left( i \right)} \right)$ uniformly on $\mathcal{D}$ and within a distance $r^{\left( i \right)}$ from the boundaries.

**Step 1.4**: If there exists $j\in\left\{ 1,\cdots,i-1 \right\}$ such that the circle of radius $r^{\left( i \right)}$ centered at $\left( v_{x}^{\left( i \right)},v_{y}^{\left( i \right)} \right)$ intersects with $Z_{j}$, set $n_{Try}=n_{Try}+1$. If $n_{Try}=100$, go back to **Step 0**. Otherwise, go back to step **1.3**. If there exists no such $j$, then $Z_{i}=\left( v_{x}^{\left( i \right)},v_{y}^{\left( i \right)},r^{\left( i \right)} \right)$ and $i=i+1$. If $i>n_{High}^{\left( R \right)}$, all the attracting regions have been defined and go to **Step 2**. Otherwise, go to **Step 1** to define the next void.

**Step 2**: Set $i=1$ and $g=1$.

**Step 2.1**: If $i>N_{g}^{\left( R \right)}$, set $g=g+1$.

**Step 2.2**: Create a point $x_{i}$ of type $g$. Set $\mathrm{bes}t_{\mathrm{pos}}=\left( 0,0 \right)$ and $\mathrm{bes}t_{dist-to-neighbour}=-1$.

**Step 2.3**: Set $n_{Try}=1$.

**Step 2.4**: Sample a position $\left( x,y \right)$ uniformly on $\mathcal{D}$.

**Step 2.5**: [*Check attractive regions*]

If $\left( x,y \right)$ falls outside of all the attracting regions $Z_{High}^{\left( R \right)}$, set $n_{Try}=n_{Try}+1$ and go to **Step** **2.8**. Otherwise, go to **Step 2.6**.

**Step 2.6**: [*Check distance to other points*]

If $\left( x,y \right)$ is within a distance less than $h$ from a point $x_{j}$ ($j=1,\cdots,i-1$), set $n_{Try}=n_{Try}+1$ and go to **Step 2.7**. Otherwise, set $\mathrm{bes}t_{\mathrm{pos}}=\left( x,y \right)$ go to **Step 2.9**.

**Step 2.7**: [*Store best position*]

If the minimal distance between $\left( x,y \right)$ and a point $x_{j}$ ($j=1,\cdots,i-1$) is greater than $\mathrm{bes}t_{dist-to-neighbour}$, set $\mathrm{bes}t_{dist-to-neighbour}$ to that minimal distance and $\mathrm{bes}t_{\mathrm{pos}}=\left( x,y \right)$.

**Step 2.8**: If $n_{Try}\leq1000$, go back to **Step 2.4**. Otherwise, go to **Step 2.9**.

**Step 2.9**: Place point $x_{i}$ at position $\mathrm{bes}t_{\mathrm{pos}}$ and go to **Step 2.10**.

**Step 2.10**: Set $i=i+1$. If $i\leq N^{\left( R \right)}$, go to **Step 2.1**. Otherwise, all points have been placed.

M5. Comparing Pairwise Occurrences of Agent Types to Randomness with Ripley’s $\boldsymbol{K}$-Function

Ripley’s standard univariate, $K_{g}$, and multivariate, $K_{gk}$, $K$-functions are standard ways of measuring correlation between points of types $g$ and $k$ in a given point pattern (autocorrelation if $g=k$)^4^. If $g=k$, $K_{g}\left( r \right)$ is defined by the expected number of points of type $g$ within a distance $r$ of a typical same-type point divided by the leave-one-out intensity of type $g$ (i.e., $\frac{N_{g}-1}{A\left( \mathcal{D} \right)}$)^5–7^. In a point pattern $X$, it is estimated by

$$\hat{K}_{g}(r)=\frac{A\mathcal{(D)}}{N_{g}\cdot\left( N_{g}-1 \right)}\sum_{x_{i}\in X_{g}} \sum_{\begin{aligned} x_{j}\in X_{g} \\ x_{j}\neq x_{i} \end{aligned}} \frac{1}{w_{ij}}\cdot\mathbf{1}_{\left( d_{ij}\leq r \right)}.$$

(S3)

In Eq. (S3), $d_{ij}$ is the distance between two distinct points $x_{i}$ and $x_{j}$ from $X_{g}$. $\frac{1}{w_{ij}}$ is an edge correction for points close to the boundary of $\mathcal{D}$. Here, we use isotropic edge correction, with $w_{ij}$ equal to $1$ when the circle centered at point $x_{i}$ and with radius $d_{ij}$ is completely inside the domain. Otherwise, $w_{ij}$ equals the proportion of the circumference of that circle that falls within $\mathcal{D}$, which can be estimated via Monte Carlo simulations.

Similarly, $K_{gk}\left( r \right)$ ($g\neq k$) is defined as the expected number of points of type $k$ within a distance $r$ of a typical point of type $g$, divided by the intensity of type $k$ ^8^:

$$\hat{K}_{gk}\left( r \right)=\frac{1}{\hat{\lambda}_{g}\hat{\lambda}_{k}A\left( \mathcal{D} \right)}\sum_{x_{i}\in X_{g}} \sum_{x_{j}\in X_{k}} \frac{1}{w_{ij}}\cdot\mathbf{1}_{\left( d_{ij}\leq r \right)},$$

(S4)

with $\hat{\lambda}_{g}=\frac{N_{g}}{A}$ and $\hat{\lambda}_{k}=\frac{N_{k}}{A}$.

In a multitype point pattern, $K$-functions, and more specifically their comparison to null envelopes, cannot uncover specific mechanisms (i.e., repulsion or attraction) behind clustering patterns. Instead, we use them to compare pairwise occurrences between all pairs of types $\left( g,k \right)\in\left[ 1,\cdots,G \right]^{2}$ to what is expected under randomness^4^. We accomplish this using envelope tests based on Monte Carlo simulations under random labelling and a two-sided global test based on maximum absolute deviation, following Myllymäki et al.^2^ (Figure S3). In each Monte Carlo simulation, point locations from $X$ are preserved, but types are permuted randomly. This approach tests the null hypothesis that type assignment is independent of spatial location after conditioning on the observed underlying point pattern^4^. This null is consistent with our assumption that spatial inhomogeneity is not type-specific (see Main Text, Step 0.1 of SHAPPE pseudocode). Under this assumption, standard rather than inhomogeneous $K$-functions are sufficient, even for inhomogeneous point patterns.

To perform this test, we generate a set $\mathcal{R}$ of equally spaced distances between $r_{min}$ and $r_{max}$. We note that this set $\mathcal{R}$ does not have to be the same set that is used when testing inhomogeneity with the $F$-function (Section M2). Instead, $\mathcal{R}$ can be restricted to values that are more relevant for clustering observations, such as values within the range of influence between two points.

Then, we

1. Evaluate function $\hat{K}_{gk}\left( r \right)$ for all $r\mathcal{\in R}$ on $X$ for every pair $\left( g,k \right)\in\left[ 1,\cdots,G \right]^{2}$ and store this as $K_{gk}^{\left( Obs \right)}$,
2. Generate 1000 null configurations $C_{R}^{(s)}$, $s=1,\cdots,1000$, by randomly assigning type to the observed point locations in $X$ while keeping constant the number of points of each type,
3. For $s=1,\cdots,1000$, evaluate function $\hat{K}_{gk}\left( r \right)$ for all $r\mathcal{\in R}$ on $C_{R}^{(s)}$ and save this as $K_{gk}^{\left( s \right)}$, for every pair $\left( g,k \right)\in\left[ 1,\cdots,G \right]^{2}$,
4. Using the set $\left\{ K_{gk}^{\left( s \right)} \right\}_{s=1,\cdots,1000}$ as the null set for pair $\left( g,k \right)\in\left[ 1,\cdots,G \right]^{2}$, we test for significant departure of $K_{gk}^{\left( Obs \right)}$ from what is expected under intensity-constrained randomness using a two-sided global test based on maximum absolute deviation:

$$\mu\left( r \right)=mean\left( \left\{ K_{gk}^{\left( s \right)} \right\}_{s=1,\cdots,1000} \right),$$

$$\sigma\left( r \right)=std\left( \left\{ K_{gk}^{\left( s \right)} \right\}_{s=1,\cdots,1000} \right),$$

$$Z_{obs}\left( r \right)=\frac{K_{gk}^{\left( Obs \right)}\left( r \right)-\mu\left( r \right)}{\sigma\left( r \right)},$$

$$Z^{\left( s \right)}\left( r \right)=\frac{K_{gk}^{\left( s \right)}\left( r \right)-\mu\left( r \right)}{\sigma\left( r \right)} (\forall s=1,\cdots,1000),$$

$$T_{obs}=\max_{r\mathcal{\in R}} \left| Z_{obs}\left( r \right) \right|,$$

$$T^{\left( s \right)}=\max_{r\mathcal{\in R}} \left| Z^{\left( s \right)}\left( r \right) \right| (\forall s=1,\cdots,1000),$$

$$p=\frac{1+\sum_{s=1}^{1000} \mathbf{1}_{\left( T^{\left( s \right)}\geq T_{obs} \right)}}{1000+1}.$$

When $g=k$, $\hat{K}_{gk}\left( r \right)$ reduces to $\hat{K}_{g}\left( r \right)$ described in the previous steps. Because we test spatial correlations for multiple type-pairs within the same point pattern $X$, we apply a Benjamini-Hochberg correction for the multiple comparison problem, with false discovery rate FDR$=0.05$. If the Benjamini-Hochberg-adjusted p-value is less than $0.05$ for $K_{gk}^{\left( Obs \right)}$, we conclude points of type $g$ encounter points of type $k$ more or less often than expected under intensity-constrained randomness.

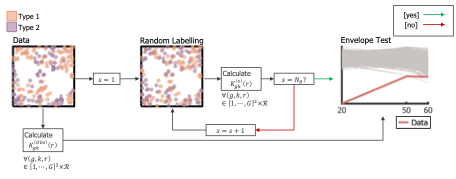

Figure S3. Assessing intertype correlations using random labelling null and an envelope test.

For every pair of type $\left( g,k \right)\in\left[ 1,\cdots, G \right]^{2}$, $K_{gk}^{\left( Obs \right)}\left( r \right)$ (for $r\mathcal{\in R}$) is estimated on the reference point pattern $X^{\left( Obs \right)}$. A null envelope for $K_{gk}$ is built by randomly assigning types to point locations in $X^{\left( Obs \right)}$ and estimating $K_{gk}^{\left( Obs \right)}$ on the new configuration. This process is repeated $N_{R}$ times. Significant deviation from the envelope is assessed using a two-sided global test based on maximum absolute deviation.

M6. Inhomogeneous Ripley’s $\boldsymbol{K}$-function

Ripley’s $K$-function between types $g$ and $k$ for an inhomogeneous point pattern^9,10^ $X$ is given by

$$\hat{K}_{gk}\left( r \right)=\frac{1}{A\left( \mathcal{D} \right)}\cdot\sum_{x_{i}\in X_{g}} \sum_{\begin{aligned} x_{j}\in X_{k} \\ x_{i}\neq x_{j} \end{aligned}} \frac{1}{\hat{\lambda}_{g}\left( x_{i} \right)}\cdot\frac{1}{\hat{\lambda}_{k}\left( x_{j} \right)}\cdot\frac{1}{w_{ij}}\cdot\mathbf{1}_{\left( d_{ij}\leq r \right)}.$$

(S5)

Eq. (S5) extends (S4) by replacing global intensities $\hat{\lambda}_{g}$ and $\hat{\lambda}_{k}$ with the intensity of types $g$ and $k$ at positions $x_{i}$ and $x_{j}$, respectively $\hat{\lambda}_{g}\left( x_{i} \right)$ and $\hat{\lambda}_{k}\left( x_{j} \right)$. In an inhomogeneous context, we estimate those with a circular moving window of radius $h_{Moving}$ with a leave-one-out rule and a uniform-kernel intensity estimator^11^. More specifically, $\hat{\lambda}_{g}\left( x_{i} \right)$ is the number of points of type $g$ that fall within a distance of $h_{Moving}$ from $x_{i}$, divided by the overlapping area between $\mathcal{D}$ and a disk of radius $h_{Moving}$ centered at $x_{i}$. $w_{ij}$ and $\boldsymbol{1}_{\left( d_{ij}\leq r \right)}$ are given as before.

M7. Detailed Sequential Hooke’s Law Point Pattern Reconstruction with Environmental Constraints
    M7.1. Ghost points

To use periodic or symmetric boundary conditions if points move out of the domain $\mathcal{D}$, we include ghost points in our search of neighbours^12^. To avoids ghost points outnumbering real points when updating positions, we only consider ghosts of points within a distance $d_{B}$ of the boundary of $\mathcal{D}$. If $\mathcal{D}$ is a square (Figure S4A), then each point at position $\left( x,y \right)$ has at most three ghost points of the same mark as itself at positions $\left( x_{1},y_{1} \right)$, $\left( x_{2},y_{2} \right)$ and $\left( x_{3},y_{3} \right)$, respectively:

$$\left( x_{1},y_{1} \right)=\left( x-sgn\left( x \right)\cdot2\cdot R,y \right),$$

$$\left( x_{2},y_{2} \right)=\left( x,y-sgn\left( y \right)\cdot2\cdot R \right),$$

$$\left( x_{3},y_{3} \right)=\left( x-sgn\left( x \right)\cdot2\cdot R,y-sgn\left( y \right)\cdot2\cdot R \right).$$

Here, $sgn\left( x \right)$ is the sign function that is equal to $-1$ if $x<0$ and to $1$ otherwise. If $\mathcal{D}$ is a disk (Figure S4B), then each point at position at $\left( x,y \right)$ has at most one ghost point of the same mark as itself at position $\left( x_{1},y_{1} \right)$:

$$\left( x_{1},y_{1} \right)=\left( x,y \right)-2\cdot R\cdot\frac{\left( x,y \right)}{\sqrt{x^{2}+y^{2}}}=\left( x-\frac{2\cdot R\cdot x}{\sqrt{x^{2}+y^{2}}},y-\frac{2\cdot R\cdot y}{\sqrt{x^{2}+y^{2}}} \right).$$

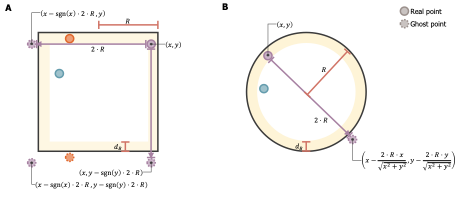

Figure S4. Definition of ghost points used in SHAPPE algorithm.

**A.** If $\mathcal{D}$ is a square, we implement periodic boundary conditions. Points can have at most three ghost points, but only ghost of points within $d_{B}$ distance units from a side of $\mathcal{D}$ (yellow region) are considered. The purple point at position $\left( x,y \right)$ is within $d_{B}$ distance units from right and upper walls and, as such, has three ghost points. The orange point is further away than $d_{B}$ distance units from both the right and left walls, but within $d_{B}$ distance units of the upper wall, which means it has only one ghost point. The blue point is not in the yellow region and has no ghost point. **B.** If $\mathcal{D}$ is a disk of radius $R$, we implement symmetric boundary conditions. If a point is within $d_{B}$ units from the distance boundary (yellow region), it has a ghost point along the same radial line, but on the opposite side of the boundary. Here, the purple point has a ghost point, but the blue point does not because it is outside the yellow region.

M7.2. Sampling target distances

We assume types are placed sequentially in the order $1,\cdots,G$. When placing points of type $g$, each point $x_{i}\in X_{g}$ is attributed target distances to neighbours of types $1,\cdots,g$. More specifically, $s_{i}^{\left( k,l \right)}$ refers to the target distance assigned to $x_{i}$’s $l$-th neighbour of type $k$, for $k\in\left\{ 1,\cdots,g \right\}$ and $l\in\left\{ 1,\cdots,L \right\}$. For $l=1$, $s_{i}^{\left( k,l \right)}$ is sampled from the nearest-neighbour distribution from type $g$ to type $k$. For $l>1$, $s_{i}^{\left( k,l \right)}$ is set recursively as $s_{i}^{(k,l)}=s_{i}^{(k,l-1)}+u$ where $u$ is sampled from the distribution of differences between the distances to the $l$-th and the $(l-1)$-th neighbours of type $k$ for points of type $g$. This distinction $l=1$ versus $l>1$ is important to sample target distances satisfying the ordering constraint $s_{i}^{\left( k,l \right)}>s_{i}^{\left( k,l-1 \right)}$.

If no significant clustering pattern is detected between types $g$ and $k$ (i.e., $\Omega_{gk}=0$), target distances are sampled independently from the corresponding distributions in the random baseline (i.e., $X^{\left( R \right)}$) using the inverse transform sampling method (Section M7.3**)**. If clustering is detected for one or more types (i.e., if there exists $k$ such that $\Omega_{gk}=1$), we use a copula (Gaussian or Student’s $t$) to jointly sample target distances from distributions derived from $X^{\left( Obs \right)}$and preserve their correlations as observed in the original point pattern. Following Demarta and McNeil^13^, we calculate the correlation matrix for both copulas using Kendall’s $\tau$ and map it to the correlation coefficient $\rho^{*}$ where

$$\rho^{*}=\sin\left( \frac{\pi}{2}\cdot\tau\right).$$

For the $t$-copulas, we additionally fit $\nu$, the degrees of freedom, by maximising the $t$-copula log-likelihood on rank-transformed observations. We then sample random numbers between 0 and 1 with linear correlation parameters $\rho^{*}$ (and degrees of freedom $\nu$ if using a Student’s $t$ copula) for all $s_{i}^{\left( k,l \right)}$ such that $\Omega_{gk}=1$ and $l=1,\cdots, L$ using the MATLAB function copularnd. Sampling is then performed with the inverse transform sampling method (Section M7.3), with $s_{i}^{(k,l)}=s_{i}^{(k,l-1)}+u$ for $l>1$, as before.

M7.3. Sampling from a distribution

To sample from a distribution, we use the cumulative distribution function (CDF) and the inverse transform sampling method (Figure S5). Assume we have a distribution $D$ of $n$ values, in ascending order, where values can appear more than once in $D$ (Figure S5A). Let $n_{Distinct}$ be the number of *distinct* values in $D$. Then, the CDF for D, $D_{Cumul}$ (Figure S5B), is given by

$$D_{Cumul}\left( x_{1} \right)=\frac{\sum_{j=1}^{n} \mathbf{1}_{\left( D\left( j \right)=x_{1} \right)}}{n},$$

$$D_{Cumul}\left( x_{i} \right)=D_{Cumul}\left( x_{i-1} \right)+\frac{\sum_{j=1}^{n} \mathbf{1}_{\left( D\left( j \right)=x_{i} \right)}}{n} (if i>1),$$

where $x_{i}$ is the $i$-th *distinct* value in $D$. $D\left( j \right)$ refers to the $j$-th element in $D$. $\sum_{j=1}^{n} \boldsymbol{1}_{\left( D\left( j \right)=x_{i} \right)}$ counts the number of times $x_{i}$ appears in $D$. To sample randomly a value from $D$, we

1. Sample a number $p$ uniformly on $\left[ 0,1 \right]$.
2. If $p\leq D_{Cumul}\left( x_{1} \right)$, set $i=1$. Otherwise, Find $i\in\left\{ 1,\cdots,n_{Distinct}-1 \right\}$ such that

$D_{Cumul}\left( x_{i} \right)<p\leq D_{Cumul}\left( x_{i+1} \right)$.

3. Sample $u$ uniformly between $x_{i}$ and $x_{i+1}$ and return $u$ to introduce variability in the new distributions.

Note that when target distances are sampled jointly with a copula (Section M7.2), the value for $p$ in Step 1 is taken from copularnd instead of sampling uniformly on $\left[ 0,1 \right]$.

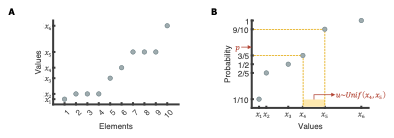

Figure S5. Sampling from a distribution with the inverse transform method.

**A**. Representation of a distribution $D$ containing $n=10$ non-unique elements. $D$ contains $n_{Distinct}=6$ distinct values. **B.** To sample from $D$, we first generate its cumulative distribution. Then, we sample a number $p$ uniformly on $[0,1]$ and find the two elements $x_{i}$ and $x_{i+1}$ such that $D_{Cumul}\left( x_{i} \right)<p\leq D_{Cumul}\left( x_{i+1} \right)$. Here, $x_{i}=x_{4}$. $u$ is then sample uniformly between $x_{i}$ and $x_{i+1}$ to introduce variability.

M7.4. Additional Environmental Constraints

**Step 0.3** of SHAPPE consists of generating a random baseline configuration $X^{\left( R \right)}$ for a synthetic reconstruction $X^{\left( Art \right)}$. For inhomogeneous point patterns, $X^{\left( R \right)}$ is obtained from ECSR, which can be defined through repulsive or attractive regions (Section M4). To better preserve distinct shapes observed in the reference point pattern $X^{\left( Obs \right)}$, such as points forming a blob, we modify Eq. (3) (Main text) in **Step 4.1** to add attraction towards or repulsion from specific regions, following the same scenario used to generate $X^{\left( R \right)}$.

M7.4.1. Attraction

Let $V=\left\{ V_{j}=\left( v_{x}^{\left( j \right)},v_{y}^{\left( j \right)},r^{\left( j \right)} \right) \right\}_{j=1,\cdots,n_{High}^{\left( R \right)}}$be the set of non-overlapping *attractive* regions in $X^{\left( R \right)}$. Suppose we are updating the position of the point $x_{i}\in X_{g}$ currently at position $\left( x,y \right)$. If $x_{i}$ is already inside one of the attractive regions, Eq. (3) in Step 4.1 remains unchanged. Otherwise, let $V_{j^{\star}}$ be the attractive region whose boundary is closest to $x_{i}$. $V_{j^{\star}}$ is the disk of radius $r^{\left( j^{\star} \right)}$ centered at $\left( v_{x}^{\left( j^{\star} \right)},v_{y}^{\left( j^{\star} \right)} \right)$. Eq. (3) in Step 4.1 becomes

$$d_{i}\left( n \right)=\sum_{k=1}^{g} \sum_{l=1}^{L} \lambda\cdot S_{i}^{\left( k,l \right)}\left( n \right)+r_{W}\cdot\frac{\left( v_{x}^{\left( j^{\star} \right)},v_{y}^{\left( j^{\star} \right)} \right)-\left( x,y \right)}{\left\| \left( v_{x}^{\left( j^{\star} \right)},v_{y}^{\left( j^{\star} \right)} \right)-\left( x,y \right) \right\|}\cdot\left( \frac{r^{\left( j^{\star} \right)}}{\left\| \left( v_{x}^{\left( j^{\star} \right)},v_{y}^{\left( j^{\star} \right)} \right)-\left( x,y \right) \right\|} \right).$$

$\left\| \left( v_{x}^{\left( j^{\star} \right)},v_{y}^{\left( j^{\star} \right)} \right)-\left( x,y \right) \right\|$ is the distance between $x_{i}$ and the center of $V_{j^{\star}}$. In that case, $x_{i}$ is attracted towards the center of $V_{j^{\star}}$ with strength $\frac{r^{\left( j^{\star} \right)}}{\left\| \left( v_{x}^{\left( j^{\star} \right)},v_{y}^{\left( j^{\star} \right)} \right)-\left( x,y \right) \right\|}$.

M7.4.2. Repulsion

Let $V=\left\{ V_{j}=\left( v_{x}^{\left( j \right)},v_{y}^{\left( j \right)},r^{\left( j \right)} \right) \right\}_{j=1,\cdots,n_{Low}}$ be the set of non-overlapping *repulsive* regions in $X^{\left( R \right)}$ and suppose we are updating the position of the point $x_{i}\in X_{g}$ currently at position $\left( x,y \right)$. If $x_{i}$ is outside of the repulsive regions, then Eq. (3) in Step 4.1 remains unchanged. Otherwise, let $V_{j^{\star}}$ be the repulsing region in which $x_{i}$ lies, such that $x_{i}\in V_{j^{\star}}$. $V_{j^{\star}}$ is the disk of radius $r^{\left( j^{\star} \right)}$ centered at $\left( v_{x}^{\left( j^{\star} \right)},v_{y}^{\left( j^{\star} \right)} \right)$. If $\left( x,y \right)\neq\left( v_{x}^{\left( j^{\star} \right)},v_{y}^{\left( j^{\star} \right)} \right)$, Eq. (3) in Step 4.1 becomes

$$d_{i}\left( n \right)=\sum_{k=1}^{g} \sum_{l=1}^{L} \lambda\cdot S_{i}^{\left( k,l \right)}\left( n \right)+r_{W}\cdot\frac{\left( x,y \right)-\left( v_{x}^{\left( j^{\star} \right)},v_{y}^{\left( j^{\star} \right)} \right)}{\left\| \left( x,y \right)-\left( v_{x}^{\left( j^{\star} \right)},v_{y}^{\left( j^{\star} \right)} \right) \right\|}\cdot\left( 1-\frac{\left\| \left( x,y \right)-\left( v_{x}^{\left( j^{\star} \right)},v_{y}^{\left( j^{\star} \right)} \right) \right\|}{r^{\left( j^{\star} \right)}} \right).$$

If $\left( x,y \right)=\left( v_{x}^{\left( j^{\star} \right)},v_{y}^{\left( j^{\star} \right)} \right)$, then $\frac{\left( x,y \right)-\left( v_{x}^{\left( j^{\star} \right)},v_{y}^{\left( j^{\star} \right)} \right)}{\left\| \left( x,y \right)-\left( v_{x}^{\left( j^{\star} \right)},v_{y}^{\left( j^{\star} \right)} \right) \right\|}$ is replaced by $\frac{\left( x^{'},y^{'} \right)-\left( v_{x}^{\left( j^{\star} \right)},v_{y}^{\left( j^{\star} \right)} \right)}{\left\| \left( x^{'},y^{'} \right)-\left( v_{x}^{\left( j^{\star} \right)},v_{y}^{\left( j^{\star} \right)} \right) \right\|}$, where $\left( x^{'},y^{'} \right)$ is a random point on the boundary of $V_{j^{\star}}$. In both cases, $x_{i}$ is repulsed from the center of $V_{\left( j^{\star} \right)}$ with strength $\left( 1-\frac{\left\| \left( x,y \right)-\left( v_{x}^{\left( j^{\star} \right)},v_{y}^{\left( j^{\star} \right)} \right) \right\|}{r^{\left( j^{\star} \right)}} \right)$.

M7.5. Boundary conditions for square and circular domains

It is necessary to define boundary conditions in case the position $p_{i}\left( n+1 \right)$ for point $x_{i}$ (Eq. (4), Main Text) fell outside of the domain $\mathcal{D}^{\left( Art \right)}$. We propose periodic boundary conditions if $\mathcal{D}^{\left( Art \right)}$ is a square domain (Figure S6A) and an approximation of symmetric boundary conditions^12^ if $\mathcal{D}^{\left( Art \right)}$ is a disk (Figure S6B). These boundaries are coherent with the definition of ghost points for updating cell positions. Similar boundary conditions can be established if $\mathcal{D}^{\left( Art \right)}$ has a more complex shape.

If $\mathcal{D}^{\left( Art \right)}=\left[ -R^{\left( Art \right)},R^{\left( Art \right)} \right]^{2}\subset\mathbb{R}^{2}$ and $p_{i}\left( n+1 \right)=\left( x,y \right)\notin\mathcal{D}^{\left( Art \right)}$, then

$$p_{i}\left( n+1 \right)=\left( \mathbf{1}_{\left( \left| x \right|>R^{\left( Art \right)} \right)}\cdot x-sign\left( x \right)\cdot2\cdot R^{\left( Art \right)},\mathbf{1}_{\left( \left| y \right|>R^{\left( Art \right)} \right)}\cdot y-sign\left( y \right)\cdot2\cdot R^{\left( Art \right)} \right).$$

If $\mathcal{D}^{\left( Art \right)}$ is a disk of radius $R^{\left( Art \right)}$ on $\mathbb{R}^{2}$, then $\mathcal{D}^{\left( Art \right)}=\left\{ \left( x,y \right) :x^{2}+y^{2}\leq R^{\left( Art \right)} \right\}$. If $p_{i}\left( n+1 \right)=\left( x,y \right)\notin\mathcal{D}^{\left( Art \right)}$, then

$$p_{i}\left( n+1 \right)=\left( x-2\cdot R^{\left( Art \right)}\cdot\frac{x}{\sqrt{x^{2}+y^{2}}},y-2\cdot R^{\left( Art \right)}\cdot\frac{y}{\sqrt{x^{2}+y^{2}}} \right).$$

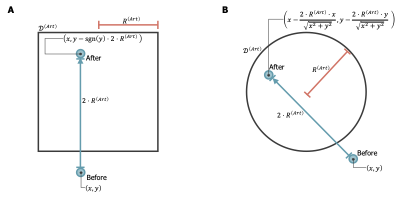

Figure S6. Boundary conditions for different domain types.

**A.** Square and **B.** circular domains.

M7.6. Density grid as a proxy for environmental constraints

For inhomogeneous point pattern reconstruction algorithms using energy minimisation, Końasová and Dvorák^14^ suggest including the difference between the estimated intensity functions (or their empirical distributions) computed from the reference data and a synthetic configuration. Here, we extend this idea to SHAPPE by imposing environmental constraints every $n_{Env}$ iteration through a comparison between density grids derived from $X^{\left( Obs \right)}$ and $X^{\left( Art \right)}$. For this, we:

**Step 1**: Build a density grid for the current point placement, $E^{\left( n,t \right)}$.

Suppose we are placing points of type $g$. Then, we consider all points of type $1,\cdots,g$. $E^{\left( n,t \right)}$ has bins of side length $bL_{Env}$, chosen such that bins have an area similar to the minimal area of zero-density region attributed to environmental factors (i.e., $A_{0}$, see Section M3).

**Step 2**: Build the corresponding data-derived grid, $E^{\left( Obs,t \right)}$.

Using $X^{\left( Obs \right)}$, we build grid $E^{\left( Obs,t \right)}$ with bins of length $bL_{Env}$ as a proxy for the environmental constraints. We again consider only points of type $1,\cdots,g$. See Section M7.7 if $X^{\left( Art \right)}$ and $X^{\left( Obs \right)}$ do not have the same domain area.

**Step 3**: Assign target point counts to bins in $E^{\left( n,t \right)}$.

We rank and pair bins in $E^{\left( n,t \right)}$ and $E^{\left( Obs,t \right)}$ according to their point count density and denote by $b_{i}^{\left( n \right)}$ and $b_{i}^{\left( X \right)}$ the bin in $E^{\left( n,t \right)}$ and $E^{\left( Obs,t \right)}$ with the $i$-th lowest point density, respectively. We then convert observed densities, $b_{i}^{\left( X \right)}$, into target counts by multiplying them by the area of their paired synthetic bin in $E^{\left( n,t \right)}$ that overlaps with $\mathcal{D}^{\left( Art \right)}$. This does not necessarily preserve the total number of points in $E^{\left( n,t \right)}$, so the resulting target counts are renormalized to ensure that their sum matches the current total number of points considered in $E^{\left( n,t \right)}$.

**Step 4**: Redistribute points from overloaded bins to underloaded neighbouring bins.

Let $c_{i}^{\left( n \right)}$ be the target count for bin $b_{i}^{\left( n \right)}$ and $N_{i}^{\left( n \right)}$ be the current number of points it contains. Its excess is given by:

$$e_{i}=\left\{ \begin{matrix} 0 & \mathrm{if}N_{i}^{\left( n \right)}\leq c_{i}^{\left( n \right)} \\ \left\lfloor N_{i}^{\left( n \right)}-c_{i}^{\left( n \right)} \right\rfloor& \mathrm{otherwise} \end{matrix} \right..$$

For all $i$ such that $e_{i}>0$, we move $e_{i}$ points of type $g$ out of bin $b_{i}^{\left( n \right)}$ and place them randomly in an underfull neighbouring bin. If there are fewer than $e_{i}$ points of type $g$ in the bin, we move all the points of type $g$ out of the bin and no longer consider this bin during the current environmental constraint update. The neighbouring bin is chosen considering Moore’s neighbourhood one layer at a time, while enforcing the exclusion distance $h$. If no position satisfying the exclusion distance is found within inside the neighbouring bin after 100 tries, we accept the best position out of these attempts. The best position in that case is defined as the position with the largest distance to the closest neighbour. Excess in bins is recalculated every time a point is moved.

M7.7. Estimating Density Grids for Different Areas and Shapes

Let $N_{Env}^{\left( Art \right)}$ and $N_{Env}^{\left( Obs \right)}$ be the number of bins in $\left[ -R^{\left( Art \right)},R^{\left( Art \right)} \right]\times\left[ -R^{\left( Art \right)},R^{\left( Art \right)} \right]$ and $\left[ -R^{\left( Obs \right)},R^{\left( Obs \right)} \right]\times\left[ -R^{\left( Obs \right)},R^{\left( Obs \right)} \right]$ that intersect with $\mathcal{D}^{\left( Art \right)}$ and $\mathcal{D}^{\left( Obs \right)}$, respectively. If $\mathcal{D}^{\left( Art \right)}$ has a larger domain than $\mathcal{D}^{\left( Obs \right)}$ or is of a different shape, then we might have $N_{Env}^{\left( Art \right)}>N_{Env}^{\left( Obs \right)}$. In that case, let $E_{raw}^{\left( Obs,g \right)}$ be the density grid built directly from $X^{\left( Obs \right)}$. $E^{\left( Obs,g \right)}$ is sampled from $E_{raw}^{\left( Obs,g \right)}$ to ensure it contains as many bins as $E^{\left( n,g \right)}$: we include all the bins from $E_{raw}^{\left( Obs,g \right)}$ to $E^{\left( Obs,g \right)}$ and sample $N_{Env}^{\left( Art \right)}-N_{Env}^{\left( Obs \right)}$ additional bins from $E_{raw}^{\left( Obs,g \right)}$ with replacement.

These steps are only performed once in the entire algorithm; when $g$ changes, the same bins are sampled, but their respective density is updated to include the new type.

M8. Overlapping Index

The overlapping index between two distributions, $\Phi_{1}$ and $\Phi_{2}$, measures the similarity between them Let $f_{1}$ and $f_{2}$ be the probability density functions associated with $\Phi_{1}$ and $\Phi_{2}$, respectively. Then, the overlapping index is given by

$$ov\left( \Phi_{1},\Phi_{2} \right)=\int_{-\infty}^{\infty} \min\left( f_{1}\left( \theta\right),f_{2}\left( \theta\right) \right)d\theta.$$

The values of $ov\left( \Phi_{1},\Phi_{2} \right)$ vary between $0$ ($\Phi_{1}$ and $\Phi_{2}$ are disjoint) and $1$ ($\Phi_{1}$ and $\Phi_{2}$ are identical). Let $\theta_{min}$ and $\theta_{max}$ be the global minimum and maximum values over all values in $\Phi_{1}$ and $\Phi_{2}$, respectively. $f_{1}\left( \theta\right)$ and $f_{2}\left( \theta\right)$ are estimated and evaluated using the function ksdensity in MATLAB on a fine grid of 1024 equally spaced values of $\theta$ between $\theta_{min}$ and $\theta_{max}$. We approximate $ov\left( \Phi_{1},\Phi_{2} \right)$ with trapezoidal numerical integration using function trapz in MATLAB, using the same grid for $\theta$ as for $f_{1}$ and $f_{2}$.

M9. Tumour Microenvironment Reconstructions Using SHAPPE

To assess inhomogeneity in $X^{\left( IMC \right)}$, the empty-space function $F\left( r \right)$ (Eq. (S1)) is evaluated on a set $\mathcal{R}$ of $5\mu m$-spaced distances $r$ between $5\mu m$ and the maximum distance $r_{max}$ in $\mathcal{D}^{\left( IMC \right)}$, i.e., $r_{max}=\sqrt{2\cdot\left( 2\cdot R^{\left( IMC \right)} \right)^{2}}\mu m$. Bull et al.^15^ observed correlations in a mouse intestinal tumour image at distance $r=50\mu m$, which corresponds to a length scale of two to three cell diameters. We thus estimate the cell-cell interaction scale to be $50\mu m$, and we set the area threshold for a zero-density region, $A_{0}$, to the area of a disk of radius $50\mu m$. To identify and quantify empty regions, we chose $bL$ by performing a visual inspection of zero-density identification results for $bL\in\left\{ 25,40,80 \right\} \mu m$ which led to $bL=40\mu m$. To assess spatial correlations in $X^{\left( IMC \right)}$, Ripley’s $K$-functions were evaluated on the set of $5\mu m$-spaced distances between $15$ and $60\mu m.$ SHAPPE parameters used for our tumour microenvironment reconstructions are given in Table S3. Hyperparameters were optimised to minimise the absolute clustering errors at distances $r=20$, $40$ and $60\mu m$. (Section M10).

| Parameter | Value | Justification |
| --- | --- | --- |
| $N_{min}$ | $15$ | Practical choice, chosen to exceed the largest value of $L$ tried in the case study |
| $h$ | $0.8202\mu m$ | Minimal distance between two cells over all 87 IMC images |
| $d_{B}$ | $50\mu m$ | Interaction scale estimated from Bull et al.^15^ |
| $n_{Env}$ | $50$ | Practical choice |
| ${bL}_{Env}$ | $80\mu m$ | Set to twice the value of $bL$ used in the zero-density region identification step |
| $d_{Med}$ | $0.1075\mu m$ | 1% of the largest cell radius |
| $d_{90}$ | $10.75\mu m$ | Largest cell radius |
| $d_{Net}$ | $10.75\mu m$ | Largest cell radius |
| ${Max}_{n}$ | $10,000$ | Practical choice |
| $n_{Eq}$ | $210$ | 2% of ${Max}_{n}$ + 10 iterations to move cells around depending on their new neighbourhood |

Table S3. Parameter values in the large-scale IMC case study.

M10. Tumour Microenvironment Reconstructions: Hyperparameter Optimisation in SHAPPE

We set $\lambda=\frac{0.01}{0.133}\approx0.075$ following Jenner et al.^12^. $\phi_{min}$ was set to the largest cell radius of $10.75\mu m$. We varied the number of neighbours, $L$, the weight assigned to uncorrelated cell types, $w_{NoCorr}$, the copula family, $C$, the neighbour influence weight, $\alpha$, the repulsion weight, $r_{W}$ and the baseline step limit, $\phi$. over the values indicated in Table S4.

| Hyperparameter | Description | Values |
| --- | --- | --- |
| $L$ | Number of closest neighbours considered when updating the position of a point. | $\left\{ 1,5,10 \right\}$ |
| $w_{NoCorr}$ | Weight attributed to uncorrelated types, relative to weight attributed to correlated types | $\left\{ 0,0.5,1 \right\}$ |
| $C$ | Copula family to sample $s_{i}^{\left( k,l \right)}$ for all $k\in\left\{ 1,\cdots,g \right\}$ such that $A_{gk}=1$ and for all $l\in\left\{ 1,\cdots,L \right\}$, for a given type $g$. | {Gaussian, Student’s $t$} |
| $\alpha$ | Neighbour influence weight | $\left\{ 0,0.5,1 \right\}$ |
| $r_{W}$ | Repulsion weight to enforce an exclusion distance | $\left\{ 1,5,10 \right\}$ |
| $\phi$ | Baseline step limit | $\left\{ 2,5,10 \right\}\cdot$largest cell radius |

Table S4. SHAPPE Hyperparameter values.

Values of hyperparameters from the optimisation of SHAPPE hyperparameters for 87 reference point patterns from as many IMC images.

Let $\Theta$ be the set of all hyperparameter combinations given by Table S4 and $\Omega$ be the set of IMC point patterns. Since $\alpha$ is irrelevant when $L=1$ (Eqs. (2) and (3) in the Main Text), we remove combinations where $L=1$ and $\alpha>0$ from $\Theta$. We perform Pareto optimisation^16^ of the absolute clustering error at distances $20$, $40$ and $60\mu m$ on $\Omega$ to select the optimised hyperparameter set $\theta^{\star}$. For inhomogeneous point patterns, we set the moving window bandwidth, $h_{Moving}$ (Section M6), to $100\mu m$. This choice is consistent with the suggestion of Illian et al.^11^ who recommend having the intensity bandwidth larger than the scale at which we calculate second-order statistics.

Specifically, for every $\theta\in\Theta$ and every $\omega\in\Omega$, we first compute the absolute clustering errors

$$e_{H}^{\left( \theta,\omega\right)}\left( 20 \right), e_{H}^{\left( \theta,\omega\right)}\left( 40 \right), e_{H}^{\left( \theta,\omega\right)}\left( 60 \right).$$

Then, for each distance $r=20, 40,$ and $60\mu m$, we min-max normalise the scores using the theoretical lower bound $0$ and the maximum observed score over all hyperparameter combinations and all IMC point patterns:

$$v_{min}=0, v_{max}\left( r \right)=\max_{\theta\in\Theta,\omega\in\Omega} e_{H}^{\left( \theta,\omega\right)}\left( r \right),$$

$$\tilde{e}_{H}^{\left( \theta,\omega\right)}\left( r \right)=\frac{e_{H}^{\left( \theta,\omega\right)}\left( r \right)-v_{min}}{v_{max}\left( r \right)-v_{min}}=\frac{e_{H}^{\left( \theta,\omega\right)}\left( r \right)}{v_{max}\left( r \right)}.$$

The average normalised absolute clustering error at distance $r$ for hyperparameter combination $\theta$ is given by:

$$\bar{e}_{H}^{\left( \theta\right)}\left( r \right)=\frac{\sum_{\omega\in\Omega} \tilde{e}_{H}^{\left( \theta,\omega\right)}\left( r \right)}{\left| \Omega\right|}.$$

(S6)

Our vector-valued objective function is defined as

$$\begin{matrix} f: & \Theta& \longrightarrow& \mathbb{R}^{3} \\ & \theta& ⟼ & \left( \begin{matrix} f_{1}\left( \theta\right)=\bar{e}_{H}^{\left( \theta\right)}\left( 20 \right) \\ f_{2}\left( \theta\right)=\bar{e}_{H}^{\left( \theta\right)}\left( 40 \right) \\ f_{3}\left( \theta\right)=\bar{e}_{H}^{\left( \theta\right)}\left( 60 \right) \end{matrix} \right) \end{matrix}.$$

(S7)

We build the Pareto front $\mathcal{P\subseteq}\Theta$ using function find_pareto_frontier^17^ in MATLAB. Since all scores are theoretically bounded below by $0$, the ideal objective vector is defined as

$$z^{ideal}=\left( \begin{matrix} 0 \\ 0 \\ 0 \end{matrix} \right).$$

We choose the optimal hyperparameter set $\theta^{\star}\mathcal{\in P}$ as the one minimising the $L^{2}$-distance to $z^{ideal}$:

$$\theta^{\star}=\arg\min_{\theta\mathcal{\in P}} \left\| f\left( \theta\right)-z^{ideal} \right\|_{2}.$$

(S8)

The optimisation routine can also be done for every point pattern. The $\omega$-dependent vector-valued objective function is defined as

$$\begin{matrix} f: & \Theta\times\Omega& \longrightarrow& \mathbb{R}^{3} \\ & \left( \theta,\omega\right) & ⟼ & \left( \begin{matrix} f_{1}\left( \theta,\omega\right)=\tilde{e}_{H}^{\left( \theta,\omega\right)}\left( 20 \right) \\ f_{2}\left( \theta,\omega\right)=\tilde{e}_{H}^{\left( \theta,\omega\right)}\left( 40 \right) \\ f_{3}\left( \theta,\omega\right)=\tilde{e}_{H}^{\left( \theta,\omega\right)}\left( 60 \right) \end{matrix} \right) \end{matrix}.$$

The $\omega$-Pareto front $\mathcal{P}\left( \omega\right)\subseteq\Theta$ is built using the function find_pareto_frontier in MATLAB^17^. The hyperparameter combination optimised for point pattern $\omega$, i.e. $\theta^{\star}\left( \omega\right)\mathcal{\in P}\left( \omega\right)$, is chosen as the one minimising the $L^{2}$-distance to $z^{ideal}$:

$$\theta^{\star}\left( \omega\right)=\arg\min_{\theta\mathcal{\in P}\left( \omega\right)} \left\| f\left( \theta,\omega\right)-z^{ideal} \right\|_{2}.$$

M11. Assessment of Hyperparameter Importance in SHAPPE

We assess the importance of SHAPPE’s hyperparameters using the hyperparameter optimisation software framework Optuna^18^ and its function get_param_importances. Optuna uses the fANOVA importance evaluator that fits a random forest regression model to predict the objective value given the hyperparameter combination. The objective value used in this assessment for a hyperparameter combination $\theta$ is the $L_{2}$-norm of our vector-valued function $f$ used in the hyperparameter optimisation step (Eq. (S8)). We also assess hyperparameter importance for each of the three average normalised absolute clustering errors included in our multi-objective function $f$ (i.e. $\bar{e}_{H}^{\left( \theta\right)}\left( 20 \right)$, $\bar{e}_{H}^{\left( \theta\right)}\left( 40 \right)$ and $\bar{e}_{H}^{\left( \theta\right)}\left( 60 \right)$, Eq. (S6)).

To compare the distributions of hyperparameter values amongst optimised hyperparameter sets of homogeneous and inhomogeneous point patterns, we perform Fisher’s exact test using the MATLAB function FisherExactTest^19^. We use a significance value of $\alpha=0.05$.

M12. Marked Point Bootstrapping

To define the 95% confidence interval of Ripley’s $K$-function $K_{gk}\left( r \right)$ for a reference point pattern, we follow the marked point bootstrapping method with blocks established by Loh and Stein^20,21^. Specifically, for each bootstrap sample,

1. We randomly sample $\phi_{B}$ square blocks of side length $\phi_{I}$,
2. We sum the marks of all the points of type $g$ that fall within the blocks, with toroidal wrapping,
3. We divide this sum by the total number of points selected.

We note that a point can fall within multiple boxes in the second step, in which case it is counted multiple times. Let $x_{i}$ be a point of type $g$. Its mark $m_{i}$ depending on the homogeneity status and if $g=k$ (univariate case) is indicated in Table S5.

|  |  | **Homogeneity Status** | |
| --- | --- | --- | --- |
|  |  | Homogeneous | Inhomogeneous |
| **Ripley’s**  $\boldsymbol{K}$**-function** | Univariate $\left( g=k \right)$ | $\frac{A\mathcal{(D)}}{\left( N_{g}-1 \right)}\sum_{\begin{aligned} x_{j}\in X_{g} \\ x_{j}\neq x_{i} \end{aligned}} \frac{1}{w_{ij}}\cdot\mathbf{1}_{\left( d_{ij}\leq r \right)}$ | $\sum_{\begin{aligned} x_{j}\in X_{g} \\ x_{j}\neq x_{i} \end{aligned}} \frac{1}{\hat{\lambda}_{g}\left( x_{i} \right)}\cdot\frac{1}{\hat{\lambda}_{k}\left( x_{j} \right)}\cdot\frac{1}{w_{ij}}\cdot\mathbf{1}_{\left( d_{ij}\leq r \right)}$ |
|  | Multivariate $\left( g\neq k \right)$ | $\frac{1}{\hat{\lambda}_{k}}\sum_{x_{j}\in X_{k}} \frac{1}{w_{ij}}\cdot\mathbf{1}_{\left( d_{ij}\leq r \right)}$ | $\sum_{x_{j}\in X_{k}} \frac{1}{\hat{\lambda}_{g}\left( x_{i} \right)}\cdot\frac{1}{\hat{\lambda}_{k}\left( x_{j} \right)}\cdot\frac{1}{w_{ij}}\cdot\mathbf{1}_{\left( d_{ij}\leq r \right)}$ |

Table S5. Marks used in marked point bootstrapping with blocks.

Let $K_{gk}^{\left( b \right)}\left( r \right)$ be the $b$-th bootstrap $K_{gk}\left( r \right)$ value ($b=1,\cdots,B$; $B=1000$). Then, the $95\%$ confidence interval for $K_{gk}\left( r \right)$ is given by

$$CI_{95}\left( K_{gk}\left( r \right) \right)=\left[ Q_{0.025}\left( \left\{ K_{gk}^{\left( b \right)}\left( r \right) \right\}_{b=1,\cdots,B} \right),Q_{0.975}\left( \left\{ K_{gk}^{\left( b \right)}\left( r \right) \right\}_{b=1,\cdots,B} \right) \right],$$

where $Q_{\alpha}\left( \left\{ K_{gk}^{\left( b \right)}\left( r \right) \right\}_{b=1,\cdots,B} \right)$ is the $\alpha$-th quantile of the bootstrap values. The 95% confidence interval for Ripley’s $H$-function $H_{gk}\left( r \right)$ is simply the $H$-transformation of $CI_{95}\left( K_{gk}\left( r \right) \right)$:

$$CI_{95}\left( H_{gk}\left( r \right) \right)=\sqrt{\frac{CI_{95}\left( K_{gk}\left( r \right) \right)}{\pi}}-r.$$

Here, we set $\phi_{I}$ as the estimated interaction length between points. Loh and Stein^20^ have found marked point bootstrapping to be minimally sensitive to block size. They suggest that the total coverage area of blocks should equal to the total area of the domain (i.e. $A\left( \mathcal{D} \right)$). This means $\phi_{B}=\left[ \frac{A\left( \mathcal{D} \right)}{\phi_{I}^{2}} \right]$.

M13. Quantification of Exposure to Cell Types

In an ABM, cells interact with other cells localised within their local neighbourhood. In this context, it is important that SHAPPE adequately reproduces this intra- and inter-cell-type exposure. We first define the exposure matrix $\Sigma$ of size $G\times G$, where $G=6$ in our glioblastoma case study. $\Sigma\left( g,k \right)$ (i.e., the element at the $g$-th row and the $k$-th column) contains the fraction of cells of type $g$ that have at least one cell of type $k$ in their neighbourhood. These cells are considered *exposed* to type $k$. We define the neighbourhood of a cell as the disk of radius $50\mu m$ centered around the cell. This is equal to our estimated interaction scale and to approximately $4.7$ times the radius of a glioblastoma cell. This choice is also coherent with Jenner and al.^12^ where they define the radius of a cell neighbourhood as four times the average radius of a glioblastoma cell. Let $\Sigma^{\left( IMC \right)}$ and $\Sigma^{\left( s \right)}$ ($s=1,\cdots,N_{rep}$) be the exposure matrices $\Sigma$ evaluated on $X^{\left( IMC \right)}$ and $N_{rep}$ synthetic reconstructions $X^{\left( s \right)}$. The dissimilarity in exposure is given by

$$e_{\Sigma}=\sum_{g=1}^{G} \mathbf{1}_{\left( N_{g}^{\left( IMC \right)}\geq N_{min} \right)}\cdot\sum_{k=1}^{G} \mathbf{1}_{\left( N_{k}^{\left( IMC \right)}\geq N_{min} \right)}\cdot\left( \frac{\sum_{s=1}^{N_{rep}} \Sigma^{\left( s \right)}\left( g,k \right)}{N_{rep}}-\Sigma^{\left( IMC \right)}\left( g,k \right) \right).$$

Similarly, we define the exposure level matrix $\Pi$ of size $G\times G$ to be the average number of cells of type $k$ in the neighbourhood of a cell of type $g$. For cells of type $g$, this quantifies the average exposure level to type $k$. Let $\Pi^{\left( IMC \right)}$ and $\Pi^{\left( s \right)}$ ($s=1,\cdots,N_{rep}$) be the exposure level matrix $\Pi$ evaluated on $X^{\left( IMC \right)}$ and $N_{rep}$ synthetic reconstructions $X^{\left( s \right)}$. The dissimilarity in exposure level is given by

$$e_{\Pi}=\sum_{g=1}^{G} \mathbf{1}_{\left( N_{g}^{\left( IMC \right)}\geq N_{min} \right)}\cdot\sum_{k=1}^{G} \mathbf{1}_{\left( N_{k}^{\left( IMC \right)}\geq N_{min} \right)}\cdot\left( \frac{\sum_{s=1}^{N_{rep}} \Pi^{\left( s \right)}\left( g,k \right)}{N_{rep}}-\Pi^{\left( IMC \right)}\left( g,k \right) \right).$$

M14. Efficacy and Validity Assessment of SHAPPE

To validate the efficacy of our novel algorithm and its validity, we compared it to CSR for homogeneous point patterns and ECSR for inhomogeneous point patterns as baselines. Let $m$ be a dissimilarity metric evaluated on $N_{rep}$ synthetic reproductions generated with SHAPPE from an original IMC point pattern $X^{\left( IMC \right)}$ (for example, we could consider $m=e_{NN}$). We denote $m^{\left( R \right)}$ to be the same metric evaluated on $N_{rep}$ configurations of the random baseline method. The change in metric $m$ is therefore calculated by:

$$\Delta\left( m \right)=m^{\left( R \right)}-m,$$

while the relative change is given by

$$\frac{\Delta\left( m \right)}{m^{\left( R \right)}}=\frac{m^{\left( R \right)}-m}{m^{\left( R \right)}}.$$

(S9)

The relative change is only defined if $m^{\left( R \right)}\neq0$. We use relative change for $e_{NN}$, $e_{D}$, $e_{\Sigma}$ and $e_{\Pi}$, and unnormalised change for $e_{H}\left( r \right)$: $H$-function differences under the random baseline can be close to zero for point patterns close to spatial randomness at distance $r$, which would inflate small deviations if using relative change. A positive change value indicates that SHAPPE produces an improvement in $m$ as compared to the random baseline.

M15. Agent-Based Modelling of Untreated Glioblastoma Growth

We derived an ABM from our previous ODE model describing glioblastoma growth and interactions between tumour and immune cells^22^.

M15.1. Cell Types and Interaction Rules

Our ABM considers glioblastoma cells, CD8+ T cells, stromal cells and TAMs (M0, M1 and M2). The main rules for each agent are described in Table S6.

| Agent type | Rules | Cell cycle schematic |
| --- | --- | --- |
| Glioblastoma cell | - Proliferates following a logistic growth with baseline proliferation rate $\beta^{\star}$. Its proliferation rate at time $t$ is given by   $\beta=\beta^{\star}\cdot\left( 1-\frac{D}{D^{\star}} \right)$  where $D$ is the local cell density in its neighbourhood and $D^{\star}$ is cell density at carrying capacity. $D$ is calculated according to  $D=\sum_{\begin{aligned} k\in\{GBM,CD8+T cells, \\ Stromal cells, TAMs) \end{aligned}} \frac{r_{k}}{r_{GBM}}\cdot N_{k}$  where $N_{k}$ and $r_{k}$ are the number of cells of type $k$ in the neighbourhood and their radius, respectively.   - Moves according to a stop-or-go mechanism with sampled persistence times and maximum speed from data-estimated distributions^12,23^. | 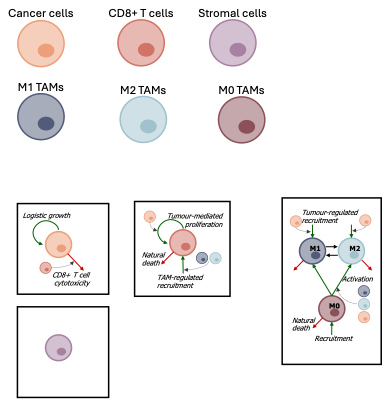 |
| Stromal cell | - Does not proliferate. - Does not move. - Occupies space. | 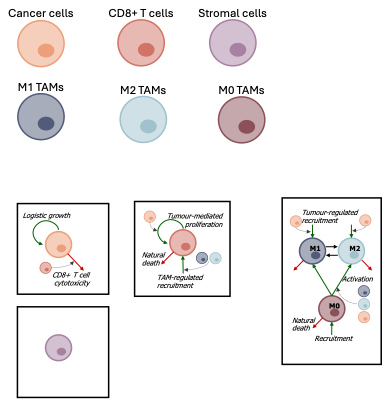 |
| CD8+ T cell | - Dies naturally at a constant rate $\delta_{T}$. - Proliferates at rate $\alpha\cdot I_{Local}\left( PD1,PDL1 \right)$ if there is at least one tumour cell in its neighbourhood. $I\left( PD1_{Local},PDL1_{Local} \right)$ is an inhibitory function given by   $I\left( PD1_{Local},PDL1_{Local} \right)=\frac{1}{1+\frac{PD1_{Local}\cdot PDL1_{Local}}{k_{YQ}}},$  where $PD1_{Local}$ and $PDL1_{Local}$ are respectively the concentration of PD1 and PDL1 receptors in its neighbourhood.   - Is recruited at rate $\eta_{1}\cdot\frac{\frac{M1}{M1+M2}}{\eta_{2}+\frac{M2}{M1+M2}}\cdot I\left( PD1_{TME},PDL1_{TME} \right)$ where $M1$ and $M2$ are the number of M1 and M2 TAMs in the TME and $I\left( PD1_{TME},PDL1_{TME} \right)$ is the PD1-PDL1 inhibitory function calculated with the concentrations of PD1 and PDL1 receptors in the entire TME. - Can attach to a tumour cell in its neighbourhood. If attached, it kills the cell at rate $\alpha_{C}$. - Moves at maximal speed $4\frac{\mu m}{\min}$. | 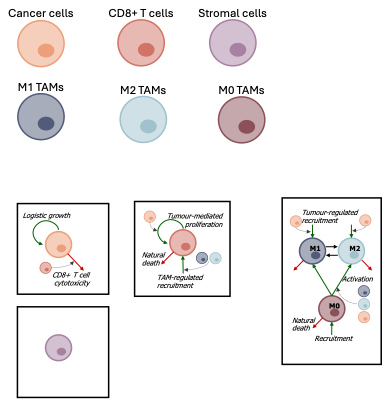 |
| TAMs | - Can be resident (M0) or activated with an anti-tumoral (M1) or pro-tumoral (M2) phenotype. - Dies naturally at rate $\delta_{M0}$ (if resident) or $\delta_{M}$ (if M1 or M2). - M1 switches to M2 phenotype at rate $\mu_{M1}$. - M2 switches to M1 phenotype at rate $\mu_{M2}$. - Resident TAMs can be activated by dead tumour cells, M1 TAMs and M2 TAMs in their neighbourhood. Upon activation, it acquires the M2 (M1) phenotype with probability $q$ ($1-q$). $q$ is the fraction of M1 and M2 TAMs that are pro-tumoral (i.e., M2) at the start of the simulations. - Resident TAMs are recruited in the TME and directly polarised with an M2 (M1) phenotype with probability $q$ ($1-q$) at rate $\lambda_{M0}\cdot\left( f-1 \right)\cdot\frac{G}{K_{G}+G}$. $K_{G}$ is calculated such that, at the start of the simulations, $\frac{G}{K_{G}+G}=0.99$, assuming the TME has reached its tumour-equilibrium state. - Resident TAMs that remain resident are recruited at rate $\lambda_{M0}$ in the TME. - Moves at maximal speed $4\frac{\mu m}{\min}$. | 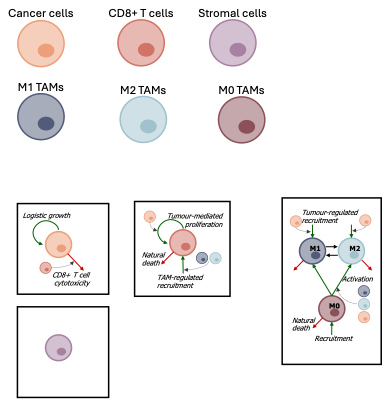 |

Table S6. Description of agents in our glioblastoma-immune system ABM.

Parameters were calculated from our previous models and from literature (Table S7).

| Parameter | Description | Units | Value | Reference |
| --- | --- | --- | --- | --- |
| $\beta^{\star}$ | Maximal proliferation rate of tumour cells | min^-1^ | $1.4014\cdot{10}^{-5}$ | Calculated from Mongeon et al.^23^ |
| $D^{\star}$ | Cell density at carrying capacity | cell | $58.09$ | Calculated from Machado et al.^24^ |
| $\alpha$ | Maximum proliferation rate of CD8+ T cells | min^-1^ | $2.6\cdot{10}^{-5}$ | Calculated from Mongeon and Craig^22^ |
| $\delta_{T}$ | Natural death rate of CD8+ T cells | min^-1^ | $2.8\cdot{10}^{-4}$ | Calculated from Mongeon and Craig^22^ |
| $k_{YQ}$ | Half-saturation constant of PD1 and PDL1 inhibition function | $\left( pg\cdot\mu m^{-2} \right)^{2}$ | $6.31\cdot{10}^{-16}$ | Calculated from Lai and Friedman^25^ |
| $\eta_{1}$ | Maximal recruitment rate of CD8+ T cells by TAMs | min^-1^ | $4.31\cdot{10}^{-2}$ | Calculated from Mongeon and Craig^22^ |
| $\eta_{2}$ | Half-saturation constant of CD8+ T cell recruitment by TAMs | unit-less | $0.1$ | Louzoun et al.^26^ |
| $\alpha_{C}$ | Killing rate of tumour cells by CD8+ T cells | min^-1^ | $6.94\cdot{10}^{-4}$ | Calculated from Storey et al.^27^ |
| $\delta_{M0}$ | Natural death rate of M0 TAMs | min^-1^ | $1.375\cdot{10}^{-5}$ | Calculated from Mongeon and Craig^22^ |
| $\delta_{M}$ | Natural death rate of M1 and M2 TAMs | min^-1^ | $1.39\cdot{10}^{-4}$ | Calculated from Mongeon and Craig^22^ |
| $\mu_{M1}$ | M1-to-M2 phenotype switch rate | min^-1^ | $5.21\cdot{10}^{-5}$ | Calculated from Mongeon and Craig^22^ |
| $\mu_{M2}$ | M2-to-M1 phenotype switch rate | min^-1^ | $3.47\cdot{10}^{-5}$ | Calculated from Mongeon and Craig^22^ |
| $\lambda_{M0}$ | TAM recruitment rate | min^-1^ | $4.23\cdot{10}^{-3}$ | Calculated from Mongeon and Craig^22^ |
| $f$ | Tumour-regulated increase in TAM recruitment | unit-less | $226.7987$ | Calculated from Husain et al.^28^ |

Table S7. Parameters in our glioblastoma-immune system ABM derived from our previous ODE model.

M15.2. Domain and Tumour Radius Increase

In our ABM, the tumour region is defined by the disk of radius $R\mu m$ centered at $\left( 0,0 \right)$. The TME radius is increased following proliferative pressure by tumour cells at its border. For this, we increase the TME radius by $10\mu m$ when more than $50\%$ of the area of the annulus of inner and outer radii $R$ and $R+10 \mu m$ is occupied by tumour cells.

M15.3. Cell movement

To capture more subtle nuances in movement than if cells where restricted to move from one lattice to another,^29^ cells move off-lattice in our model. In PhysiCell, the position of a cell is updated following the second-order Adam’s Bashforth discretisation every $\Delta_{Mech}$ min. Specifically, at time $t+\Delta_{Mech}$, we calculate the cell’s motility vector $\boldsymbol{v}_{mot}\left( t+\Delta_{Mech} \right)$. Let $\boldsymbol{p}$ be the position of the cell at time $t+\Delta_{Mech}$, before updating. We define here two types of cell movement implemented in our ABM, random and spring, to calculate $\boldsymbol{v}_{mot}\left( t+\Delta_{Mech} \right)$.

In random cell movement, we randomly sample a unit vector $\boldsymbol{\phi}_{\boldsymbol{r}}$ on the unit-circle centered at $\boldsymbol{p}$. Let $\Gamma$ be the set of cells in the neighbourhood of the cell of interest and $\boldsymbol{d}_{\boldsymbol{n}}$ the vector from the cell to its neighbour $n\in\Gamma$. Then,

$$\boldsymbol{\phi}_{\boldsymbol{raw}}=\boldsymbol{\phi}_{\boldsymbol{r}}-\sum_{n\in\Gamma} 1_{\left\{ \left\| \boldsymbol{d}_{\boldsymbol{n}} \right\|<{max}_{d} \right\}}\cdot10\cdot\frac{\boldsymbol{d}_{\boldsymbol{n}}}{\left\| \boldsymbol{d}_{\boldsymbol{n}} \right\|}\cdot\left( 1-\frac{\left\| \boldsymbol{d}_{\boldsymbol{n}} \right\|}{{max}_{d}} \right)^{2}$$

$$\boldsymbol{\phi}\left( \boldsymbol{t}\mathbf{+}\boldsymbol{\Delta}_{\boldsymbol{Mech}} \right)=\frac{\boldsymbol{\phi}_{\boldsymbol{raw}}}{\left\| \phi_{raw} \right\|}.$$

(S10)

In Eq. (S10), ${max}_{d}$ is the minimum distance between two cells, set to the sum of the two cells’ radii. Hence, the sum in Eq. (S10) is a repulsion between two cells that are too close from one to another. We then set

$$\boldsymbol{v}_{mot}\left( t+\Delta_{Mech} \right)=\psi\left( t+\Delta_{Mech} \right)\cdot\boldsymbol{\phi}\left( t+\Delta_{Mech} \right),$$

(S11)

where $\psi\left( t+\Delta_{Mech} \right)$ is the cell’s maximum intrinsic speed.

In spring cell movement, we define two types of forces exerted on cells: repulsion forces from neighbours that are too close and spring forces from first neighbours of each type. The latter are analogous to how we place cells in our SHAPPE algorithm. We first calculate a repulsion movement $\nu_{Rep}$ given by

$$\boldsymbol{\nu}_{\boldsymbol{Rep},\boldsymbol{Raw}}=-\sum_{n\in\Gamma} 1_{\left\{ \left\| \boldsymbol{d}_{\boldsymbol{n}} \right\|<{max}_{d} \right\}}\cdot10\cdot\left\| \boldsymbol{d}_{\boldsymbol{n}} \right\|\cdot\left( 1-\frac{\left\| \boldsymbol{d}_{\boldsymbol{n}} \right\|}{{max}_{d}} \right)^{2}$$

$$\boldsymbol{\nu}_{\boldsymbol{Rep}}=\min\left( 0.5\cdot r,\left\| \boldsymbol{\nu}_{\boldsymbol{Rep},\boldsymbol{Raw}} \right\| \right)\cdot\frac{\boldsymbol{\nu}_{\boldsymbol{Rep},\boldsymbol{Raw}}}{\left\| \boldsymbol{\nu}_{\boldsymbol{Rep},\boldsymbol{Raw}} \right\|}.$$

Here, $\Gamma$, $\boldsymbol{d}_{\boldsymbol{n}}$ and ${max}_{d}$ are as in Eq. (S10). The magnitude of $\nu_{Rep}$ is capped to be half the radius of the cell (i.e., $r$). Then, similarly to Eqs. (1) and (2) in the Main Text, let $\boldsymbol{p}^{\left( \boldsymbol{k,1} \right)}$, $d^{\left( k,1 \right)}$ and $s^{\left( k,1 \right)}$ be the position of the cell’s first neighbour of type $k$, the distance to this neighbour and the corresponding target distance sampled from data distributions, respectively. Then, we calculate the spring movement as

$$\boldsymbol{\nu}_{\boldsymbol{Spring}\mathbf{,}\boldsymbol{Raw}}=\sum_{\begin{aligned} k\in\{GBM,CD8+T cells, \\ Stromal cells, TAMs) \end{aligned}} \lambda\cdot\frac{\boldsymbol{p}^{\left( \boldsymbol{k}\mathbf{,1} \right)}-\boldsymbol{p}}{d^{\left( k,1 \right)}}\cdot\left( d^{\left( k,1 \right)}-s^{\left( k,1 \right)} \right)$$

$$\boldsymbol{\nu}_{\boldsymbol{Spring}}=\min\left( \psi\left( t+\Delta_{Mech} \right),\left\| \boldsymbol{\nu}_{\boldsymbol{Spring}\mathbf{,}\boldsymbol{Raw}} \right\| \right)\cdot\frac{\boldsymbol{\nu}_{\boldsymbol{Spring}\mathbf{,}\boldsymbol{Raw}}}{\left\| \boldsymbol{\nu}_{\boldsymbol{Spring}\mathbf{,}\boldsymbol{Raw}} \right\|}.$$

The value of $\lambda$ is the same as in SHAPPE (Section M10). The final motility vector at time $t+\Delta_{Mech}$ is given by

$$\boldsymbol{\nu}_{mot}\left( t+\Delta_{Mech} \right)=\boldsymbol{\nu}_{\boldsymbol{Rep}}+\boldsymbol{\nu}_{\boldsymbol{Spring}}.$$

M15.4. Boundary conditions

We use reflecting boundary conditions when cells exit the TME domain. Specifically, a cell at position $\boldsymbol{p}$ outside the disk of radius $R+10 \mu m$, with $R$ being the current TME radius, will have a moving direction $\boldsymbol{\phi}\left( t+\Delta t \right)$ equal to $-\frac{\boldsymbol{p}}{\left\| \boldsymbol{p} \right\|}$ and we calculate its motility vector using Eq. (S11), independently of the implemented cell movement type.

M15.5. Environmental Inhomogeneity

For inhomogeneous configurations, we extend our ECSR definition and implement environmental constraints by accepting cells in predefined voids with small probability. Here, for each initial configuration obtained using SHAPPE, we use the set of repulsive regions sampled during the generation of random baseline (**Step 0.3**). We also generate additional repulsive regions outside of the initial tumour radius since the latter will be increased with time, following the steps in Section M4.1.

After updating the position of the cell, at every mechanical step $\Delta_{Mech}$, we check if cells enter repulsive regions and, if so, either accept their position or repulse (push) them from the region. Specifically, for $\mathbf{p}=\left( p_{x},p_{y} \right)$ (i.e., the position of the cell that entered a repulsive region $Z$ centered at $\mathbf{v}=\left( v_{x},v_{y} \right)$ and of radius $r_{v}$), we accept the cell within region $Z$ with probability $p_{accept}$:

$$p_{accept}=\underset{Eligibility}{\underbrace{p_{E}}}\cdot\underset{\begin{aligned} Baseline \\ Acceptance Rate \end{aligned}}{\underbrace{p_{B}}}\cdot\underset{Occupancy}{\underbrace{\frac{1}{1+\exp\left( \alpha\cdot\left( \frac{N_{V}}{K_{V}}-1 \right) \right)}}},$$

based on the eligibility of the cell, on the baseline acceptance rate of void regions, and of the current occupancy in $Z$. This probability arises from the assumptions that:

1. ($Eligibility$): A cell that was not just born can only enter a void if it close to its border. Hence,

$$p_{E}=\left\{ \begin{matrix} 1 & \mathrm{if}x was just born \\ \mathbf{1}_{\left( r_{v}-\left\| \mathbf{p}-\mathbf{v} \right\|\leq\delta\right)} & \mathrm{else} \end{matrix} \right..$$

We use $\delta=0.1\cdot r_{V}$ as a boundary threshold. This accounts for cells entering void regions by consuming resources and space at their boundary, except for when a cell that was in a void divided.

2. ($Baseline Acceptance Rate$): We consider $p_{B}$ to be the proportion of cells in void regions in the initial configuration. This accounts for resource availability in repulsive regions compared to the rest of the TME.
3. ($Occupancy$) Let $N_{Z}$ be the current number of cells in the repulsive region $Z$ and let $K_{Z}$ be its carrying capacity, equal to the average cell density in repulsive regions in the initial configuration times the area of $Z$ that intersects with the current tumour region at the current mechanical step. We set $\alpha=3$ to relax the hard boundary of $K_{Z}$ (Figure S7). The term $\frac{1}{1+\exp\left( \alpha\cdot\left( \frac{N_{Z}}{K_{Z}}-1 \right) \right)}$ simulates stronger competition for resources in void regions.

If the cell is rejected from $Z$, it gets repulsed from the center of $Z$ with magnitude $1-\frac{\left\| \mathbf{p}-\mathbf{v} \right\|}{r_{v}}$ and its new position becomes

| $\left( p_{x}+\left( 1-\frac{\left\Vert\mathbf{p}-\mathbf{v} \right\Vert}{r_{v}} \right)\cdot\frac{p_{x}-v_{x}}{\left\Vert\mathbf{p}-\mathbf{v} \right\Vert},p_{y}+\left( 1-\frac{\left\Vert\mathbf{p}-\mathbf{v} \right\Vert}{r_{v}} \right)\cdot\frac{p_{y}-v_{y}}{\left\Vert\mathbf{p}-\mathbf{v} \right\Vert} \right)$ |
| --- |

at every following mechanical update step until it exits $Z$; $\mathbf{p}=\left( p_{x},p_{y} \right)$ is updated as it moves.

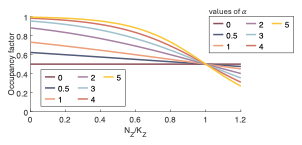

Figure S7. Occupancy factor depending on the value of $\boldsymbol{\alpha}$.

The occupancy factor is defined as $\frac{1}{1+\exp\left( \alpha\cdot\left( \frac{N_{Z}}{K_{Z}}-1 \right) \right)}$, where $N_{Z}$ and $K_{Z}$ are respectively the number of cells in a repulsive region and its cell capacity. $\alpha$ was set to $3$ in simulations.

### Supplementary Results

R1. Two-Step Process is Necessary to Identify Inhomogeneity Associated with Environmental Heterogeneity

We define inhomogeneity as spatial variation in the first-order intensity function that results in regions where points are intrinsically less likely to appear regardless of clustering patterns (see **Step 0.1** of SHAPPE pseudocode in the Main Text). Clustering patterns reflect second-order spatial structures, while the first-order intensity function can be attributed to environmental constraints (e.g., resource availability), among other things. We tested our two-step inhomogeneity identification process by generating three types (A, B and C) of synthetic point patterns. Types A and B both had homogeneous intensity functions, but different point densities. Type C had an inhomogeneous intensity function and the same point density as Type A. The domain for all three types was the square of side length $1000\mu m$ and centered at the origin, i.e. $\mathcal{D=}\left\{ \left( x,y \right)\in\mathbb{R}^{2}:-500\leq x\leq500 \mathrm{and}-500\leq y\leq500 \right\}$. This domain corresponds to the same domain than in most of the IMC samples.

$N$ points were placed sequentially on $\mathcal{D}$ by sampling positions uniformly until acceptance. For type C scenarios with inhomogeneous first-order intensity function, we randomly placed three non-overlapping disks with radii chosen randomly between $50\mu m$ and $100\mu m$ and rejected points falling inside these disks. Otherwise, for points other than the first one, we rejected points with positions with a nearest neighbour further away than $20\mu$m, with probability $s$. Here, $s$ serves to mimic the clustering strength such that $s=0$ implies no clustering at length $20\mu m$ while $s=1$ implies maximal clustering. Clustering strength was randomly chosen from a uniform distribution between $s_{min}$ and $s_{max}$ to introduce variability. Details for each synthetic point pattern type is given in Table S8.

|  | Point Pattern Type | | |
| --- | --- | --- | --- |
|  | A | B | C |
| Number of points $\boldsymbol{N}$ | $2500$ | $500$ | $2500$ |
| Clustering strength range $\left[ \boldsymbol{s}_{\boldsymbol{min}}\boldsymbol{,}\boldsymbol{s}_{\boldsymbol{max}} \right]$ | $\left[ 0.4,0.7 \right]$  (Medium clustering) | $\left[ 0,0.2 \right]$  (Low clustering) | $\left[ 0.4,0.7 \right]$  (Medium clustering) |

Table S8. Three types of point patterns to test our two-step process to identify first-order inhomogeneity.

The chosen number of points were based on the lowest number of points in the IMC samples (486) and on the average number of points (2490.7). The clustering strength was chosen to be of medium strength. For each type of point pattern, we generated 100 different point patterns and classified each of them as homogeneous or inhomogeneous following our two-step process:

**Step 1:**	Test if the point pattern has an empty-space distribution significantly different from what is expected under CSR.

**Step 2:**	Test if the point pattern has at least one zero-density region with an area greater or equal to $A_{0}$.

For the second step, as for the IMC dataset, the density grid was built with bins of side length $bL\approx40\mu m$. We also used the same area threshold, $A_{0}$, equal to the area of a disk of radius $50\mu m$. Point patterns of type A, with a homogeneous first-order intensity function, satisfied the first step in 100% of cases and correctly failed the second step in 89% of cases, resulting in a final homogeneity identification accuracy score of 89% (Figure S8A). The 11% false inhomogeneity identification could be attributed to stronger clustering producing gaps large enough to meet the area threshold $A_{0}$. Point patterns of type B, which also have homogeneous first-order intensity function, failed the first step in 94% of cases (Figure S8B). This false-positive rate (6%) is likely attributed to the significance level used for the global envelope test with the $F$-function ($\alpha=0.05$). 100% of point patterns of type B passed the second step of our identification process, as expected due to the low point density. Taken together, this translates into a homogeneity identification accuracy score of 94% (Figure S8B). Finally, all point patterns of type C (homogeneous intensity functions) passed both steps and were correctly identified as inhomogeneous (Figure S8C). These results show that our two-step process can successfully identify spatial inhomogeneity most likely associated with inhomogeneity in a first-order intensity function and not clustering patterns.

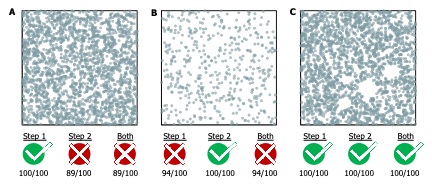

Figure S8. Assessment of the accuracy of our two-step inhomogeneity identification process.

Three types of point patterns were defined as follows: type **A** represents point patterns with medium point density, first-order inhomogeneity and medium clustering between points. Type **B** refers to point patterns with low point density, but no clustering between points and first-order inhomogeneity. Type **C** represents point patterns with average point density, first-order inhomogeneity and medium clustering between points. Indicated are the expected result of each step of our inhomogeneity identification process, as well as the accuracy score estimated by generating 100 point patterns of each type and classifying them as homogeneous or inhomogeneous.

R2. Spatial Correlations in Glioblastoma Tumour Microenvironment

Significant spatial correlation assessed with Ripley’s $K$-function between types $g$ and $k$ at distance $r$ can indicate either clustering or dispersion patterns. In the first case, points of type $g$ see more points of type $k$ in their surroundings than what is expected under randomness, whereas they see fewer points of type $k$ than expected in the second case. To label a significant spatial relationship as clustering or dispersion, we calculated the area between $K_{gk}\left( r \right)$ and the upper bound of the random labeling envelope, $AUC_{UB}$, and the area between $K_{gk}\left( r \right)$ and the lower bound of the random labeling envelope, $AUC_{LB}$. $AUC_{UB}$ and $AUC_{LB}$ were estimated using $r\in\left\{ 40,45,50 \right\}\mu m$, with $50\mu m$ being the estimated interaction scale in our glioblastoma case study. If $AUC_{UB}>AUC_{LB}$, the spatial relationship was classified as a clustering pattern. If $AUC_{UB}<AUC_{LB}$, the spatial relationship was classified as a dispersion pattern.

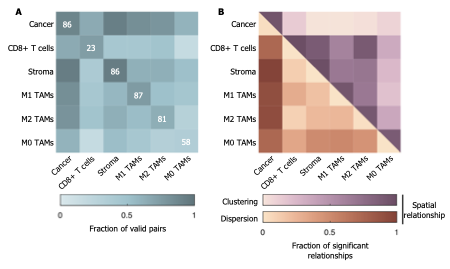

Figure S9. Spatial relationships in glioblastoma tumour microenvironment.

Significant spatial correlations were assessed using a random labeling test. **A.** Fraction of valid pairs of cell types with a significant deviation from the random envelope. A minimum number of cells set to $N_{min}=15$ was enforced on both cell types of a pair to assess correlations. Numbers of the diagonal indicate how many reference point patterns satisfy this criterion. **B.** Fraction of the significant spatial relationships that were classified as clustering patterns (purples, upper right triangle) or dispersion patterns (oranges, lower left triangle). Spatial relationships were classified as clustering if the observed function had an area over the random envelope greater than its area under the random envelope. If the area over the random envelope was less than that under the random envelope, then the spatial relationship was classified as dispersion.

R3. Tumour Microenvironment Reconstructions with SHAPPE
    R3.1. Computational Performance and Optimised Hyperparameters

TME reconstructions were first obtained by generating $N_{rep}=10$ synthetic configurations of the same size and shape as $X^{\left( IMC \right)}$ using SHAPPE. The optimal hyperparameter combination was defined as the combination which minimised average differences with Ripley’s $H$-function at distances $20$, $40$ and $60\mu m$ across the complete dataset. It was found to be $\theta^{\star}=\left[ L,w_{NoCorr},C,\alpha,r_{W},\phi\right]=\left[ 10,0,\mathrm{Gaussian},0,1,10 \right]$. Running Steps 1-6 took a median of 1.17 minutes of execution time (IQR: $0.79$–$1.92$). Equilibrium (converging criteria 1 and 2, **Step 6**) was reached in 99.45% of cases. Each of the SHAPPE reproductions for every IMC point pattern is available in the document “Supplementary SHAPPE Results”.

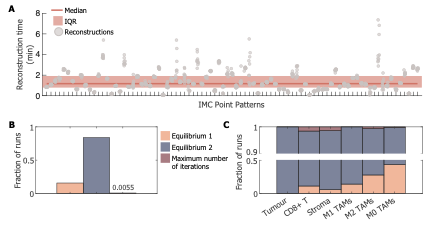

Figure S10. Execution times and convergence of SHAPPE algorithm.

**A.** Execution times of all 10 runs of SHAPPE (same domain size and shape), for all 87 point patterns ($x$-axis). Solid line indicates the median across all runs, and shaded area indicates the interquartile range. **B-C.** A cell type is fixed when the median displacement for points is less than $d_{med}$ units and the 90^th^ quantile is less than $d_{90}$ units (Equilibrium 1, orange), the median net displacement over the last 25 iterations is less than $d_{Net}$ units and $n\geq n_{Eq}$ (Equilibrium 2, dark blue), or when the maximum number of iterations was reached (dark red). **B.** Fraction of cell type placement call that ended in each criterion, all cell types included. **C.** Fractions of cell type placement call that ended in each criterion, depending on the cell type.

R3.2. Importance Analysis of SHAPPE’s Hyperparameters

SHAPPE includes up to eight hyperparameters (Table 1, Main Text). In our case study, we fixed $\lambda$ to a value from the literature and $\phi_{min}$ to the largest cell radius. The other six hyperparameters were varied across ranges of values. We first assessed our hyperparameter optimisation routine by analysing the performance on our Pareto front. Figure S11A shows the performance of all tested hyperparameter combinations with regards to the three metrics included in the loss function (average normalised absolute clustering errors $\bar{e}_{H}\left( 20 \right)$, $\bar{e}_{H}\left( 40 \right)$ and $\bar{e}_{H}\left( 60 \right)$, see Eq. (S6)). The seven Pareto-optimal solutions differed only in copula type, step limit ($\phi$) and repulsion weight ($r_{W}$), three hyperparameters with low estimated importance (Figure S11B). They also had very similar values for $\bar{e}_{H}\left( 20 \right)$, $\bar{e}_{H}\left( 40 \right)$ and $\bar{e}_{H}\left( 60 \right)$. This suggests that the solutions are probably functionally interchangeable. Hyperparameter importance was estimated using the Optuna fANOVA framework. We observe that for the combined score, the number of neighbours $L$ is the most important hyperparameter, which is also true for $\bar{e}_{H}\left( 40 \right)$ and $\bar{e}_{H}\left( 60 \right)$, suggesting that including more neighbours in the positional update is important to reproduce longer-range clustering patterns.

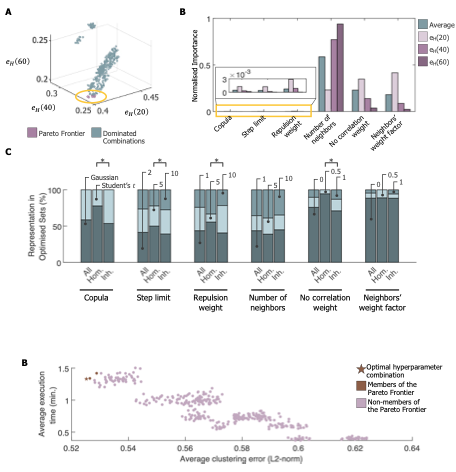

Figure S11. SHAPPE hyperparameter importance analysis.

**A.** Average normalised absolute clustering errors $\bar{e}_{H}\left( 20 \right)$, $\bar{e}_{H}\left( 40 \right)$ and $\bar{e}_{H}\left( 60 \right)$ for all the hyperparameter combinations. **B.** Normalised importance of hyperparameter estimated using the Optuna fANOVA framework. **C.** Distribution of hyperparameter values in combinations optimised for each point pattern. Fisher’s exact test was performed to assess significant associations between hyperparameters and the homogeneity status. $\boldsymbol{*}$ indicates a $p$-value less than $0.05$.

We also observed that Pareto front members have low scores for $\bar{e}_{H}\left( 20 \right)$, $\bar{e}_{H}\left( 40 \right)$ and $\bar{e}_{H}\left( 60 \right)$ (Figure S11A). Because Ripley’s $H$-function is cumulative across radii, the three metrics were found to be significantly positively correlated (Table S9), as expected. However, the correlation was found to be stronger between $\bar{e}_{H}\left( 40 \right)$ and $\bar{e}_{H}\left( 60 \right)$ than between $\bar{e}_{H}\left( 20 \right)$ and the other two metrics. This is coherent with our estimation of the interaction scale $(50\mu m$): capturing clustering patterns in ranges of $40\mu m$ is sufficient to capture those in ranges exceeding this interaction distance. Hence, optimisation at a few well-chosen biologically relevant radii, such as $20$ and $40\mu m$ for this IMC dataset, may be sufficient to adequately reproduce clustering patterns, while most likely reducing computation costs.

| Pearson correlation coefficient ($\boldsymbol{p}$-value) | ${\bar{\boldsymbol{e}}}_{\boldsymbol{H}}\left( \mathbf{20} \right)$ | ${\bar{\boldsymbol{e}}}_{\boldsymbol{H}}\left( \mathbf{40} \right)$ | ${\bar{\boldsymbol{e}}}_{\boldsymbol{H}}\left( \mathbf{60} \right)$ |
| --- | --- | --- | --- |
| ${\bar{\boldsymbol{e}}}_{\boldsymbol{H}}\left( \mathbf{20} \right)$ |  | $0.6985$  $\left( <{10}^{-16} \right)$ | $0.4097$  $\left( <{10}^{-16} \right)$ |
| ${\bar{\boldsymbol{e}}}_{\boldsymbol{H}}\left( \mathbf{40} \right)$ | $0.6985$  $\left( <{10}^{-16} \right)$ |  | $0.9270$  $\left( <{10}^{-16} \right)$ |
| ${\bar{\boldsymbol{e}}}_{\boldsymbol{H}}\left( \mathbf{60} \right)$ | $0.4097$  $\left( <{10}^{-16} \right)$ | $0.9270$  $\left( <{10}^{-16} \right)$ |  |

Table S9. Pearson correlation coefficients between the three metrics used in the optimisation process.

We also ran the optimisation routine separately for each sample. To test whether some hyperparameter values were selected more often than expected by chance, we performed a chi-square goodness-of-fit test for each hyperparameter, with expected counts proportional to the frequency of each value in the hyperparameter search grid. We applied a Benjamini–Hochberg correction on $p$-values with a false discovery rate FDR = $0.05$. Since $\alpha$ is undefined for $L=1$ and was set to $0$ in that case, $\alpha$ was tested on the optimised sets with $L>1$ only ($n=49$). The number of neighbours, $L$, was already identified by Optuna as the most important hyperparameter (Figure S11B), and deviated significantly from expectation (adjusted $p$-value $<{10}^{-13}$; Figure S11C). Indeed, $L=1$ was selected far more often than expected (38/87) and $L=5$ less often (18/87), while $L=10$ was selected at a frequency close to the expectation under random (31/87). This suggests that the neighbourhood size required to reproduce clustering patterns depends on the point pattern. However, since all seven Pareto-optimal solutions use $L=10$, overestimating the neighbourhood size is probably less harmful to performance than underestimating it. The no-correlation weight, $w_{NoCorr}=0$, was also found to be significantly overrepresented (adjusted $p$-value $<{10}^{-14}$), indicating that uncorrelated types (i.e., types that did not appear more or less frequently than under CSR or ECSR) could be ignored when placing points without affecting clustering reproduction in most cases. When more than one neighbour was considered, $\alpha=0$ was significantly overrepresented (adjusted $p$-value $<{10}^{-10}$), suggesting that distant neighbours should contribute as much as nearby neighbours to adequately reproduce clustering patterns.

Finally, we looked at the association between the hyperparameter values and the inhomogeneity status using the sample-specific optimised sets (Figure S11C) and Fisher’s exact test. We again adjusted $p$-values using the Benjamini-Hochberg correction with false discovery rate FDR = 0.05. Significant associations were observed for the repulsion weight, $r_{W}$ ($p=0.0302$), and the weight attributed to uncorrelated pairs of cell types, $w_{NoCorr}$ ($p=0.0471$). A greater proportion of inhomogeneous point patterns require $r_{W}=5$ in their optimised set than homogeneous point patterns, with $r_{W}$ being included in the repulsion from exclusion zones (or attraction to inclusion zones, depending on the ECSR scenario). Homogeneous cases tend to include less uncorrelated pairs of agents in SHAPPE (i.e. $w_{NoCorr}=0$) than inhomogeneous point patterns.

R4. Raw Metrics

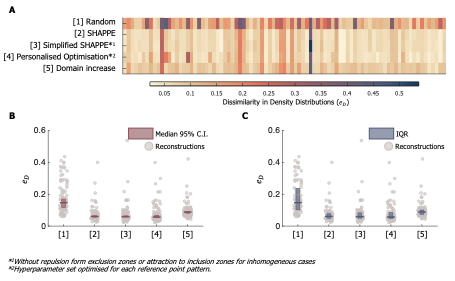

Figure S12. Dissimilarity in density distributions.

**A.** Dissimilarity in density distributions ($e_{D}$) for all 87 reference point patterns ($x$-axis). The rows indicate the reconstruction method. Random refers to CSR (ECSR) for (in)homogeneous point patterns. The darker the color, the higher the dissimilarity is. **B.** 95% confidence interval (C.I.) of the median $e_{D}$ score across 87 reference point patterns, obtained by bootstrapping. **C.** Interquartile range (IQR) of the $e_{D}$ score across 87 reference point patterns. **B-C.** Solid lines: median value.

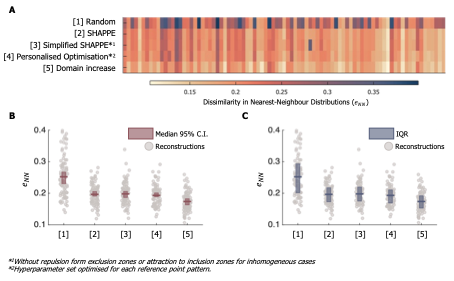

Figure S13. Dissimilarity in nearest-neighbour distributions.

**A.** Dissimilarity in nearest-neighbour distributions ($e_{NN}$) for all 87 reference point patterns ($x$-axis). The rows indicate the reconstruction method. Random refers to CSR (ECSR) for (in)homogeneous point patterns. The darker the color, the higher the dissimilarity is. **B.** 95% confidence interval (C.I.) of the median $e_{NN}$ score across 87 reference point patterns, obtained by bootstrapping. **C.** Interquartile range (IQR) of the $e_{NN}$ score across 87 reference point patterns. **B-C.** Solid lines: median value.

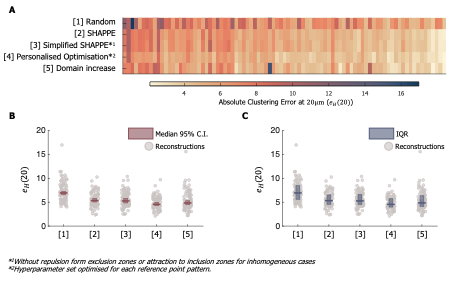

Figure S14. Absolute clustering error at $\boldsymbol{20}\boldsymbol{\mu m}$.

**A.** Absolute clustering error at $20\mu m$ ($e_{H}\left( 20 \right)$) for all 87 reference point patterns ($x$-axis). The rows indicate the reconstruction method. Random refers to CSR (ECSR) for (in)homogeneous point patterns. The darker the color, the higher the dissimilarity is. **B.** 95% confidence interval (C.I.) of the median $e_{H}(20)$ score across 87 reference point patterns, obtained by bootstrapping. **C.** Interquartile range (IQR) of the $e_{H}(20)$ score across 87 reference point patterns. **B-C.** Solid lines: median value.

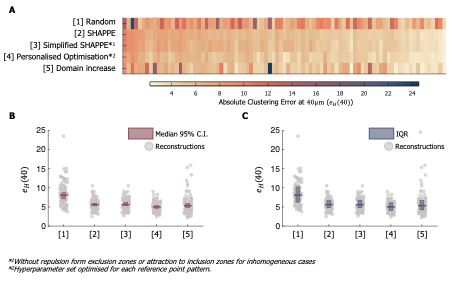

Figure S15. Absolute clustering error at $\boldsymbol{40}\boldsymbol{\mu m}$.

**A.** Absolute clustering error at $40\mu m$ ($e_{H}\left( 40 \right)$) for all 87 reference point patterns ($x$-axis). The rows indicate the reconstruction method. Random refers to CSR (ECSR) for (in)homogeneous point patterns. The darker the color, the higher the dissimilarity is. **B.** 95% confidence interval (C.I.) of the median $e_{H}(40)$ score across 87 reference point patterns, obtained by bootstrapping. **C.** Interquartile range (IQR) of the $e_{H}(40)$ score across 87 reference point patterns. **B-C.** Solid lines: median value.

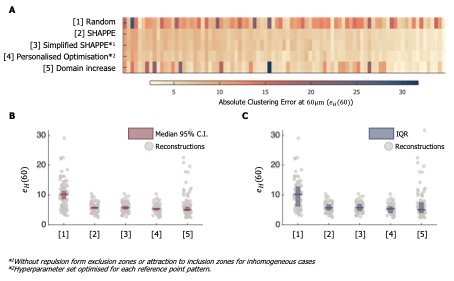

Figure S16. Absolute clustering error at $\boldsymbol{60}\boldsymbol{\mu m}$.

**A.** Absolute clustering error at $60\mu m$ ($e_{H}\left( 60 \right)$) for all 87 reference point patterns ($x$-axis). The rows indicate the reconstruction method. Random refers to CSR (ECSR) for (in)homogeneous point patterns. The darker the color, the higher the dissimilarity is. **B.** 95% confidence interval (C.I.) of the median $e_{H}(60)$ score across 87 reference point patterns, obtained by bootstrapping. **C.** Interquartile range (IQR) of the $e_{H}(60)$ score across 87 reference point patterns. **B-C.** Solid lines: median value.

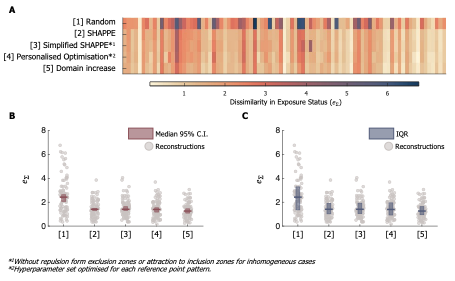

Figure S17. Dissimilarity in exposure status.

**A.** Dissimilarity in exposure status ($e_{\Sigma}$) for all 87 reference point patterns ($x$-axis). The rows indicate the reconstruction method. Random refers to CSR (ECSR) for (in)homogeneous point patterns. The darker the color, the higher the dissimilarity is. **B.** 95% confidence interval (C.I.) of the median $e_{\Sigma}$ score across 87 reference point patterns, obtained by bootstrapping. **C.** Interquartile range (IQR) of the $e_{\Sigma}$ score across 87 reference point patterns. **B-C.** Solid lines: median value.

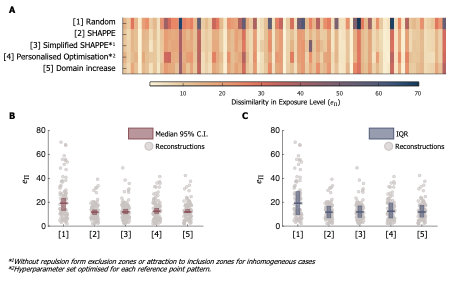

Figure S18. Dissimilarity in exposure level.

**A.** Dissimilarity in exposure status ($e_{\Pi}$) for all 87 reference point patterns ($x$-axis). The rows indicate the reconstruction method. Random refers to CSR (ECSR) for (in)homogeneous point patterns. The darker the color, the higher the dissimilarity is. **B.** 95% confidence interval (C.I.) of the median $e_{\Pi}$ score across 87 reference point patterns, obtained by bootstrapping. **C.** Interquartile range (IQR) of the $e_{\Pi}$ score across 87 reference point patterns. **B-C.** Solid lines: median value.

R5. Variability Accuracy Scores in IMC Reconstructions

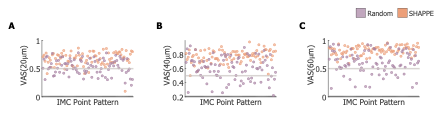

Figure S19. Variability accuracy scores in IMC point pattern reconstructions.

Reference point patterns ordered according to their average $e_{H}\left( r \right)$ score for $r=20$, $40$ and $60\mu m$. The variability accuracy score at distance $r$, $VAS\left( r \right)$, reports the fraction of pairwise Ripley’s $H$-functions at distance $r$ that fell between the 95% confidence interval bounds for cell types satisfying the minimal number of cells threshold (15 cells) for 10 replicates. **A.** $r=20\mu m$. **B.** $r=40\mu m$. **C.** $r=60\mu m$.

R6. Sequential Hooke’s Law Reproduces Exposure Patterns

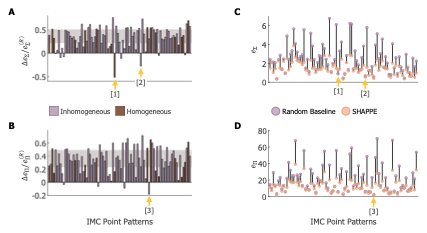

Figure S20. SHAPPE algorithm outperforms random baseline at reproduction of exposure patterns.

**A-B.** Relative change in dissimilarity between SHAPPE and the random baseline for **A.** the exposure matrix $\Sigma$ ($\Delta e_{\Sigma}/e_{\Sigma}^{\left( R \right)}$) and **B.** the exposure level matrix $\Pi$ ($\Delta e_{\Pi}/e_{\Pi}^{\left( R \right)}$). A positive change indicates an improvement with SHAPPE relative to the random baseline. Shaded gray regions show the interquartile range across point patterns. **C-D.** Dissimilarities in **C.** the exposure matrix and **D.** the exposure level matrix for the random baseline (purple) and SHAPPE (orange). Vertical lines connect the two scores for the same point pattern. When a reference point pattern closely agrees with CSR (if homogeneous) or ECSR (if inhomogeneous), the random baseline achieves a low dissimilarity score, which can inflate the magnitude of the relative change if SHAPPE has a slightly higher dissimilarity score. Arrows in A and B indicate reference point patterns with large negative relative changes, and the corresponding arrows in C and D show that both SHAPPE and the random baseline perform well in these cases.
