## Supplementary SHAPPE Results for "SHAPPEing agent-based models to spatial data: Sequential Hooke’s law Algorithm for Point Pattern reconstruction with Environmental constraints"

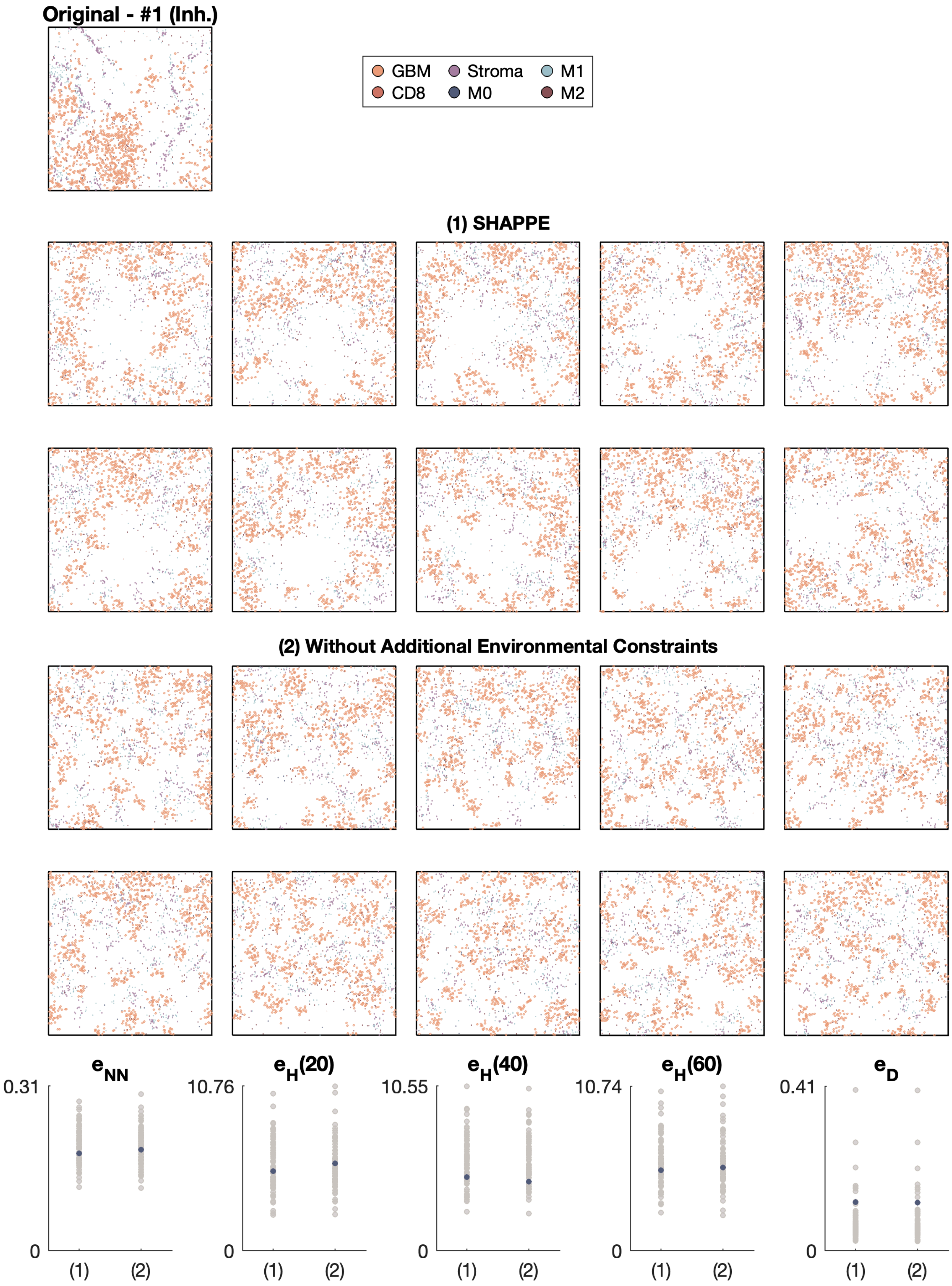

Original - #1 (Inh.)
GBM
Stroma
M1
CD8
M0
M2
(1) SHAPPE
(2) Without Additional Environmental Constraints
$e_{NN}$
$e_H(20)$
$e_H(40)$
$e_H(60)$
$e_D$
0.31
10.76
10.55
10.74
0.41
0
(1)
(2)

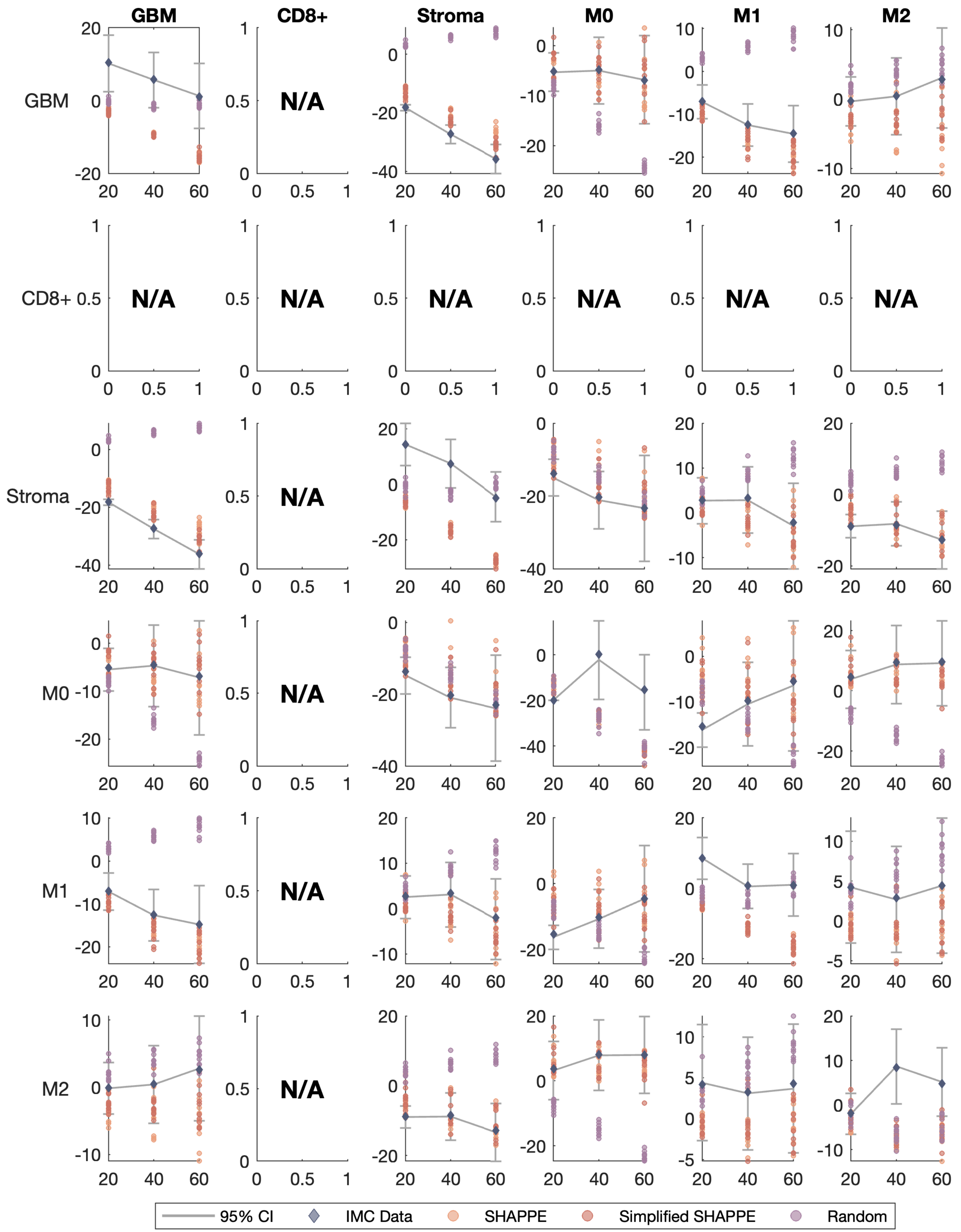
GBM
CD8+
Stroma
M0
M1
M2
N/A
95% CI
IMC Data
SHAPPE
Simplified SHAPPE
Random

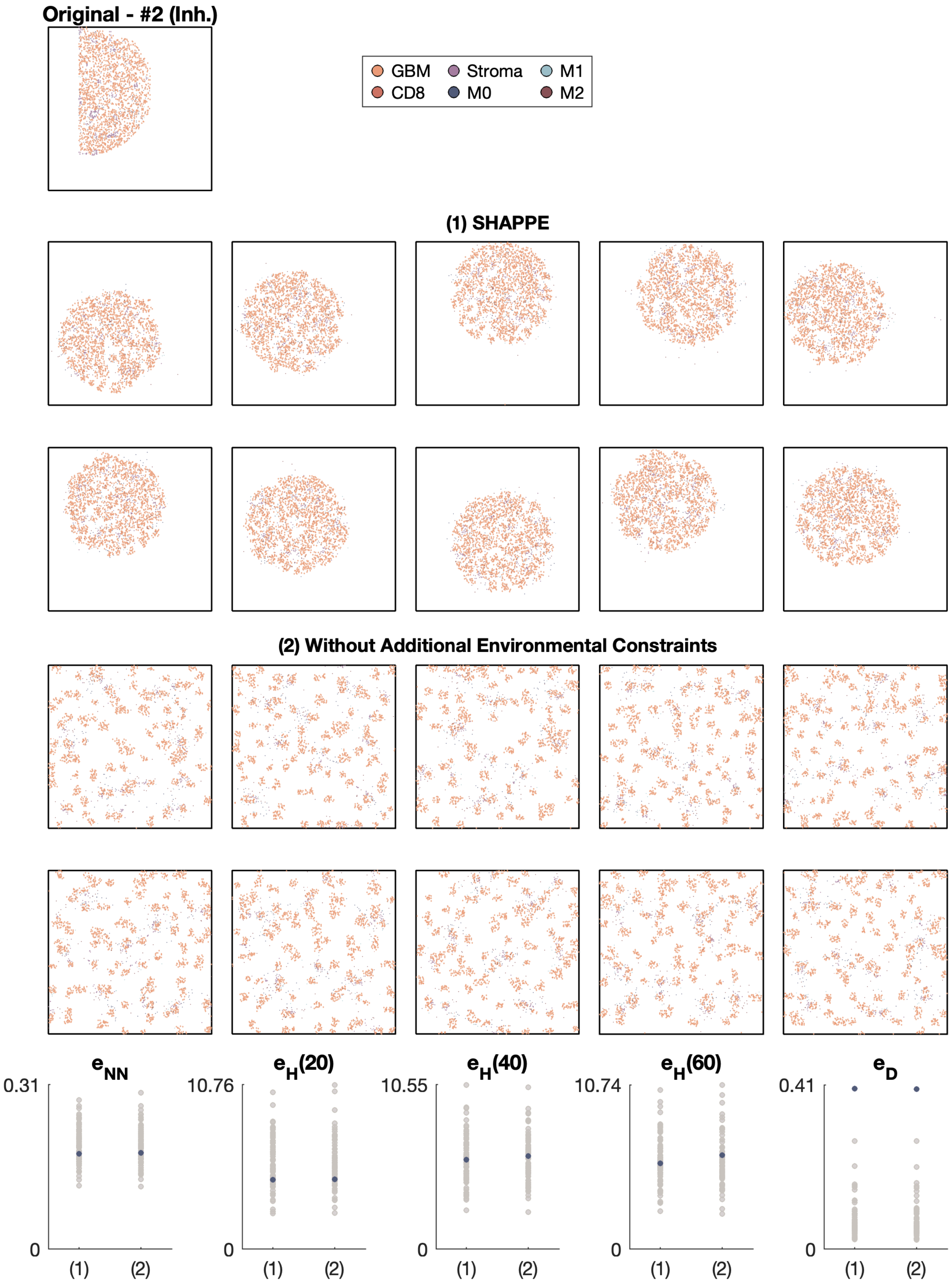
Original - #2 (Inh.)
GBM
Stroma
M1
CD8
M0
M2
(1) SHAPPE
(2) Without Additional Environmental Constraints
e_NN
0.31
0
(1)
(2)
e_H(20)
10.76
0
(1)
(2)
e_H(40)
10.55
0
(1)
(2)
e_H(60)
10.74
0
(1)
(2)
e_D
0.41
0
(1)
(2)

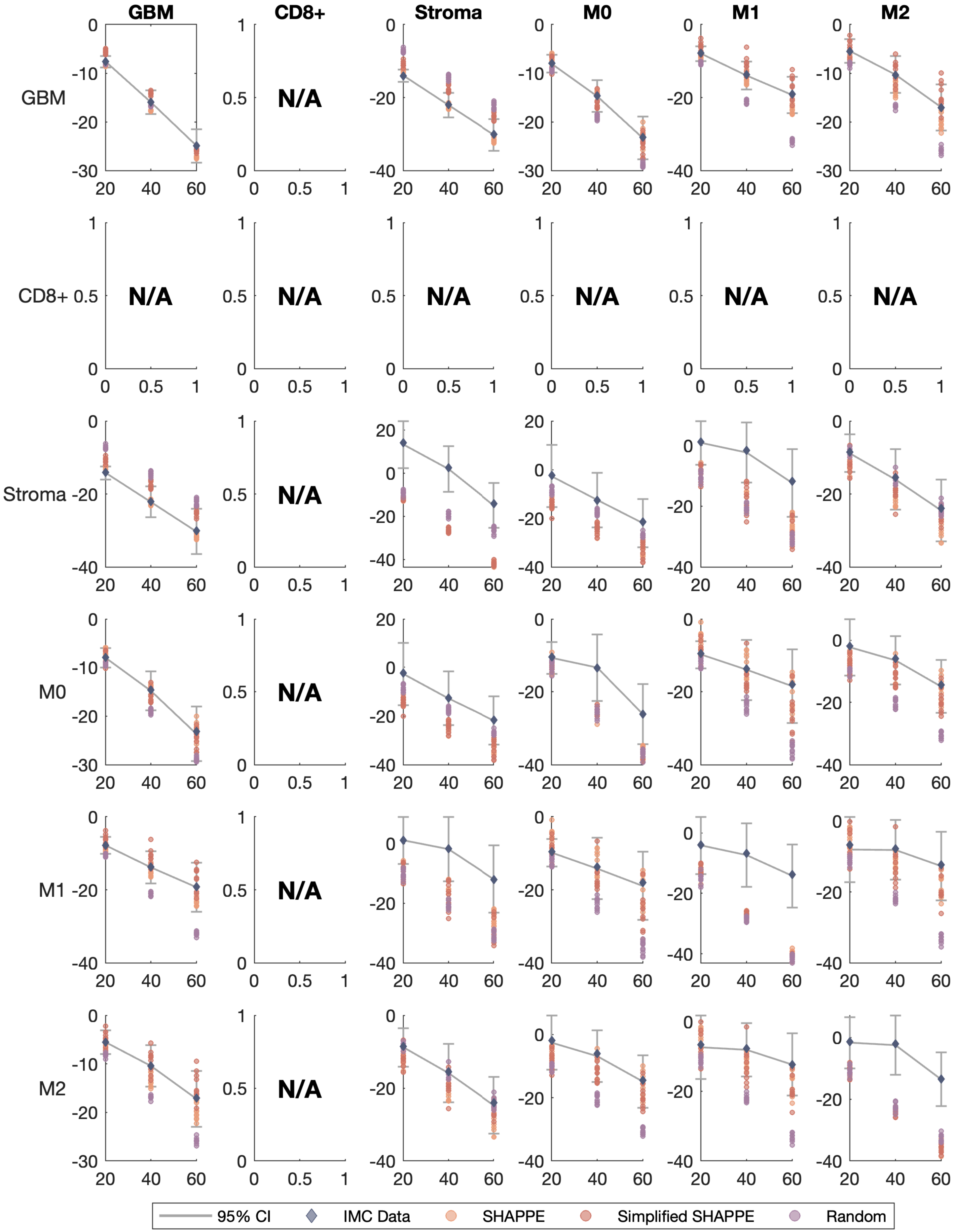

GBM
CD8+
Stroma
M0
M1
M2
N/A
95% CI
IMC Data
SHAPPE
Simplified SHAPPE
Random

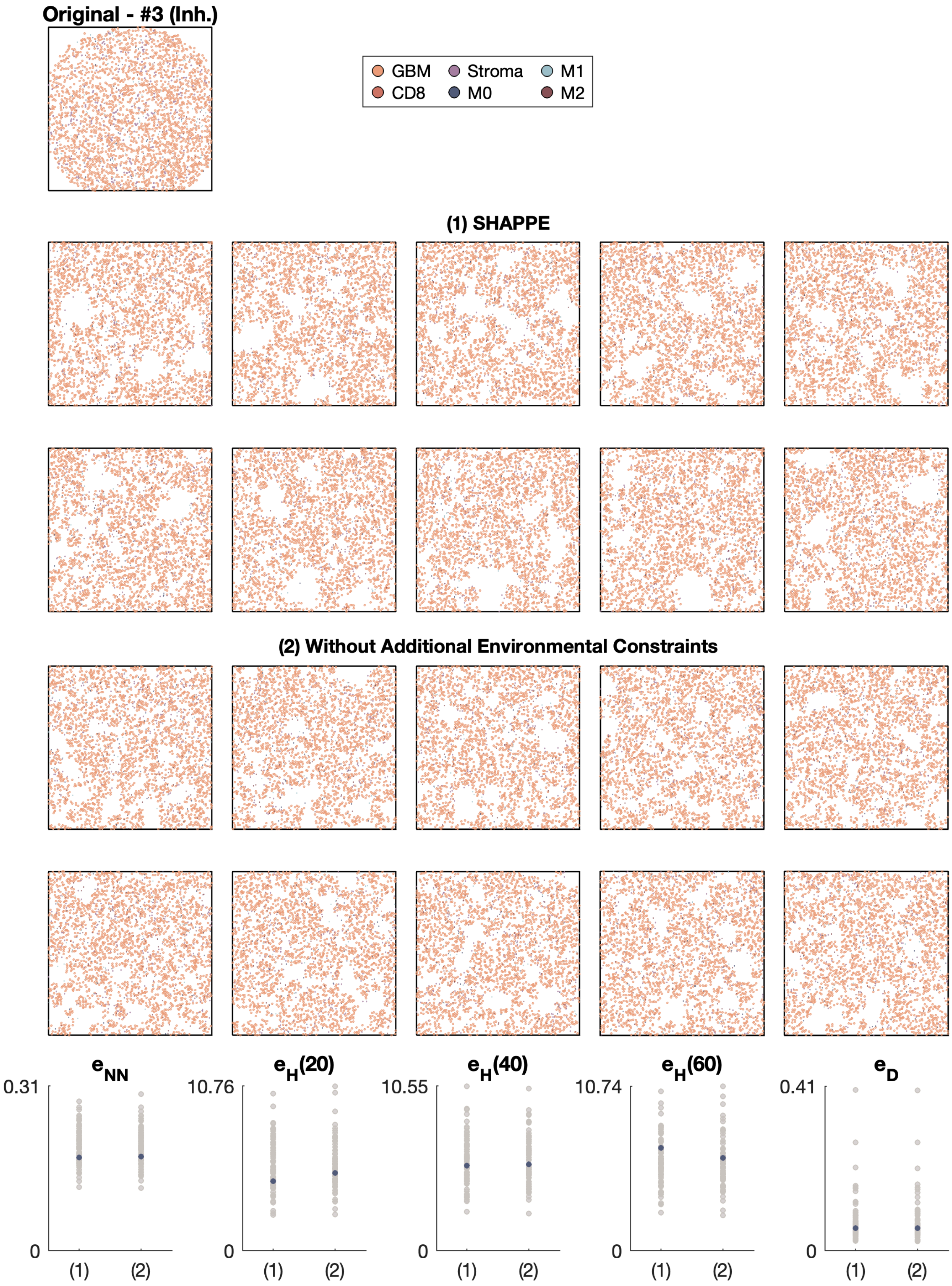
Original - #3 (Inh.)
GBM
Stroma
M1
CD8
M0
M2
(1) SHAPPE
(2) Without Additional Environmental Constraints
$e_{NN}$
$e_H(20)$
$e_H(40)$
$e_H(60)$
$e_D$
0.31
10.76
10.55
10.74
0.41
0
(1)
(2)

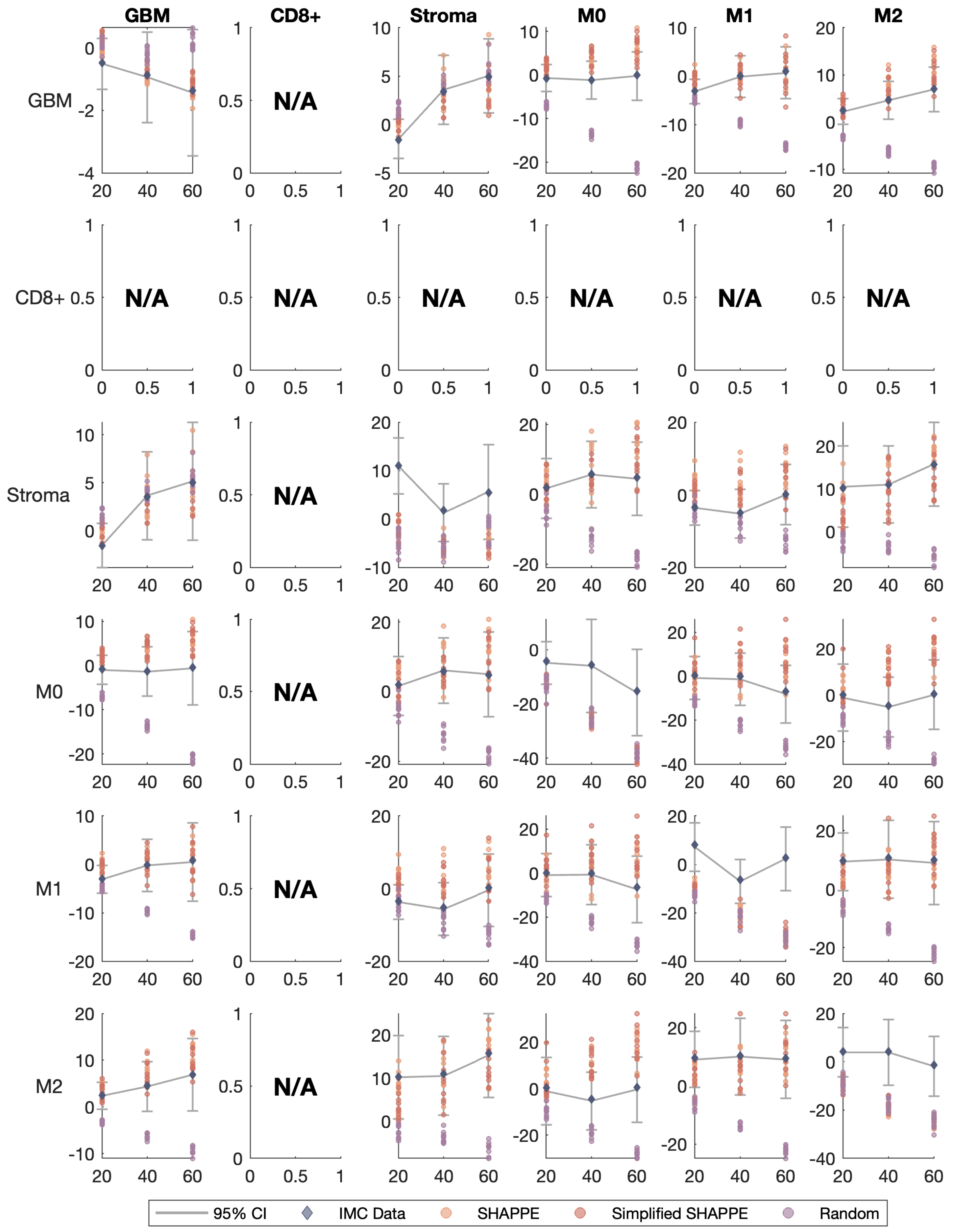
GBM
CD8+
Stroma
M0
M1
M2
GBM
CD8+
Stroma
M0
M1
M2
N/A
95% CI
IMC Data
SHAPPE
Simplified SHAPPE
Random

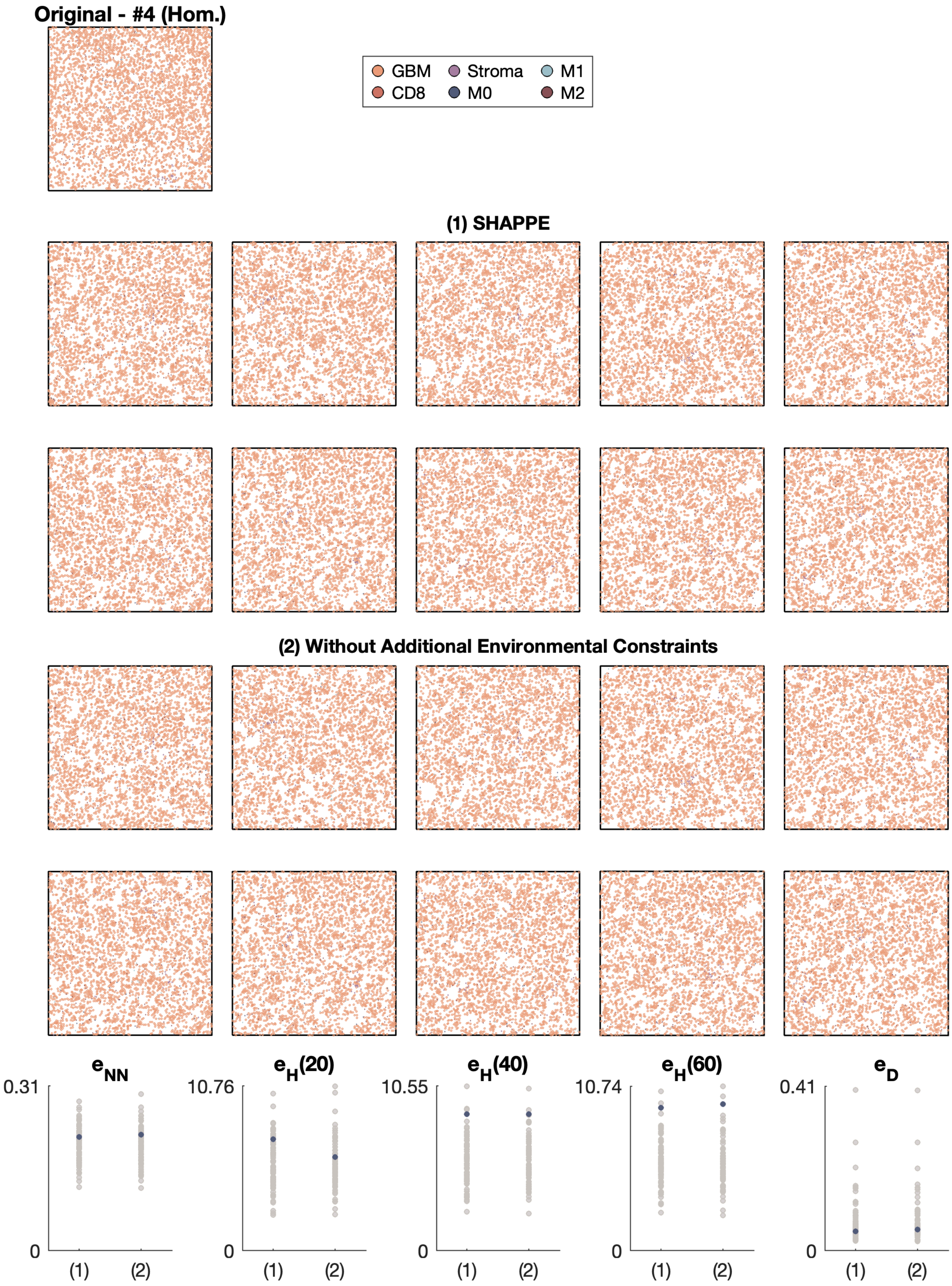

Original - #4 (Hom.)
GBM
Stroma
M1
CD8
M0
M2
(1) SHAPPE
(2) Without Additional Environmental Constraints
$e_{NN}$
$e_H(20)$
$e_H(40)$
$e_H(60)$
$e_D$
0.31
10.76
10.55
10.74
0.41
0
(1)
(2)

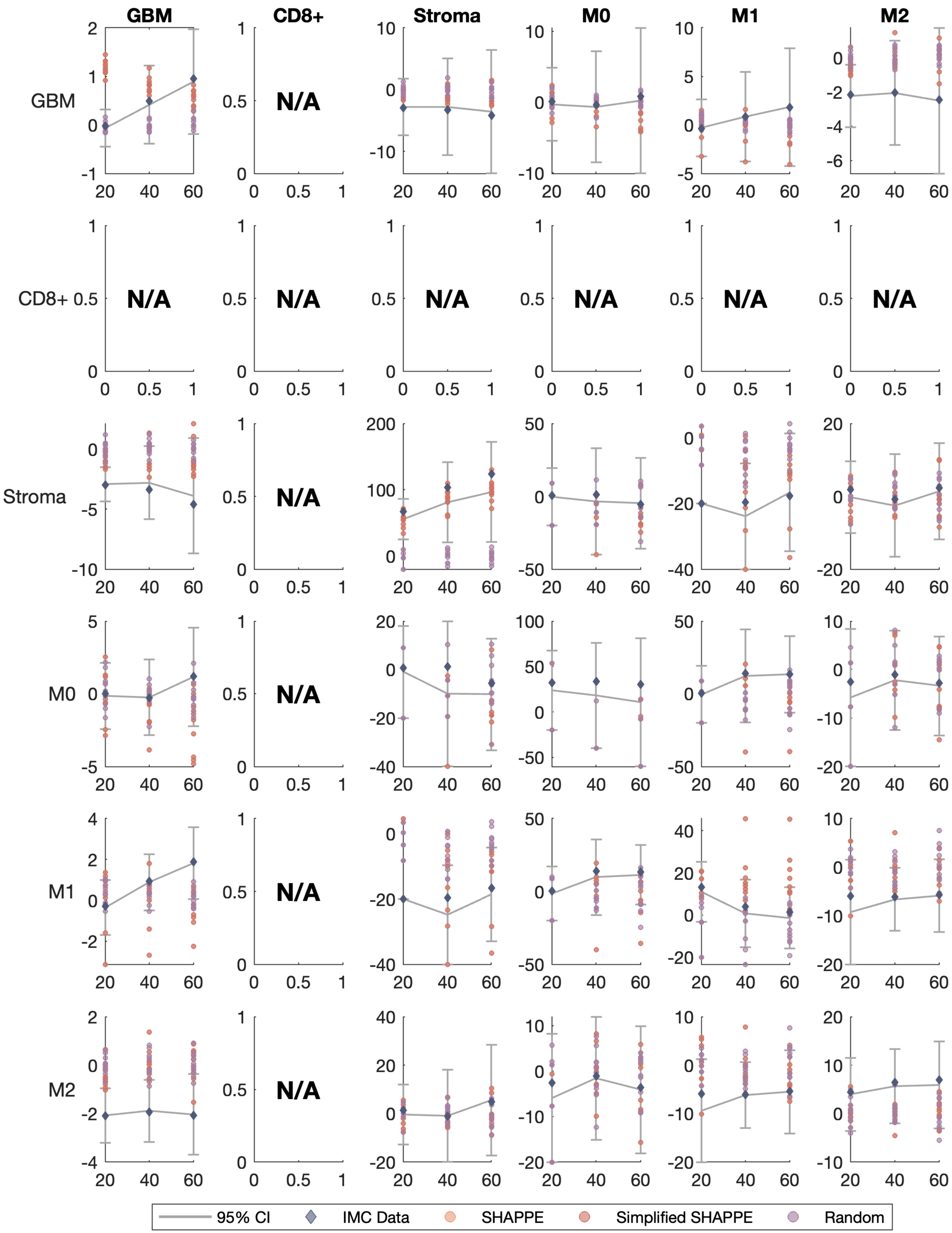

GBM
CD8+
Stroma
M0
M1
M2
N/A
95% CI
IMC Data
SHAPPE
Simplified SHAPPE
Random

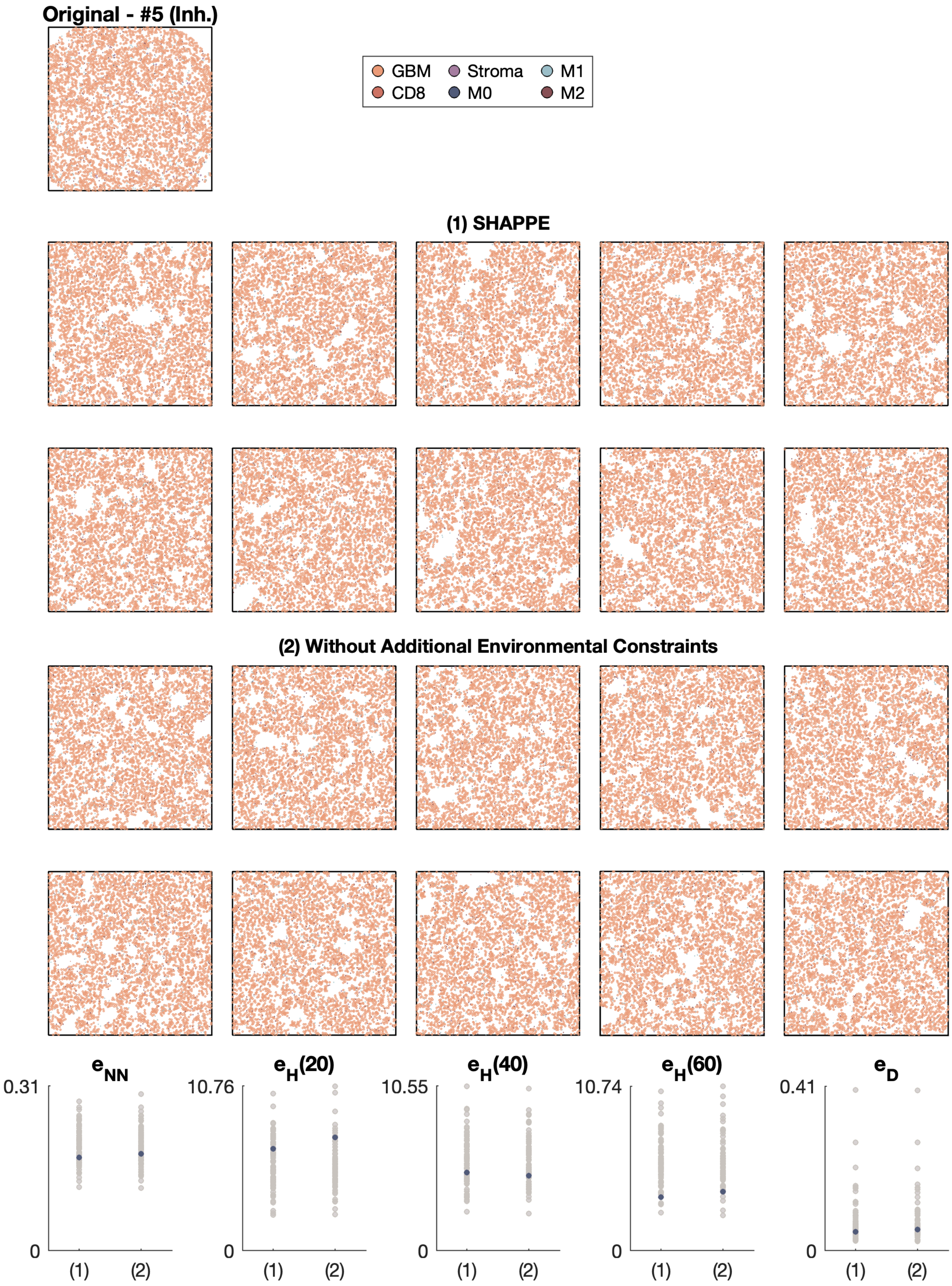
Original - #5 (Inh.)
GBM
Stroma
M1
CD8
M0
M2
(1) SHAPPE
(2) Without Additional Environmental Constraints
$e_{NN}$
0.31
0
(1)
(2)
$e_H(20)$
10.76
0
(1)
(2)
$e_H(40)$
10.55
0
(1)
(2)
$e_H(60)$
10.74
0
(1)
(2)
$e_D$
0.41
0
(1)
(2)

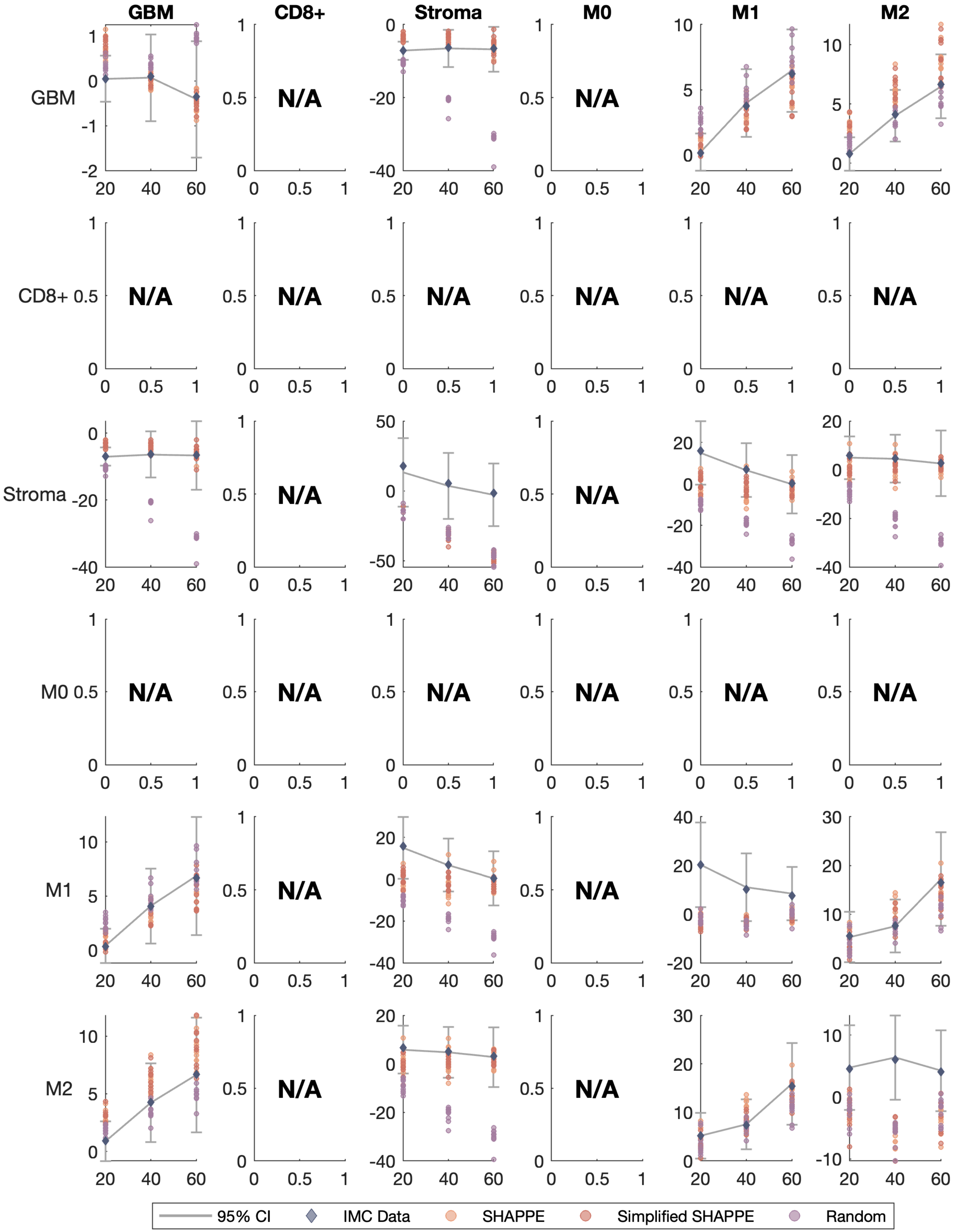
GBM
CD8+
Stroma
M0
M1
M2
N/A
95% CI
IMC Data
SHAPPE
Simplified SHAPPE
Random

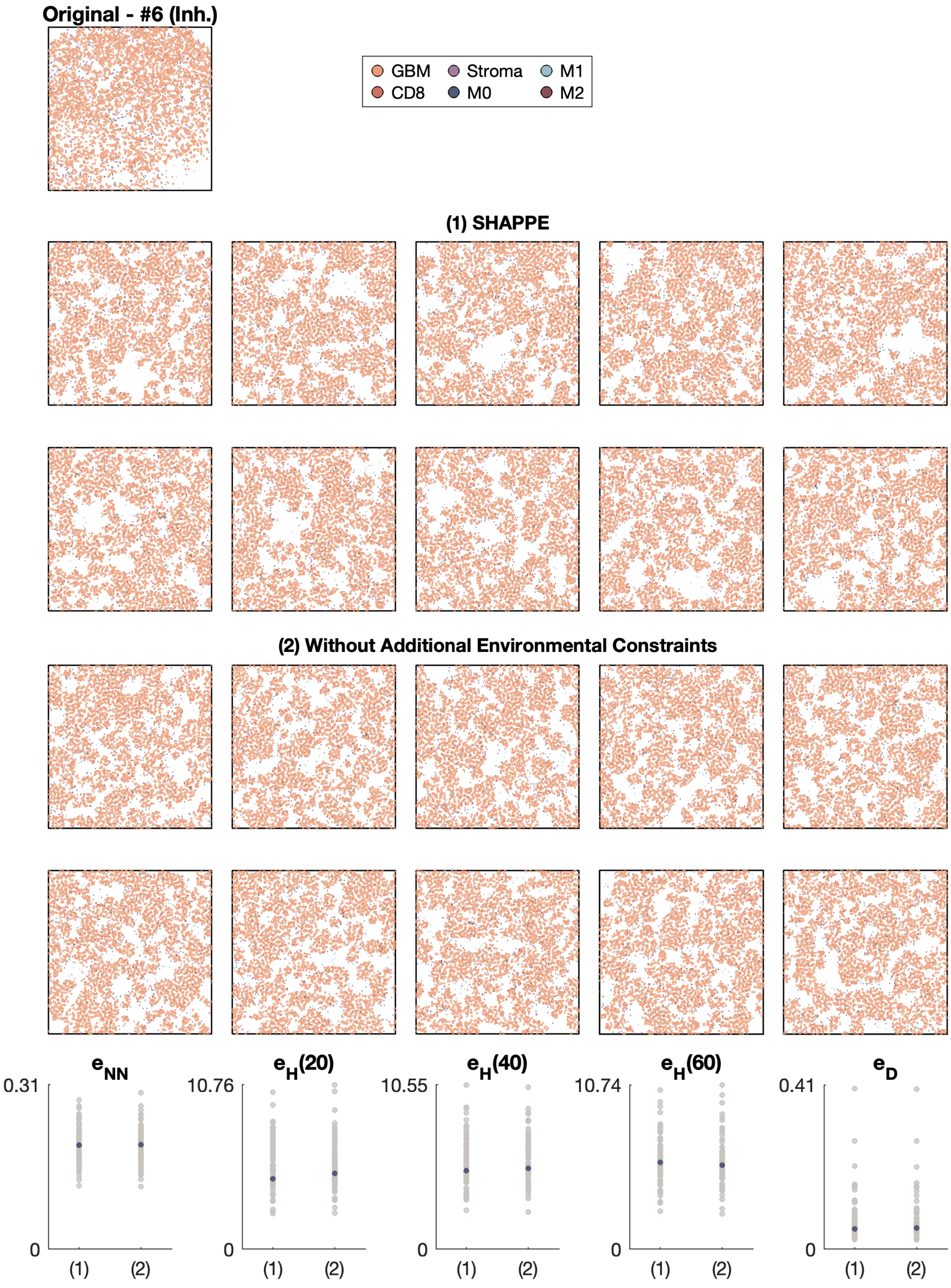

Original - #6 (Inh.)
GBM
CD8
Stroma
M0
M1
M2
(1) SHAPPE
(2) Without Additional Environmental Constraints
$e_{NN}$
0.31
0
(1)
(2)
$e_H(20)$
10.76
0
(1)
(2)
$e_H(40)$
10.55
0
(1)
(2)
$e_H(60)$
10.74
0
(1)
(2)
$e_D$
0.41
0
(1)
(2)

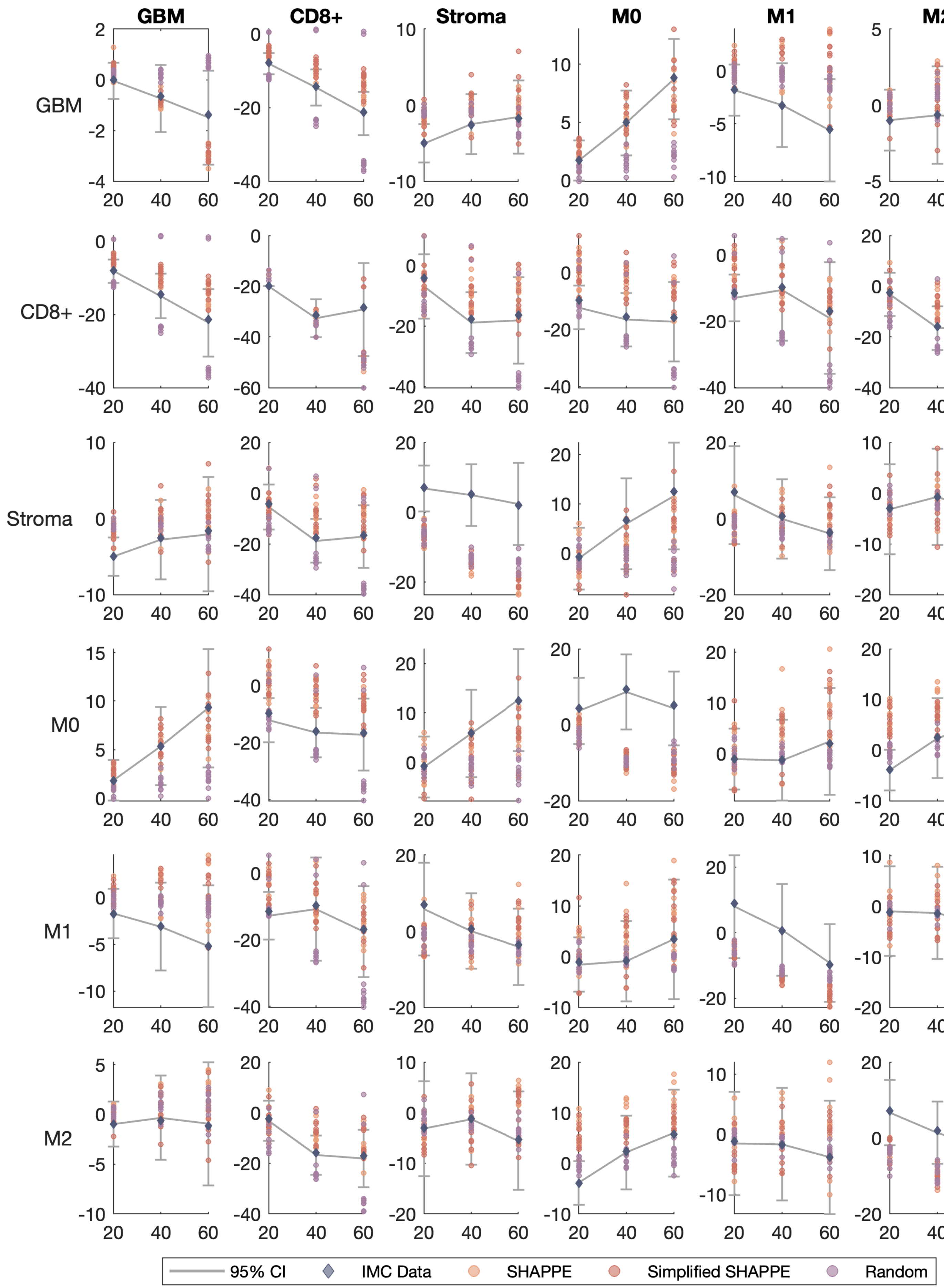
GBM
CD8+
Stroma
M0
M1
M2
95% CI
IMC Data
SHAPPE
Simplified SHAPPE
Random

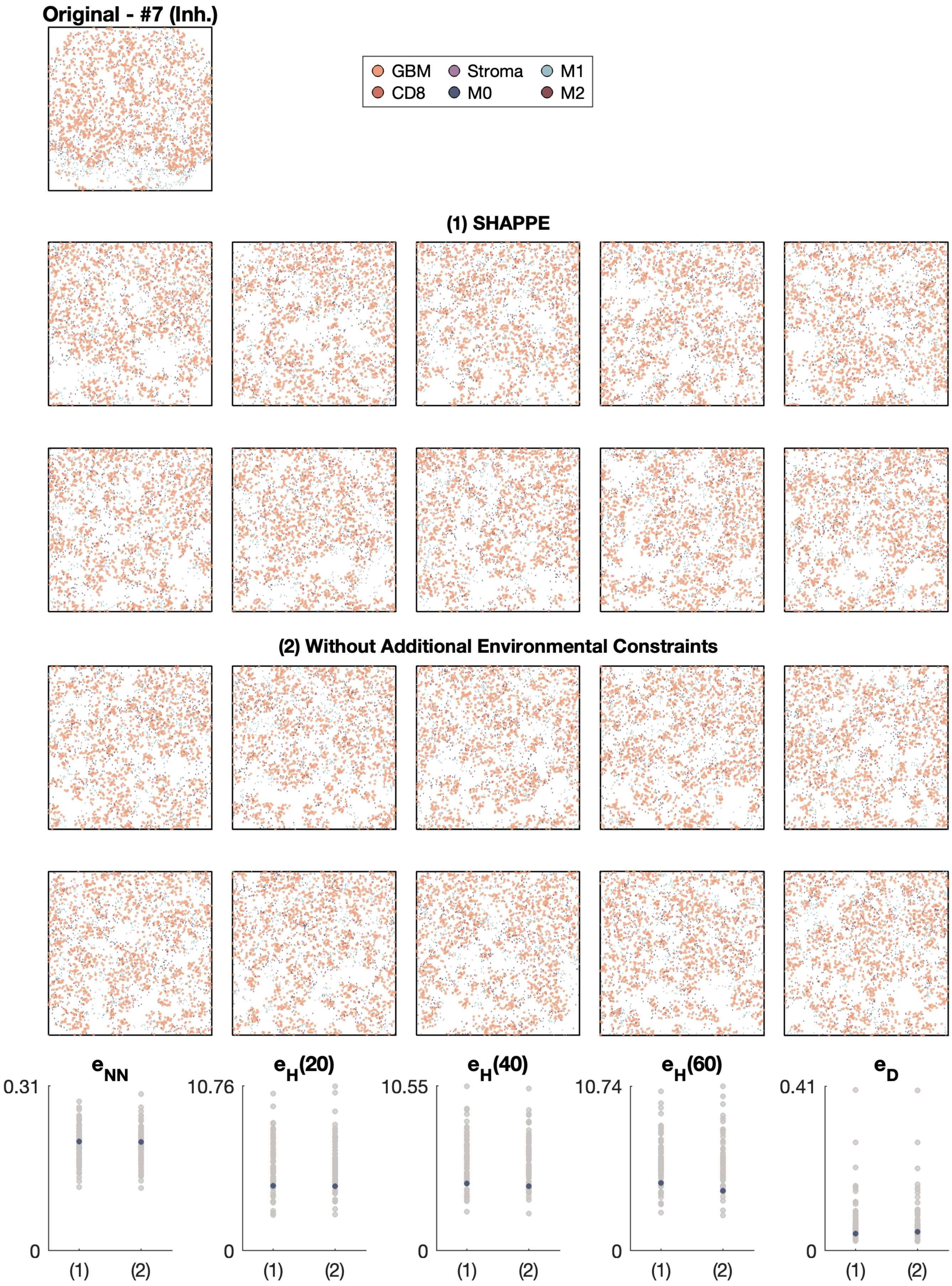

Original - #7 (Inh.)
GBM
CD8
Stroma
M0
M1
M2
(1) SHAPPE
(2) Without Additional Environmental Constraints
$e_{NN}$
$e_H(20)$
$e_H(40)$
$e_H(60)$
$e_D$
0.31
10.76
10.55
10.74
0.41
0
(1)
(2)

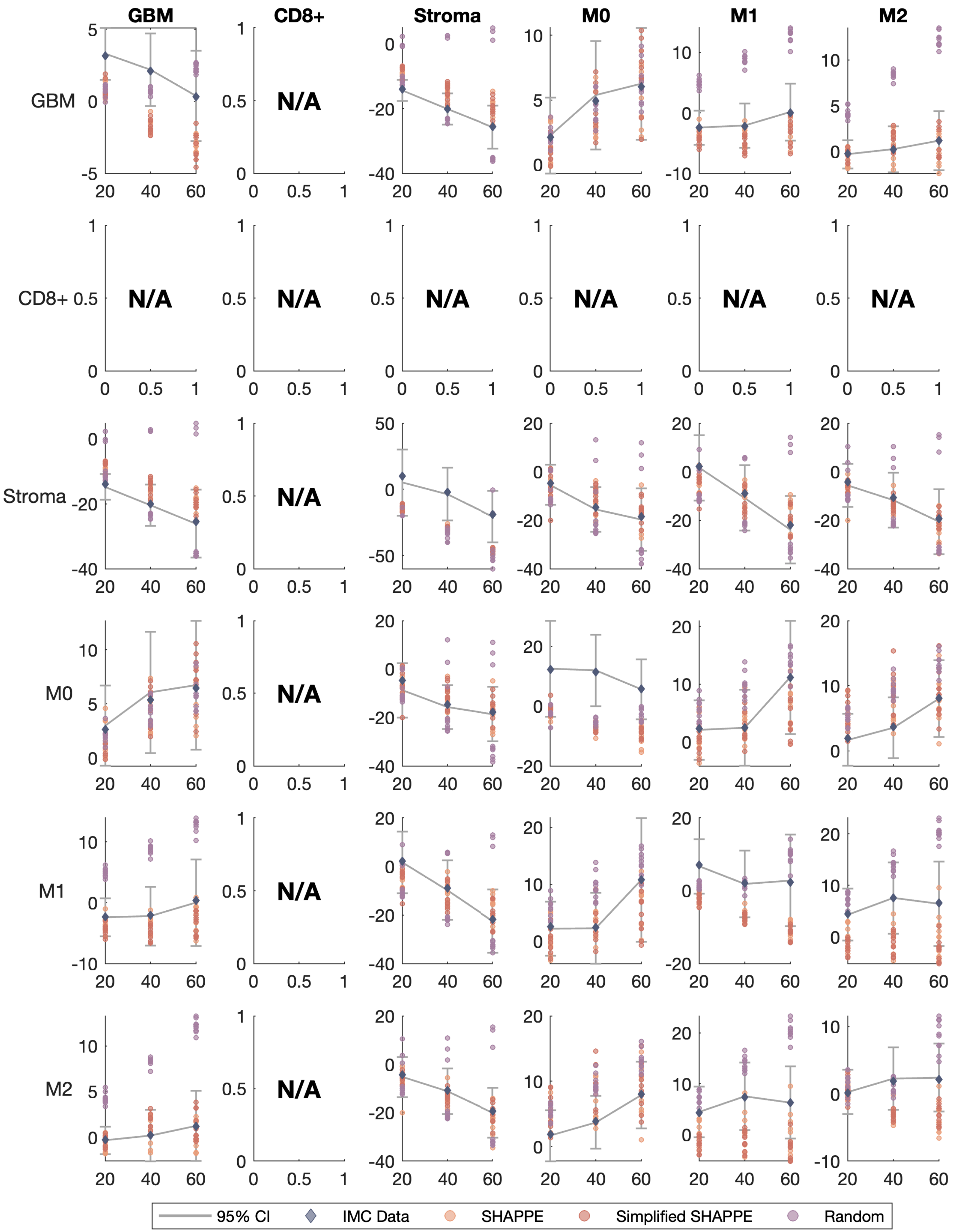

GBM
CD8+
Stroma
M0
M1
M2
N/A
95% CI
IMC Data
SHAPPE
Simplified SHAPPE
Random

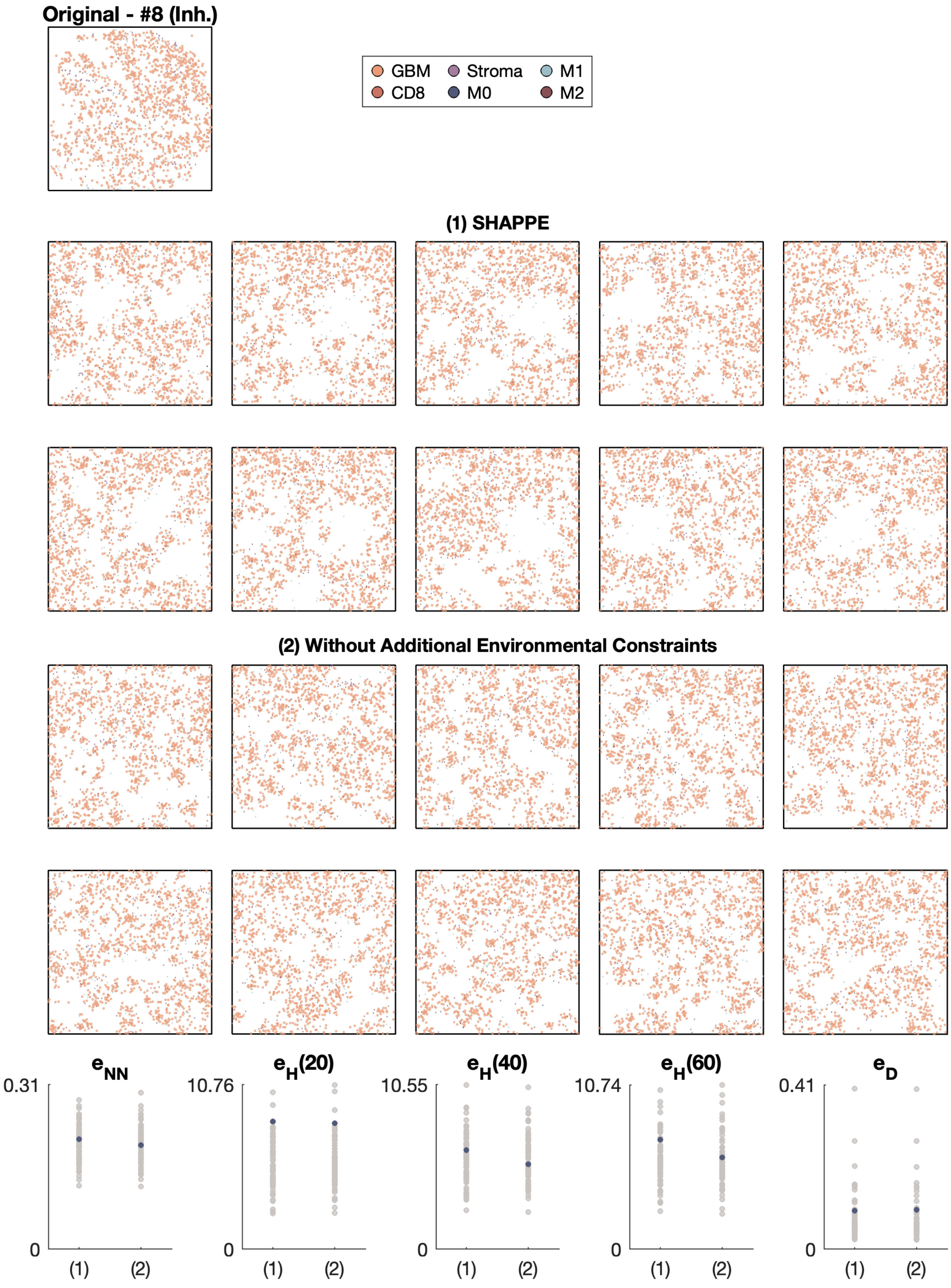

Original - #8 (Inh.)
GBM
CD8
Stroma
M0
M1
M2
(1) SHAPPE
(2) Without Additional Environmental Constraints
$e_{NN}$
$e_H(20)$
$e_H(40)$
$e_H(60)$
$e_D$
0.31
10.76
10.55
10.74
0.41
0
(1)
(2)

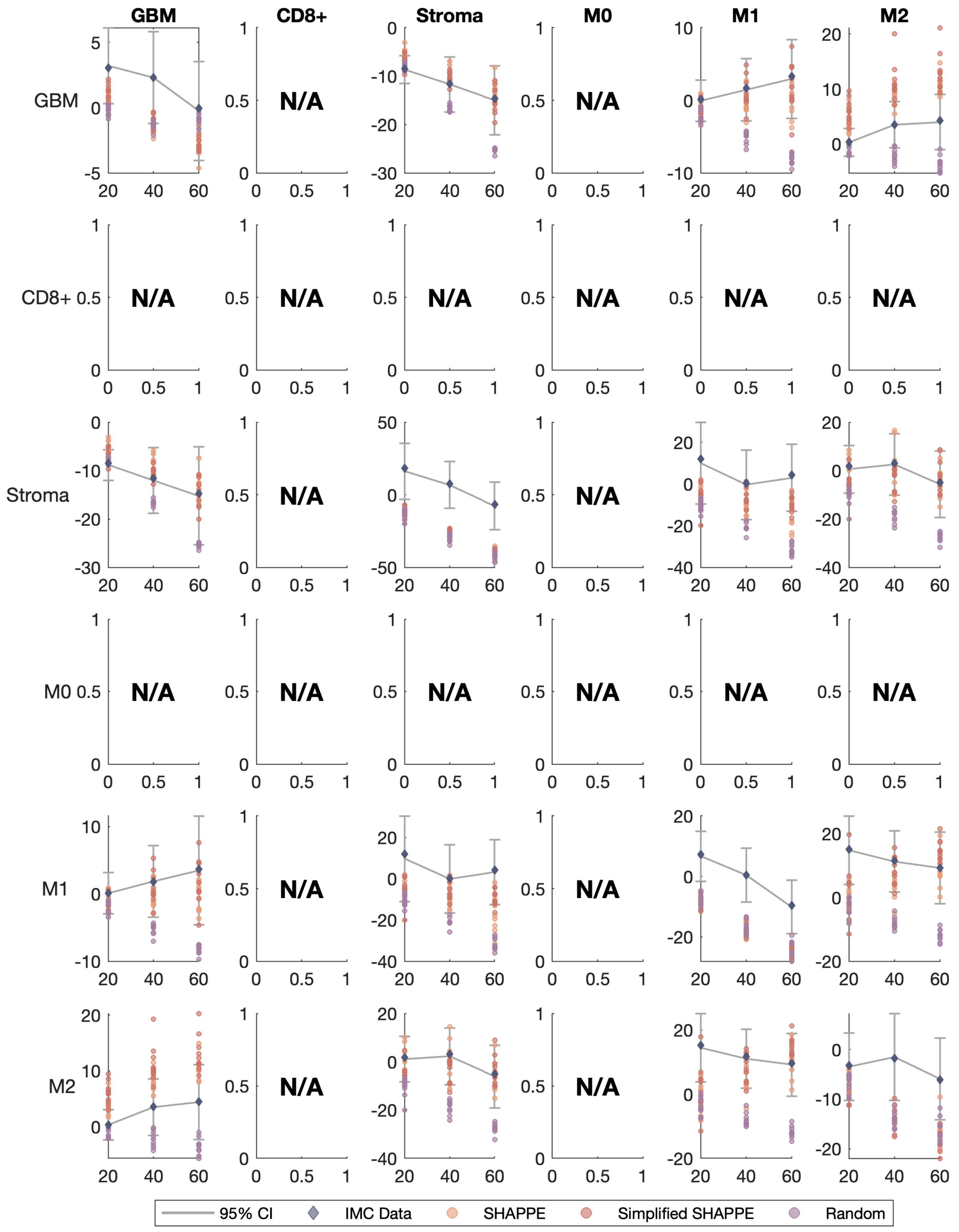

GBM
CD8+
Stroma
M0
M1
M2
GBM
CD8+
Stroma
M0
M1
M2
N/A
95% CI
IMC Data
SHAPPE
Simplified SHAPPE
Random

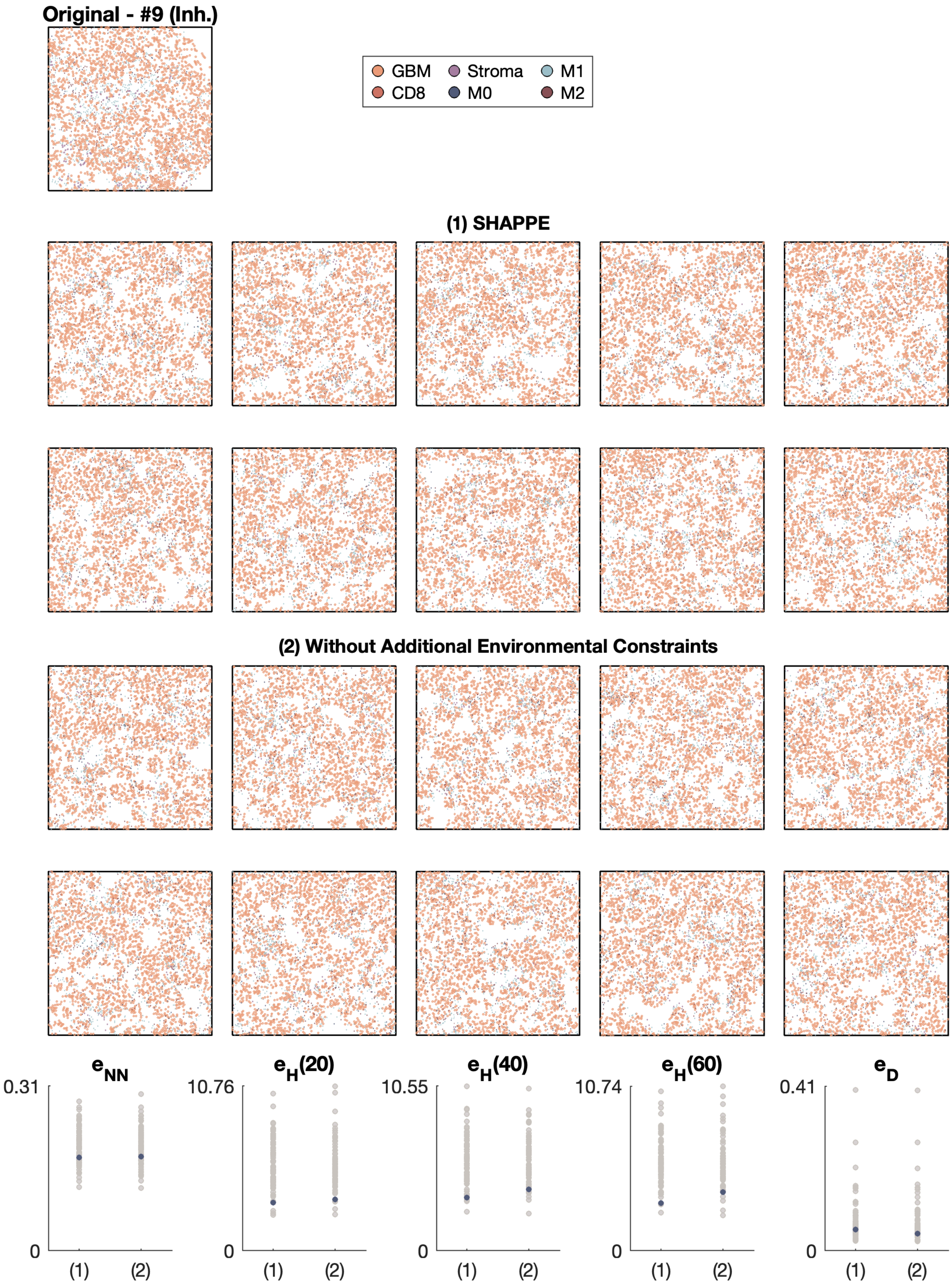
Original - #9 (Inh.)
GBM
Stroma
M1
CD8
M0
M2
(1) SHAPPE
(2) Without Additional Environmental Constraints
$e_{NN}$
0.31
0
(1)
(2)
$e_H(20)$
10.76
0
(1)
(2)
$e_H(40)$
10.55
0
(1)
(2)
$e_H(60)$
10.74
0
(1)
(2)
$e_D$
0.41
0
(1)
(2)

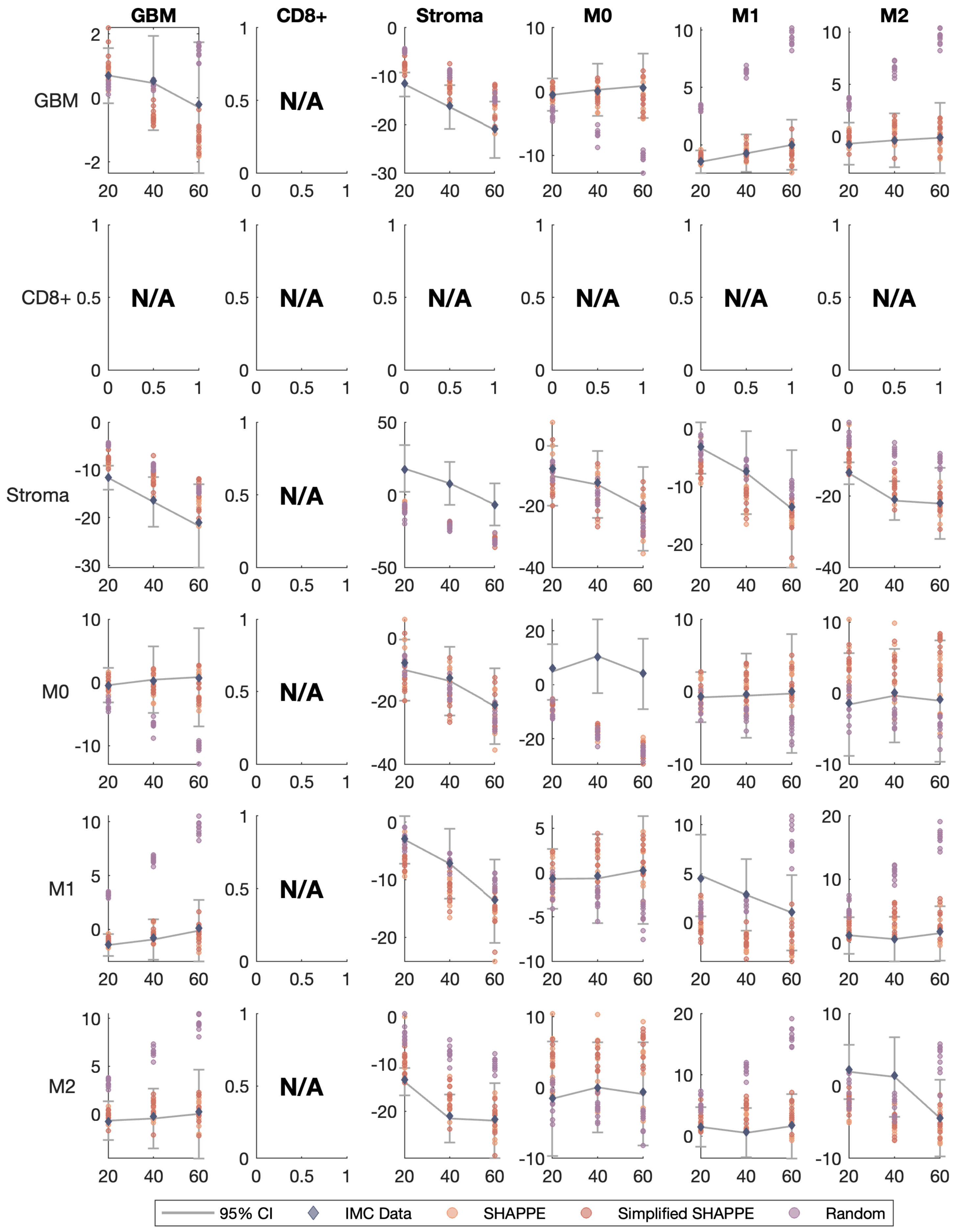

GBM
CD8+
Stroma
M0
M1
M2
N/A
95% CI
IMC Data
SHAPPE
Simplified SHAPPE
Random

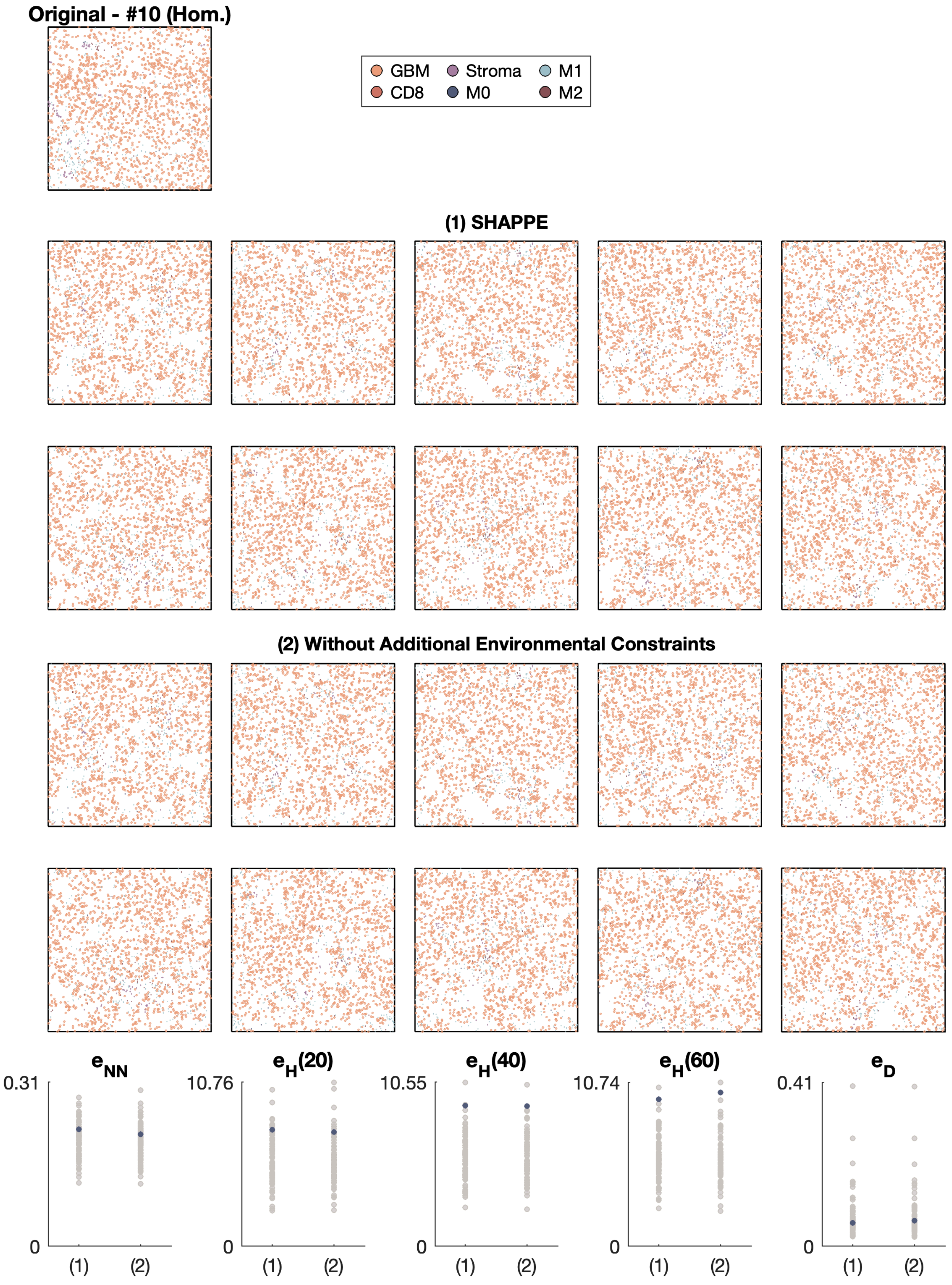

Original - #10 (Hom.)
GBM
Stroma
M1
CD8
M0
M2
(1) SHAPPE
(2) Without Additional Environmental Constraints
$e_{NN}$
0.31
0
(1)
(2)
$e_H(20)$
10.76
0
(1)
(2)
$e_H(40)$
10.55
0
(1)
(2)
$e_H(60)$
10.74
0
(1)
(2)
$e_D$
0.41
0
(1)
(2)

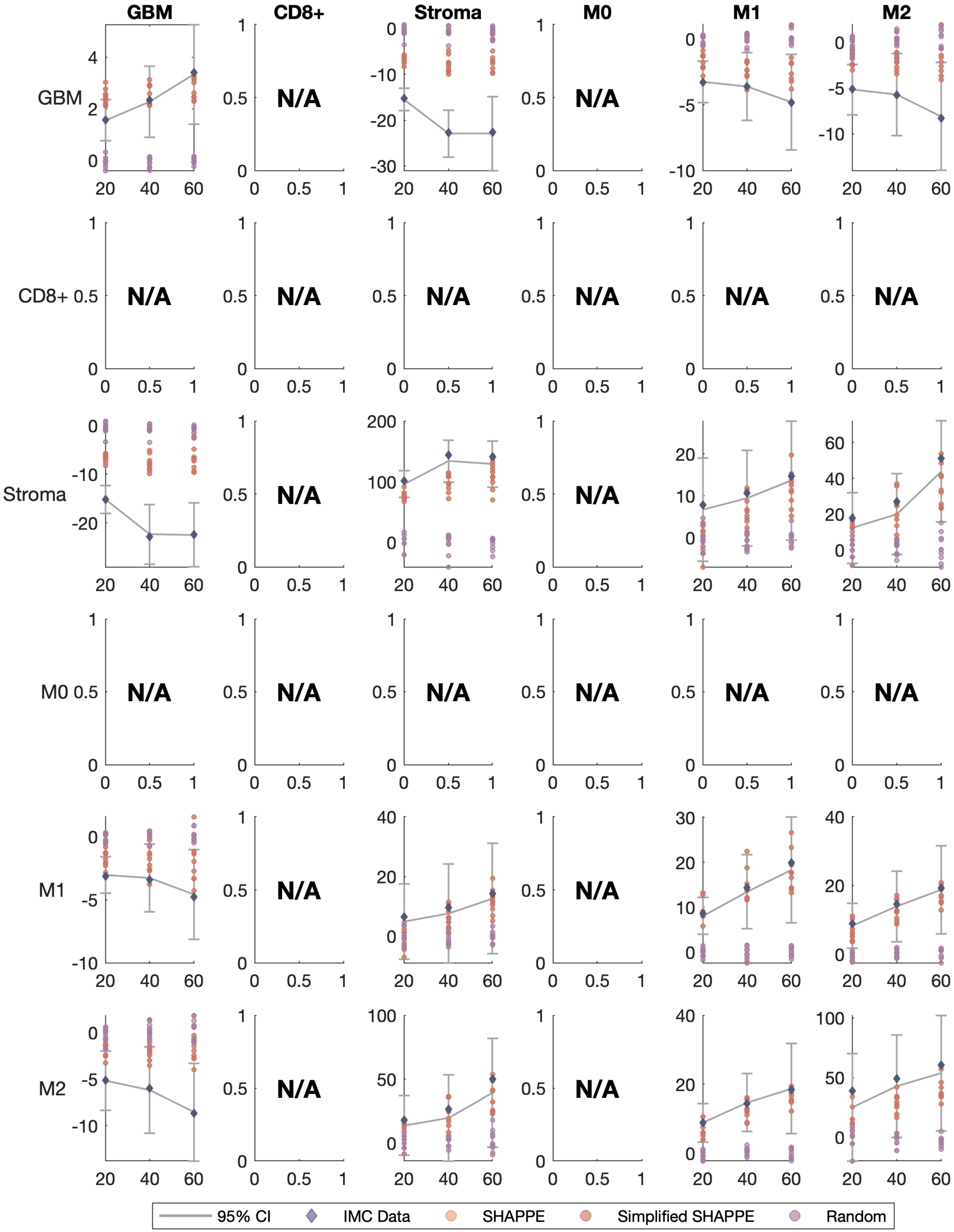
GBM
CD8+
Stroma
M0
M1
M2
N/A
95% CI
IMC Data
SHAPPE
Simplified SHAPPE
Random

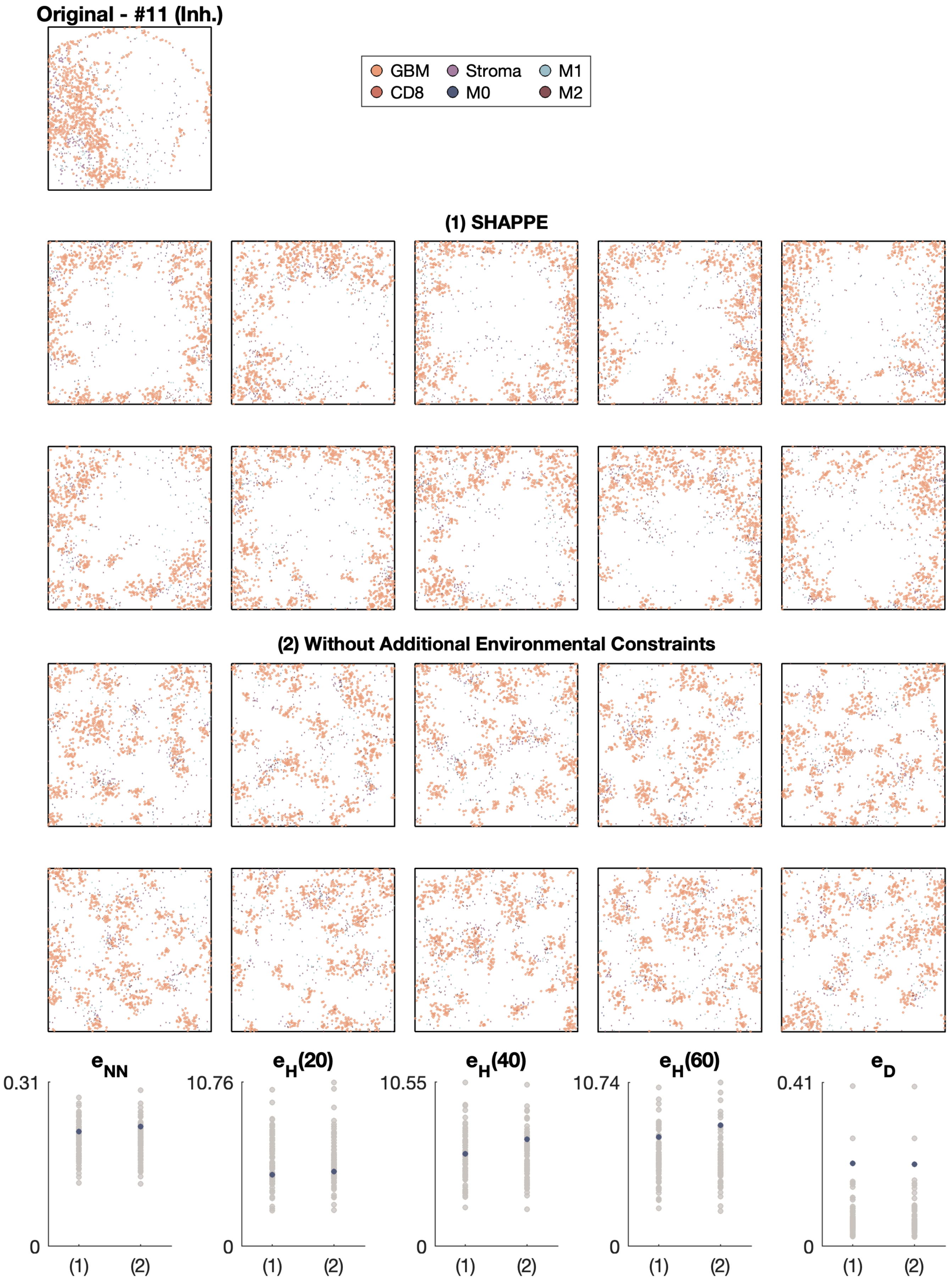
Original - #11 (Inh.)
GBM
CD8
Stroma
M0
M1
M2
(1) SHAPPE
(2) Without Additional Environmental Constraints
$e_{NN}$
$e_H(20)$
$e_H(40)$
$e_H(60)$
$e_D$
0.31
10.76
10.55
10.74
0.41
0
(1)
(2)

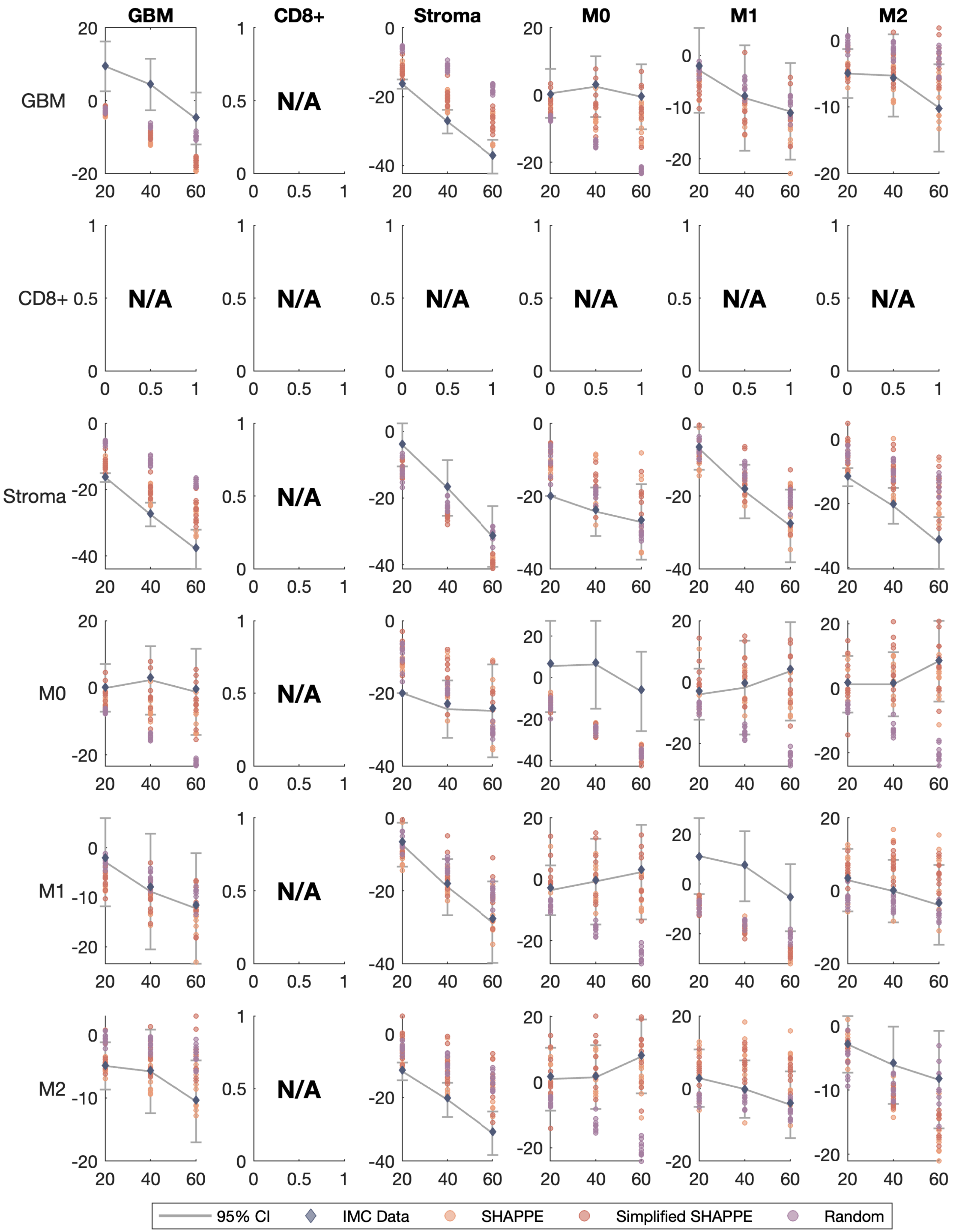

GBM
CD8+
Stroma
M0
M1
M2
N/A
95% CI
IMC Data
SHAPPE
Simplified SHAPPE
Random

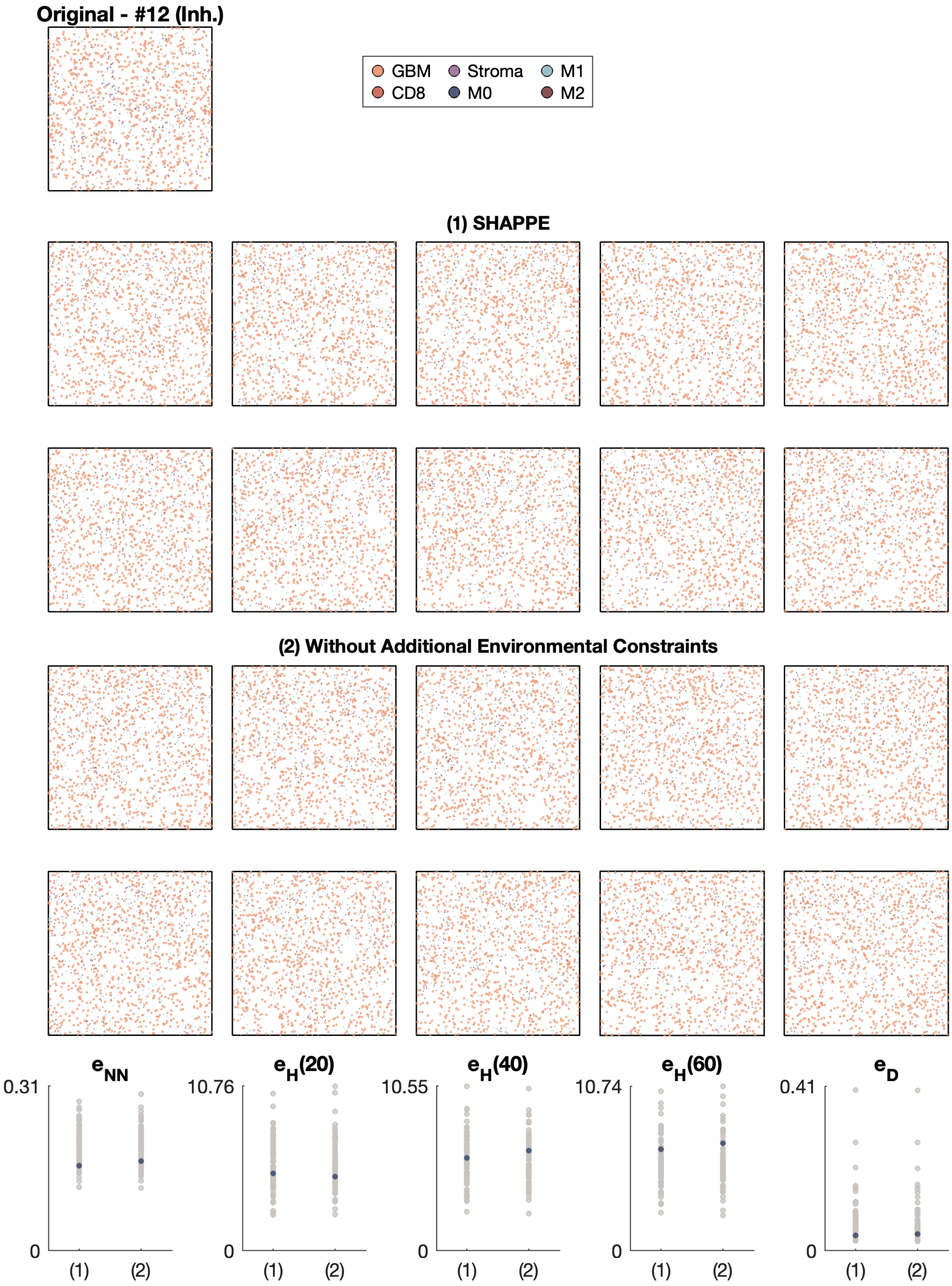
Original - #12 (Inh.)
GBM
Stroma
M1
CD8
M0
M2
(1) SHAPPE
(2) Without Additional Environmental Constraints
e_NN
0.31
0
(1)
(2)
e_H(20)
10.76
0
(1)
(2)
e_H(40)
10.55
0
(1)
(2)
e_H(60)
10.74
0
(1)
(2)
e_D
0.41
0
(1)
(2)

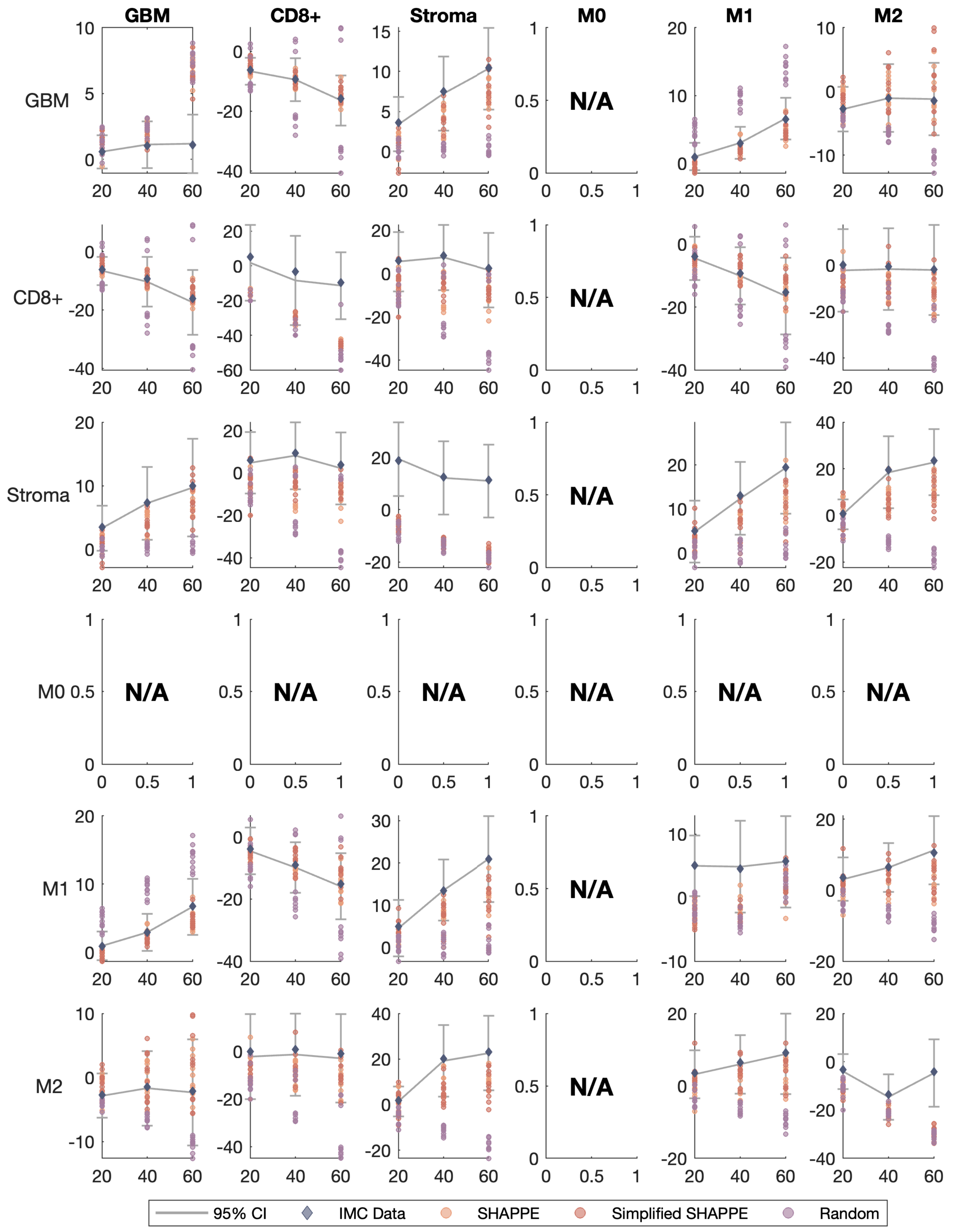

GBM
CD8+
Stroma
M0
M1
M2
N/A
95% CI
IMC Data
SHAPPE
Simplified SHAPPE
Random

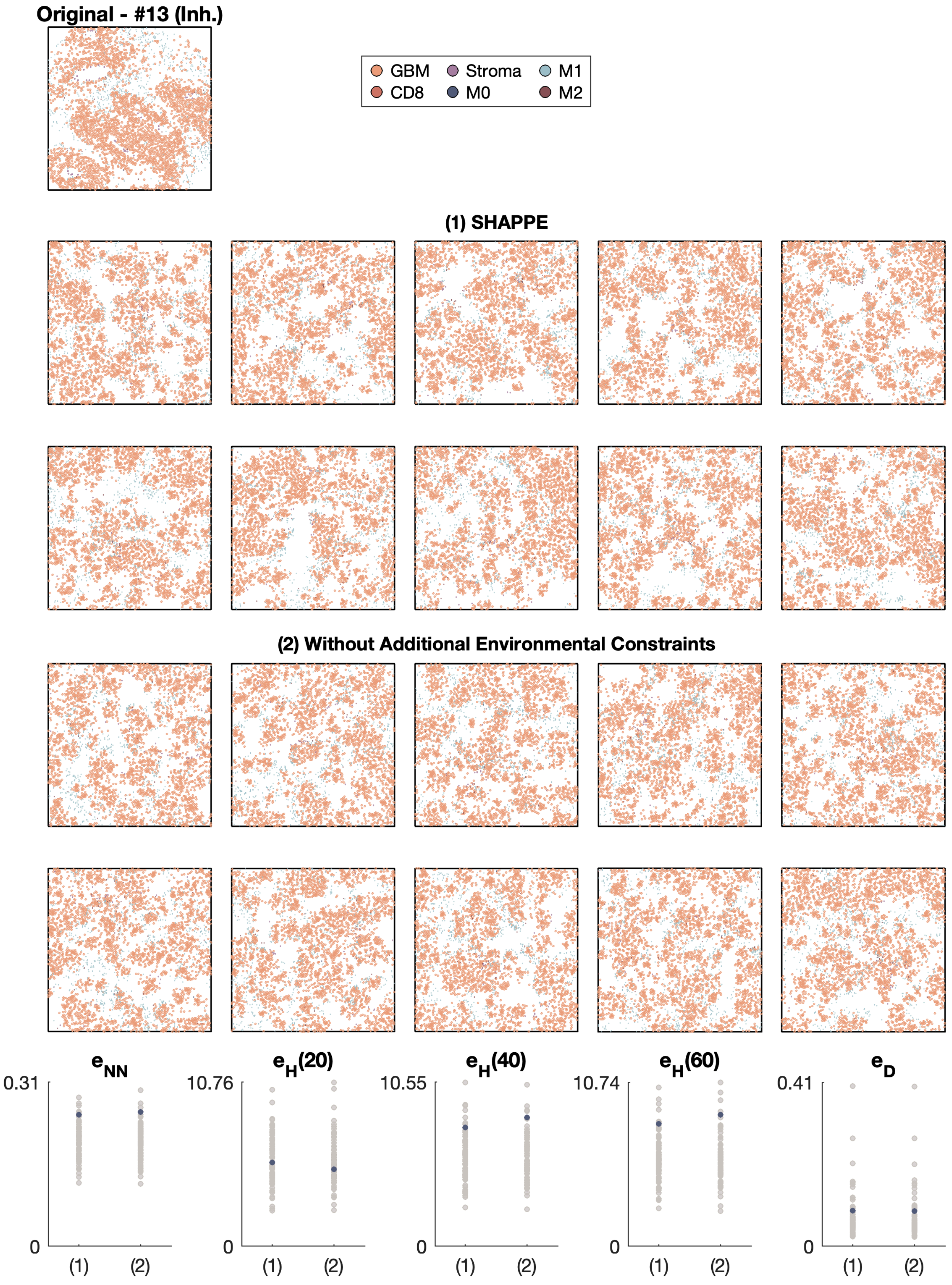

Original - #13 (Inh.)
GBM
CD8
Stroma
M0
M1
M2
(1) SHAPPE
(2) Without Additional Environmental Constraints
$e_{NN}$
$e_H(20)$
$e_H(40)$
$e_H(60)$
$e_D$
0.31
10.76
10.55
10.74
0.41
0
(1)
(2)

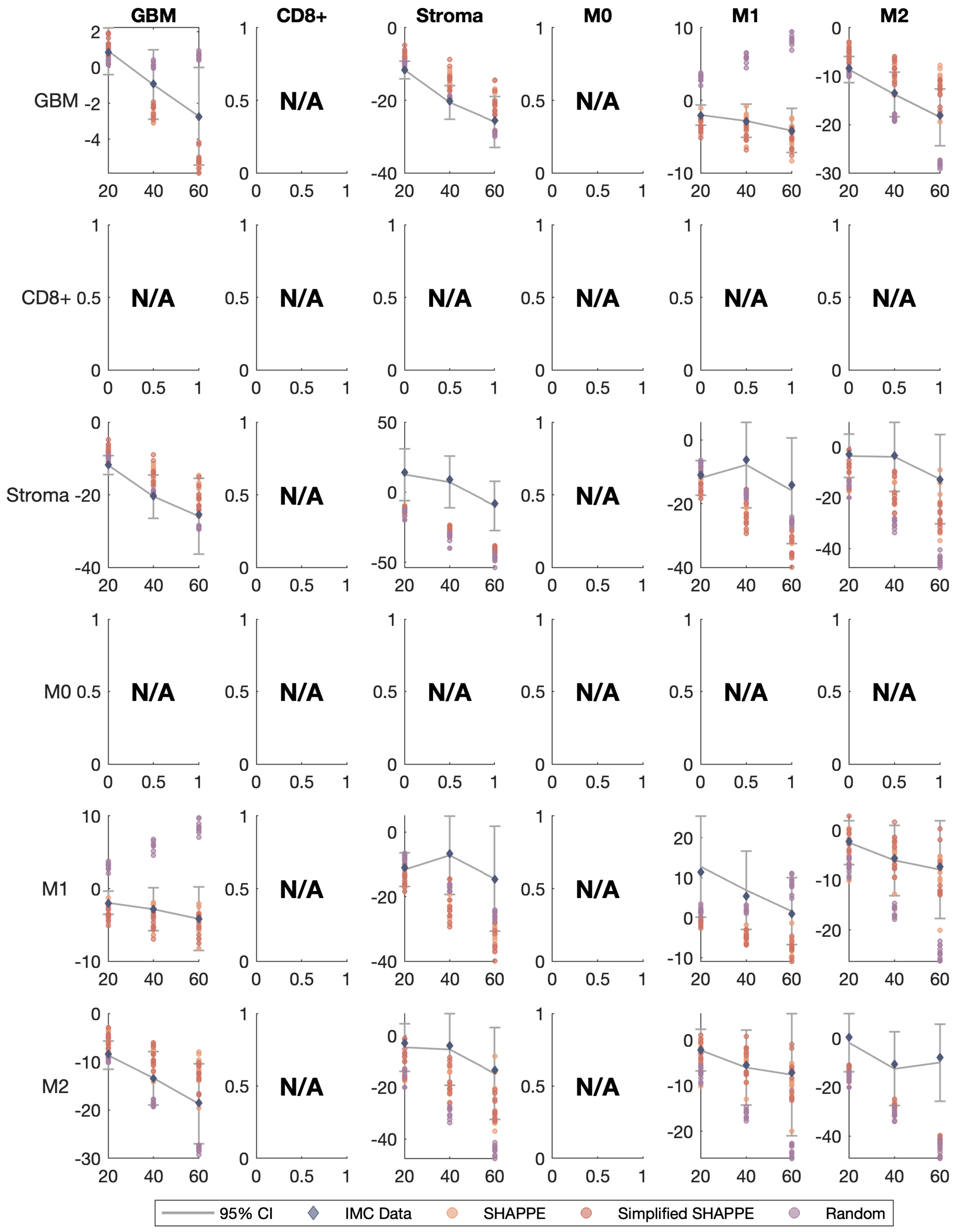

GBM
CD8+
Stroma
M0
M1
M2
N/A
95% CI
IMC Data
SHAPPE
Simplified SHAPPE
Random

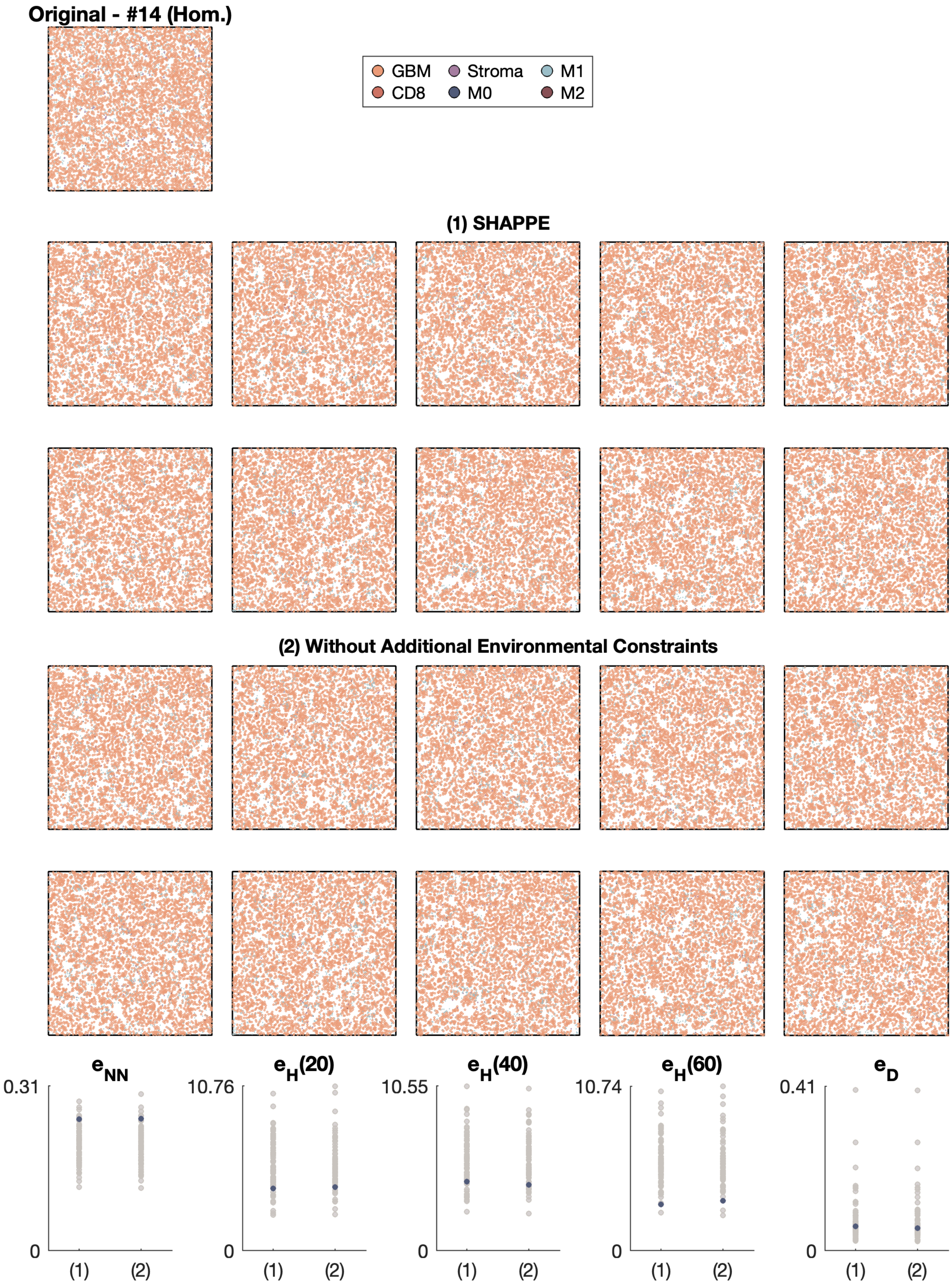

Original - #14 (Hom.)
GBM
Stroma
M1
CD8
M0
M2
(1) SHAPPE
(2) Without Additional Environmental Constraints
e_NN
e_H(20)
e_H(40)
e_H(60)
e_D
0.31
10.76
10.55
10.74
0.41
0
(1)
(2)

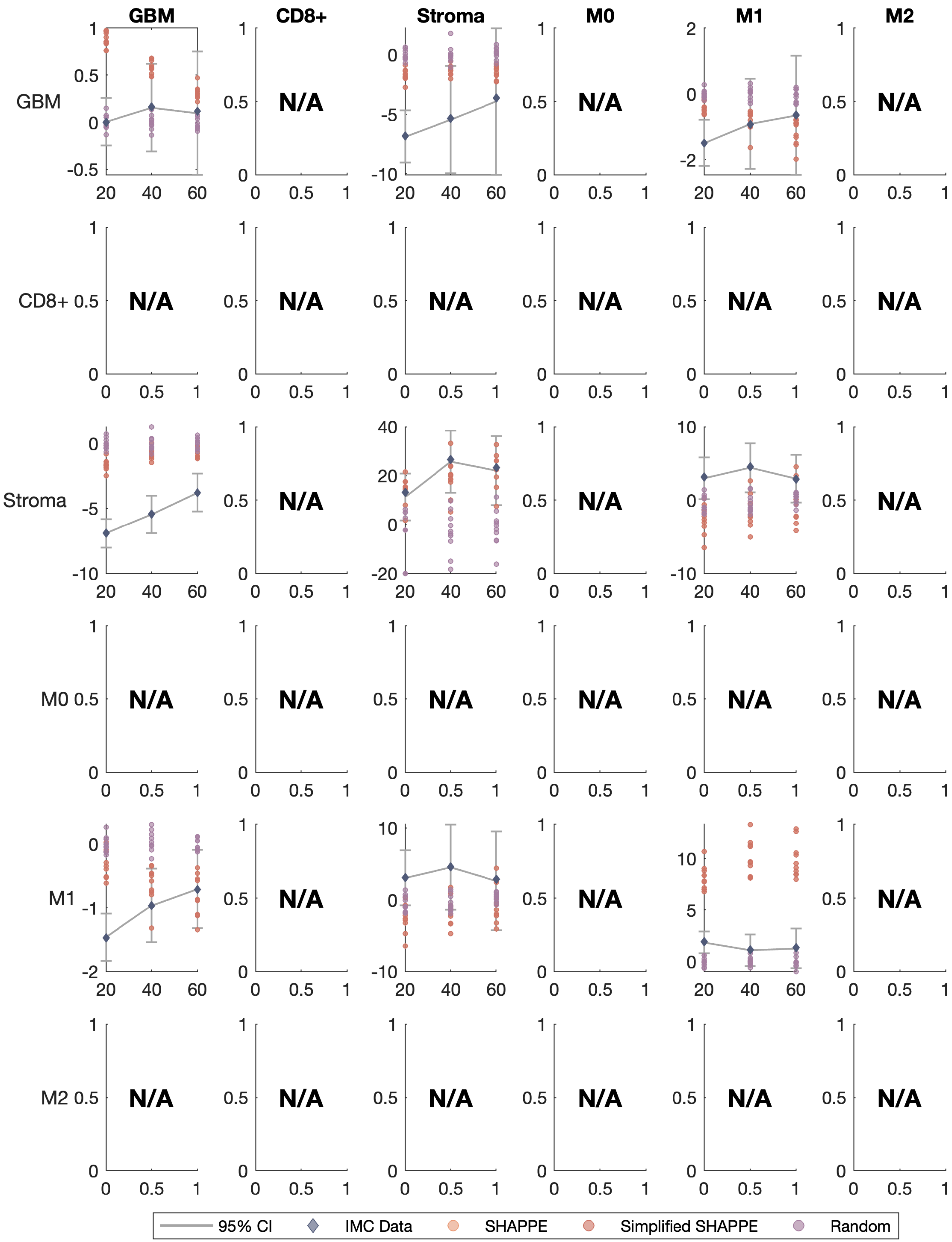

GBM
CD8+
Stroma
M0
M1
M2
N/A
95% CI
IMC Data
SHAPPE
Simplified SHAPPE
Random

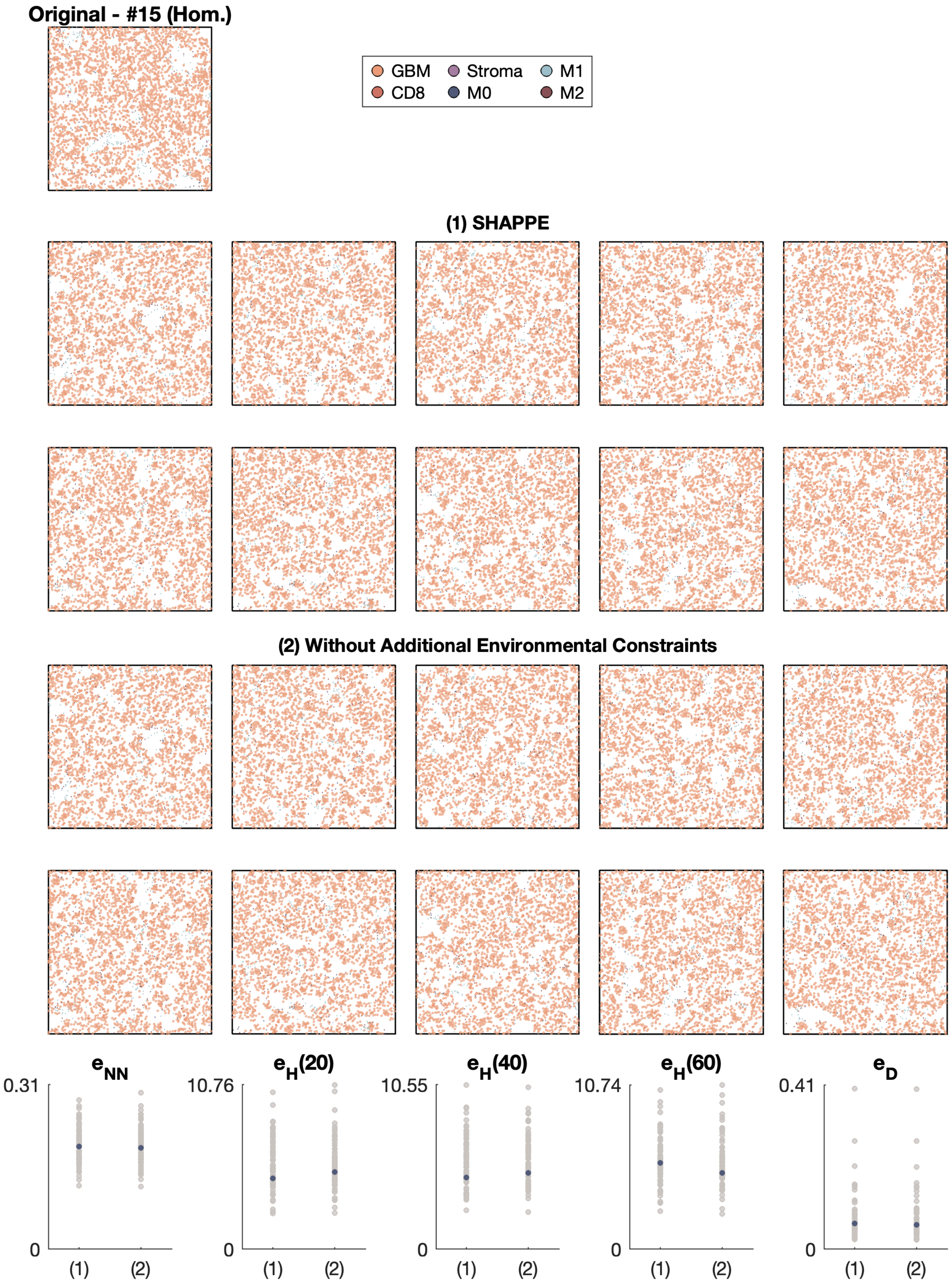

Original - #15 (Hom.)
GBM
Stroma
M1
CD8
M0
M2
(1) SHAPPE
(2) Without Additional Environmental Constraints
e_NN
0.31
0
(1)
(2)
e_H(20)
10.76
0
(1)
(2)
e_H(40)
10.55
0
(1)
(2)
e_H(60)
10.74
0
(1)
(2)
e_D
0.41
0
(1)
(2)

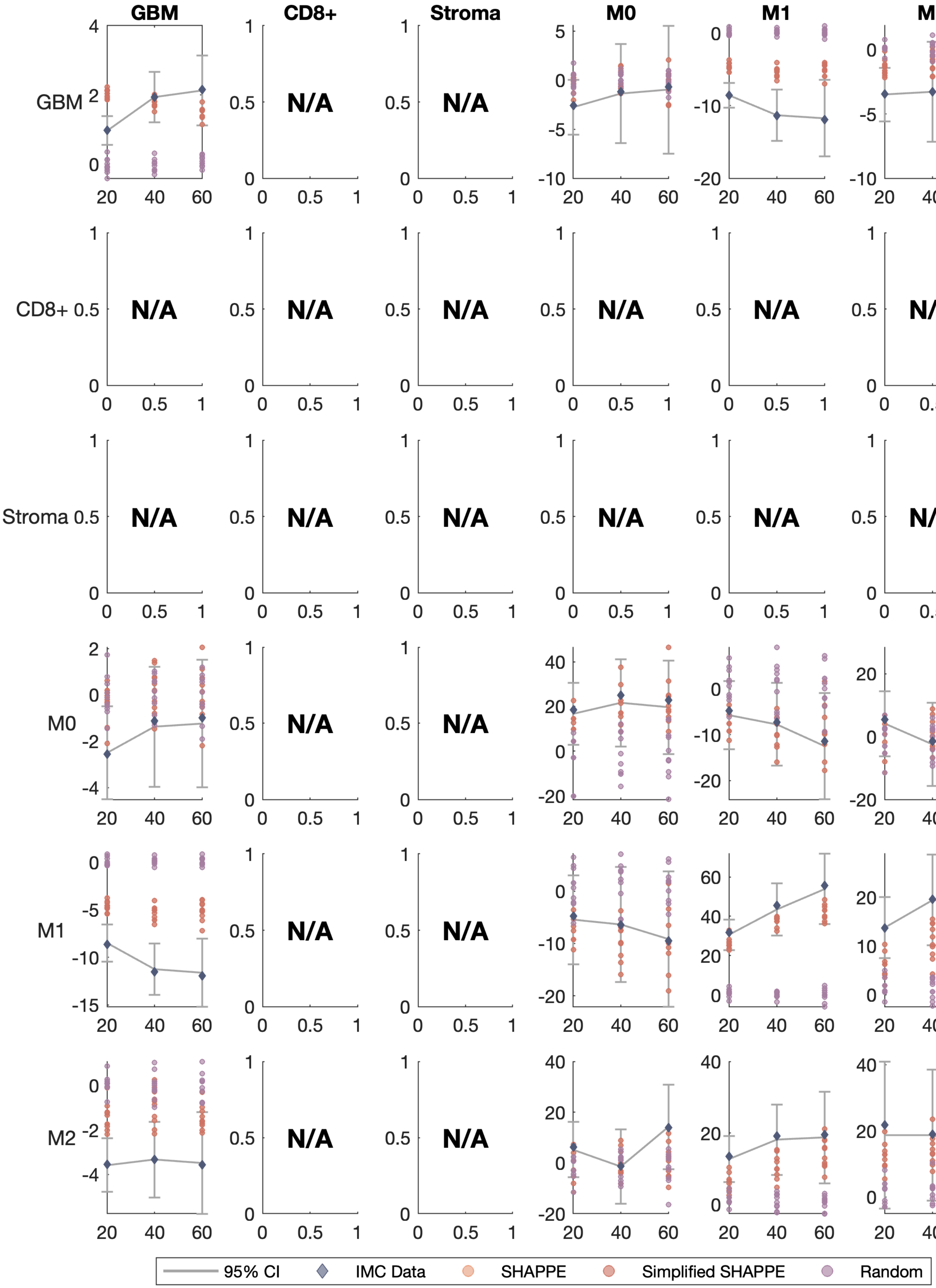
GBM
CD8+
Stroma
M0
M1
M2
GBM
CD8+
Stroma
M0
M1
M2
N/A
95% CI
IMC Data
SHAPPE
Simplified SHAPPE
Random

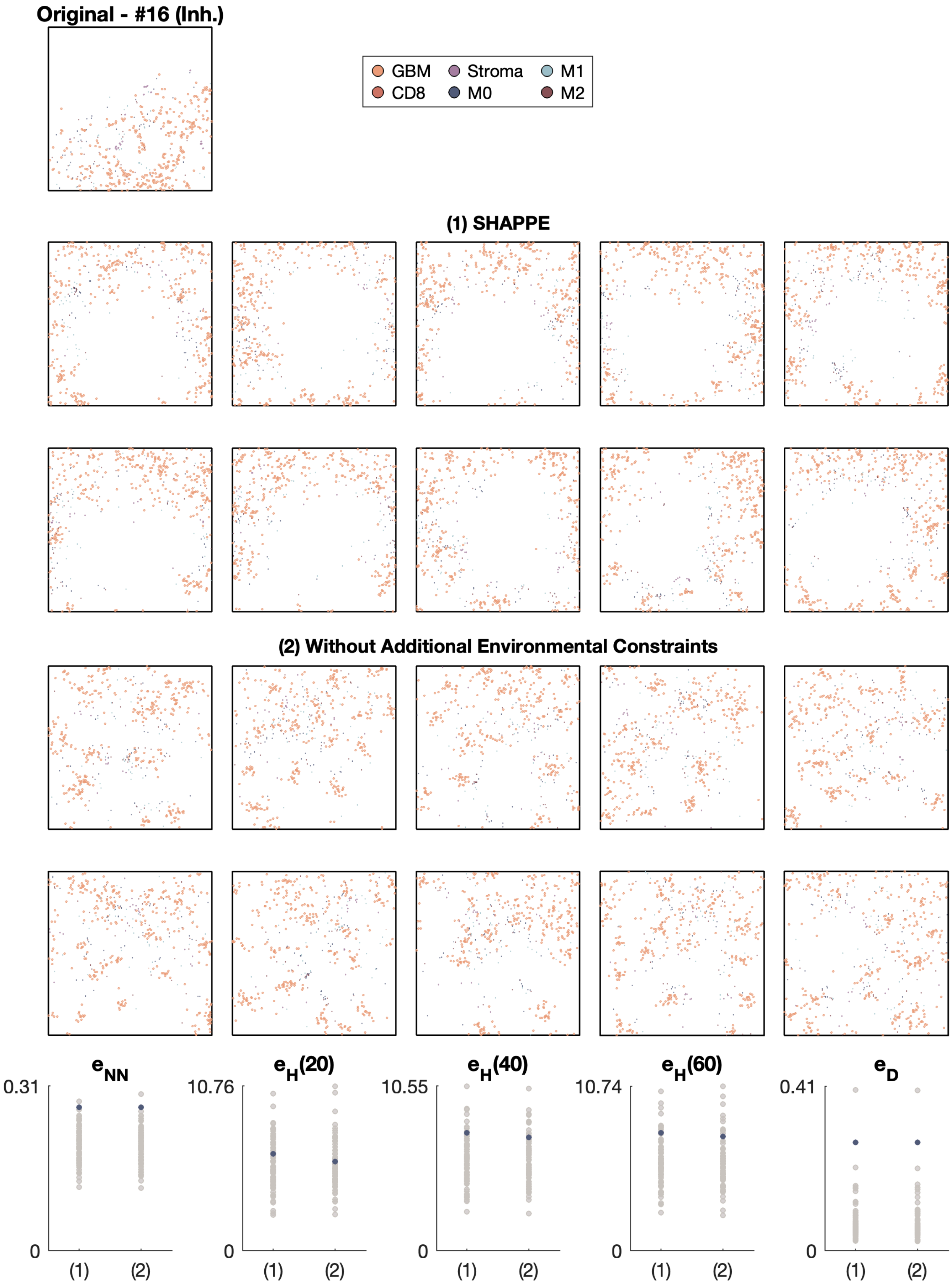

Original - #16 (Inh.)
GBM
Stroma
M1
CD8
M0
M2
(1) SHAPPE
(2) Without Additional Environmental Constraints
$e_{NN}$
$e_H(20)$
$e_H(40)$
$e_H(60)$
$e_D$
0.31
10.76
10.55
10.74
0.41
0
(1)
(2)

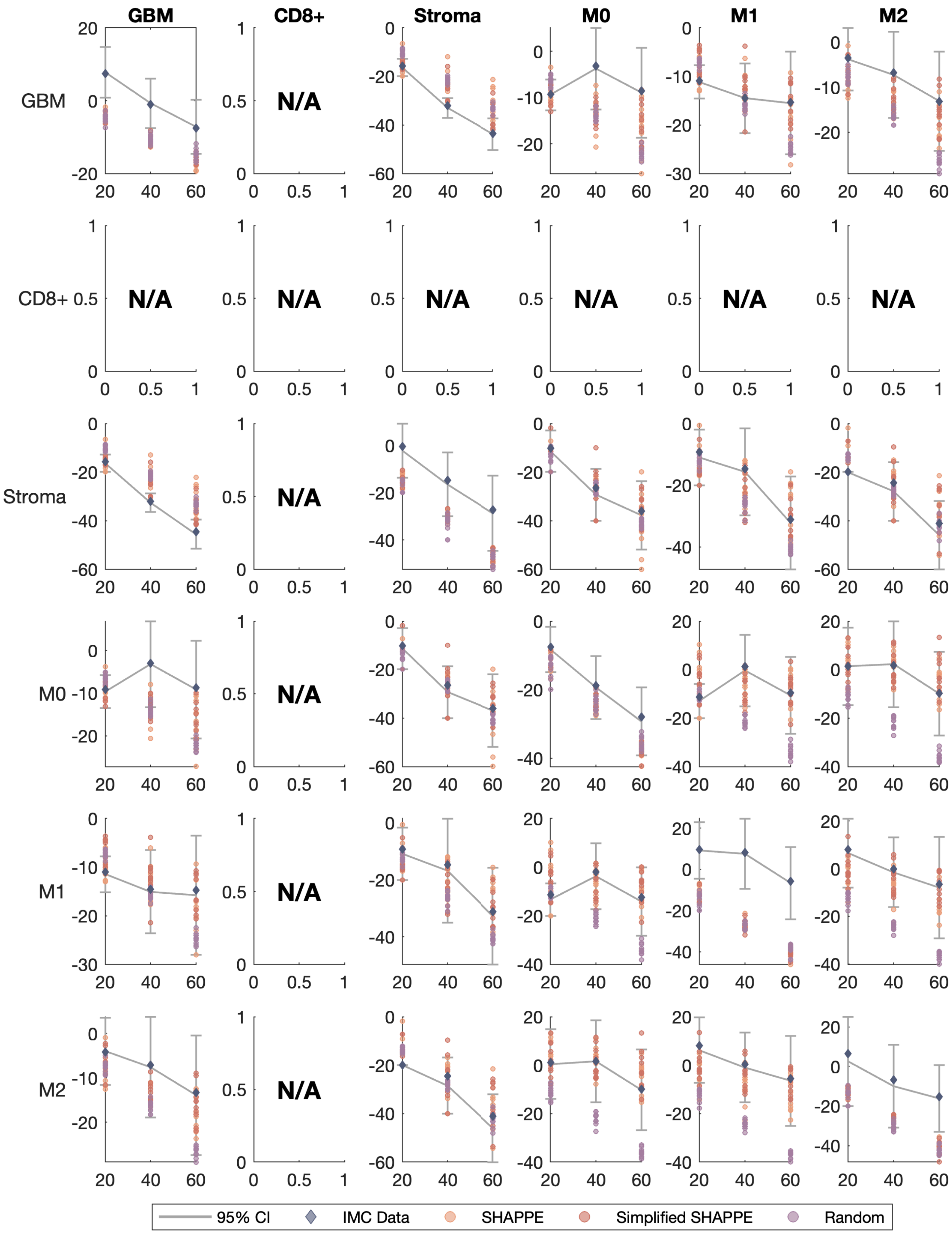

GBM
CD8+
Stroma
M0
M1
M2
N/A
95% CI
IMC Data
SHAPPE
Simplified SHAPPE
Random

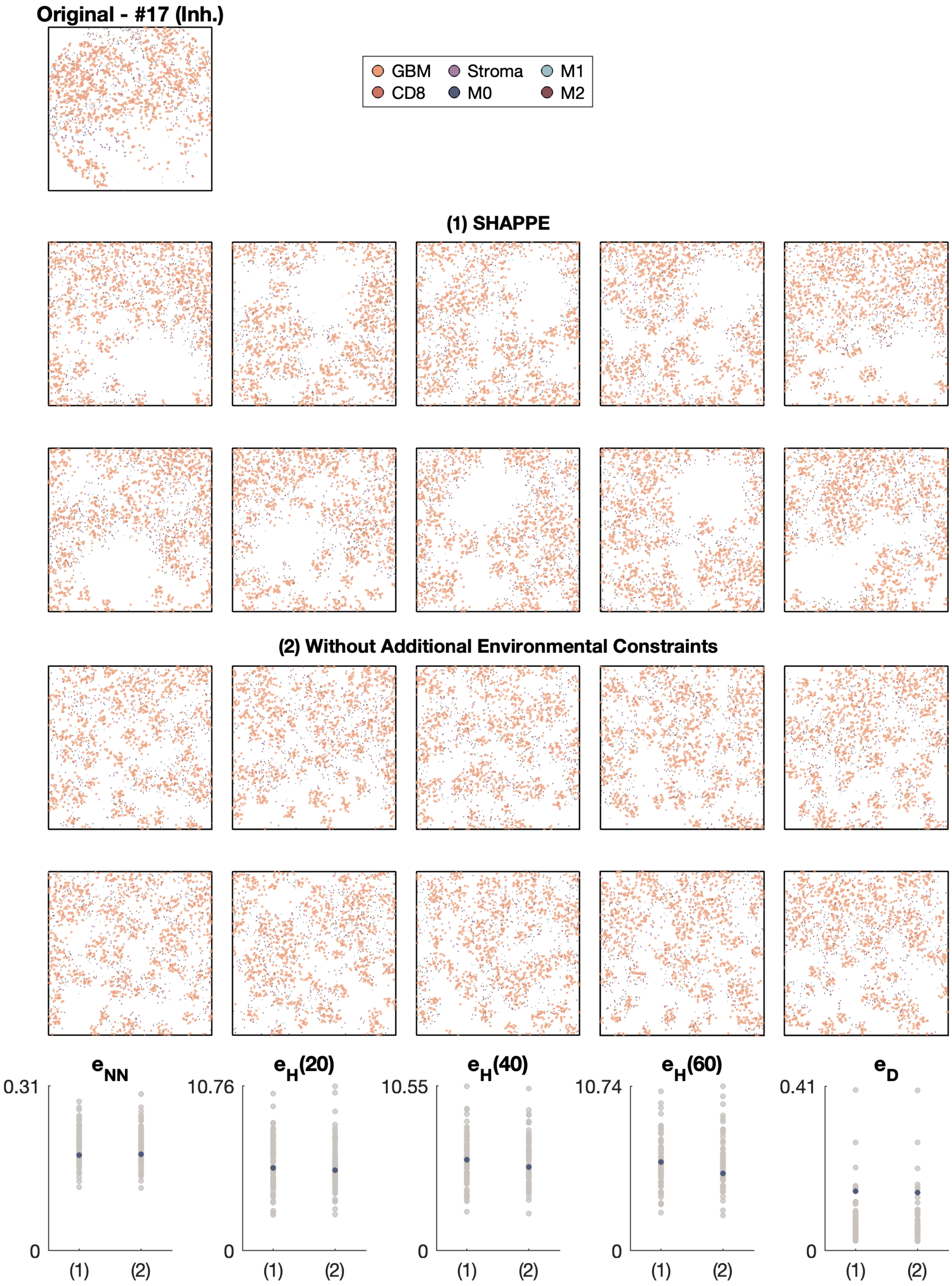

Original - #17 (Inh.)
GBM
Stroma
M1
CD8
M0
M2
(1) SHAPPE
(2) Without Additional Environmental Constraints
e_NN
e_H(20)
e_H(40)
e_H(60)
e_D
0.31
10.76
10.55
10.74
0.41
0
(1)
(2)

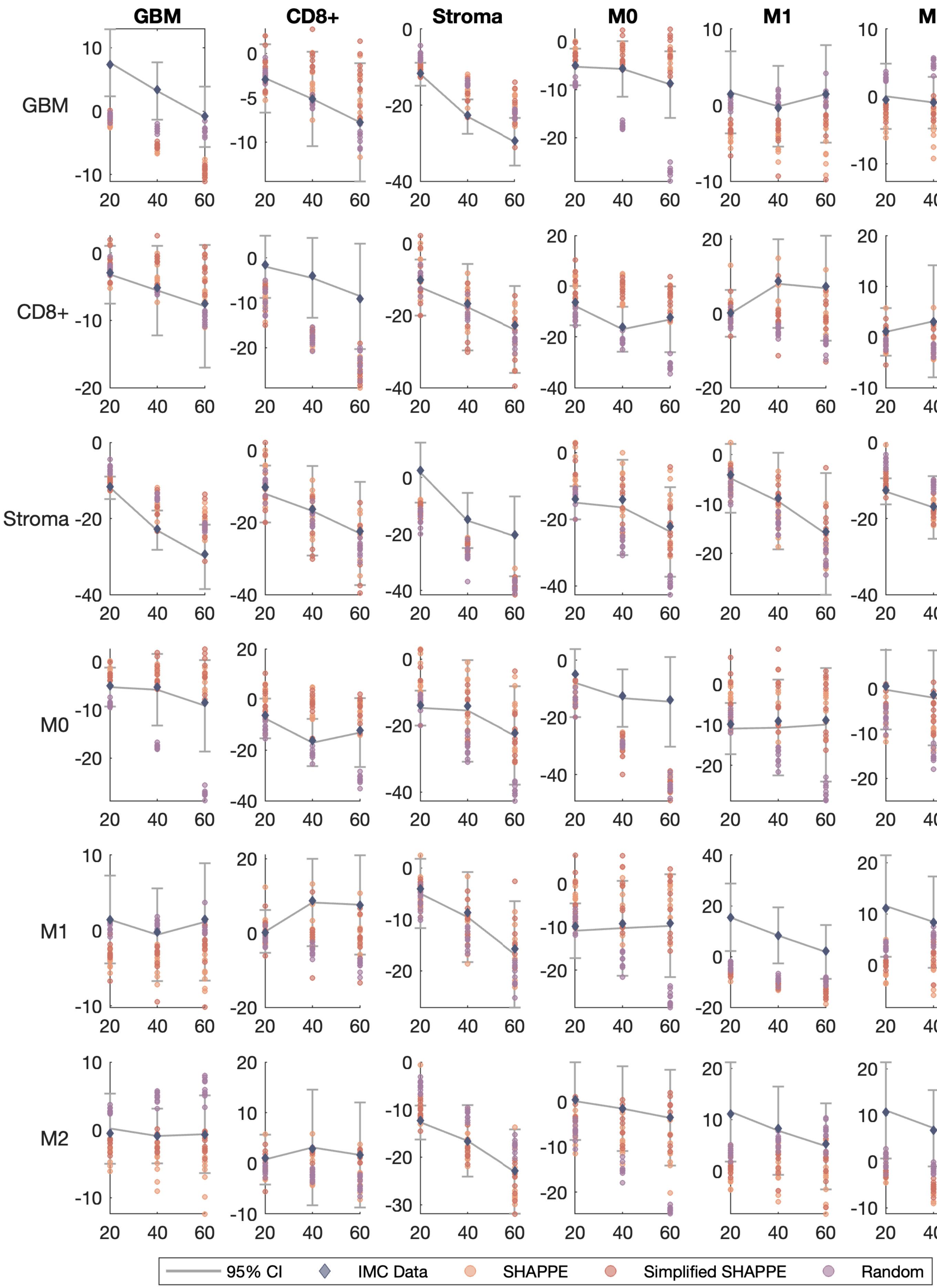

GBM
CD8+
Stroma
M0
M1
M2
95% CI
IMC Data
SHAPPE
Simplified SHAPPE
Random

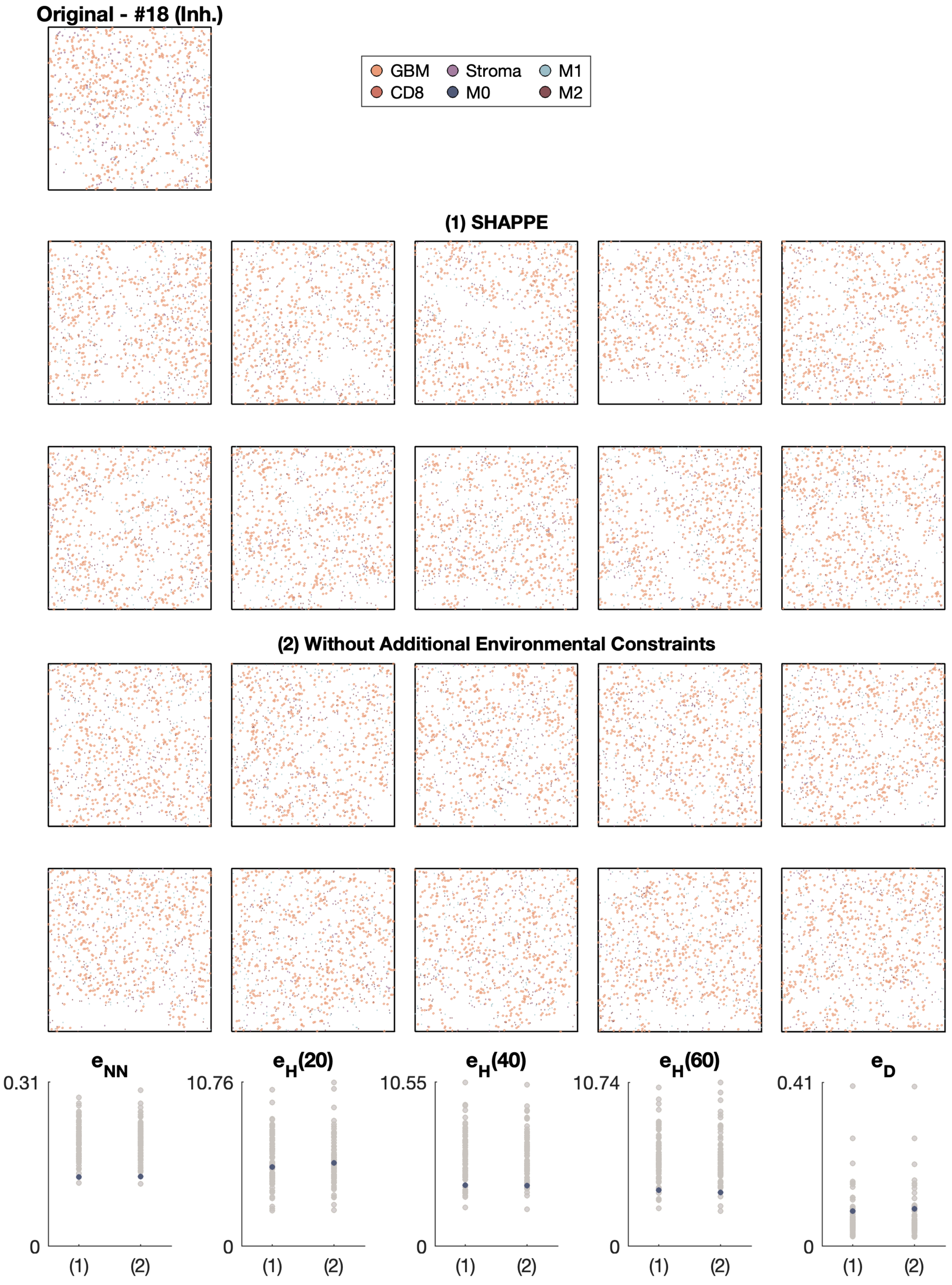

Original - #18 (Inh.)
GBM
CD8
Stroma
M0
M1
M2
(1) SHAPPE
(2) Without Additional Environmental Constraints
e_NN
0.31
0
e_H(20)
10.76
e_H(40)
10.55
e_H(60)
10.74
e_D
0.41
(1)
(2)

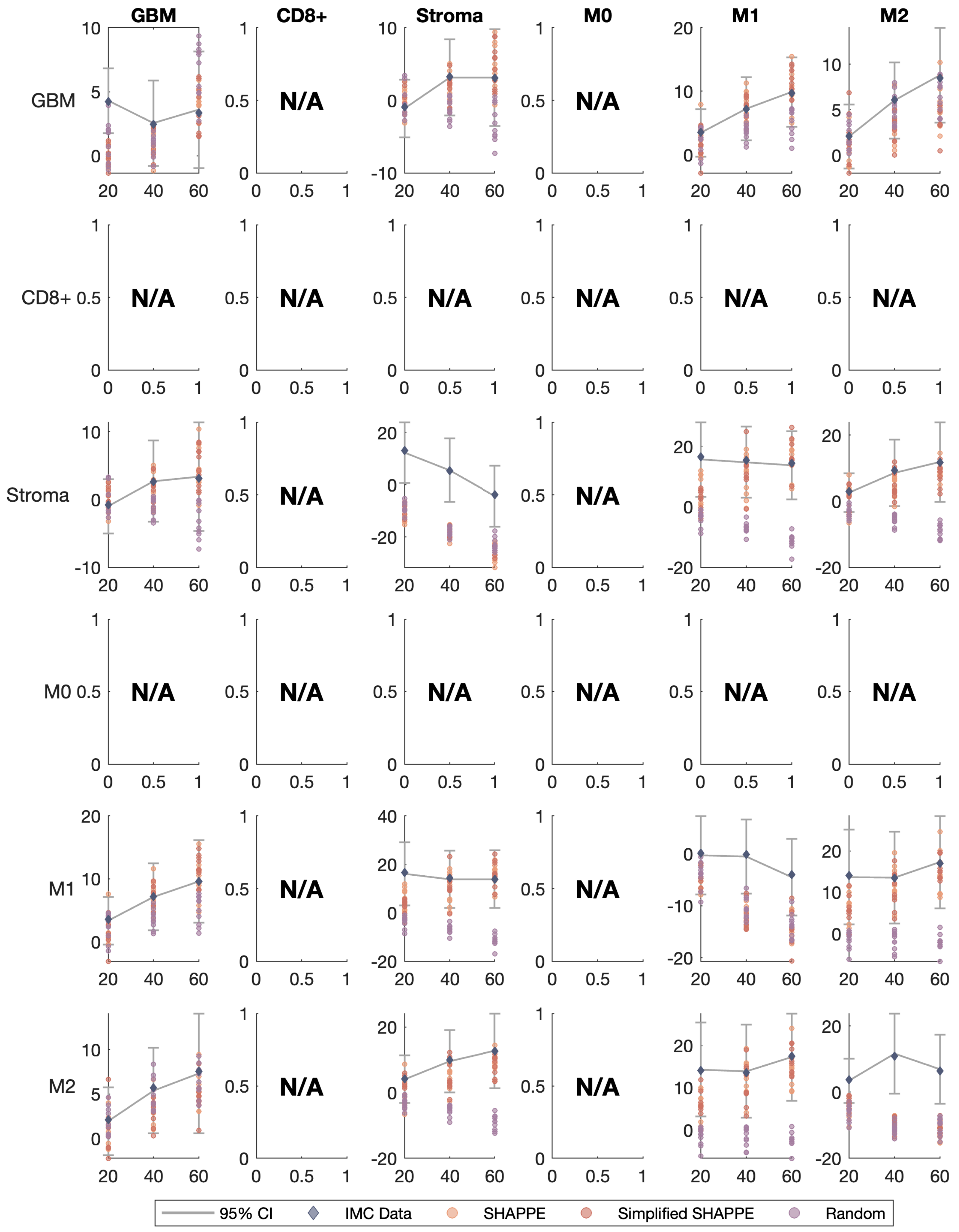

GBM
CD8+
Stroma
M0
M1
M2
N/A
95% CI
IMC Data
SHAPPE
Simplified SHAPPE
Random

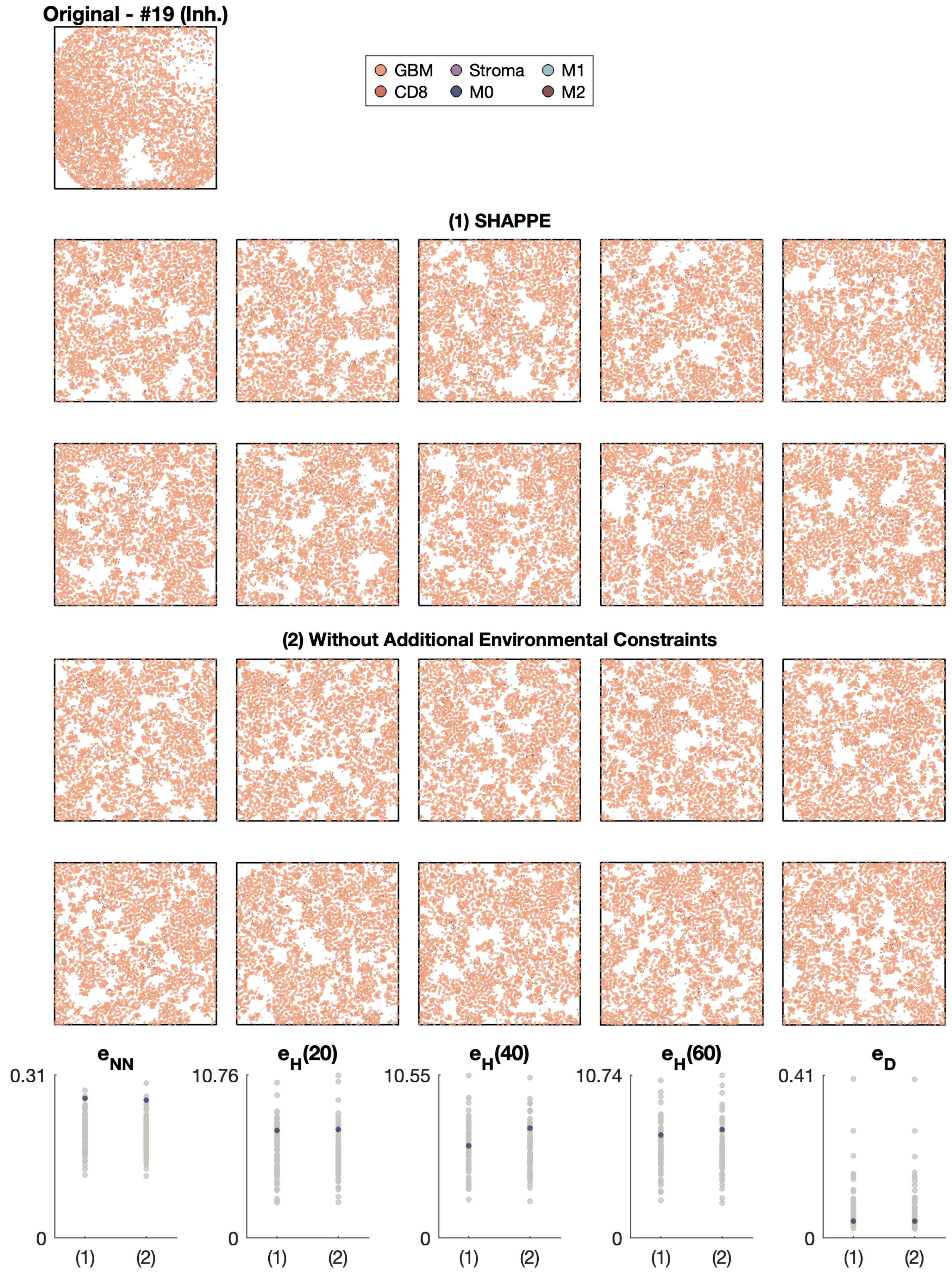
Original - #19 (Inh.)
GBM
CD8
Stroma
M0
M1
M2
(1) SHAPPE
(2) Without Additional Environmental Constraints
$e_{NN}$
$e_H(20)$
$e_H(40)$
$e_H(60)$
$e_D$
0.31
10.76
10.55
10.74
0.41
0
(1)
(2)

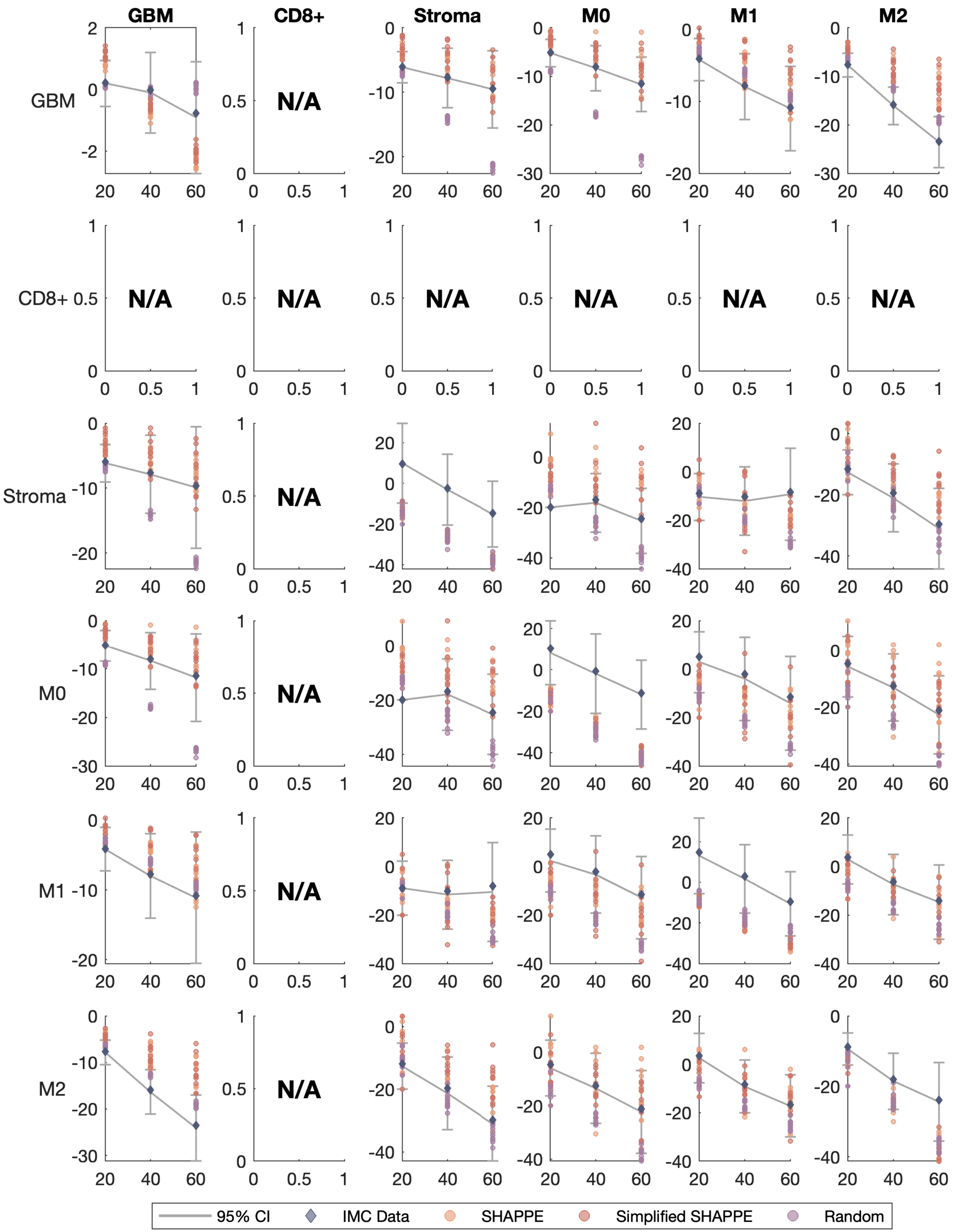

GBM
CD8+
Stroma
M0
M1
M2
N/A
95% CI
IMC Data
SHAPPE
Simplified SHAPPE
Random

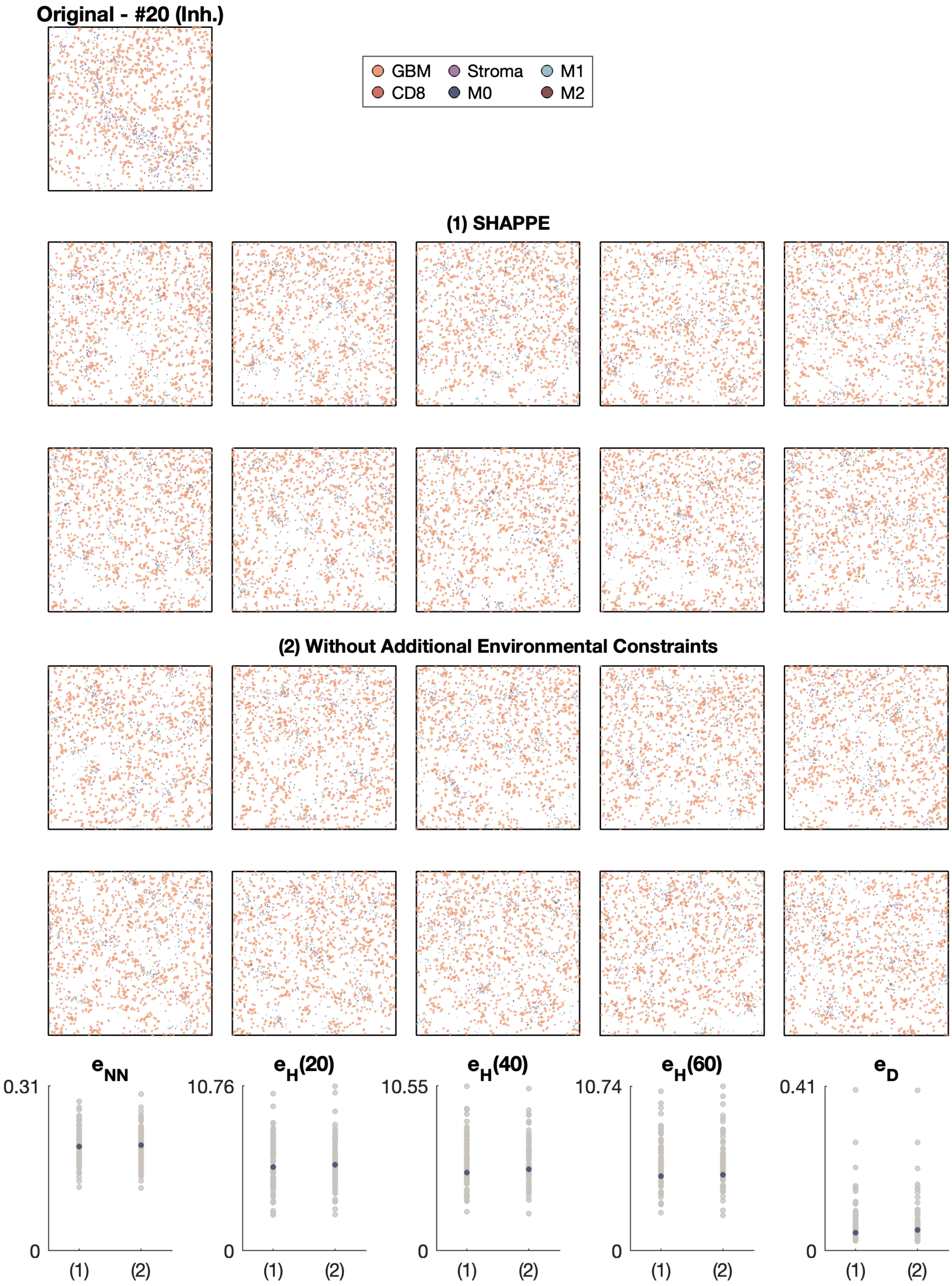

Original - #20 (Inh.)
GBM
Stroma
M1
CD8
M0
M2
(1) SHAPPE
(2) Without Additional Environmental Constraints
$e_{NN}$
0.31
0
(1)
(2)
$e_H(20)$
10.76
0
(1)
(2)
$e_H(40)$
10.55
0
(1)
(2)
$e_H(60)$
10.74
0
(1)
(2)
$e_D$
0.41
0
(1)
(2)

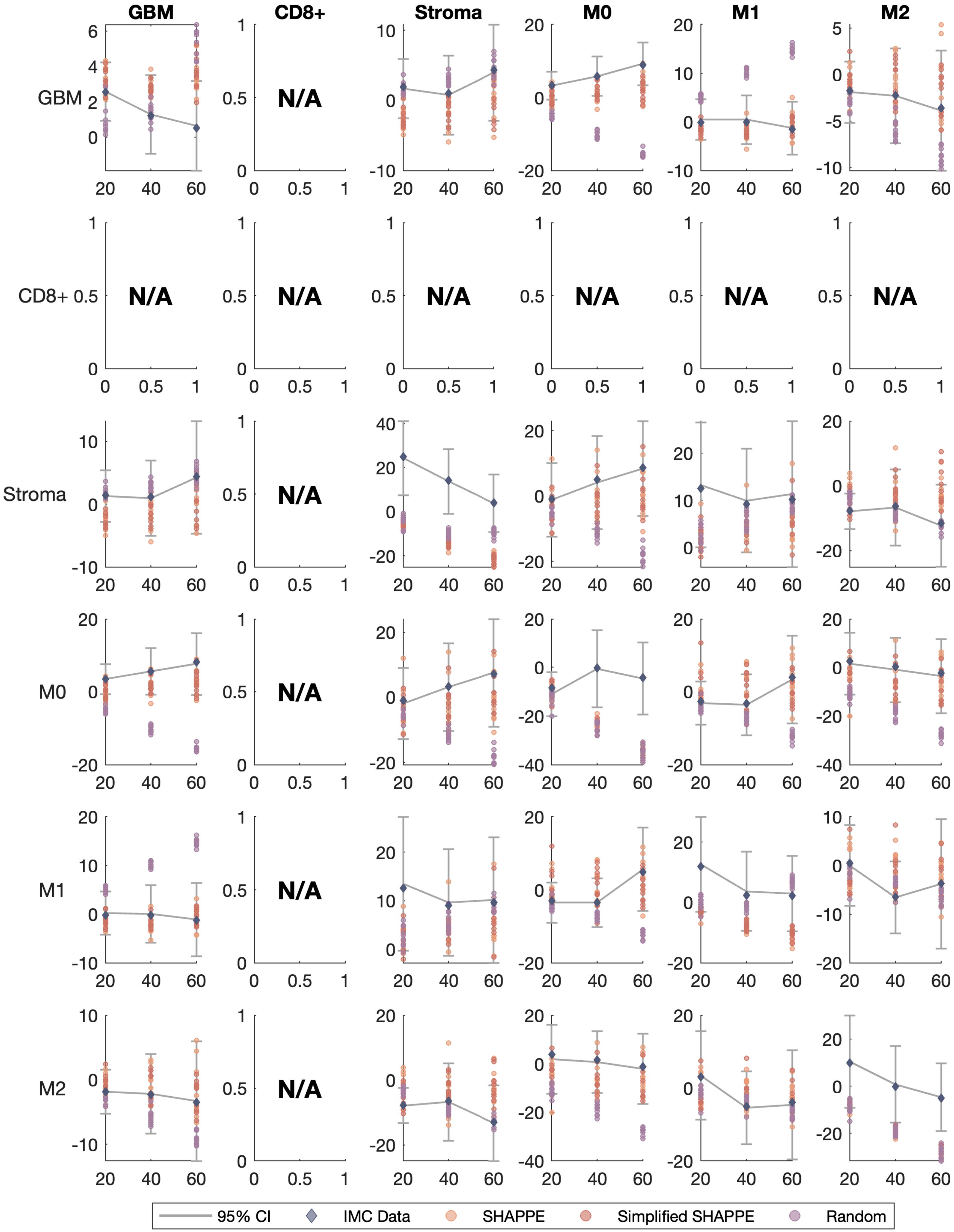
GBM
CD8+
Stroma
M0
M1
M2
N/A
95% CI
IMC Data
SHAPPE
Simplified SHAPPE
Random

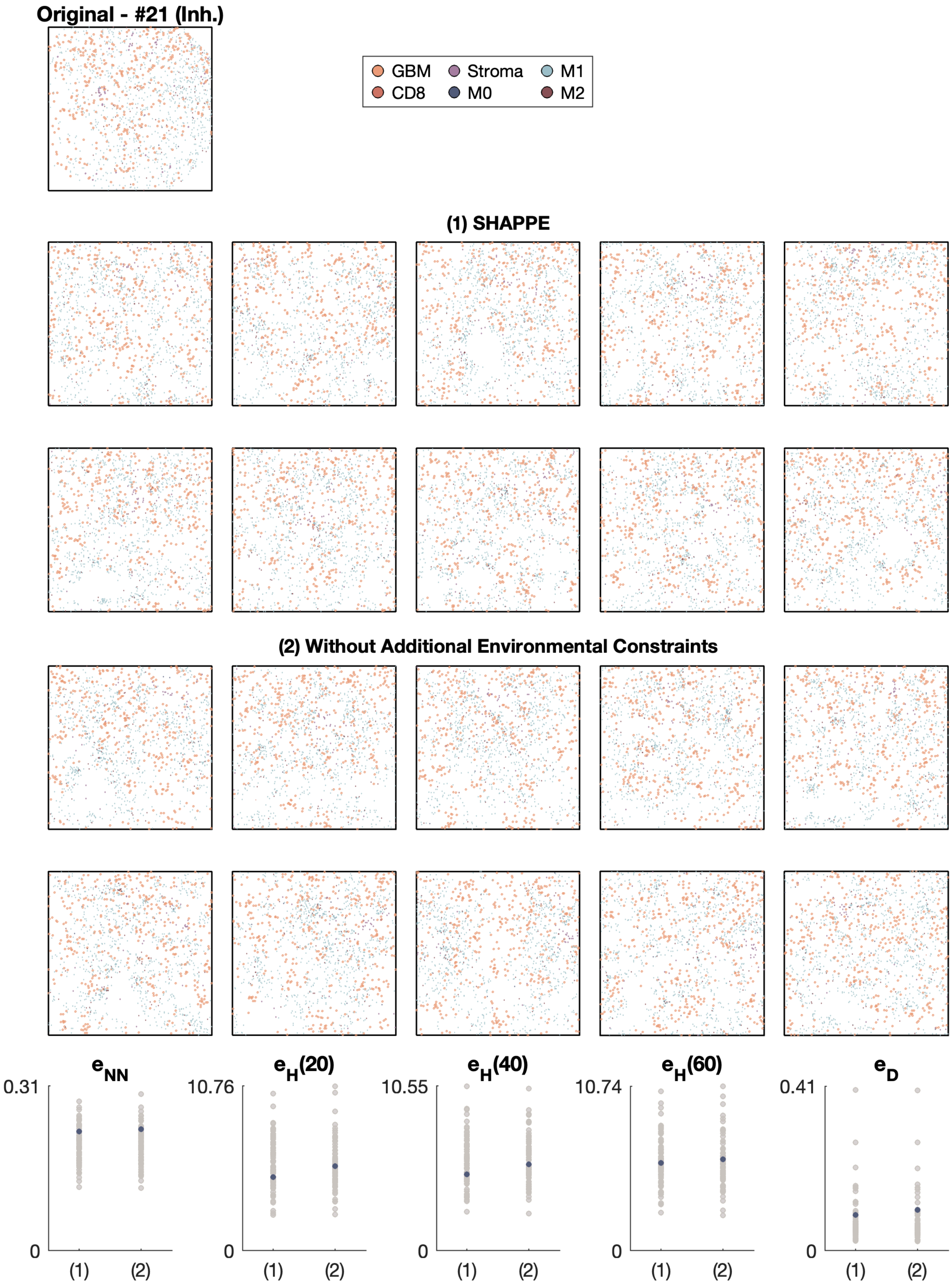

Original - #21 (Inh.)
GBM
Stroma
M1
CD8
M0
M2
(1) SHAPPE
(2) Without Additional Environmental Constraints
e_NN
0.31
0
(1)
(2)
e_H(20)
10.76
0
(1)
(2)
e_H(40)
10.55
0
(1)
(2)
e_H(60)
10.74
0
(1)
(2)
e_D
0.41
0
(1)
(2)

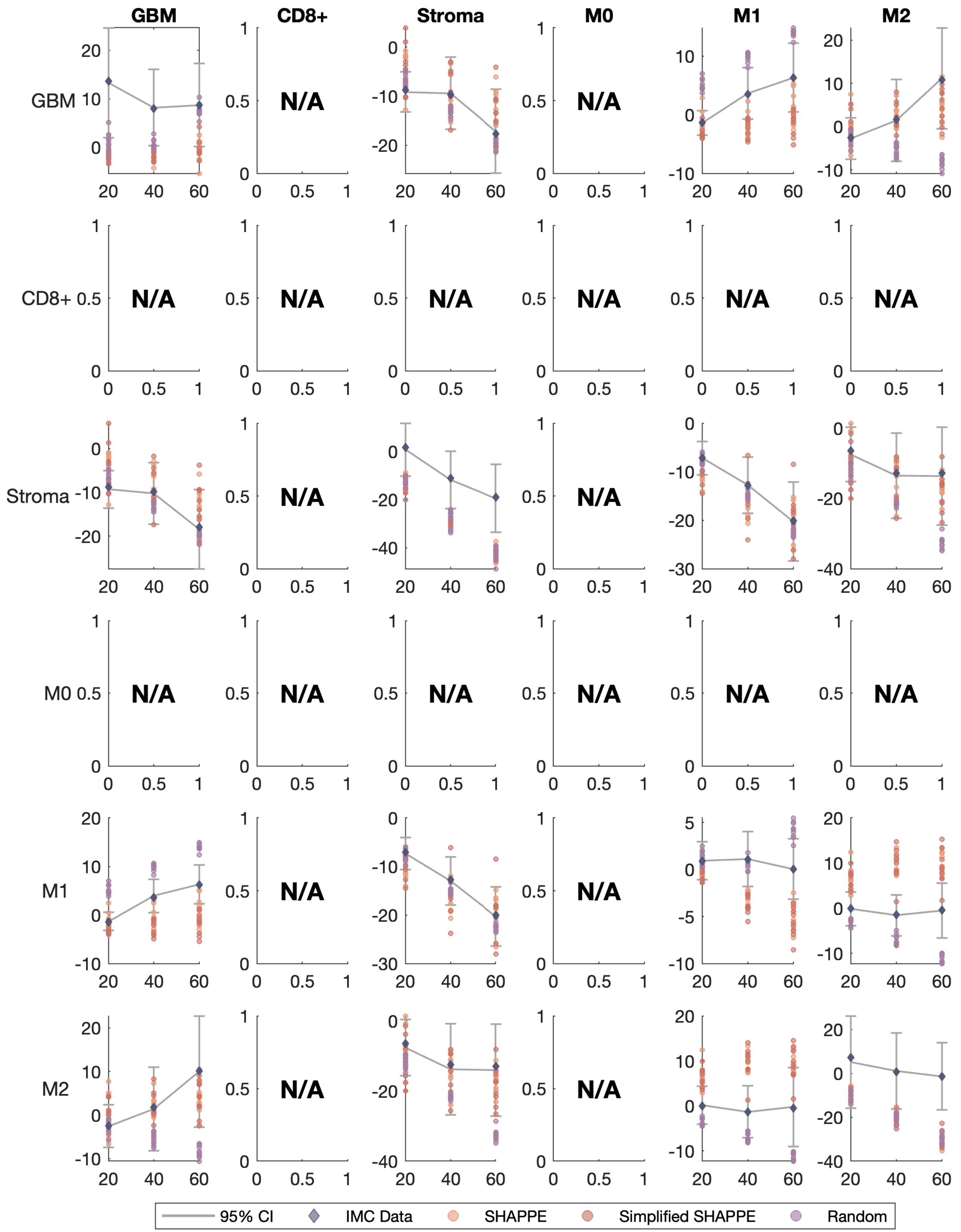

GBM
CD8+
Stroma
M0
M1
M2
GBM
CD8+
Stroma
M0
M1
M2
N/A
95% CI
IMC Data
SHAPPE
Simplified SHAPPE
Random

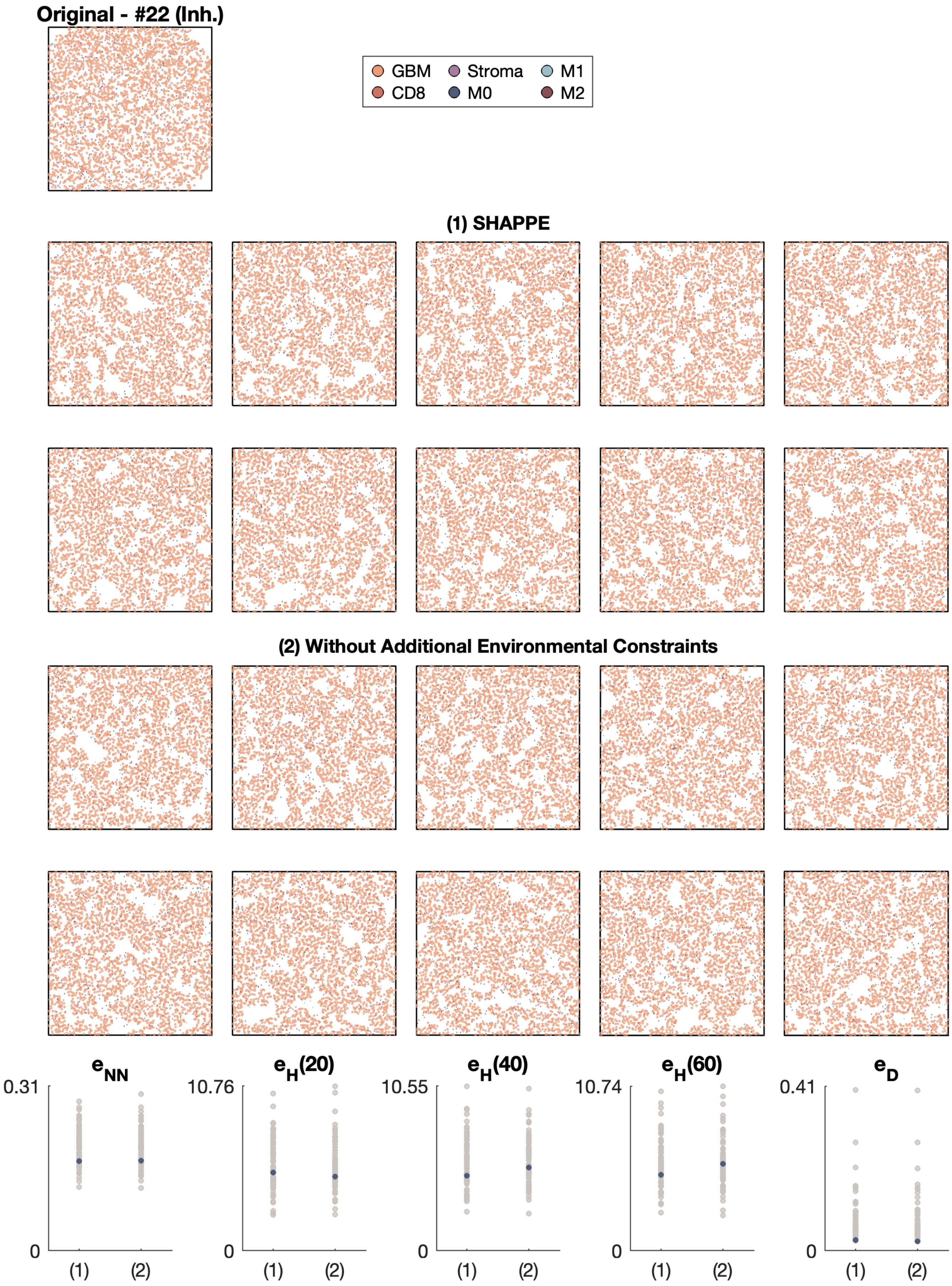

Original - #22 (Inh.)
GBM
Stroma
M1
CD8
M0
M2
(1) SHAPPE
(2) Without Additional Environmental Constraints
$e_{NN}$
$e_H(20)$
$e_H(40)$
$e_H(60)$
$e_D$
0.31
10.76
10.55
10.74
0.41
0
(1)
(2)

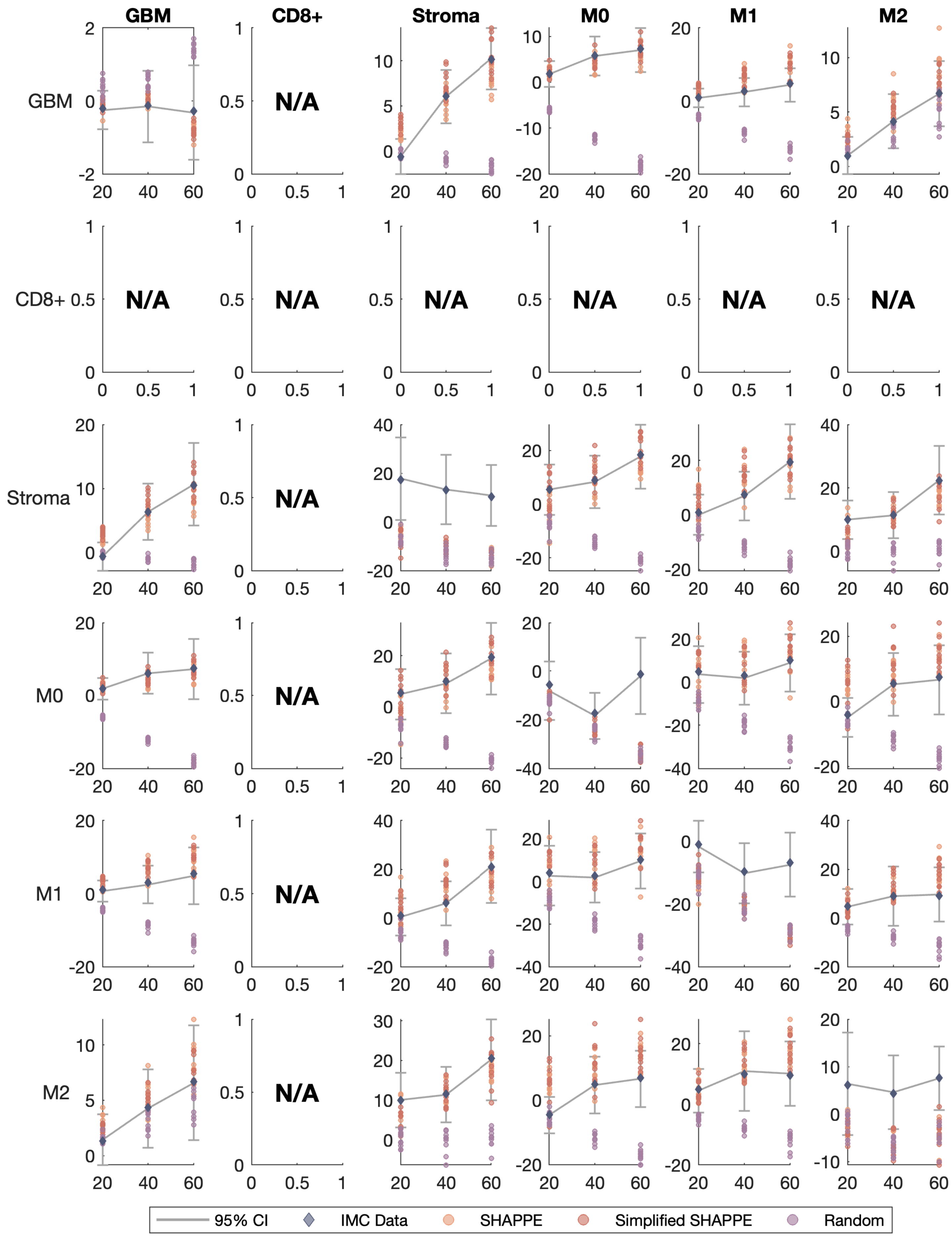
GBM
CD8+
Stroma
M0
M1
M2
N/A
95% CI
IMC Data
SHAPPE
Simplified SHAPPE
Random

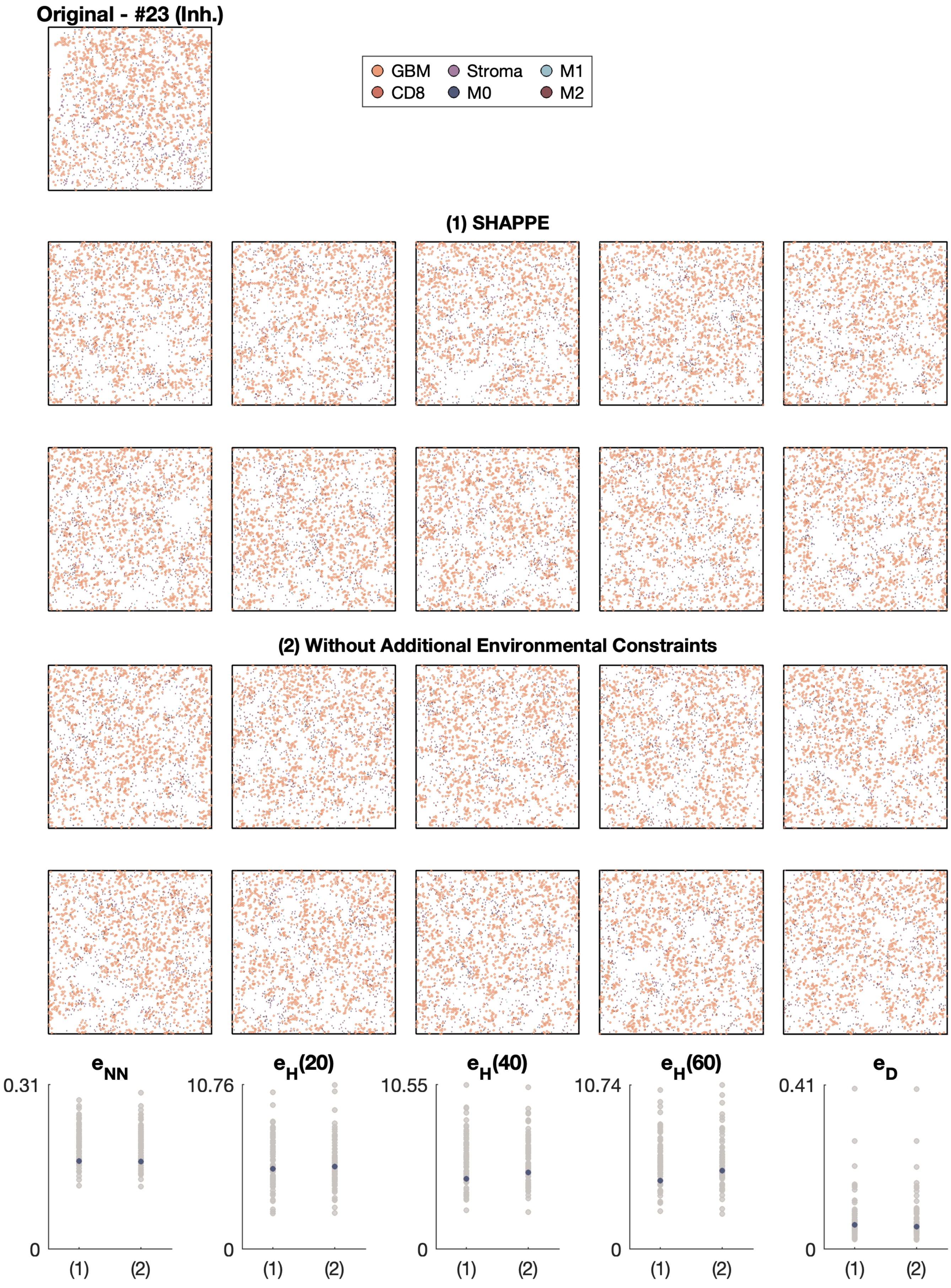
Original - #23 (Inh.)
GBM
CD8
Stroma
M0
M1
M2
(1) SHAPPE
(2) Without Additional Environmental Constraints
$e_{NN}$
$e_H(20)$
$e_H(40)$
$e_H(60)$
$e_D$
0.31
10.76
10.55
10.74
0.41
0
(1)
(2)

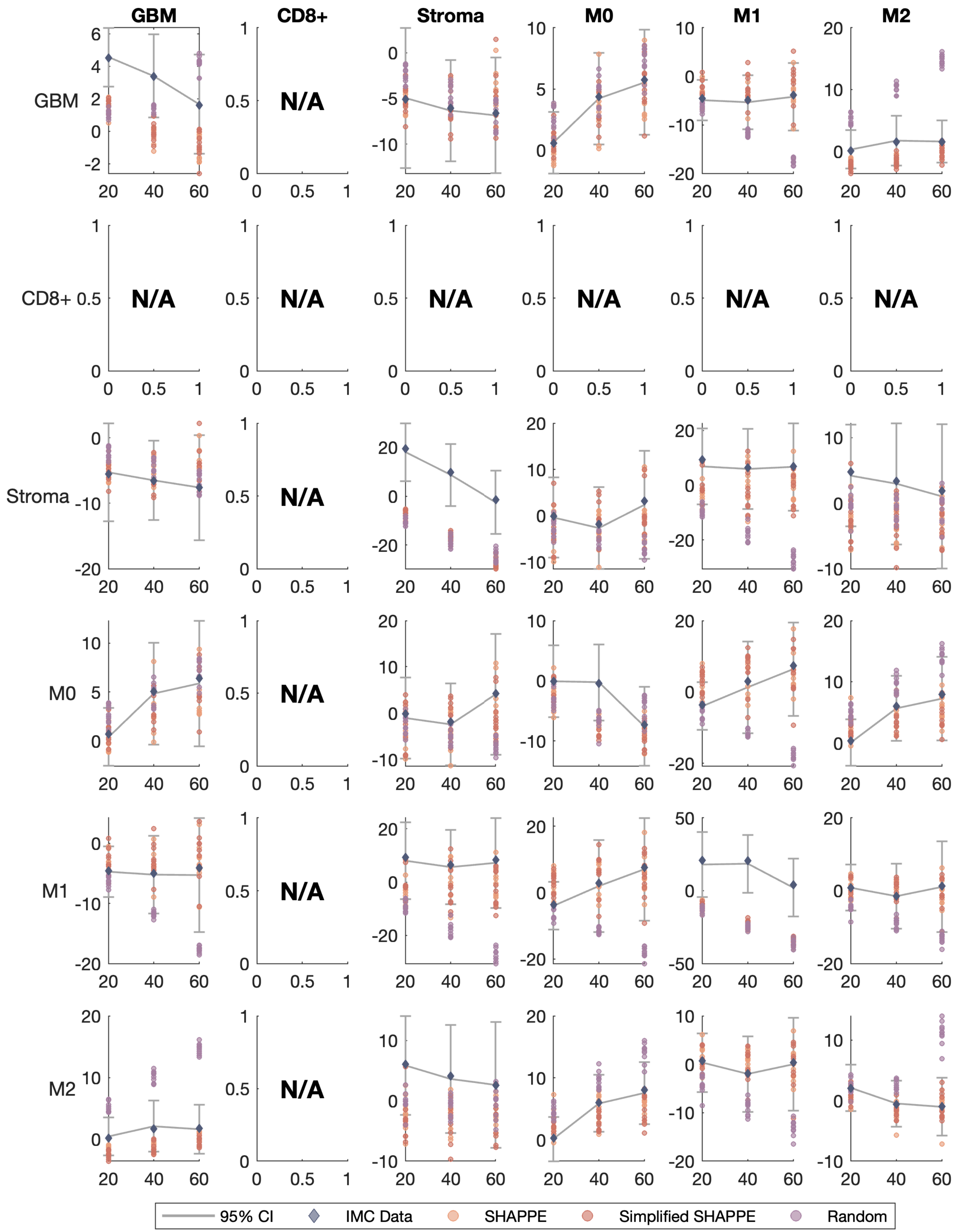
GBM
CD8+
Stroma
M0
M1
M2
N/A
95% CI
IMC Data
SHAPPE
Simplified SHAPPE
Random

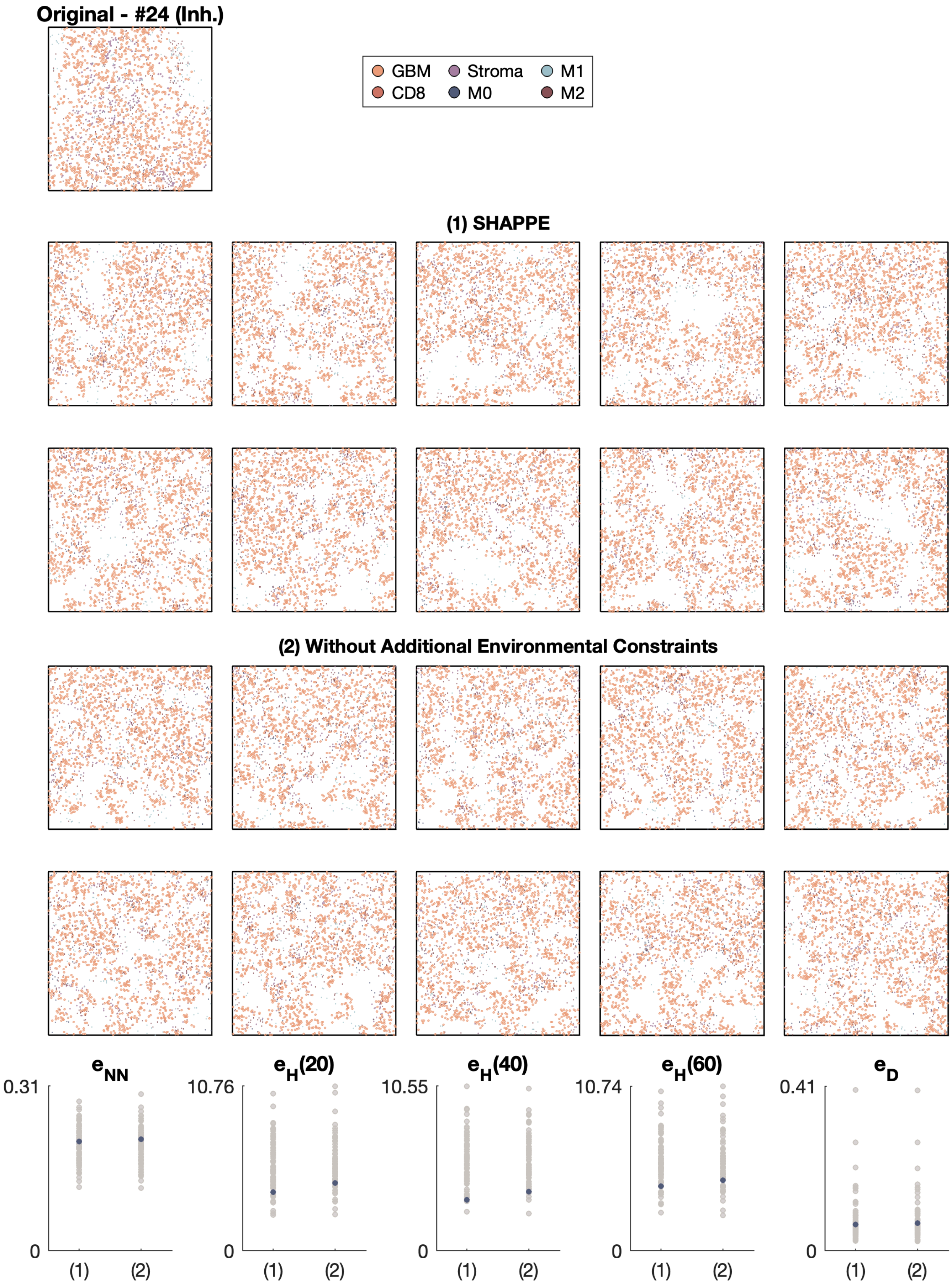

Original - #24 (Inh.)
GBM
Stroma
M1
CD8
M0
M2
(1) SHAPPE
(2) Without Additional Environmental Constraints
$e_{NN}$
$e_H(20)$
$e_H(40)$
$e_H(60)$
$e_D$
0.31
10.76
10.55
10.74
0.41
0
(1)
(2)

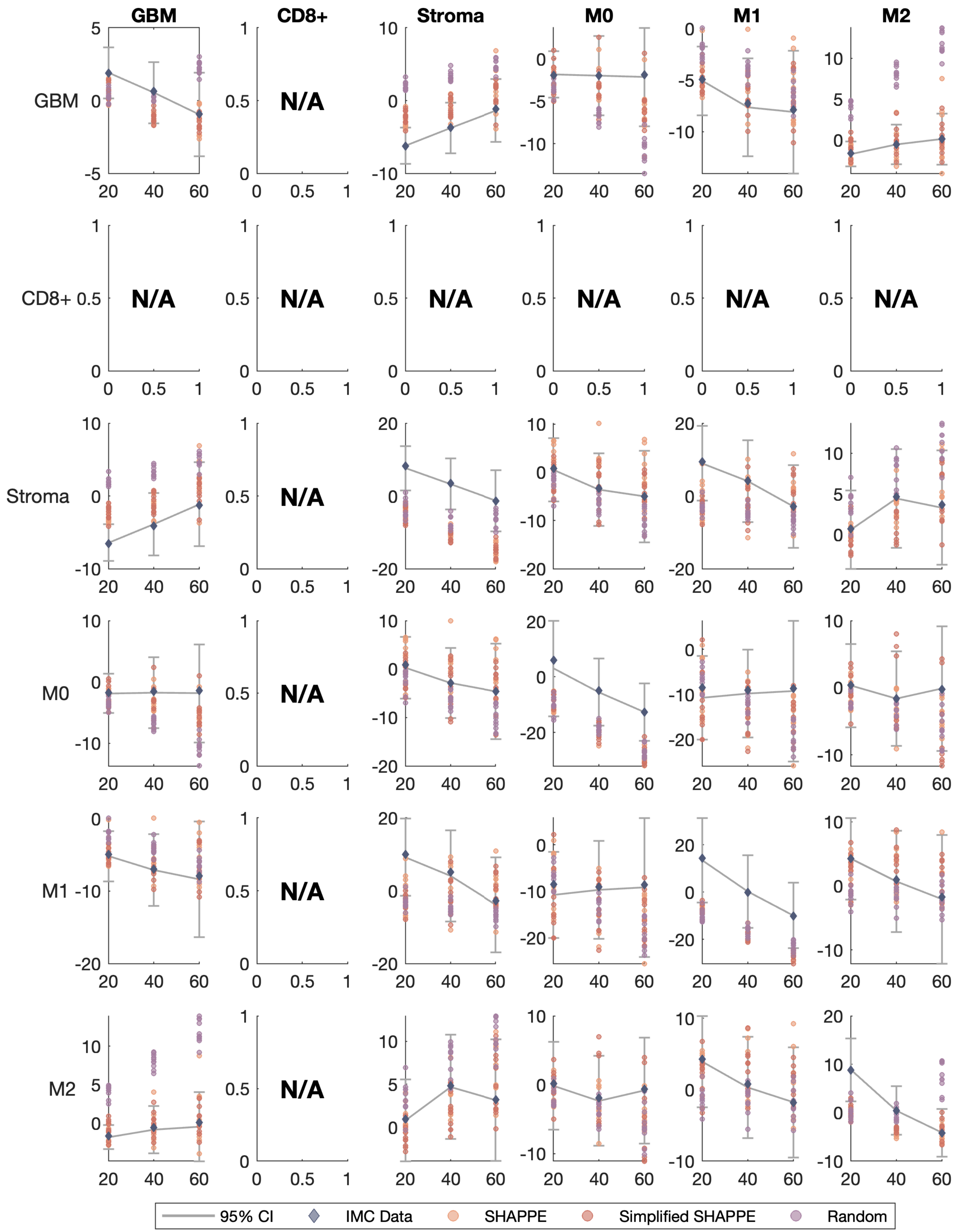
GBM
CD8+
Stroma
M0
M1
M2
N/A
95% CI
IMC Data
SHAPPE
Simplified SHAPPE
Random

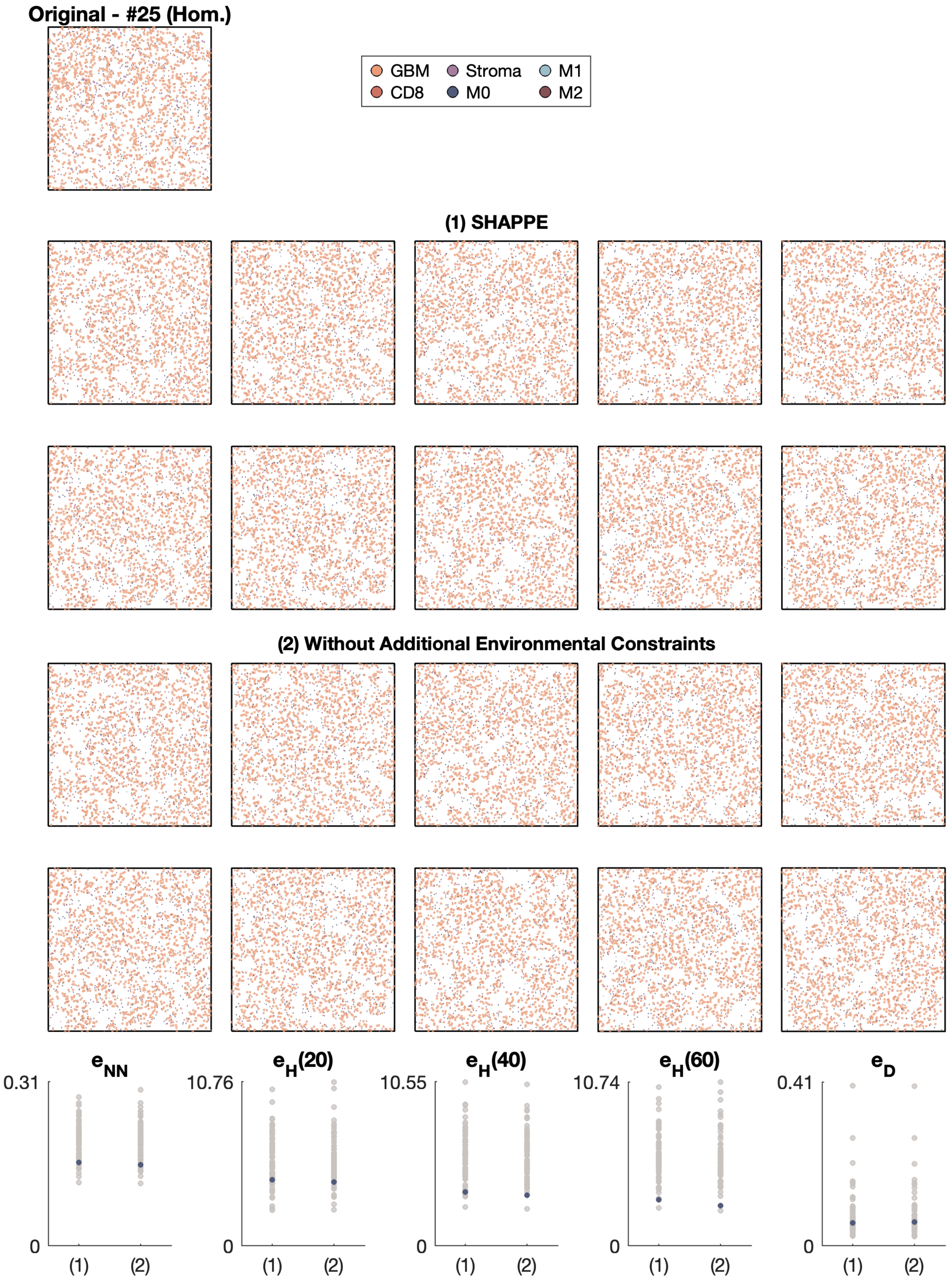
Original - #25 (Hom.)
GBM
Stroma
M1
CD8
M0
M2
(1) SHAPPE
(2) Without Additional Environmental Constraints
$e_{NN}$
$e_H(20)$
$e_H(40)$
$e_H(60)$
$e_D$
0.31
10.76
10.55
10.74
0.41
0
(1)
(2)

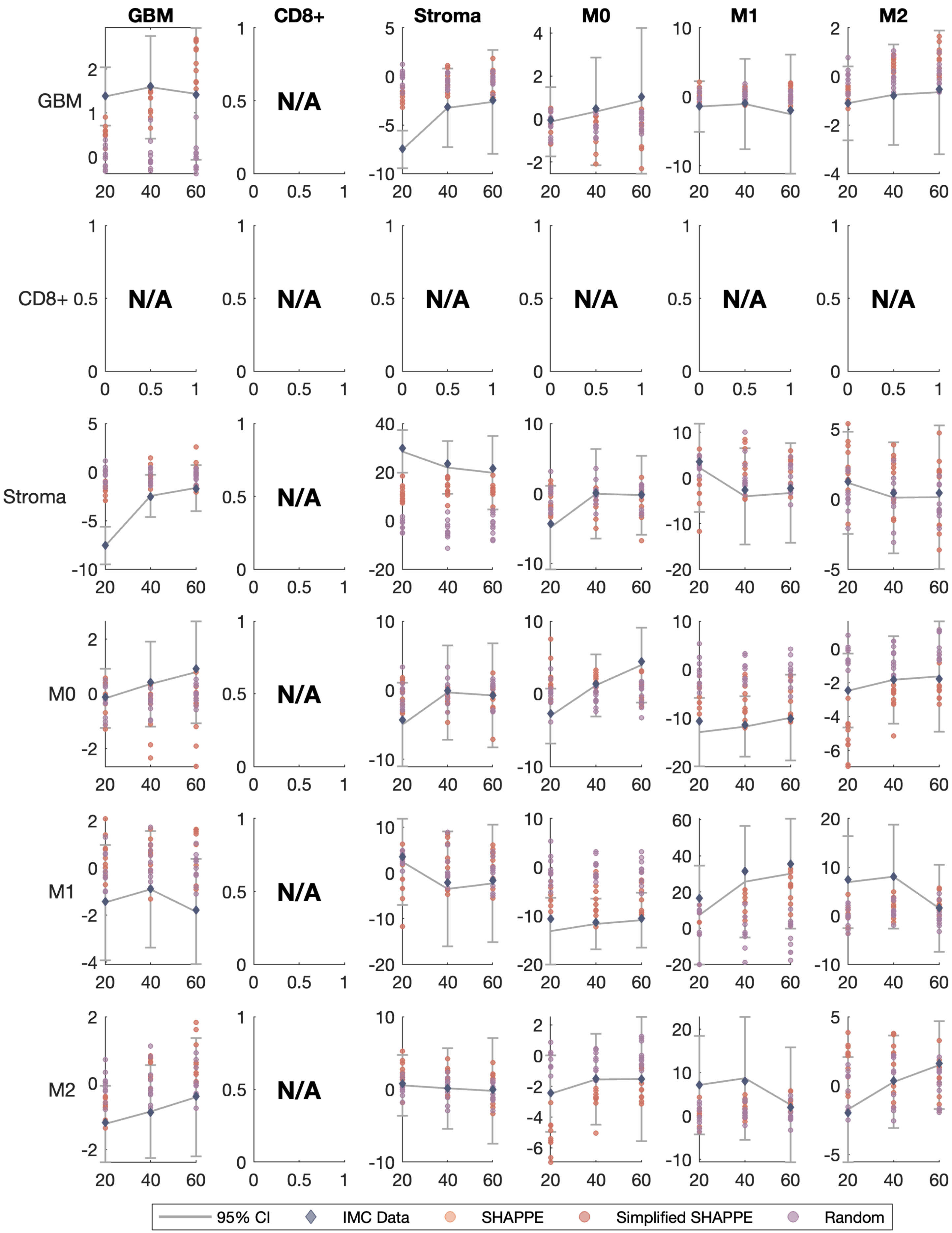

GBM
CD8+
Stroma
M0
M1
M2
N/A
95% CI
IMC Data
SHAPPE
Simplified SHAPPE
Random

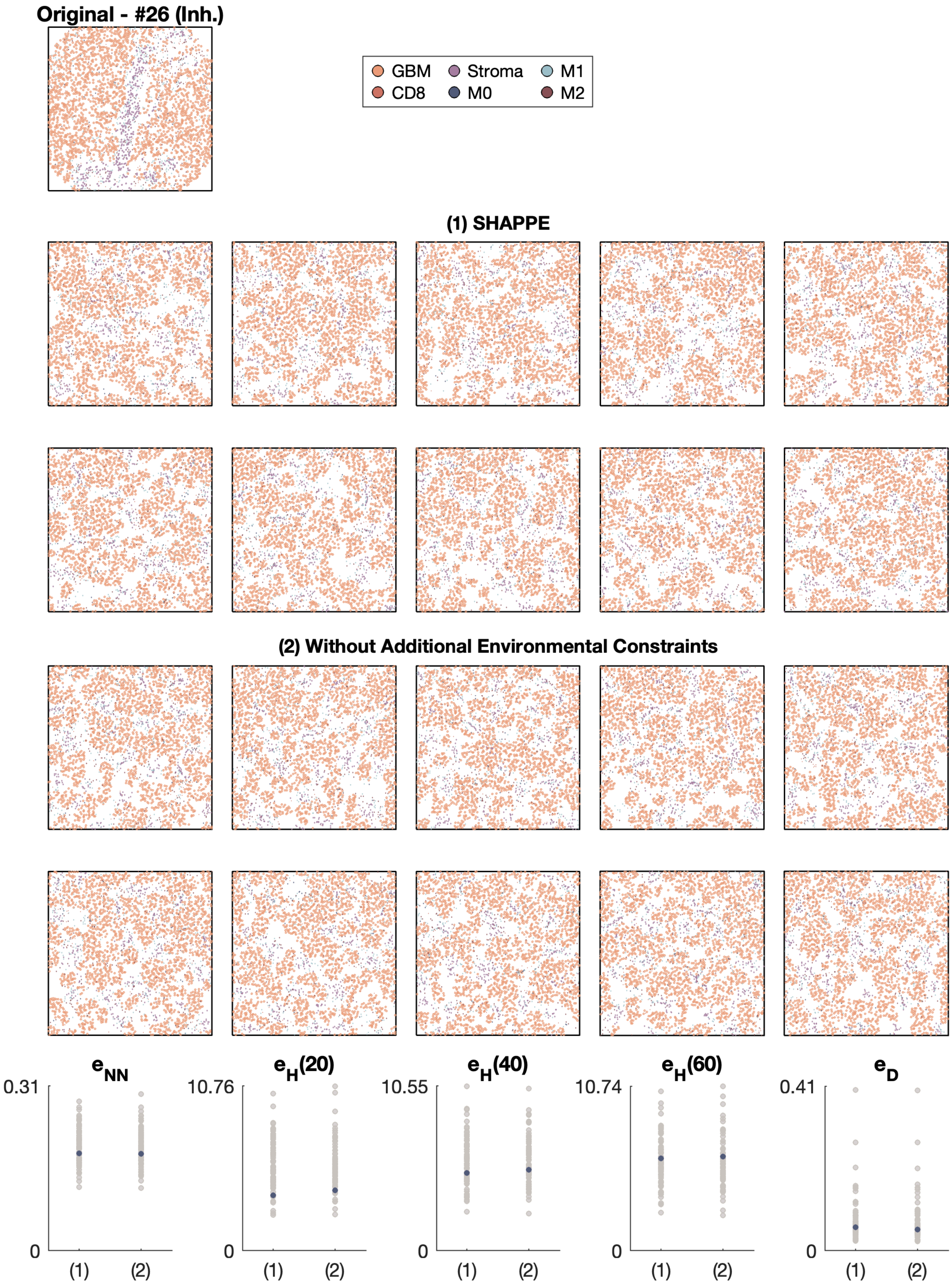

Original - #26 (Inh.)
GBM
Stroma
M1
CD8
M0
M2
(1) SHAPPE
(2) Without Additional Environmental Constraints
$e_{NN}$
$e_H(20)$
$e_H(40)$
$e_H(60)$
$e_D$
0.31
10.76
10.55
10.74
0.41
0
(1)
(2)

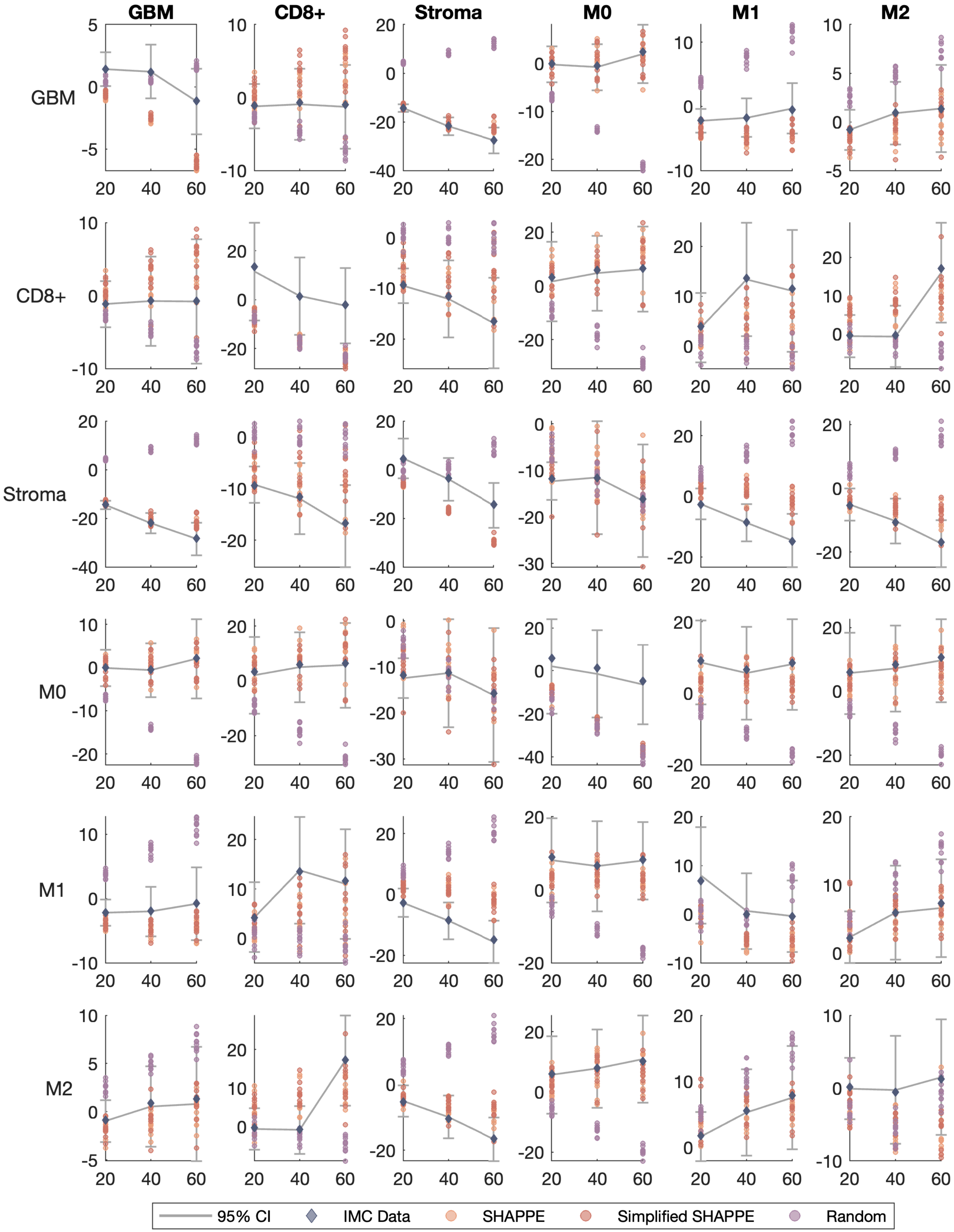

GBM
CD8+
Stroma
M0
M1
M2
GBM
CD8+
Stroma
M0
M1
M2
95% CI
IMC Data
SHAPPE
Simplified SHAPPE
Random

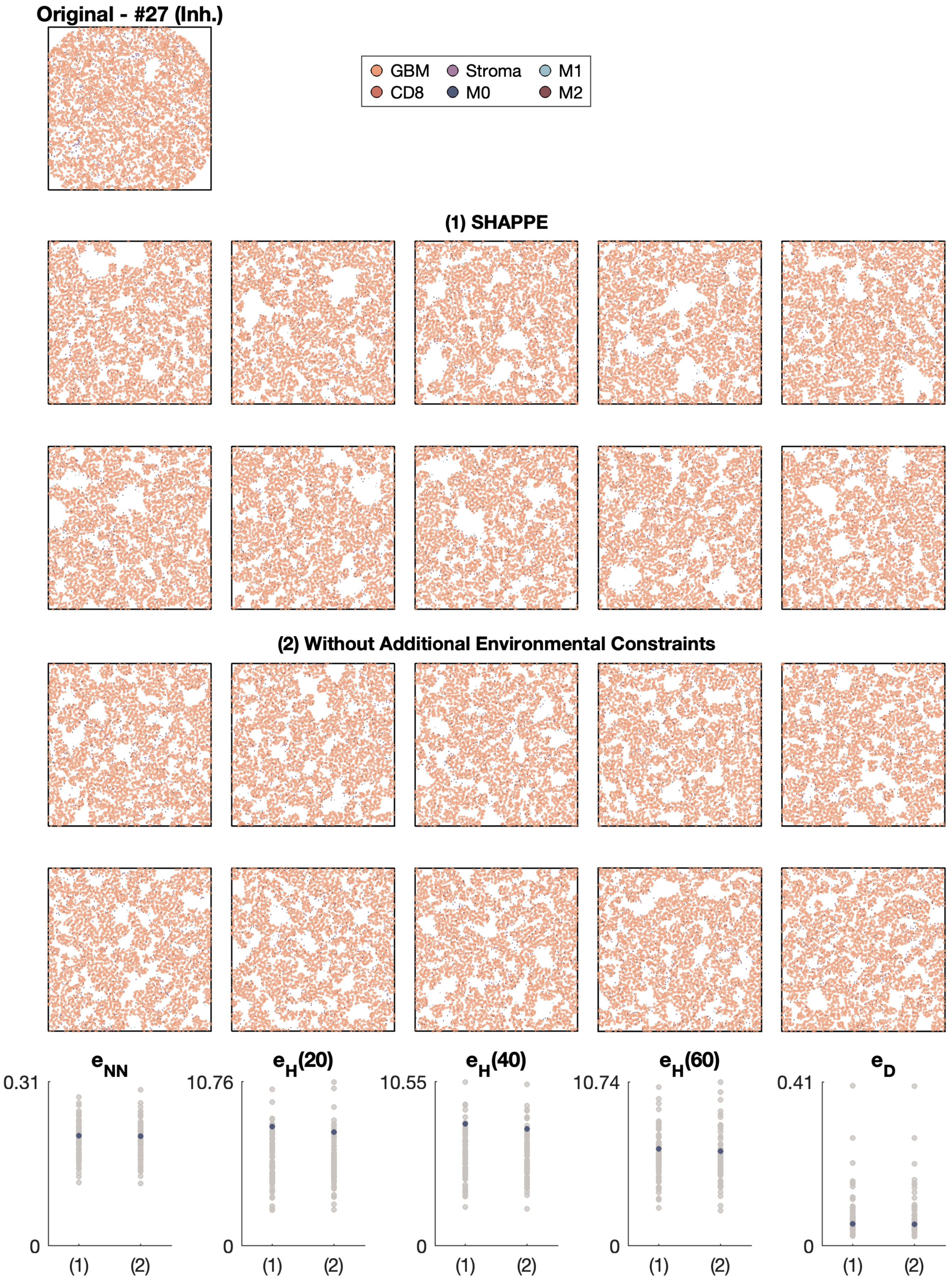

Original - #27 (Inh.)
GBM
CD8
Stroma
M0
M1
M2
(1) SHAPPE
(2) Without Additional Environmental Constraints
$e_{NN}$
$e_H(20)$
$e_H(40)$
$e_H(60)$
$e_D$
0.31
10.76
10.55
10.74
0.41
0
(1)
(2)

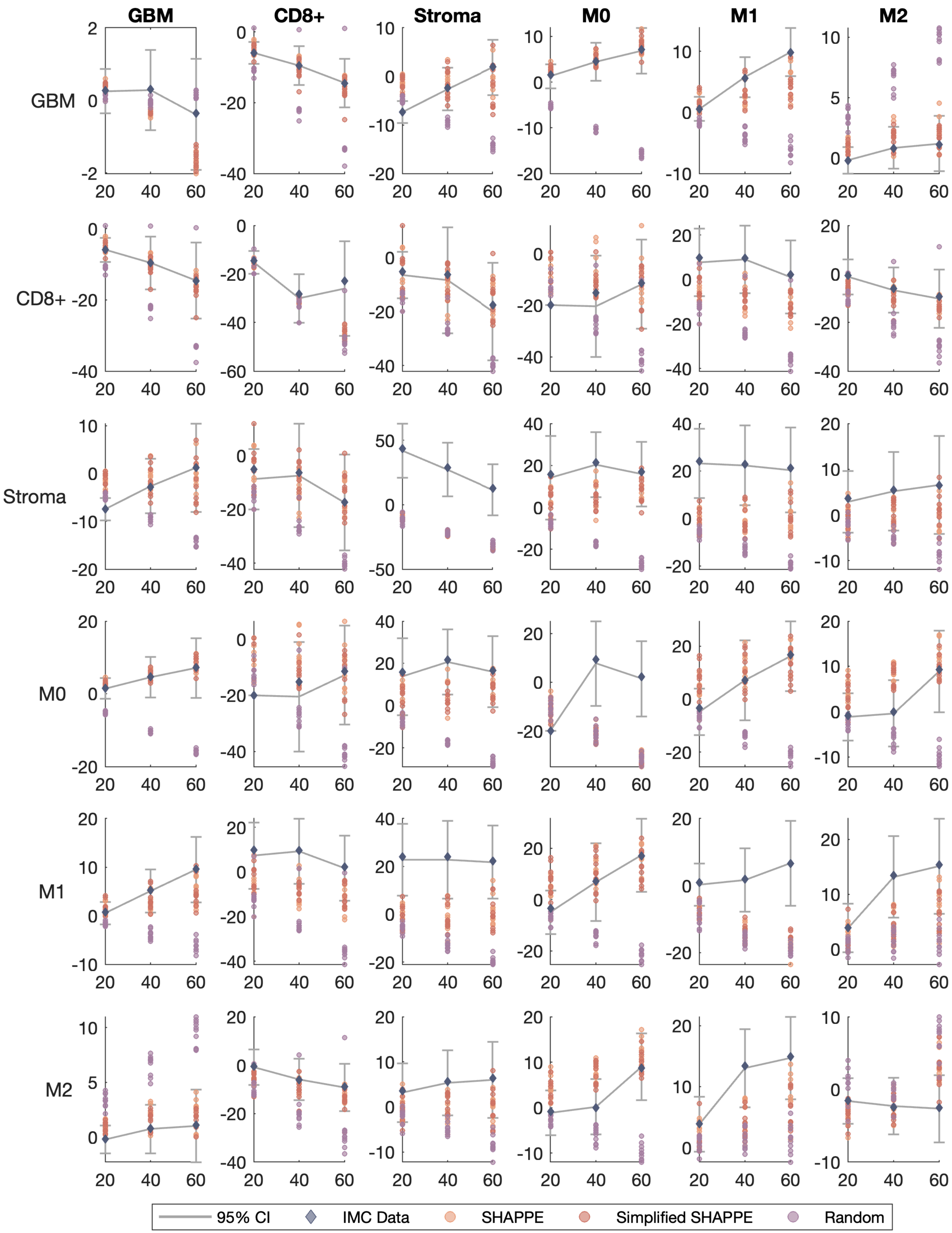

GBM
CD8+
Stroma
M0
M1
M2
GBM
CD8+
Stroma
M0
M1
M2
95% CI
IMC Data
SHAPPE
Simplified SHAPPE
Random

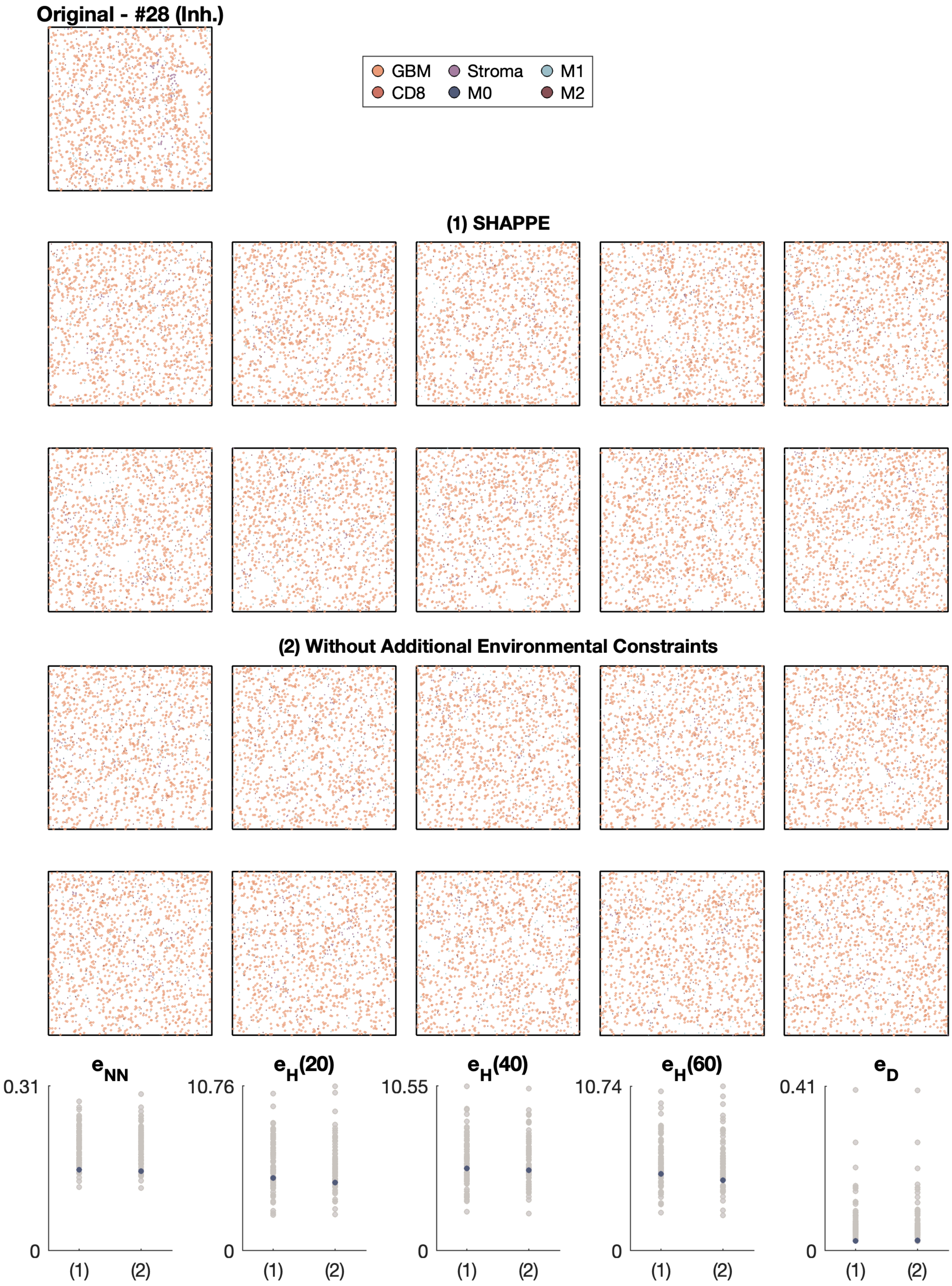

Original - #28 (Inh.)
GBM
Stroma
M1
CD8
M0
M2
(1) SHAPPE
(2) Without Additional Environmental Constraints
e_NN
e_H(20)
e_H(40)
e_H(60)
e_D
0.31
10.76
10.55
10.74
0.41
0
(1)
(2)

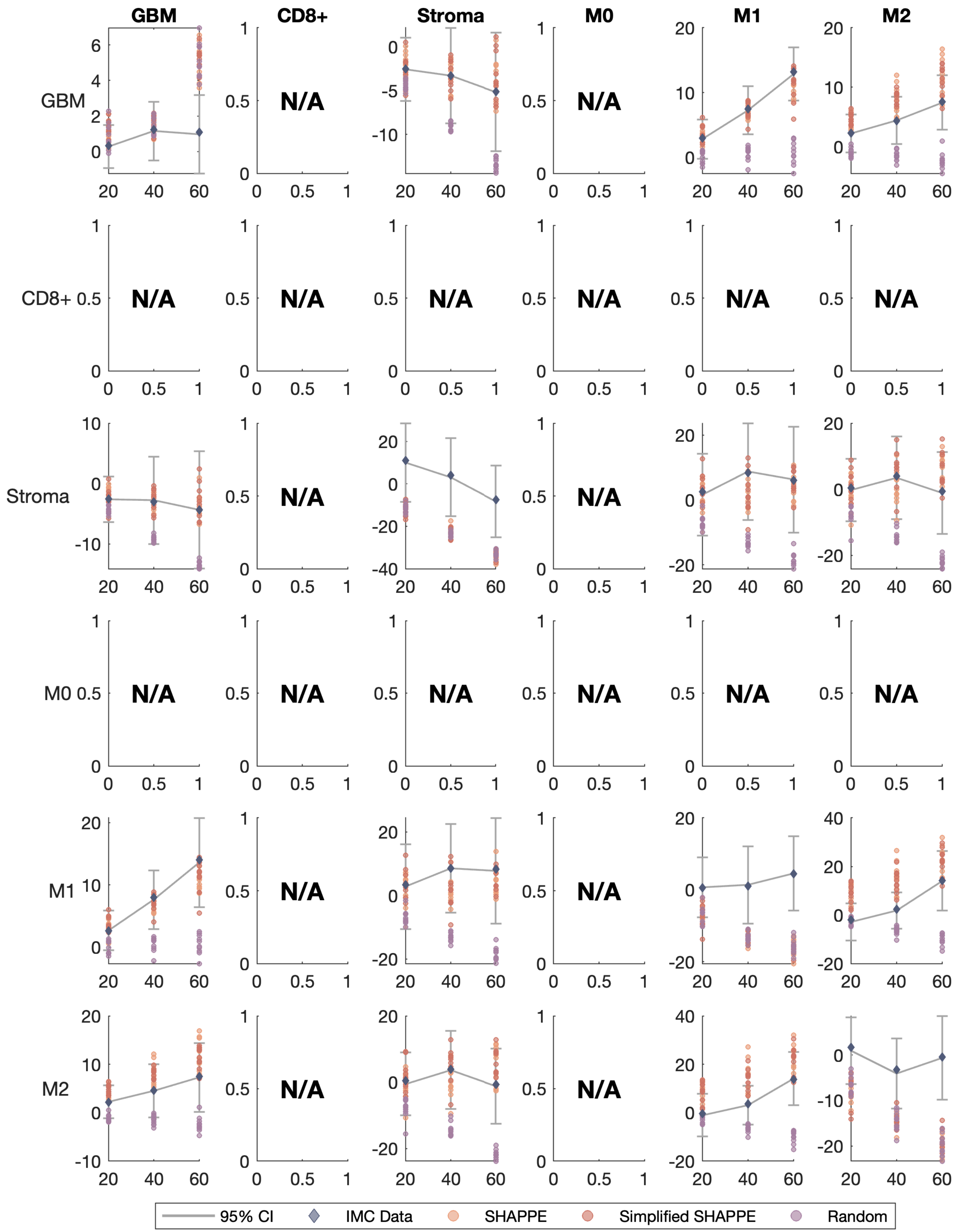
GBM
CD8+
Stroma
M0
M1
M2
GBM
CD8+
Stroma
M0
M1
M2
N/A
95% CI
IMC Data
SHAPPE
Simplified SHAPPE
Random

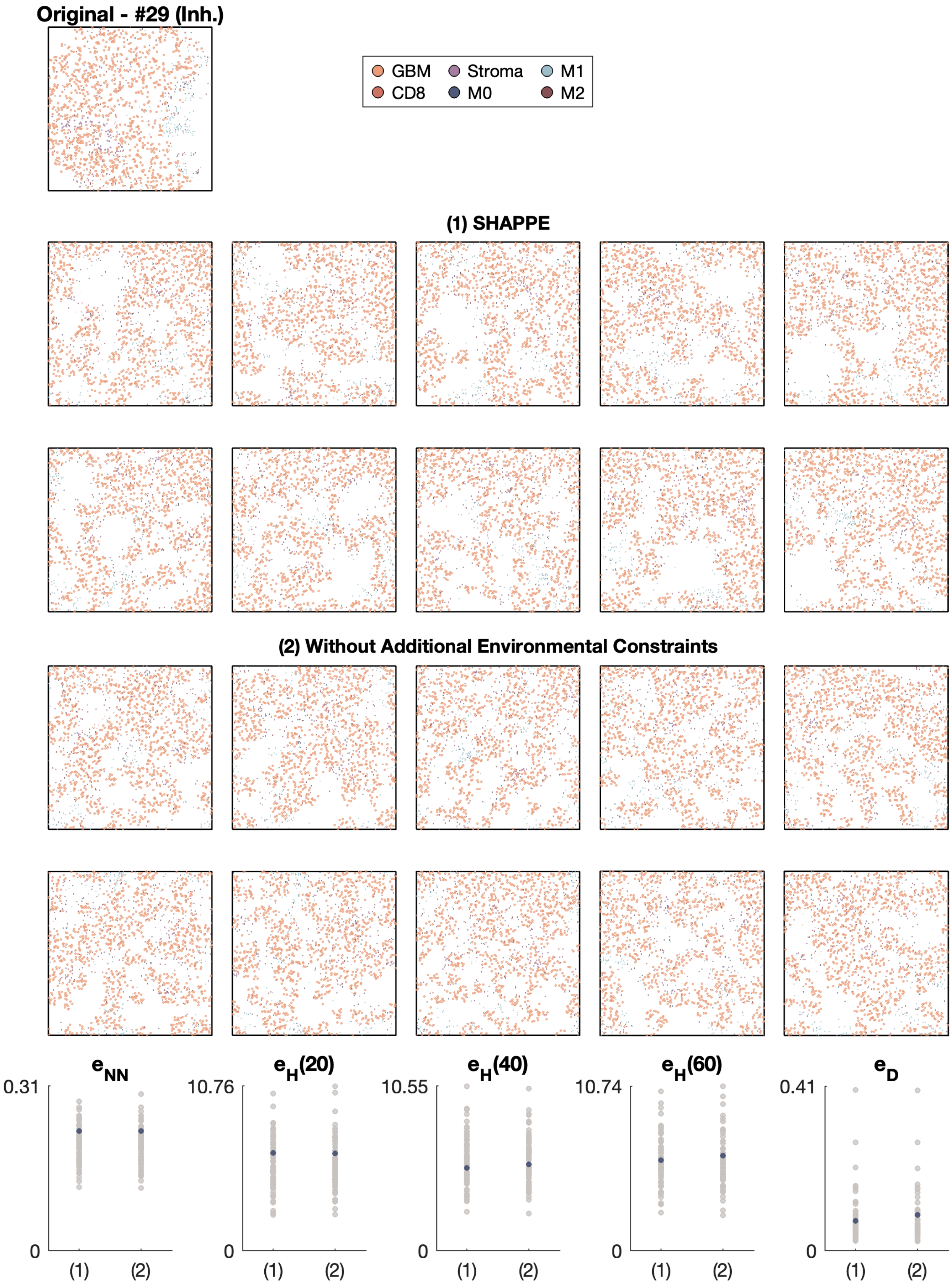

Original - #29 (Inh.)
GBM
Stroma
M1
CD8
M0
M2
(1) SHAPPE
(2) Without Additional Environmental Constraints
e_NN
0.31
0
(1)
(2)
e_H(20)
10.76
0
(1)
(2)
e_H(40)
10.55
0
(1)
(2)
e_H(60)
10.74
0
(1)
(2)
e_D
0.41
0
(1)
(2)

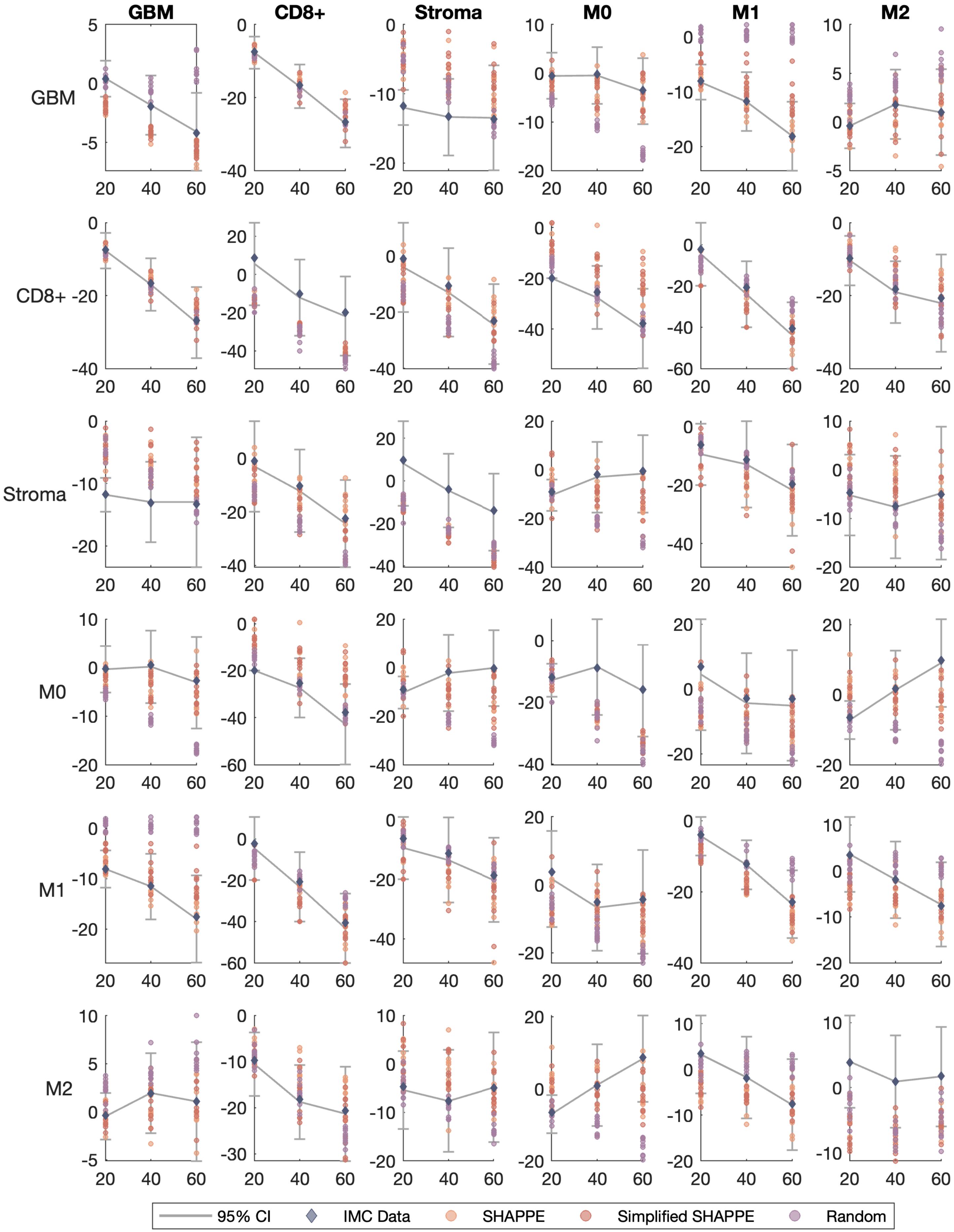

GBM
CD8+
Stroma
M0
M1
M2
95% CI
IMC Data
SHAPPE
Simplified SHAPPE
Random

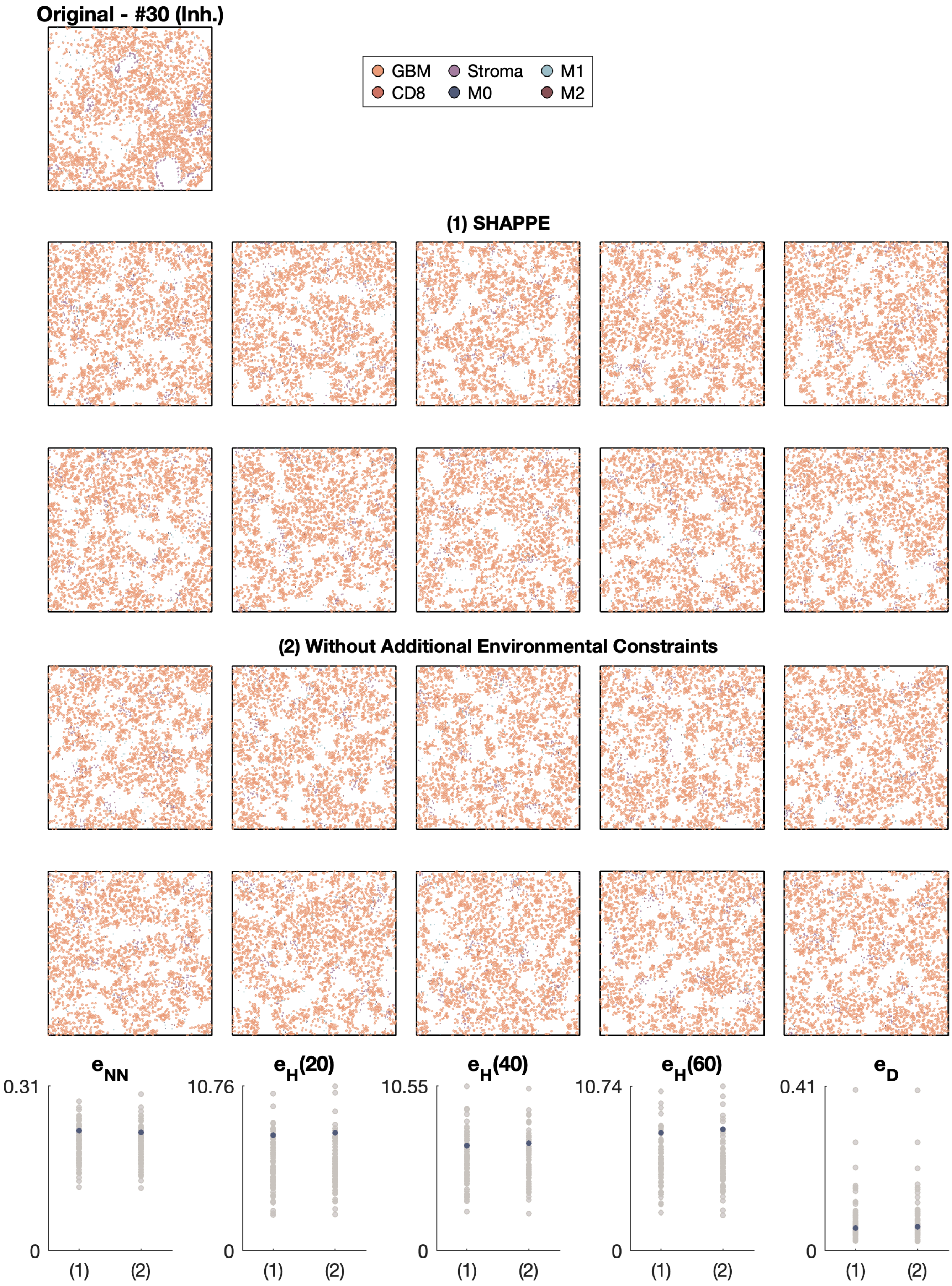

Original - #30 (Inh.)
GBM
Stroma
M1
CD8
M0
M2
(1) SHAPPE
(2) Without Additional Environmental Constraints
e_NN
e_H(20)
e_H(40)
e_H(60)
e_D
0.31
10.76
10.55
10.74
0.41
0
(1)
(2)

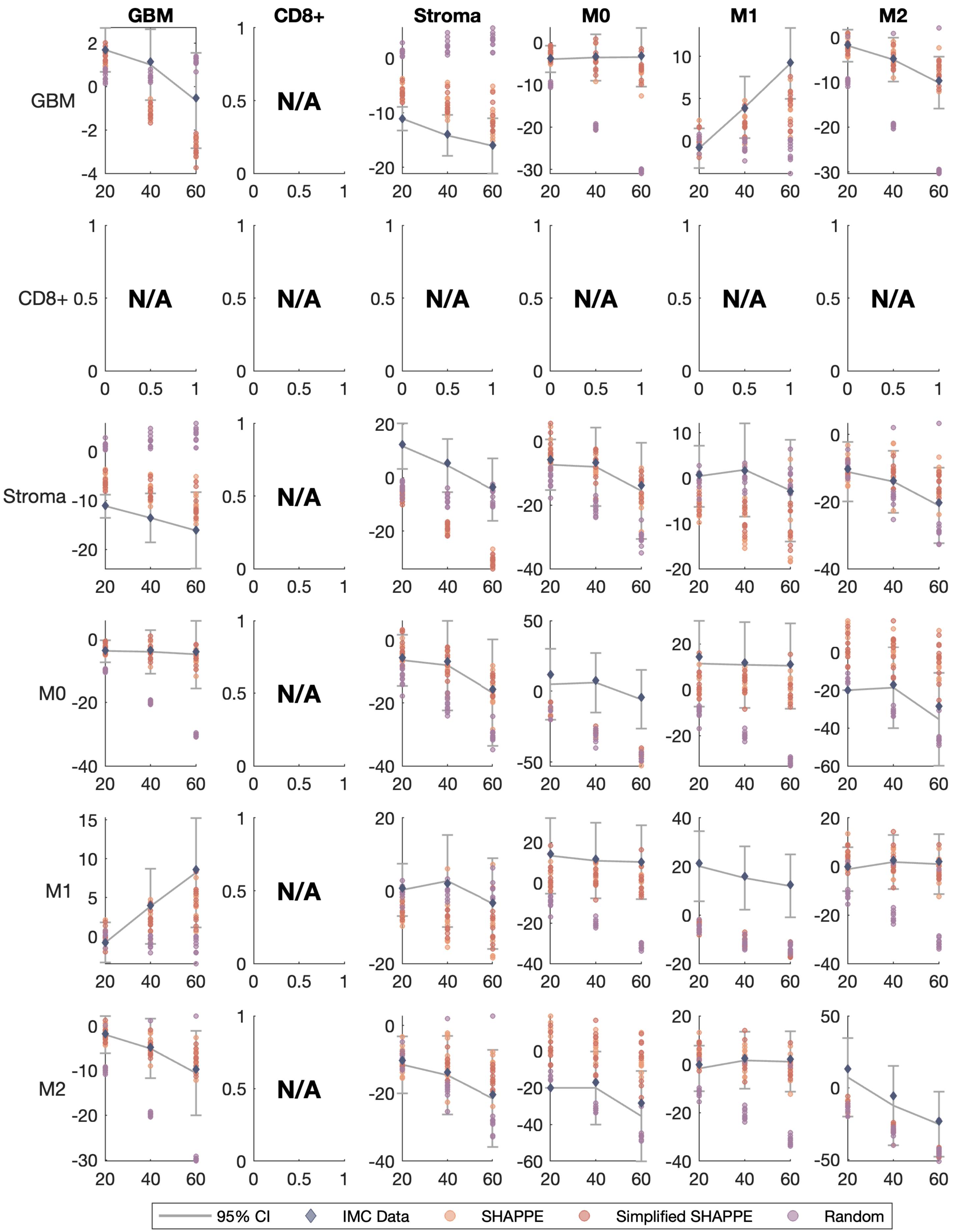

GBM
CD8+
Stroma
M0
M1
M2
N/A
95% CI
IMC Data
SHAPPE
Simplified SHAPPE
Random

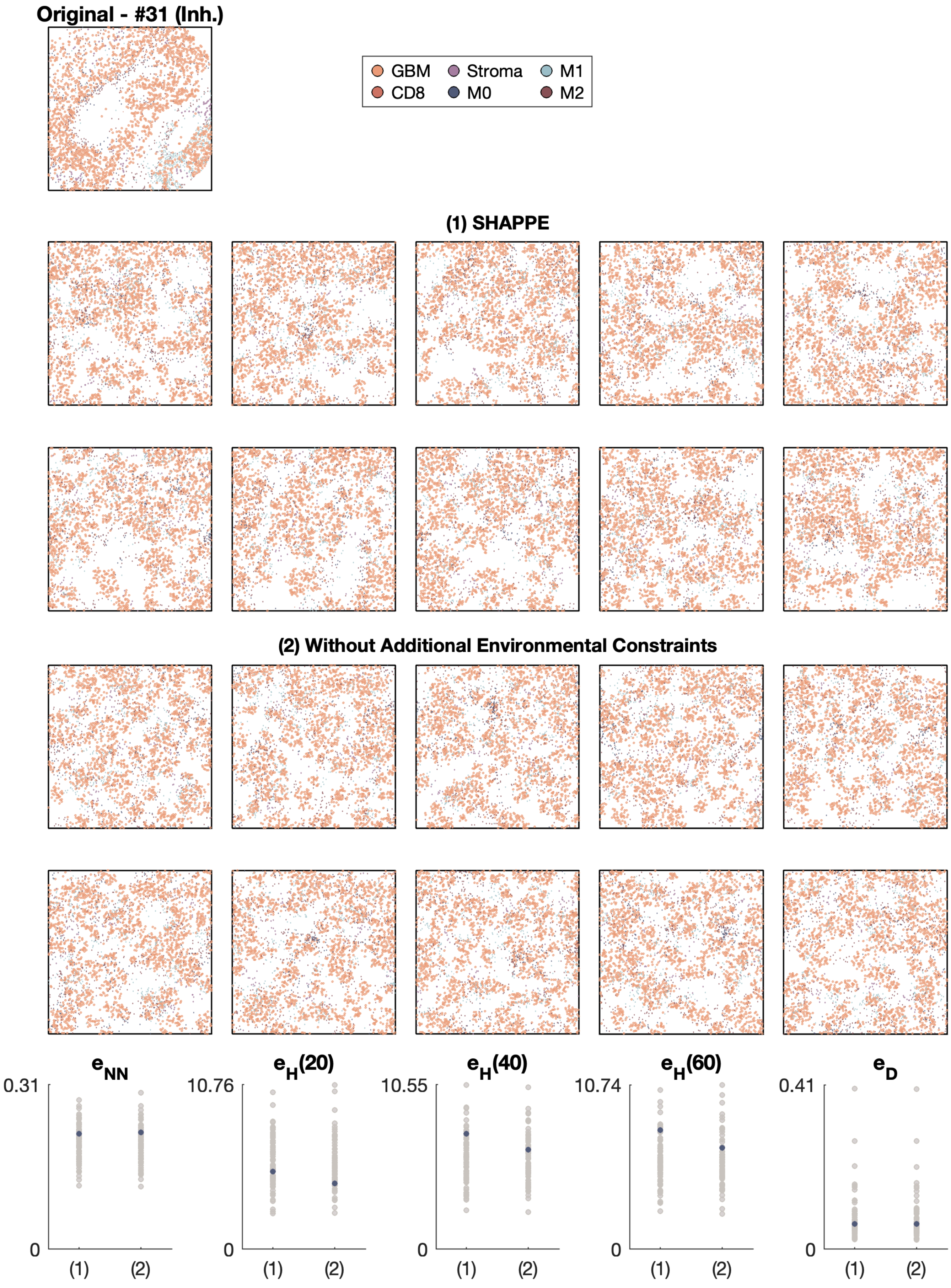

Original - #31 (Inh.)
GBM
Stroma
M1
CD8
M0
M2
(1) SHAPPE
(2) Without Additional Environmental Constraints
e_NN
e_H(20)
e_H(40)
e_H(60)
e_D
0.31
10.76
10.55
10.74
0.41
0
(1)
(2)

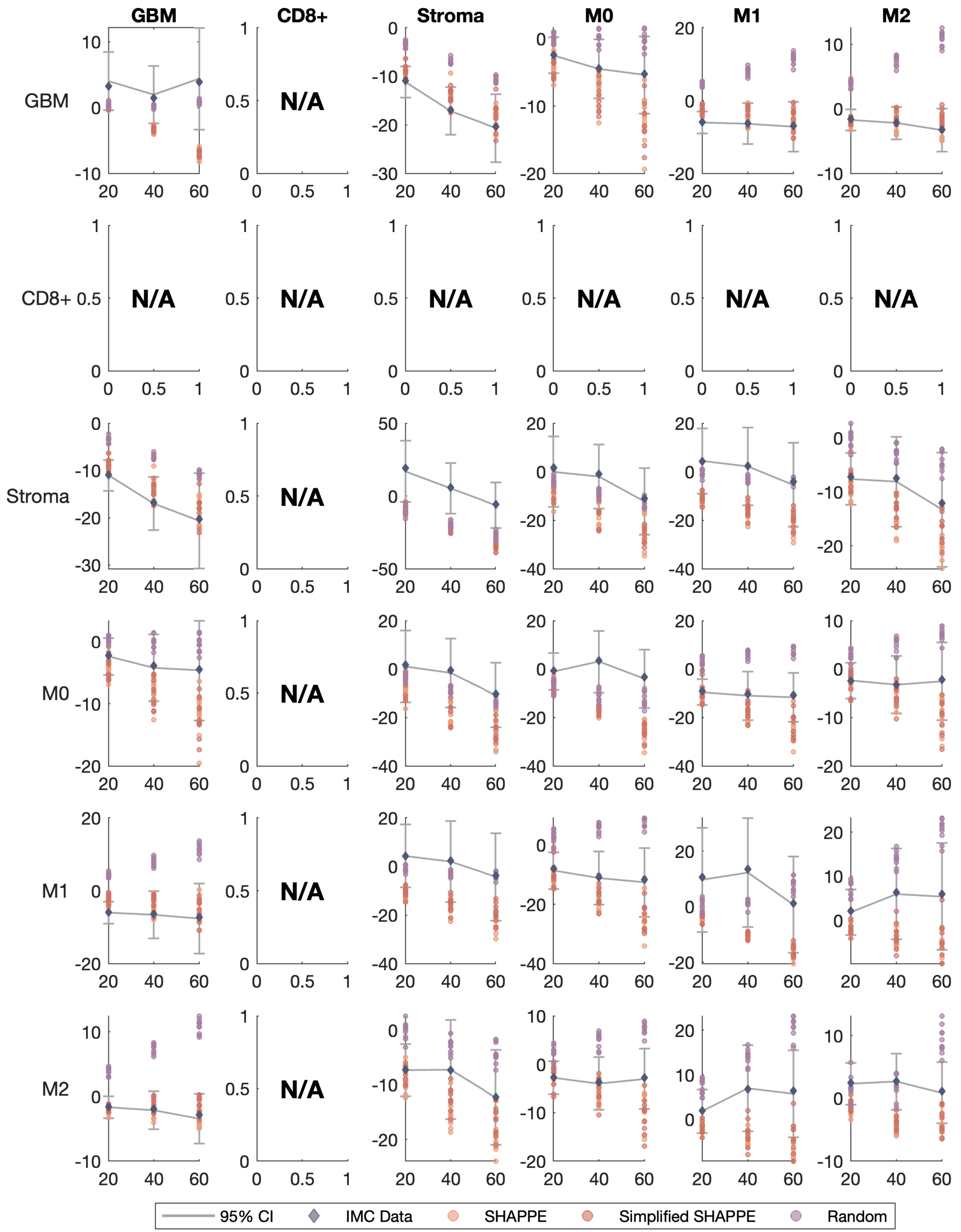

GBM
CD8+
Stroma
M0
M1
M2
N/A
95% CI
IMC Data
SHAPPE
Simplified SHAPPE
Random

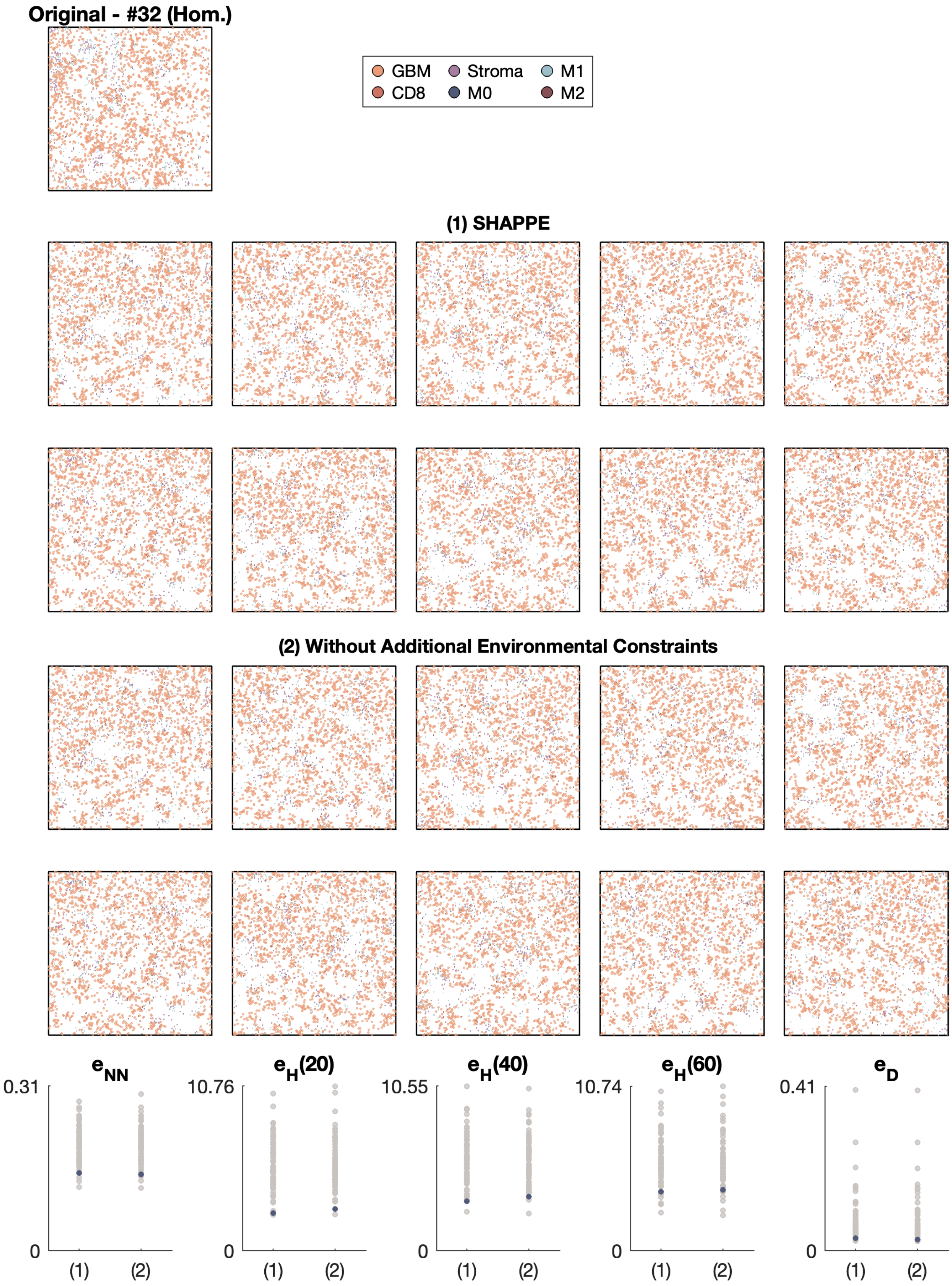

Original - #32 (Hom.)
GBM
Stroma
M1
CD8
M0
M2
(1) SHAPPE
(2) Without Additional Environmental Constraints
$e_{NN}$
$e_H(20)$
$e_H(40)$
$e_H(60)$
$e_D$
0.31
10.76
10.55
10.74
0.41
0
(1)
(2)

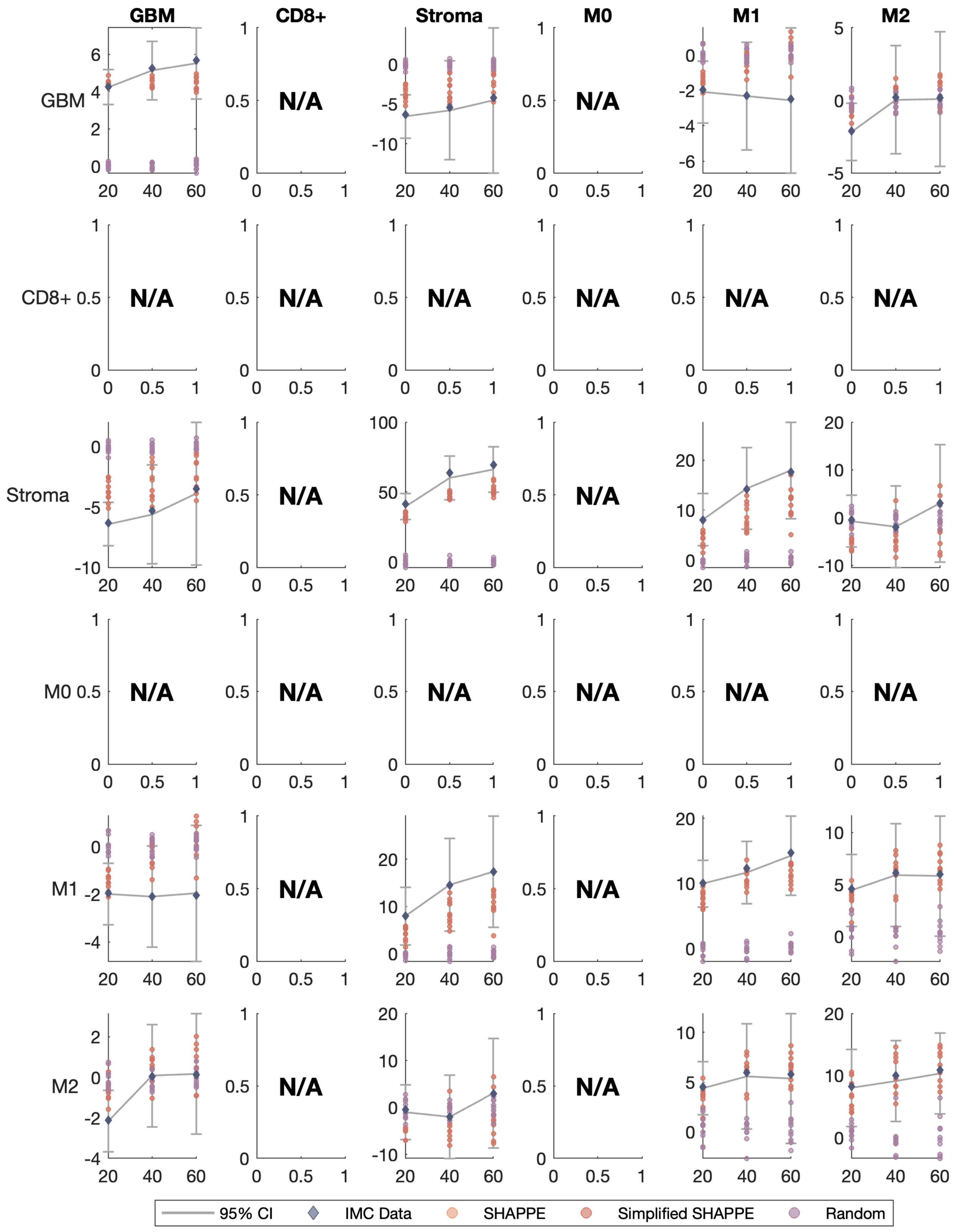
GBM
CD8+
Stroma
M0
M1
M2
GBM
CD8+
Stroma
M0
M1
M2
N/A
95% CI
IMC Data
SHAPPE
Simplified SHAPPE
Random

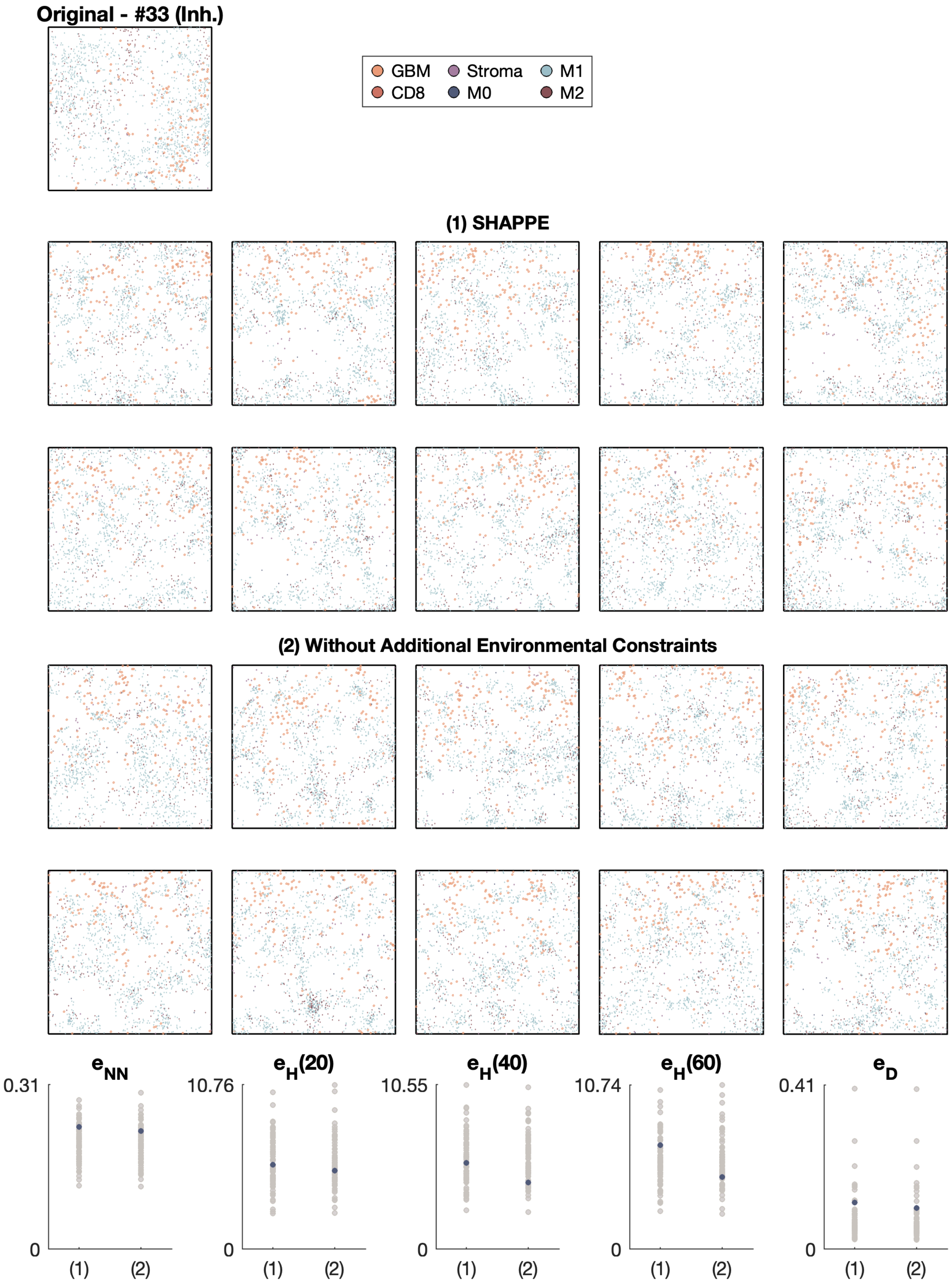

Original - #33 (Inh.)
GBM
CD8
Stroma
M0
M1
M2
(1) SHAPPE
(2) Without Additional Environmental Constraints
$e_{NN}$
0.31
0
(1)
(2)
$e_H(20)$
10.76
0
(1)
(2)
$e_H(40)$
10.55
0
(1)
(2)
$e_H(60)$
10.74
0
(1)
(2)
$e_D$
0.41
0
(1)
(2)

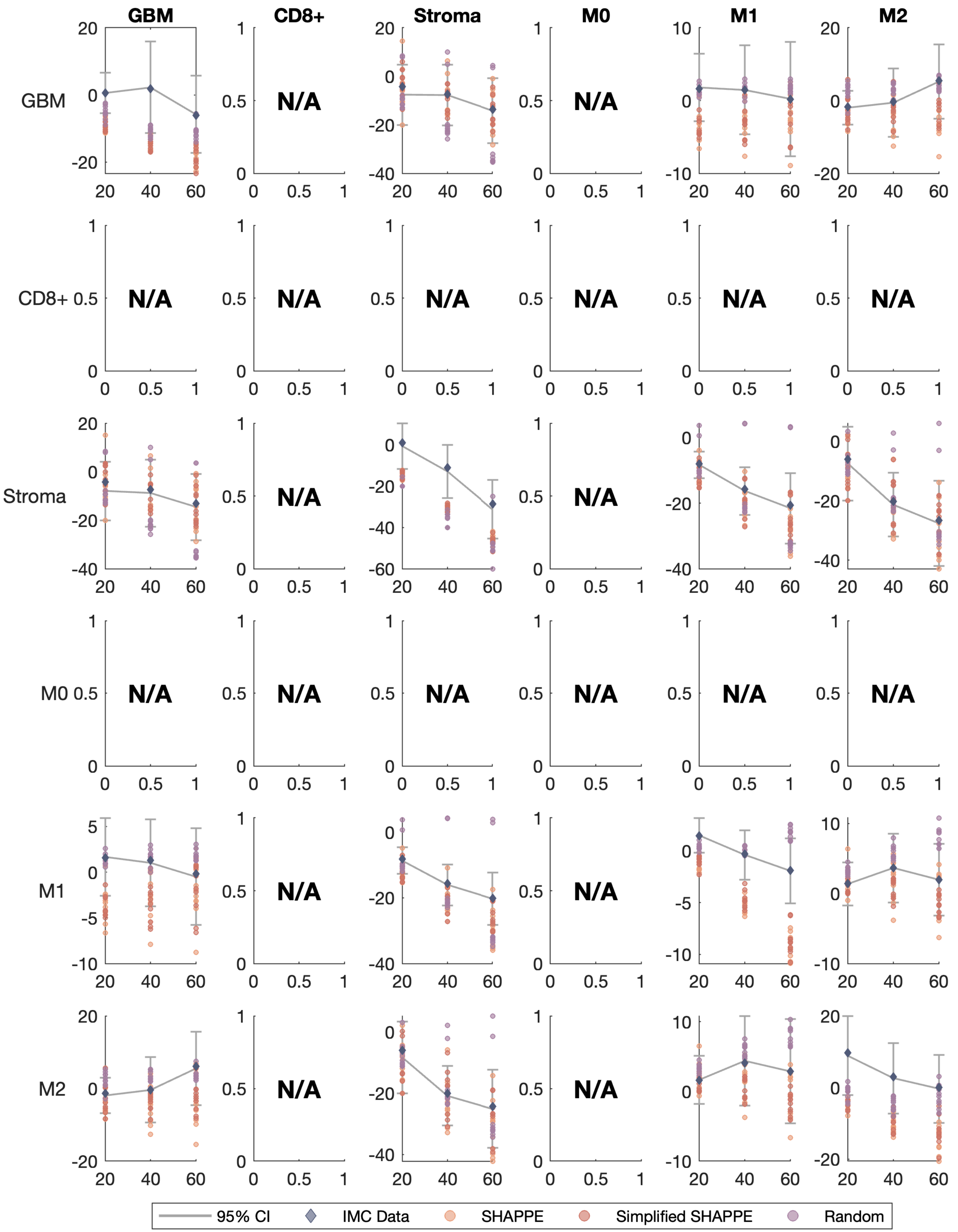
GBM
CD8+
Stroma
M0
M1
M2
GBM
CD8+
Stroma
M0
M1
M2
N/A
95% CI
IMC Data
SHAPPE
Simplified SHAPPE
Random

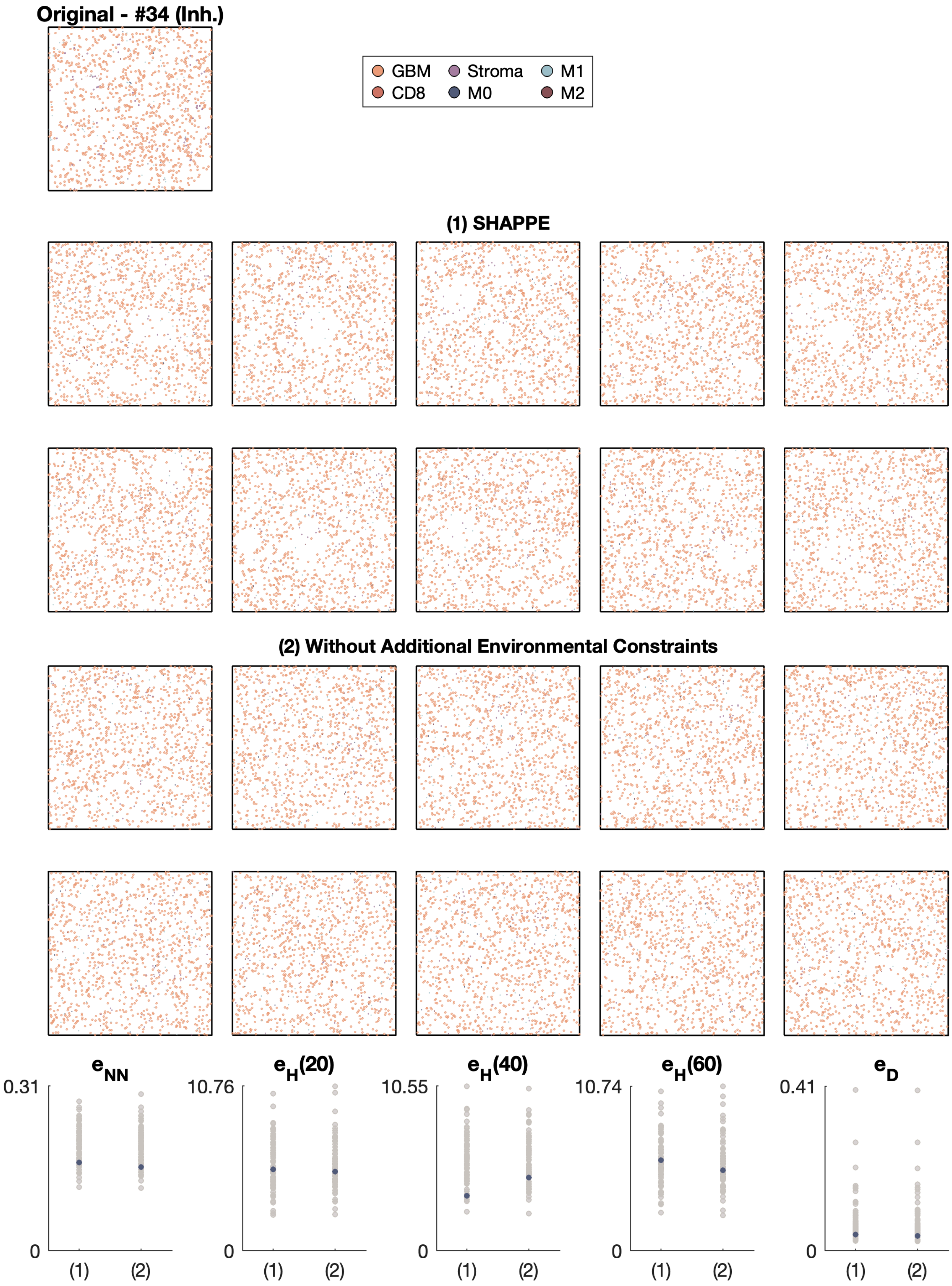
Original - #34 (Inh.)
GBM
CD8
Stroma
M0
M1
M2
(1) SHAPPE
(2) Without Additional Environmental Constraints
$e_{NN}$
0.31
0
(1)
(2)
$e_H(20)$
10.76
0
(1)
(2)
$e_H(40)$
10.55
0
(1)
(2)
$e_H(60)$
10.74
0
(1)
(2)
$e_D$
0.41
0
(1)
(2)

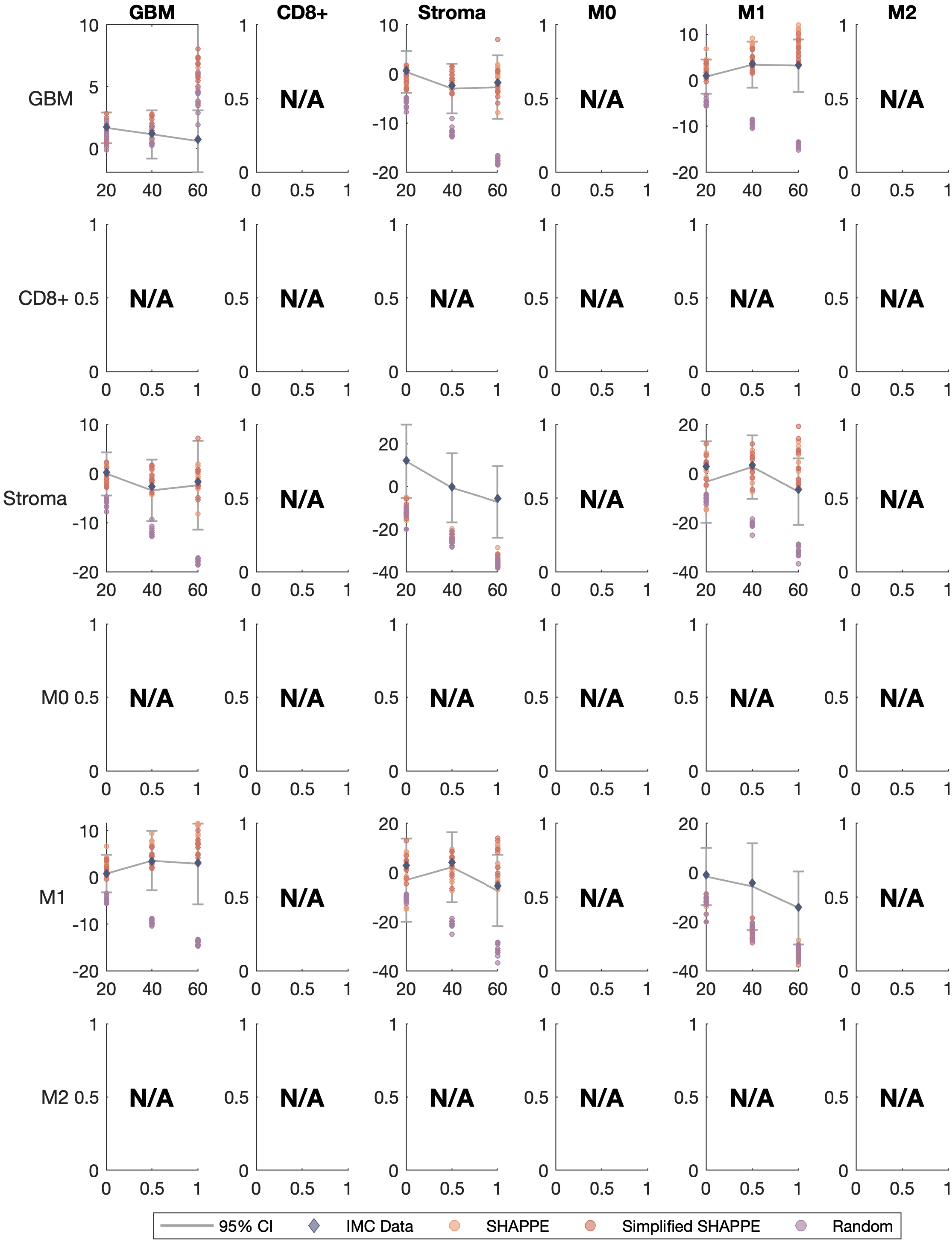

GBM
CD8+
Stroma
M0
M1
M2
N/A
95% CI
IMC Data
SHAPPE
Simplified SHAPPE
Random

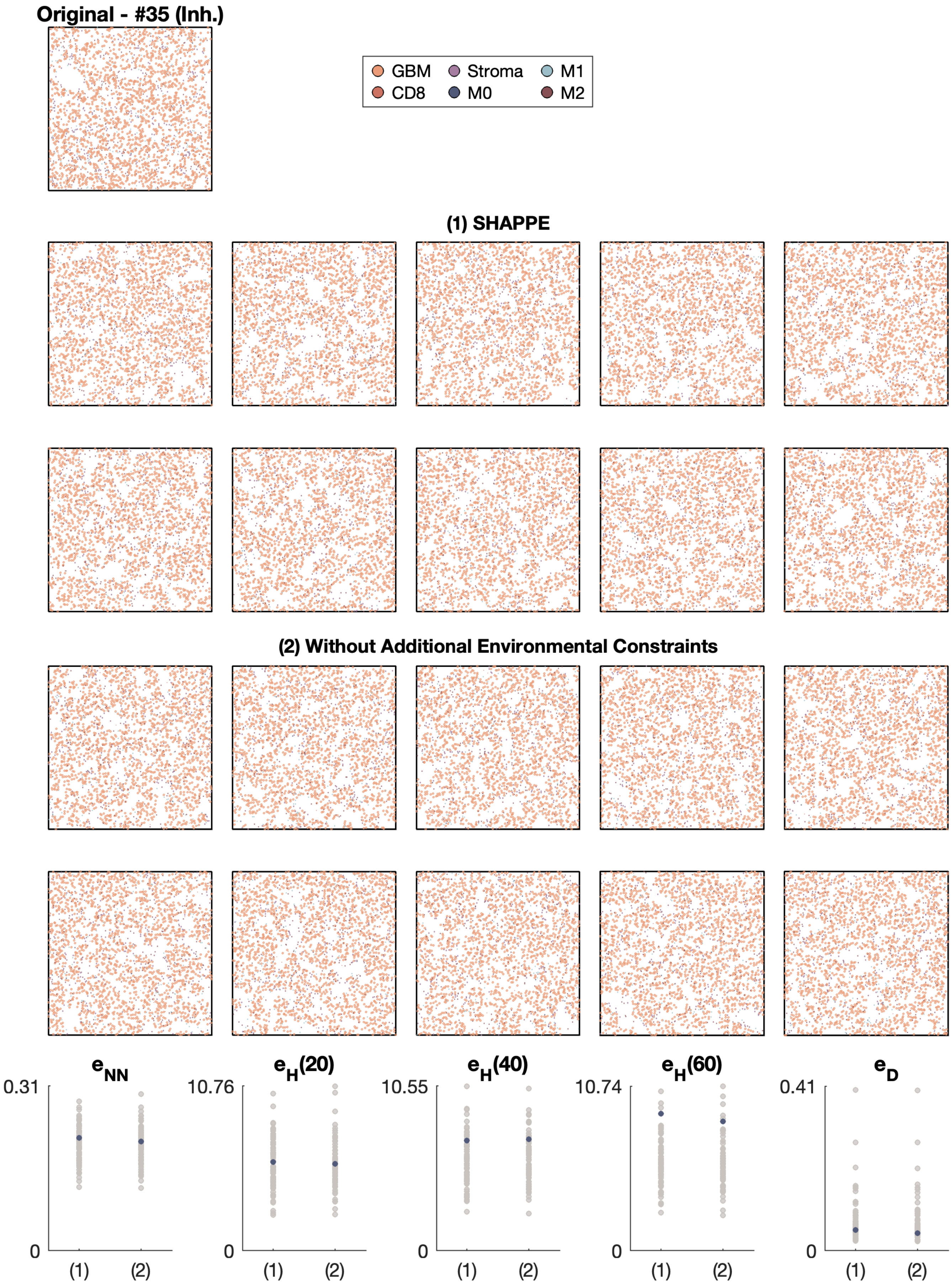
Original - #35 (Inh.)
GBM
Stroma
M1
CD8
M0
M2
(1) SHAPPE
(2) Without Additional Environmental Constraints
$e_{NN}$
0.31
0
(1)
(2)
$e_H(20)$
10.76
0
(1)
(2)
$e_H(40)$
10.55
0
(1)
(2)
$e_H(60)$
10.74
0
(1)
(2)
$e_D$
0.41
0
(1)
(2)

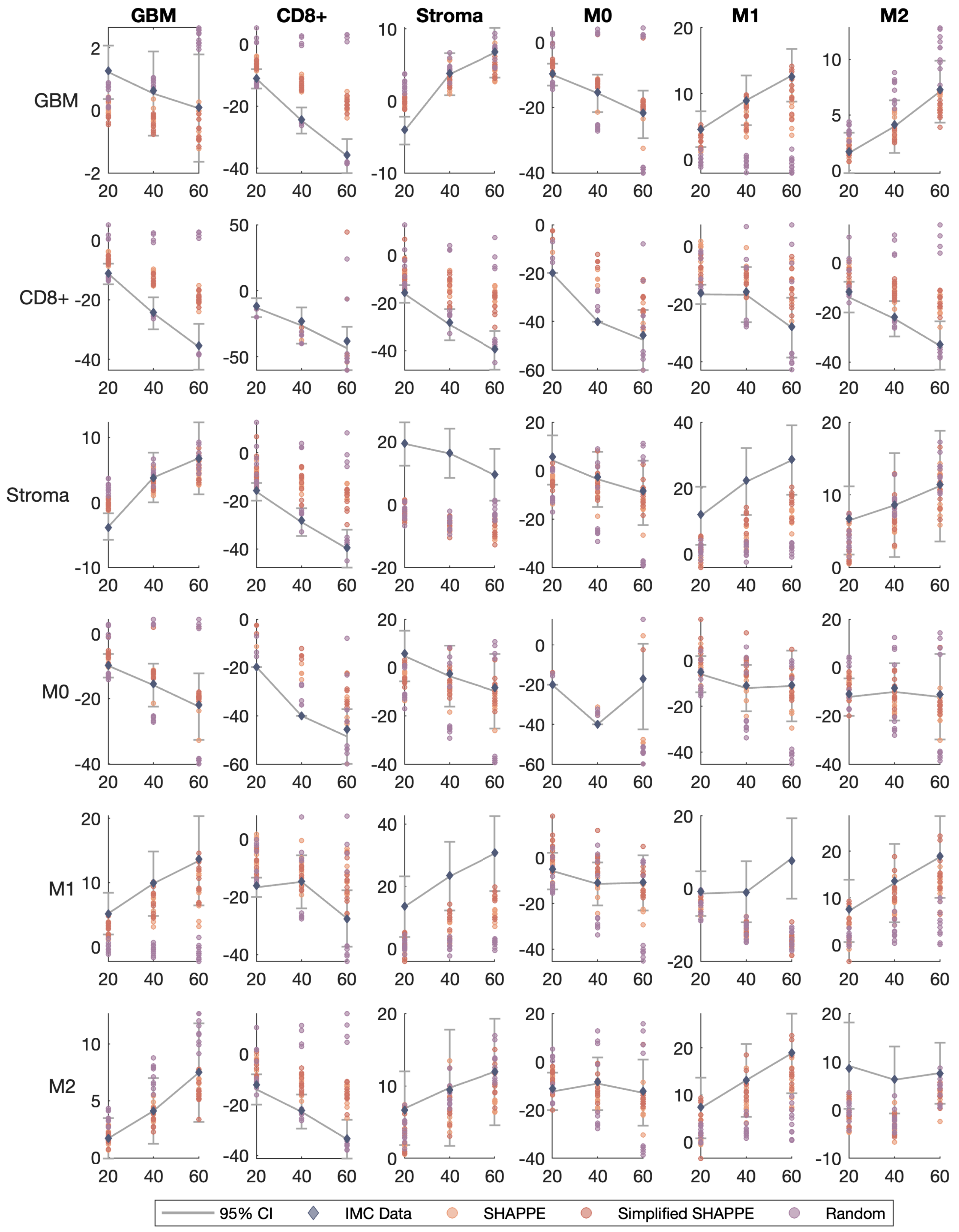
GBM
CD8+
Stroma
M0
M1
M2
GBM
CD8+
Stroma
M0
M1
M2
95% CI
IMC Data
SHAPPE
Simplified SHAPPE
Random

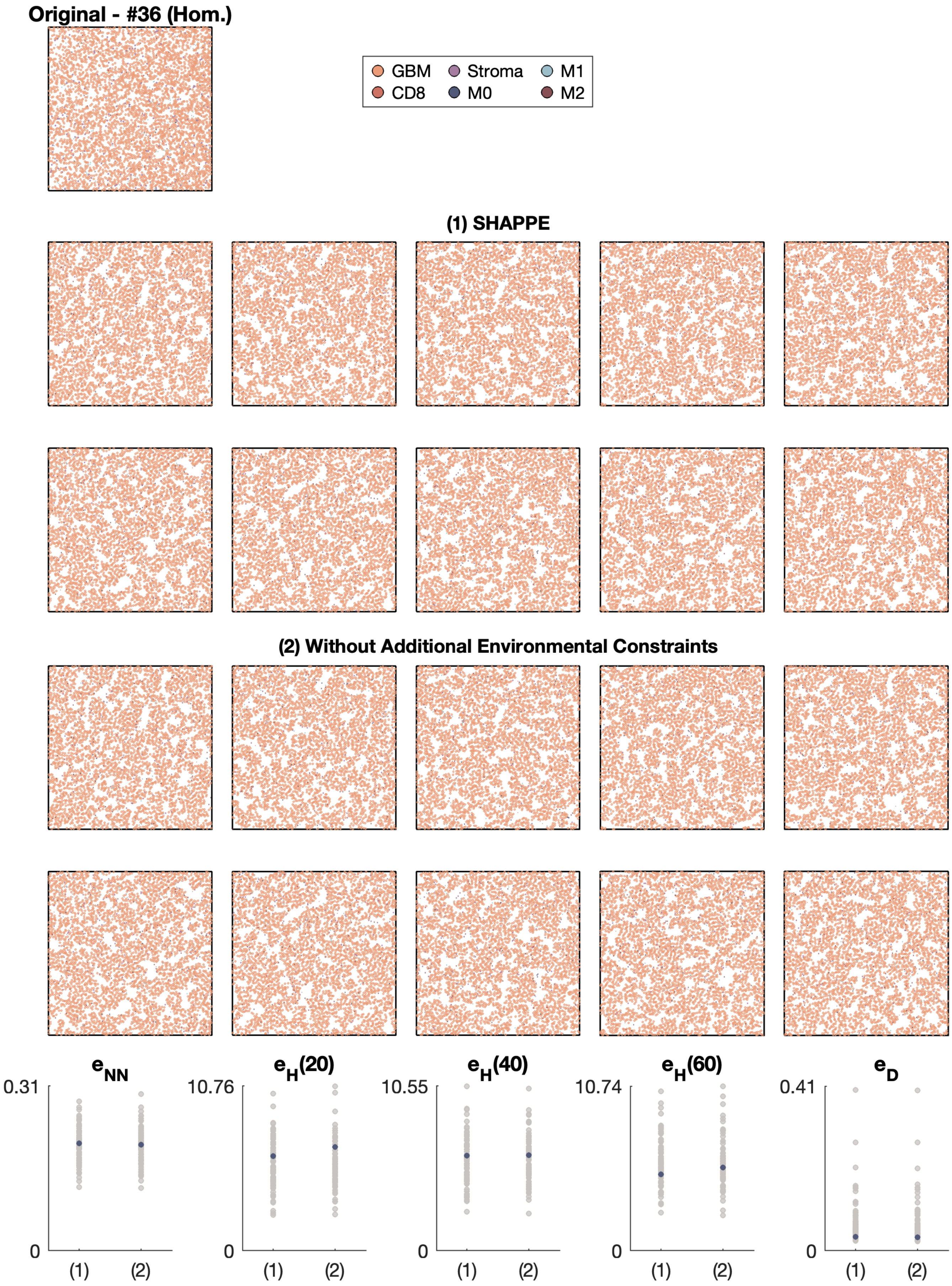
Original - #36 (Hom.)
GBM
CD8
Stroma
M0
M1
M2
(1) SHAPPE
(2) Without Additional Environmental Constraints
$e_{NN}$
0.31
0
$e_H(20)$
10.76
$e_H(40)$
10.55
$e_H(60)$
10.74
$e_D$
0.41
(1)
(2)

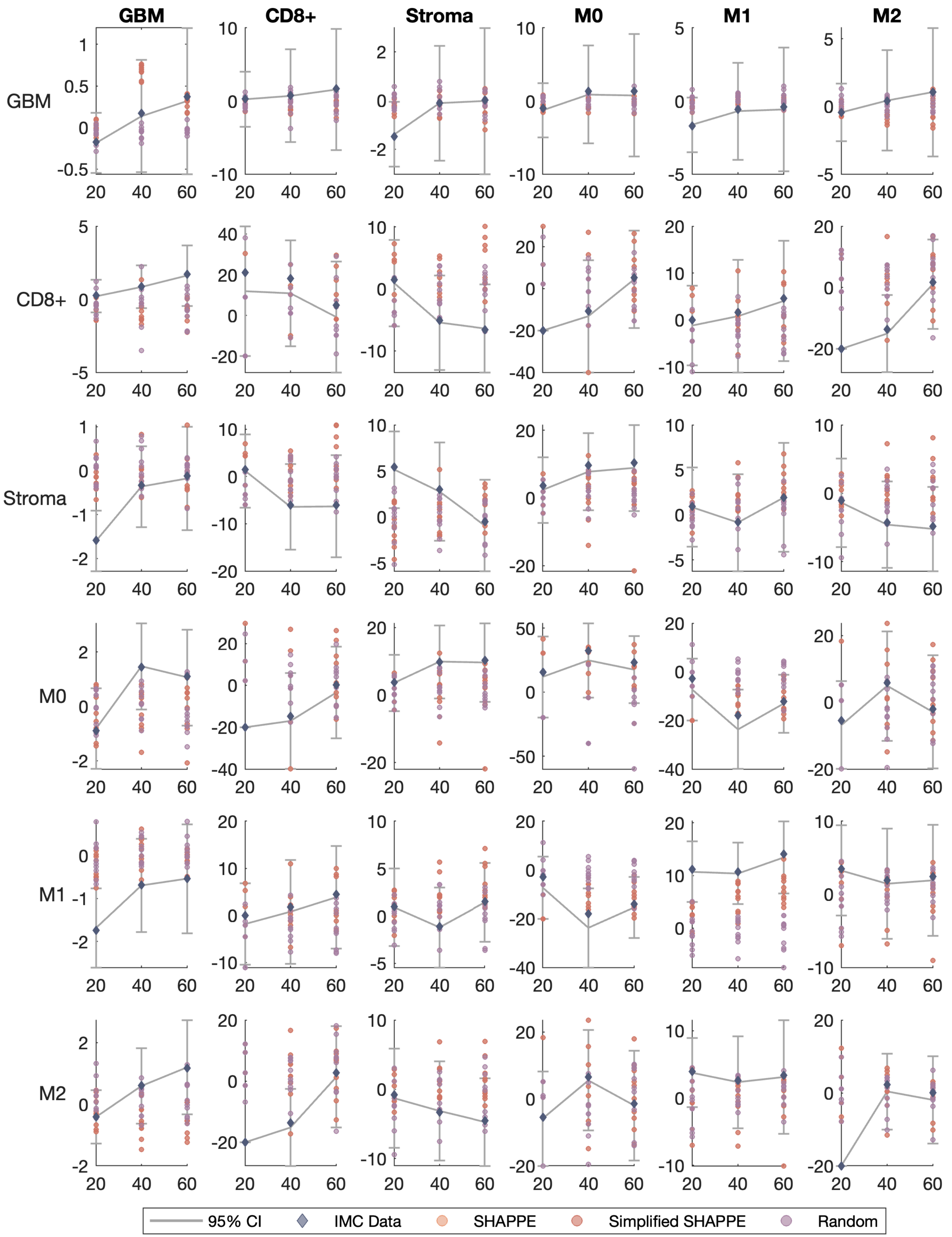
GBM
CD8+
Stroma
M0
M1
M2
GBM
CD8+
Stroma
M0
M1
M2
95% CI
IMC Data
SHAPPE
Simplified SHAPPE
Random

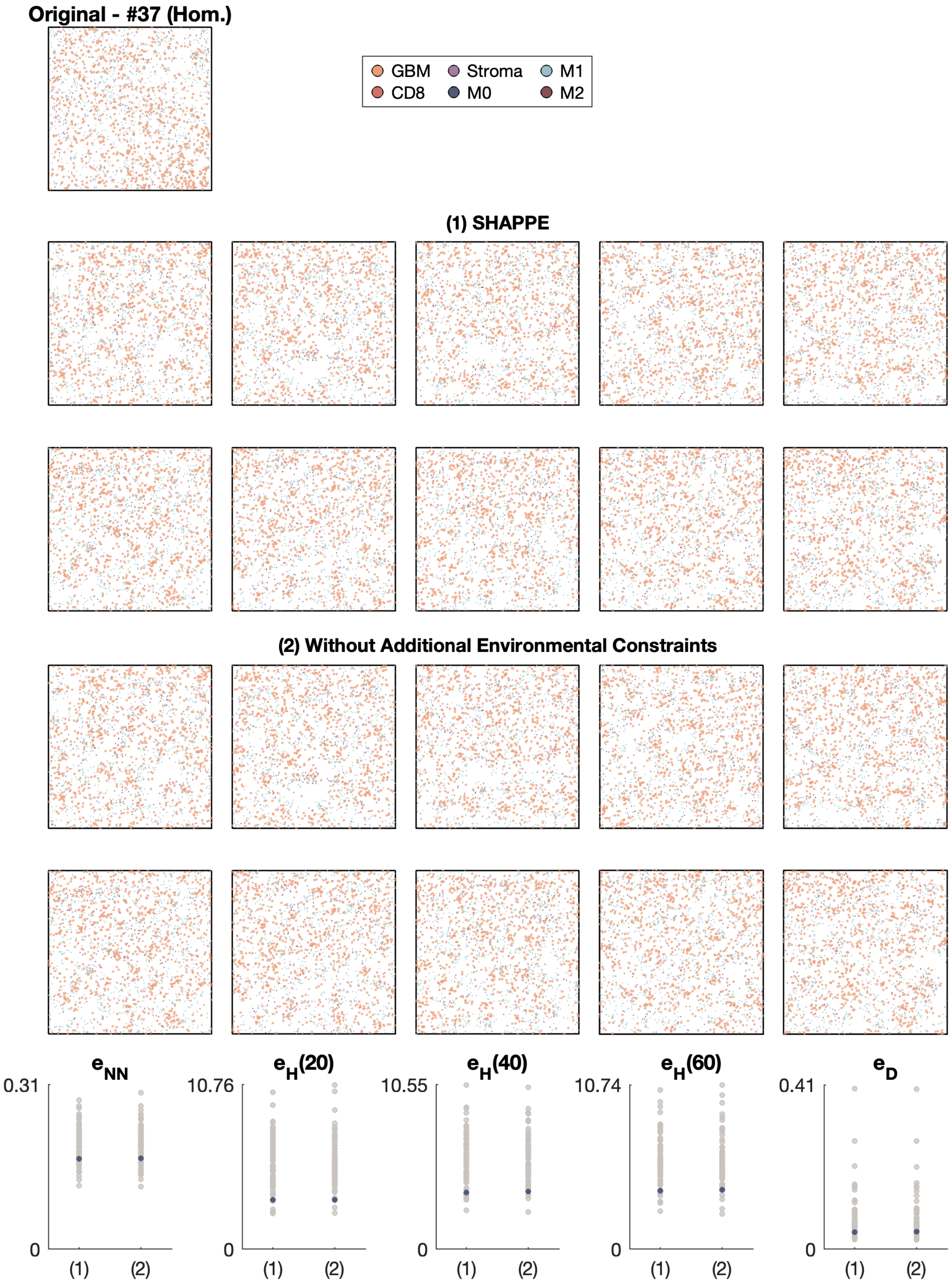
Original - #37 (Hom.)
GBM
CD8
Stroma
M0
M1
M2
(1) SHAPPE
(2) Without Additional Environmental Constraints
$e_{NN}$
$e_H(20)$
$e_H(40)$
$e_H(60)$
$e_D$
0.31
10.76
10.55
10.74
0.41
0
(1)
(2)

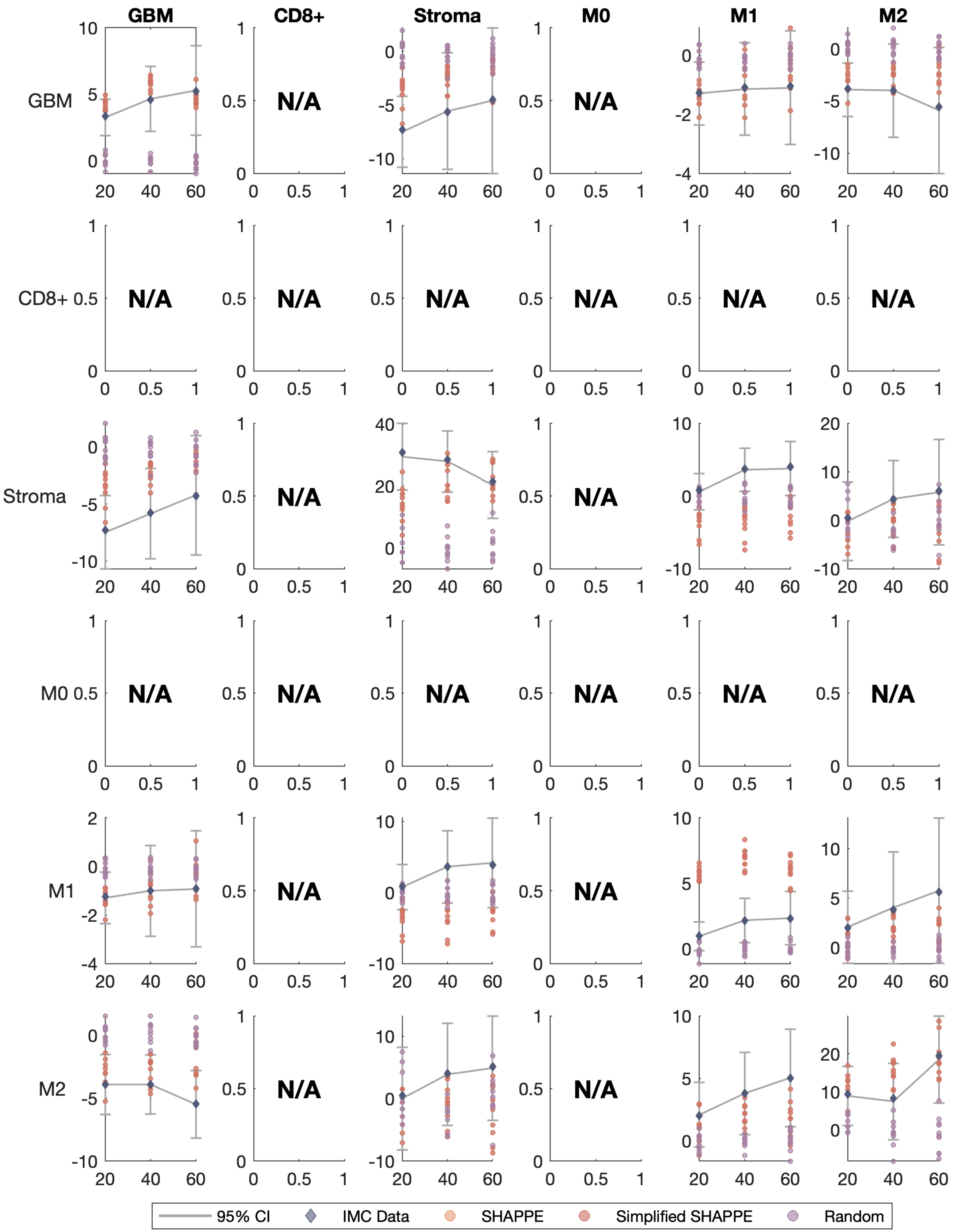

GBM
CD8+
Stroma
M0
M1
M2
N/A
95% CI
IMC Data
SHAPPE
Simplified SHAPPE
Random

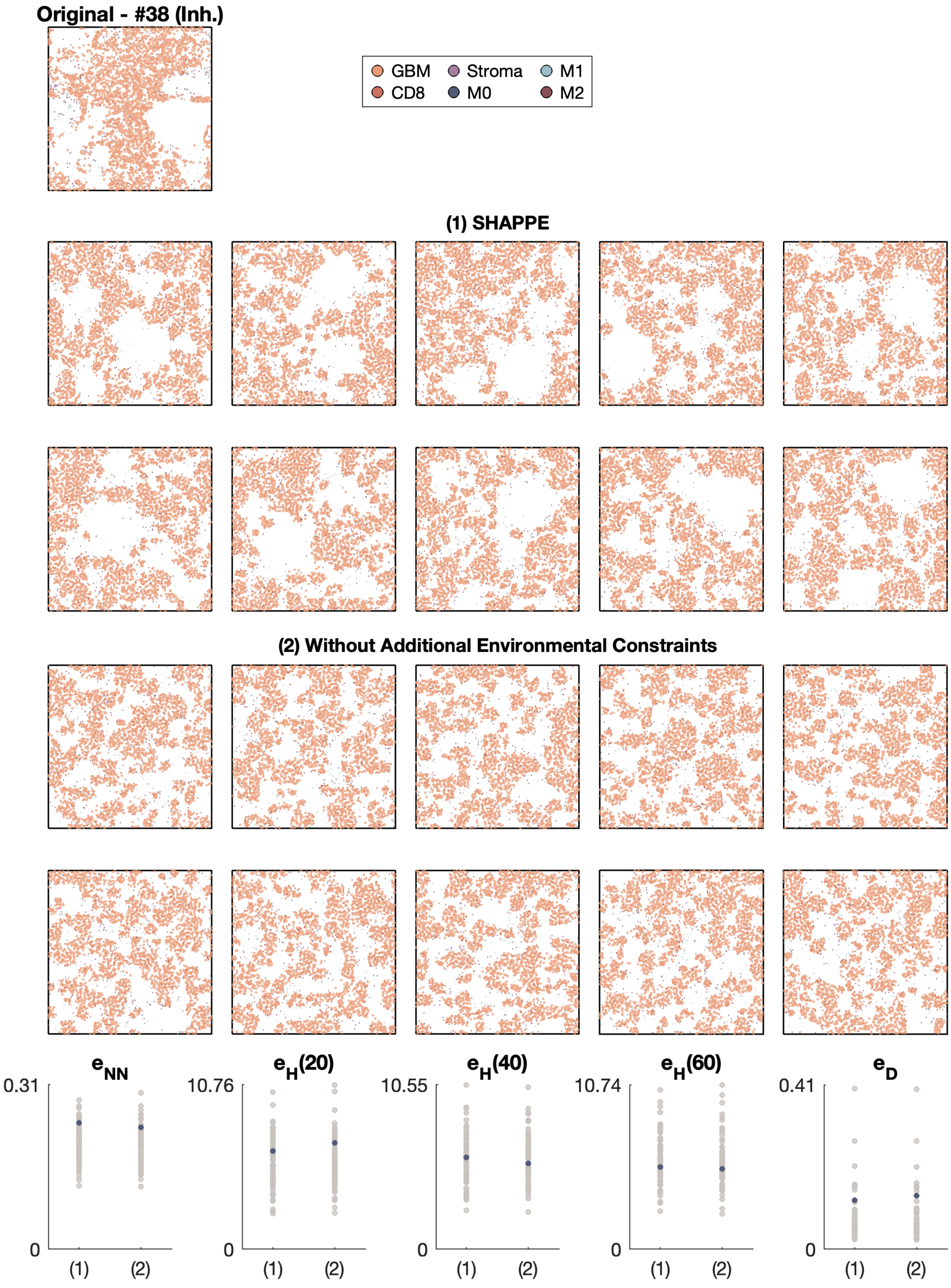
Original - #38 (Inh.)
GBM
Stroma
M1
CD8
M0
M2
(1) SHAPPE
(2) Without Additional Environmental Constraints
e_NN
0.31
0
(1)
(2)
e_H(20)
10.76
0
(1)
(2)
e_H(40)
10.55
0
(1)
(2)
e_H(60)
10.74
0
(1)
(2)
e_D
0.41
0
(1)
(2)

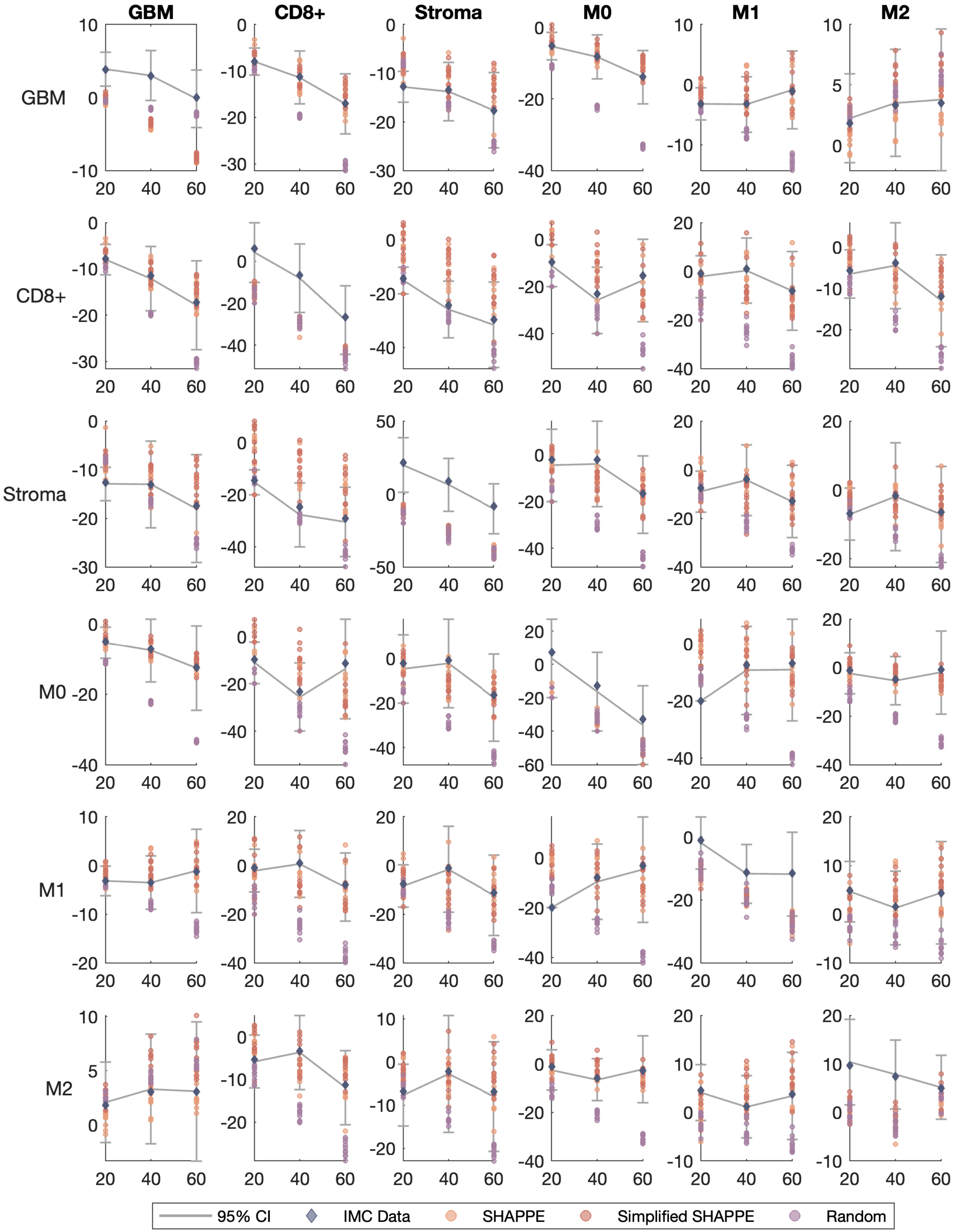
GBM
CD8+
Stroma
M0
M1
M2
95% CI
IMC Data
SHAPPE
Simplified SHAPPE
Random

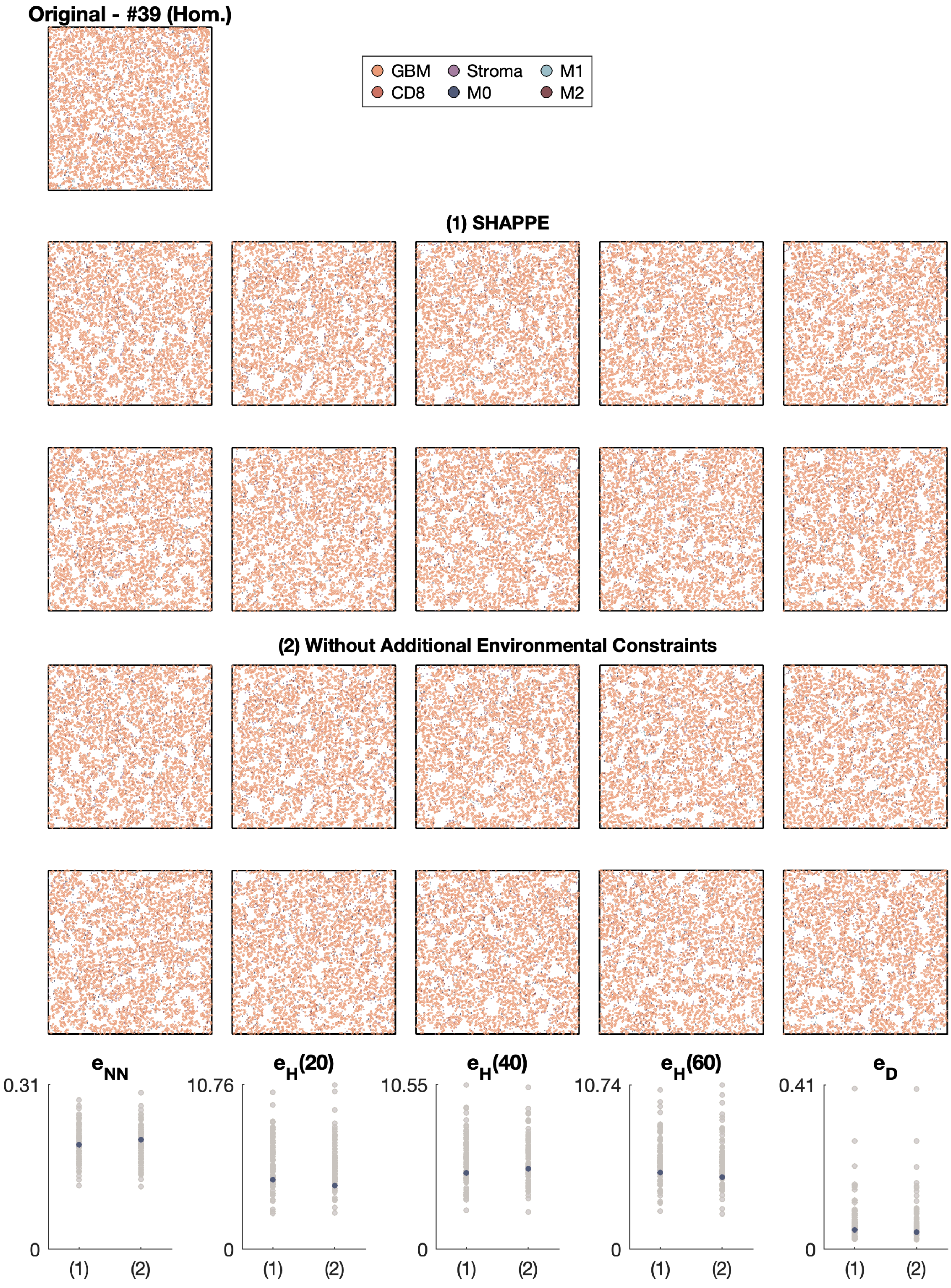
Original - #39 (Hom.)
GBM
CD8
Stroma
M0
M1
M2
(1) SHAPPE
(2) Without Additional Environmental Constraints
$e_{NN}$
0.31
0
(1)
(2)
$e_H(20)$
10.76
0
(1)
(2)
$e_H(40)$
10.55
0
(1)
(2)
$e_H(60)$
10.74
0
(1)
(2)
$e_D$
0.41
0
(1)
(2)

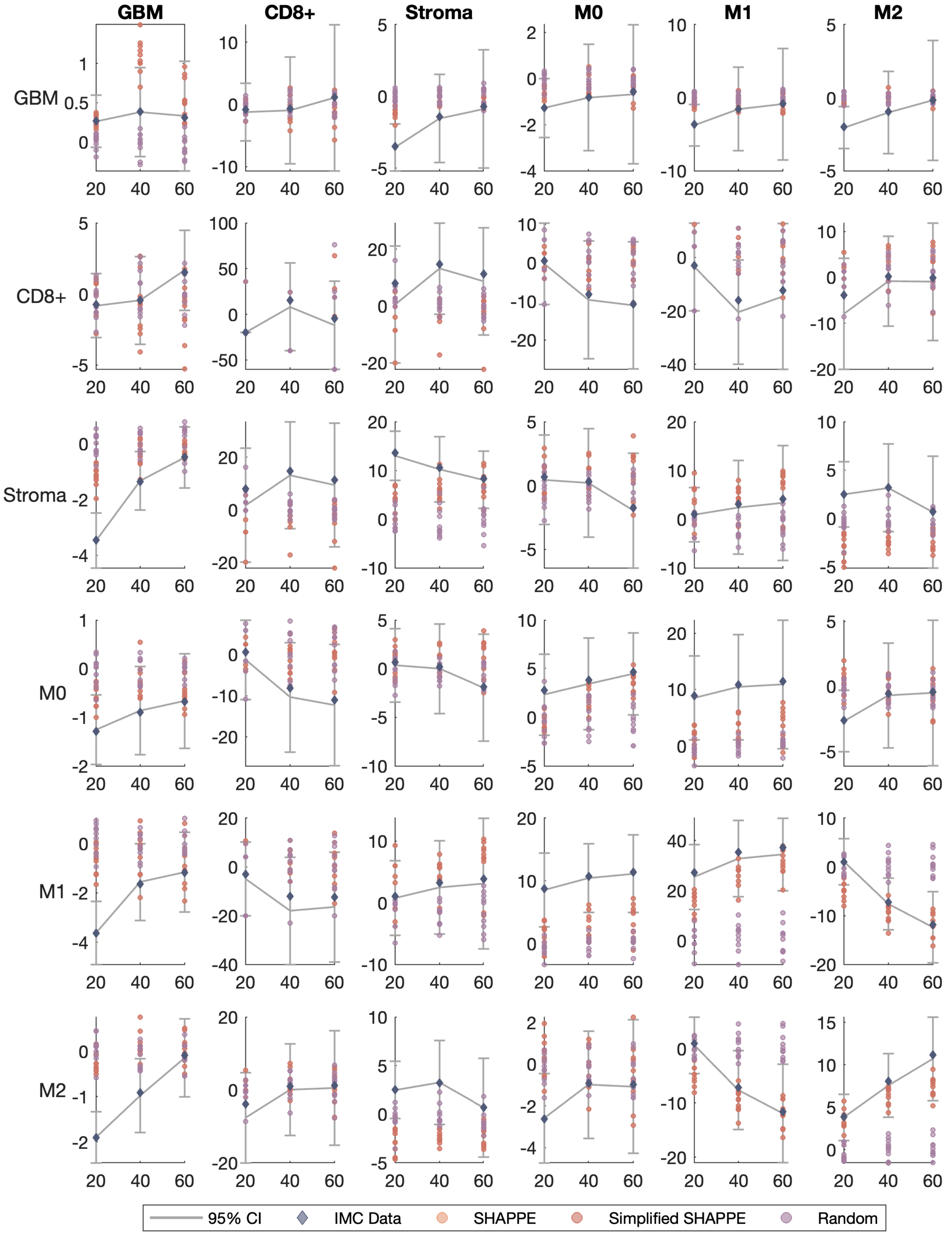

GBM
CD8+
Stroma
M0
M1
M2
95% CI
IMC Data
SHAPPE
Simplified SHAPPE
Random

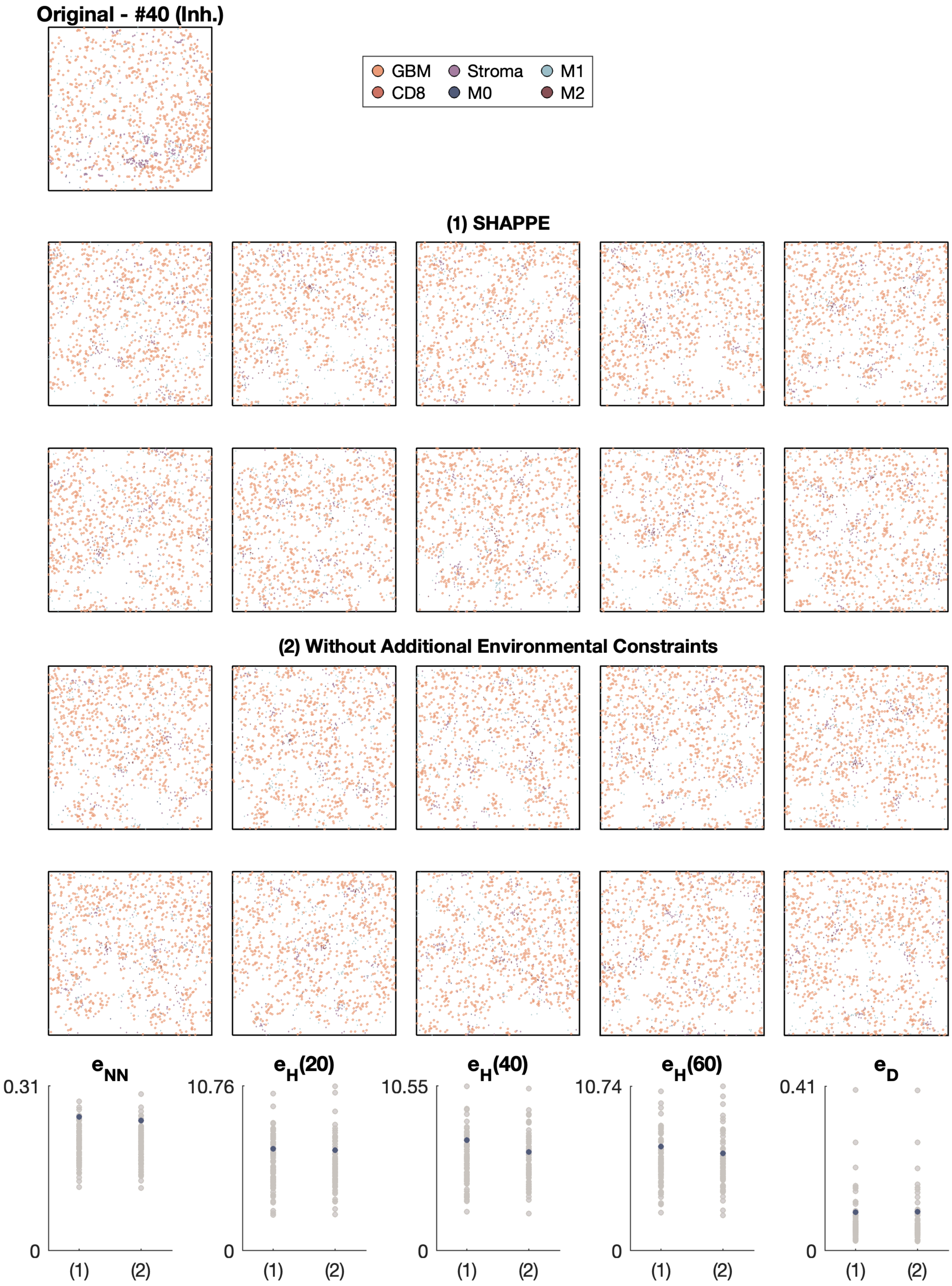

Original - #40 (Inh.)
GBM
CD8
Stroma
M0
M1
M2
(1) SHAPPE
(2) Without Additional Environmental Constraints
$e_{NN}$
$e_H(20)$
$e_H(40)$
$e_H(60)$
$e_D$
0.31
10.76
10.55
10.74
0.41
0
(1)
(2)

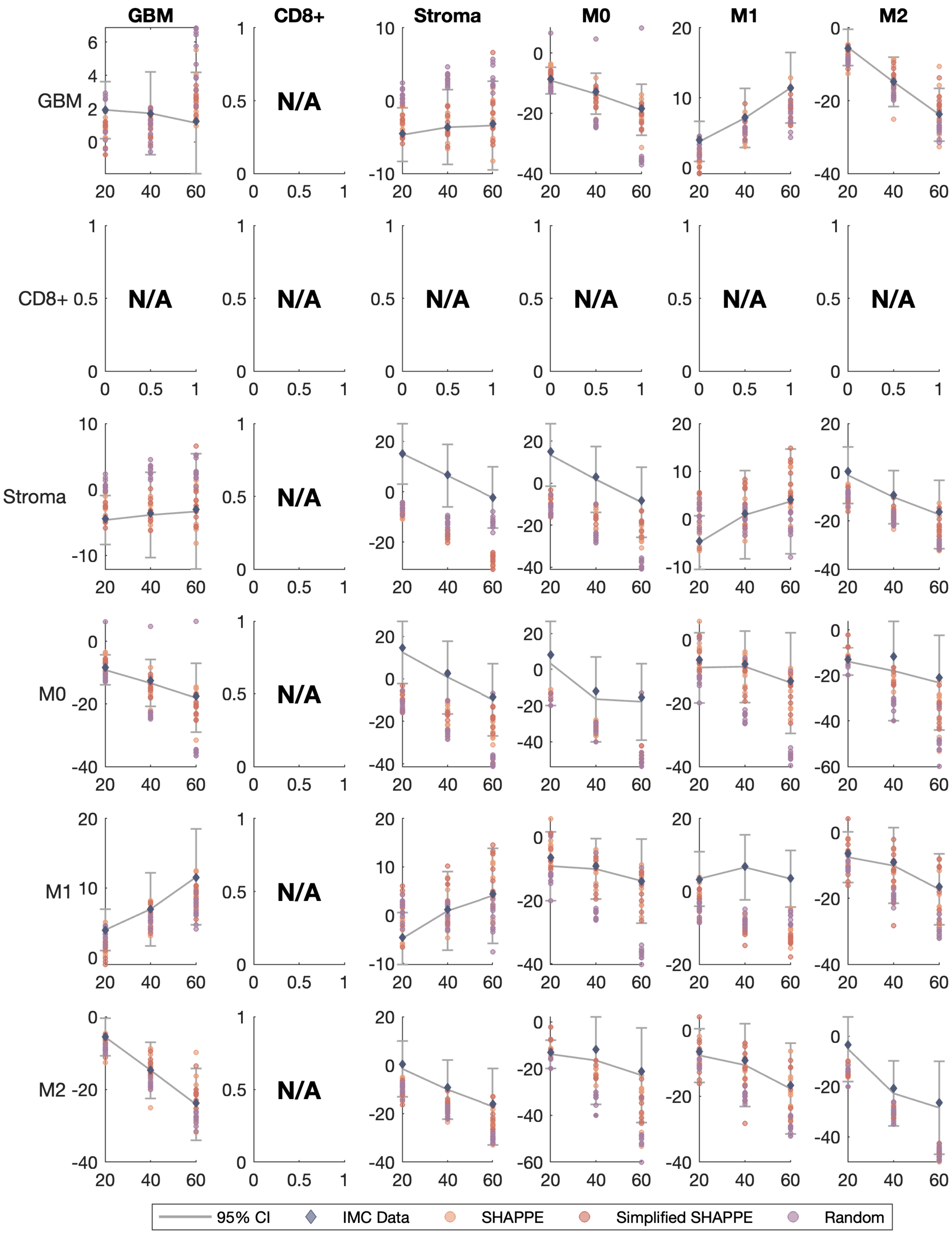
GBM
CD8+
Stroma
M0
M1
M2
N/A
95% CI
IMC Data
SHAPPE
Simplified SHAPPE
Random

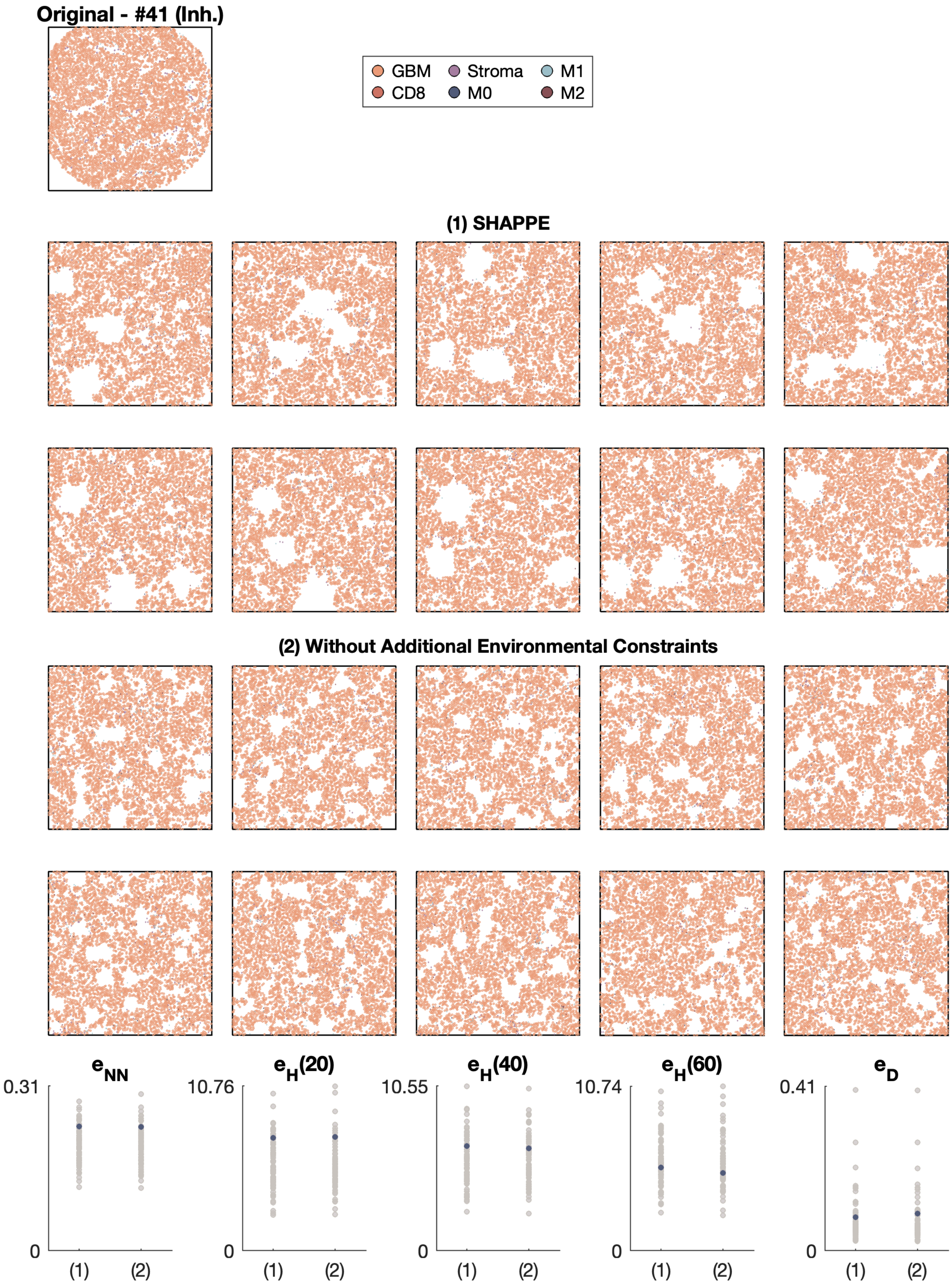

Original - #41 (Inh.)
GBM
CD8
Stroma
M0
M1
M2
(1) SHAPPE
(2) Without Additional Environmental Constraints
e_NN
e_H(20)
e_H(40)
e_H(60)
e_D
0.31
10.76
10.55
10.74
0.41
0
(1)
(2)

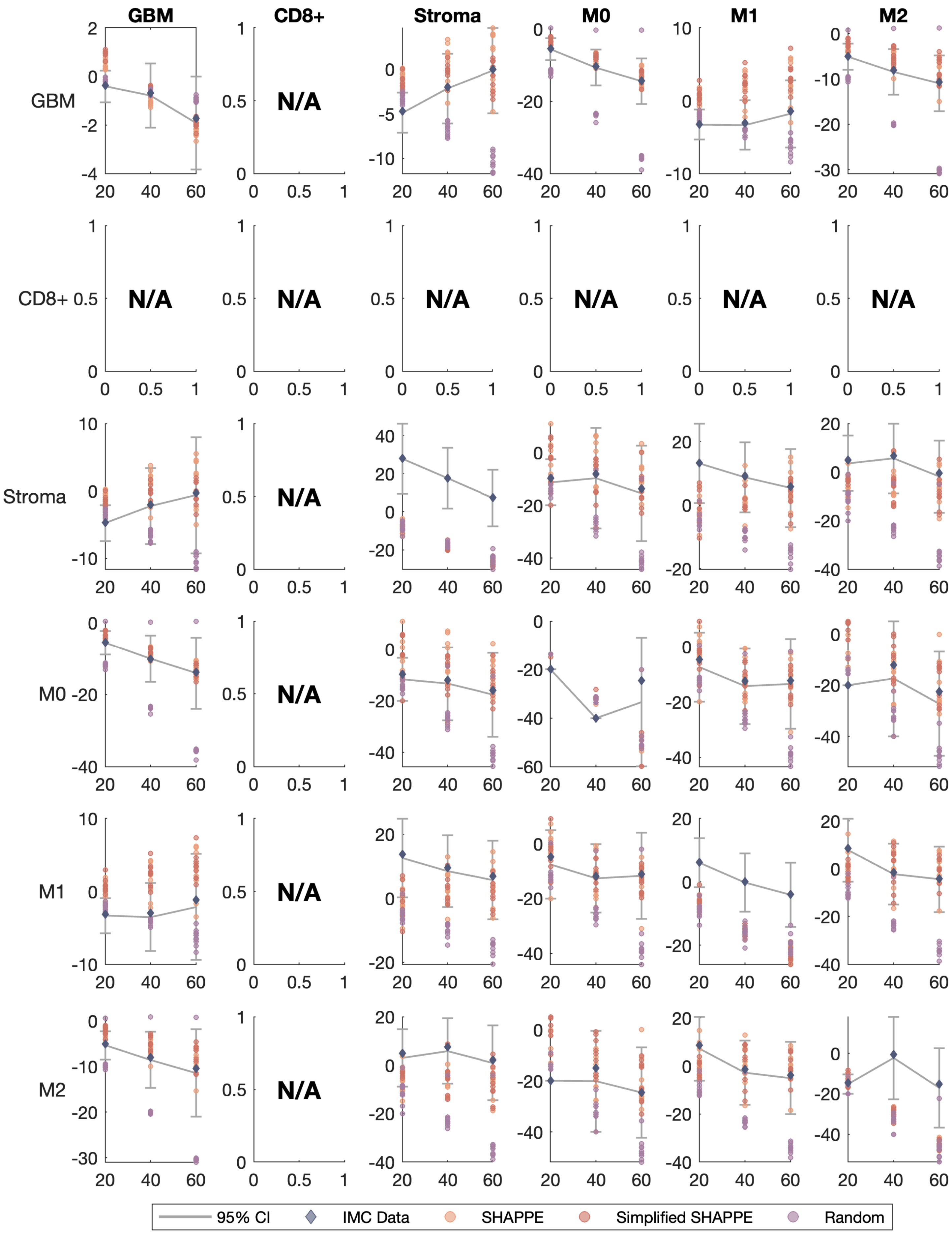

GBM
CD8+
Stroma
M0
M1
M2
GBM
CD8+
Stroma
M0
M1
M2
N/A
95% CI
IMC Data
SHAPPE
Simplified SHAPPE
Random

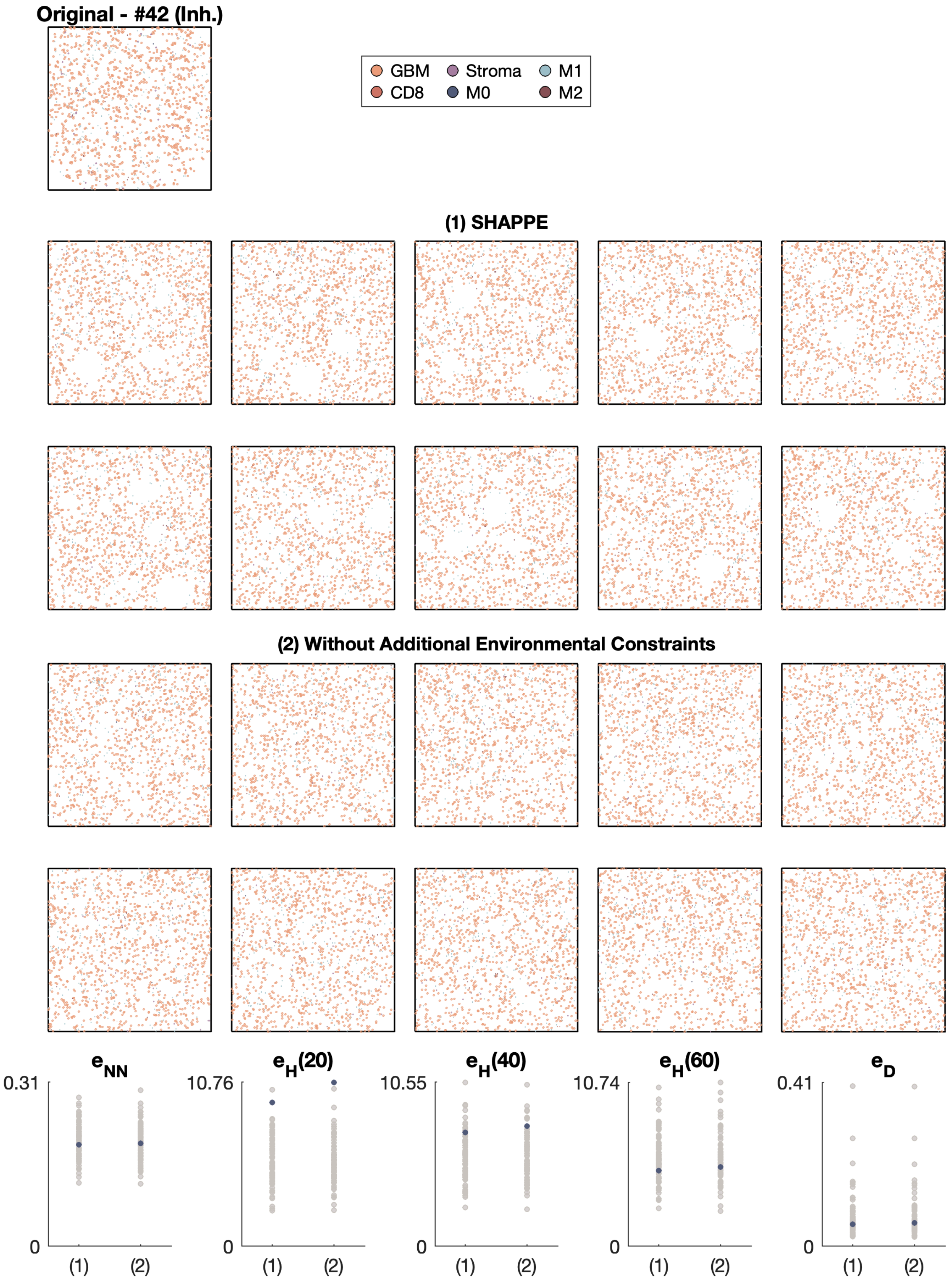
Original - #42 (Inh.)
GBM
Stroma
M1
CD8
M0
M2
(1) SHAPPE
(2) Without Additional Environmental Constraints
$e_{NN}$
$e_H(20)$
$e_H(40)$
$e_H(60)$
$e_D$
0.31
10.76
10.55
10.74
0.41
0
(1)
(2)

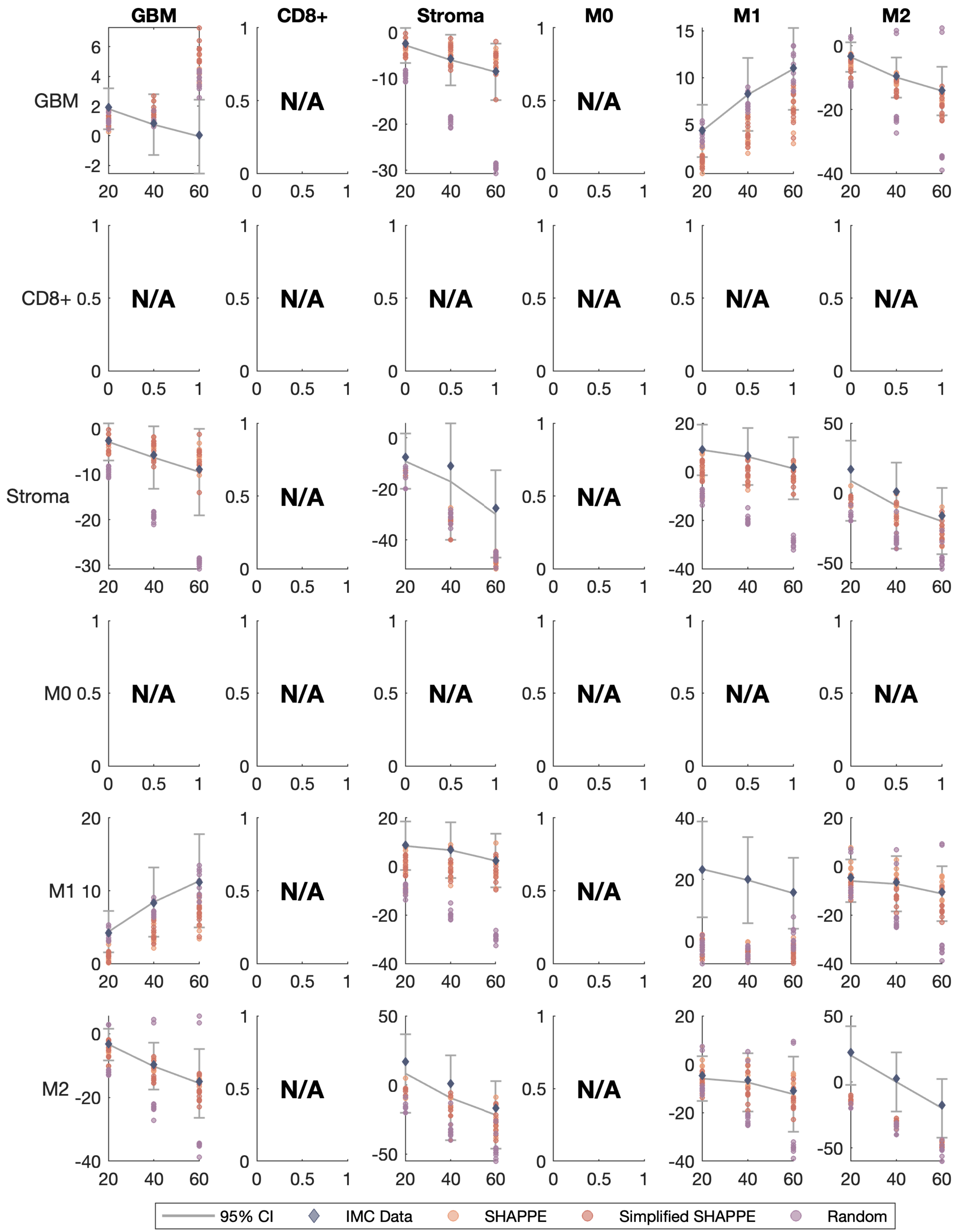

GBM
CD8+
Stroma
M0
M1
M2
N/A
95% CI
IMC Data
SHAPPE
Simplified SHAPPE
Random

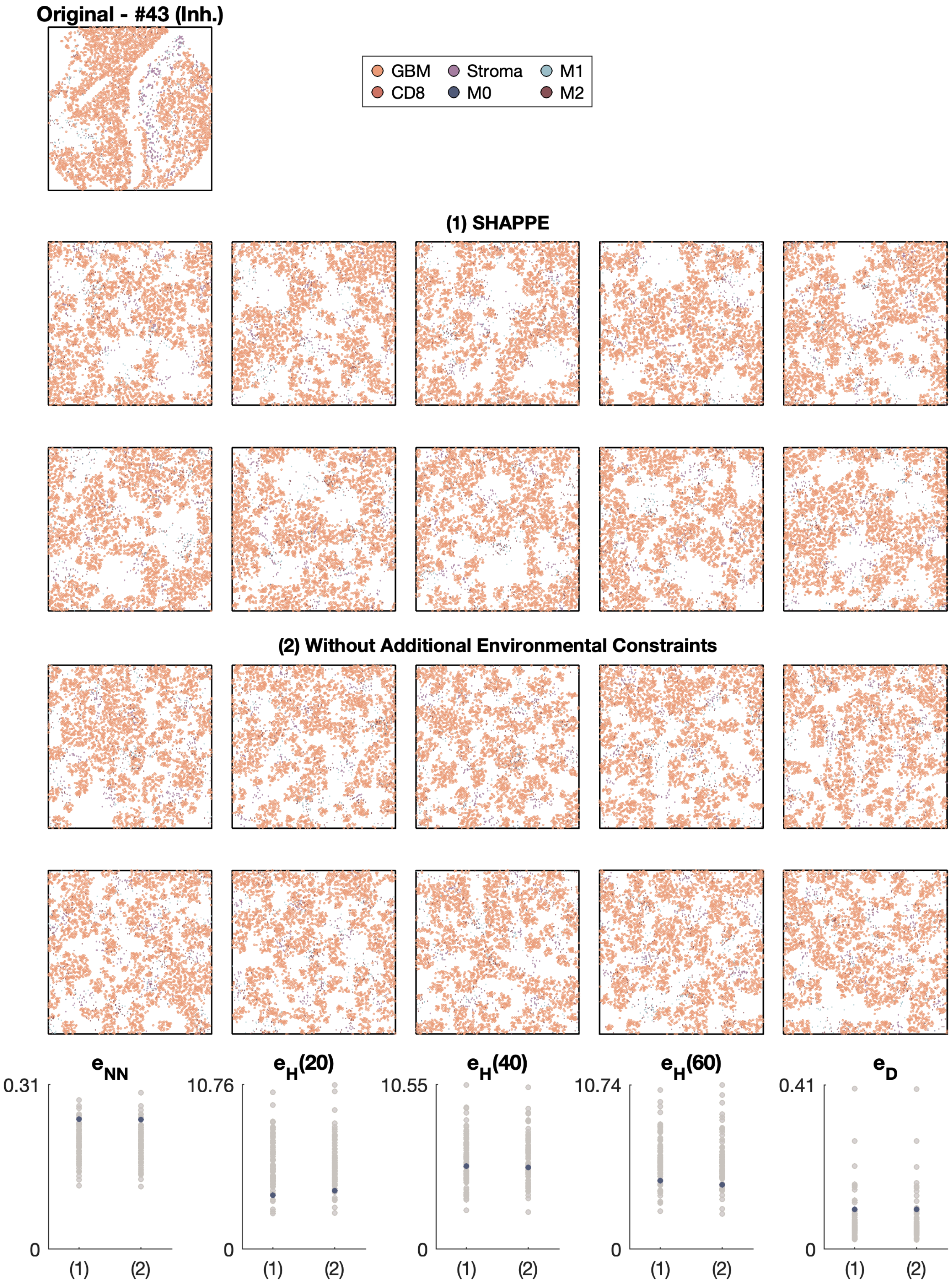
Original - #43 (Inh.)
GBM
Stroma
M1
CD8
M0
M2
(1) SHAPPE
(2) Without Additional Environmental Constraints
$e_{NN}$
0.31
0
(1)
(2)
$e_H(20)$
10.76
0
(1)
(2)
$e_H(40)$
10.55
0
(1)
(2)
$e_H(60)$
10.74
0
(1)
(2)
$e_D$
0.41
0
(1)
(2)

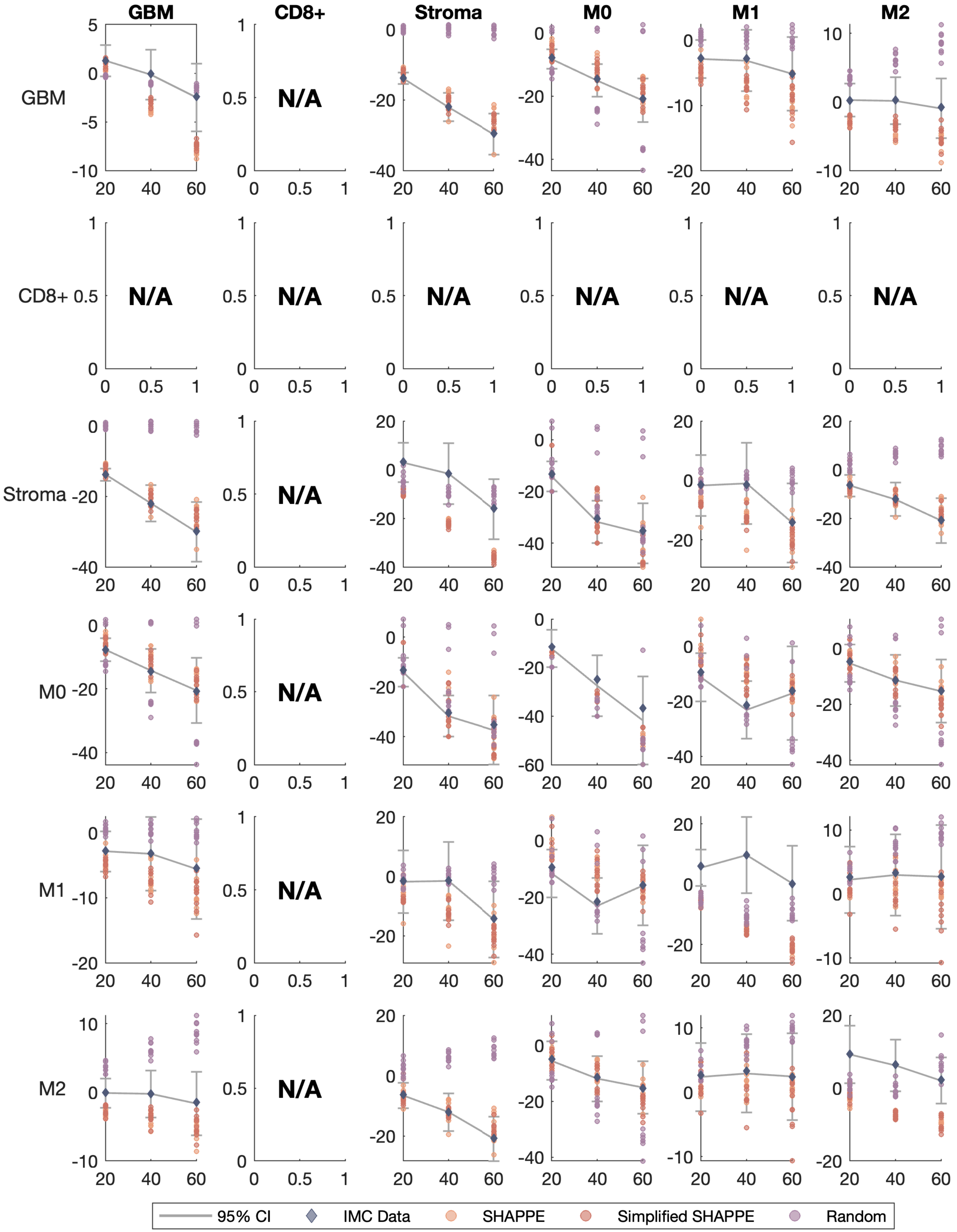
GBM
CD8+
Stroma
M0
M1
M2
GBM
CD8+
Stroma
M0
M1
M2
N/A
95% CI
IMC Data
SHAPPE
Simplified SHAPPE
Random

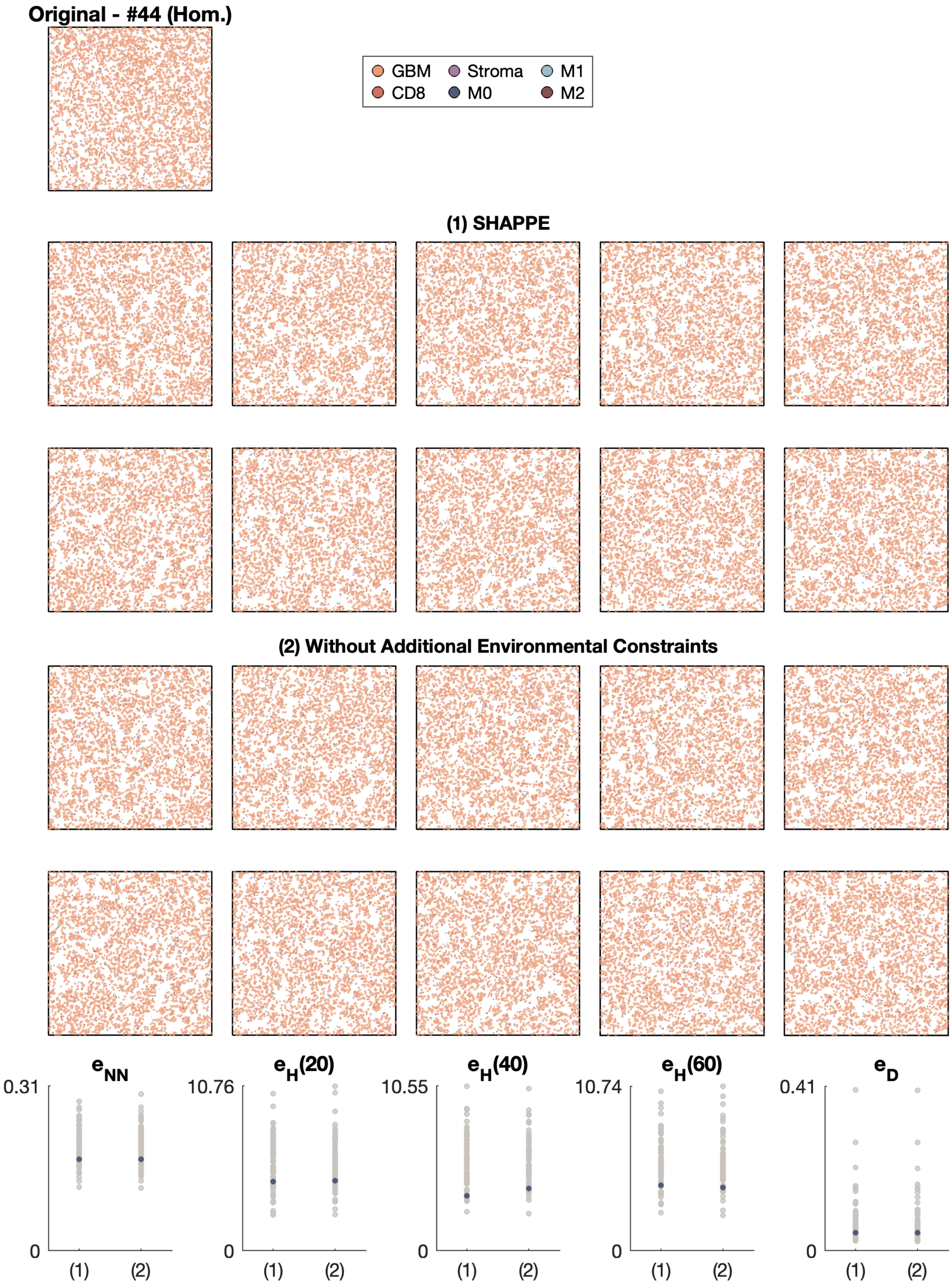

Original - #44 (Hom.)
GBM
Stroma
M1
CD8
M0
M2
(1) SHAPPE
(2) Without Additional Environmental Constraints
$e_{NN}$
0.31
0
(1)
(2)
$e_H(20)$
10.76
0
(1)
(2)
$e_H(40)$
10.55
0
(1)
(2)
$e_H(60)$
10.74
0
(1)
(2)
$e_D$
0.41
0
(1)
(2)

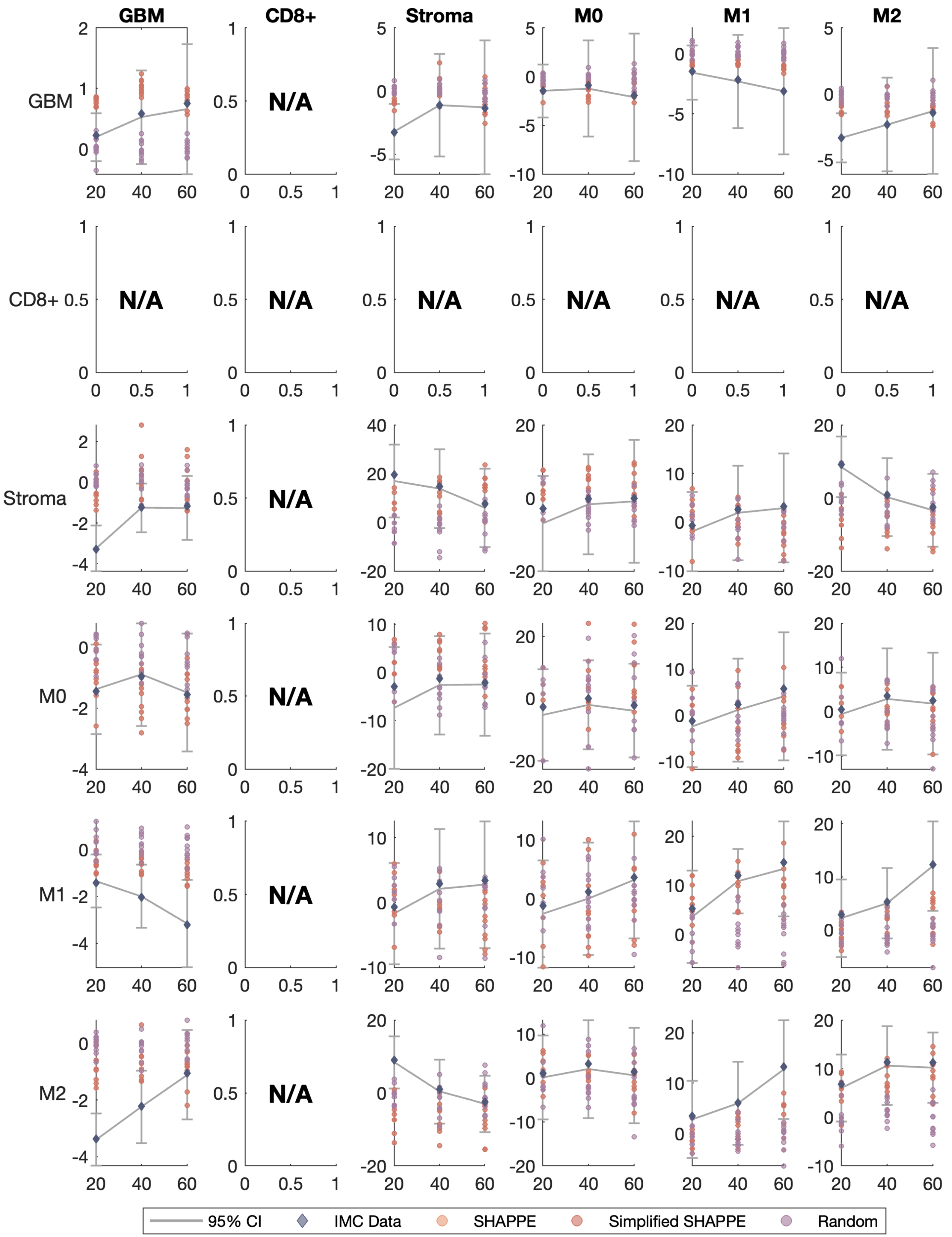

GBM
CD8+
Stroma
M0
M1
M2
GBM
CD8+
Stroma
M0
M1
M2
N/A
95% CI
IMC Data
SHAPPE
Simplified SHAPPE
Random

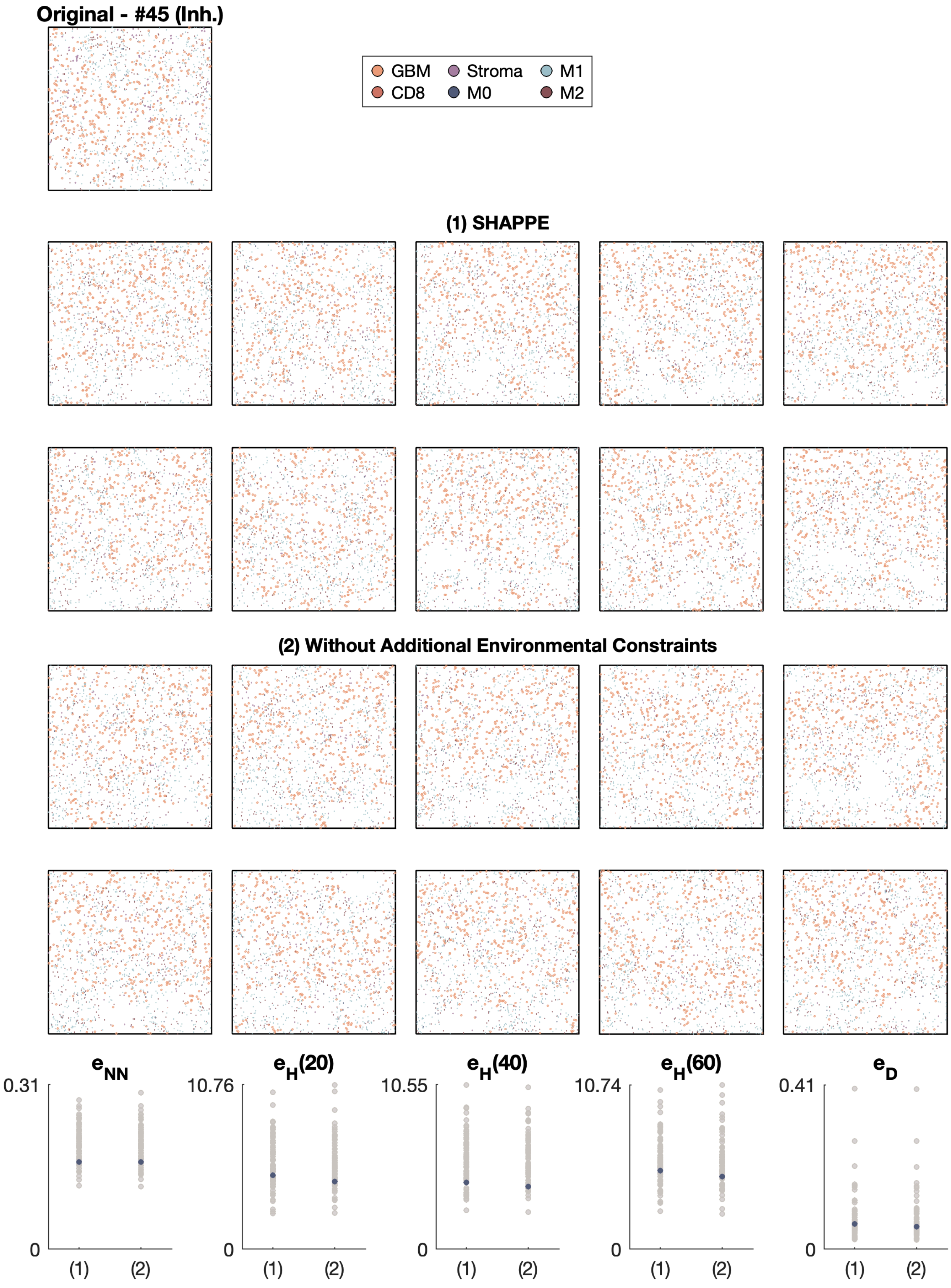
Original - #45 (Inh.)
GBM
CD8
Stroma
M0
M1
M2
(1) SHAPPE
(2) Without Additional Environmental Constraints
$e_{NN}$
$e_H(20)$
$e_H(40)$
$e_H(60)$
$e_D$
0.31
10.76
10.55
10.74
0.41
0
(1)
(2)

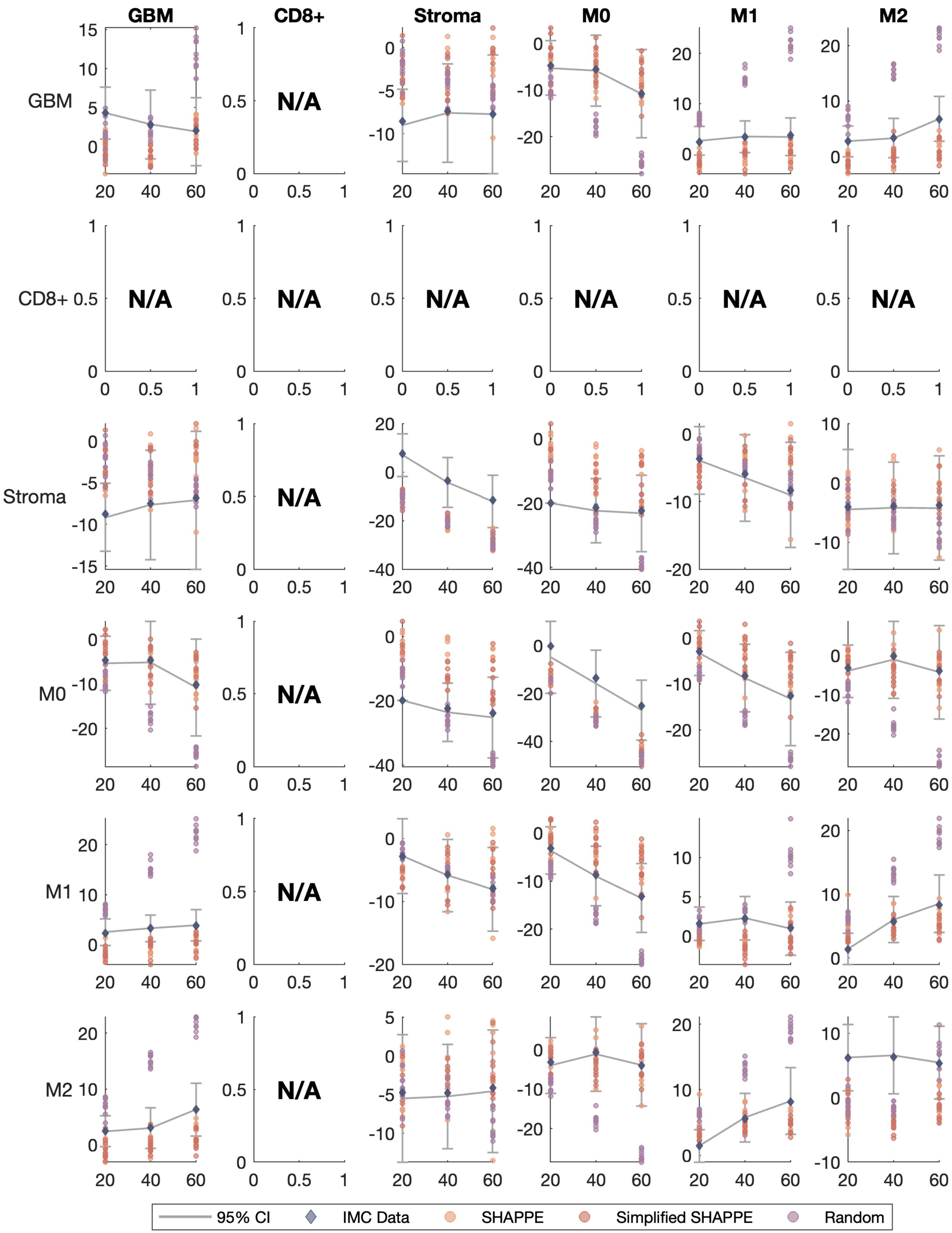

GBM
CD8+
Stroma
M0
M1
M2
N/A
95% CI
IMC Data
SHAPPE
Simplified SHAPPE
Random

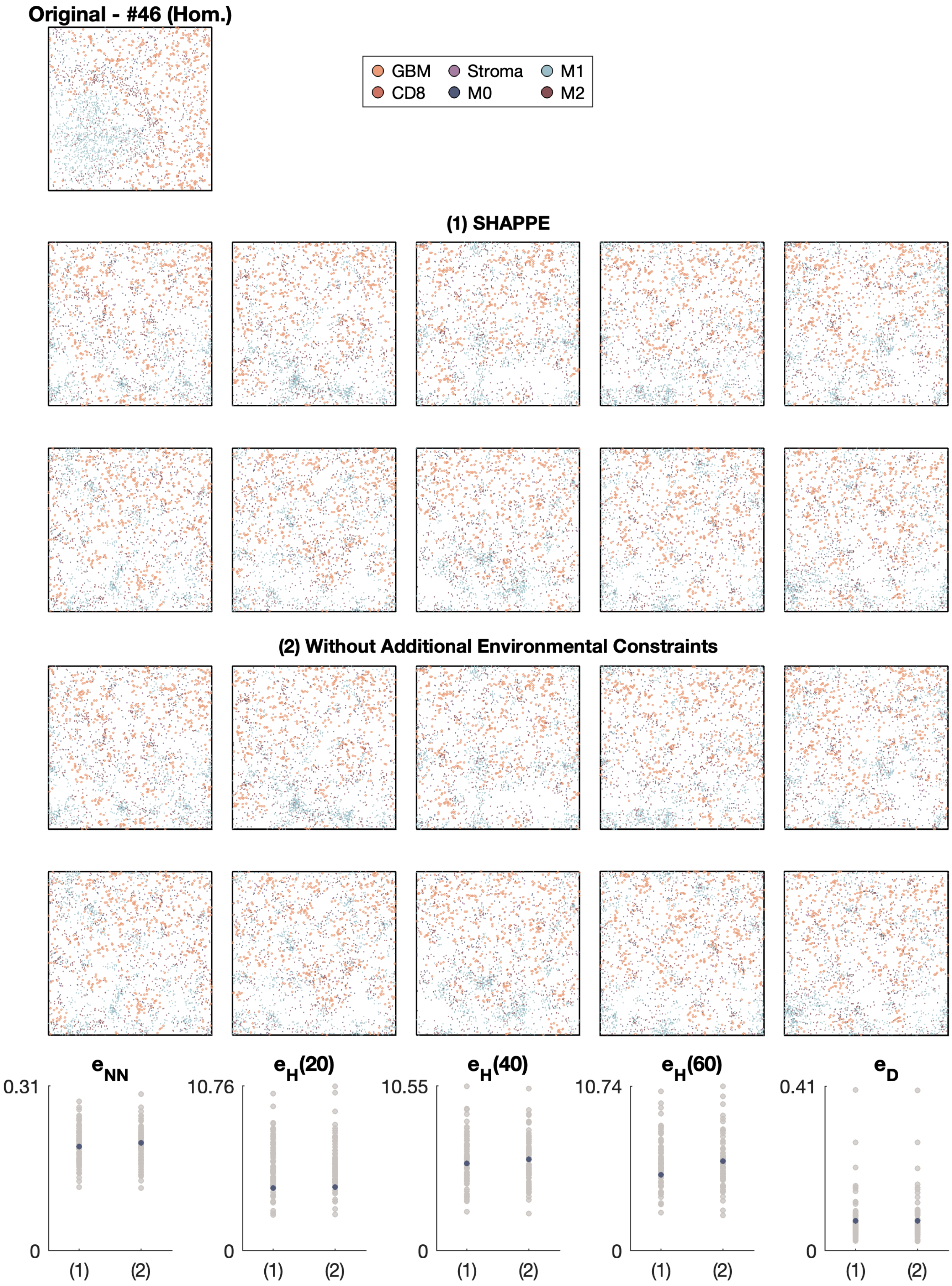
Original - #46 (Hom.)
GBM
CD8
Stroma
M0
M1
M2
(1) SHAPPE
(2) Without Additional Environmental Constraints
$e_{NN}$
0.31
0
(1)
(2)
$e_H(20)$
10.76
0
(1)
(2)
$e_H(40)$
10.55
0
(1)
(2)
$e_H(60)$
10.74
0
(1)
(2)
$e_D$
0.41
0
(1)
(2)

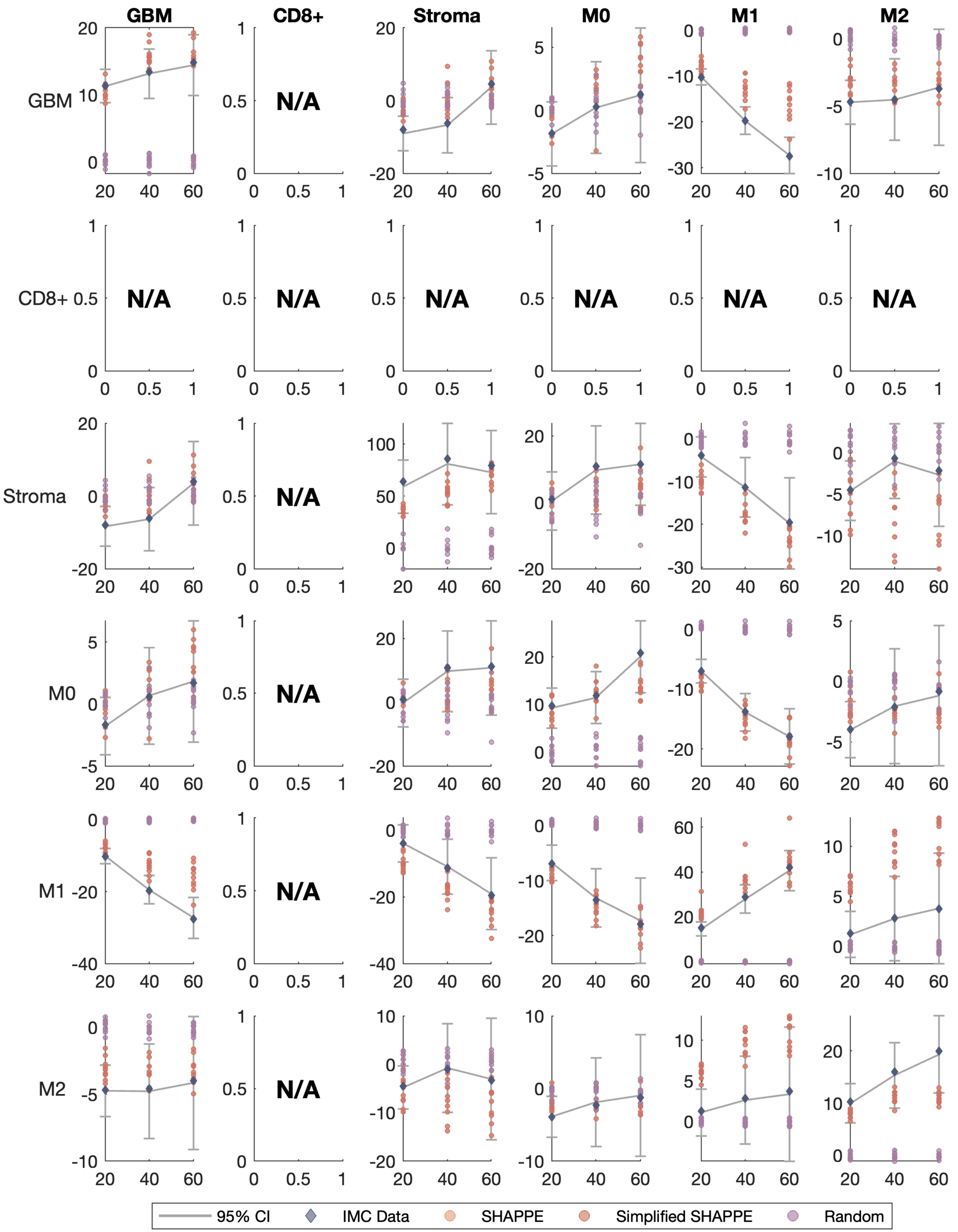
GBM
CD8+
Stroma
M0
M1
M2
N/A
95% CI
IMC Data
SHAPPE
Simplified SHAPPE
Random

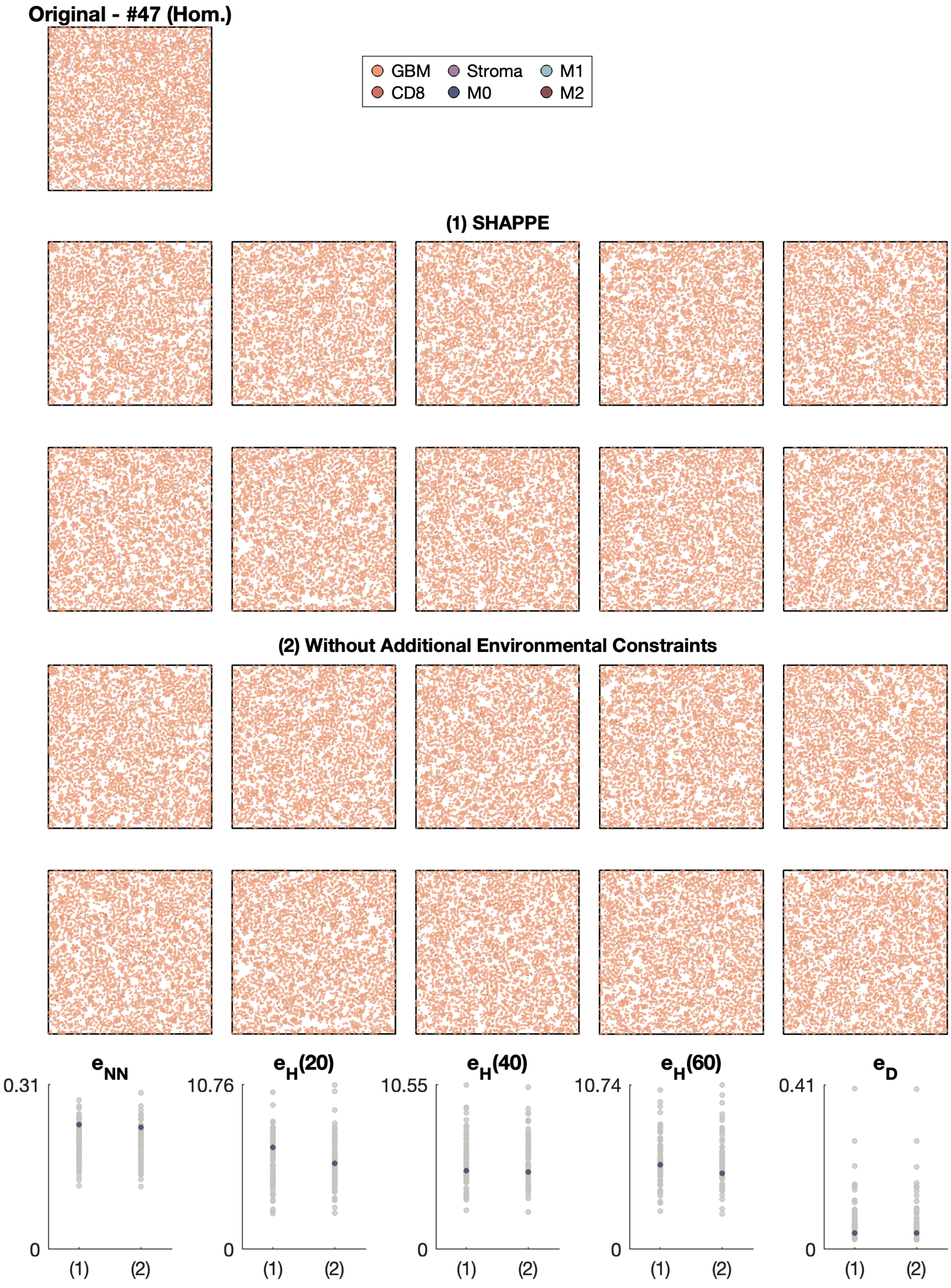

Original - #47 (Hom.)
GBM
CD8
Stroma
M0
M1
M2
(1) SHAPPE
(2) Without Additional Environmental Constraints
$e_{NN}$
$e_H(20)$
$e_H(40)$
$e_H(60)$
$e_D$
0.31
10.76
10.55
10.74
0.41
0
(1)
(2)

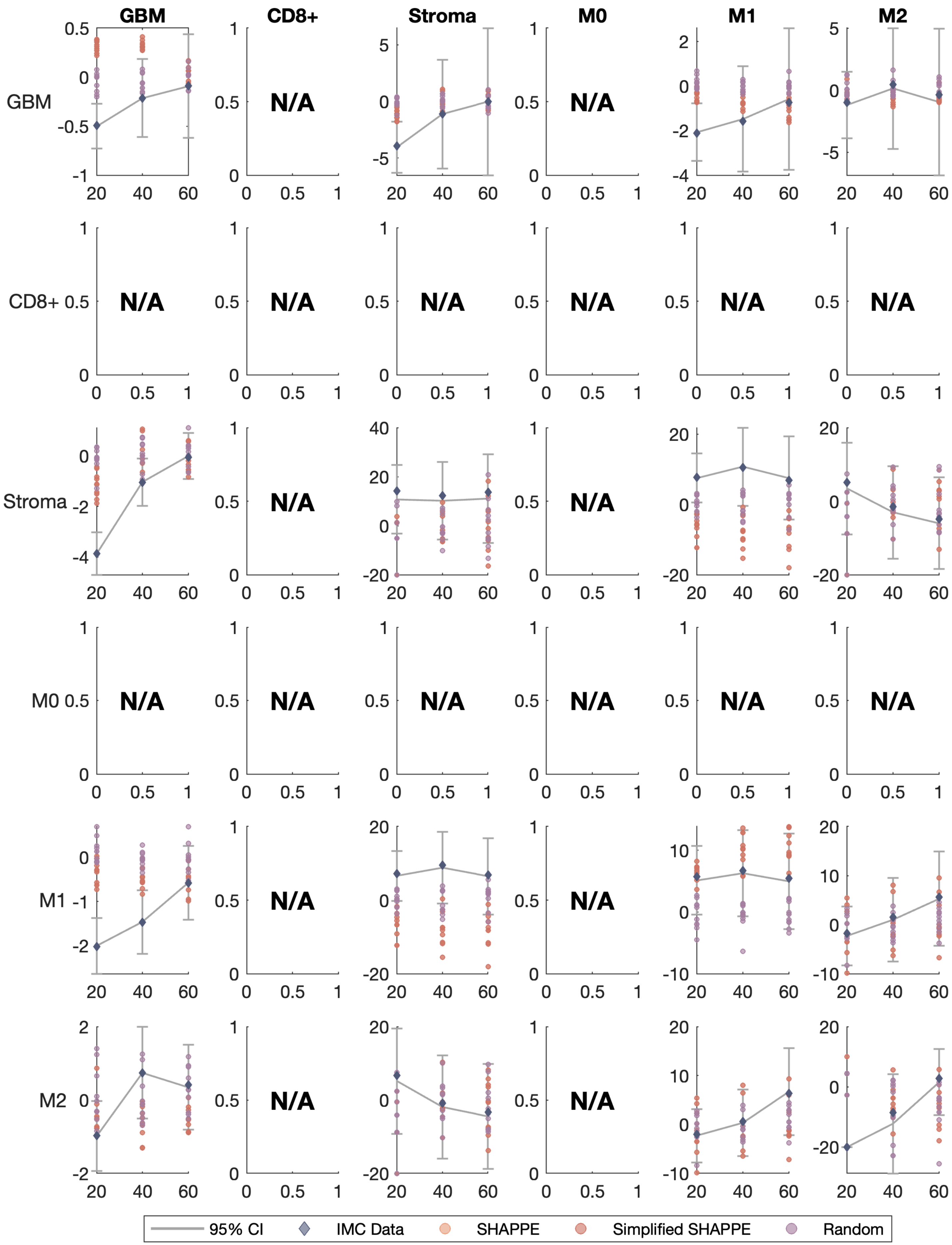

GBM
CD8+
Stroma
M0
M1
M2
N/A
95% CI
IMC Data
SHAPPE
Simplified SHAPPE
Random

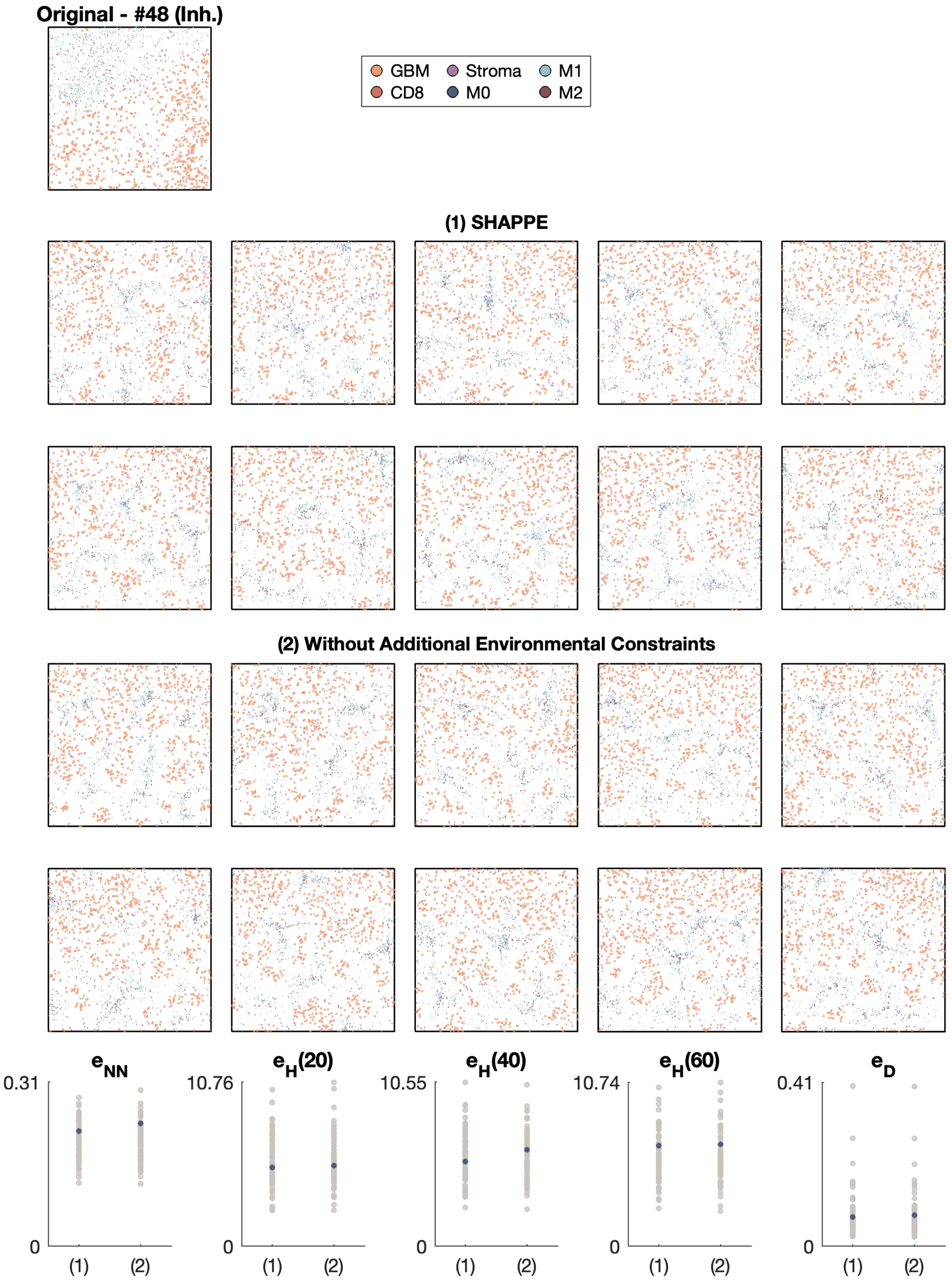

Original - #48 (Inh.)
GBM
Stroma
M1
CD8
M0
M2
(1) SHAPPE
(2) Without Additional Environmental Constraints
e_NN
e_H(20)
e_H(40)
e_H(60)
e_D
0.31
10.76
10.55
10.74
0.41
0
(1)
(2)

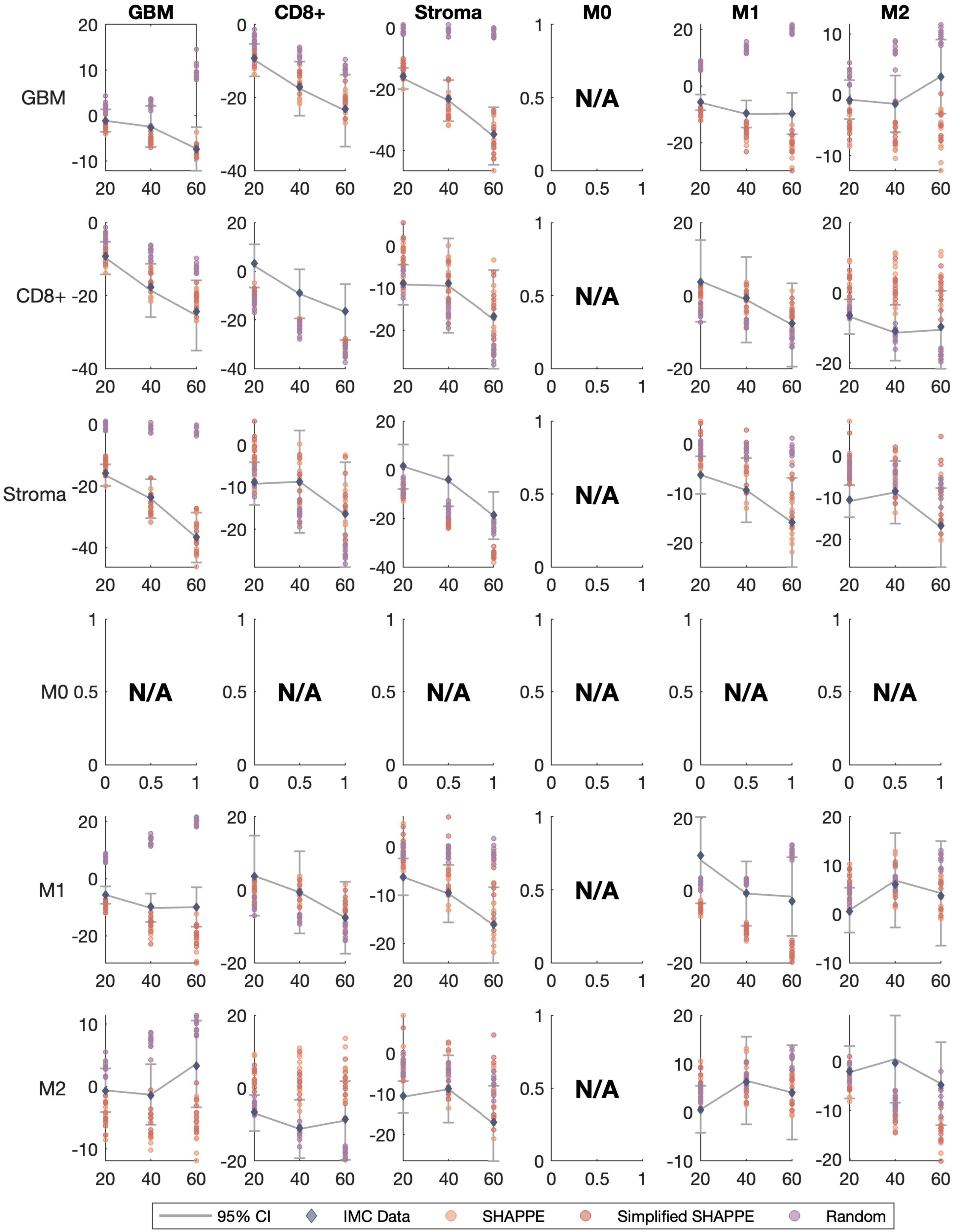
GBM
CD8+
Stroma
M0
M1
M2
N/A
95% CI
IMC Data
SHAPPE
Simplified SHAPPE
Random

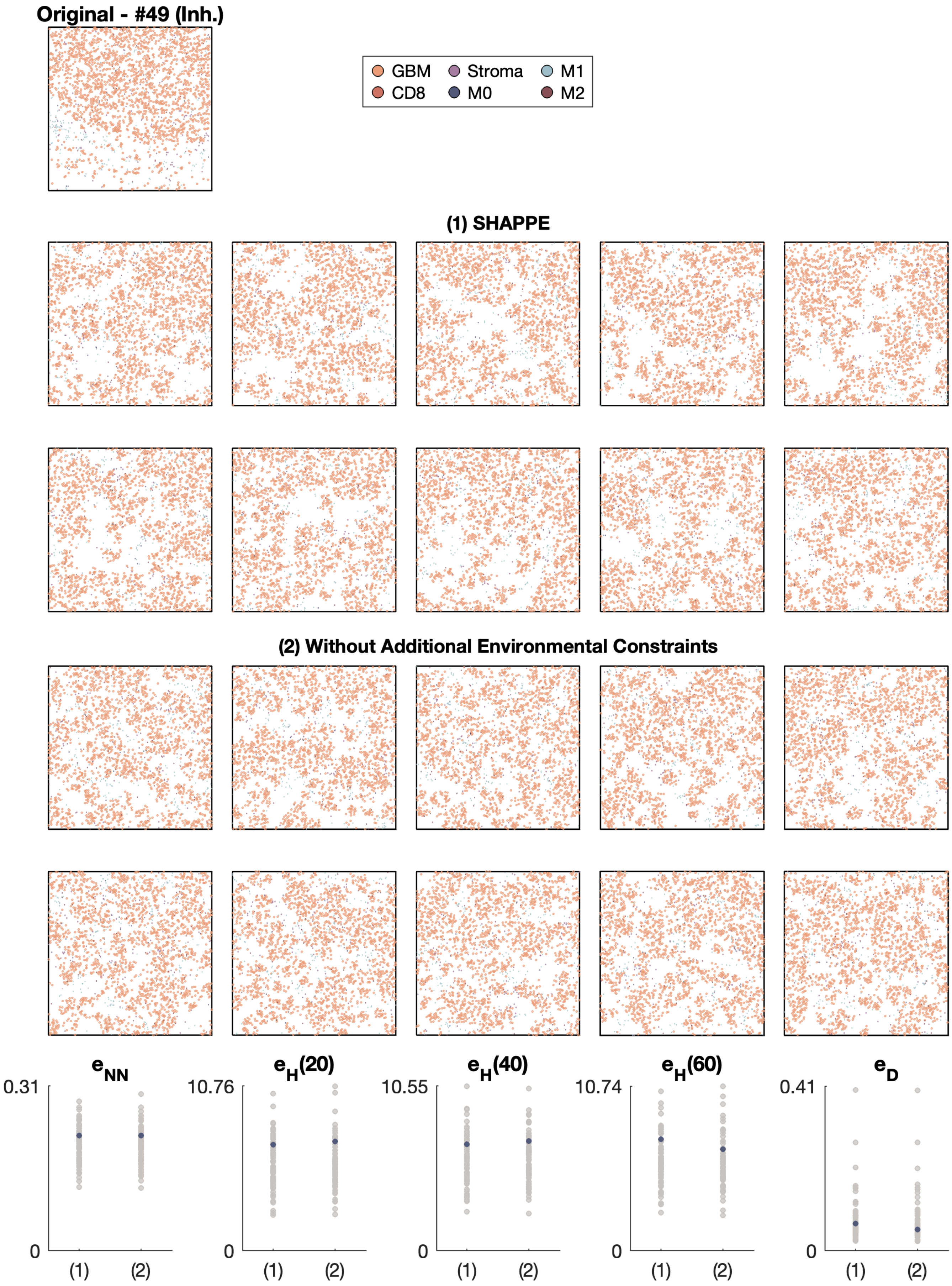

Original - #49 (Inh.)
GBM
Stroma
M1
CD8
M0
M2
(1) SHAPPE
(2) Without Additional Environmental Constraints
e_NN
e_H(20)
e_H(40)
e_H(60)
e_D
0.31
10.76
10.55
10.74
0.41
0
(1)
(2)

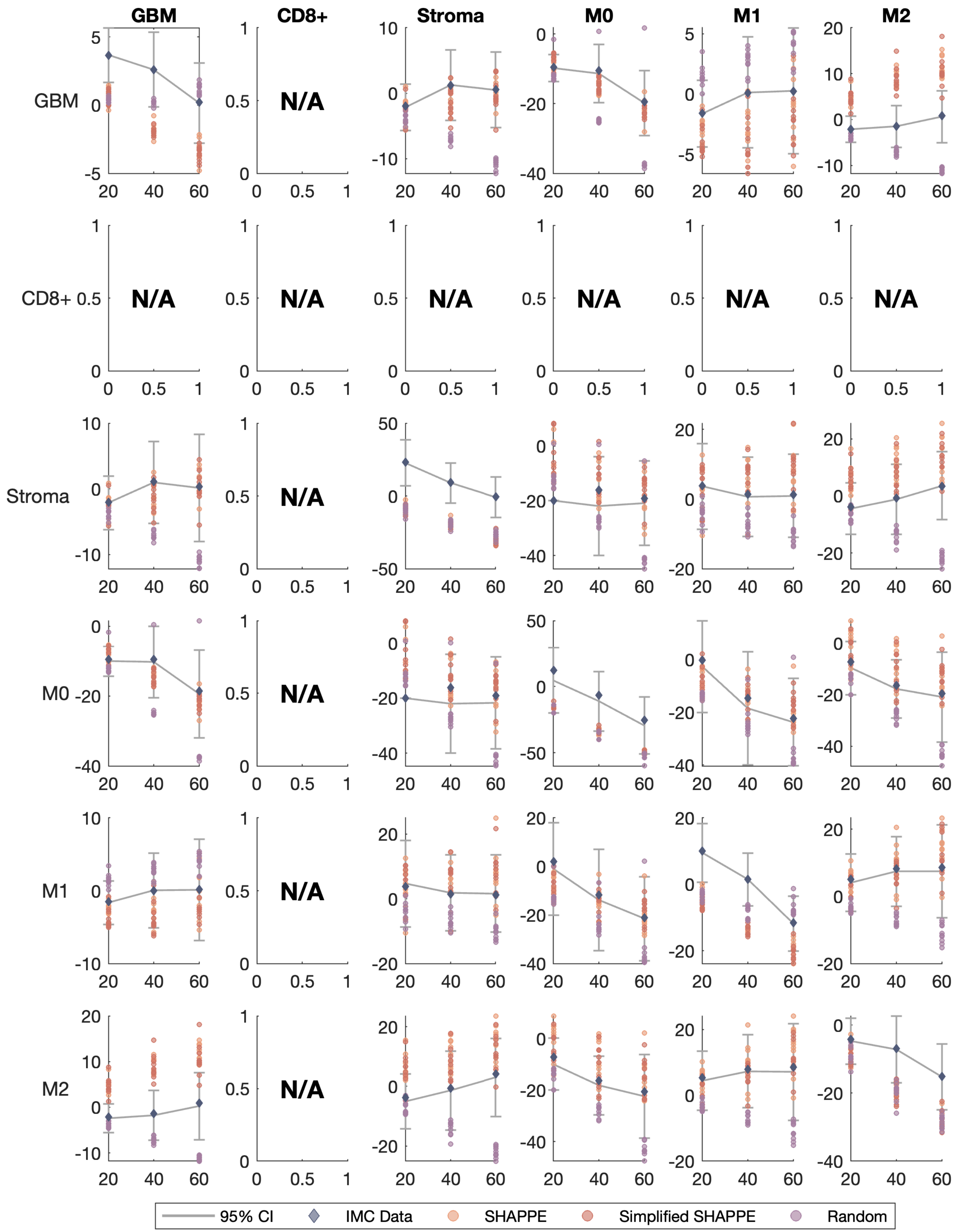

GBM
CD8+
Stroma
M0
M1
M2
N/A
95% CI
IMC Data
SHAPPE
Simplified SHAPPE
Random

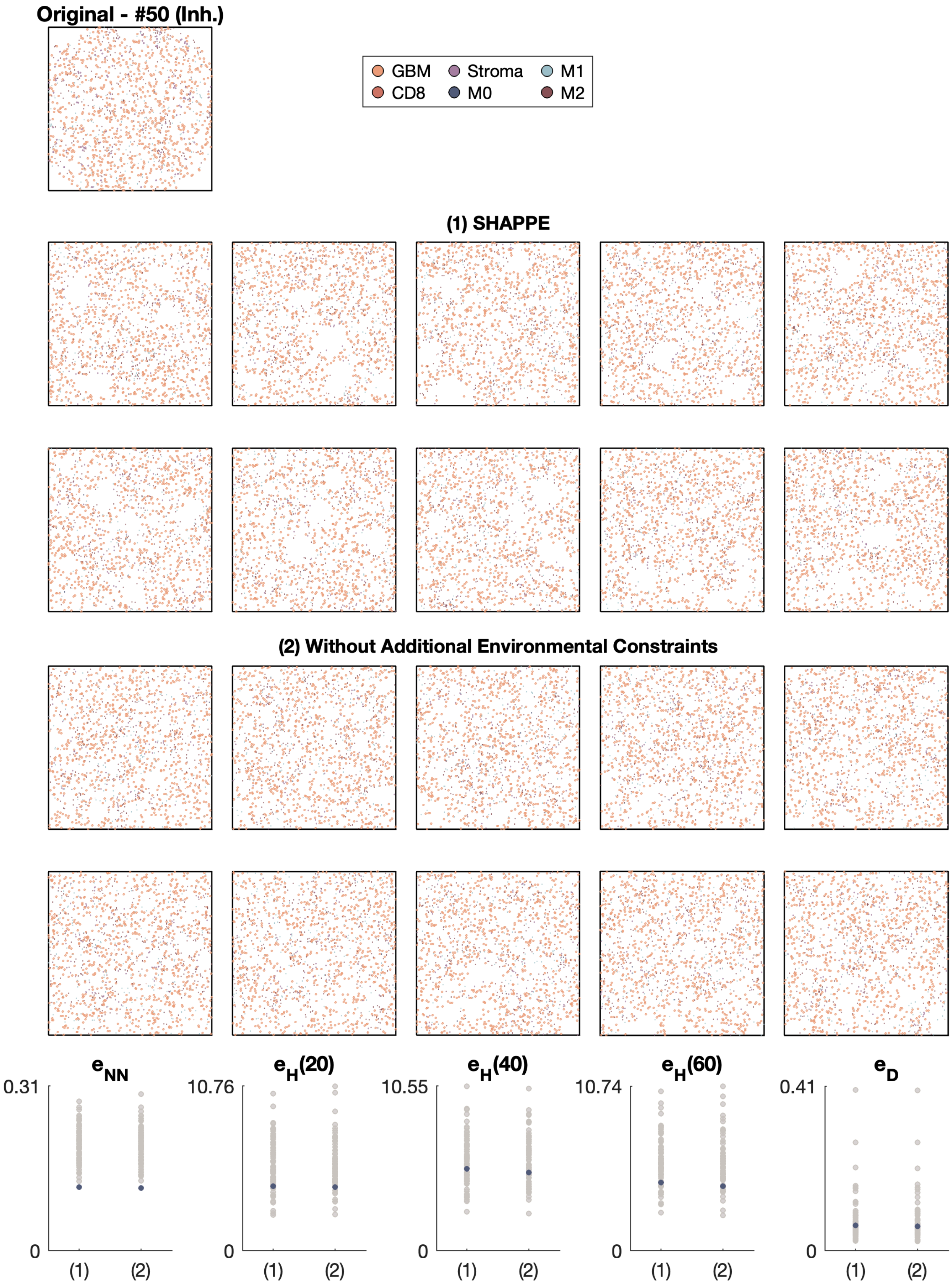

Original - #50 (Inh.)
GBM
Stroma
M1
CD8
M0
M2
(1) SHAPPE
(2) Without Additional Environmental Constraints
$e_{NN}$
0.31
0
(1)
(2)
$e_H(20)$
10.76
0
(1)
(2)
$e_H(40)$
10.55
0
(1)
(2)
$e_H(60)$
10.74
0
(1)
(2)
$e_D$
0.41
0
(1)
(2)

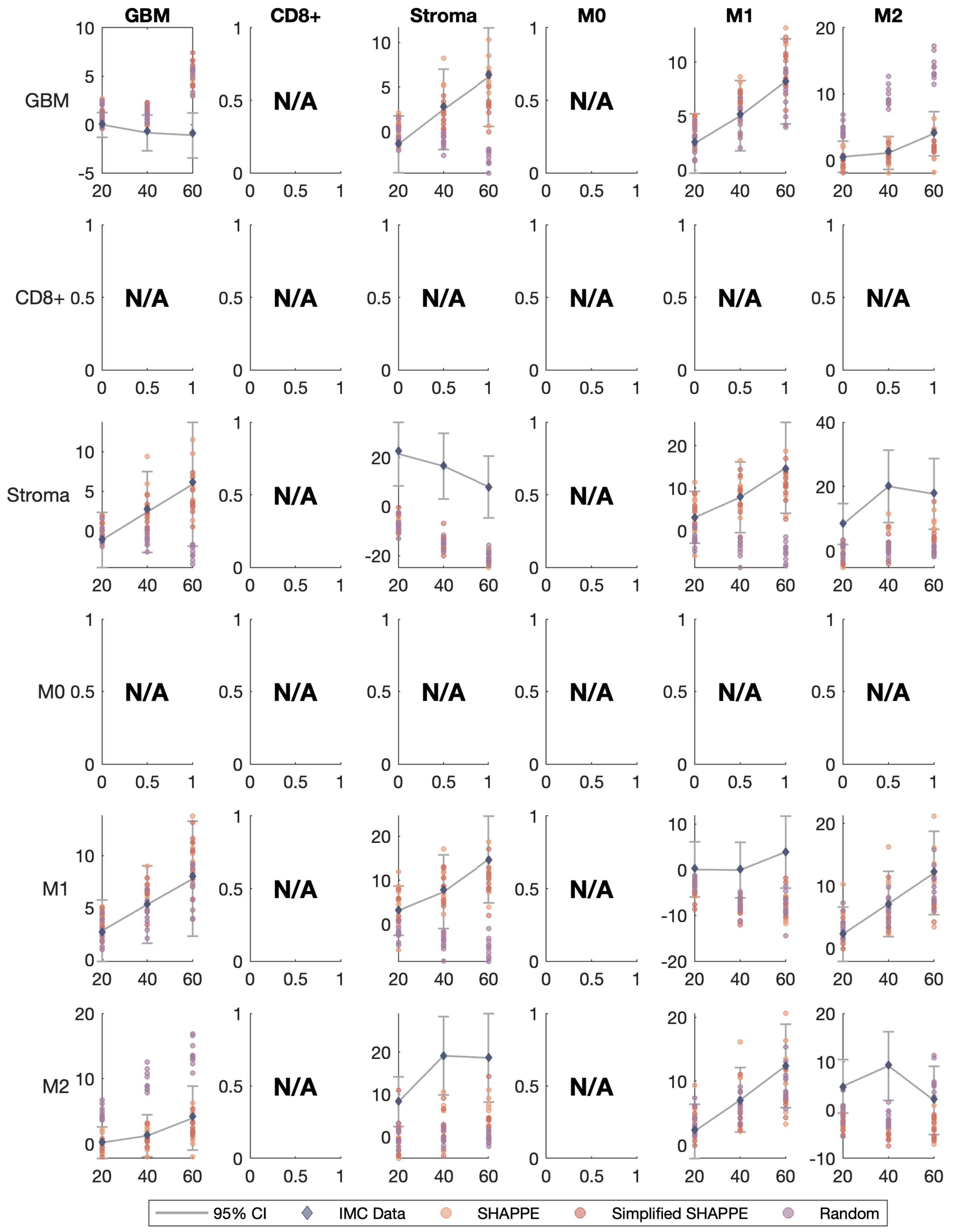

GBM
CD8+
Stroma
M0
M1
M2
N/A
95% CI
IMC Data
SHAPPE
Simplified SHAPPE
Random

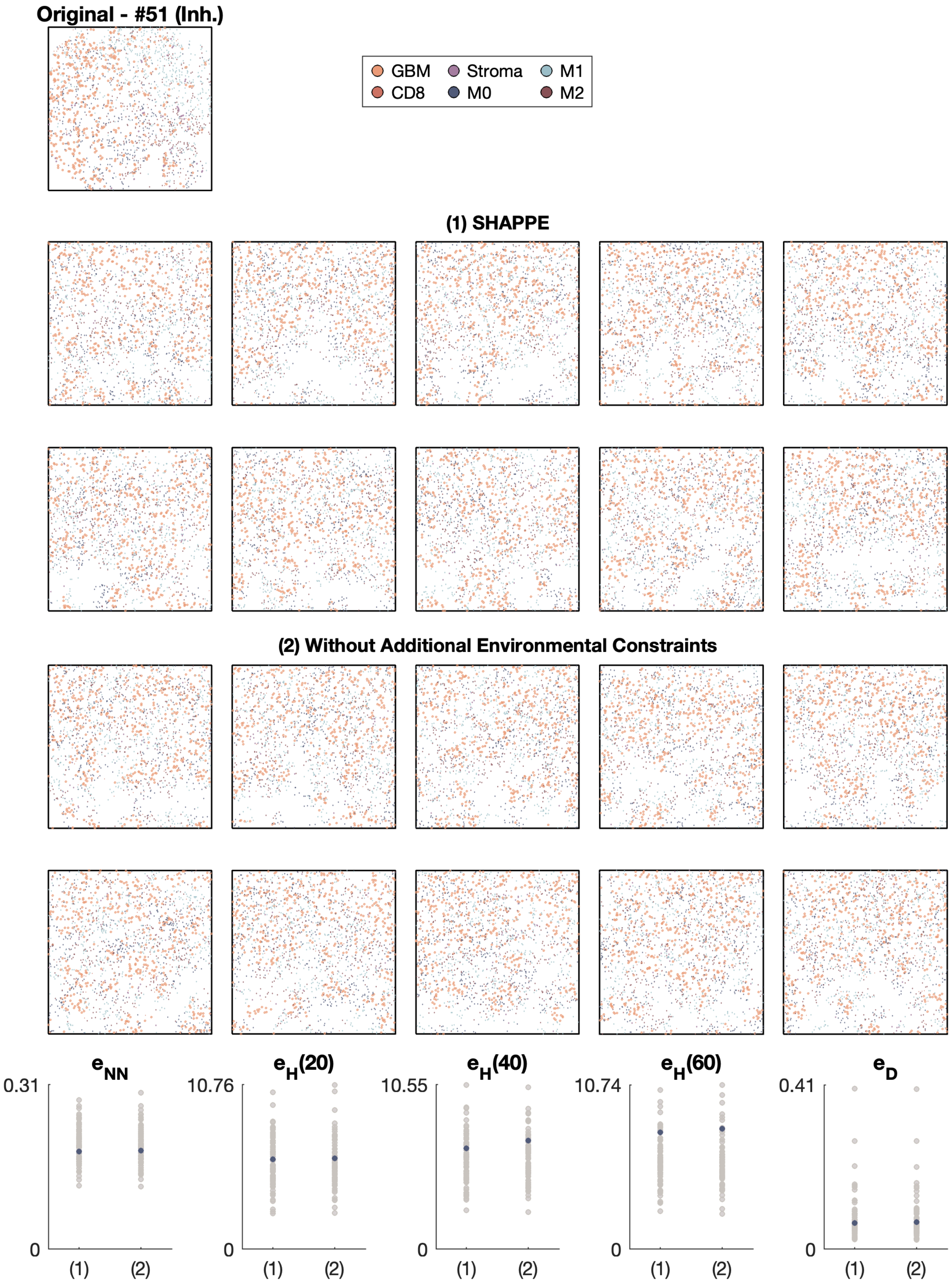

Original - #51 (Inh.)
GBM
CD8
Stroma
M0
M1
M2
(1) SHAPPE
(2) Without Additional Environmental Constraints
$e_{NN}$
0.31
0
(1)
(2)
$e_H(20)$
10.76
0
(1)
(2)
$e_H(40)$
10.55
0
(1)
(2)
$e_H(60)$
10.74
0
(1)
(2)
$e_D$
0.41
0
(1)
(2)

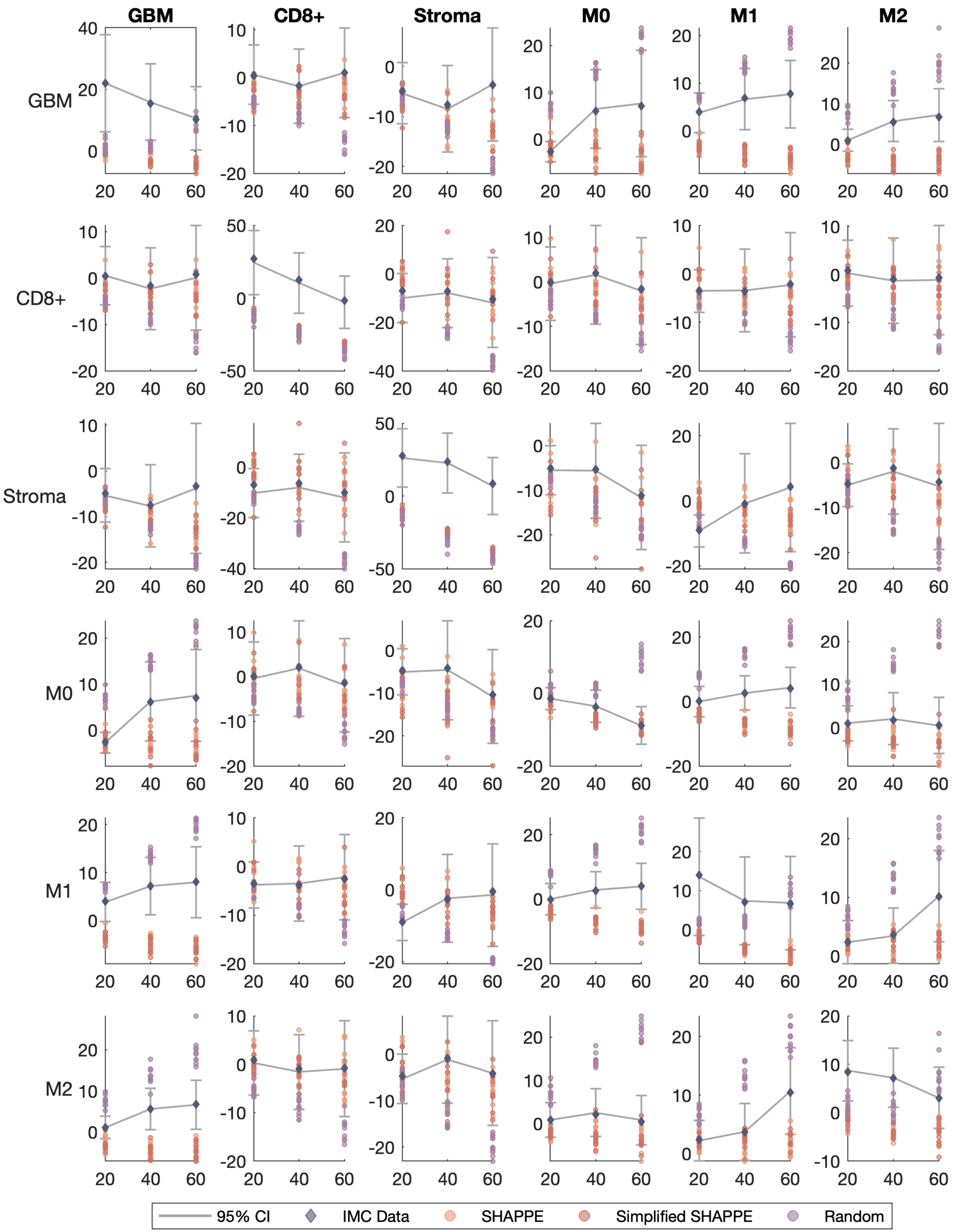
GBM
CD8+
Stroma
M0
M1
M2
GBM
CD8+
Stroma
M0
M1
M2
95% CI
IMC Data
SHAPPE
Simplified SHAPPE
Random

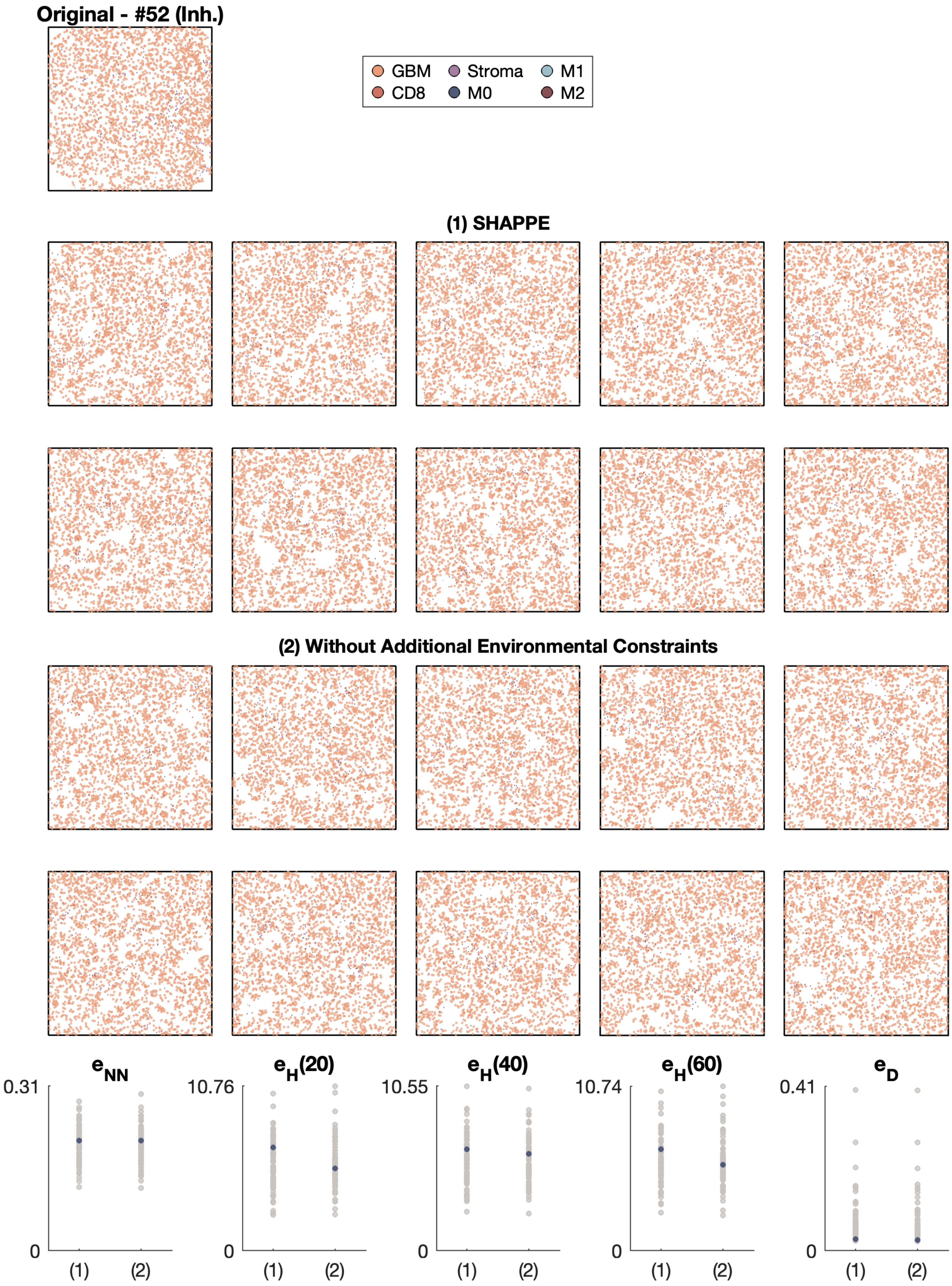

Original - #52 (Inh.)
GBM
Stroma
M1
CD8
M0
M2
(1) SHAPPE
(2) Without Additional Environmental Constraints
$e_{NN}$
$e_H(20)$
$e_H(40)$
$e_H(60)$
$e_D$
0.31
10.76
10.55
10.74
0.41
0
(1)
(2)

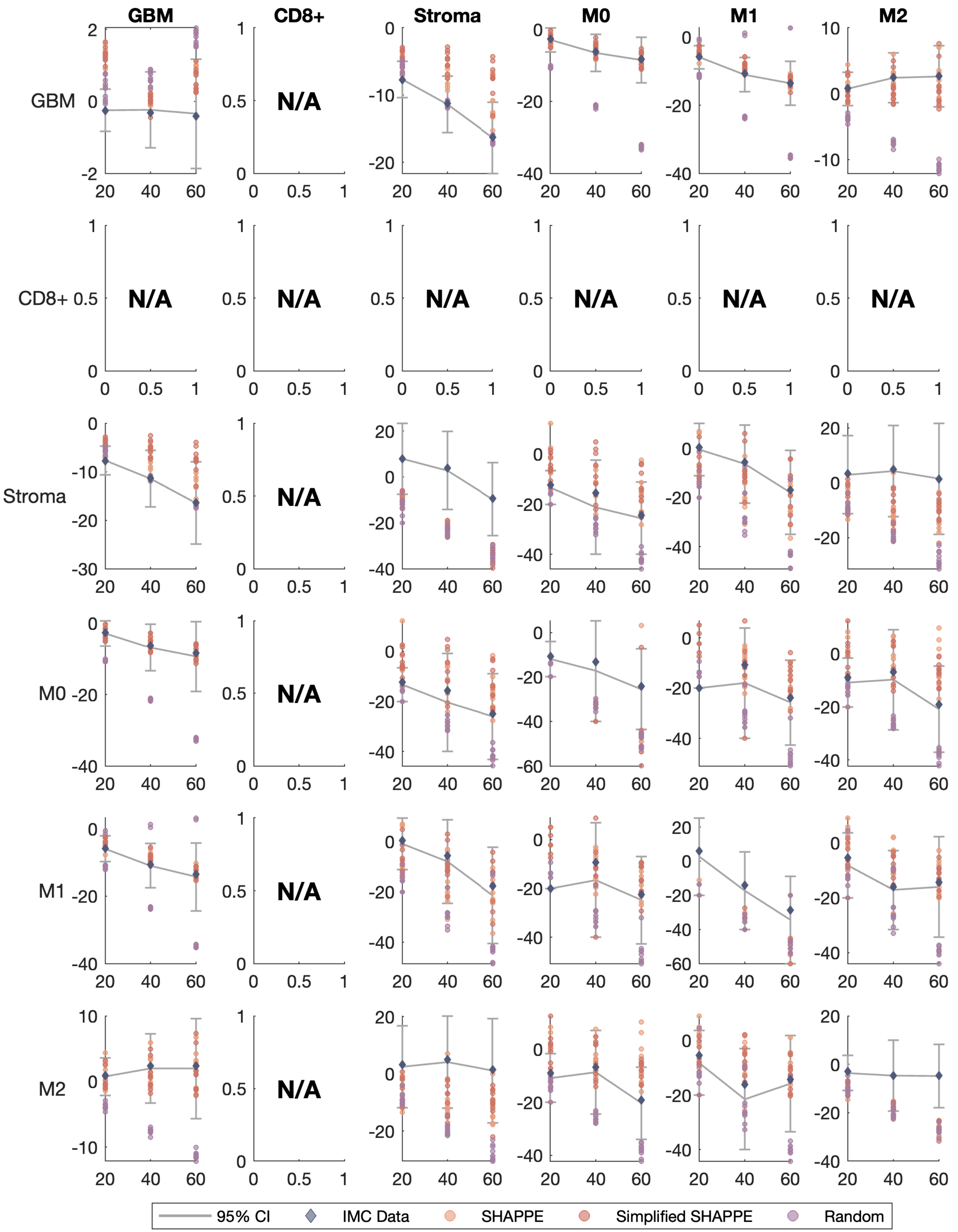
GBM
CD8+
Stroma
M0
M1
M2
N/A
95% CI
IMC Data
SHAPPE
Simplified SHAPPE
Random

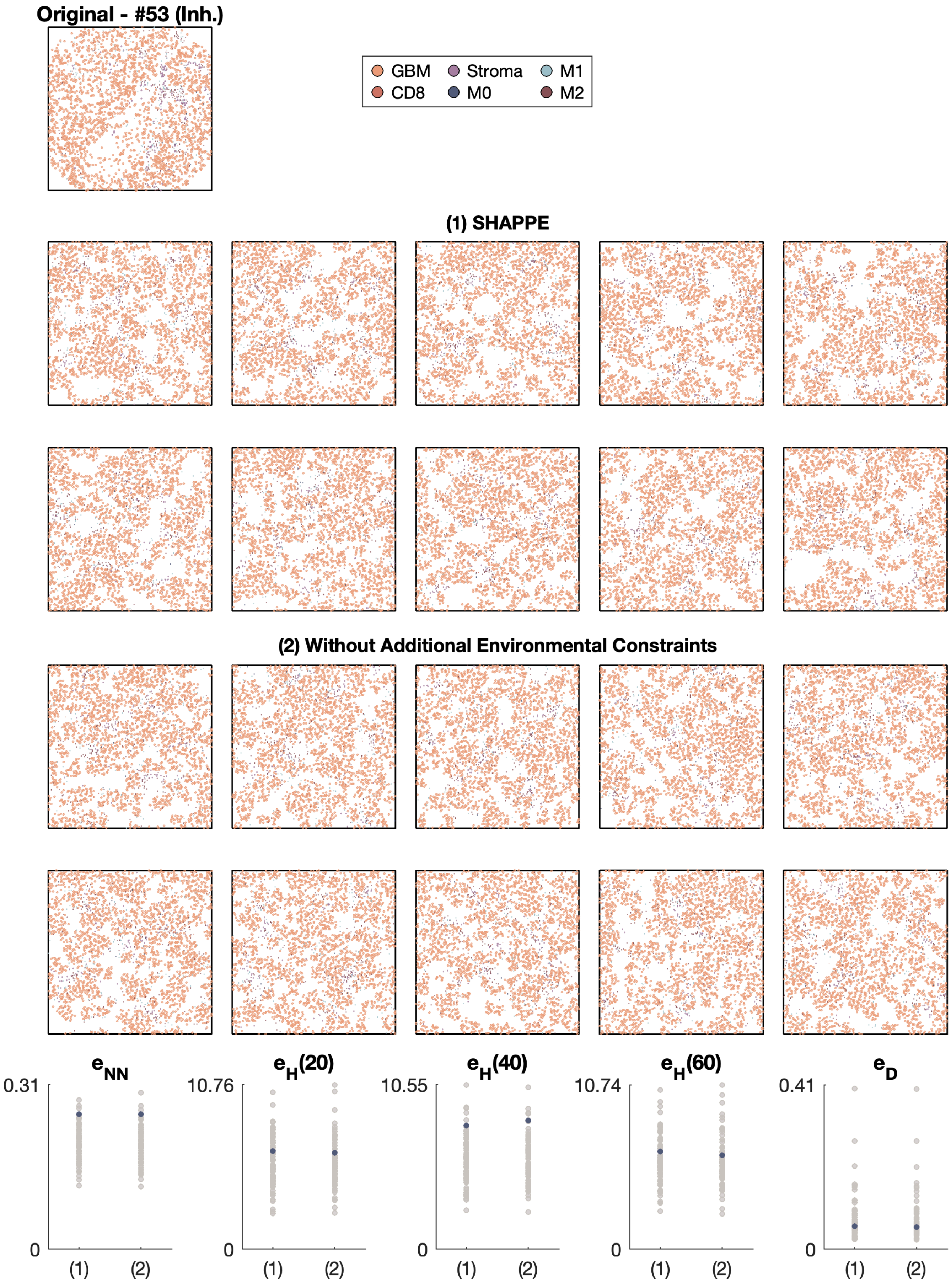

Original - #53 (Inh.)
GBM
CD8
Stroma
M0
M1
M2
(1) SHAPPE
(2) Without Additional Environmental Constraints
$e_{NN}$
$e_H(20)$
$e_H(40)$
$e_H(60)$
$e_D$
0.31
10.76
10.55
10.74
0.41
0
(1)
(2)

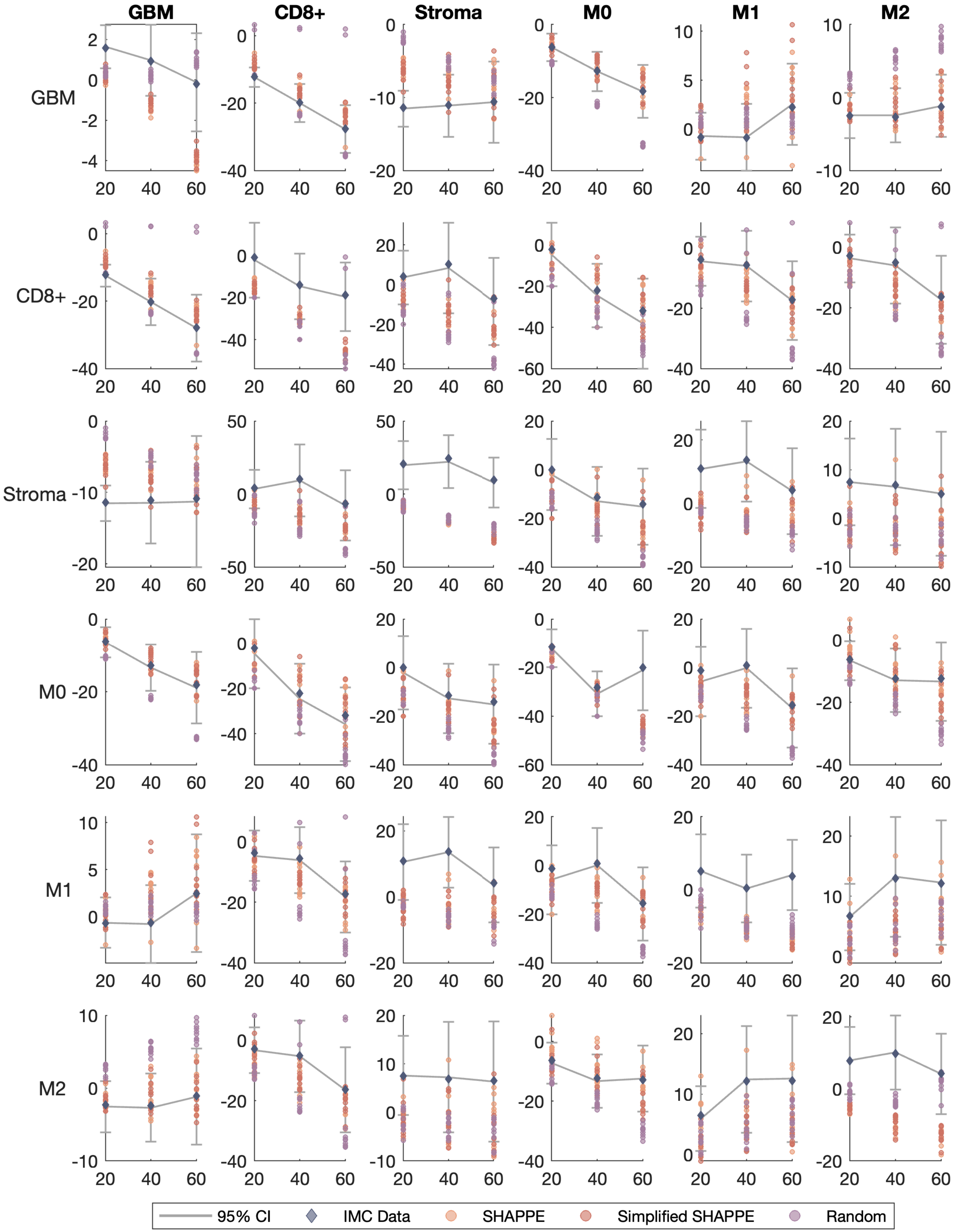
GBM
CD8+
Stroma
M0
M1
M2
GBM
CD8+
Stroma
M0
M1
M2
20
40
60
95% CI
IMC Data
SHAPPE
Simplified SHAPPE
Random

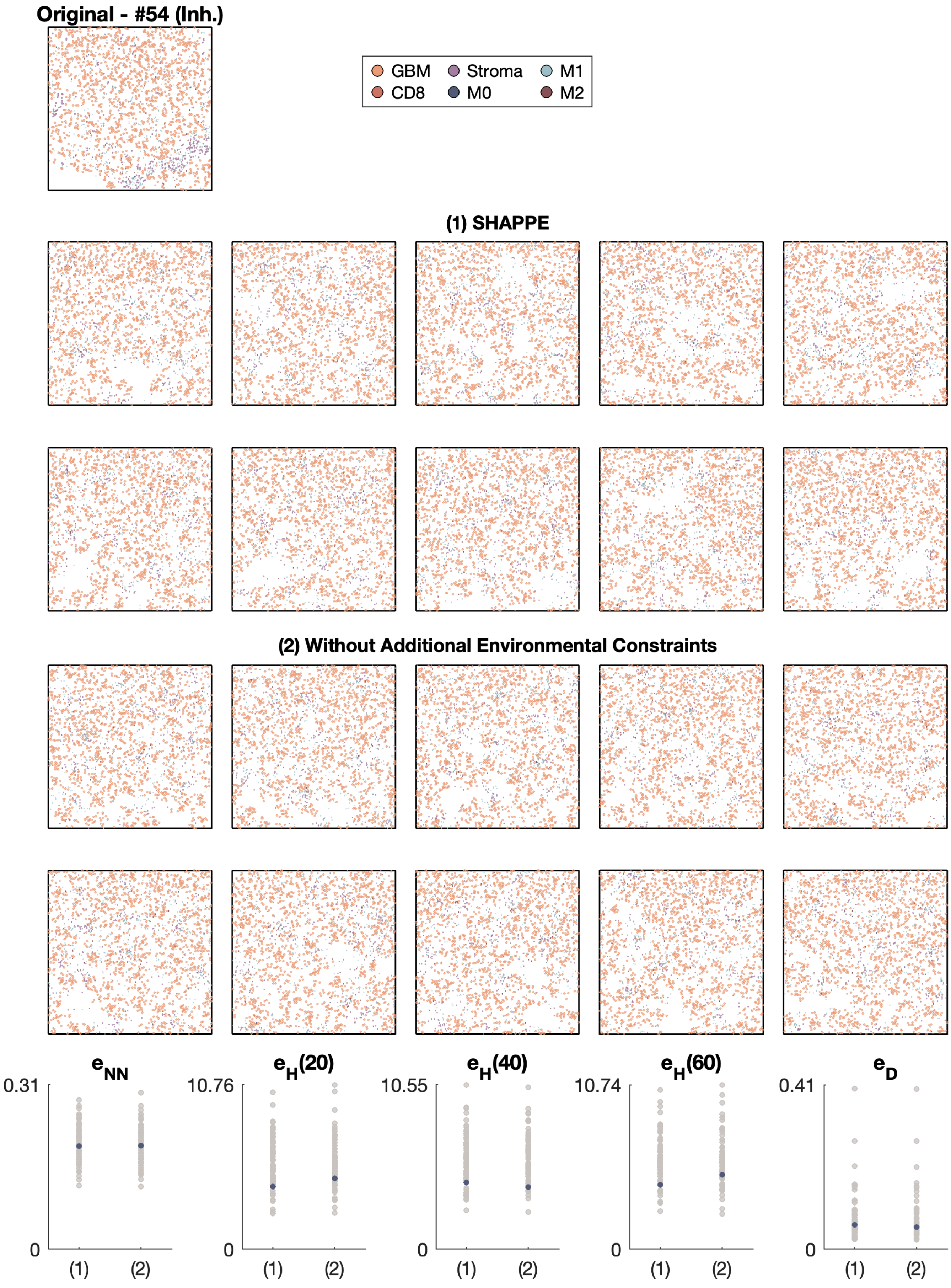
Original - #54 (Inh.)
GBM
Stroma
M1
CD8
M0
M2
(1) SHAPPE
(2) Without Additional Environmental Constraints
$e_{NN}$
$e_H(20)$
$e_H(40)$
$e_H(60)$
$e_D$
0.31
10.76
10.55
10.74
0.41
0
(1)
(2)

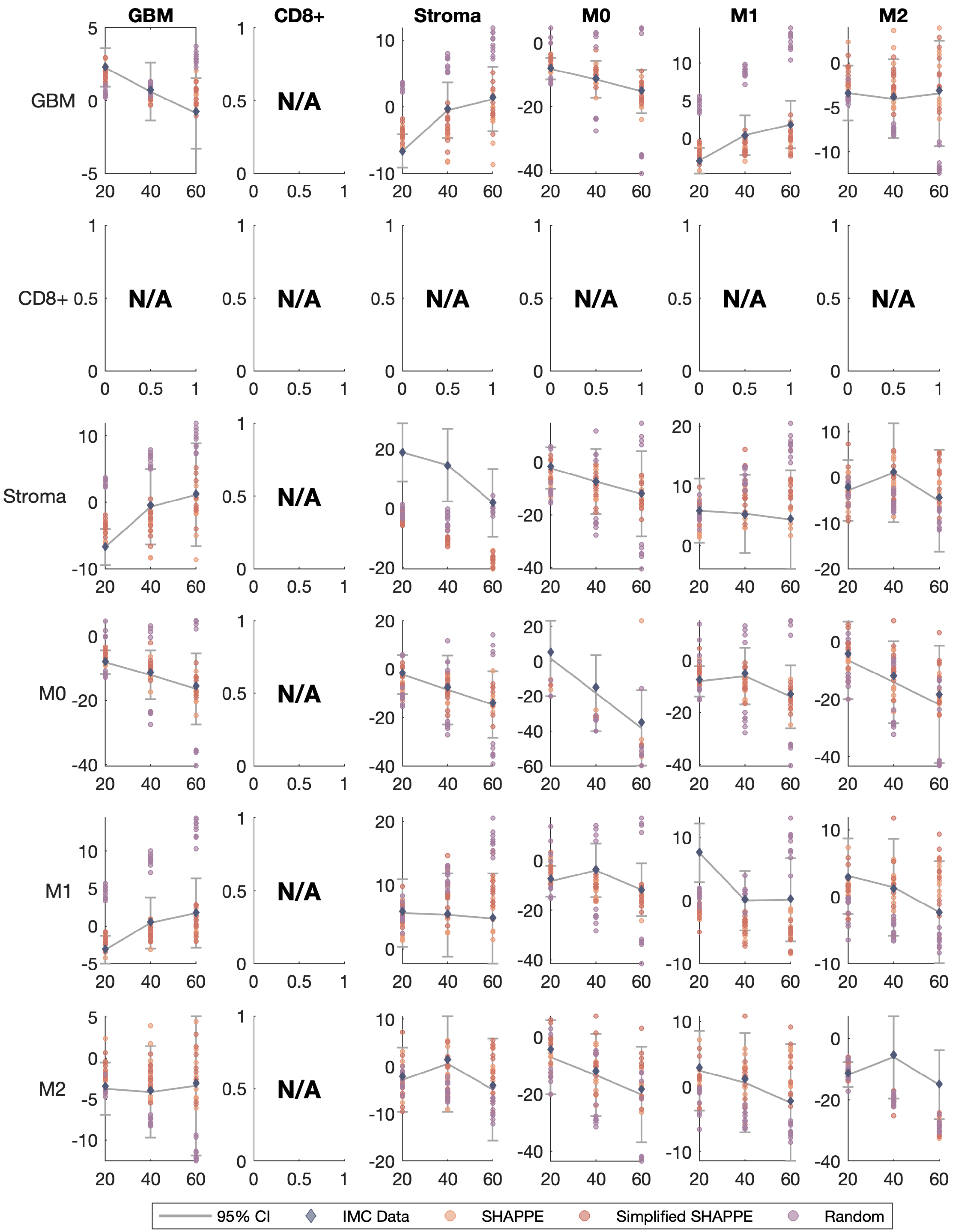
GBM
CD8+
Stroma
M0
M1
M2
GBM
CD8+
Stroma
M0
M1
M2
N/A
95% CI
IMC Data
SHAPPE
Simplified SHAPPE
Random

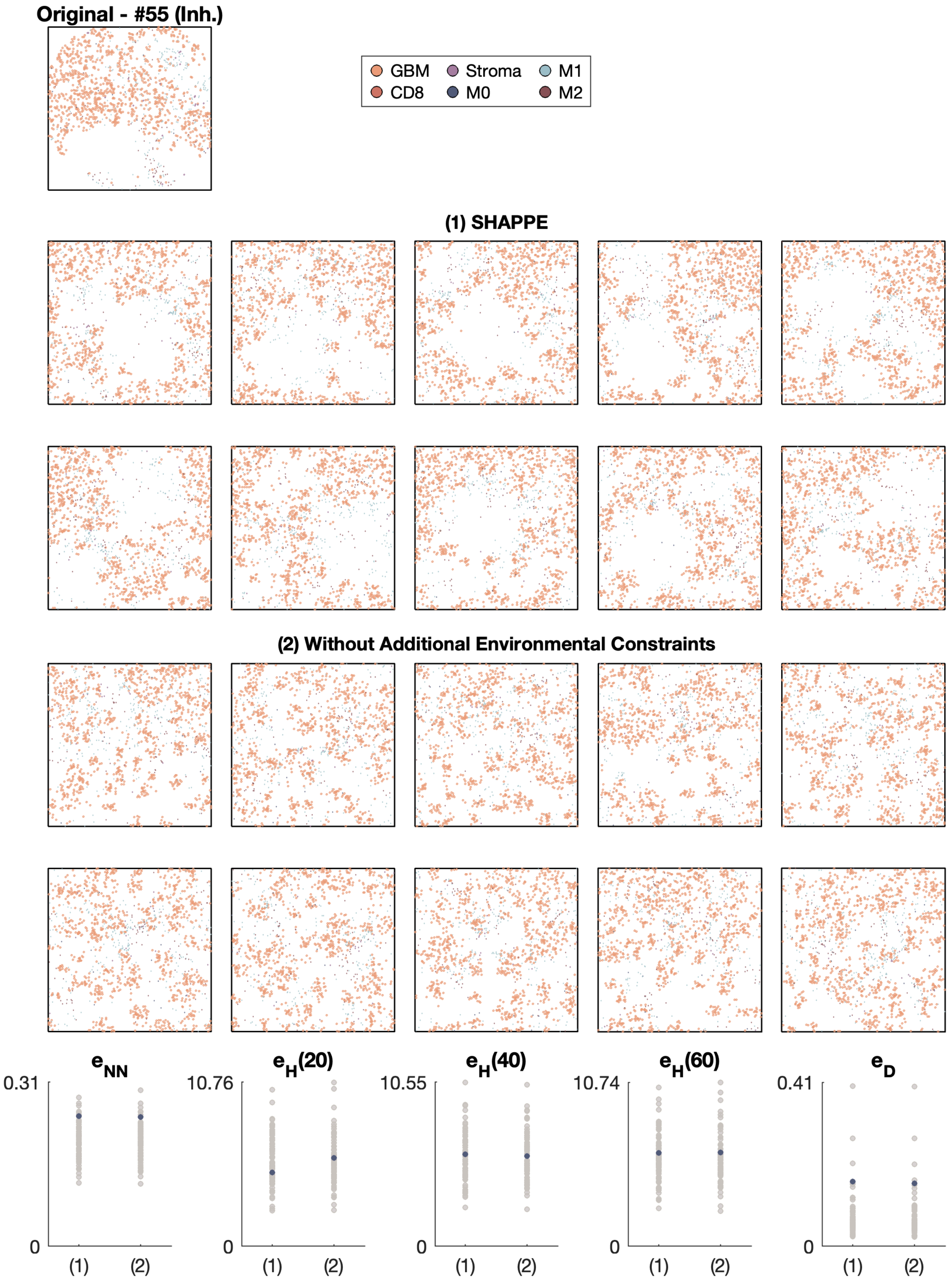

Original - #55 (Inh.)
GBM
Stroma
M1
CD8
M0
M2
(1) SHAPPE
(2) Without Additional Environmental Constraints
$e_{NN}$
$e_H(20)$
$e_H(40)$
$e_H(60)$
$e_D$
0.31
10.76
10.55
10.74
0.41
0
(1)
(2)

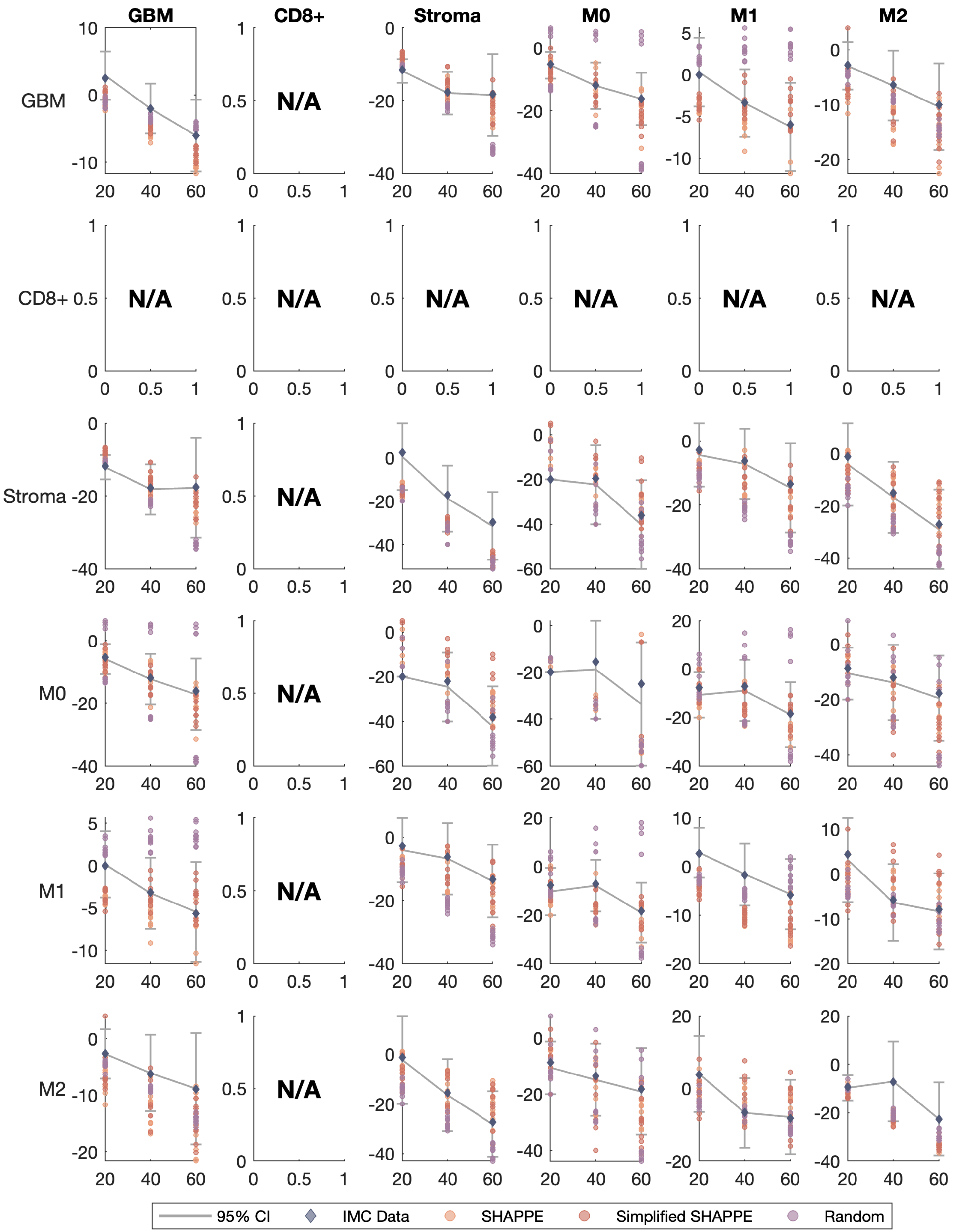
GBM
CD8+
Stroma
M0
M1
M2
GBM
CD8+
Stroma
M0
M1
M2
N/A
95% CI
IMC Data
SHAPPE
Simplified SHAPPE
Random

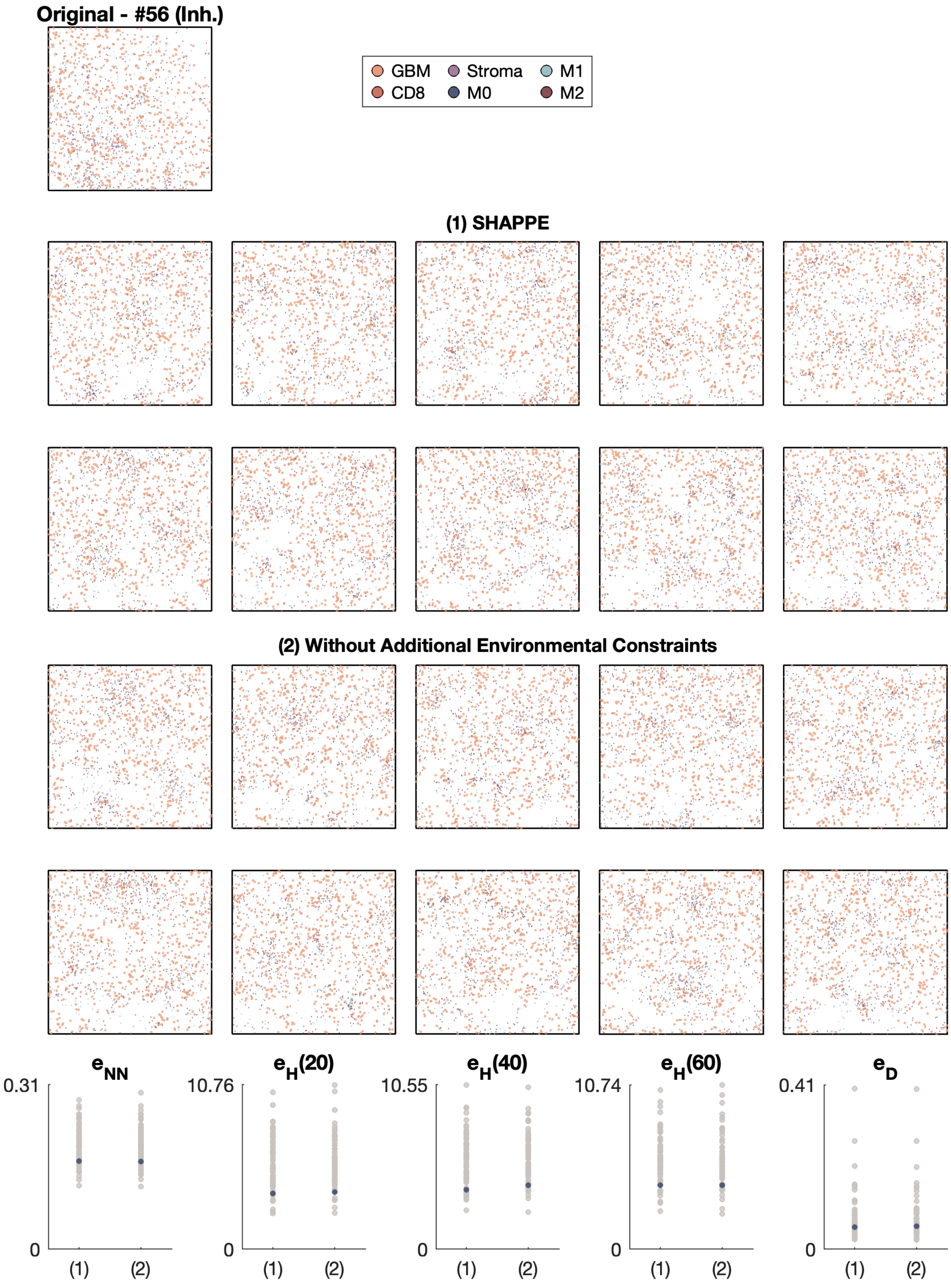

Original - #56 (Inh.)
GBM
CD8
Stroma
M0
M1
M2
(1) SHAPPE
(2) Without Additional Environmental Constraints
$e_{NN}$
$e_H(20)$
$e_H(40)$
$e_H(60)$
$e_D$
0.31
10.76
10.55
10.74
0.41
0
(1)
(2)

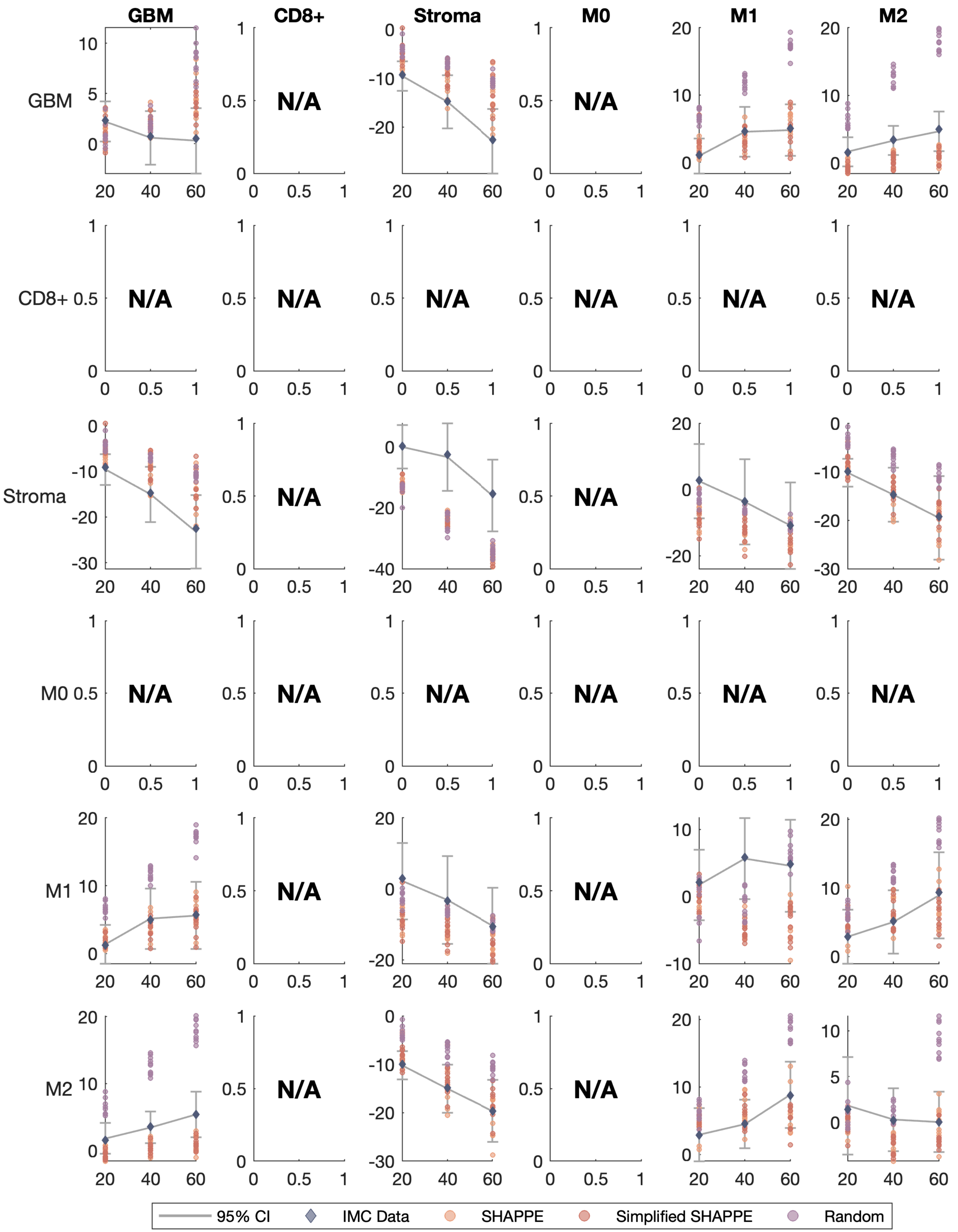

GBM
CD8+
Stroma
M0
M1
M2
N/A
95% CI
IMC Data
SHAPPE
Simplified SHAPPE
Random

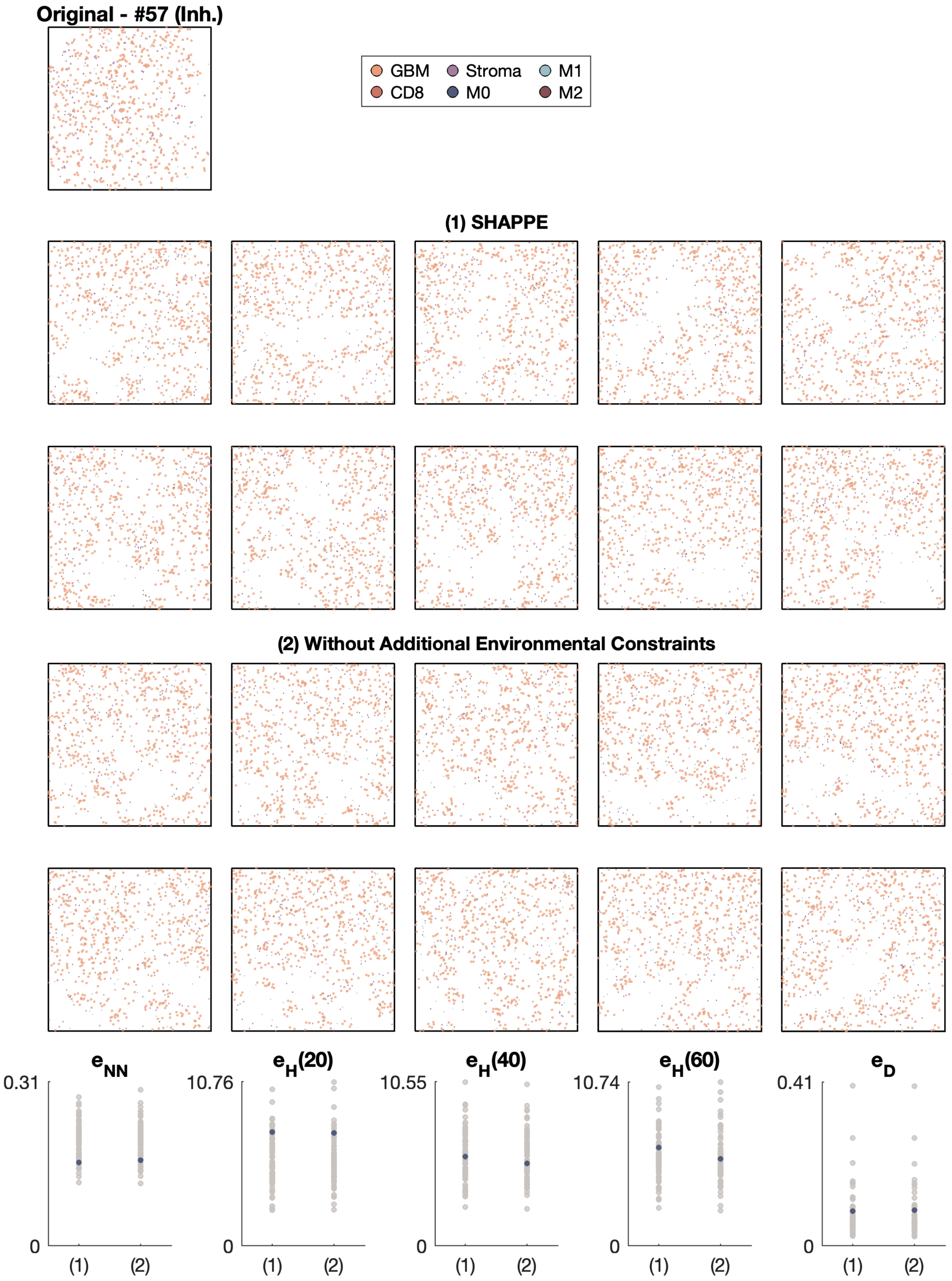

Original - #57 (Inh.)
GBM
Stroma
M1
CD8
M0
M2
(1) SHAPPE
(2) Without Additional Environmental Constraints
$e_{NN}$
0.31
0
(1)
(2)
$e_H(20)$
10.76
0
(1)
(2)
$e_H(40)$
10.55
0
(1)
(2)
$e_H(60)$
10.74
0
(1)
(2)
$e_D$
0.41
0
(1)
(2)

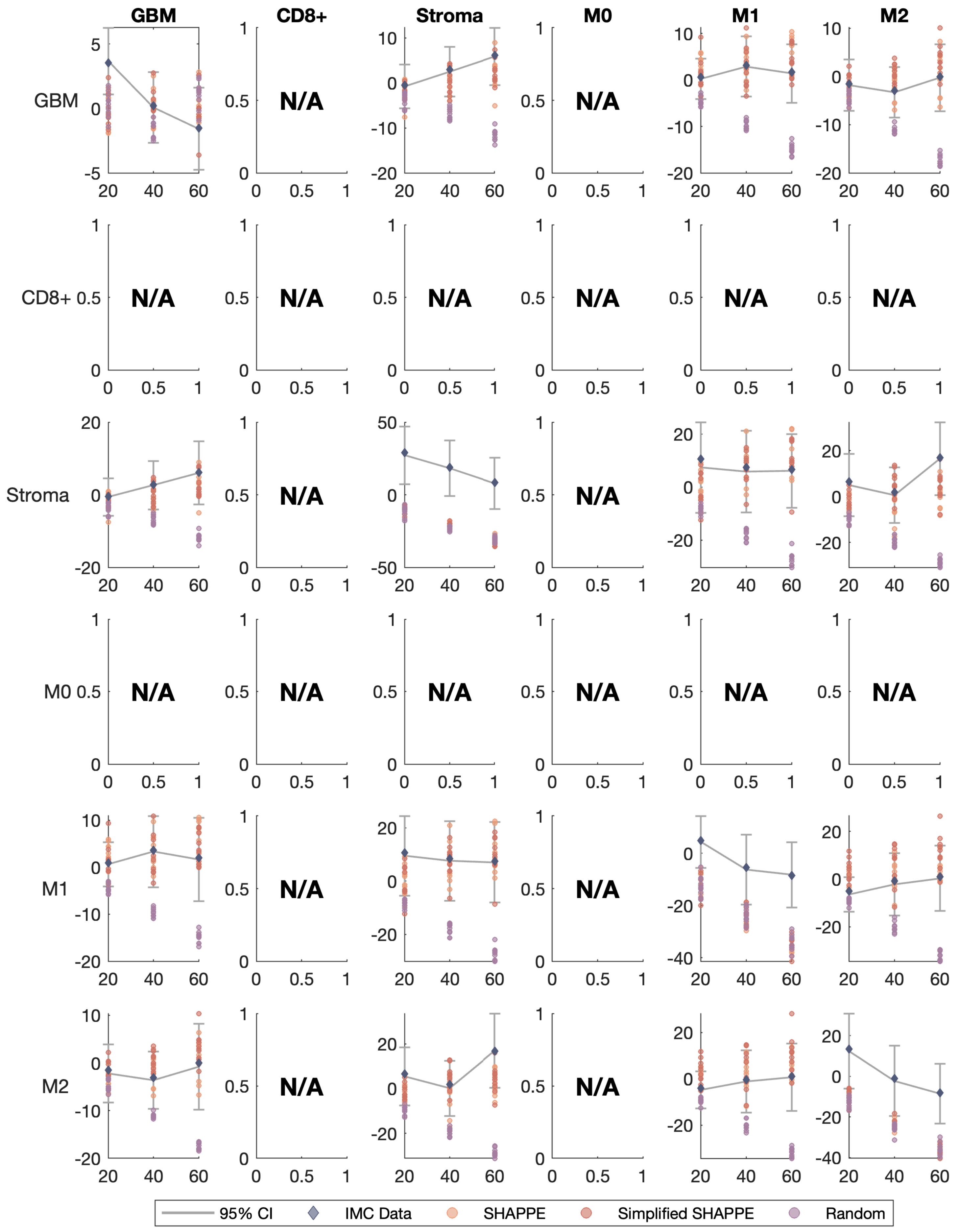
GBM
CD8+
Stroma
M0
M1
M2
N/A
95% CI
IMC Data
SHAPPE
Simplified SHAPPE
Random

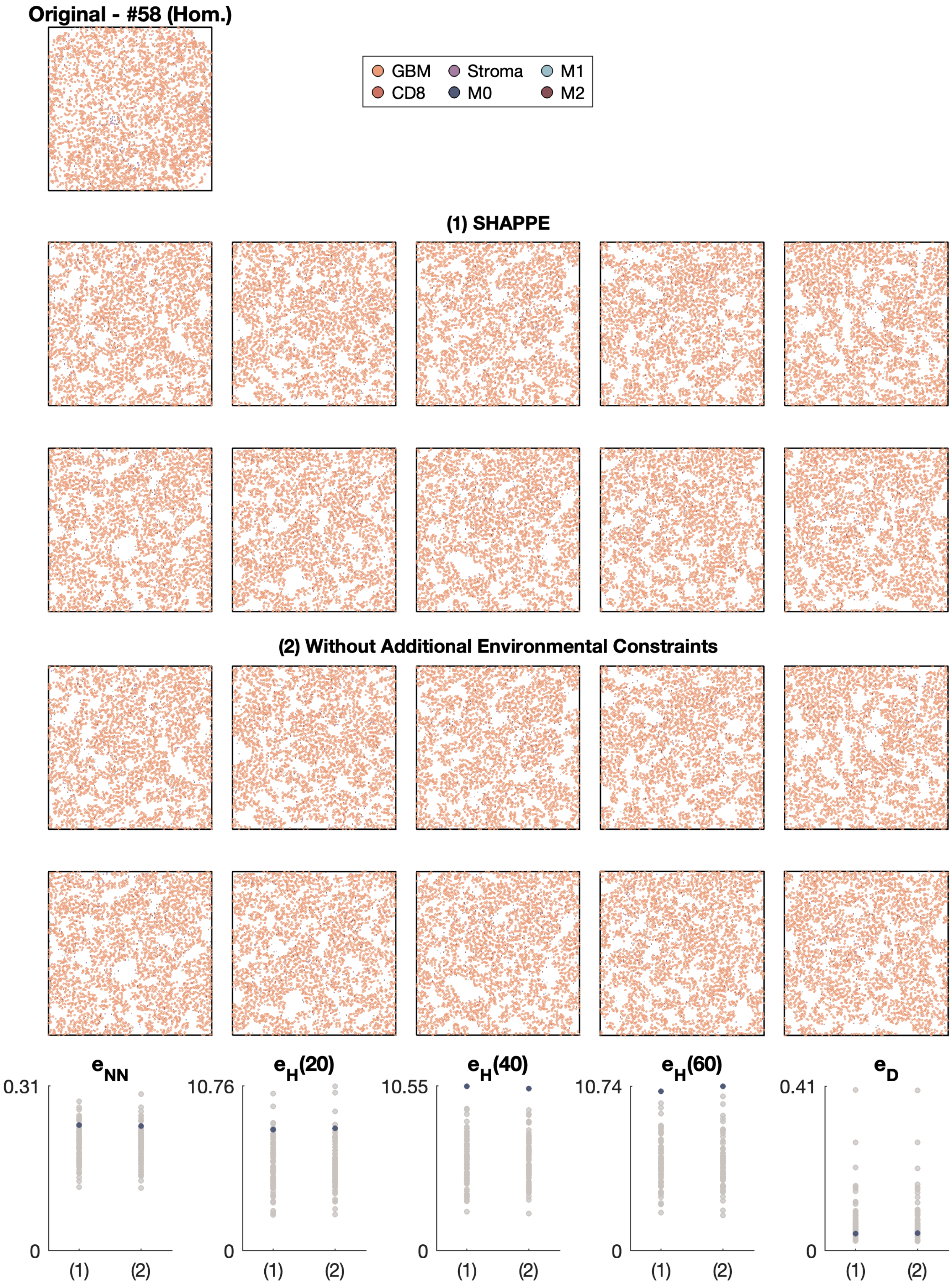

Original - #58 (Hom.)
GBM
Stroma
M1
CD8
M0
M2
(1) SHAPPE
(2) Without Additional Environmental Constraints
e_NN
0.31
0
(1)
(2)
e_H(20)
10.76
0
(1)
(2)
e_H(40)
10.55
0
(1)
(2)
e_H(60)
10.74
0
(1)
(2)
e_D
0.41
0
(1)
(2)

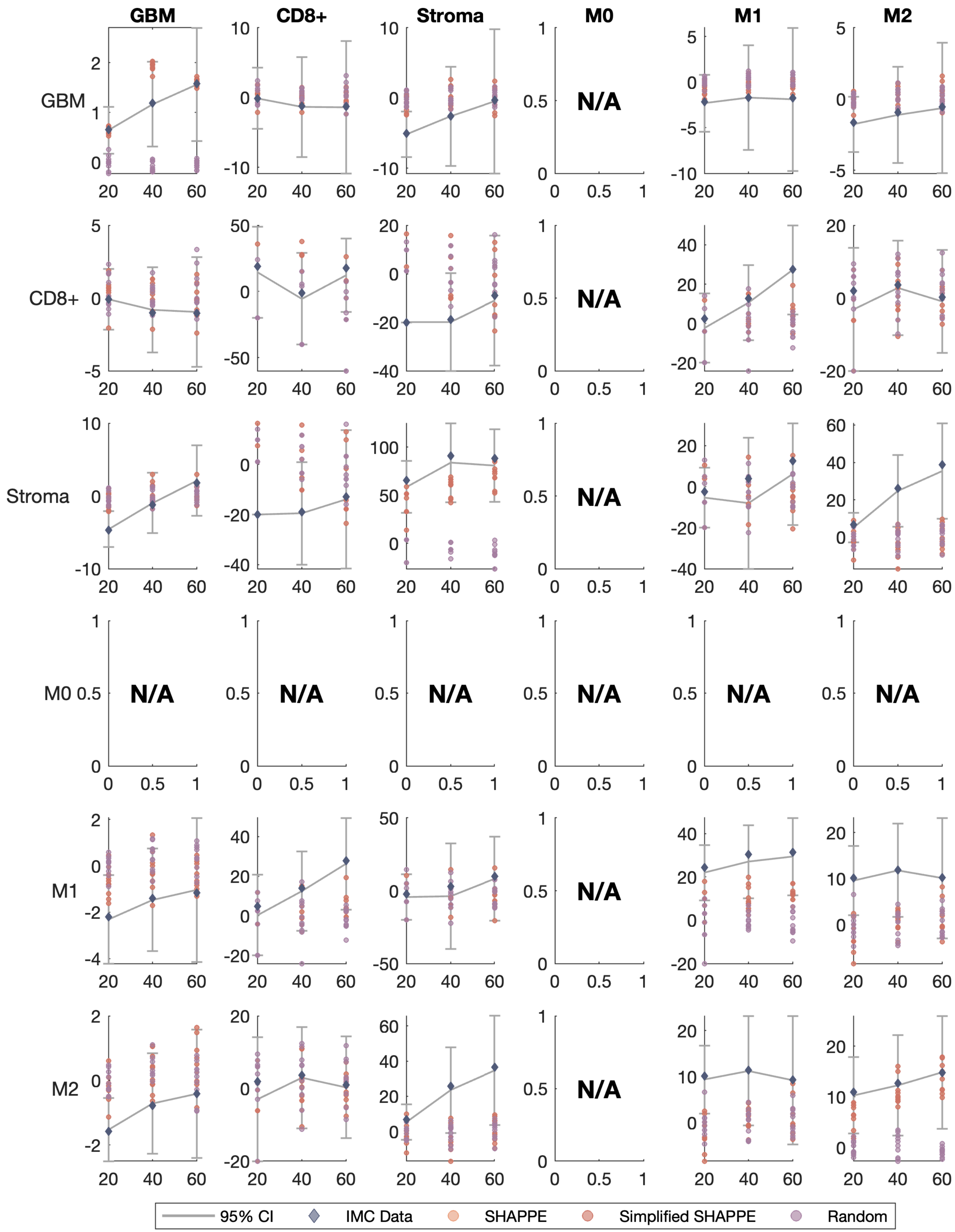
GBM
CD8+
Stroma
M0
M1
M2
N/A
95% CI
IMC Data
SHAPPE
Simplified SHAPPE
Random

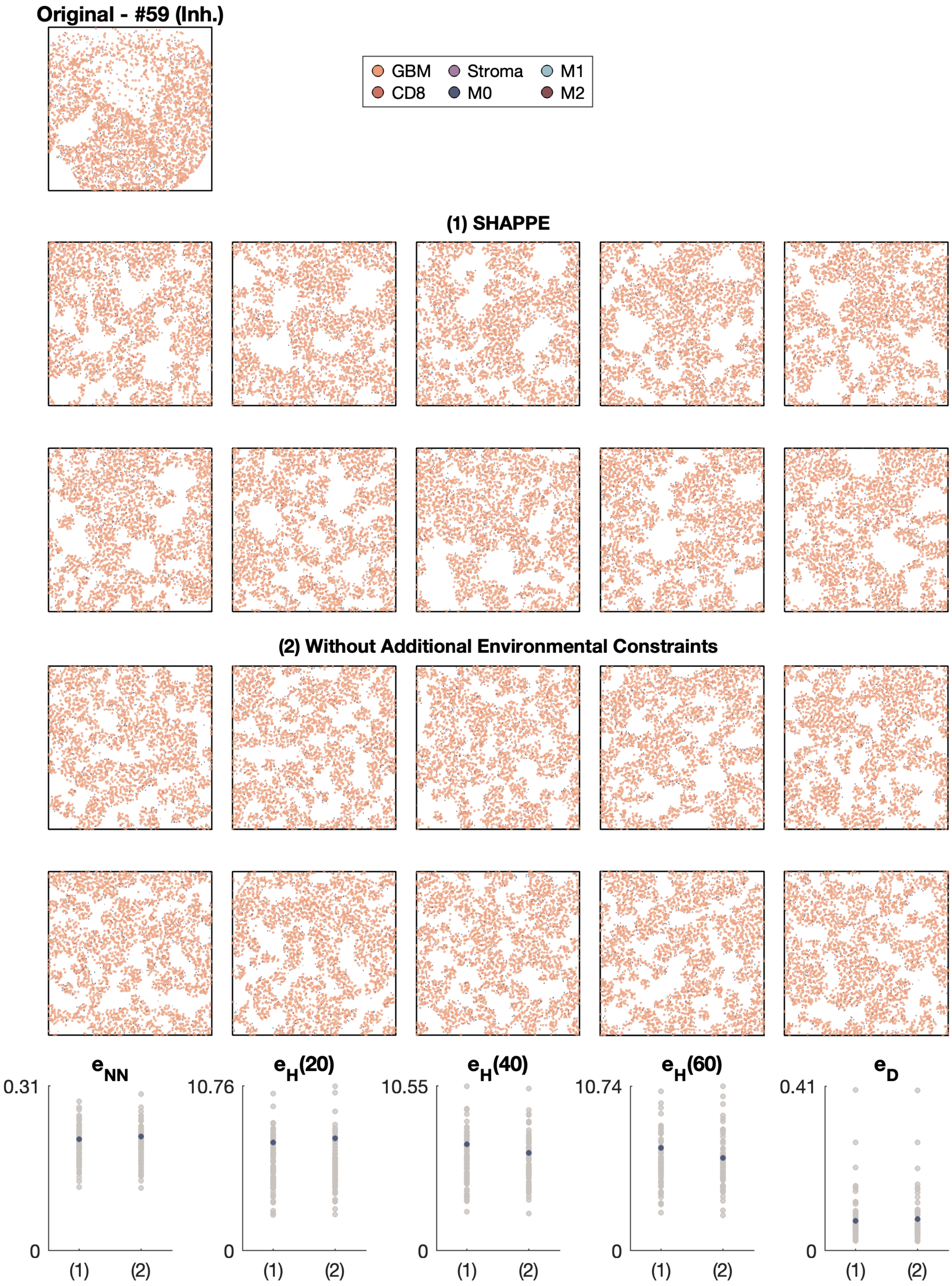

Original - #59 (Inh.)
GBM
Stroma
M1
CD8
M0
M2
(1) SHAPPE
(2) Without Additional Environmental Constraints
e_NN
0.31
0
(1)
(2)
e_H(20)
10.76
0
(1)
(2)
e_H(40)
10.55
0
(1)
(2)
e_H(60)
10.74
0
(1)
(2)
e_D
0.41
0
(1)
(2)

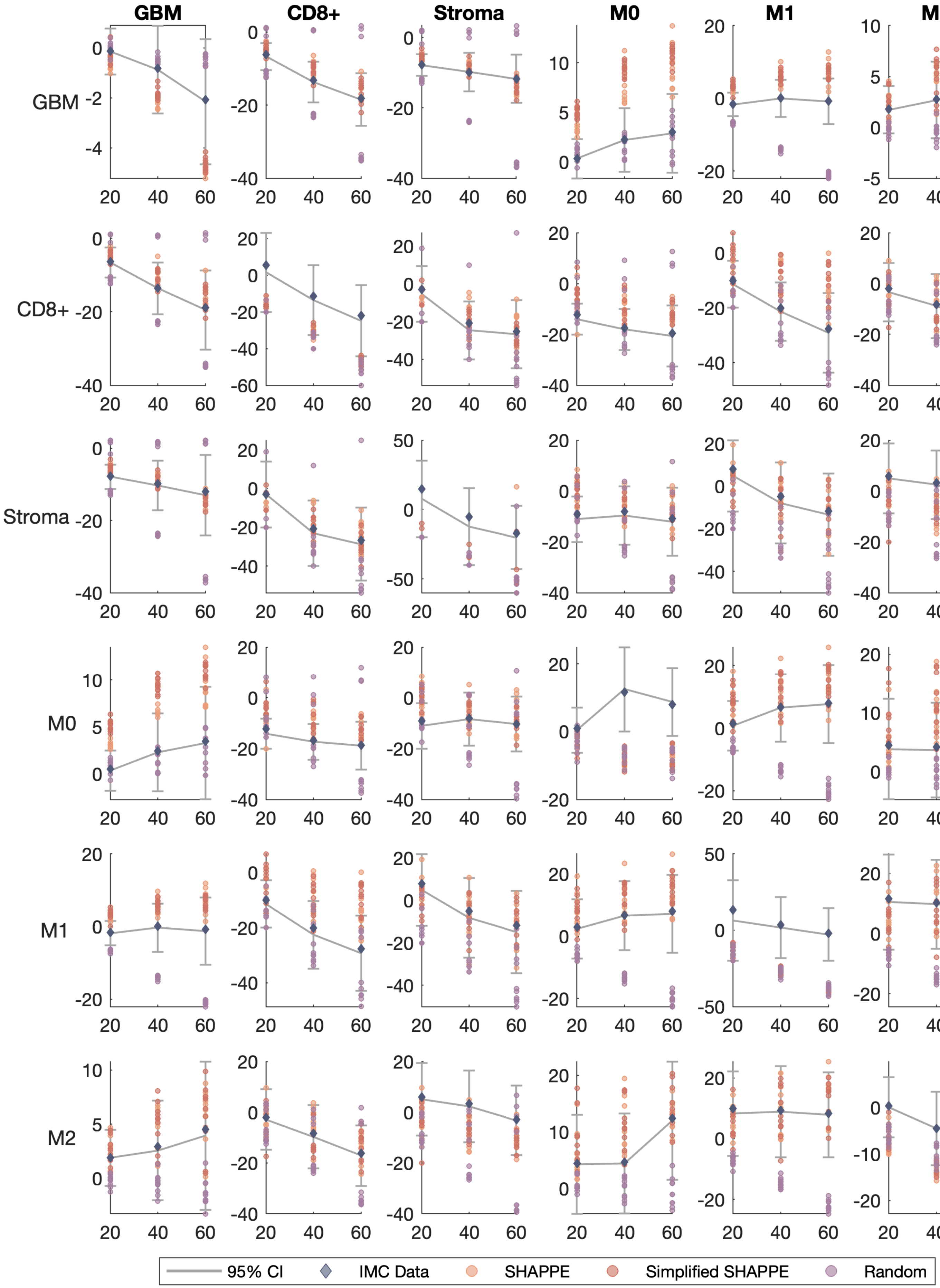

GBM
CD8+
Stroma
M0
M1
M2
95% CI
IMC Data
SHAPPE
Simplified SHAPPE
Random

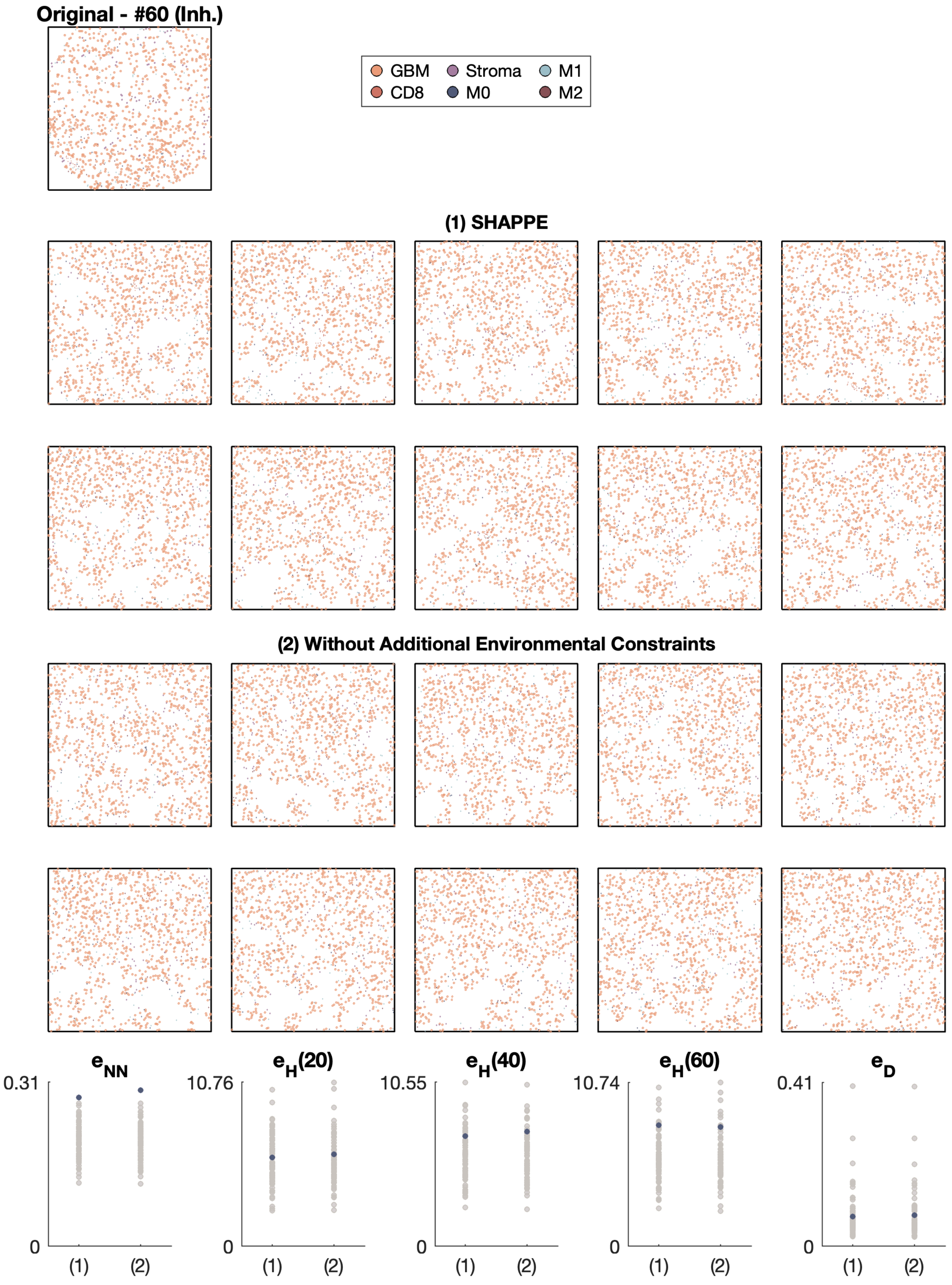

Original - #60 (Inh.)
GBM
CD8
Stroma
M0
M1
M2
(1) SHAPPE
(2) Without Additional Environmental Constraints
$e_{NN}$
0.31
0
$e_H(20)$
10.76
0
$e_H(40)$
10.55
0
$e_H(60)$
10.74
0
$e_D$
0.41
0
(1)
(2)

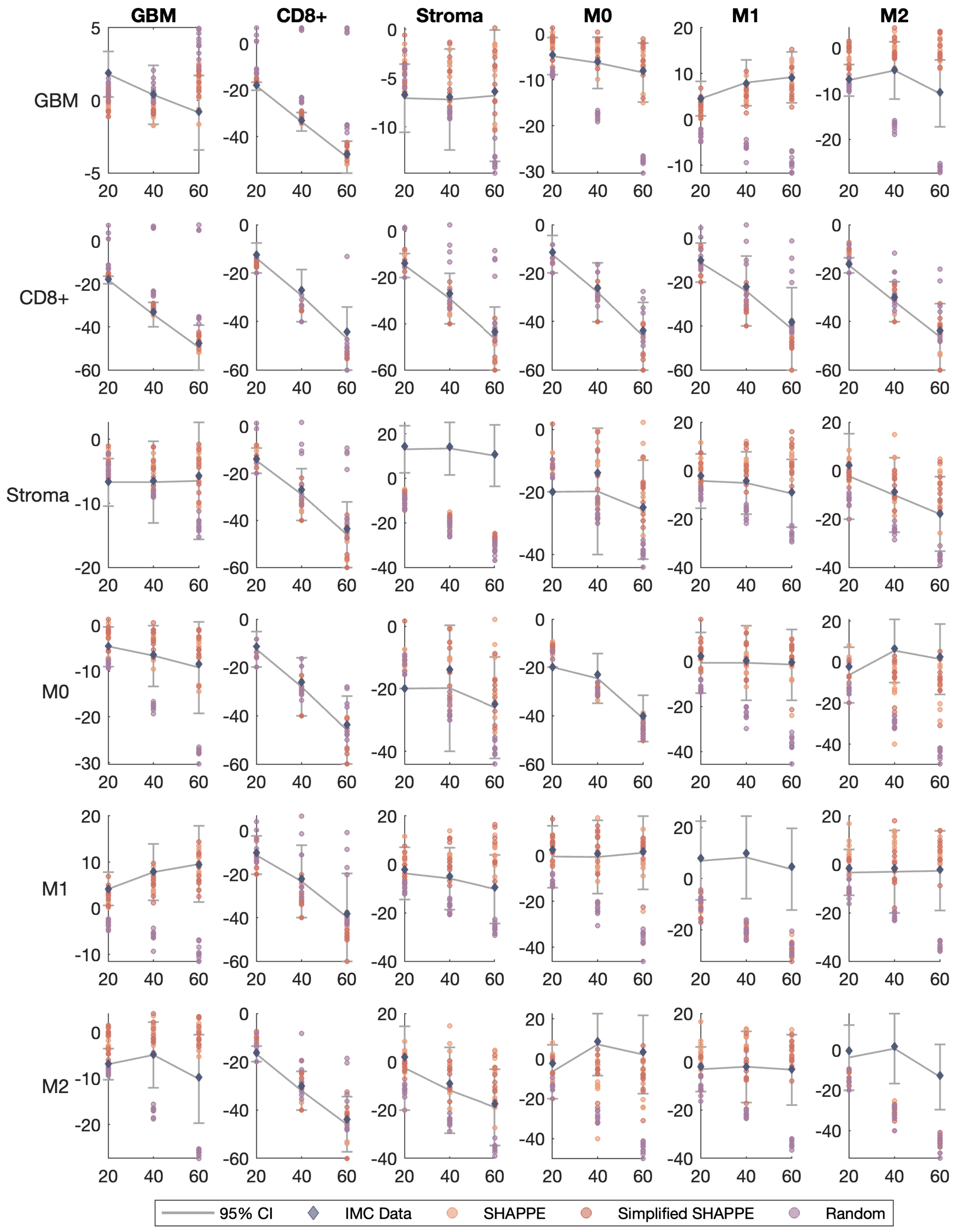
GBM
CD8+
Stroma
M0
M1
M2
95% CI
IMC Data
SHAPPE
Simplified SHAPPE
Random

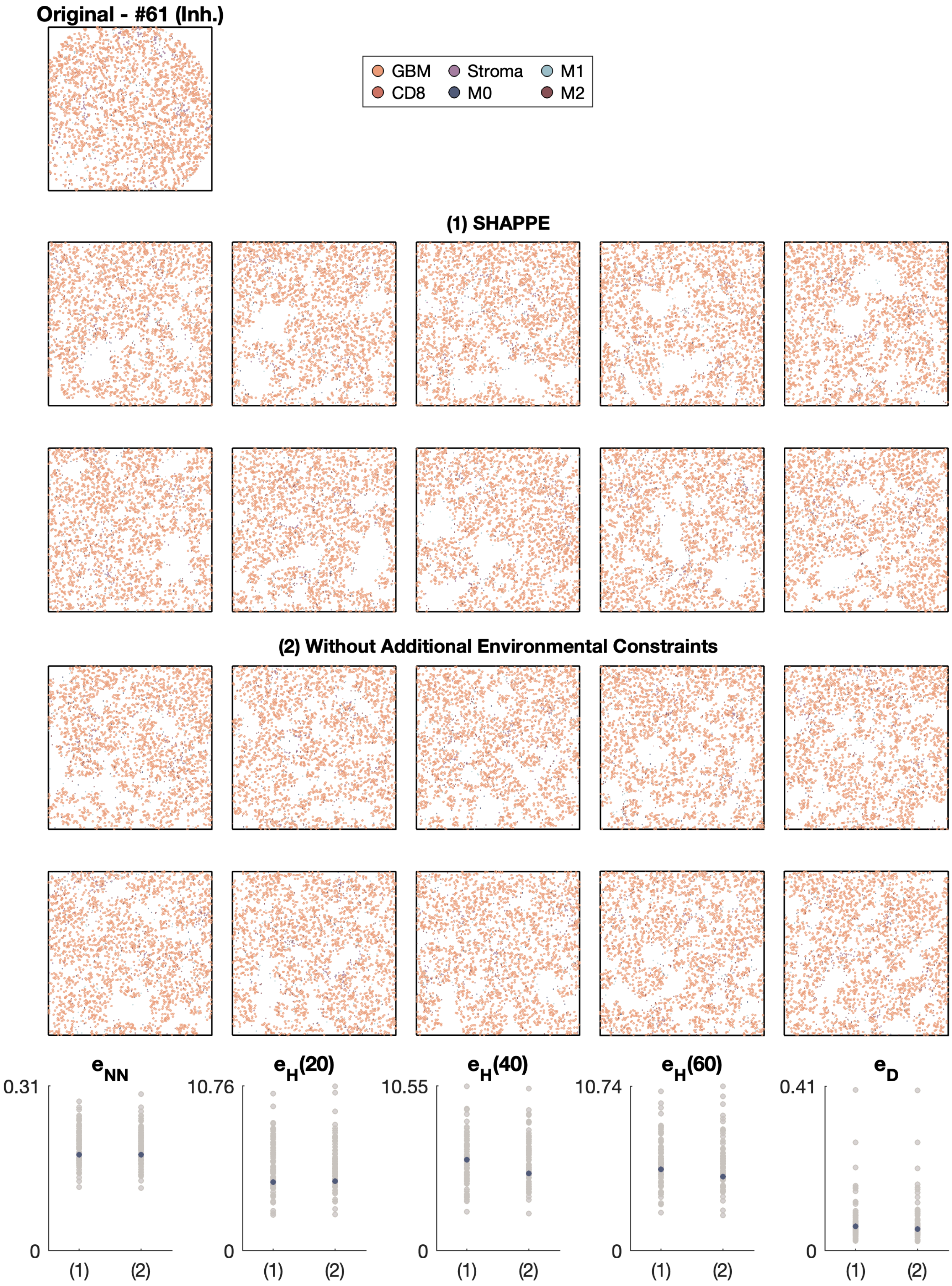
Original - #61 (Inh.)
GBM
Stroma
M1
CD8
M0
M2
(1) SHAPPE
(2) Without Additional Environmental Constraints
e_NN
e_H(20)
e_H(40)
e_H(60)
e_D
0.31
10.76
10.55
10.74
0.41
0
(1)
(2)

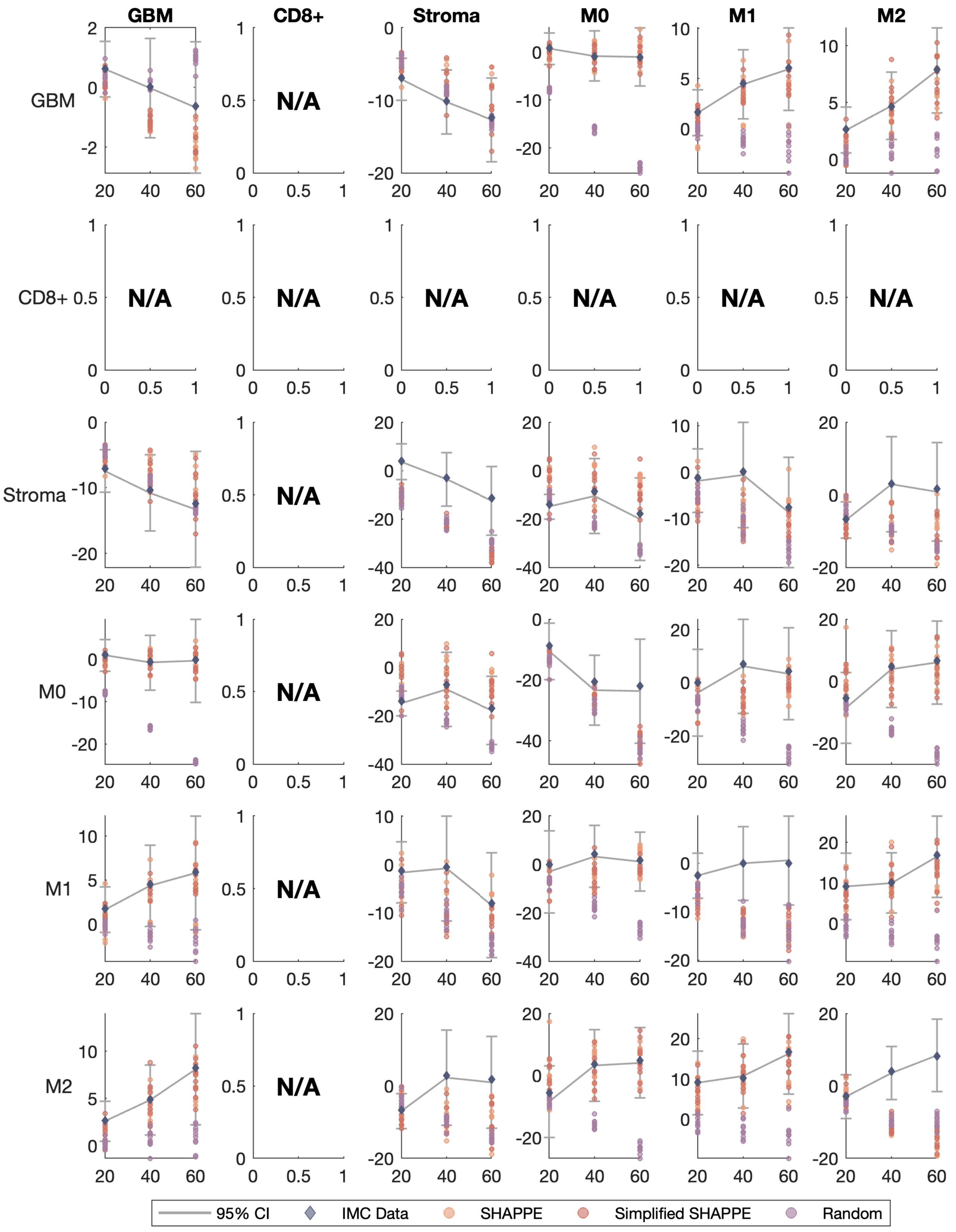
GBM
CD8+
Stroma
M0
M1
M2
GBM
CD8+
Stroma
M0
M1
M2
N/A
95% CI
IMC Data
SHAPPE
Simplified SHAPPE
Random

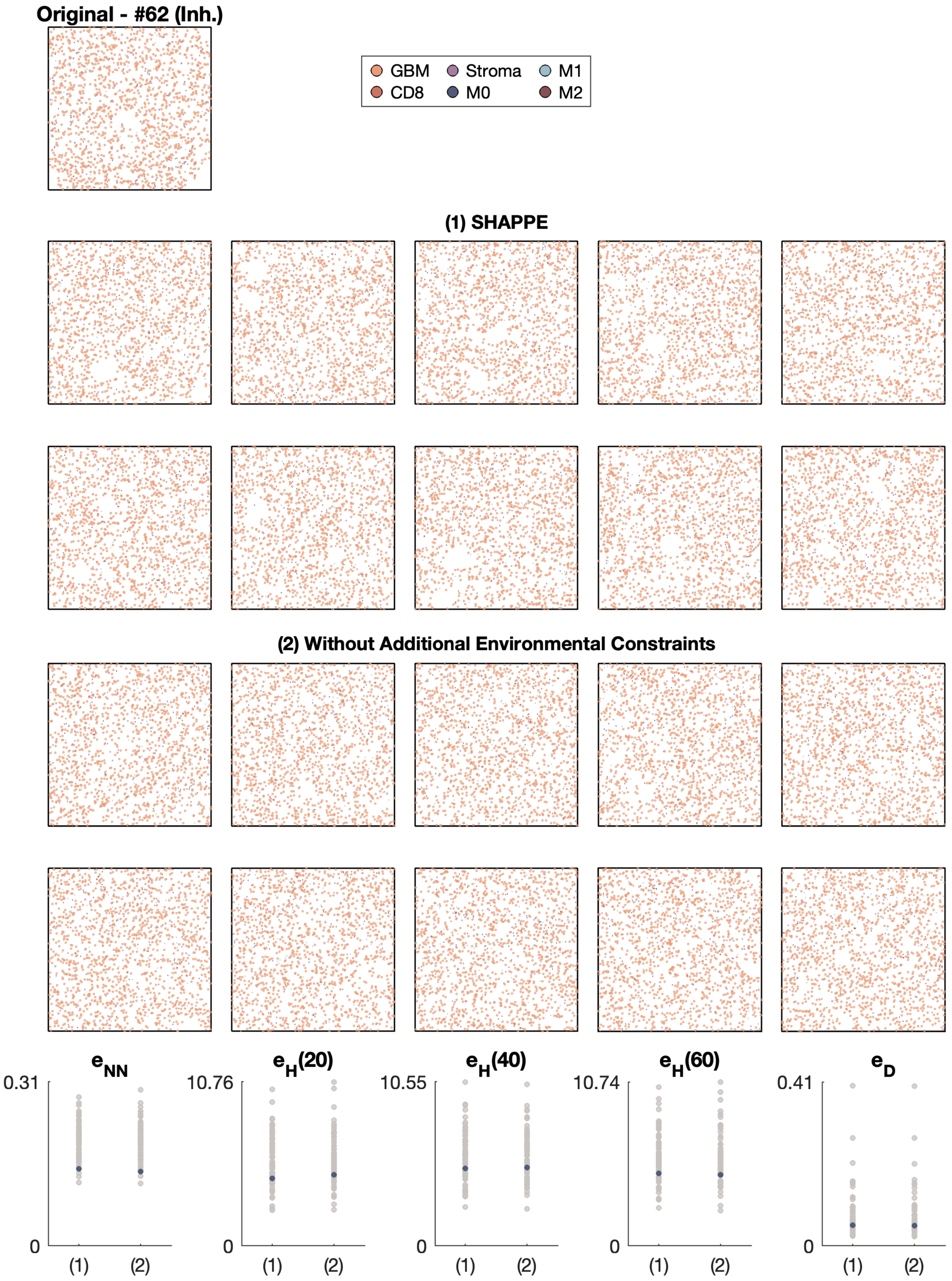
Original - #62 (Inh.)
GBM
CD8
Stroma
M0
M1
M2
(1) SHAPPE
(2) Without Additional Environmental Constraints
e_NN
0.31
0
e_H(20)
10.76
e_H(40)
10.55
e_H(60)
10.74
e_D
0.41
(1)
(2)

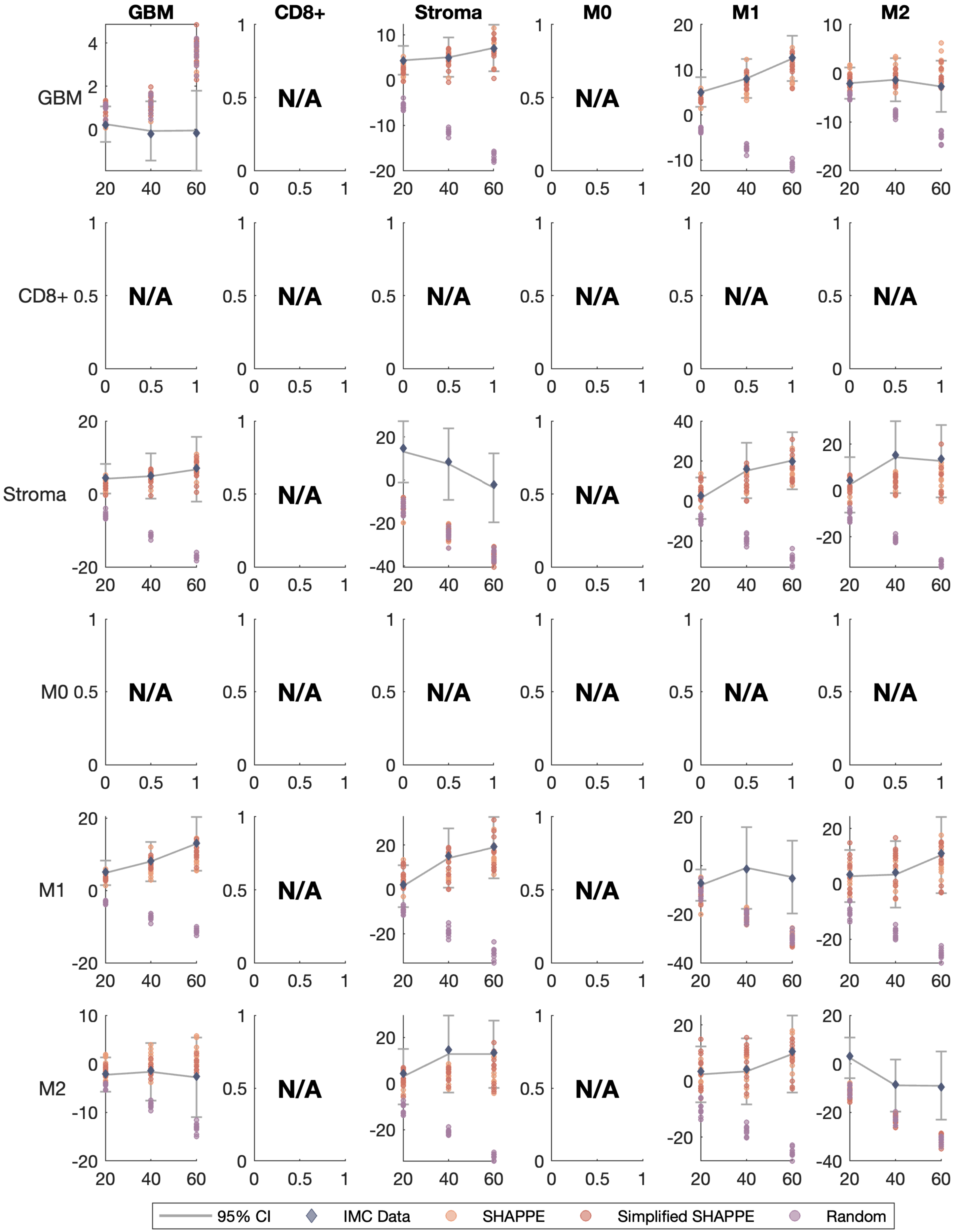

GBM
CD8+
Stroma
M0
M1
M2
GBM
CD8+
Stroma
M0
M1
M2
N/A
95% CI
IMC Data
SHAPPE
Simplified SHAPPE
Random

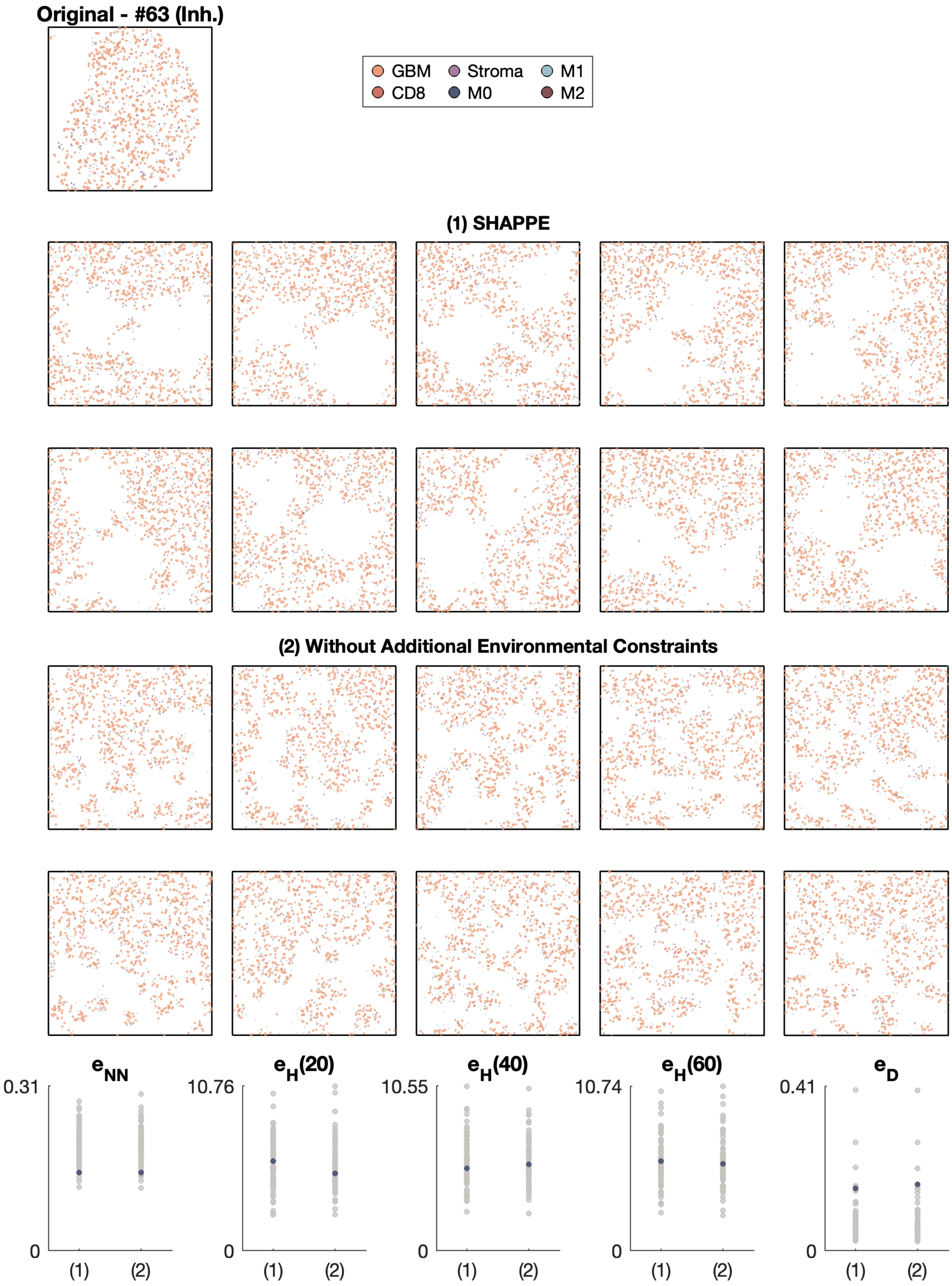

Original - #63 (Inh.)
GBM
Stroma
M1
CD8
M0
M2
(1) SHAPPE
(2) Without Additional Environmental Constraints
e_NN
0.31
0
(1)
(2)
e_H(20)
10.76
0
(1)
(2)
e_H(40)
10.55
0
(1)
(2)
e_H(60)
10.74
0
(1)
(2)
e_D
0.41
0
(1)
(2)

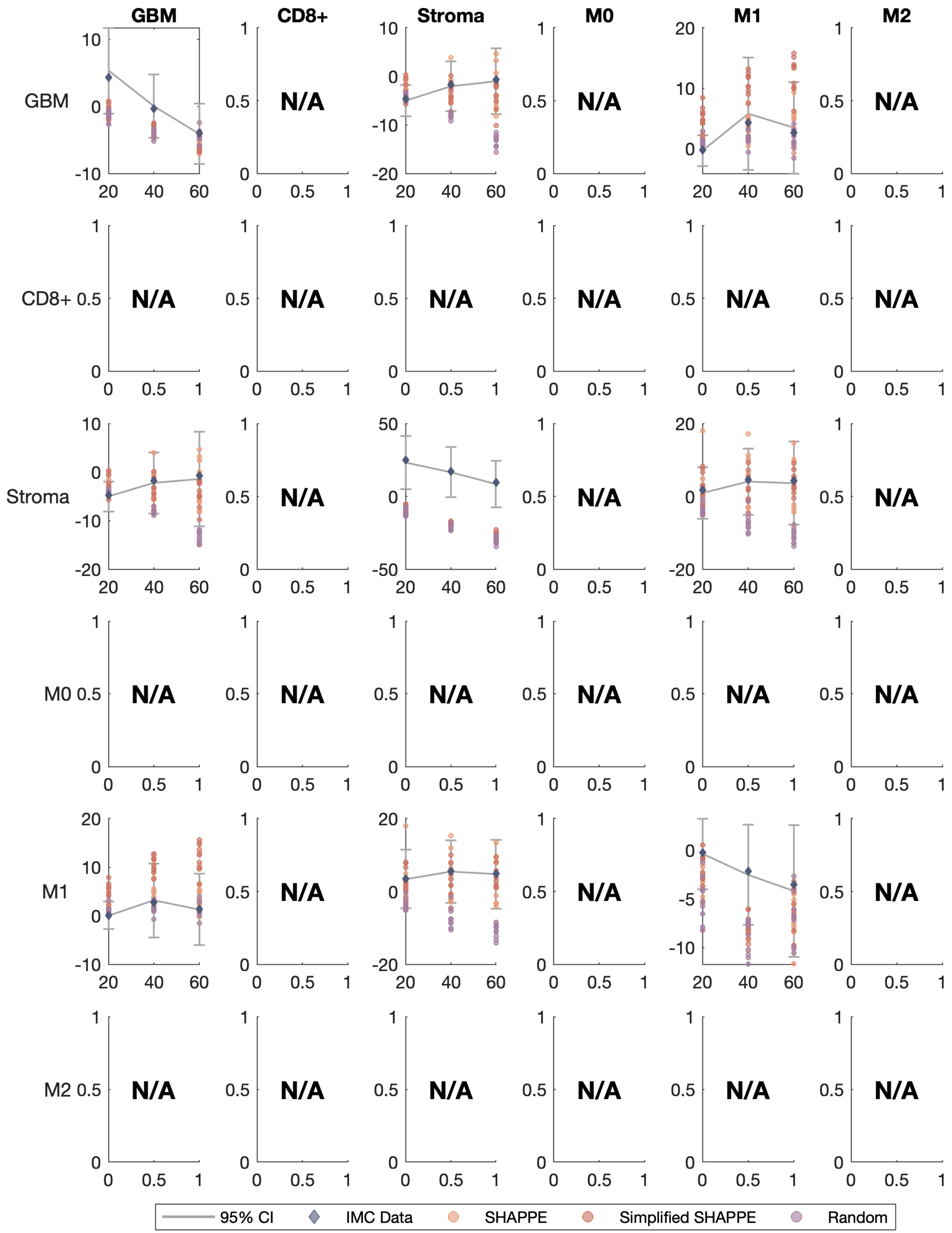
GBM
CD8+
Stroma
M0
M1
M2
GBM
CD8+
Stroma
M0
M1
M2
N/A
95% CI
IMC Data
SHAPPE
Simplified SHAPPE
Random

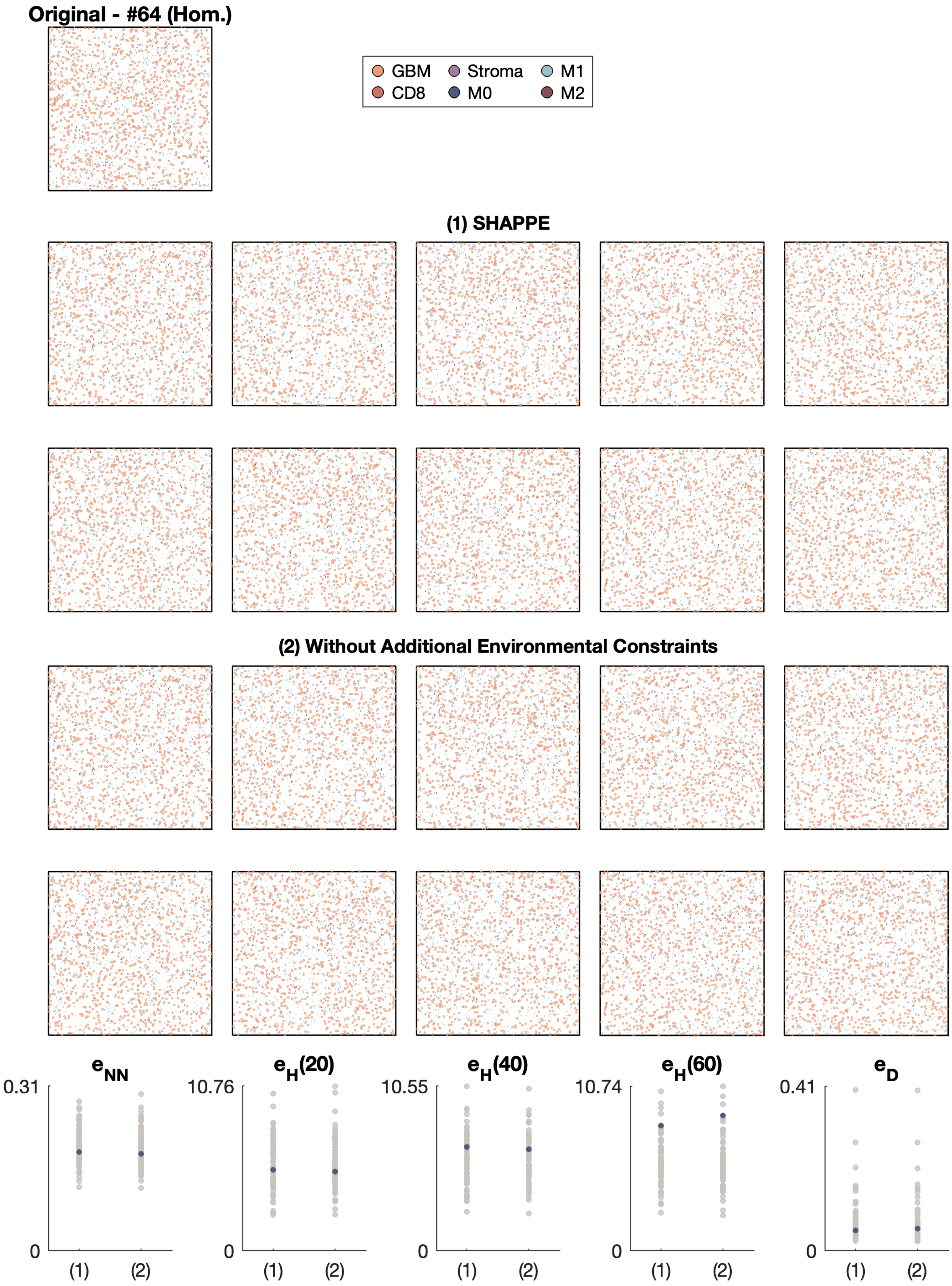
Original - #64 (Hom.)
GBM
CD8
Stroma
M0
M1
M2
(1) SHAPPE
(2) Without Additional Environmental Constraints
$e_{NN}$
0.31
0
(1)
(2)
$e_H(20)$
10.76
0
(1)
(2)
$e_H(40)$
10.55
0
(1)
(2)
$e_H(60)$
10.74
0
(1)
(2)
$e_D$
0.41
0
(1)
(2)

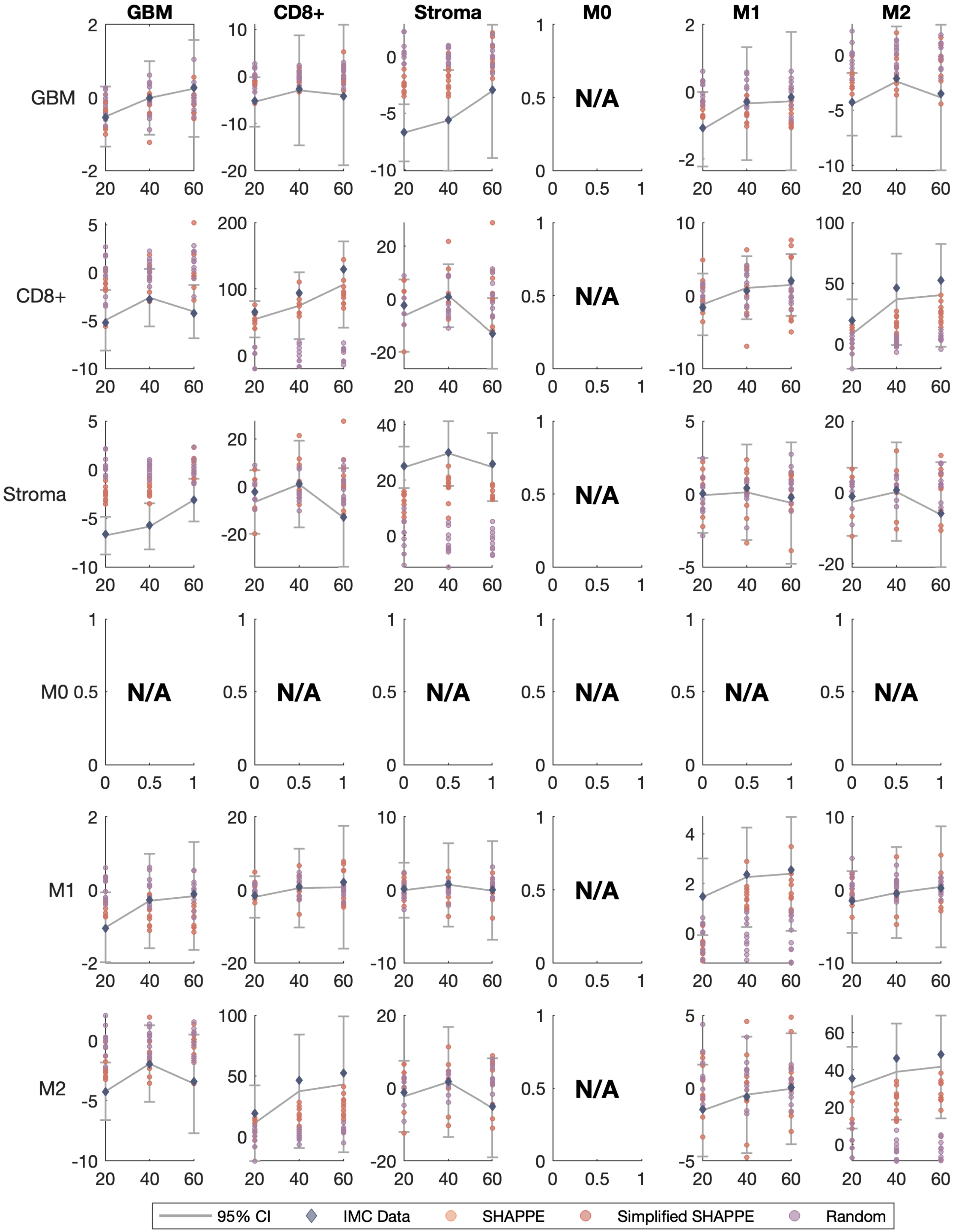
GBM
CD8+
Stroma
M0
M1
M2
N/A
95% CI
IMC Data
SHAPPE
Simplified SHAPPE
Random

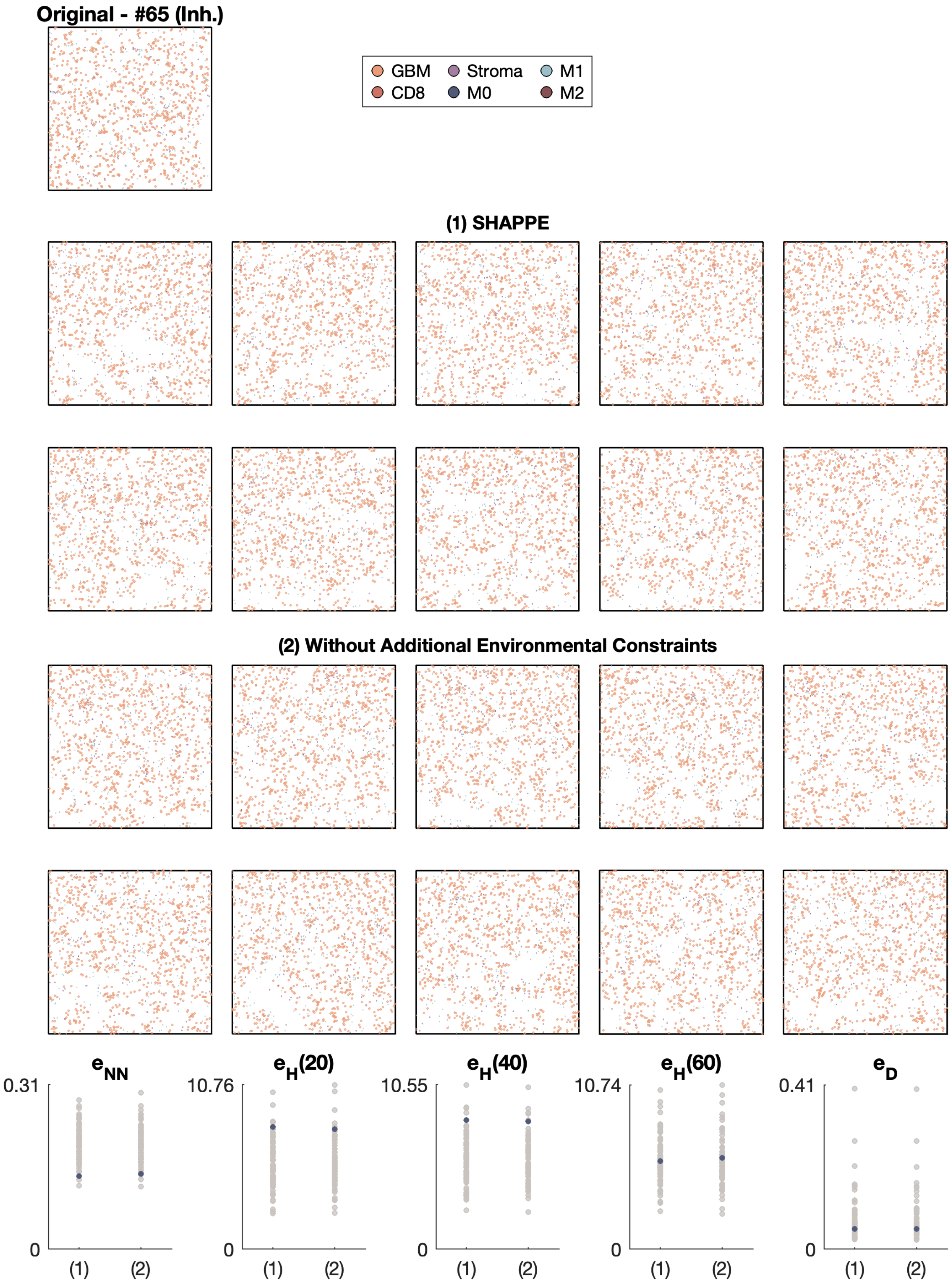

Original - #65 (Inh.)
GBM
Stroma
M1
CD8
M0
M2
(1) SHAPPE
(2) Without Additional Environmental Constraints
$e_{NN}$
$e_H(20)$
$e_H(40)$
$e_H(60)$
$e_D$
0.31
10.76
10.55
10.74
0.41
0
(1)
(2)

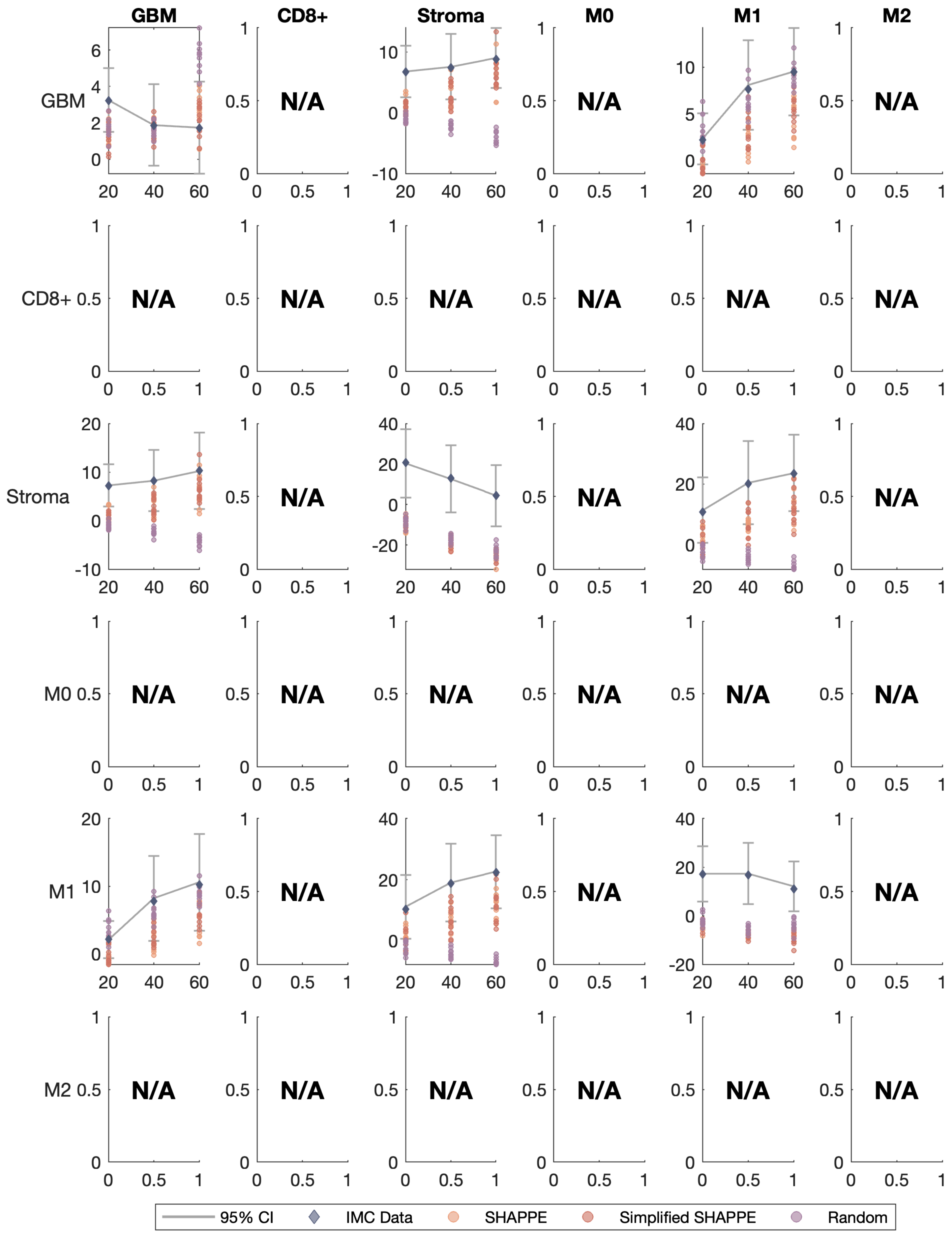

GBM
CD8+
Stroma
M0
M1
M2
N/A
95% CI
IMC Data
SHAPPE
Simplified SHAPPE
Random

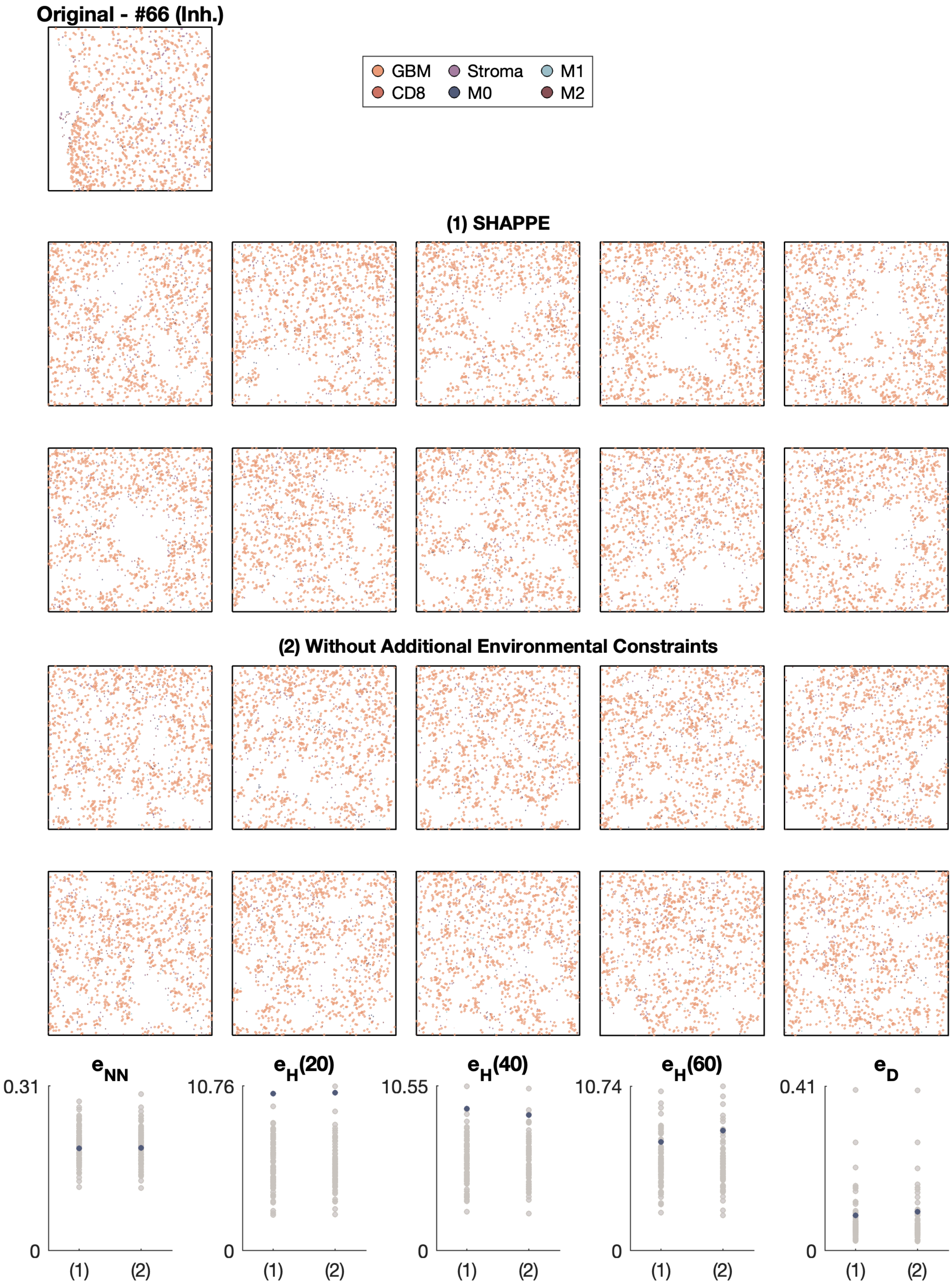

Original - #66 (Inh.)
GBM
CD8
Stroma
M0
M1
M2
(1) SHAPPE
(2) Without Additional Environmental Constraints
$e_{NN}$
$e_H(20)$
$e_H(40)$
$e_H(60)$
$e_D$
0.31
10.76
10.55
10.74
0.41
0
(1)
(2)

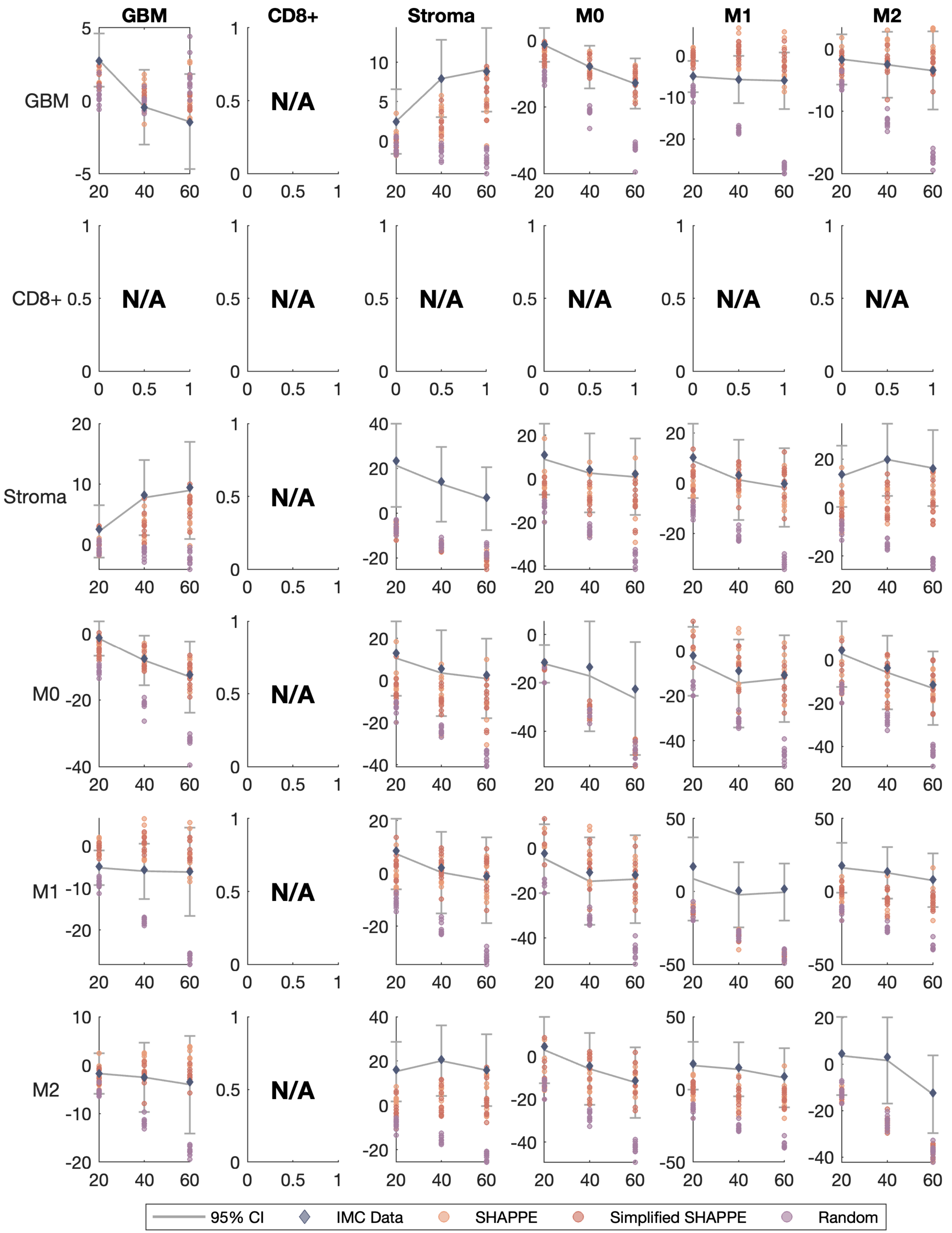

GBM
CD8+
Stroma
M0
M1
M2
N/A
95% CI
IMC Data
SHAPPE
Simplified SHAPPE
Random

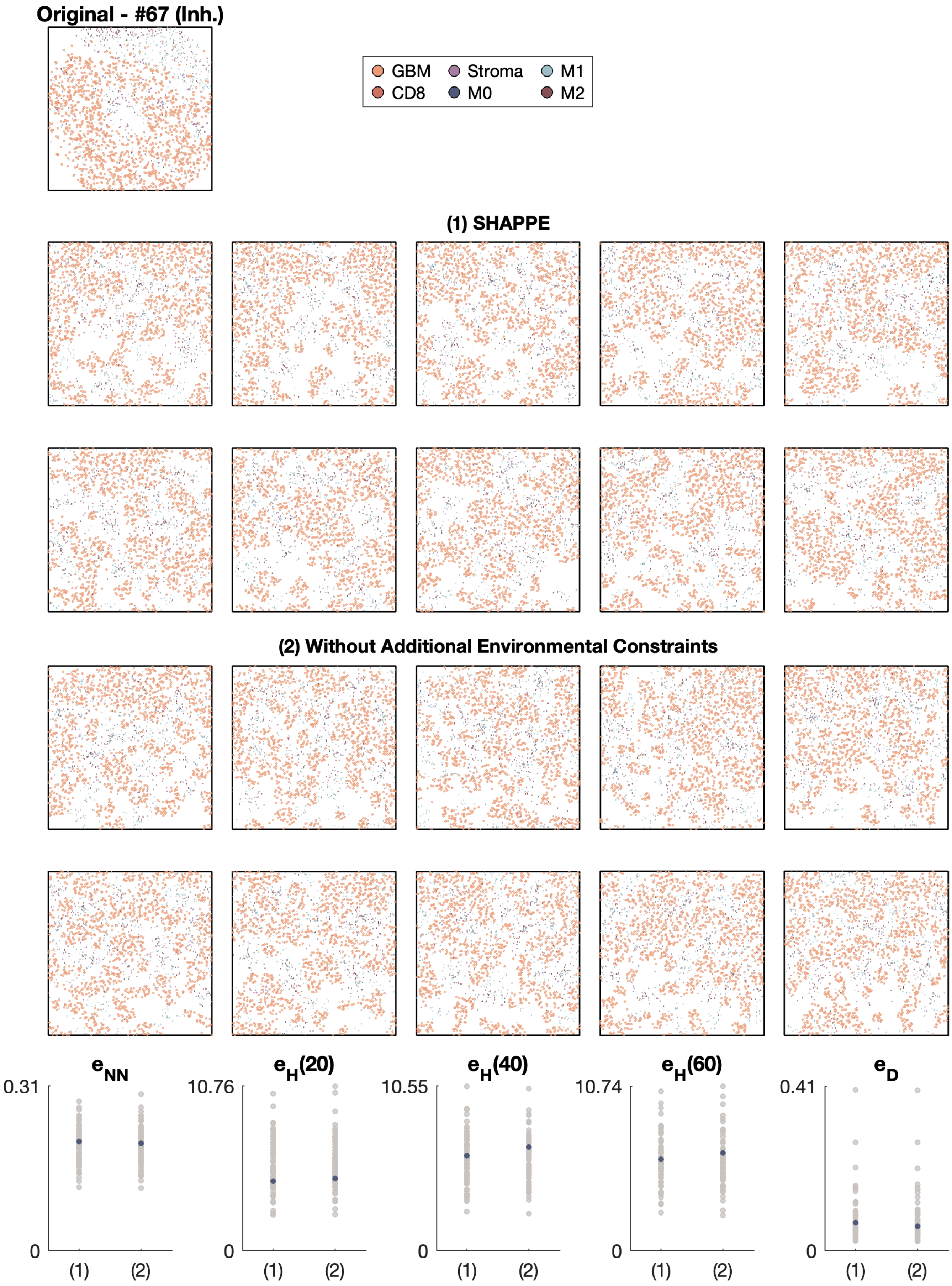

Original - #67 (Inh.)
GBM
Stroma
M1
CD8
M0
M2
(1) SHAPPE
(2) Without Additional Environmental Constraints
$e_{NN}$
$e_H(20)$
$e_H(40)$
$e_H(60)$
$e_D$
0.31
10.76
10.55
10.74
0.41
0
(1)
(2)

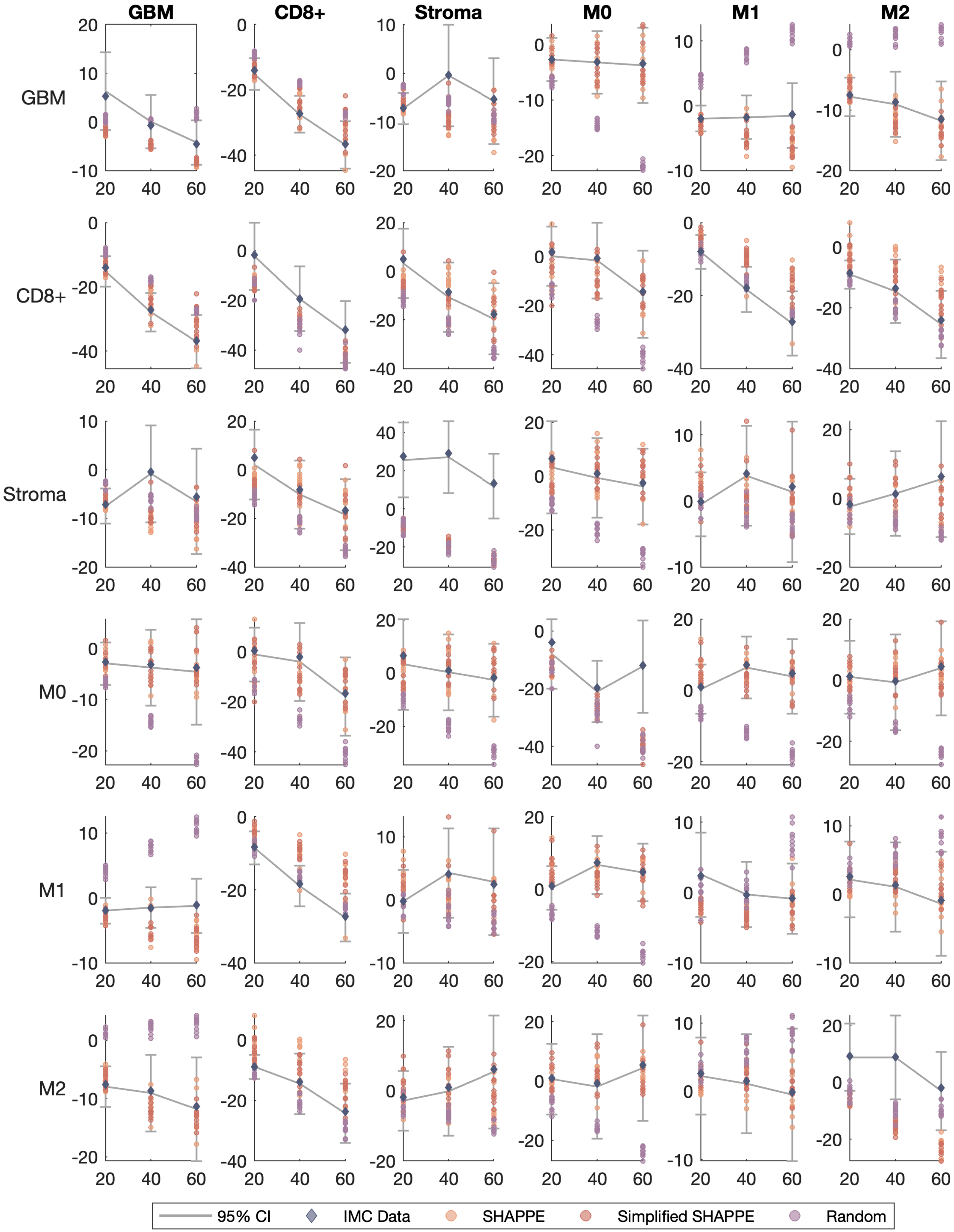
GBM
CD8+
Stroma
M0
M1
M2
95% CI
IMC Data
SHAPPE
Simplified SHAPPE
Random

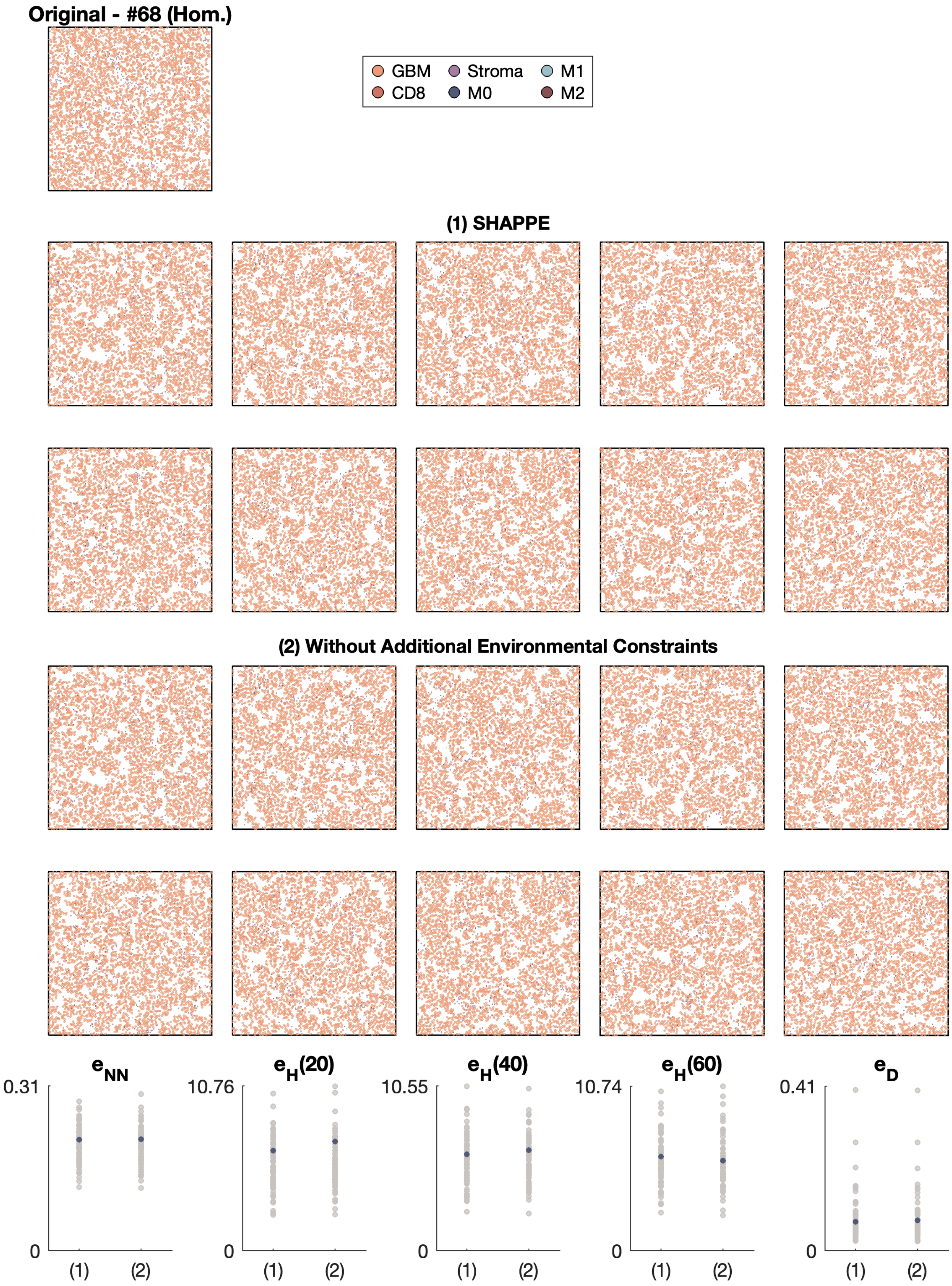
Original - #68 (Hom.)
GBM
Stroma
M1
CD8
M0
M2
(1) SHAPPE
(2) Without Additional Environmental Constraints
$e_{NN}$
0.31
0
(1)
(2)
$e_H(20)$
10.76
0
(1)
(2)
$e_H(40)$
10.55
0
(1)
(2)
$e_H(60)$
10.74
0
(1)
(2)
$e_D$
0.41
0
(1)
(2)

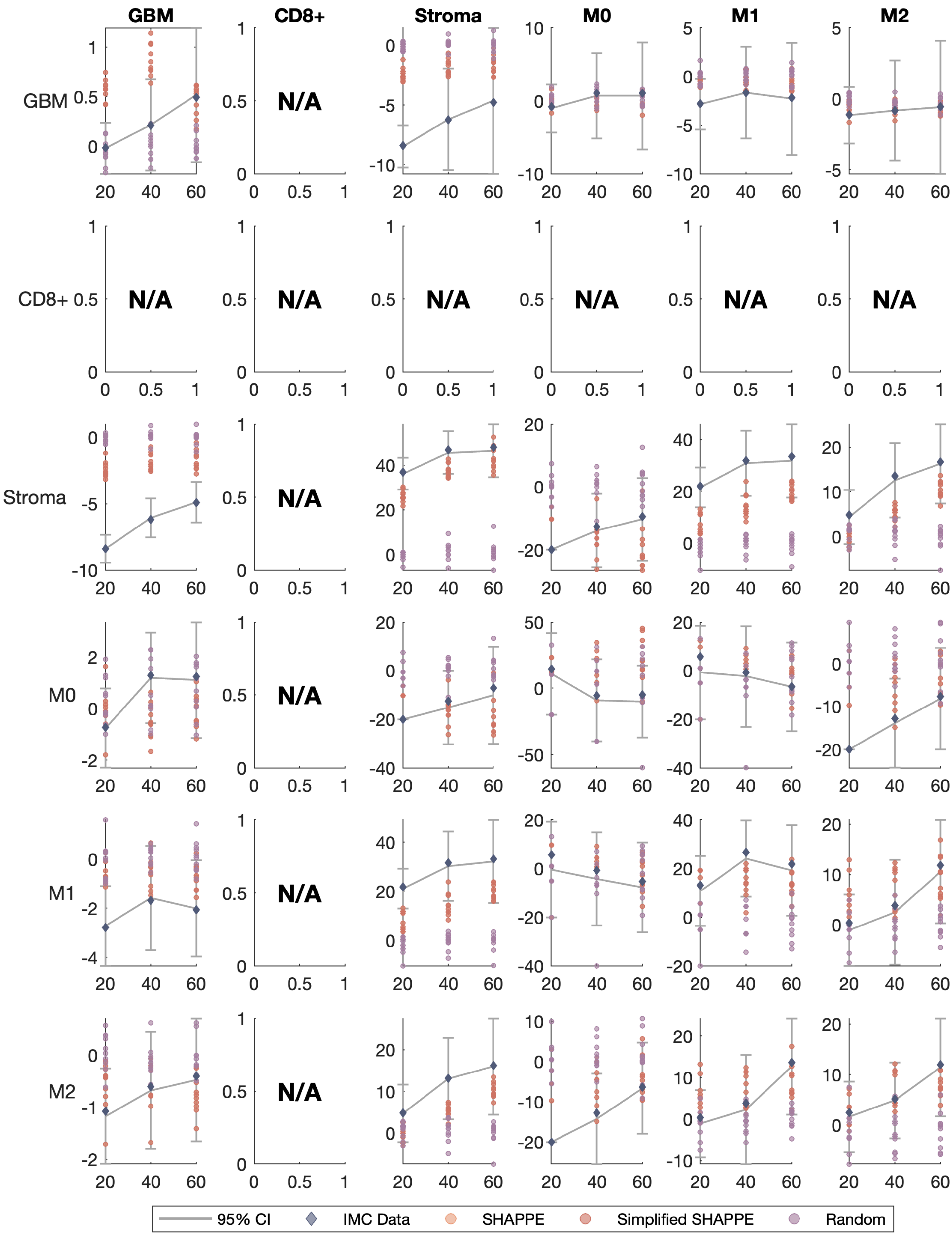
GBM
CD8+
Stroma
M0
M1
M2
N/A
95% CI
IMC Data
SHAPPE
Simplified SHAPPE
Random

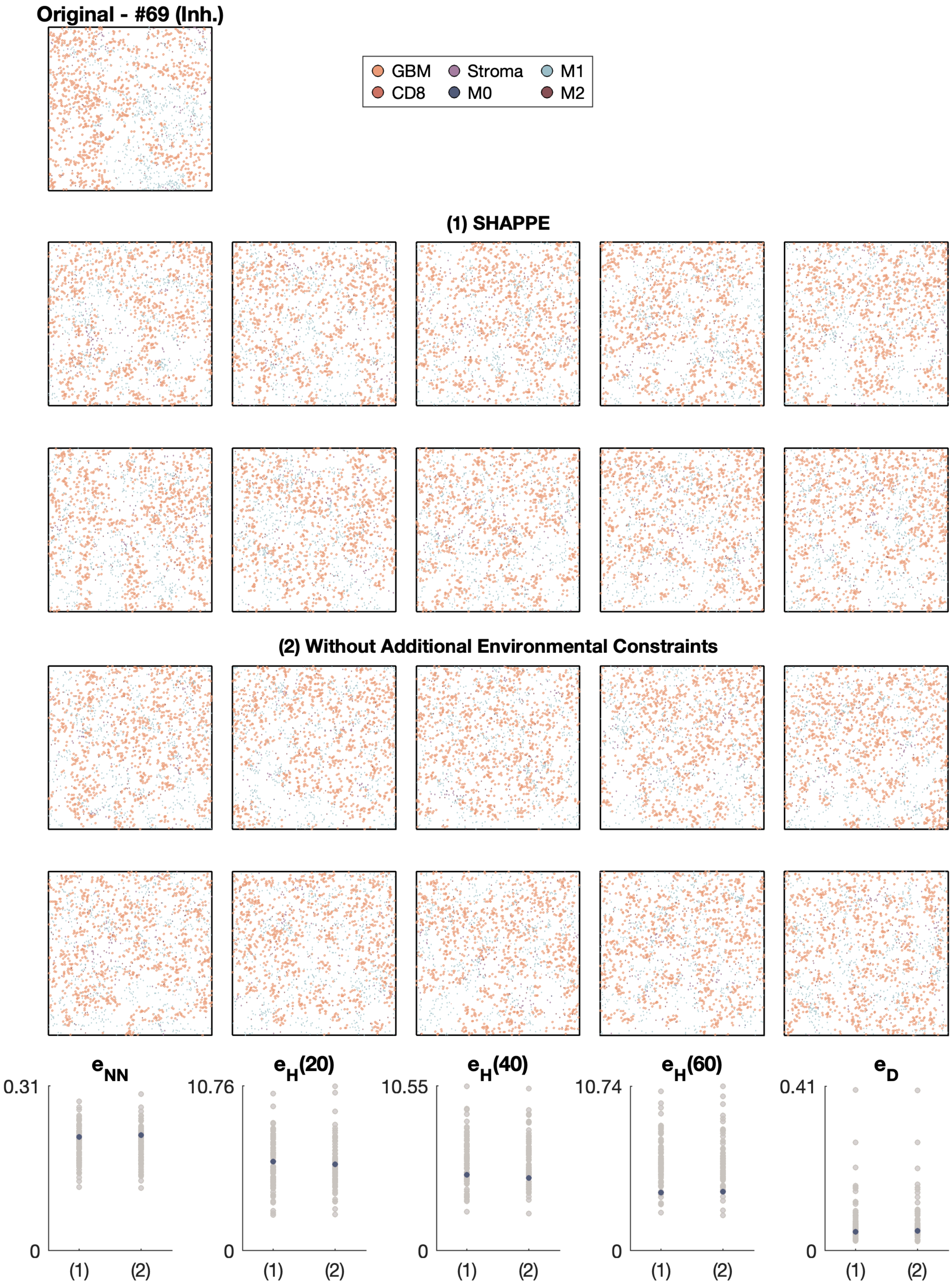

Original - #69 (Inh.)
GBM
CD8
Stroma
M0
M1
M2
(1) SHAPPE
(2) Without Additional Environmental Constraints
$e_{NN}$
$e_H(20)$
$e_H(40)$
$e_H(60)$
$e_D$
0.31
10.76
10.55
10.74
0.41
0
(1)
(2)

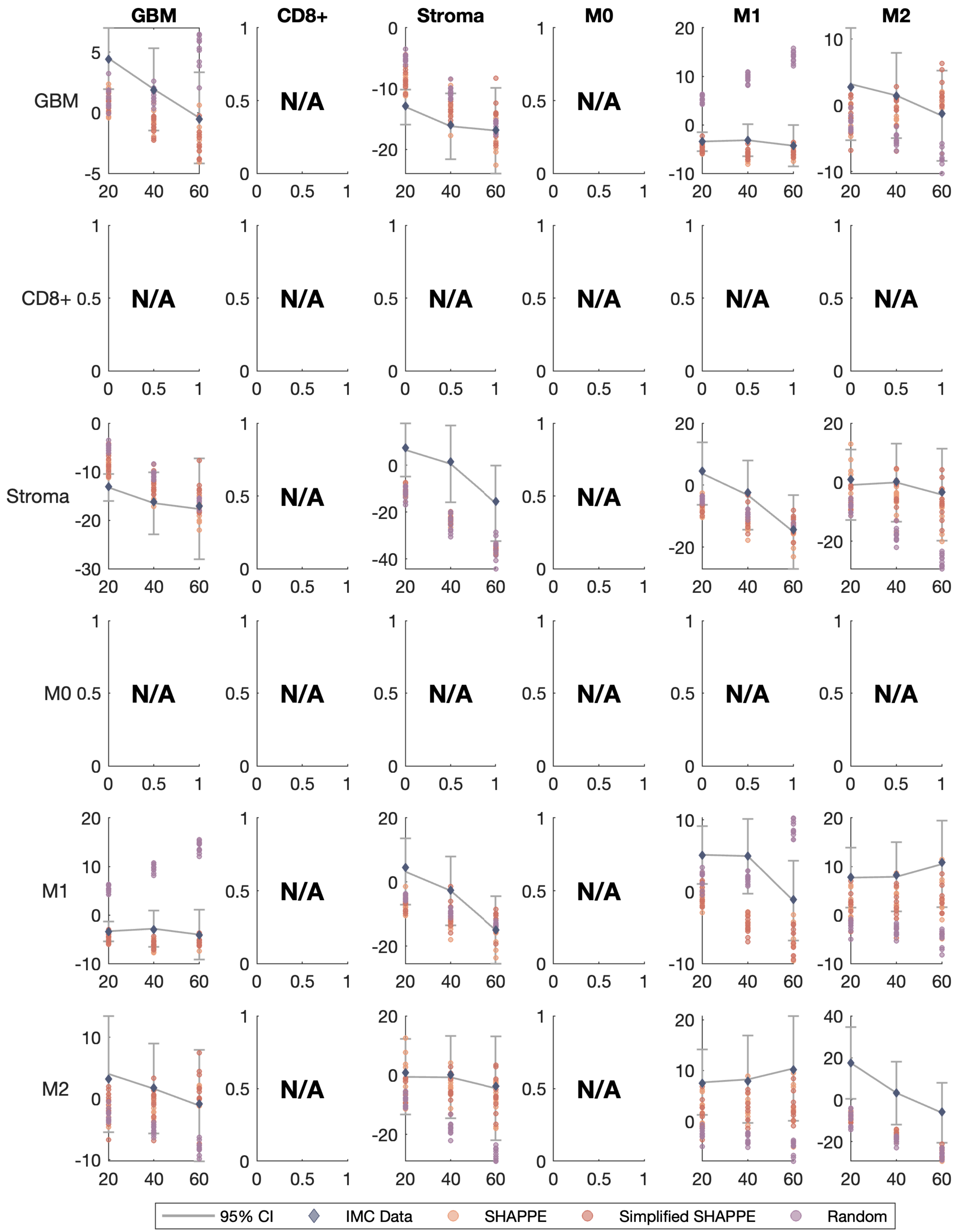
GBM
CD8+
Stroma
M0
M1
M2
N/A
95% CI
IMC Data
SHAPPE
Simplified SHAPPE
Random

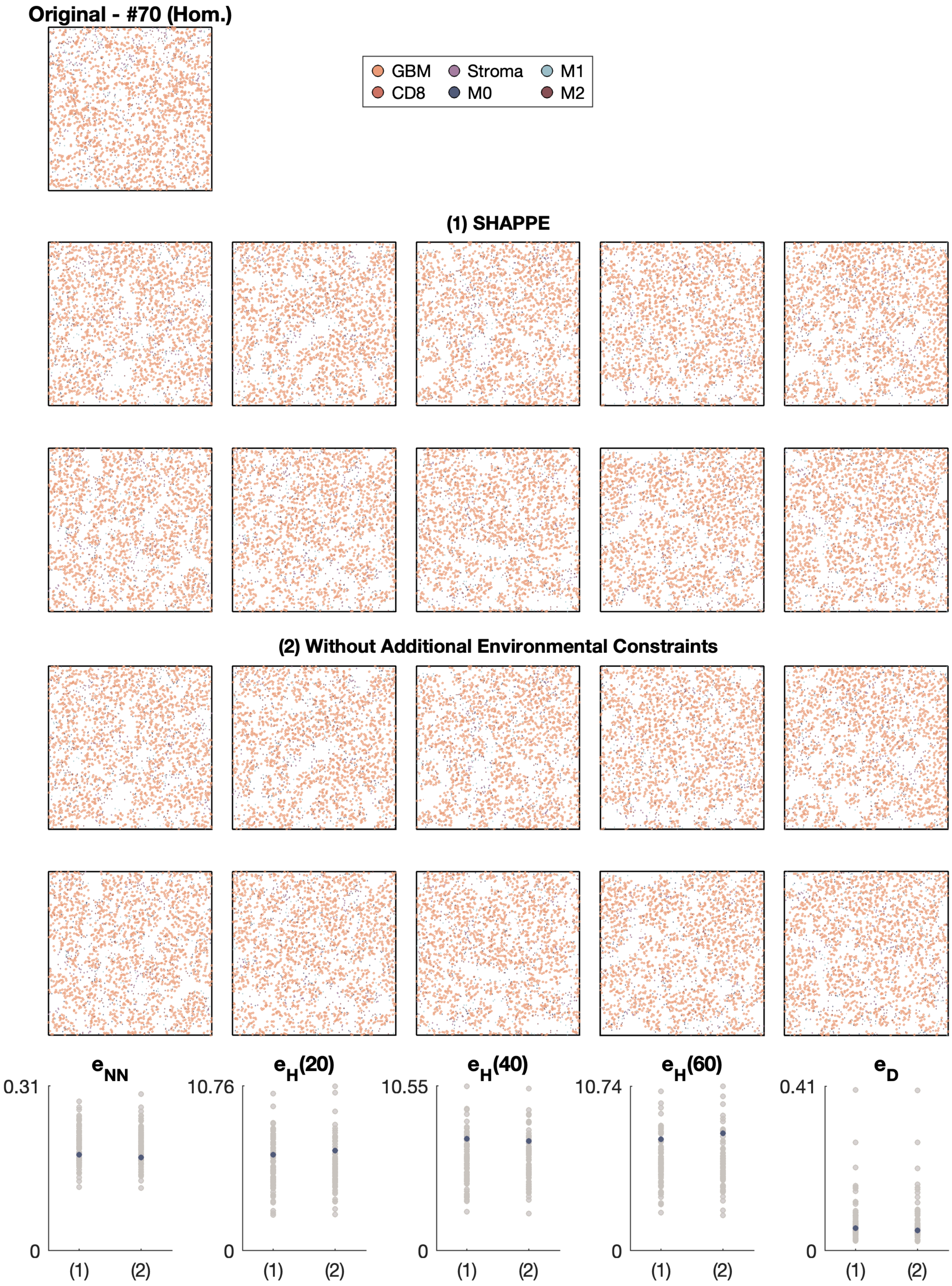

Original - #70 (Hom.)
GBM
Stroma
M1
CD8
M0
M2
(1) SHAPPE
(2) Without Additional Environmental Constraints
$e_{NN}$
$e_H(20)$
$e_H(40)$
$e_H(60)$
$e_D$
0.31
10.76
10.55
10.74
0.41
0
(1)
(2)

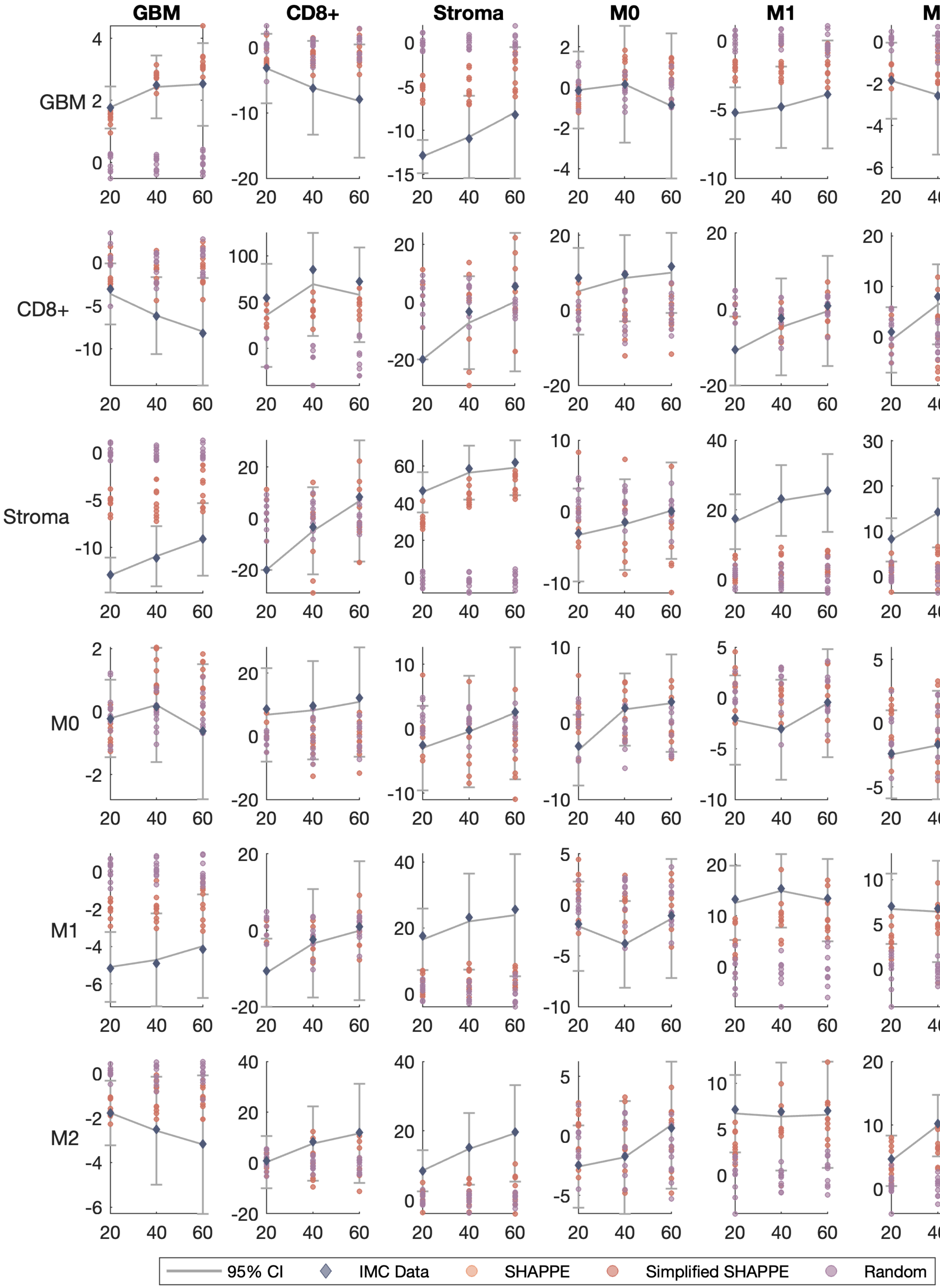
GBM
CD8+
Stroma
M0
M1
M2
GBM
CD8+
Stroma
M0
M1
M2
95% CI
IMC Data
SHAPPE
Simplified SHAPPE
Random

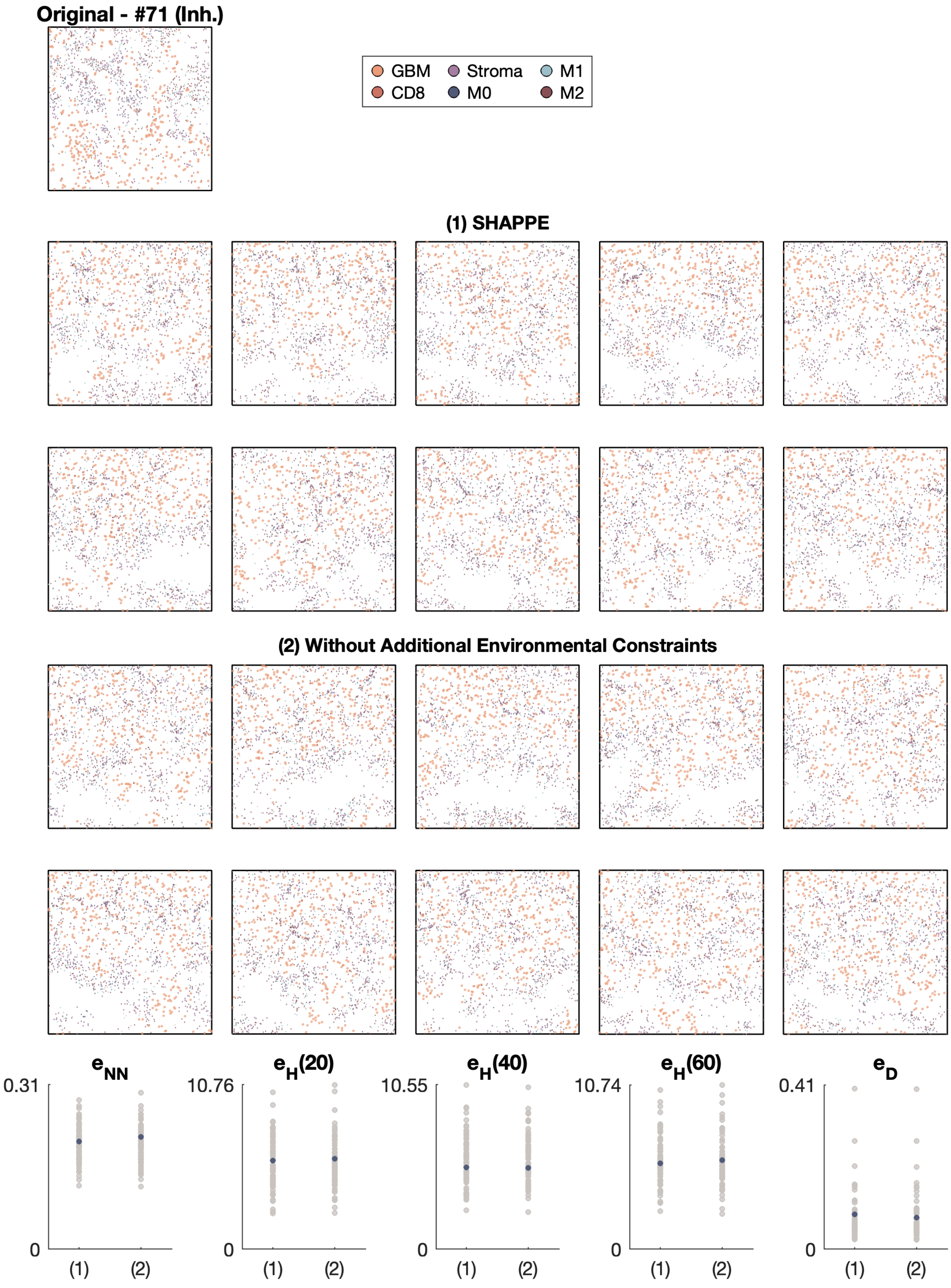

Original - #71 (Inh.)
GBM
Stroma
M1
CD8
M0
M2
(1) SHAPPE
(2) Without Additional Environmental Constraints
e_NN
e_H(20)
e_H(40)
e_H(60)
e_D
0.31
10.76
10.55
10.74
0.41
0
(1)
(2)

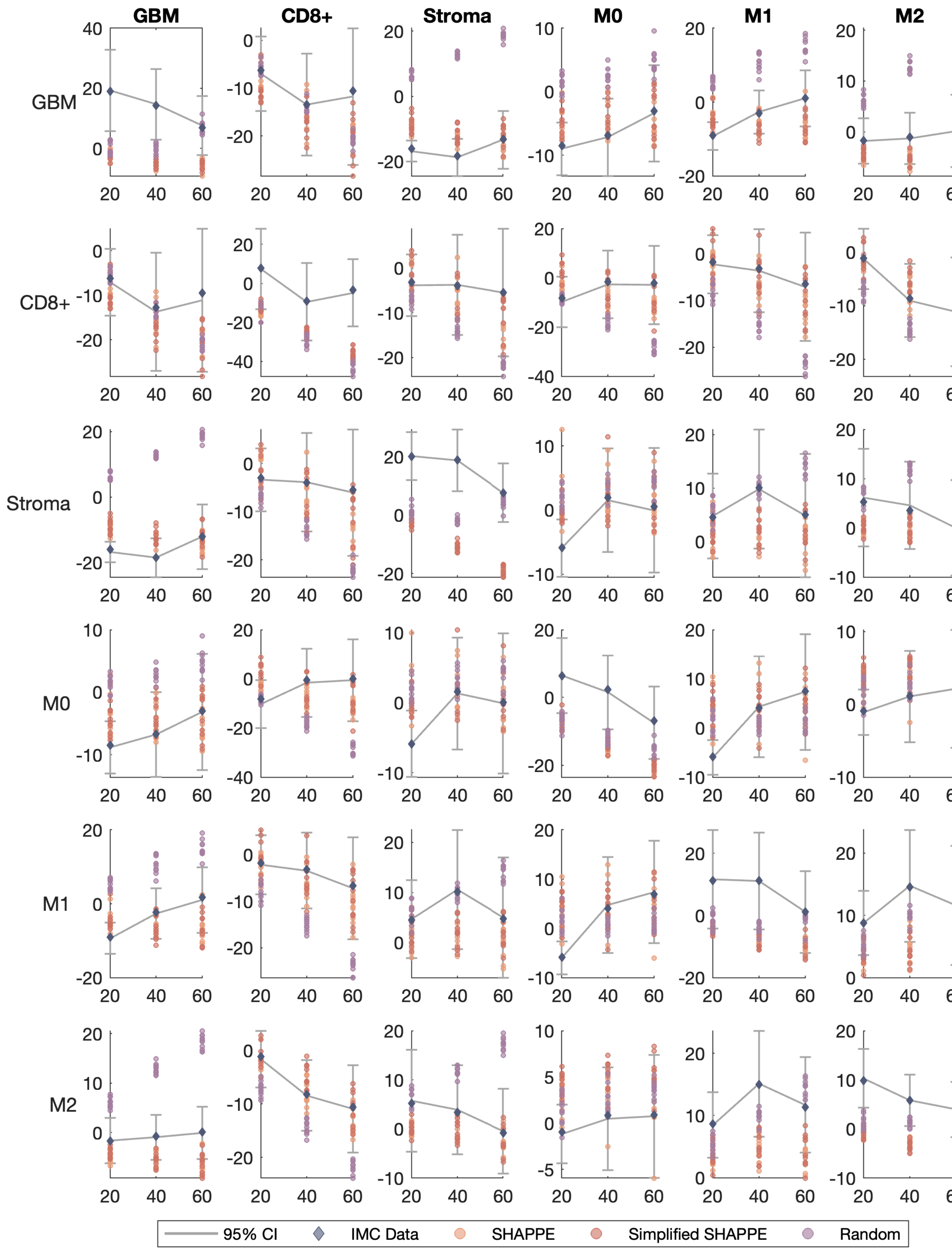

GBM
CD8+
Stroma
M0
M1
M2
GBM
CD8+
Stroma
M0
M1
M2
95% CI
IMC Data
SHAPPE
Simplified SHAPPE
Random

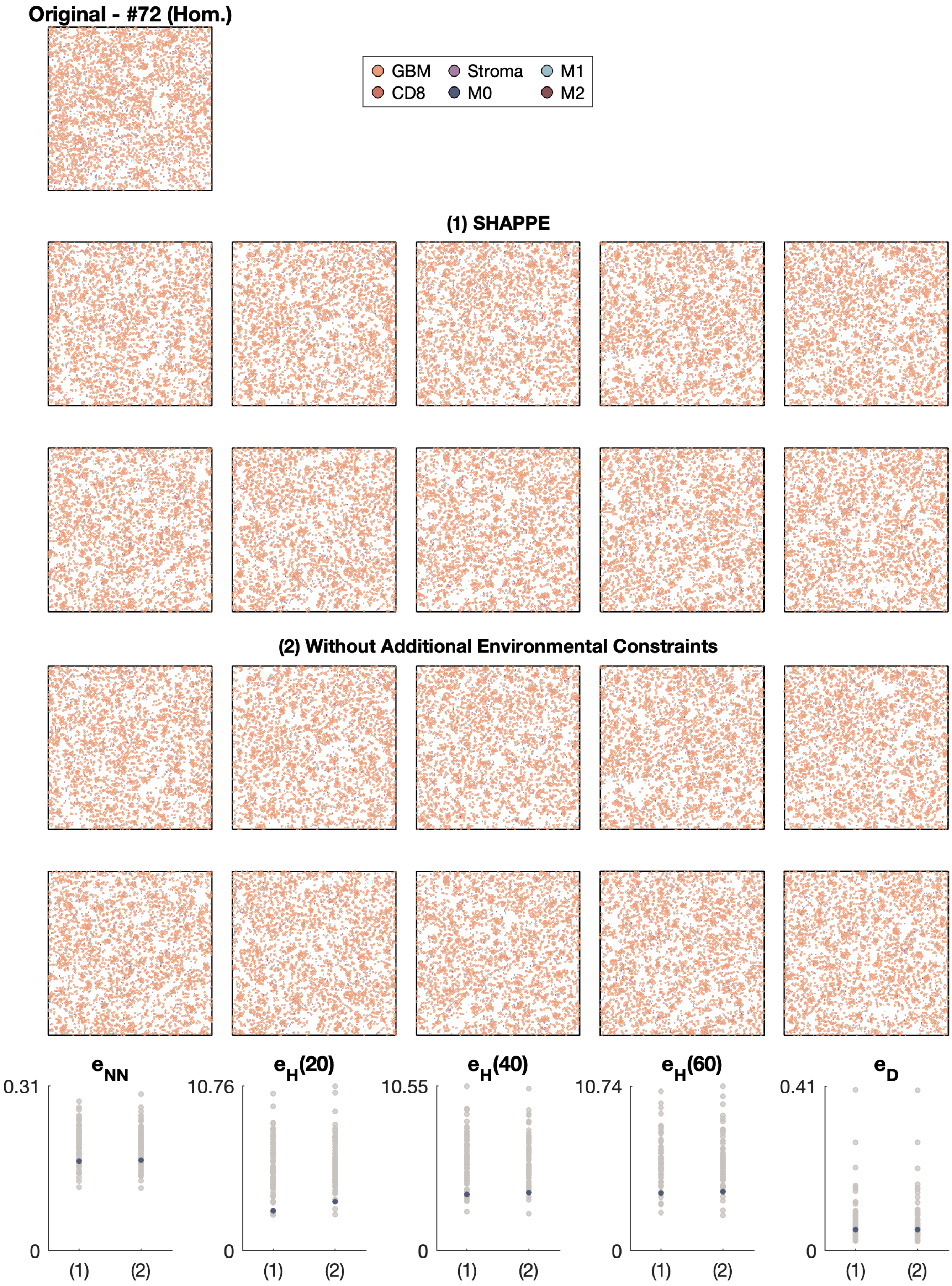
Original - #72 (Hom.)
GBM
Stroma
M1
CD8
M0
M2
(1) SHAPPE
(2) Without Additional Environmental Constraints
$e_{NN}$
$e_H(20)$
$e_H(40)$
$e_H(60)$
$e_D$
0.31
10.76
10.55
10.74
0.41
0
(1)
(2)

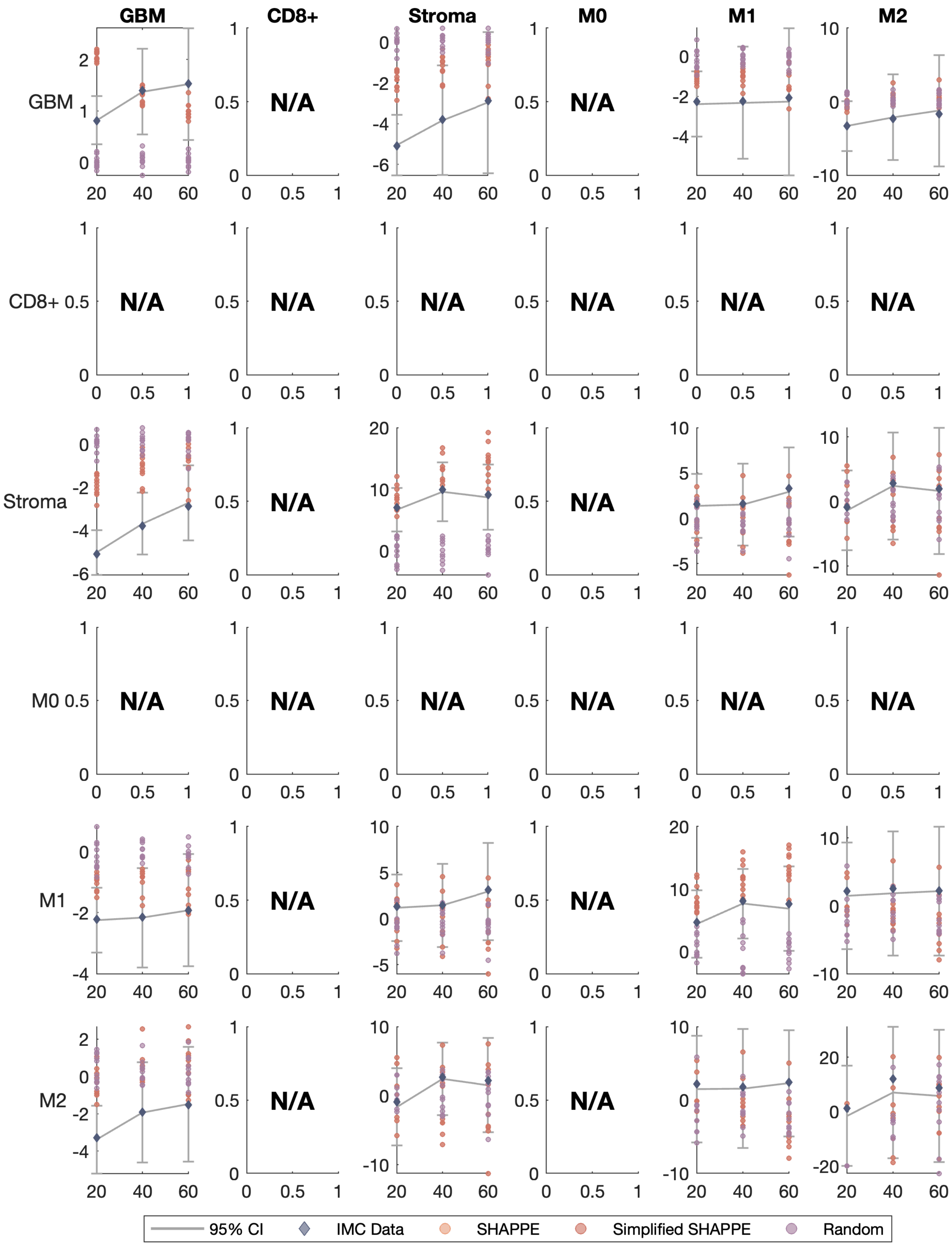
GBM
CD8+
Stroma
M0
M1
M2
N/A
95% CI
IMC Data
SHAPPE
Simplified SHAPPE
Random

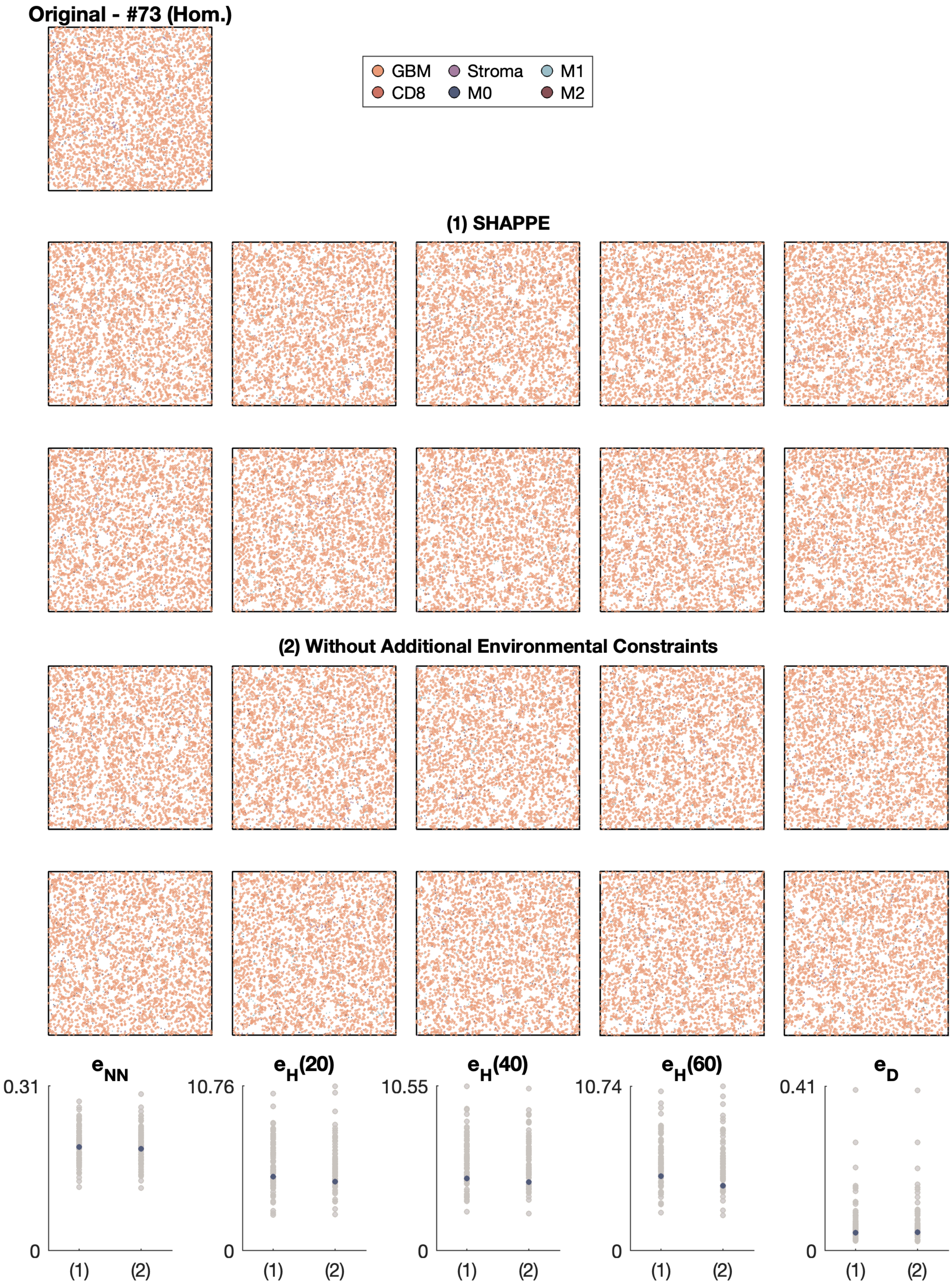
Original - #73 (Hom.)
GBM
Stroma
M1
CD8
M0
M2
(1) SHAPPE
(2) Without Additional Environmental Constraints
$e_{NN}$
$e_H(20)$
$e_H(40)$
$e_H(60)$
$e_D$
0.31
10.76
10.55
10.74
0.41
0
(1)
(2)

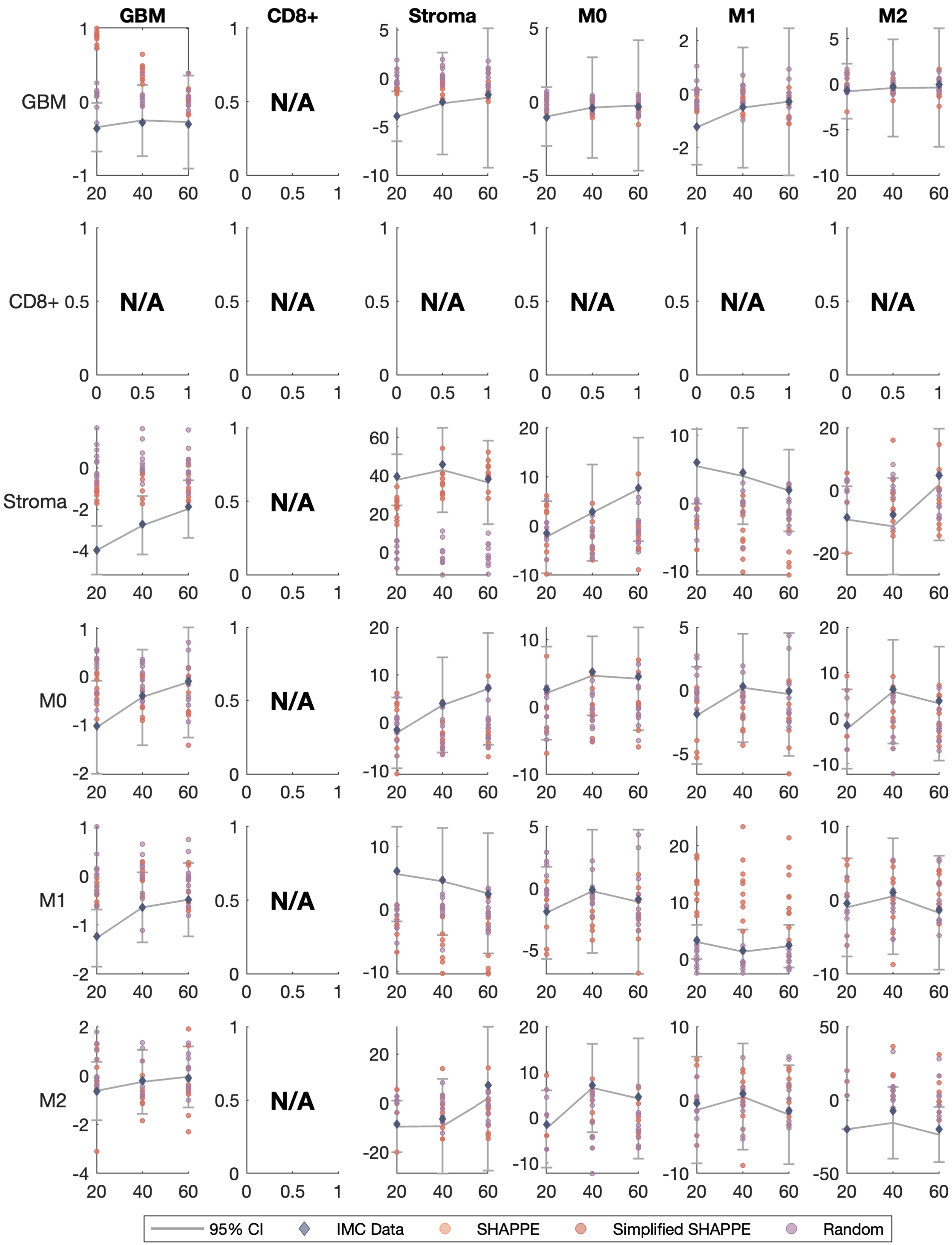

GBM
CD8+
Stroma
M0
M1
M2
N/A
95% CI
IMC Data
SHAPPE
Simplified SHAPPE
Random

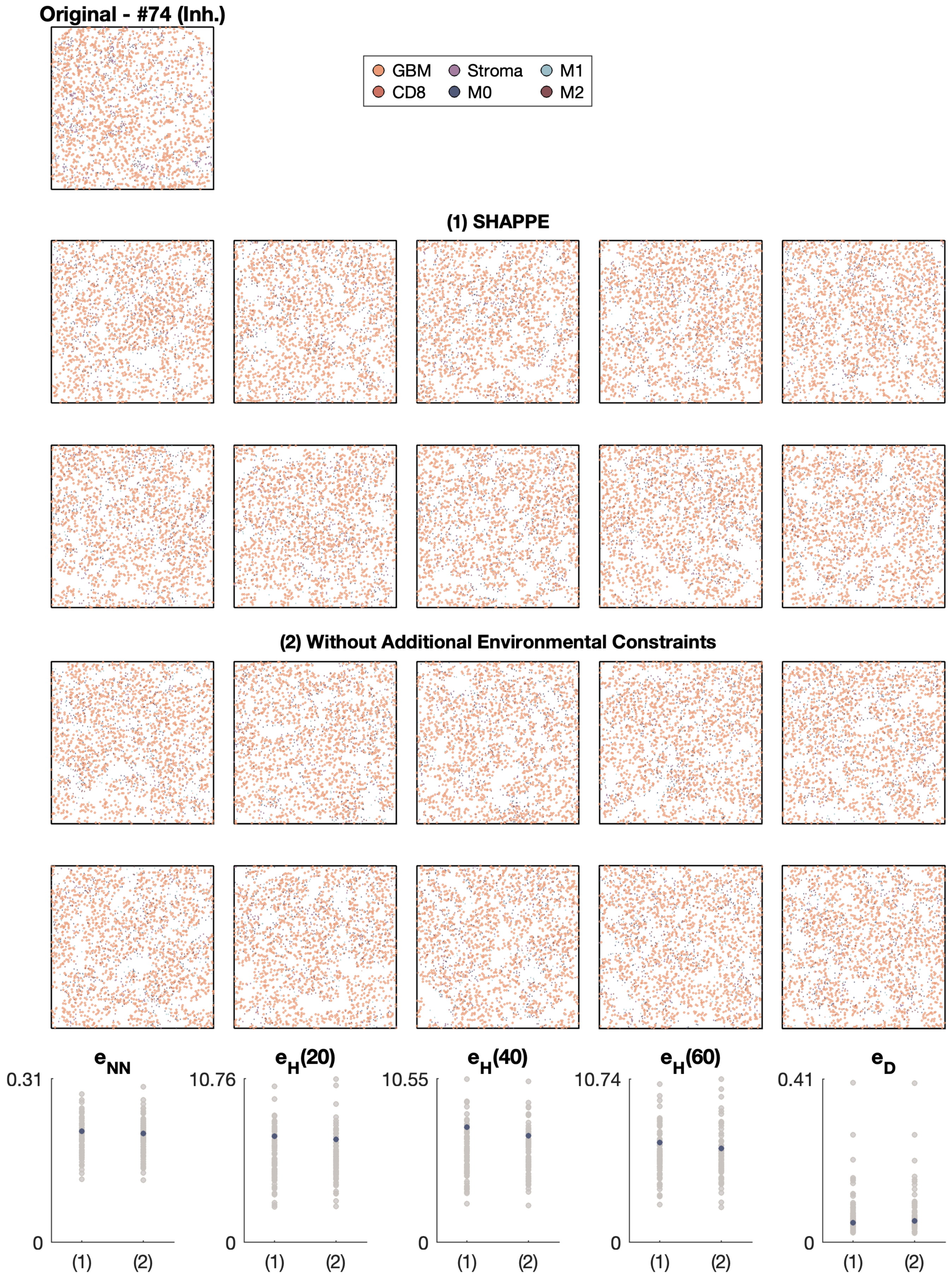

Original - #74 (Inh.)
GBM
Stroma
M1
CD8
M0
M2
(1) SHAPPE
(2) Without Additional Environmental Constraints
e_NN
e_H(20)
e_H(40)
e_H(60)
e_D
0.31
10.76
10.55
10.74
0.41
0
(1)
(2)

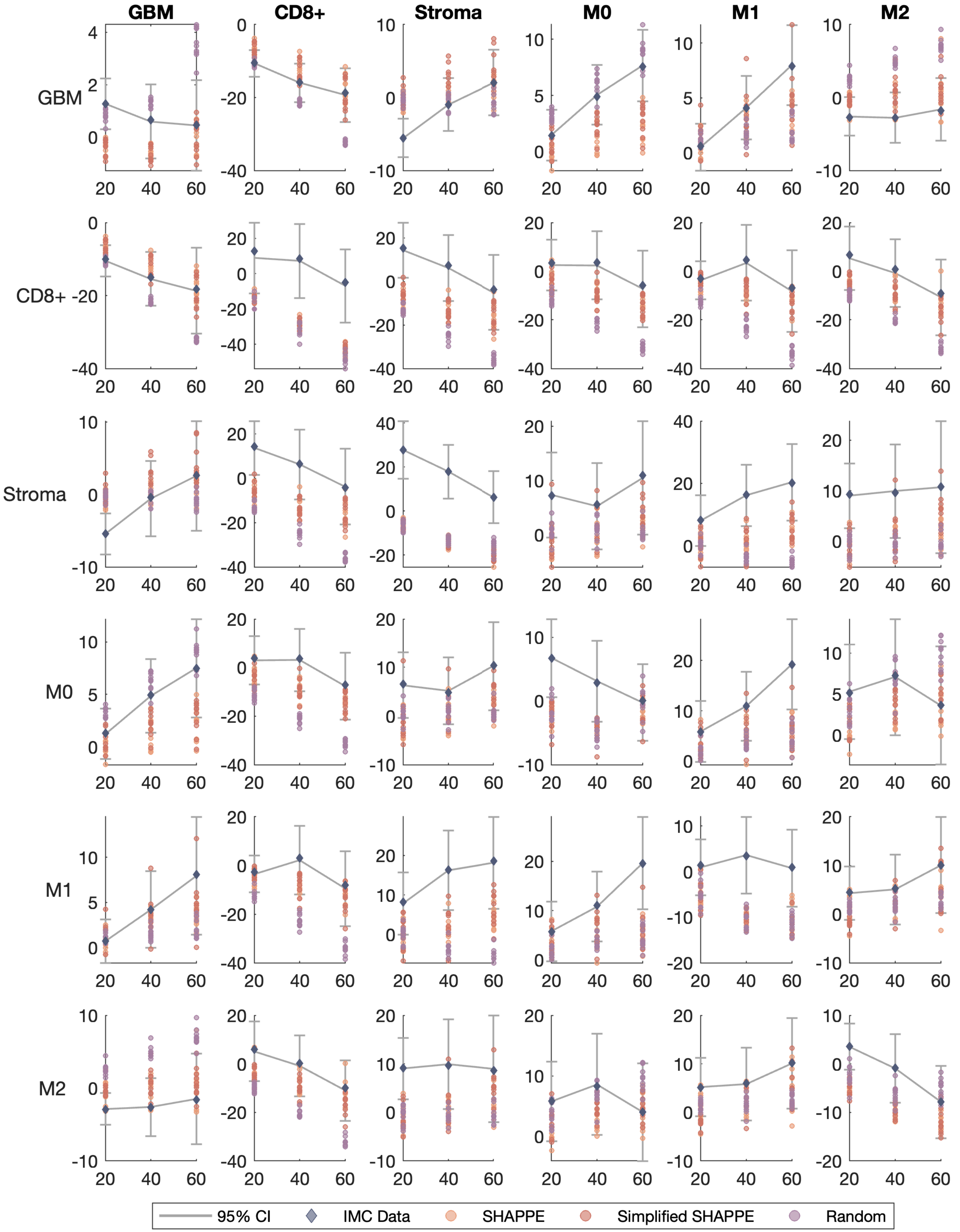
GBM
CD8+
Stroma
M0
M1
M2
95% CI
IMC Data
SHAPPE
Simplified SHAPPE
Random

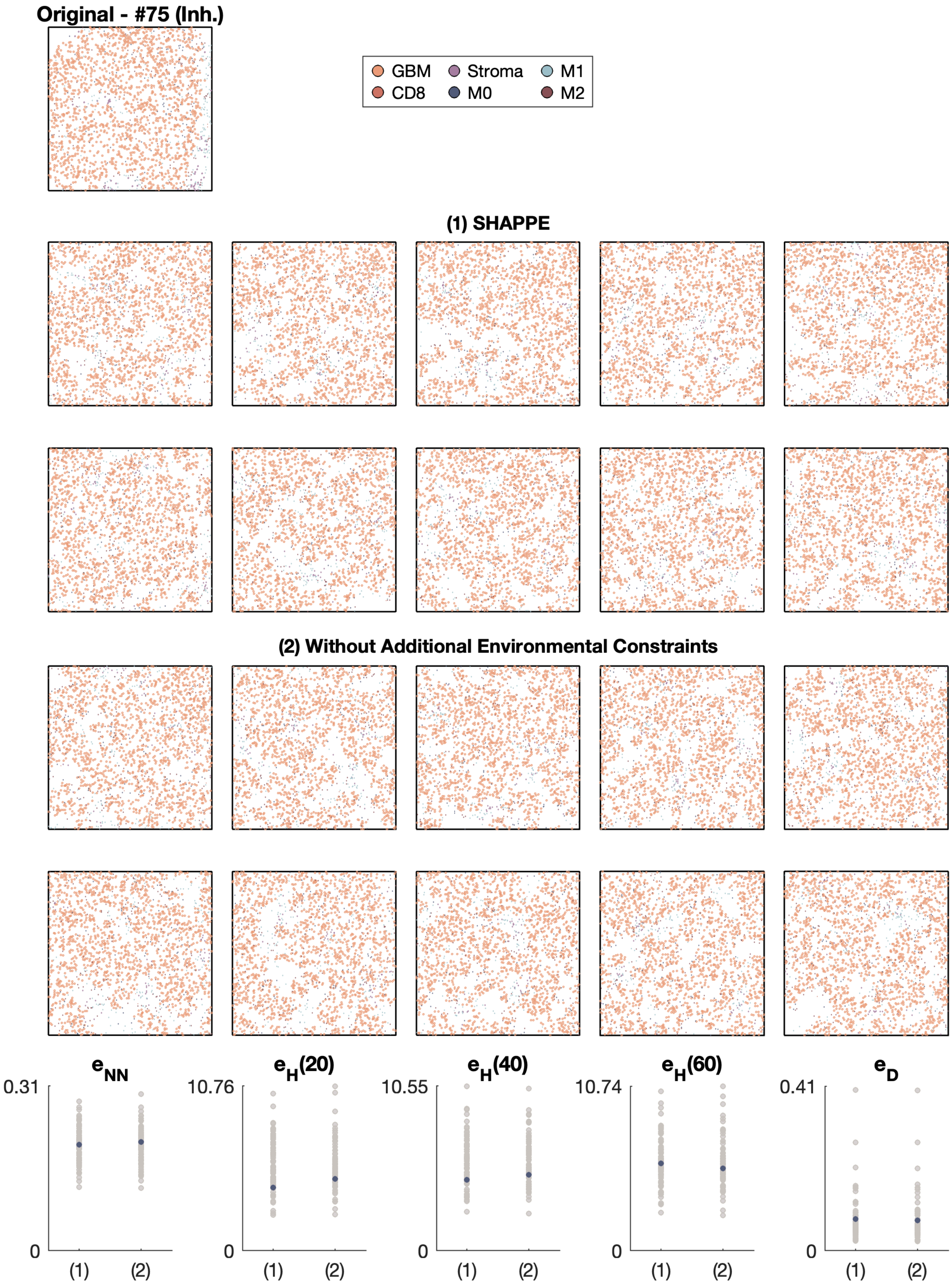
Original - #75 (Inh.)
GBM
CD8
Stroma
M0
M1
M2
(1) SHAPPE
(2) Without Additional Environmental Constraints
eNN
0.31
0
(1)
(2)
eH(20)
10.76
0
(1)
(2)
eH(40)
10.55
0
(1)
(2)
eH(60)
10.74
0
(1)
(2)
eD
0.41
0
(1)
(2)

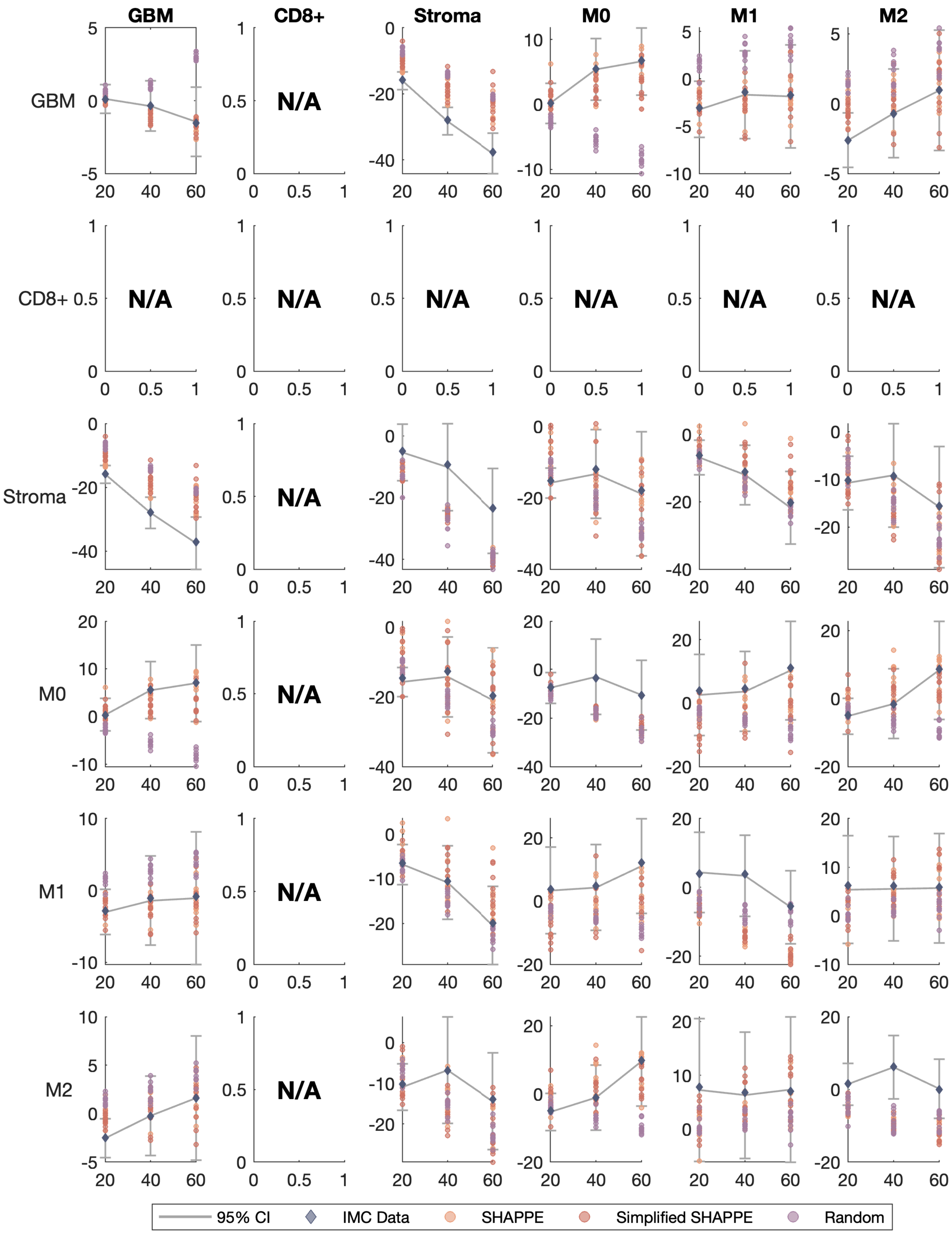

GBM
CD8+
Stroma
M0
M1
M2
N/A
95% CI
IMC Data
SHAPPE
Simplified SHAPPE
Random

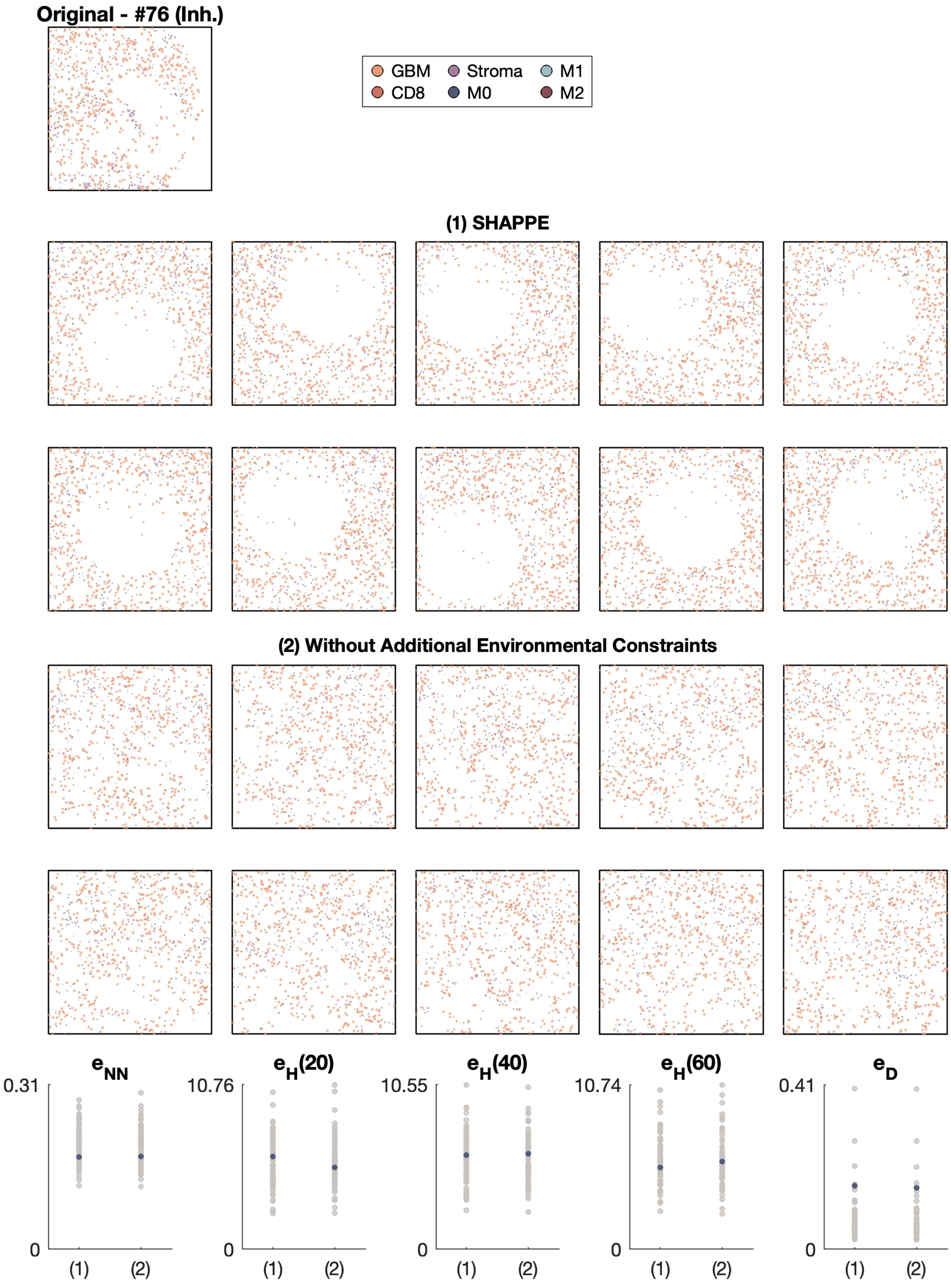
Original - #76 (Inh.)
GBM
CD8
Stroma
M0
M1
M2
(1) SHAPPE
(2) Without Additional Environmental Constraints
e_NN
0.31
0
(1)
(2)
e_H(20)
10.76
0
(1)
(2)
e_H(40)
10.55
0
(1)
(2)
e_H(60)
10.74
0
(1)
(2)
e_D
0.41
0
(1)
(2)

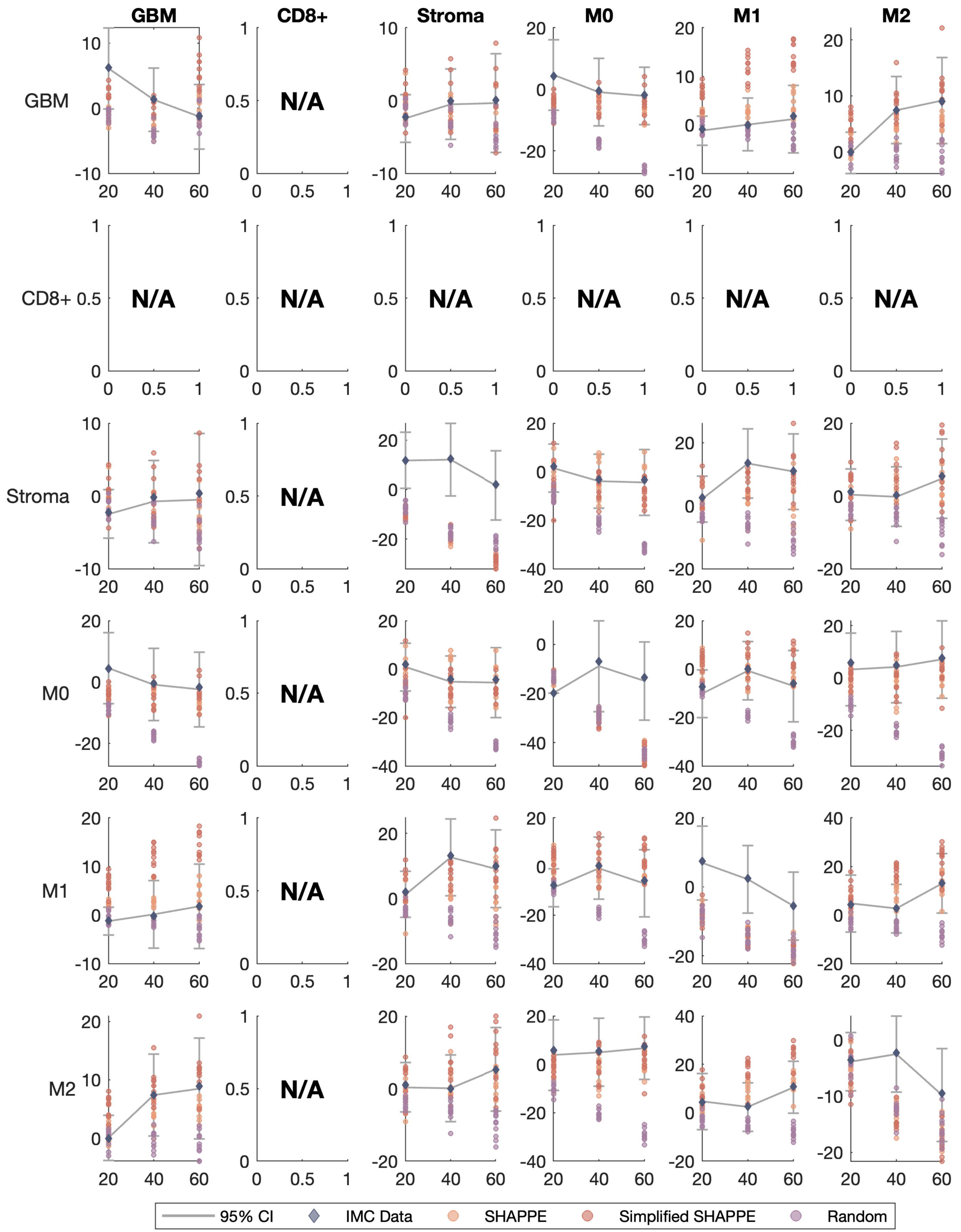
GBM
CD8+
Stroma
M0
M1
M2
GBM
CD8+
Stroma
M0
M1
M2
N/A
95% CI
IMC Data
SHAPPE
Simplified SHAPPE
Random

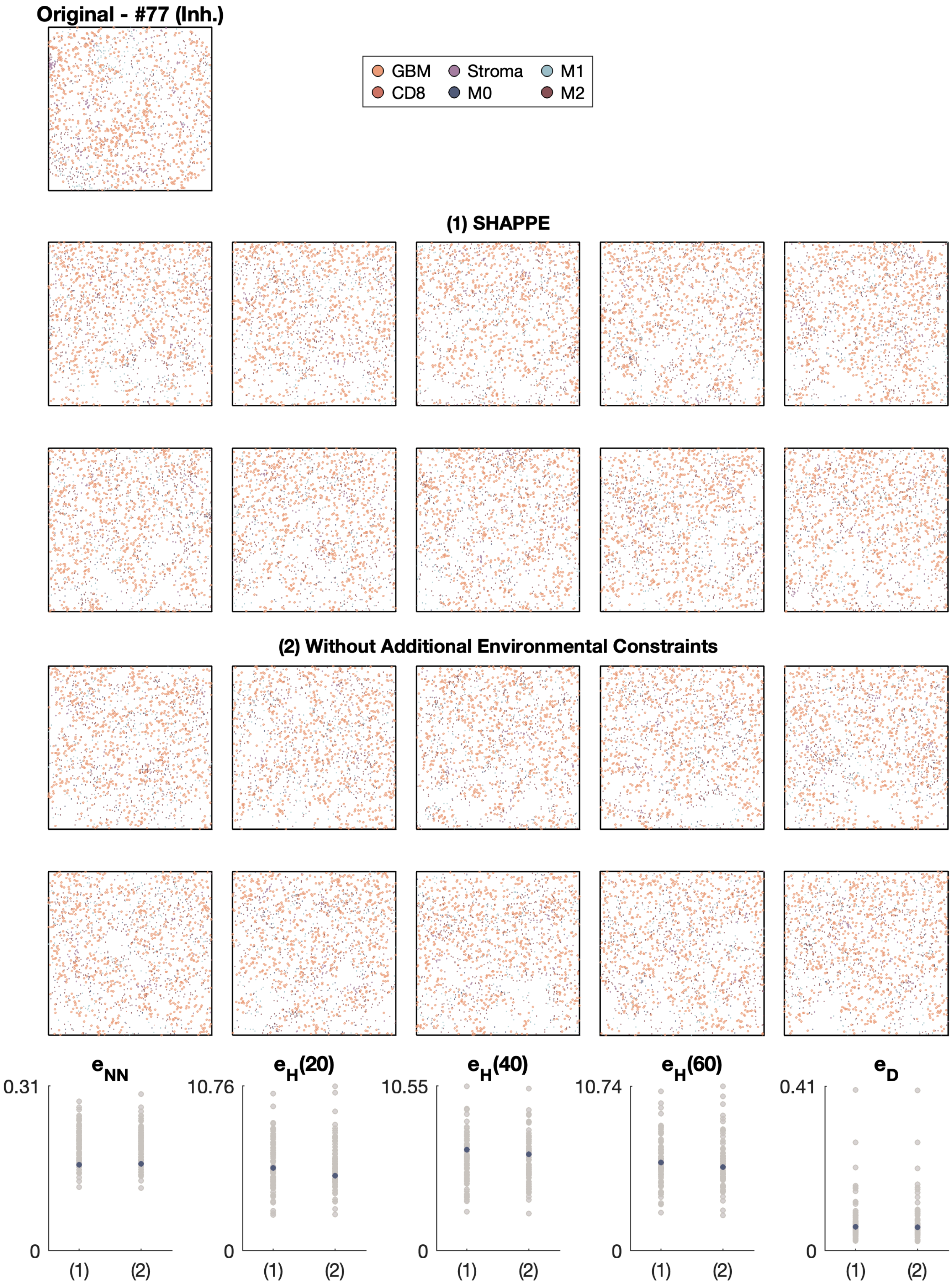
Original - #77 (Inh.)
GBM
Stroma
M1
CD8
M0
M2
(1) SHAPPE
(2) Without Additional Environmental Constraints
$e_{NN}$
0.31
0
(1)
(2)
$e_H(20)$
10.76
0
(1)
(2)
$e_H(40)$
10.55
0
(1)
(2)
$e_H(60)$
10.74
0
(1)
(2)
$e_D$
0.41
0
(1)
(2)

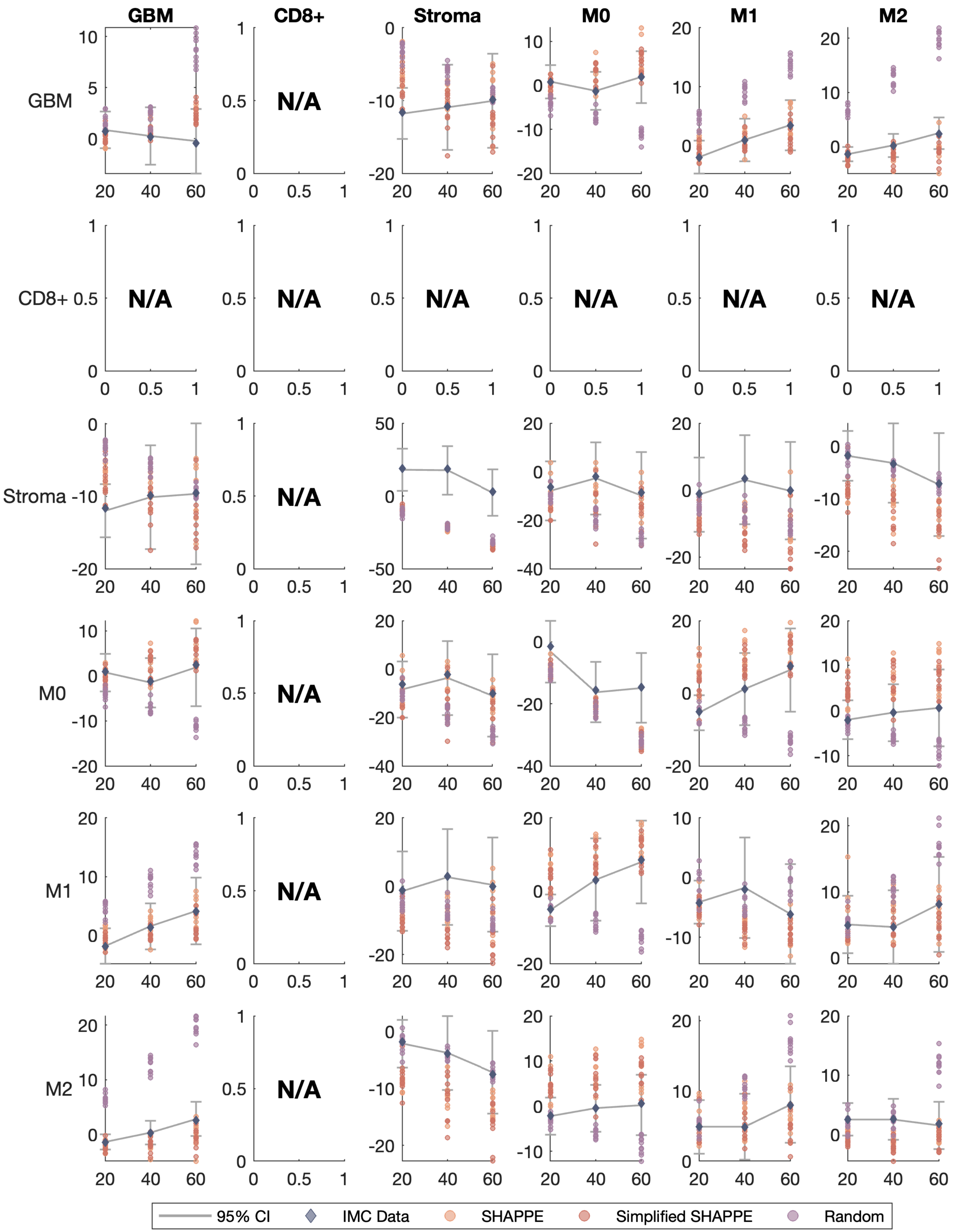

GBM
CD8+
Stroma
M0
M1
M2
N/A
95% CI
IMC Data
SHAPPE
Simplified SHAPPE
Random

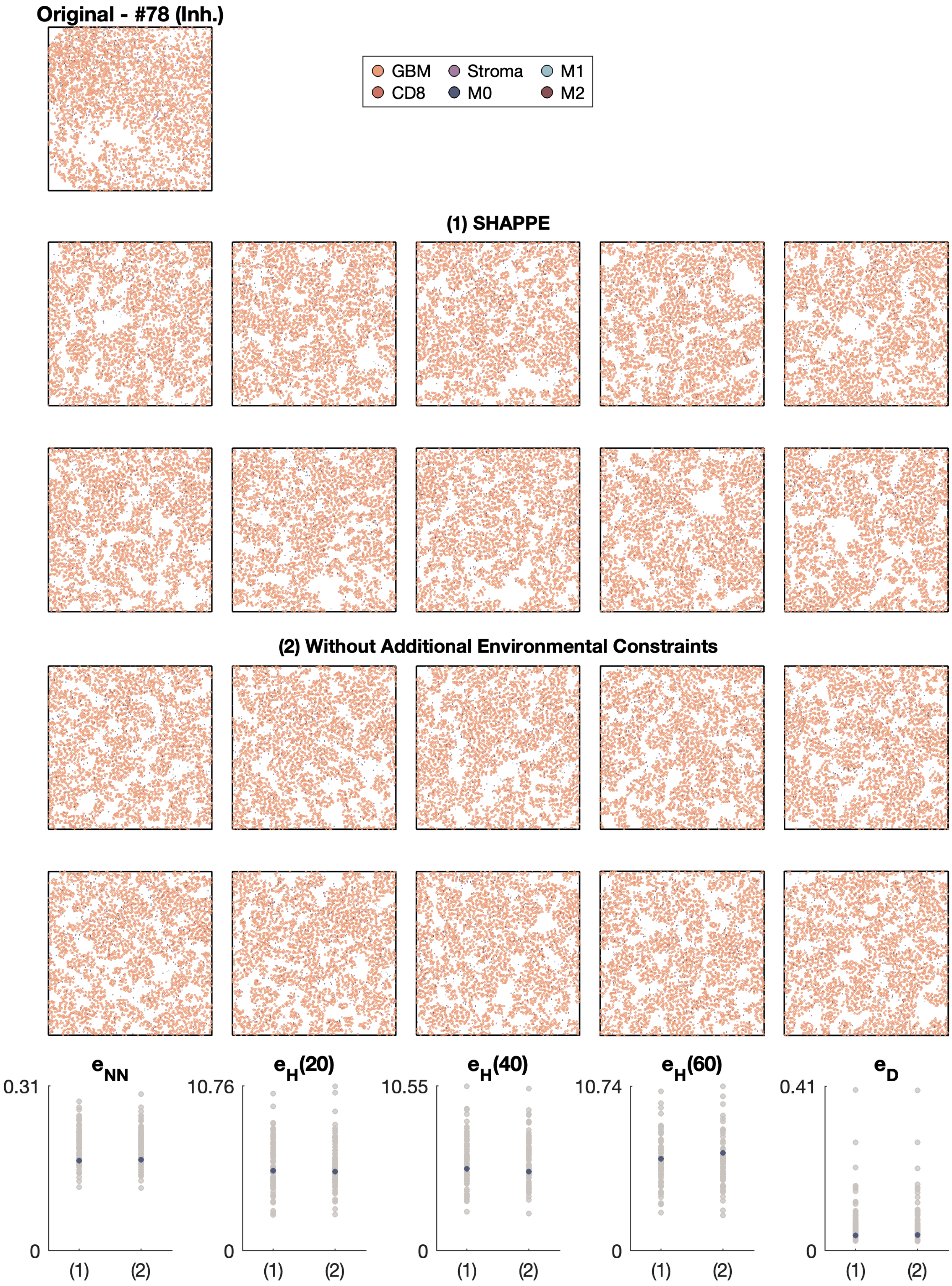

Original - #78 (Inh.)
GBM
CD8
Stroma
M0
M1
M2
(1) SHAPPE
(2) Without Additional Environmental Constraints
$e_{NN}$
0.31
0
$e_H(20)$
10.76
$e_H(40)$
10.55
$e_H(60)$
10.74
$e_D$
0.41
(1)
(2)

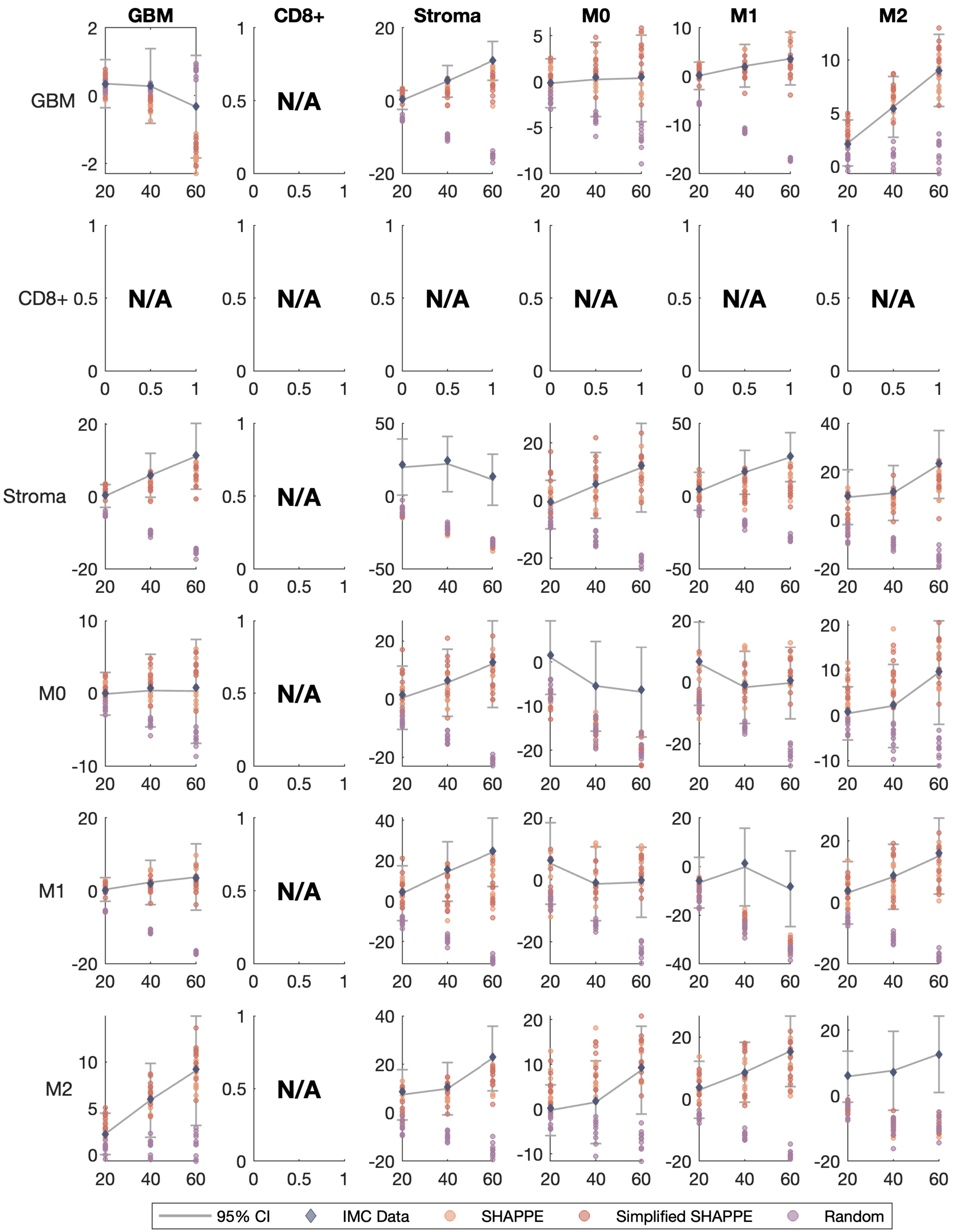

GBM
CD8+
Stroma
M0
M1
M2
N/A
95% CI
IMC Data
SHAPPE
Simplified SHAPPE
Random

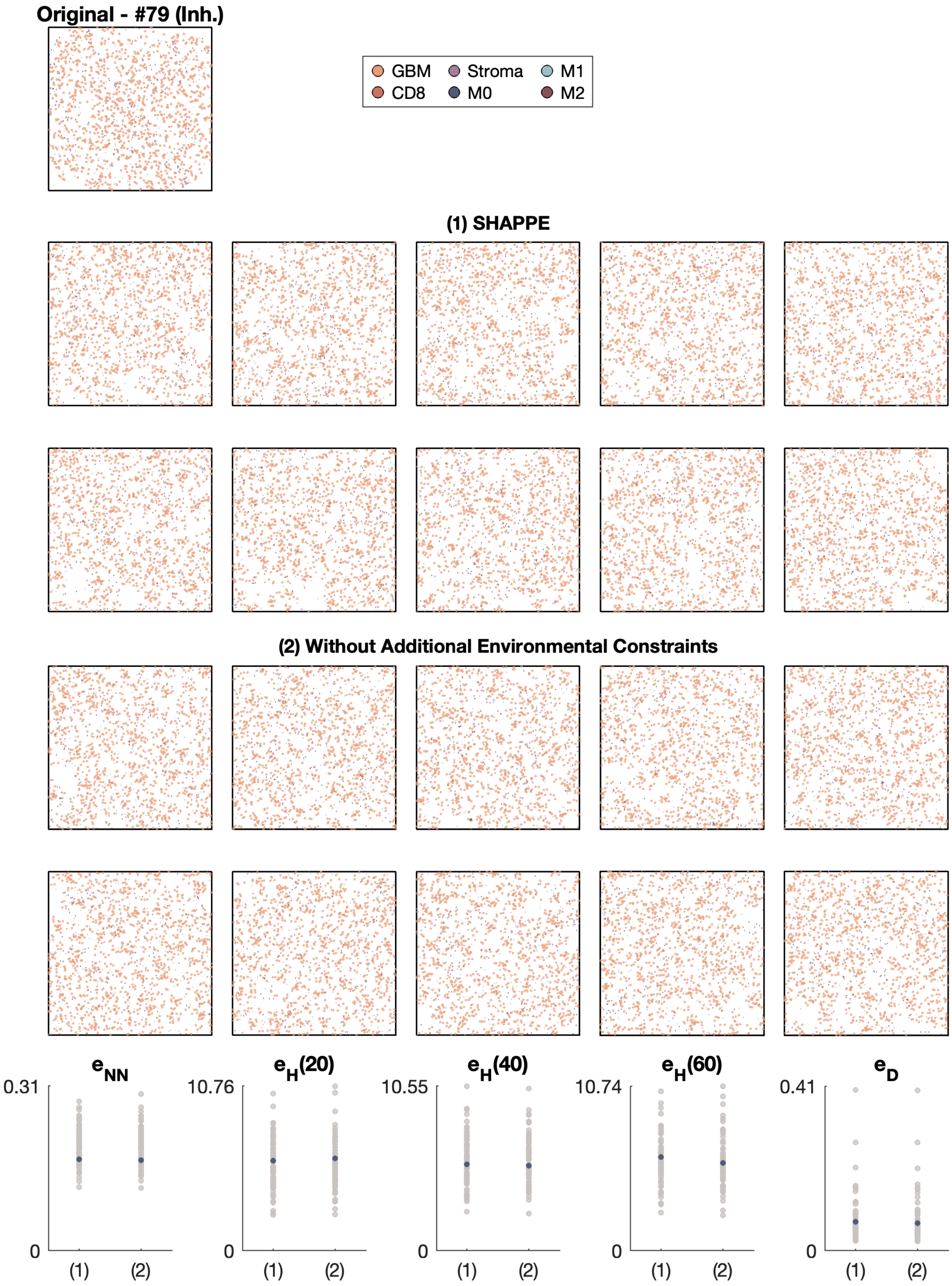
Original - #79 (Inh.)
GBM
CD8
Stroma
M0
M1
M2
(1) SHAPPE
(2) Without Additional Environmental Constraints
$e_{NN}$
$e_H(20)$
$e_H(40)$
$e_H(60)$
$e_D$
0.31
10.76
10.55
10.74
0.41
0
(1)
(2)

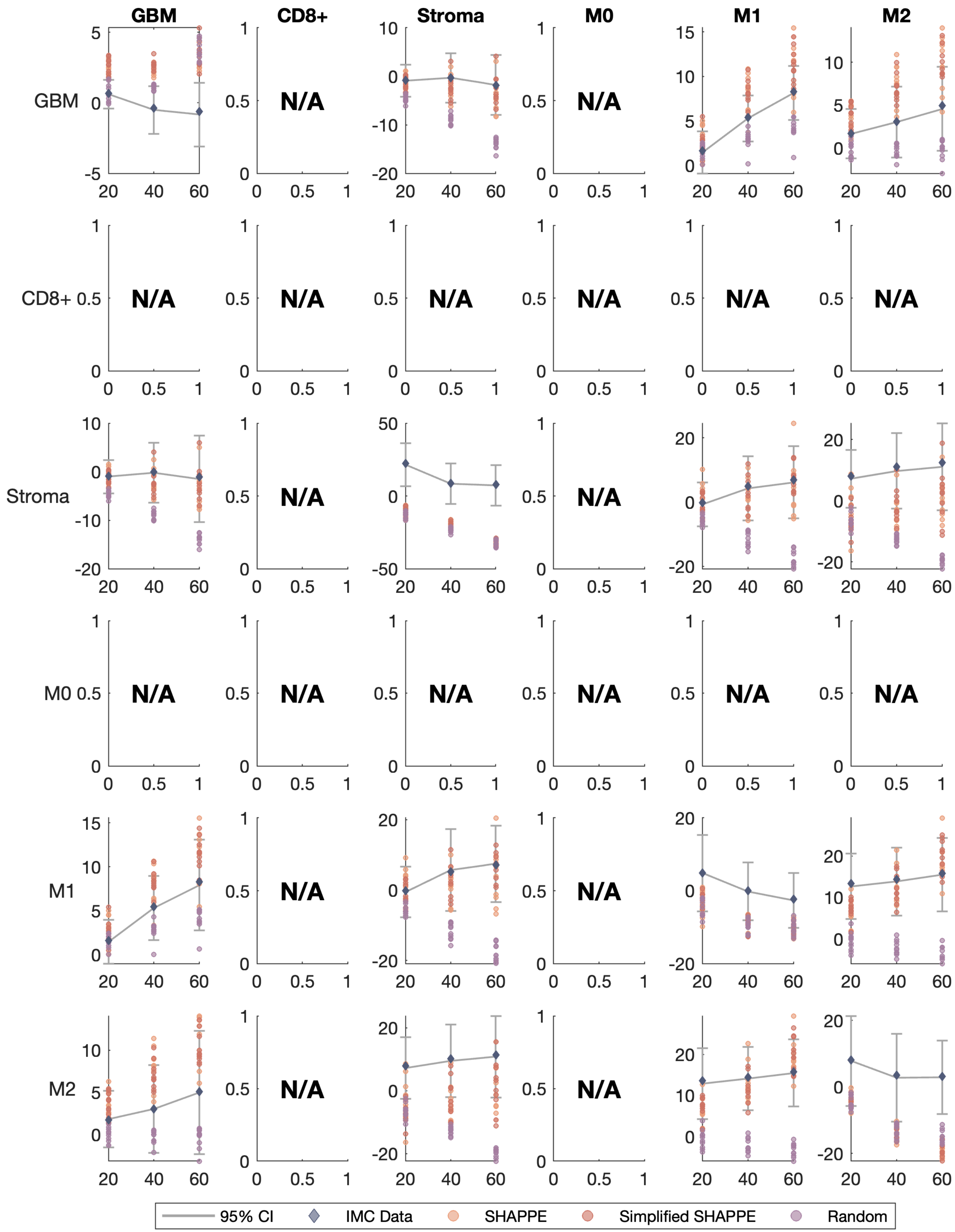

GBM
CD8+
Stroma
M0
M1
M2
N/A
95% CI
IMC Data
SHAPPE
Simplified SHAPPE
Random

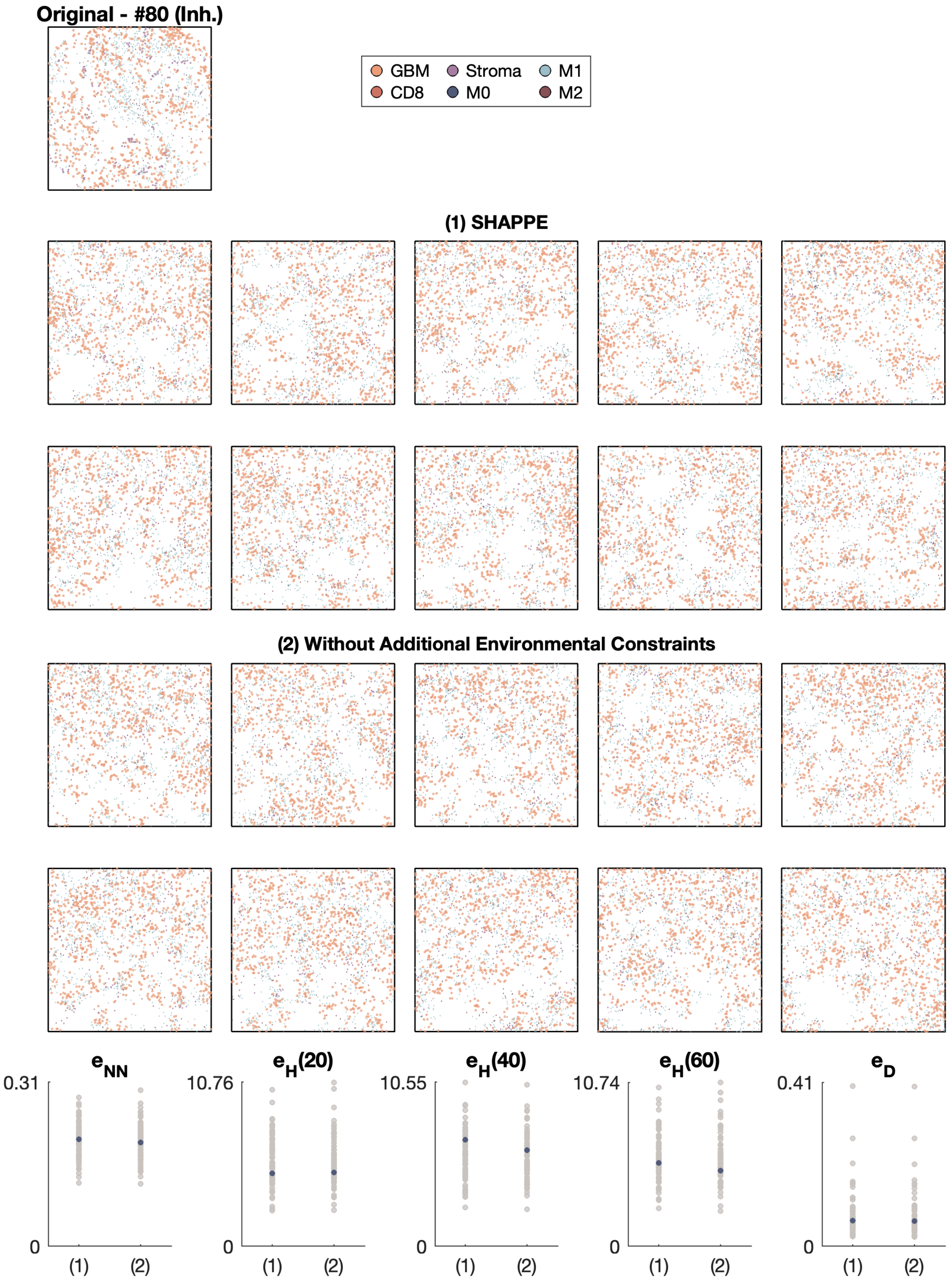

Original - #80 (Inh.)
GBM
Stroma
M1
CD8
M0
M2
(1) SHAPPE
(2) Without Additional Environmental Constraints
$e_{NN}$
$e_H(20)$
$e_H(40)$
$e_H(60)$
$e_D$
0.31
10.76
10.55
10.74
0.41
0
(1)
(2)

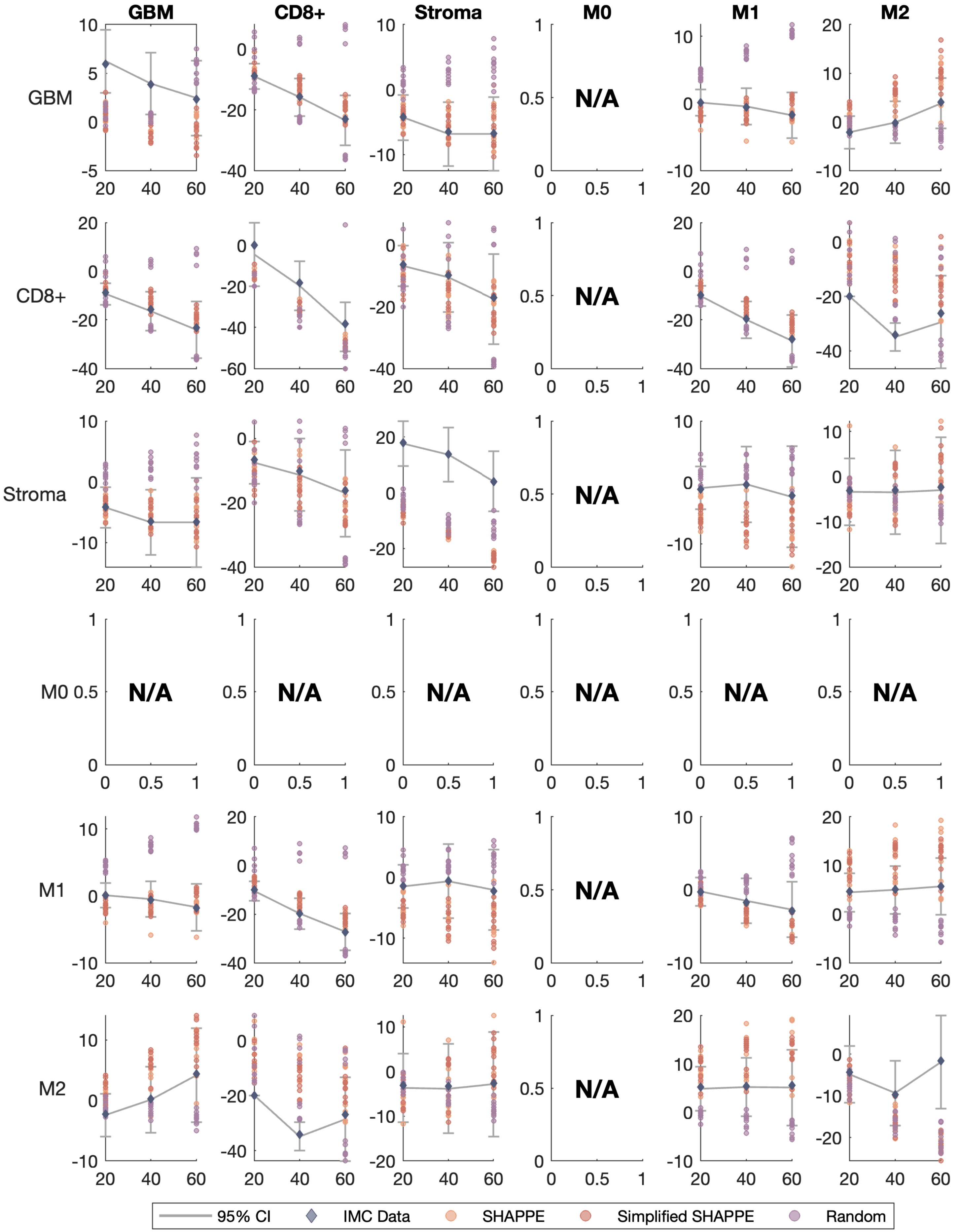
GBM
CD8+
Stroma
M0
M1
M2
N/A
95% CI
IMC Data
SHAPPE
Simplified SHAPPE
Random

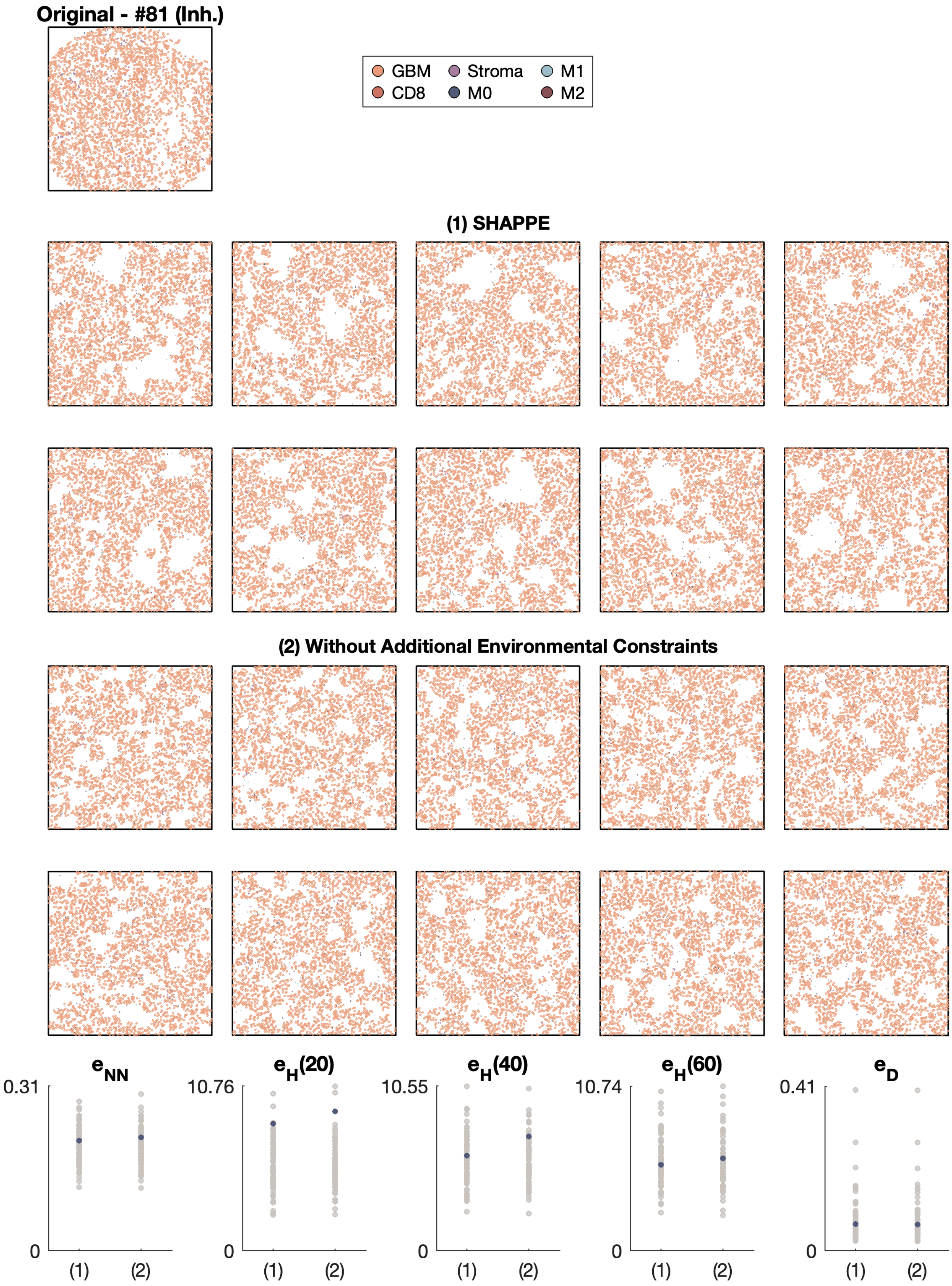
Original - #81 (Inh.)
GBM
Stroma
M1
CD8
M0
M2
(1) SHAPPE
(2) Without Additional Environmental Constraints
$e_{NN}$
$e_H(20)$
$e_H(40)$
$e_H(60)$
$e_D$
0.31
10.76
10.55
10.74
0.41
0
(1)
(2)

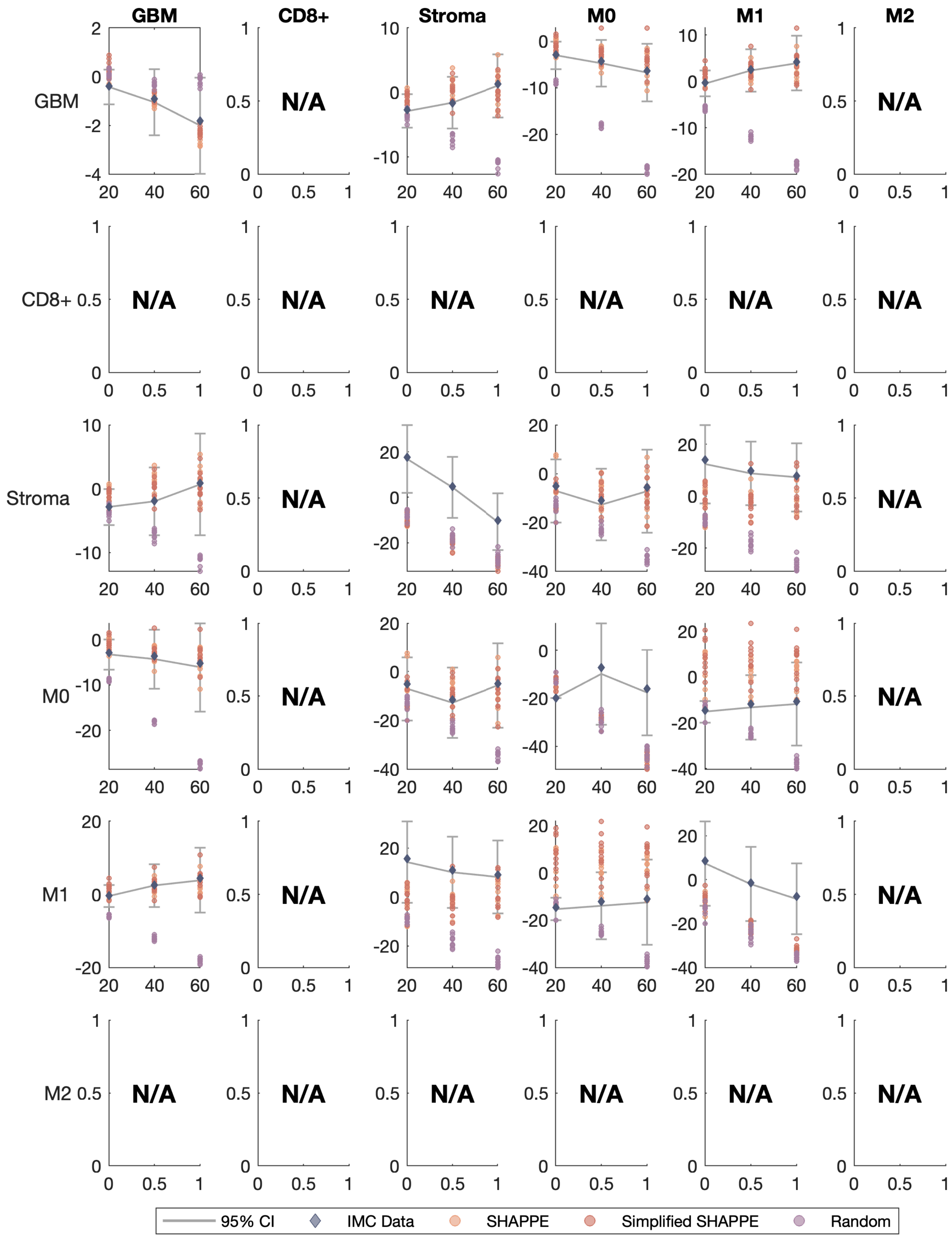

GBM
CD8+
Stroma
M0
M1
M2
N/A
95% CI
IMC Data
SHAPPE
Simplified SHAPPE
Random

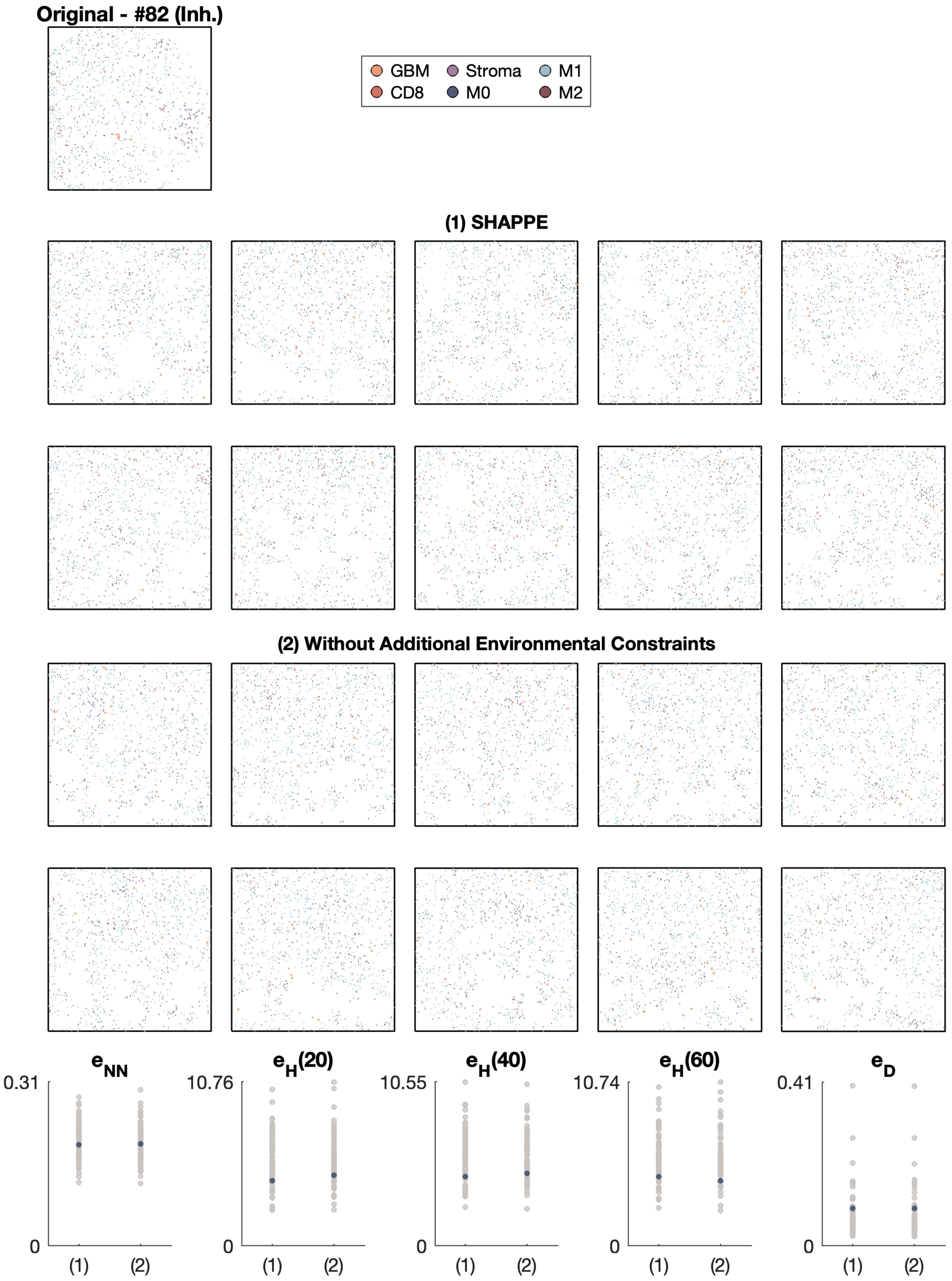
Original - #82 (Inh.)
GBM
CD8
Stroma
M0
M1
M2
(1) SHAPPE
(2) Without Additional Environmental Constraints
$e_{NN}$
$e_H(20)$
$e_H(40)$
$e_H(60)$
$e_D$
0.31
10.76
10.55
10.74
0.41
0
(1)
(2)

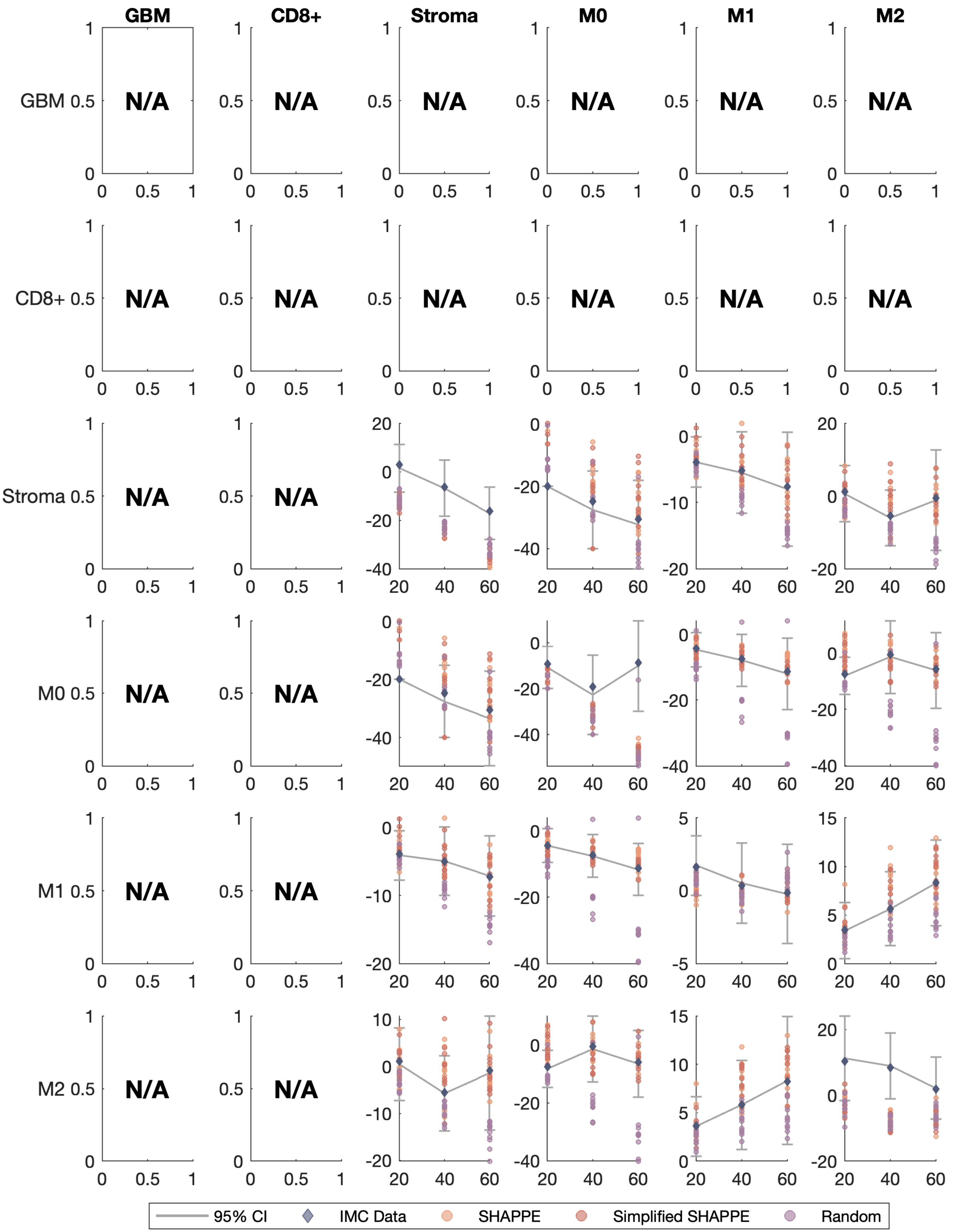

GBM
CD8+
Stroma
M0
M1
M2
GBM
CD8+
Stroma
M0
M1
M2
N/A
95% CI
IMC Data
SHAPPE
Simplified SHAPPE
Random

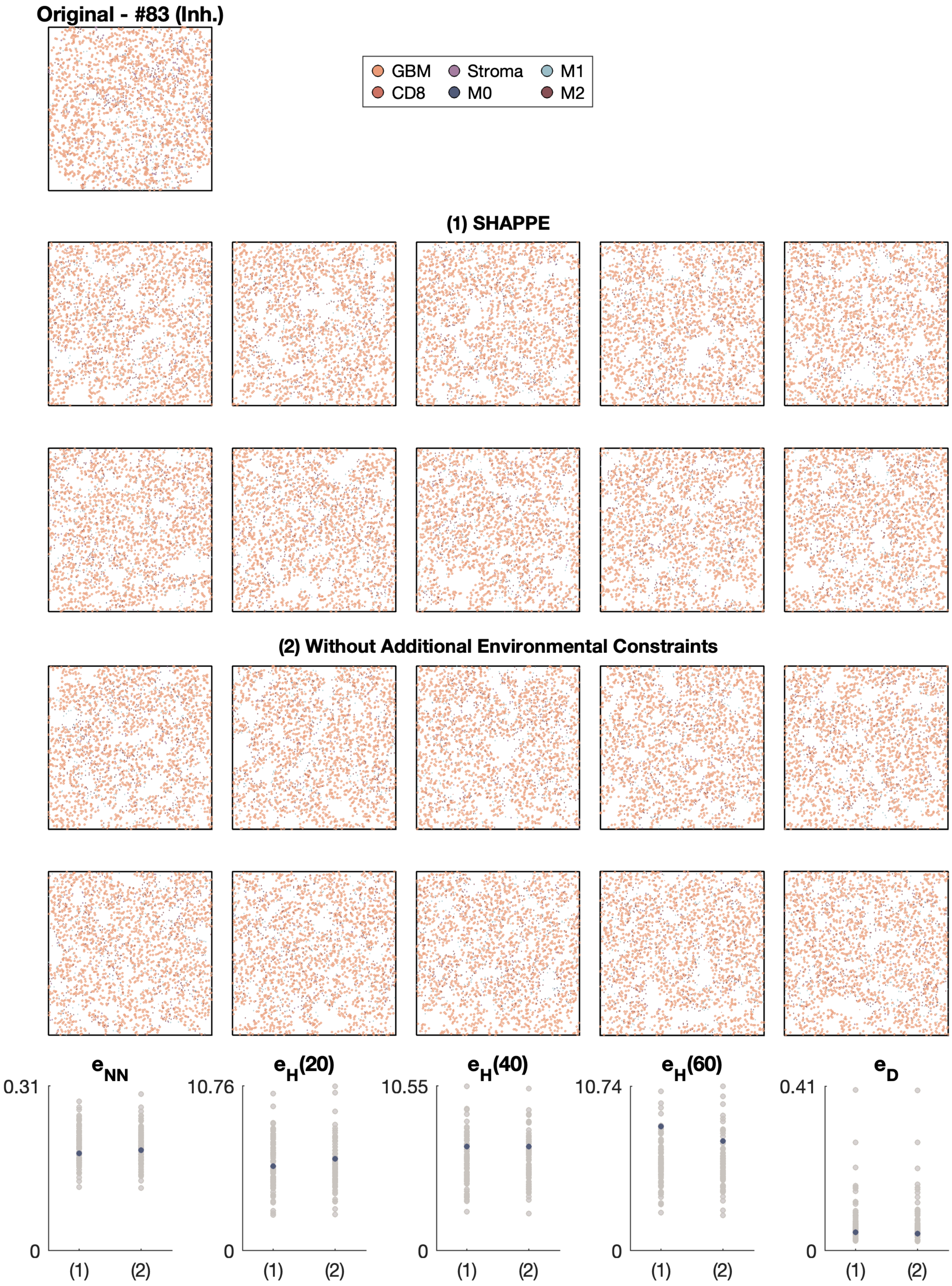
Original - #83 (Inh.)
GBM
Stroma
M1
CD8
M0
M2
(1) SHAPPE
(2) Without Additional Environmental Constraints
$e_{NN}$
0.31
0
(1)
(2)
$e_H(20)$
10.76
0
(1)
(2)
$e_H(40)$
10.55
0
(1)
(2)
$e_H(60)$
10.74
0
(1)
(2)
$e_D$
0.41
0
(1)
(2)

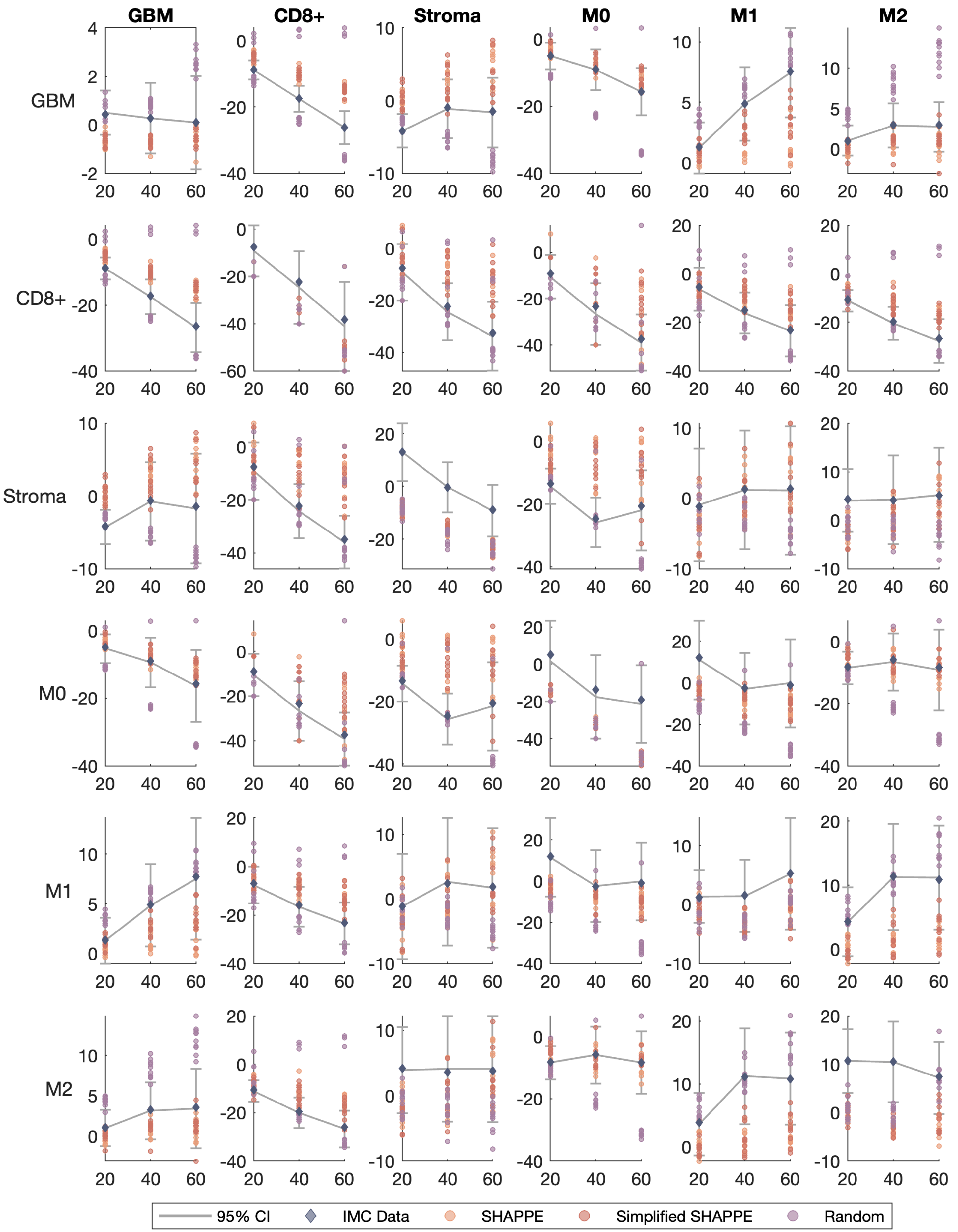

GBM
CD8+
Stroma
M0
M1
M2
95% CI
IMC Data
SHAPPE
Simplified SHAPPE
Random

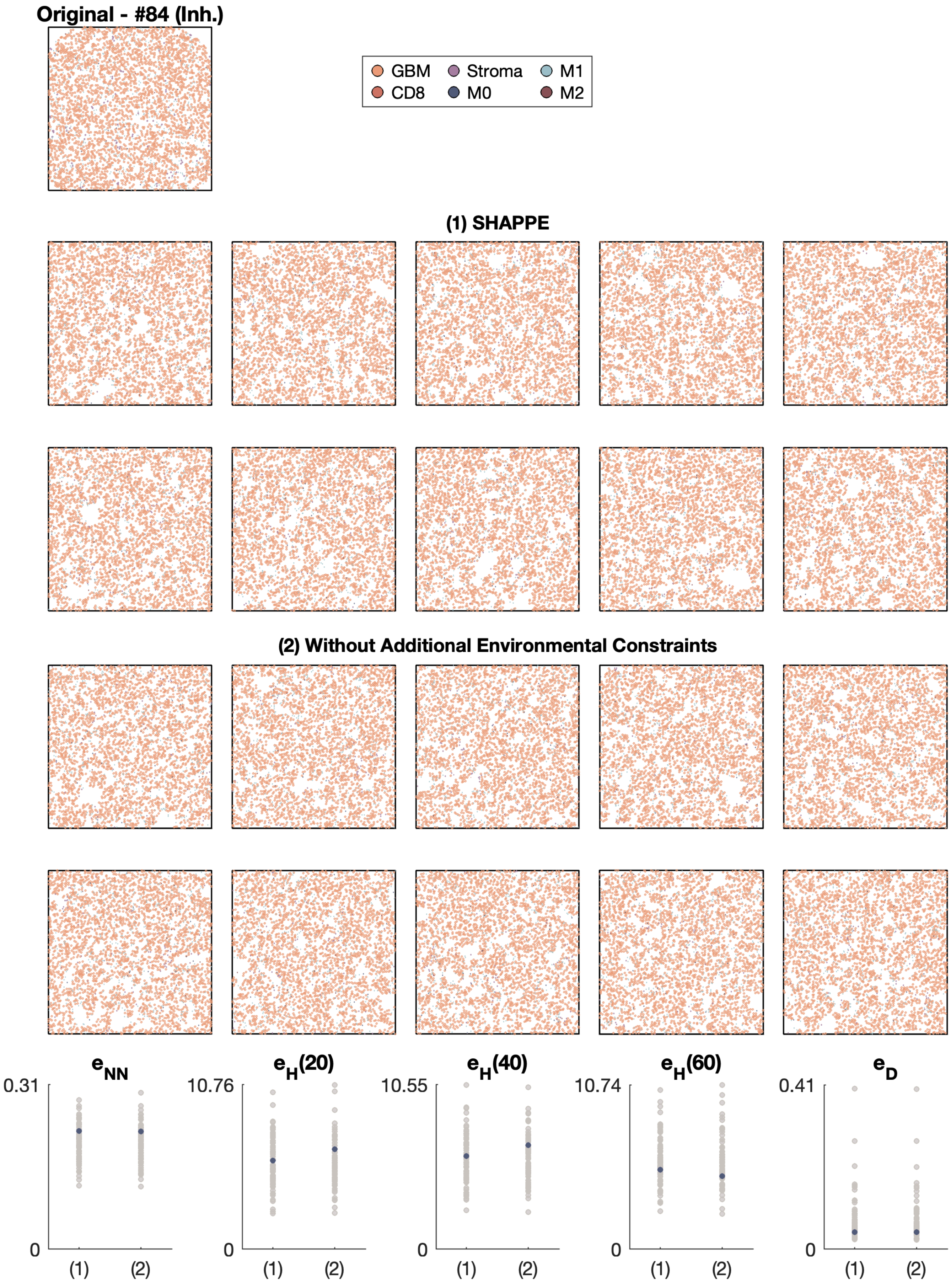
Original - #84 (Inh.)
GBM
CD8
Stroma
M0
M1
M2
(1) SHAPPE
(2) Without Additional Environmental Constraints
e_NN
0.31
0
(1)
(2)
e_H(20)
10.76
0
(1)
(2)
e_H(40)
10.55
0
(1)
(2)
e_H(60)
10.74
0
(1)
(2)
e_D
0.41
0
(1)
(2)

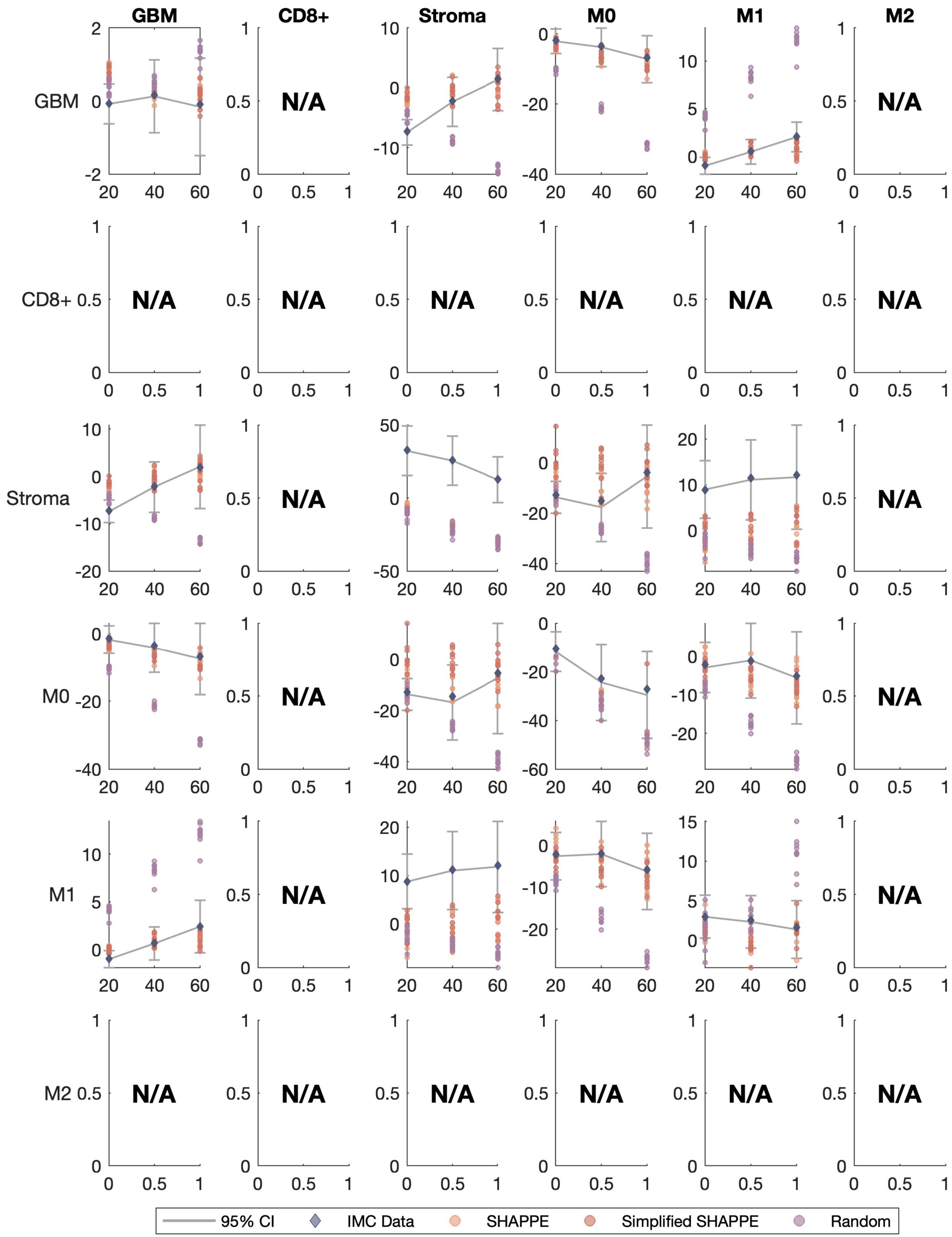
GBM
CD8+
Stroma
M0
M1
M2
N/A
95% CI
IMC Data
SHAPPE
Simplified SHAPPE
Random

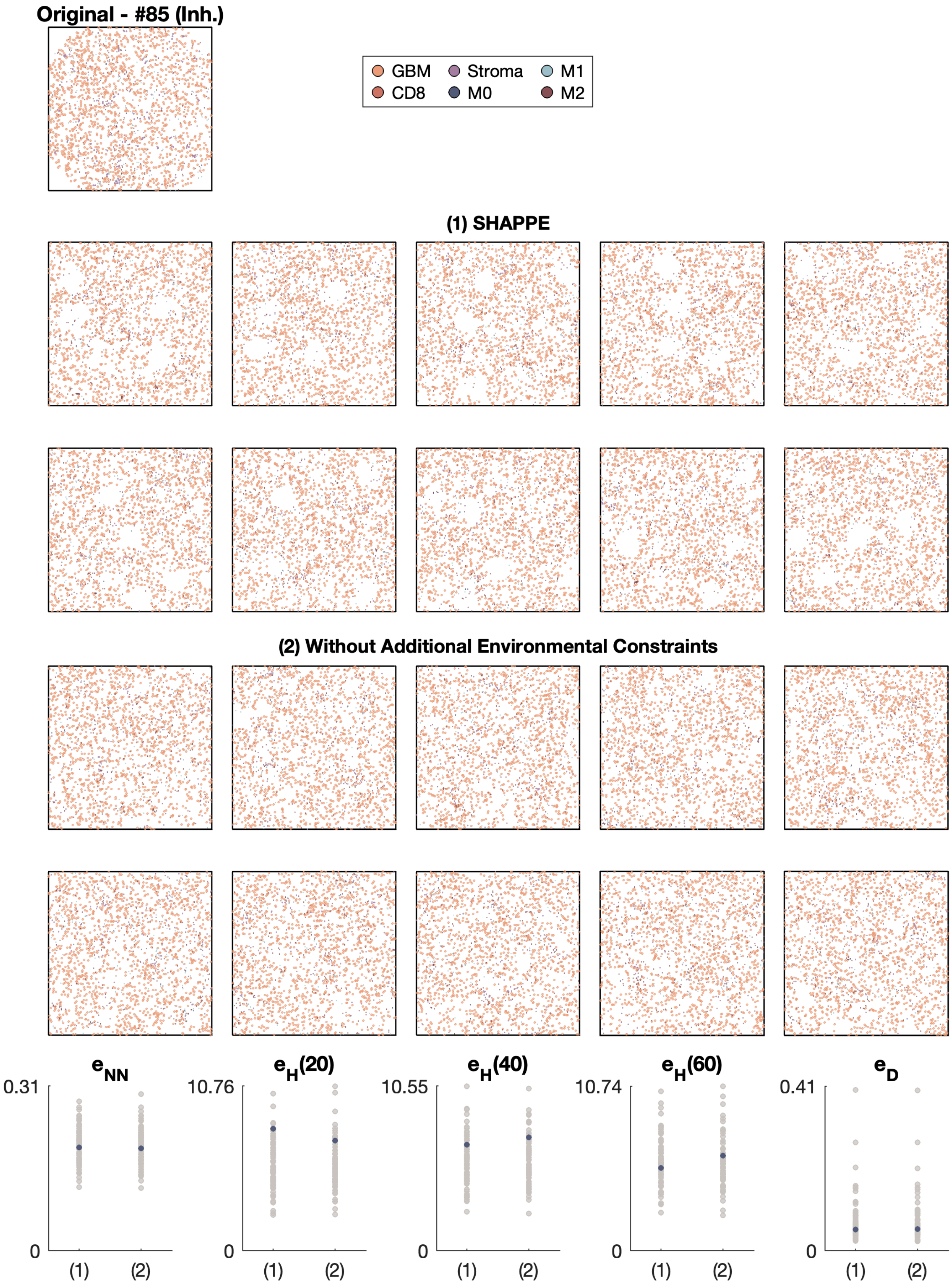

Original - #85 (Inh.)
GBM
Stroma
M1
CD8
M0
M2
(1) SHAPPE
(2) Without Additional Environmental Constraints
$e_{NN}$
0.31
0
$e_H(20)$
10.76
0
$e_H(40)$
10.55
0
$e_H(60)$
10.74
0
$e_D$
0.41
0
(1)
(2)

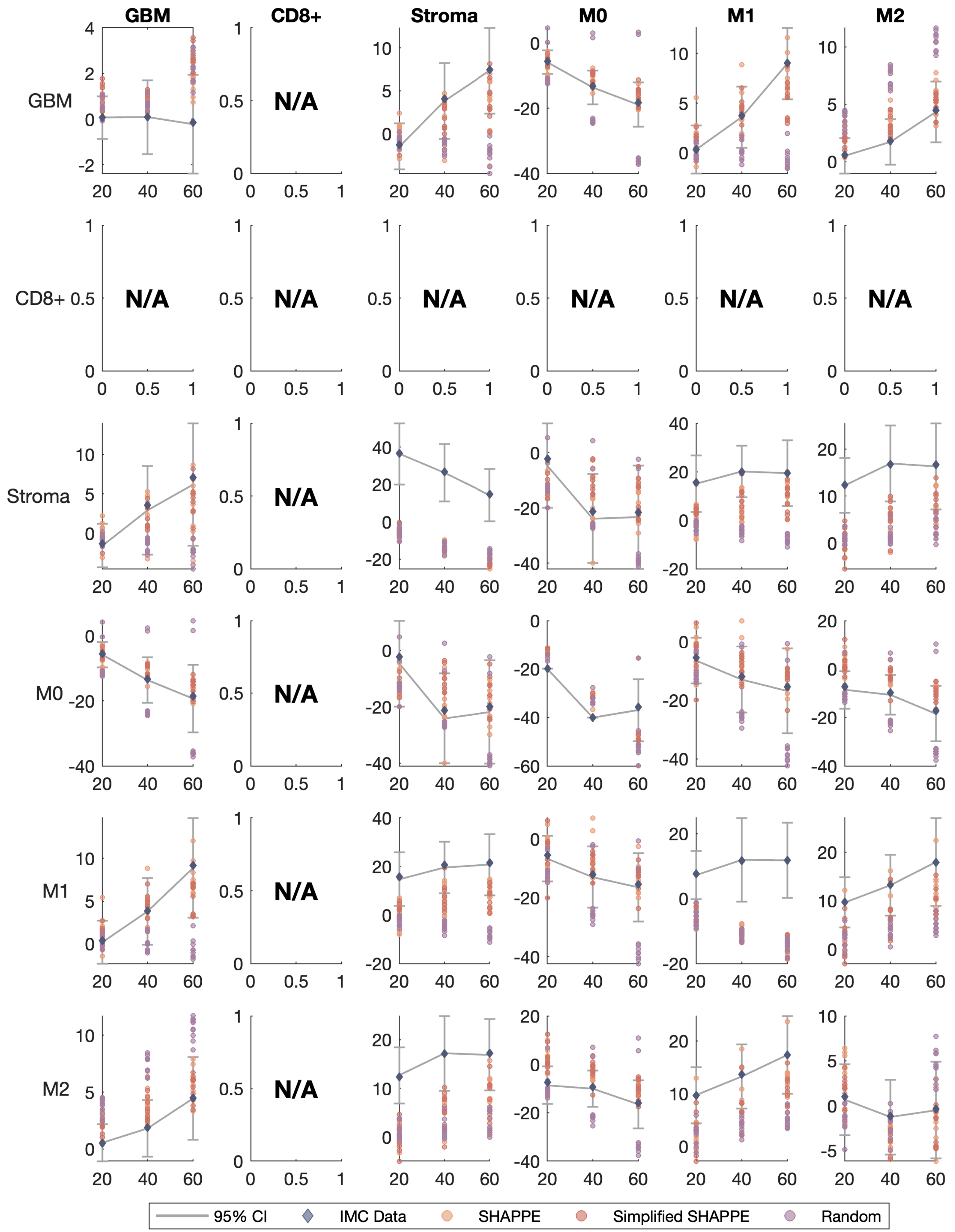

GBM
CD8+
Stroma
M0
M1
M2
N/A
95% CI
IMC Data
SHAPPE
Simplified SHAPPE
Random

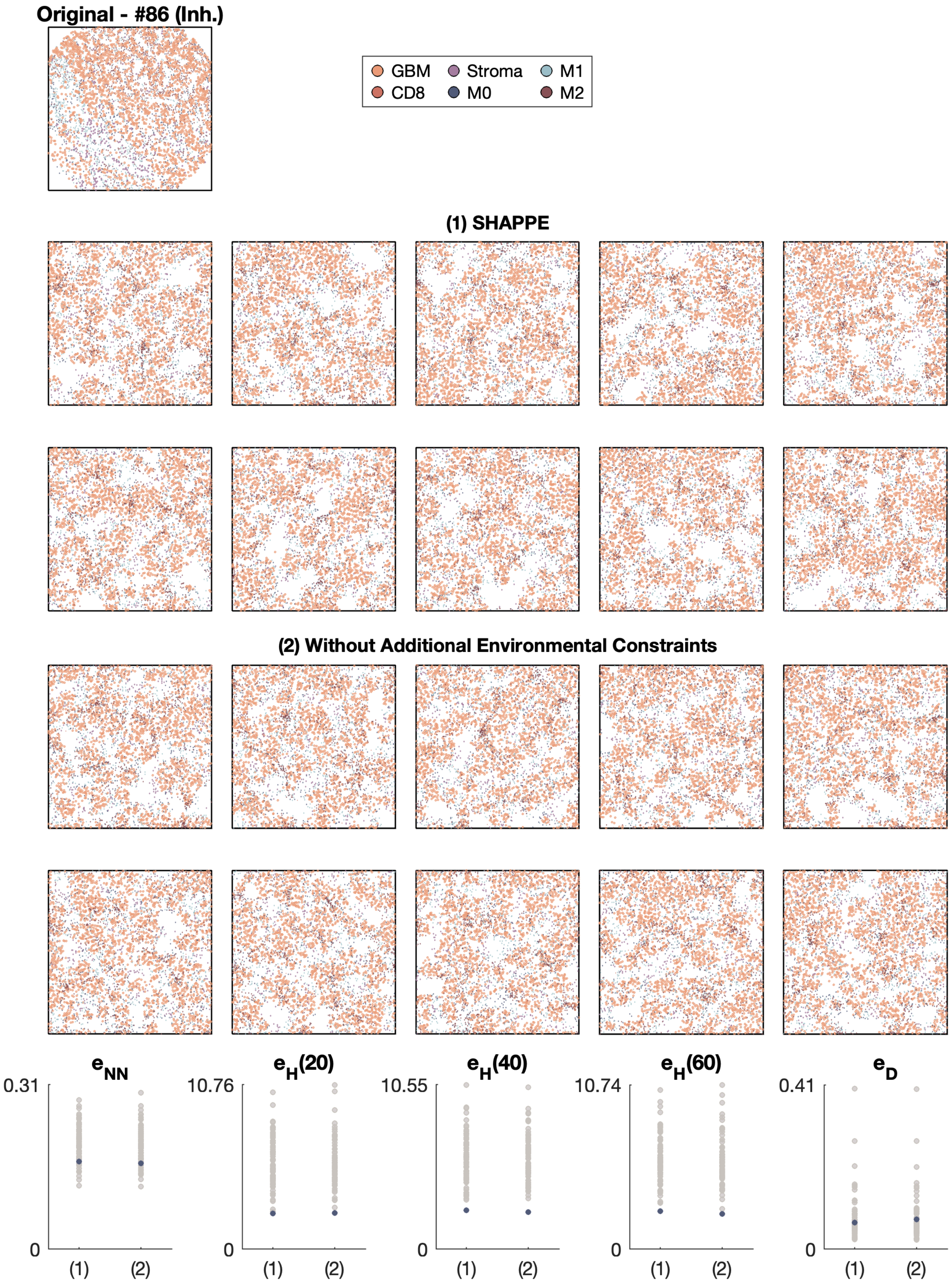

Original - #86 (Inh.)
GBM
Stroma
M1
CD8
M0
M2
(1) SHAPPE
(2) Without Additional Environmental Constraints
e_NN
e_H(20)
e_H(40)
e_H(60)
e_D
0.31
10.76
10.55
10.74
0.41
0
(1)
(2)

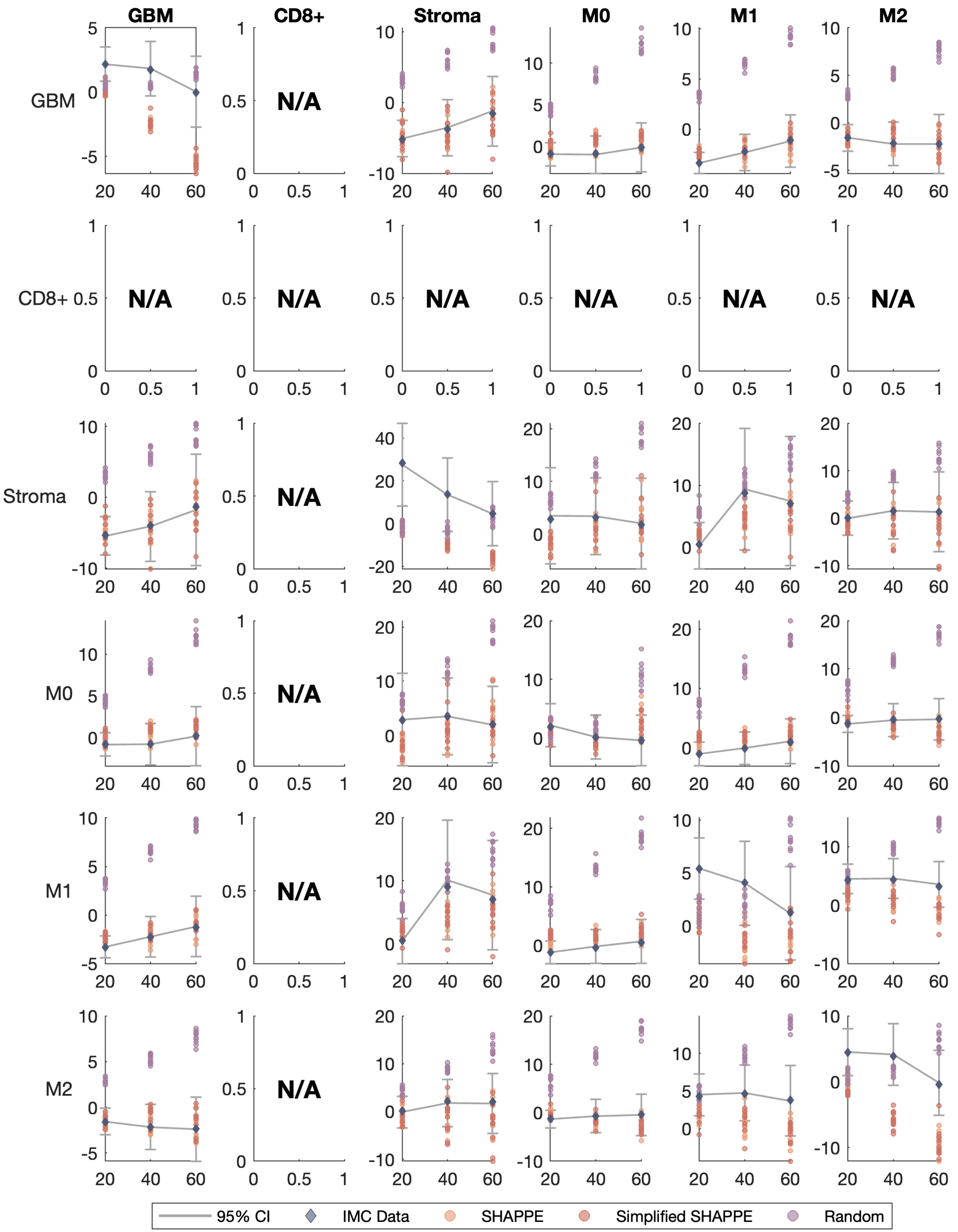
GBM
CD8+
Stroma
M0
M1
M2
N/A
95% CI
IMC Data
SHAPPE
Simplified SHAPPE
Random

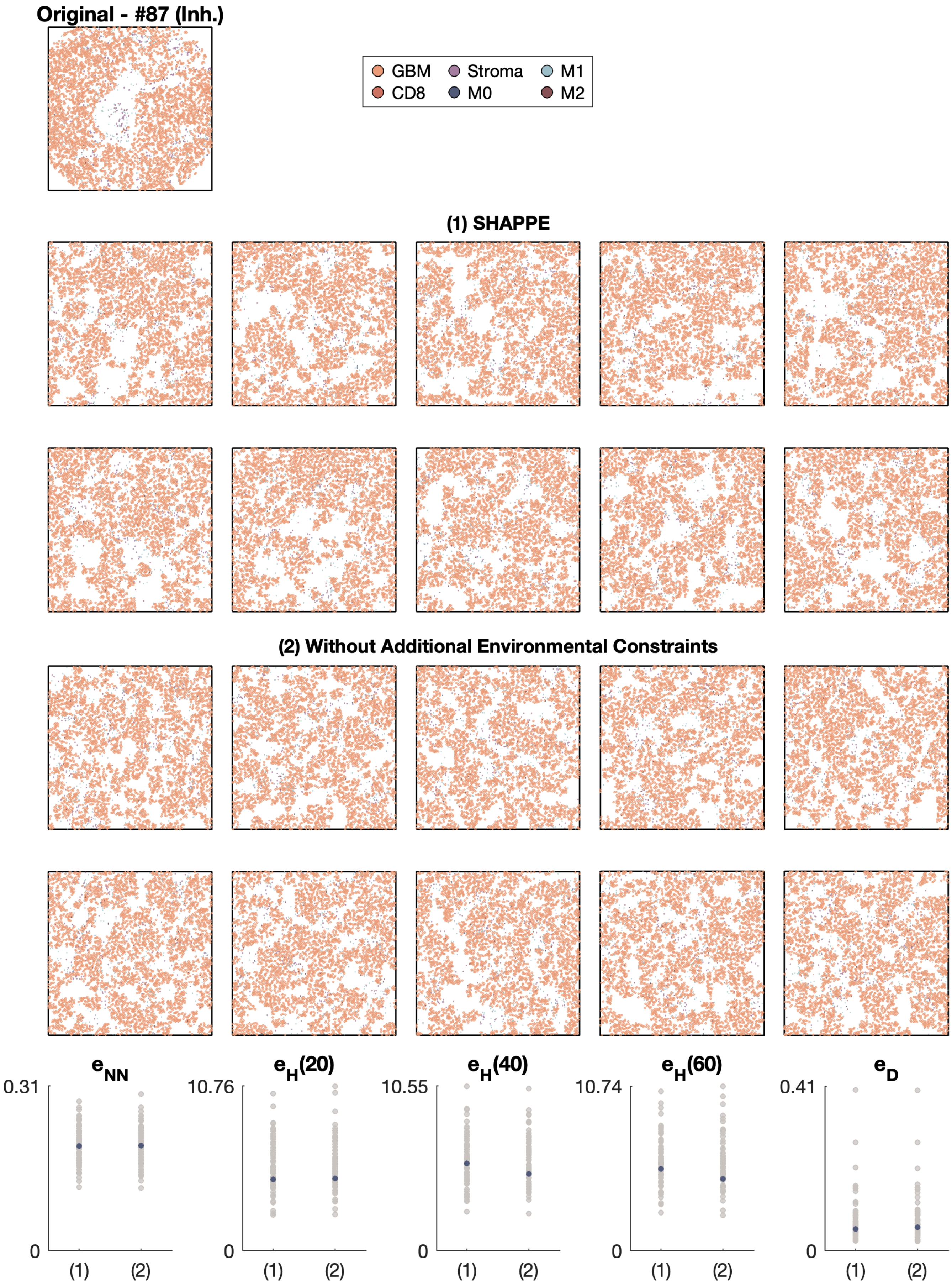
Original - #87 (Inh.)
GBM
CD8
Stroma
M0
M1
M2
(1) SHAPPE
(2) Without Additional Environmental Constraints
$e_{NN}$
$e_H(20)$
$e_H(40)$
$e_H(60)$
$e_D$
0.31
10.76
10.55
10.74
0.41
0
(1)
(2)

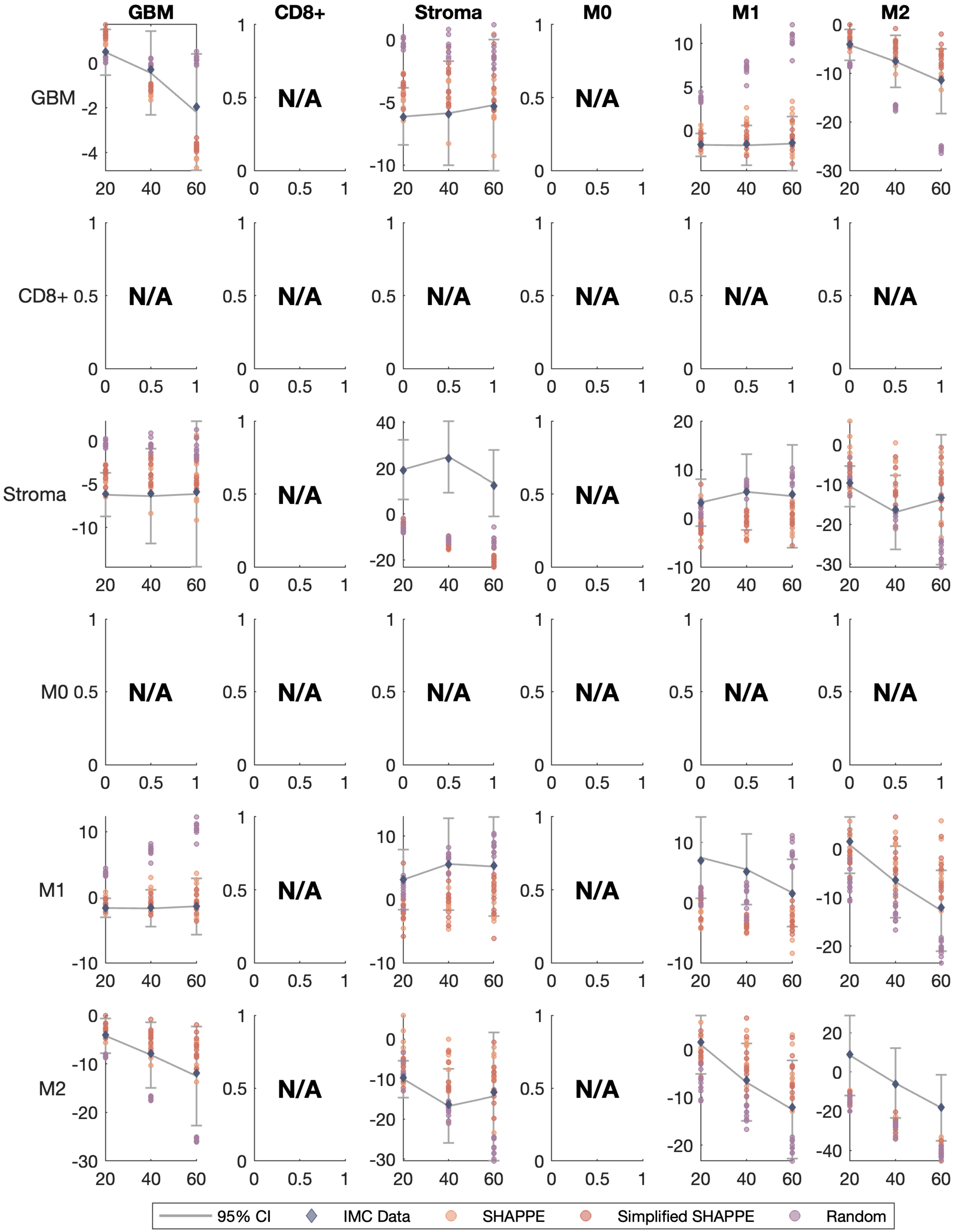

GBM
CD8+
Stroma
M0
M1
M2
N/A
95% CI
IMC Data
SHAPPE
Simplified SHAPPE
Random
